# Large scale application of species distribution models to predict future vulnerability to social wasp invasions

**DOI:** 10.64898/2026.08.10.744014

**Authors:** Thomas Hagan, Sara E. Miller

## Abstract

Social wasps (family: Vespidae) are increasingly concerning invaders and have been subject to increased detections and a growing number of invasive populations in the last few decades. As established invasive populations are challenging to eradicate, preventing introductions and prioritizing early interventions are the most cost-effective management solutions to mitigate these effects. A current challenge to this approach is that species distribution data is limited for many social wasp species, hindering our ability to accurately predict novel habitats with high suitability. To address this gap, we used MAXENT to create species distribution models (SDM) for 299 species of social vespid. We identified existing invasive populations of social wasps and incorporated their current invasive ranges to improve the transferability of our models in predicting habitat suitability in new environments. Current range sizes and habitat suitability varied widely among species and genera. We identified new species of high invasive concern, particularly in the genus *Vespa*. We also identified previously unrecognized regions that may be at high risk of future invasion primarily in Central Africa and the Indo-Australian Archipelago. Combining current and suitable ranges, we calculated an “Invasion Risk Score” to compare the relative likelihood of each species establishing a new invasive population based upon habitat suitability. To assess invasion risk in the future, we projected habitat suitability under four Shared Socioeconomic Pathway (SSP) climate change scenarios. Under all scenarios, species faced significant changes in habitat suitability for current native ranges. Habitat suitability generally shrank and shifted towards the poles, leaving equatorial species at highest risk of habitat loss. Notably, *Vespa* was the only genus whose suitable habitat expanded under these climate scenarios. Our framework demonstrates how multi-species SDMs can be applied to risk management of invasive populations.

## Introduction

Invasive species are some of the most devastating by-products of globalization, being the cause of severe economic and ecological damage (1–4). Social insects (termites, ants, some bees and wasps) number among the worst invasive species listed by the IUCN (5) and predatory wasps are of growing concern (6). The success of social insect invaders is often attributed (albeit not solely (7)) to the division of labor, which offers protection from predation to reproductive individuals (8). Despite higher intensity in detection and management practices, nascent invasive populations of insects, including social insects, are becoming increasingly common (9). As globalization and trade intensifies, increased movement between and within regions is predicted to accelerate social insect invasion rates (10). Predicting future invasion risk is complicated by the impact of climate change, which will both shift the current ranges of invasive species and make previously unsuitable environments accessible to new invaders.

Of the social Hymenoptera, wasps within the family Vespidae are the subject of growing concern (11). This is due to both a history of successful invasions (12) and a recent surge of several notable invasive hornets (13, 14). Invasive social Vespidae can be incredibly costly. For example, the common wasp (*Vespula germanica*) causes damage to human and animal health, horticulture industries, pastoral industries and apiculture industries, costing New Zealand’s Agricultural and Health industries approximately NZ$133 million (15) and could cost Australia as much as AUD$2.7 billion over 50 years if not for current management programs (16). Furthermore, the recent successful invasion of the Asian yellow-legged Hornet (*Vespa velutina*) causes losses to the apiculture industry of at least 100 million €/year in France alone (17). Unfortunately, the eradication of Vespidae species is difficult, and even promising solutions such as gene drives require additional constraints not required for other taxa (18). As cost-benefit analyses typically favoring eradication over management (16), the obvious solution is to prioritize prevention and early intervention when eradication costs remain low. Indeed, successful eradication programs can often be attributed to early intervention (19), such as the successful eradication of the Asian giant hornet (*Vespa mandarinia*) (20). However, the detection of invasive wasp species is not one size fits all and trying to monitor all possible invaders may hinder detection efficiency. A more efficient solution would be to focus on pre-identified species of concern with a high suitability to local environs.

Social Vespidae are contained to three subfamilies; Stenogastrinae (Hover wasps: typically colony sizes are small and species can be facultatively eusocial (21, 22)), Polistinae (Paper wasps: colony sizes range dramatically and species vary from facultatively eusocial to obligately eusocial (23)) and Vespinae (Yellowjackets and Hornets: colony sizes are large and species are obligately eusocial (24)). Invasive species in social Vespidae are mostly contained to Polistinae and Vespinae (12), within genera *Polistes* (Global distribution (23)), *Vespula* (North American, European and mainland Asian distribution (25)) and *Vespa* (European and Asian distribution (26)). It is unknown why these genera are more successful invaders, although one possibility is that species with larger distributions may be more likely to establish new populations by chance alone. Unfortunately, the native range of many other wasp species (particularly those of South American and Southeast Asian origins) are not yet known to a high degree of confidence, which makes testing this hypothesis difficult.

Species distribution models (SDMs) are tools used to determine regions of high climate suitability for a given species. These models typically use spatial data to extract climate variables at places of known presence and then attempt to find relationships that delineate presence and absence. These models are typically used to determine the current distributions of species (native or otherwise) but can be extended to determine regions of high climate suitability outside of a species current range. We consider these habitats to be regions of high invasive potential. SDMs are numerous, but models that use the MAXENT algorithm have been shown to be robust over many circumstances (27). MAXENT models also perform relatively well with a limited number of presence points, although increasing data still helps to successfully predict invasive potential.

Thankfully, in recent years the proliferation of online biological databases and citizen science projects has allowed for an abundance of location information that can be used in ecology and population biology (28, 29). In fact, community science initiatives are so comprehensive that they have begun to offload the burden of early detection (30, 31). These databases now offer an exciting new possibility to rapidly determine the distributions of species that were previously unknown. This not only has large implications for the management of native species but will also help inform management and eradication efforts for current and future invasive species. SDMs constructed using data from these biological databases can now pre-identify species of invasive concern, if they account for issues surrounding transferability (transferring predictions to regions outside a species current range) (32, 33). This is especially useful for species that have been previously disregarded with respect to invasive potential. These include species native to, and abundant in, regions that were previously inaccessible (e.g., in dense rainforest), where SDMs will illuminate potentially high-risk invaders that are constrained only by their location.

Here we use the wealth of distribution data available from online resources to model the current ranges of 299 species of social Vespidae. Using species with existing invasive populations, we calculate a corrective factor that is applied on models to account for transferability. We then identify suitable habitat and predict potential invasive ranges of currently non-invasive species. Using regions of invasive suitability, we comment on which species, in which regions, are likely to be of a high degree of invasive concern in the next few decades. Finally, we predict how both native and invasive populations are likely to shift under multiple climate change scenarios.

## Methods

### 2.1 Data Cleaning

We obtained distribution records of all social Vespidae through the online databases GBIF (34) and iNaturalist (accessed 22^nd^ of November 2024). Where applicable we also supplemented these records with published occurrence records (23, 35) and transcribed coordinate information of non-digitized specimens with associated spatial data from the American Natural History Museum, the Smithsonian National Museum of Natural History, Cornell University Insect Collection (CUIC), Kansas University Entomology Collection, and the University of Guelph Insect Collection. Records without Longitude or Latitude data, records at 0° Latitude and/or Longitude, records found in oceans and duplicate records with the same Latitude and Longitude coordinates as another record were removed. We then visually inspected the distribution records for each species and removed any aberrant records from continents where that species is not native and where no invasive populations have been reported (N=521 records over 106 species representing 0.15% of the total number of records). These records likely represent misidentified specimens. Some occurrence records of *Vespula* species required further cleaning due to species synonymization and revisions. In all cases these *Vespula* species were initially believed to have a Holarctic distribution but have since been revised to differ between the Pale- and Nearctic (i.e., *V. austrica*, *V. rufa* and *V. vulgaris* in North America are now *V. infernalis*, *V. intermedia* and *V. alascensis* respectively). Finally, we performed a literature search to identify known invasive populations of Vespidae species (12).

Species with fewer than 25 coordinates were removed from our dataset, and any population with less than 20 coordinates were ignored. Populations were identified visually and defined by continuous distributions within or across continents. For example, coordinate distributions that are sharply discontinuous across coastal Europe to coastal Asia are counted as two populations, but continuous distributions would be counted as one population. Island populations that were noted as being invasive were also treated as different populations. Only one species in our dataset had an invasive population that was later eradicated (*Vespa mandarinia*), but this invasive population had fewer than 20 coordinates and was thus not included in our analysis. After data cleaning, our dataset contained 299 social Vespidae species, of which 16 species had one or more invasive populations with sufficient data for analysis.

### 2.2 MAXENT Models for Current Populations

We used our cleaned data to create MAXENT models in R (MAXENT version 3.4.3, *dismo* v1.3-14, R v4.4.2) for each species in our dataset. MAXENT models were created using presence data from all populations (native and invasive), climate data (2.5” resolution tif files for 19 biological variables and elevation available on WorldClim (36)) and were trained using 10^5^ background points from across the globe (which helps to determine the probability distribution of covarying environmental variables). We opted to not perform stepwise variable selection across our models as: a) models tended to deviate little with variable removal, b) we were unsure *a priori* which variables would be important for different species across such a broad taxa and c) our aim was to keep models entirely comparable across our dataset. The RAW output of MAXENT models are tif files with a habitat suitability score in each 2.5” cell. Habitat suitability scores were then used to predict species presence, absence and potentially suitable invasive habitat.

To find discrete distributions from these outputs we established a suitability score threshold value above which species presence is assumed. This threshold was found through the model output that resulted in the highest True Skill Statistic (TSS: TSS = Sensitivity (True Positive Rate) + Specificity (True Negative Rate) - 1) for the model. Calculating Specificity relies on generating pseudo-absence data, randomly generated spatial points that represent “absent” observations. Unfortunately, global pseudo-absence coordinates may artificially inflate threshold values for species with multiple populations, particularly for species with different coordinate densities across populations. Therefore, we elected to perform these thresholds by population rather than by species. We defined a large region around these population from which 10^4^ pseudorandom background points were drawn. The region surrounding a population was created by finding the convex hull that contains all coordinates for that population using the function *st_concave_hull* (and a concavity of 1: *sf* v1.0-19) and then buffered that polygon using *st_buffer* by 36” (approx. 4000 km). We also used a similar polygon (buffered by 4.5”, approx. 500 km) to define the region over which the populations threshold value was applied, as to prevent overlap in thresholds between populations. The coordinates used as test data in *evaluate* were dependent on the number of distribution coordinates in a population. In cases where the number of coordinates in a population is high (≥ 200) the k-fold partition method is preferred. Here randomly generated folds are approximately representative of the entire dataset and a small number of models generates a generally similar set of thresholds. Instead, where the number of coordinates is low (< 200) the bootstrap method is preferred. Here the high number of models generated results in a threshold that approaches optimal. For populations that fit the k-fold partition method, we subdivided the distribution data into 5 equally sized datasets (folds). We initially retrained another model using folds 2-5 and retained fold 1 as test data. We then repeated this process such that each fold was used as test data once, and then took the average threshold found by each of the 5 folds. For populations that fit the bootstrap method, we sampled with replacement from the initial training data N samples, where N was the number of coordinates used to train the model. We used this sample as the testing data and repeated this process 100 times. We then took the average threshold value of the model as the threshold value of the population. Finally, we defined each cell with a value higher than or equal to the population specific threshold to be where the species is present, and cells with values below this threshold to be where the species is absent. Native and invasive species distributions were then collated by genus to better compare with other results.

### 2.3 Transferability Corrections

The ability of SDMs to predict regions of invasive success is termed transferability. Simple SDMs typically have higher transferability than complex SDMs, although they have a tendency towards overprediction (32). Complex SDMs likely have lower transferability as the complex relationships between environmental variables and species presence may not extend into the unique conditions experienced in a potential new range. In our study, we opted to retain complexity and to instead address transferability using real invasive populations to inform the relative differences in model predictions before and after invasion. To achieve this, we built another series of MAXENT models that excluded occurrence data from invasive populations for applicable species (Table 1). Using these models and their complete dataset counterparts, we then calculated the relative difference in prediction values for invasive populations (a diagram of this process can be found in Figure S1). In some cases, we found that the predicted values of these invasive regions prior to invasion were particularly small (usually in cases of island populations or where invasive population coordinates outnumbered native coordinates). These species were deemed as having low transferability, likely the result of island population experiencing environmental conditions not present in the native range. To prevent the bias of populations with extremely low transferability, we removed any cells from invasive population with values lower than 0.01, the approximate value of the lowest threshold used in our *Current Populations* section (∼0.011). We found the relative difference in cell value for all cells with an invasive population between models trained with and without invasive populations was ∼0.62. That is, cells of invasive populations trained in models without those populations had a value ∼0.62 smaller than of cells trained with these populations. We therefore considered any cell with a value above half (a slightly more conservative estimate than 0.62) the threshold of a population determined in the native range to allow for species survival. Transferability corrected threshold values are therefore half the threshold values predicted in section 2.2.

**Table 1:** Current populations of non-native social Vespidae. Native Area is the size of the original habitat of each species, Invaded area is the current non-native habitat, and potential invasive area is the size of non-native habitat (excluding current invaded area) that would be suitable for this species.

| Species | Native Area (10<sup>6</sup> km<sup>2</sup>) | Invaded Area (10<sup>6</sup> km<sup>2</sup>) | Potential Invasive Area (10<sup>6</sup> km<sup>2</sup>) |
| --- | --- | --- | --- |
| *Brachygastra lecheguana* | 4.081 | 0.010 | 7.271 |
| *Polistes aurifer* | 4.581 | 0.023 | 26.813 |
| *Polistes chinensis* | 2.331 | 0.580 | 7.119 |
| *Polistes dominula* | 8.762 | 6.756 | 1.152 |
| *Polistes exclamans* | 4.149 | 0.077 | 25.470 |
| *Polistes humilis* | 0.813 | 0.231 | 7.191 |
| *Polistes jokahamae* | 2.274 | 0.003 | 0.319 |
| *Polistes olivaceus* | 3.000 | 0.199 | 4.248 |
| *Polistes versicolor* | 4.198 | 0.010 | 9.061 |
| *Polistes wattii* | 5.177 | 0.005 | 1.488 |
| *Vespa crabro* | 6.195 | 3.168 | 4.223 |
| *Vespa orientalis* | 5.832 | 0.055 | 3.620 |
| *Vespa tropica* | 7.797 | 0.002 | 13.425 |
| *Vespa velutina* | 5.902 | 1.411 | 21.689 |
| *Vespula germanica* | 7.628 | 4.775 | 1.031 |
| *Vespula pensylvanica* | 3.684 | 0.005 | 16.334 |
| *Vespula vulgaris* | 5.700 | 1.105 | 7.783 |

### 2.4 Regions of Invasive Suitability

We took our original models and applied the corrected threshold values to find regions of invasive suitability. For species with multiple populations, and therefore multiple threshold values, we took the lowest threshold value for the species. We assumed species could survive in any cell with a value higher than this threshold. Some currently invasive populations in our dataset are found on the same continent as their native population (e.g., *Polistes exclamans* and *P. humilis*), however it is often difficult to distinguish these from cases of population expansion. We therefore opted to exclude any cells that shared a continent or island with an already established population (native or invasive) to prevent population expansion from obscuring our results. We also collated these populations by genus, allowing a broad look at the regions of invasive concern of a genus.

### 2.5 Predicting future suitable habitat

To investigate how the distribution of different populations might change in the future, we applied the same models produced in the *Current Populations* section of our methods to forecasts of future climate. We obtained predictions of future bioclimatic climate data through WorldClim, using the CMIP6, ACCESS-CM2 projections at 2.5” resolution (36). We modeled species distributions for the four Shared Socioeconomic Pathway (SSP) climate change scenarios (ACCESS-CM2, SSP1-2.6, SSP2-4.5, SSP3-7.0 and SSP5-8.5) over four time periods (2021-40, 2041-60, 2061-80 and 2081-2100). These represent the main pathways suggested by the CMIP6 (37), each with increasing climate change impacts. To find defined populations of species for each scenario and time period, we predicted climate suitability using these scenarios. For current populations we applied a buffer of 2000 km (using the function *buffer* in *terra* v1.7-39) and applied the same threshold found in the *Current Populations* section of our methods. This buffer prevented current populations extending to new continents but still allowed for significant population movement. The resulting regions were considered indicative of how these populations may shift under multiple climate change scenarios. Shifts in regions of invasive suitability were found by simply applying the same restrictions and thresholds as found in the *Regions of Invasive Suitability* section of our methods. We then calculated the relative increase or decrease to habitat area in both cases to investigate the impact of different SSPs on wasp populations. We also collate these predicted populations by genus, to allow a broad look at population shifts.

### 2.6 Statistical Analyses

Closely related species may exhibit similar physiological tolerances. To test if current species ranges had high habitat suitability for other members of the genus, we performed Pearson correlations between the native, invasive and potentially invasive distributions of each genus. We also performed a regression analysis using *lm* (*stats* v4.4.2) to find the relationship between the size of a genera’s native range (10^6^*km^2^) and the size of its current invasive range (10^6^*km^2^).

Finally, we calculated a simple metric to rank the relative future risk of a new invasive population of each species based upon habitat suitability. We estimated the potential propagule pressure as the size of the species currently occupied range (native and invasive; A_O_) and area at risk as the size of the species potential invasive range (A_P_). We considered the product of these two values as representative of invasive risk, which was then log transformed and standardized using the log transformed area of the terrestrial globe (A_T_ = 1.49*10^8^ km^2^) squared. Scores were multiplied by 100 to scale between 0-100. The metric is calculated like so:

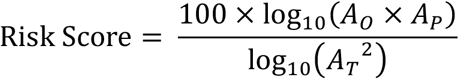

A higher score indicates that a species is at high risk of becoming invasive, either due to high propagule pressure (due to the size of A_O_) or a large area of invasive potential, or both.

## Results

### Current distributions of social Vespidae

We found the native range distributions of 299 social Vespidae (summarized by genus in Figure 1). The range size varied widely among species (Range: 0.2-11.6*10^6^ km^2^). The average range was 2.5*10^6^ km^2^, with ∼90% of species occupying an area of less than 5.7*10^6^ km^2^. Species tended to overlap with other members of the same genus, and the average area occupied by a genus was 12.4*10^6^ km^2^. Sixteen species in this dataset had current invasive populations with sufficient data to model (Figure 2; Table 1). For these species we found a positive correlation between the area of the native population and that of the invasive population(s) (Invasive Area (10^6^ km^2^) = 0.6*Native Area (10^6^ km^2^) – 1.9, F_1,15_ = 11.4, R^2^ = 0.39, p = 0.004), suggesting native range size plays some role in invasive potential. However, as many of the invasive ranges are either limited island populations or currently expanding, this relationship should be interpreted with caution.

**Figure 1:**
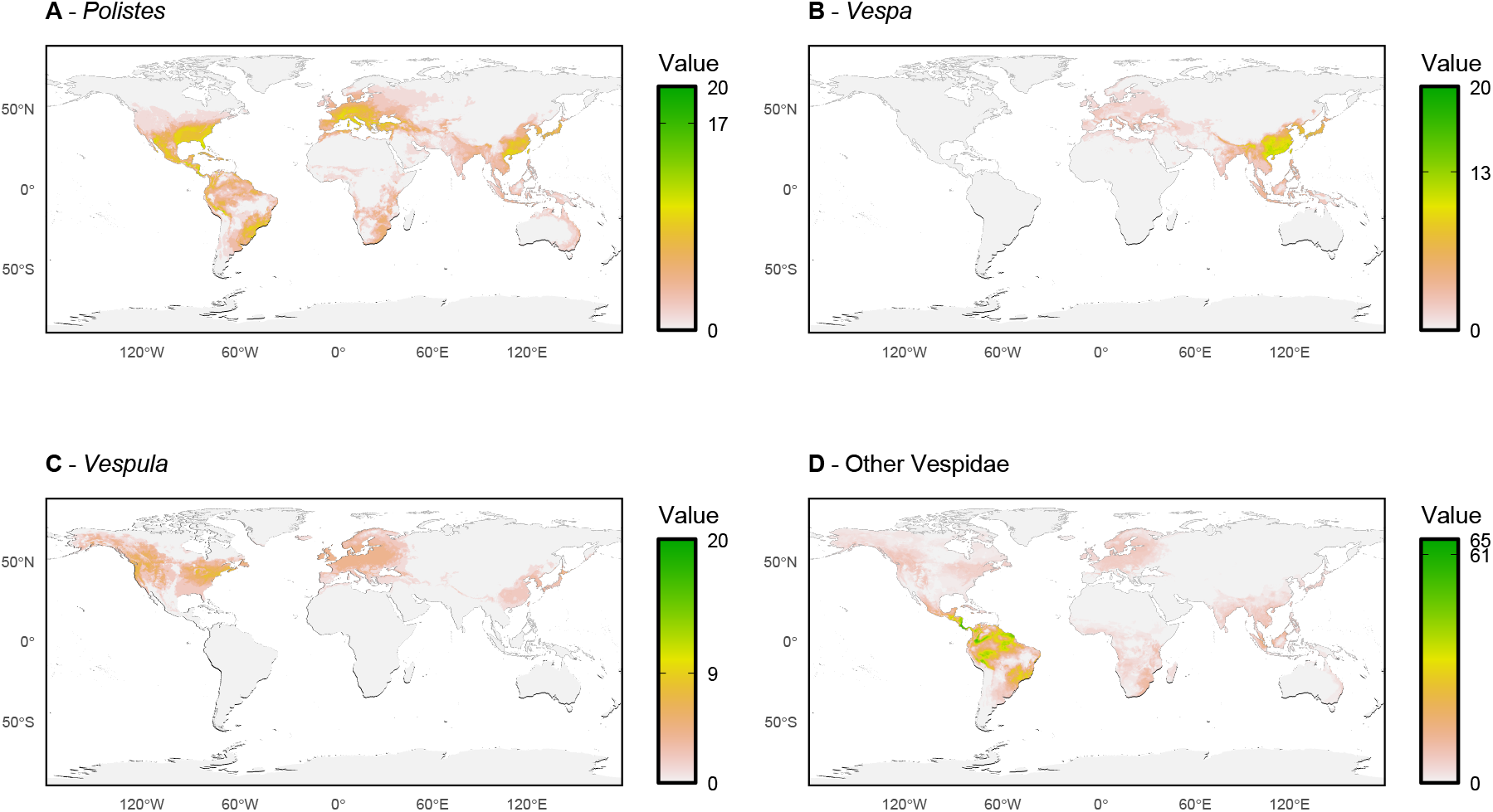
Species richness plots of native range populations for *Polistes* (A), *Vespa* (B), *Vespula* (C) and all other social Vespidae (D).

**Figure 2:**
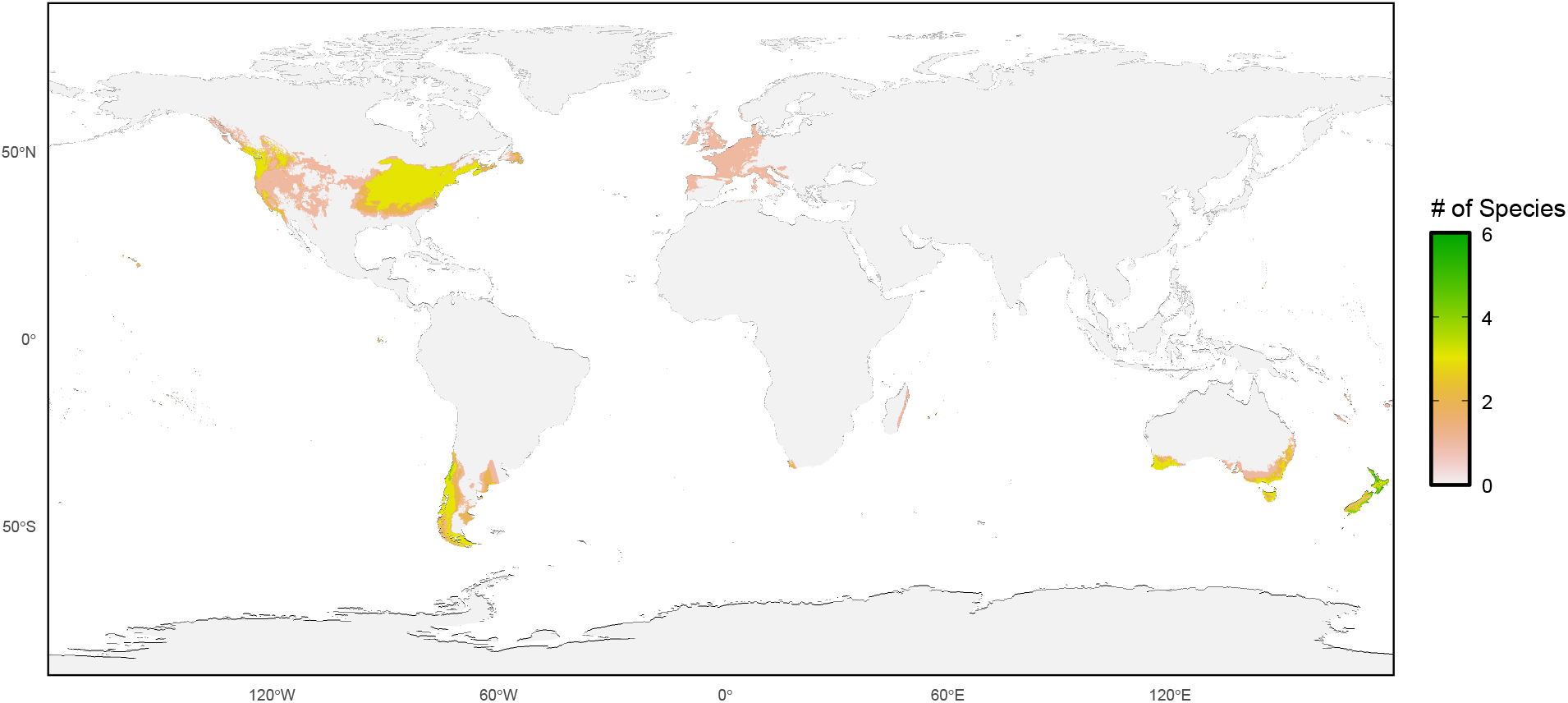
Species richness plot of invasive populations of all social Vespidae in our study.

### Invasive suitability varies by genus

Next, we identified regions of invasive suitability for all 299 species included in our study (Figure 3). The average area of suitable habitat outside of the native range was 4.10*10^6^ km^2^, roughly equivalent to the size of the South Asian subcontinent. However, this value varied greatly among species (1.68*10^4^ km^2^ to 2.68*10^7^ km^2^). Furthermore, locations of invasive suitability varied greatly among genera. These patterns are likely caused by similarities in physiological tolerances and current geographical distributions shared by species of the same genera. We outline specific regions of concern below.

**Figure 3:**
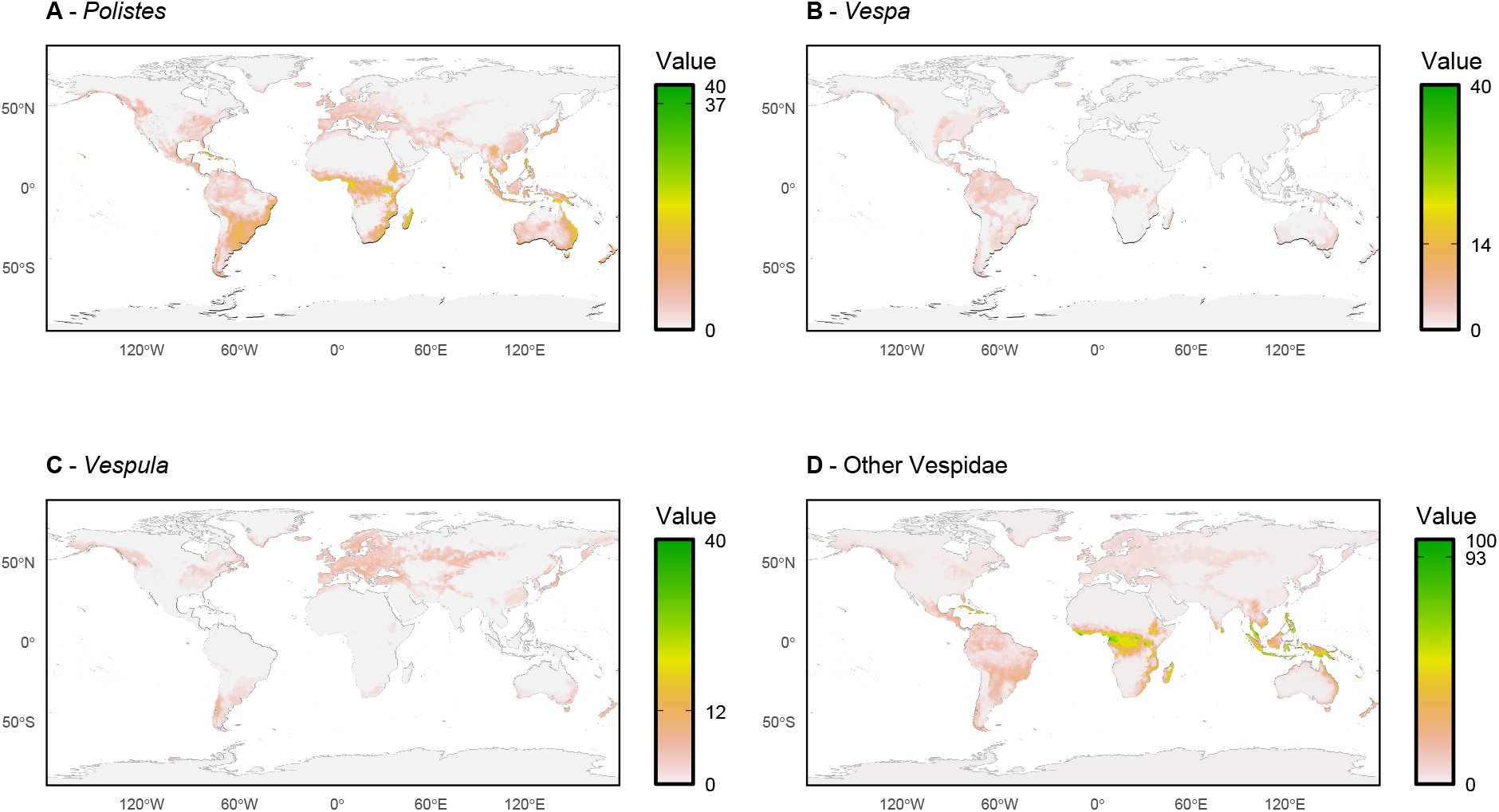
Species richness plots of invasively suitable habitat for *Polistes* (A), *Vespa* (B), *Vespula* (C) and all other social Vespidae (D). Note, these represent regions at risk of invasion, not those with current invasive populations.

### Regions at risk from Polistes

*Polistes* species are predicted to be highly suited to Central and Eastern Africa, the Río de la Plata basin of South America, South East Asian & Pacific Islands and Eastern Australia. Interestingly, the average region of concern per species is relatively small compared to other taxa (3.4*10^6^ km^2^; Table S1). Instead, a few highly plastic generalist species are the major drivers of the numerous regions of invasive risk across this large genus.

### Regions at risk from Vespa

*Vespa* species have a high suitability in Eastern Africa, Eastern Australia, the Pacific Islands, the Amazon, Japan and Eastern North America (particularly Florida). Many *Vespa* species follow a similar pattern to that of *Polistes*, however there are some species with notably larger regions of concern in Europe, Australia, New Zealand and the Southern Cone of South America.

### Regions at risk from Vespula

*Vespula* species are suited to much of the Holarctic, but this is mostly complementary. That is, North American *Vespula* species are highly suited to Europe and vice versa. However, some *Vespula* are also highly suited for other regions in South America, Australia and New Zealand (Figure 3).

### Other Social Vespidae

Finally, other social Vespidae species tend to have small regions of suitability that are equally distributed across the globe. However, both Central Africa and the Indo-Australian Archipelago have a high number of potential invasive species, indicating that these regions are at higher risk. Furthermore, species in this category do not have a history of invasiveness, so this risk is likely underappreciated.

### Estimating future invasion risk

Using the range currently occupied by a species (Invasive and Native regions) and the potential invasive range of a species, we calculated an “Invasion Risk Score”. The top 5 species with the highest score were 2 *Vespa* species (*V. velutina* and *V. tropica*) and 3 *Polistes* species (*P. stigma*, *P. aurifer* and *P. exclamans*).

We found that the native range of a genus was a highly variable predictor of invasive suitability, with Pearson correlations between native ranges and at-risk regions varying greatly (−0.03 – 0.50). Genera with a high correlation between native and suitable invasive regions have a semi-global distribution, such as the globally distributed *Polistes* (0.39) and the Holarctic distributed *Vespula* (0.43) and *Dolichovespula* (0.50). In these cases, native populations may act as a “proof of concept” of invasive potential, indicating that the climatic conditions of native ranges are at least somewhat suitable for potential invaders. However, this is more likely to indicate the large area inhabited by these taxa. As all other genera have a low Pearson correlation (< 0.24), it seems native area is usually a poor predictor of invasive suitability. The currently invaded range of a genera also tended to be a poor indicator of invasive suitability (0.00 – 0.26). In all, the regions of highest invasive suitability for social Vespidae will be regions naïve to the group, both native and currently invasive.

### Predicting future habitat suitability

Lastly, we considered how native ranges, invasive ranges and regions of invasive suitability will shift under several climate change scenarios. All scenarios resulted in significant changes in habitat suitability for current native ranges. In general, all climate change scenarios tended to both shrink ranges and shift them towards the poles (Figure S2). Species with equatorial native ranges are likely at highest risk as suitable habitat shifts often resulted in population splits. Shrinking populations, range shifts and population splits mean that native wasps are likely to experience significant pressures due to climate change in the future.

Species that are currently invasive also tended to decrease in range under most climate change scenarios. However, these ranges also moved poleward to regions currently free of invasive social Vespidae. This is notable with *Polistes dominula*, which is predicted to shift much further into Canada and southern Alaska. Regions of invasive suitability exhibited similar patterns, except when equatorial where regions tended to grow. This effect is most notable within *Vespa*, the only invasive genus that grew in invasive suitability under extreme climate change scenarios (Figure 4). This suggests that while climate change won’t result in the overall growth of globally invasive social Vespidae (current or potential), it does have the capacity to open new, vulnerable regions to these invaders.

**Figure 4:**
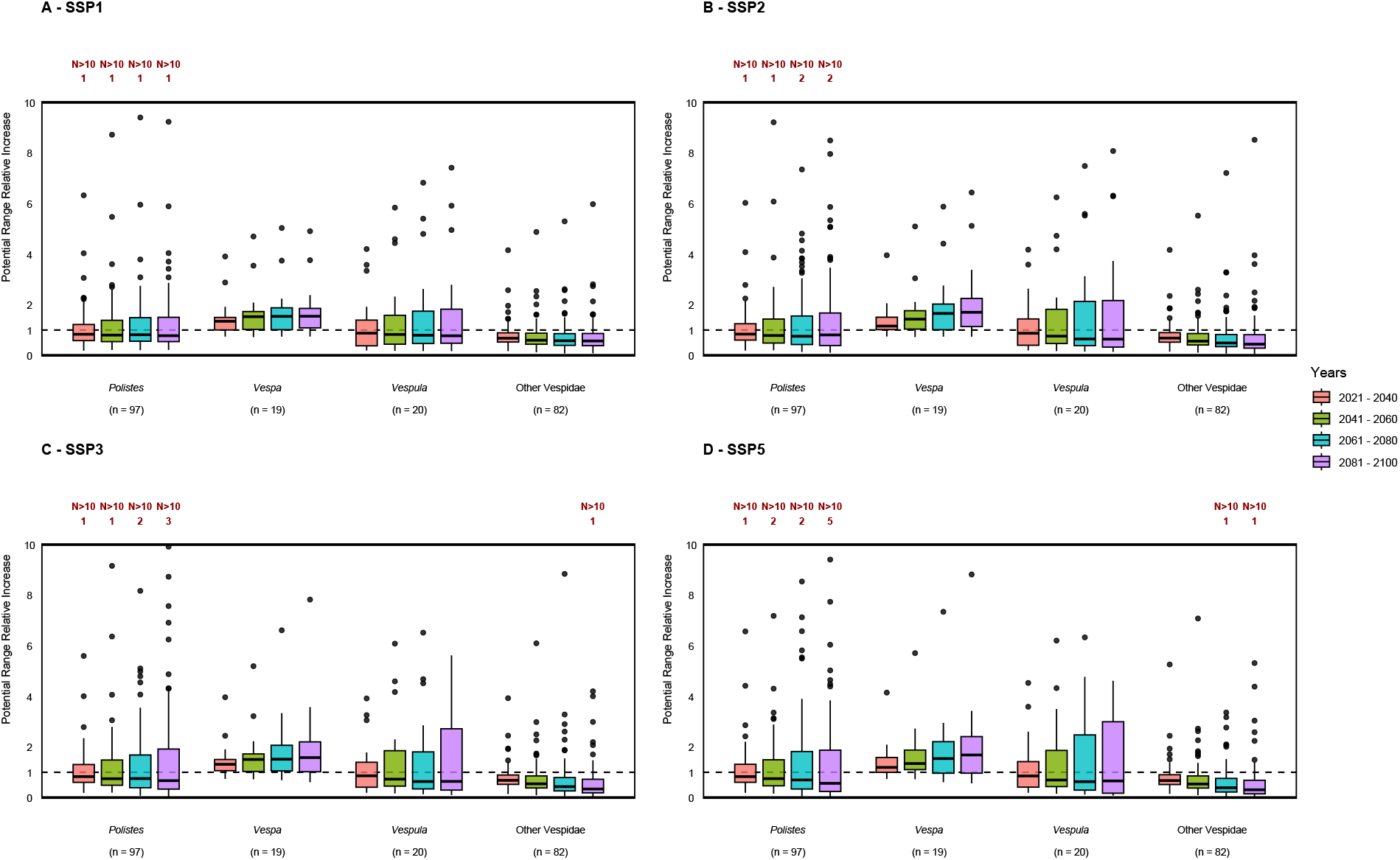
The relative increase of invasively suitable habitat under different climate change scenarios (A – Shared Socioeconomic Pathway (SSP) 1, B – SSP2, C – SSP3, D – SSP5). This is a relative increase, so the dashed line at 1 indicates no increase in suitable habitat. While many species have increases in suitable habitat, the only genus with consistent increases in habitat are *Vespa*.

## Discussion

In this paper we have successfully modelled the native ranges of 299 social Vespidae, and the invasive ranges of 16 species. We found that species distributions and range size impacts invasive suitability in predictable ways, such as larger native ranges corresponding to larger invasive ranges. We also identified new species of concern and previously unrecognized regions that may be at high risk of future invasive social wasp populations. Lastly, we show that the risk of new invasive populations is not static. Regions of suitability for both native and invasive species will change – often dramatically – with a changing climate. One notable group is the genus *Vespa*, which has high invasive potential under all climate change scenarios .

The size of a genera’s native range is predictive of the size of the size of the invasive range of a genus, but only in some cases. Conversely, the current distributions of invasive species do not predict suitable invasive habitat for other species in a genus. This could indicate two separate phenomena. Firstly, a species-rich genus is more likely to contain pairs of species with similar physiological tolerances in geographically distant places. Native species may act as a “proof of concept” for the survival of similar species within the genus. This is perhaps not surprising, there are many cases of invasive species invading in the range of closely related sister species (38, 39), and even within our own study we observe that there are some notable cases of invasive species invading into ranges of other sister species (such as *Vespula germanica* and *Polistes dominula* in the US, which is home to several native *Vespula* and *Polistes* species). While we did not observe a strong correlation between the native range of a genus and its current invasive range, this is likely due to the native range of most generas being far larger than their currently invaded range. Secondly, this result indicates that future invasions of social Vespidae will occur in naïve ecosystems that may not have appropriate countermeasures in place. Most of the invasively suitable habitat of social Vespidae (including for our taxa of greatest concern; *Polistes*, *Vespula* and *Vespa*) occurs in regions that do not have other native species in the same genus, let alone other invasive species. To prevent the devastating impact of social Vespidae invaders, we suggest preparation in regions of highest concern (Central Africa and the Indo-Australian Archipelago). Finally, these regions are at risk not only from already common invaders within *Polistes* and *Vespa*, but also from invasively naïve social Vespidae (including species within Epiponini, *Mischocyttarus* and *Ropalidia*).

Climate shifts will impact the distributions of both native and invasive Vespids. Generally, the habitats of all Vespid populations will shrink as climate change intensifies, including equatorial populations splits. Current populations of invasive Vespids showed similar trends. A simple interpretation is that the impact of existing invasive populations and the risk of future invasive Vespids will be diminished under most climate change scenarios. However, this interpretation does not consider several aspects of invasive biology. First and foremost, these models do not consider the capacity for phenotypic plasticity or adaptation. For example, the invasive European paper wasp (*P. dominula*) population in the United States initially established in a similar climate to its native range but has subsequently spread into drier regions with climates of higher seasonal variability (40). While our relaxed thresholds account for some of the phenotypic plasticity that may occur between native and invasive populations, it is unlikely to account for all. Similarly, we cannot account for the adaptations that might occur in native populations that will also increase their ability to survive in novel climates. Secondly, while climate change may reduce the environment available for social Vespid invaders writ large, it also shifts habitat suitability into regions previously inaccessible (for example, northwards movements of *Polistes* invaders into Canada). This will introduce social Vespids to species and ecosystems that are entirely naïve to such invaders, which may in turn exacerbate their impacts.

Risk of invasive species depends on both habitat suitability and propagule pressure. Propagule pressure is often defined as the quality, quantity and frequency of invading organisms (41). Most social Vespids are transplanted through lumber shipments and shipping containers (42), therefore shipping traffic is considered a reasonable approximation of propagule pressure (12). We find that regions of high volumes of shipping traffic (North America and Europe) (43), have relatively poor invasive suitability for many social Vespids. While the invaders present in these regions number among the most impactful, it seems that high propagule pressure has resulted in relatively few social Vespidae invaders to date. In contrast, regions of high invasive suitability (Central Africa and the Indo-Australian Archipelago) are experiencing growing volumes of shipping traffic (44). This is concerning, as these habitats are at growing risk for the establishment of new invasive social Vepids. Thankfully, this traffic is not yet associated with shipments from South America (43) where the species of highest invasive suitability (Epiponini and *Mischocyttarus*) are located. Despite the high potential for invasiveness in these regions, many *Mischocyttarus* and Epiponini had low global invasion risk scores, indicating enhanced detection for species in these taxa should only be needed in specific areas. Future work combining our SDM data with detailed measurements of shipping traffic may more precisely identify regions of highest concern.

*Vespa* species have already proven to be highly successful invaders that severely impact local ecosystems and agriculture (12, 45, 46). Our analyses corroborate this and show that some *Vespa* species, but not all, have the potential to be incredibly successful invaders. The four already invasive *Vespa* species are *V. crabro* (invasive in North America), *V. orientalis* (invasive in South America), *V. tropica* (invasive in Guam) and *V. velutina* (invasive in Europe). *V. crabro* and *V. velutina* are particularly successful, *V. crabro* already inhabits a large region of North America (47) and *V. velutina* is spreading rapidly across Europe (12). Here we identified *V. velutina* as the species with the highest global invasion risk. It is climatically suited to large areas of the Eastern United States, much of South America, Australia and many Pacific islands (Figure S3). Given that *V. velutina* has been detected (although not yet established) in both Georgia and South Carolina in 2024 (48), invasive populations in these regions seem inevitable without intervention. Such intervention is possible, proven by the successful eradication efforts of New Zealand early in 2026. *Vespa* species in general had consistently high risk scores, a trait not shared by any other genus in this study. In fact, *Vespa* were the only invasive taxa where suitability grew under multiple climate change scenarios (Figure 4), indicating an increased need for detection and intervention in the coming decades. Despite these concerns, not all *Vespa* species are equally problematic. For example, we found that *V. mandarinia* was only climatically suited on the North American continent in coastal Alaska, coastal western Canada and the southwestern border of the United States (although this is more restrictive than previous models (49–51)). While the discovery of the *V. mandarinia* in the state of Washington in 2019 received high publicity due to concerns of danger to human health (52), this discovery was outside our predicted range. Regardless, due to the broad suitability of *Vespa* across many habitats and the increased risk of invasion with climate change, we recommend for the development and use of *Vespa* specific monitoring in any port with high traffic volume.

Social Vespidae are going to be increasingly common invasive species. Without due preparation they will become established in regions that are hitherto naïve to such invaders. While we could not model habitat suitability for all social Vespid species due to data limitations, 299 species represent a significant portion of the group. Furthermore, the remaining species often have small native ranges that would result in low propagule pressure, and are thus of low concern for future invasiveness. Here we suggest that our models of habitat suitability (current and future) can be applied by local authorities to identify species of local concern. Further, we suggest increased caution of social Vespidae for ports in Central Africa and the Indo-Australian Archipelago, where invasion risk seems to be at its highest and most novel. We also recommend increased caution of *Vespa* species globally, particularly with respect to *Vespa velutina*, due to the large area these species find suitable. While SDMs have been widely used to predict suitable habitats, these analyses were limited by the availability of presence data for each species. Our study shows how we can leverage community science databases to build multi-species SDMs in taxa where this analysis was previously impractical. As the number of community science records continues to grow over time (53), it is likely that these tools will only increase in popularity for this purpose in the future. Finally, we suggest that these methods serve as a basis for understanding invasive potential in other taxa unrelated to social Vespids.

## References

1. Bradshaw CJA, Leroy B, Bellard C, Roiz D, Albert C, Fournier A, et al. Massive yet grossly underestimated global costs of invasive insects. Nature Communications. 2016;7(1):12986.

2. Clavero M, García-Berthou E. Invasive species are a leading cause of animal extinctions. Trends in Ecology & Evolution. 2005;20(3):110.

3. Clavero M, Brotons L, Pons P, Sol D. Prominent role of invasive species in avian biodiversity loss. Biological Conservation. 2009;142(10):2043–9.

4. Gallardo B, Clavero M, Sánchez MI, Vilà M. Global ecological impacts of invasive species in aquatic ecosystems. Global Change Biology. 2016;22(1):151–63.

5. Lowe S, Browne M, Boudjelas S, De Poorter M. 100 of the world’s worst invasive alien species: a selection from the global invasive species database: Invasive Species Specialist Group Auckland; 2000.

6. Wilson Rankin EE. Emerging patterns in social wasp invasions. Current Opinion in Insect Science. 2021;46:72–7.

7. Eyer P-A, Vargo EL. Breeding structure and invasiveness in social insects. Current Opinion in Insect Science. 2021;46:24–30.

8. Moller H. Lessons for invasion theory from social insects. Biological Conservation. 1996;78(1):125–42.

9. Seebens H, Blackburn TM, Dyer EE, Genovesi P, Hulme PE, Jeschke JM, et al. No saturation in the accumulation of alien species worldwide. Nature communications. 2017;8(1):14435.

10. Bertelsmeier C. Globalization and the anthropogenic spread of invasive social insects. Current Opinion in Insect Science. 2021;46:16–23.

11. Otis GW, Taylor BA, Mattila HR. Invasion potential of hornets (Hymenoptera: Vespidae: *Vespa* spp.). Frontiers in Insect Science. 2023;3.

12. Beggs JR, Brockerhoff EG, Corley JC, Kenis M, Masciocchi M, Muller F, et al. Ecological effects and management of invasive alien Vespidae. BioControl. 2011;56(4):505–26.

13. Carisio L, Cerri J, Lioy S, Bianchi E, Bertolino S, Porporato M. Impacts of the invasive hornet Vespa velutina on native wasp species: a first effort to understand population-level effects in an invaded area of Europe. Journal of Insect Conservation. 2022;26(4):663–71.

14. Taylor BA, Tembrock LR, Sankovitz M, Wilson TM, Looney C, Takahashi J, et al. Population genomics of the invasive Northern Giant Hornet Vespa mandarinia in North America and across its native range. Scientific Reports. 2024;14(1):10803.

15. MacIntyre P, Hellstrom J. An evaluation of the costs of pest wasps (Vespula species) in New Zealand. Wellington: Department of Conservation and Ministry for Primary Industries; 2015.

16. Hester SM, Tait P, Kwong R, Lefoe G, Kriticos D, Cacho OJ. Biological control of the invasive wasp Vespula germanica in Australia: Assessing socio-economic feasibility. Ecological Economics. 2024;224:108315.

17. Turchi L, Derijard B. Options for the biological and physical control of Vespa velutina nigrithorax (Hym.: Vespidae) in Europe: A review. Journal of Applied Entomology. 2018;142(6):553–62.

18. Meiborg AB, Faber NR, Taylor BA, Harpur BA, Gorjanc G. The suppressive potential of a gene drive in populations of invasive social wasps is currently limited. Scientific Reports. 2023;13(1):1640.

19. Simberloff D. How Much Information on Population Biology Is Needed to Manage Introduced Species? Conservation Biology. 2003;17(1):83–92.

20. APHIS. APHIS in Action: Victory Over the World’s Largest Hornet Species: U.S. Department of Agriculture; 2024 [Available from: https://www.aphis.usda.gov/news/agency-announcements/aphis-action-victory-over-worlds-largest-hornet-species.

21. Bolton A, Sumner S, Shreeves G, Casiraghi M, Field J. Colony genetic structure in a facultatively eusocial hover wasp. Behavioral Ecology. 2006;17(6):873–80.

22. Hansell M. Elements of eusociality in colonies of Eustenogaster calyptodoma (Sakagami & Yoshikawa) (Stenogastrinae, Vespidae). Animal Behaviour. 1987;35(1):131–41.

23. Miller SE, Bluher SE, Bell E, Cini A, Silva RCd, de Souza AR, et al. WASPnest: a worldwide assessment of social Polistine nesting behavior. Ecology. 2018;99(10):2405-.

24. Richards OW. THE BIOLOGY OF THE SOCIAL WASPS (HYMENOPTERA, VESPIDAE). Biological Reviews. 1971;46(4):483–528.

25. Greene A. Dolichovespula and vespula. The social biology of wasps: Cornell University Press; 1991. p. 263–305.

26. Matsuura M. Vespa and provespa. The social biology of wasps: Cornell University Press; 1991. p. 232–62.

27. Valavi R, Guillera-Arroita G, Lahoz-Monfort JJ, Elith J. Predictive performance of presence-only species distribution models: a benchmark study with reproducible code. Ecological Monographs. 2022;92(1):e01486.

28. Secretariat G. GBIF science review 2020. 2021.

29. Chandler M, See L, Copas K, Bonde AMZ, López BC, Danielsen F, et al. Contribution of citizen science towards international biodiversity monitoring. Biological Conservation. 2017;213:280–94.

30. Larson ER, Graham BM, Achury R, Coon JJ, Daniels MK, Gambrell DK, et al. From eDNA to citizen science: emerging tools for the early detection of invasive species. Frontiers in Ecology and the Environment. 2020;18(4):194–202.

31. Gallo T, Waitt D. Creating a Successful Citizen Science Model to Detect and Report Invasive Species. BioScience. 2011;61(6):459–65.

32. Werkowska W, Márquez AL, Real R, Acevedo P. A practical overview of transferability in species distribution modeling. Environmental Reviews. 2017;25(1):127–33.

33. Liu C, Wolter C, Courchamp F, Roura-Pascual N, Jeschke JM. Biological invasions reveal how niche change affects the transferability of species distribution models. Ecology. 2022;103(8):e3719.

34. GBIF Occurrence Download [Internet]. 22 November 2024.

35. Miller SE, Sheehan MJ. Ecogeographical patterns of body size differ among North American paper wasp species. Insectes Sociaux. 2021;68:109–22.

36. Fick SE, Hijmans RJ. WorldClim 2: new 1-km spatial resolution climate surfaces for global land areas. International Journal of Climatology. 2017;37(12):4302–15.

37. Riahi K, van Vuuren DP, Kriegler E, Edmonds J, O’Neill BC, Fujimori S, et al. The Shared Socioeconomic Pathways and their energy, land use, and greenhouse gas emissions implications: An overview. Global Environmental Change. 2017;42:153–68.

38. Vuillaume B, Valette V, Lepais O, Grandjean F, Breuil M. Genetic Evidence of Hybridization between the Endangered Native Species Iguana delicatissima and the Invasive Iguana iguana (Reptilia, Iguanidae) in the Lesser Antilles: Management Implications. PLOS ONE. 2015;10(6):e0127575.

39. Matthews J, Van der Velde G, Bij de Vaate A, Collas FPL, Koopman KR, Leuven RSEW. Rapid range expansion of the invasive quagga mussel in relation to zebra mussel presence in The Netherlands and Western Europe. Biological Invasions. 2014;16(1):23–42.

40. Kuinkel S, Miller SE. Niche Dynamics and Climatic Novelty Drive the Invasion Success of the European Paper Wasp Across North America. Journal of Biogeography. 2026;53(5):e70267.

41. Meffe GK, Carroll CR, Groom M. Principles of Conservation Biology, 3rd Edition. Martha J. Groom, Gary K. Meffe, C. Ronald Carroll. 2006. Sinauer Associates. Sunderland, MA2006.

42. Lester PJ, Beggs JR. Invasion Success and Management Strategies for Social Vespula Wasps. Annual Review of Entomology. 2019;64(Volume 64, 2019):51–71.

43. Xu M, Li Z, Shi Y, Zhang X, Jiang S. Evolution of regional inequality in the global shipping network. Journal of Transport Geography. 2015;44:1–12.

44. Konstantinus A, Woxenius J. Case study: Coastal shipping in sub-Saharan Africa. Case Studies on Transport Policy. 2022;10(4):2064–74.

45. Lioy S, Bergamino C, Porporato M. The invasive hornet *Vespa velutina*: distribution, impacts and management options. CABI Reviews. 2022.

46. Nave A, Godinho J, Fernandes J, Garcia AI, Ferreira Golpe MA, Branco M. *Vespa velutina*: a menace for Western Iberian fruit production. Cogent Food & Agriculture. 2024;10(1):2313679.

47. Kimsey LS, Carpenter JM. The Vespinae of North America (Vespidae, Hymenoptera). Journal of Hymenoptera Research. 2012;28:37–65.

48. Hoebeke ER, Bartlett LJ, Evans M, Freeman BE, Wares JP. First Records of *Vespa velutina* (Lepeletier) (Color form *Nigrithorax*) (Hymenoptera: Vespidae) in North America, an Invasive Pest of Domesticated Honeybees. Proceedings of the Entomological Society of Washington. 2024;126(2):193–205, 13.

49. Norderud ED, Powell SL, Peterson RKD. Risk Assessment for the Establishment of Vespa mandarinia (Hymenoptera: Vespidae) in the Pacific Northwest, United States. Journal of Insect Science. 2021;21(4).

50. Alaniz AJ, Carvajal MA, Vergara PM. Giants are coming? Predicting the potential spread and impacts of the giant Asian hornet (Vespa mandarinia, Hymenoptera: Vespidae) in the USA. Pest Management Science. 2021;77(1):104–12.

51. Nuñez-Penichet C, Osorio-Olvera L, Gonzalez VH, Cobos ME, Jiménez L, DeRaad DA, et al. Geographic potential of the world’s largest hornet, Vespa mandarinia Smith (Hymenoptera: Vespidae), worldwide and particularly in North America. PeerJ. 2021;9:e10690.

52. Wilson TM, Takahashi J, Spichiger S-E, Kim I, van Westendorp P. First Reports of Vespa mandarinia (Hymenoptera: Vespidae) in North America Represent Two Separate Maternal Lineages in Washington State, United States, and British Columbia, Canada. Annals of the Entomological Society of America. 2020;113(6):468–72.

53. Pocock MJO, Tweddle JC, Savage J, Robinson LD, Roy HE. The diversity and evolution of ecological and environmental citizen science. PLOS ONE. 2017;12(4):e0172579.

