## Supplementary material for "Large scale application of species distribution models to predict future vulnerability to social wasp invasions": Figure S1

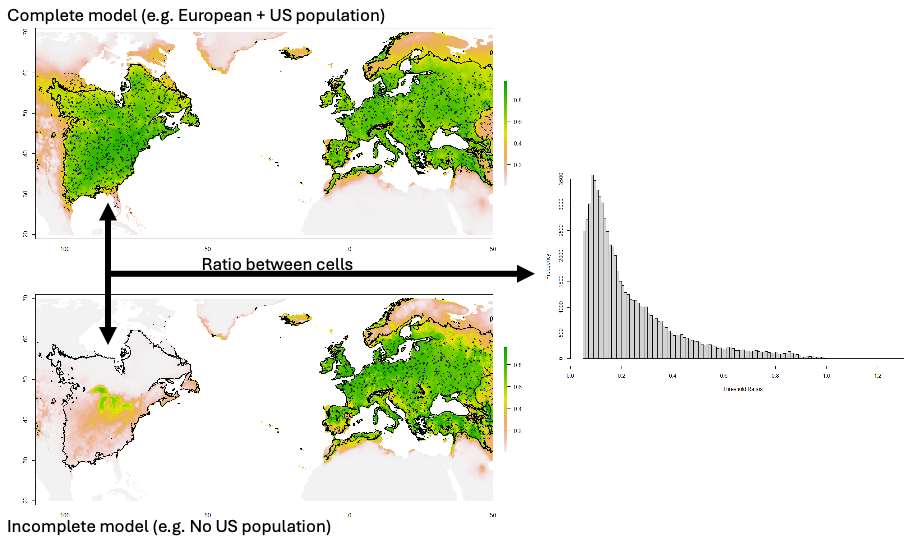
**Figure S1:** Diagram of the transferability correction factor found in Methods section 2.3.

**Figure S2:** Species richness plots by genus for changes in invasively suitable habitat under various climate change scenarios.

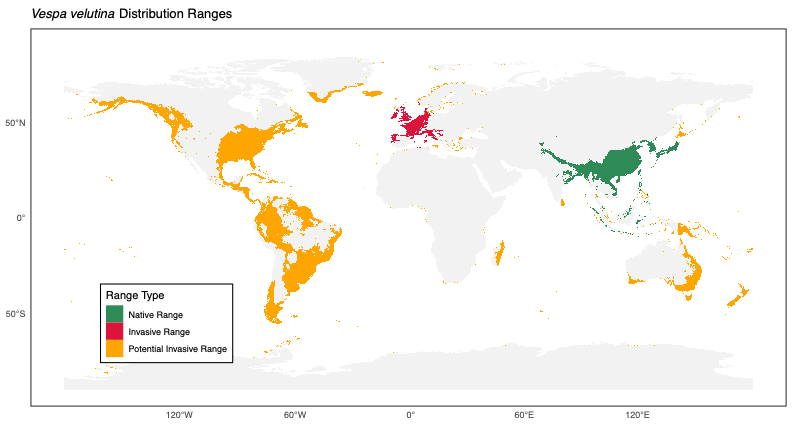
**Figure S3:** Species Distributions for all species in this study (including *Vespa velutina*). Area is delineated into native range, invasive range (if present) and potential invasive range.

**Table S1:** Risk Scores of all social Vespids in this study. The size of Native Areas, Invaded Areas and Potential Invaded Areas are used in the calculation of this score.
