## Supplementary material for "Large scale application of species distribution models to predict future vulnerability to social wasp invasions": Figure S2

*Agelaia angulata* Distribution Ranges

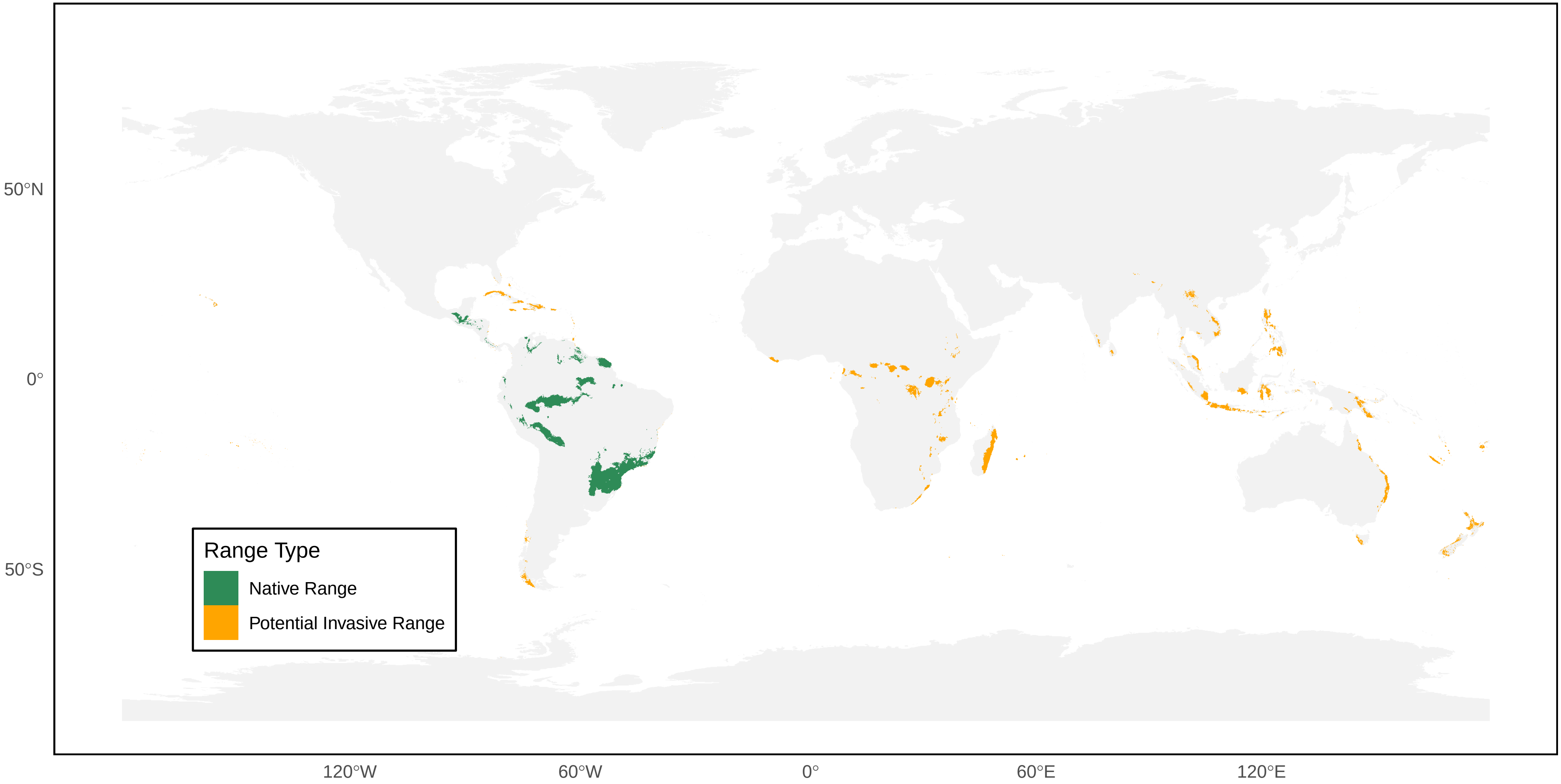

*Agelaia areata* Distribution Ranges

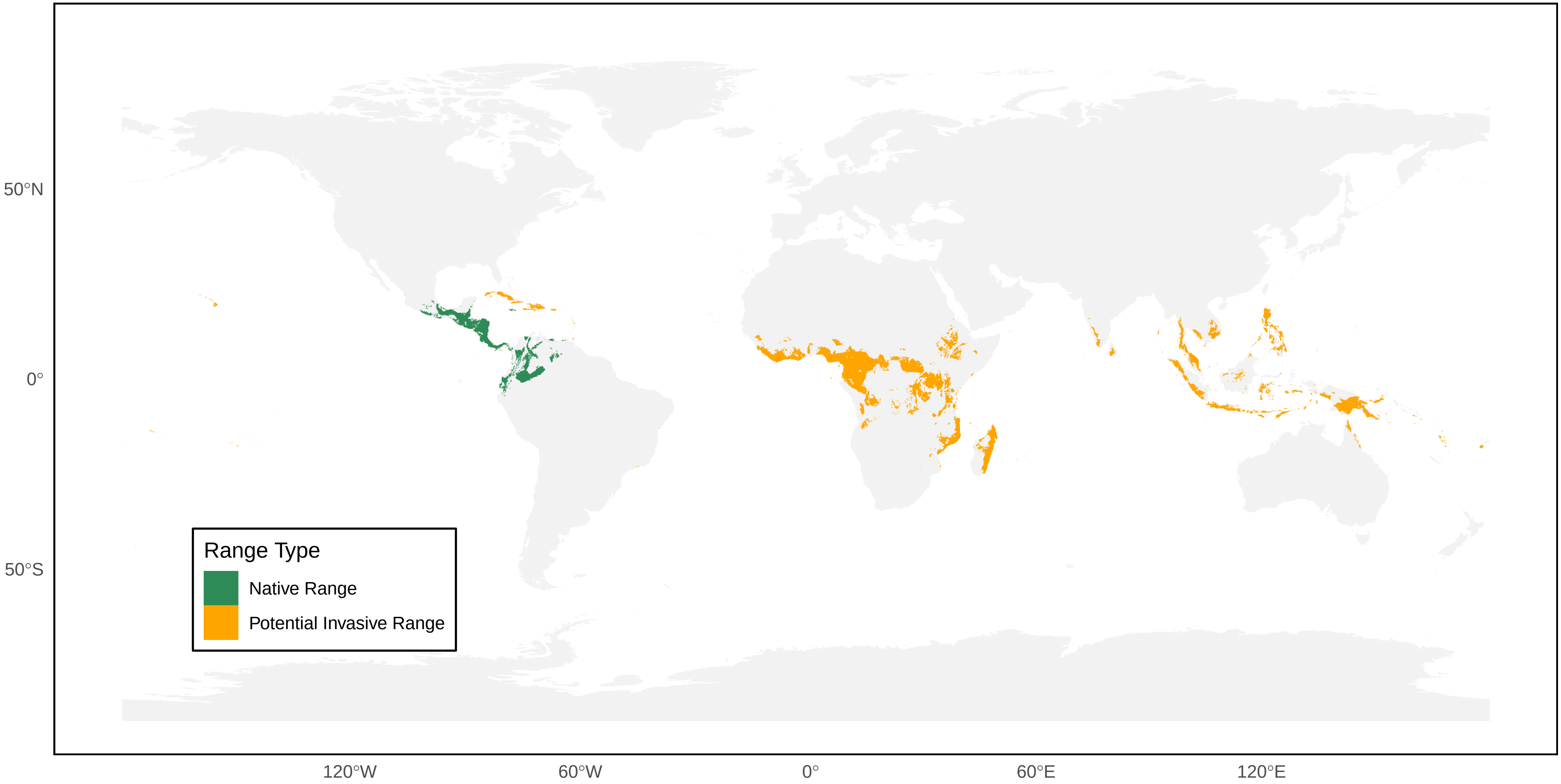

*Agelaia bequaerti* Distribution Ranges

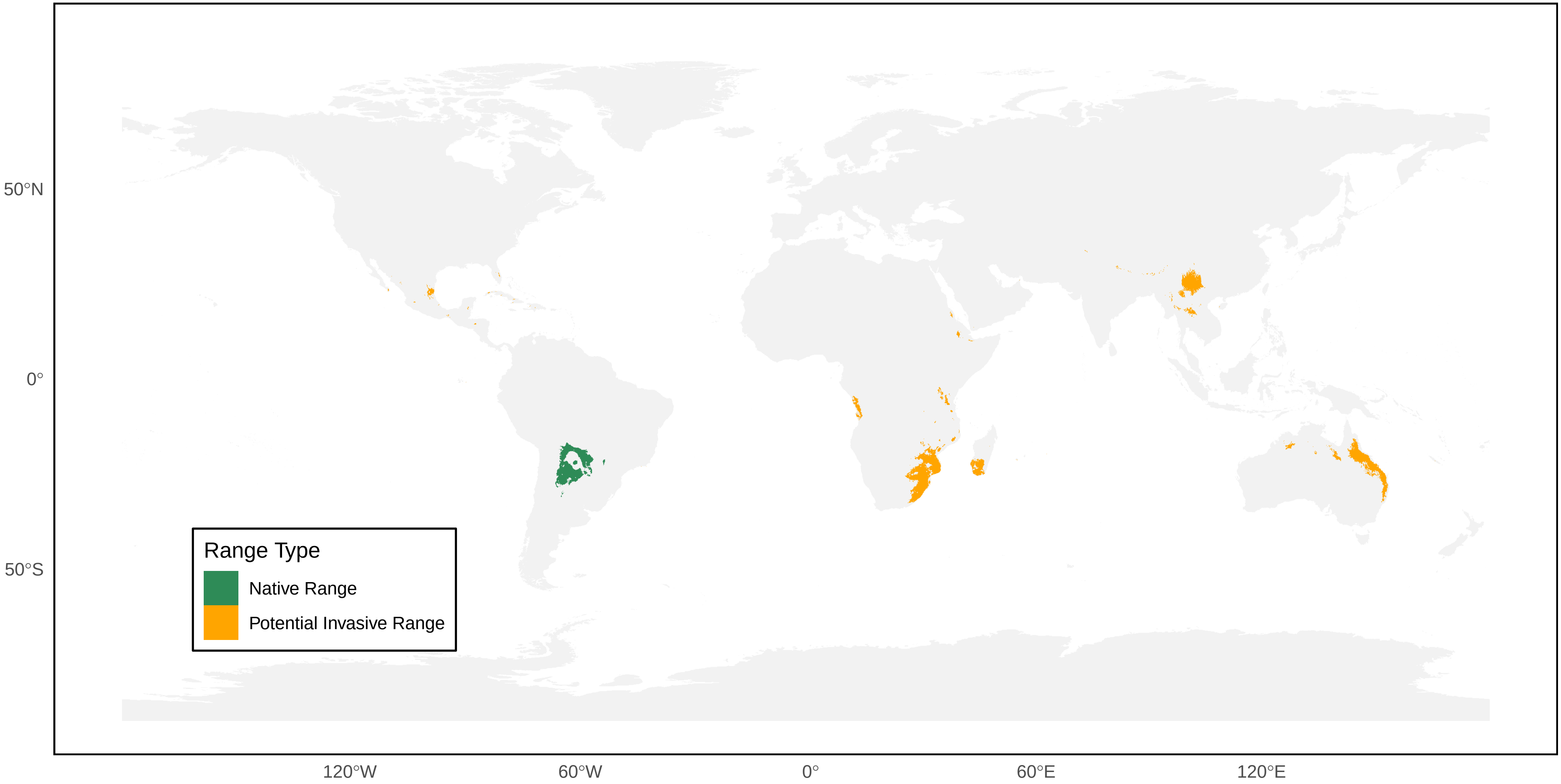

*Agelaia cajennensis* Distribution Ranges

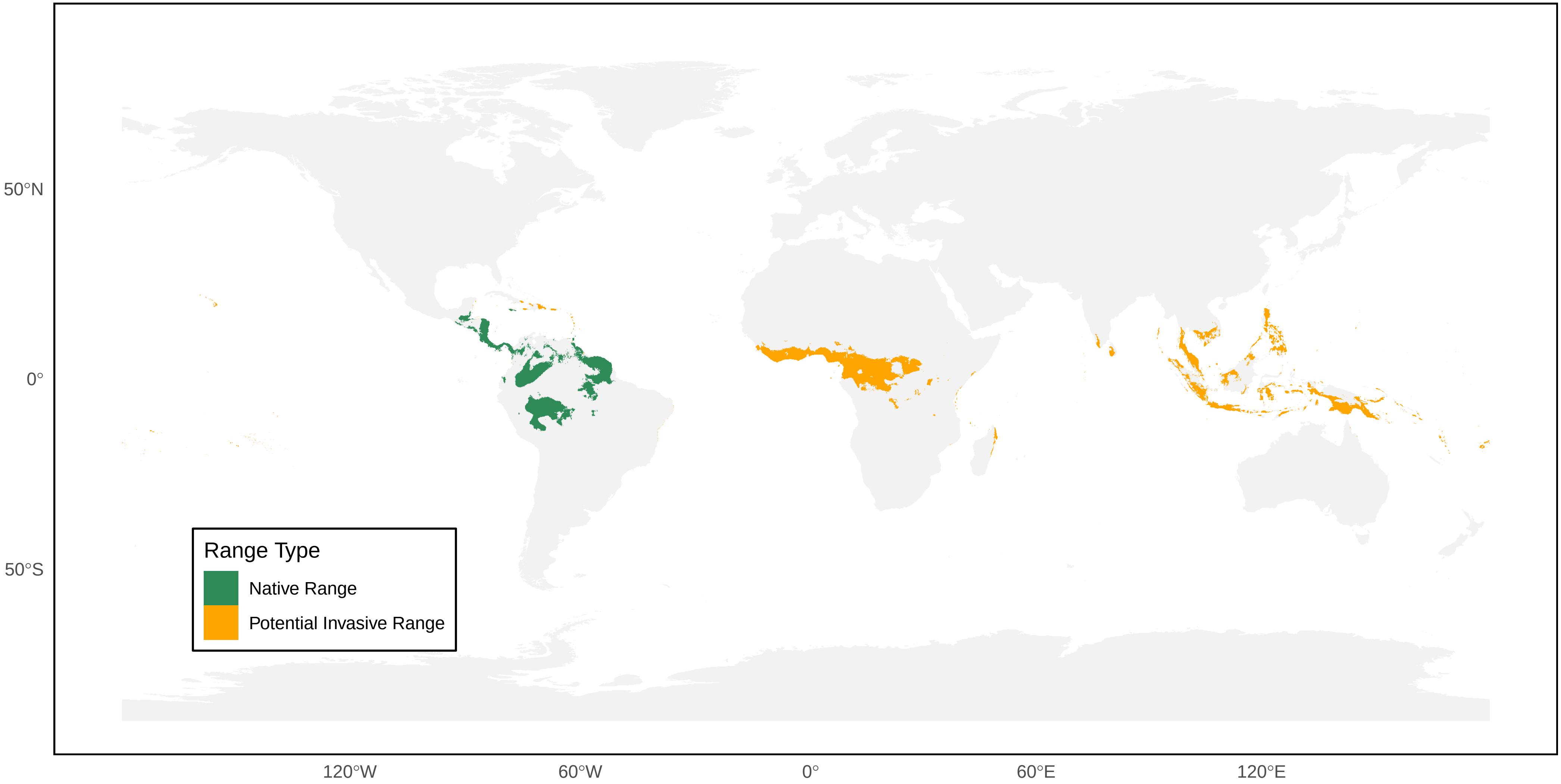

*Agelaia centralis* Distribution Ranges

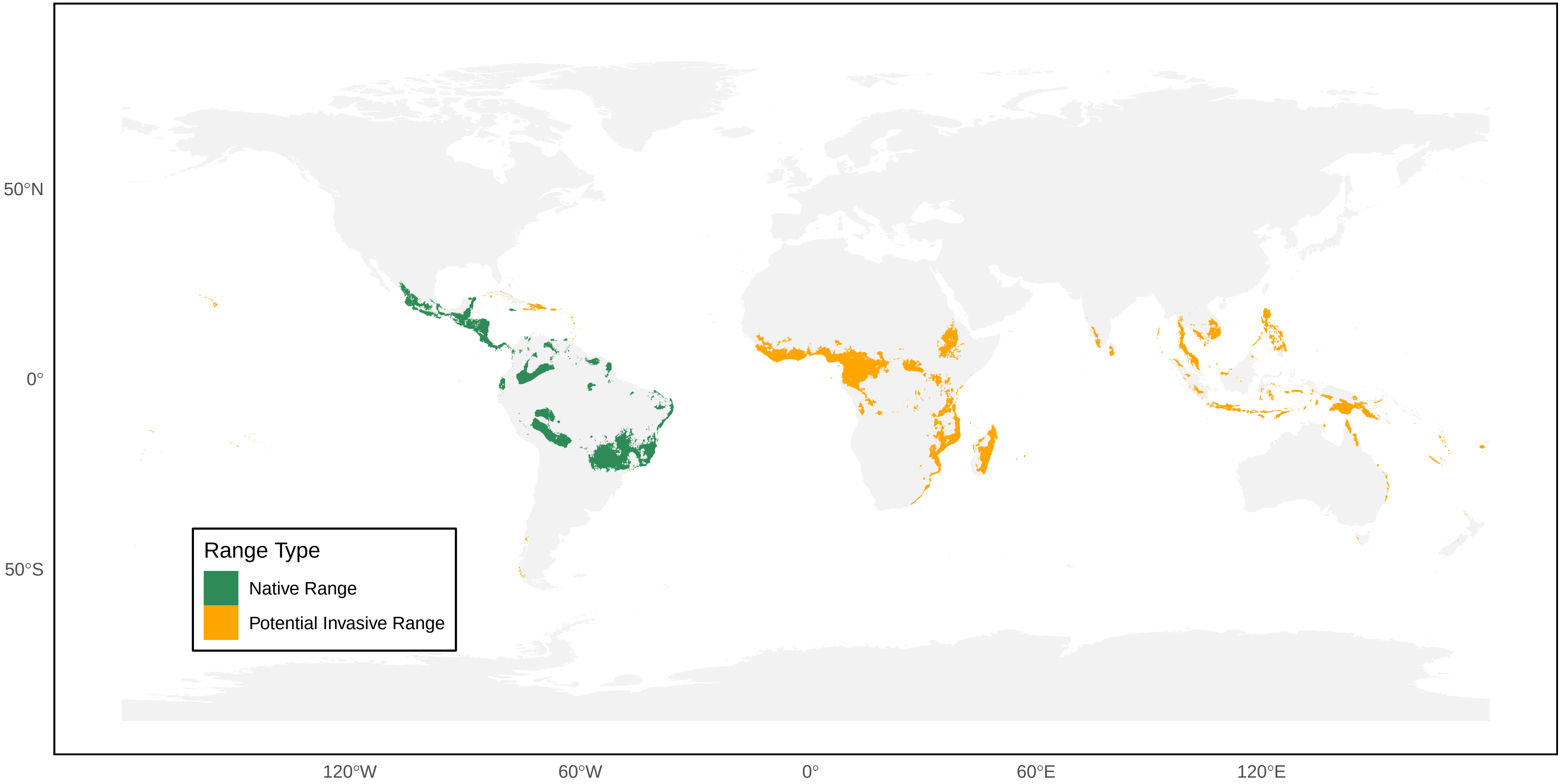

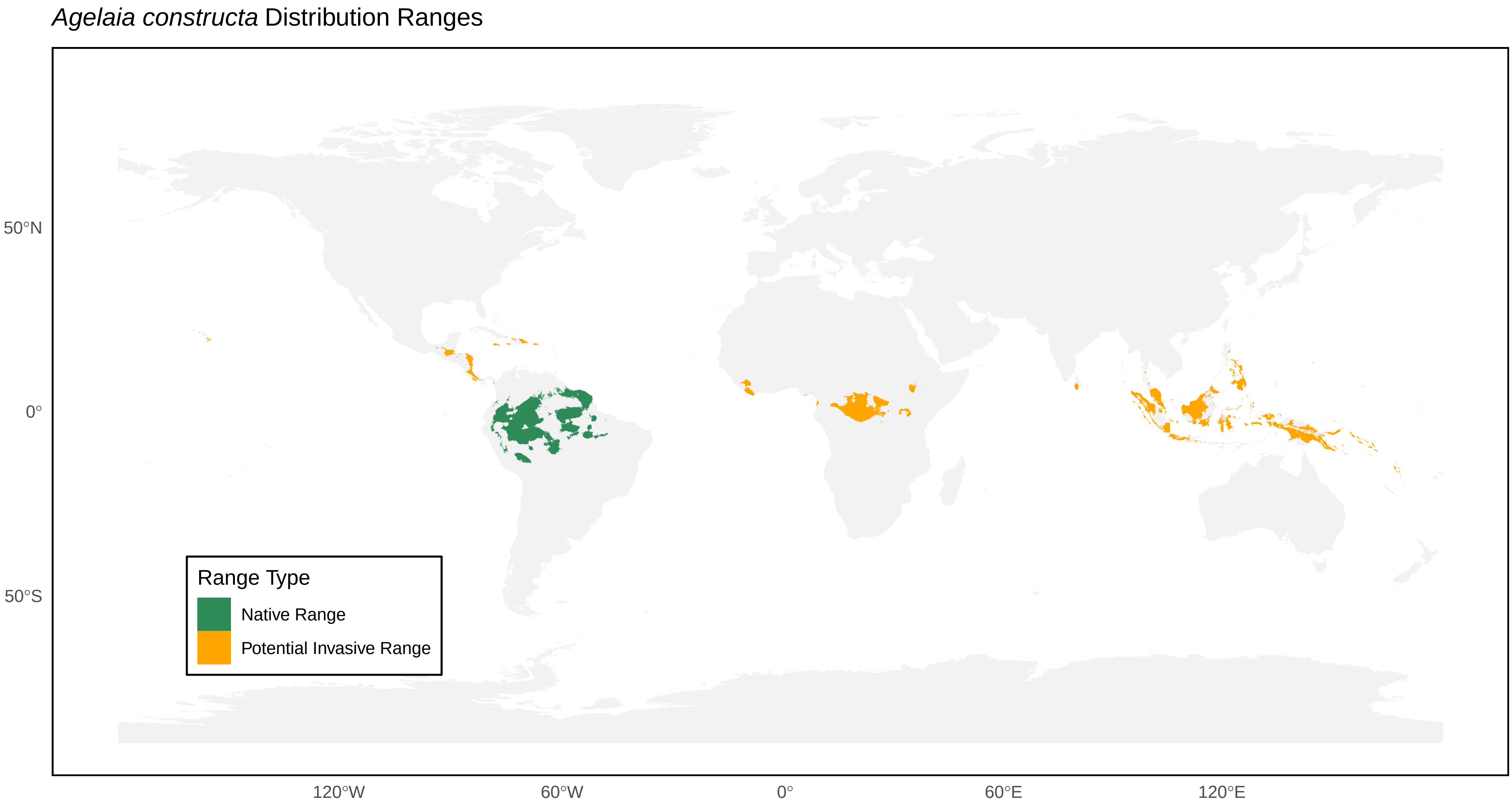

*Agelaia constructa* Distribution Ranges
50°N
0°
50°S
120°W
60°W
0°
60°E
120°E
Range Type
Native Range
Potential Invasive Range

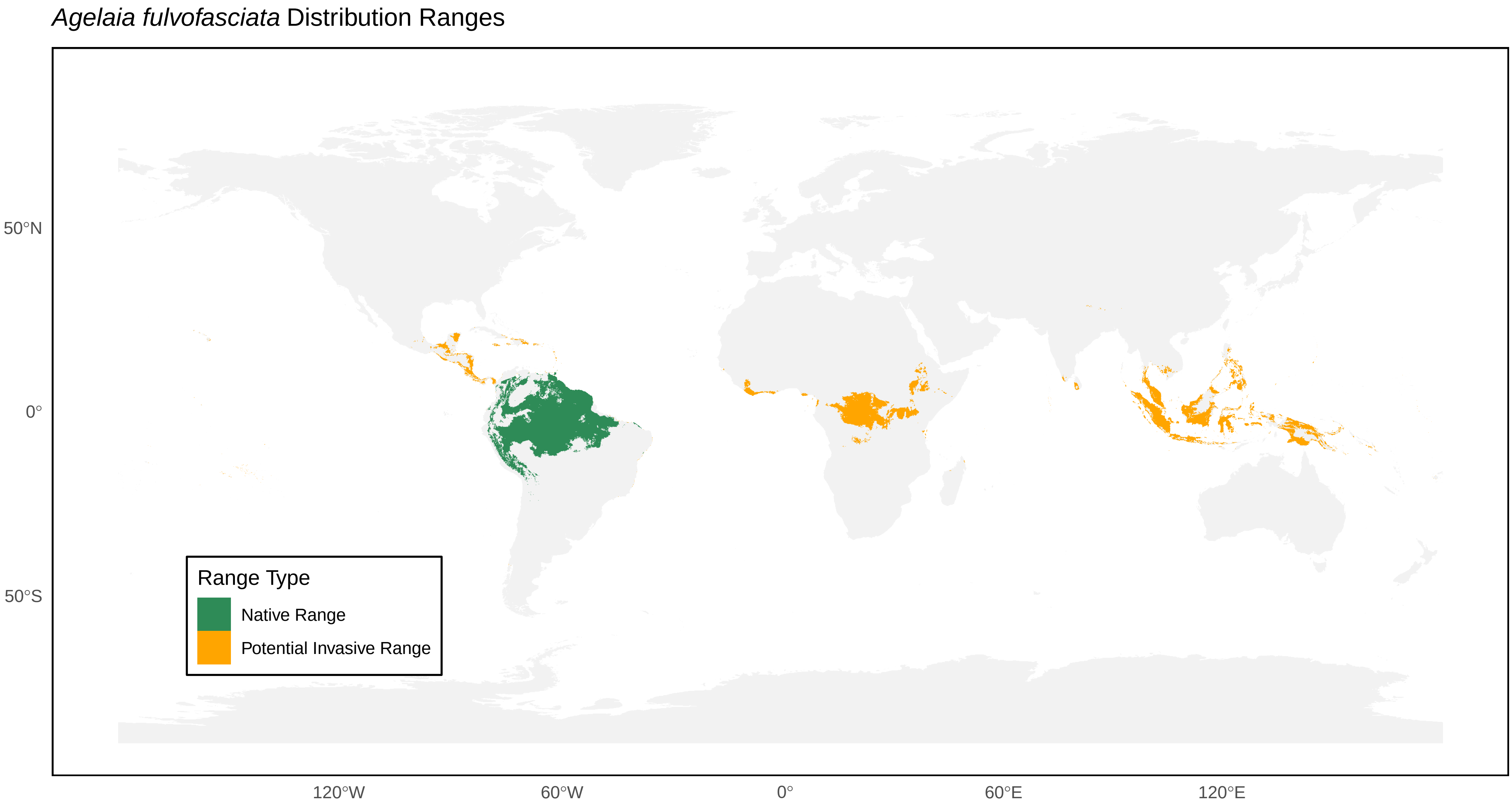
*Agelaia fulvofasciata* Distribution Ranges
50°N
0°
50°S
120°W
60°W
0°
60°E
120°E
Range Type
Native Range
Potential Invasive Range

*Agelaia melanopyga* Distribution Ranges

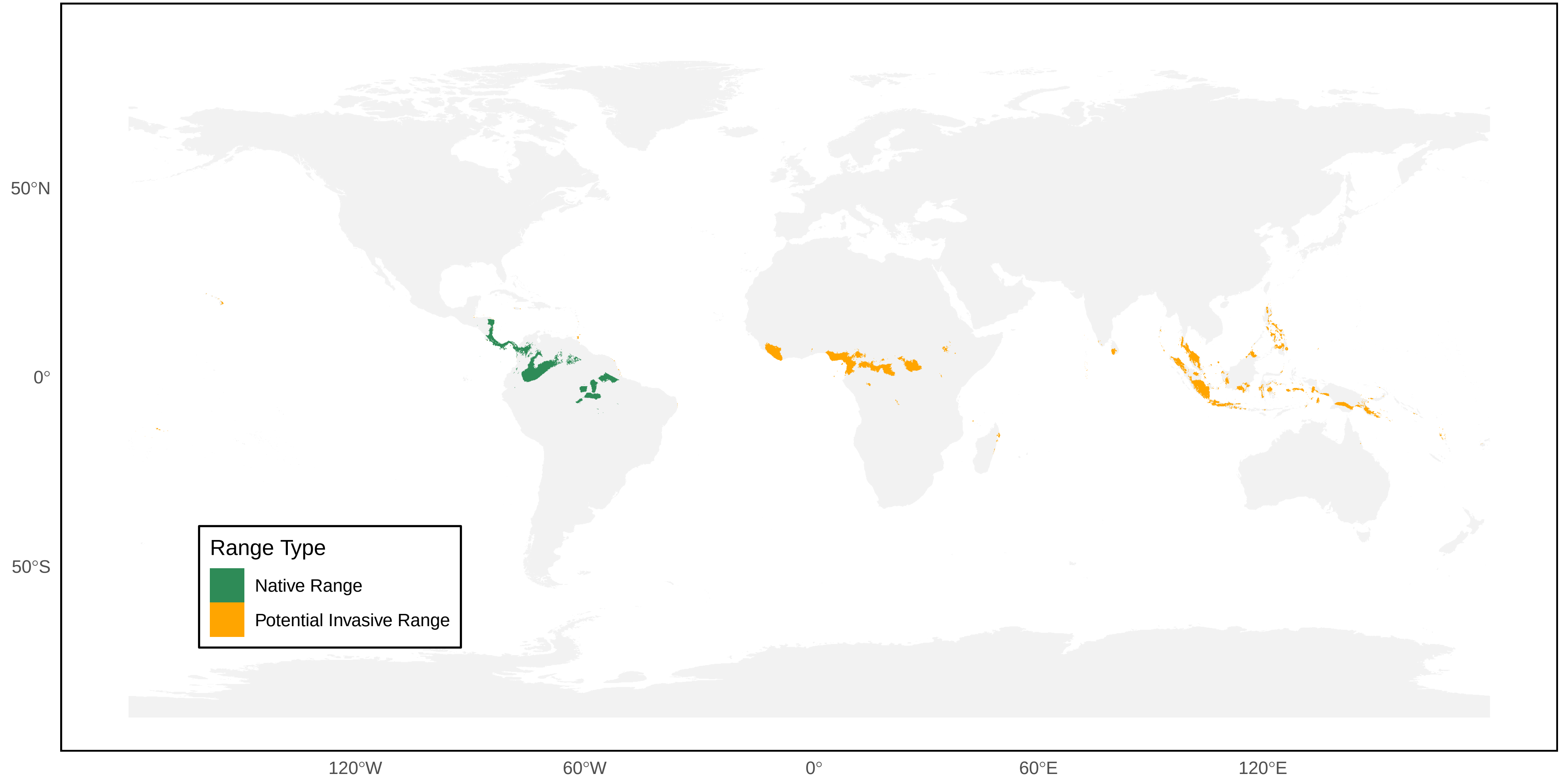

*Agelaia multipicta* Distribution Ranges

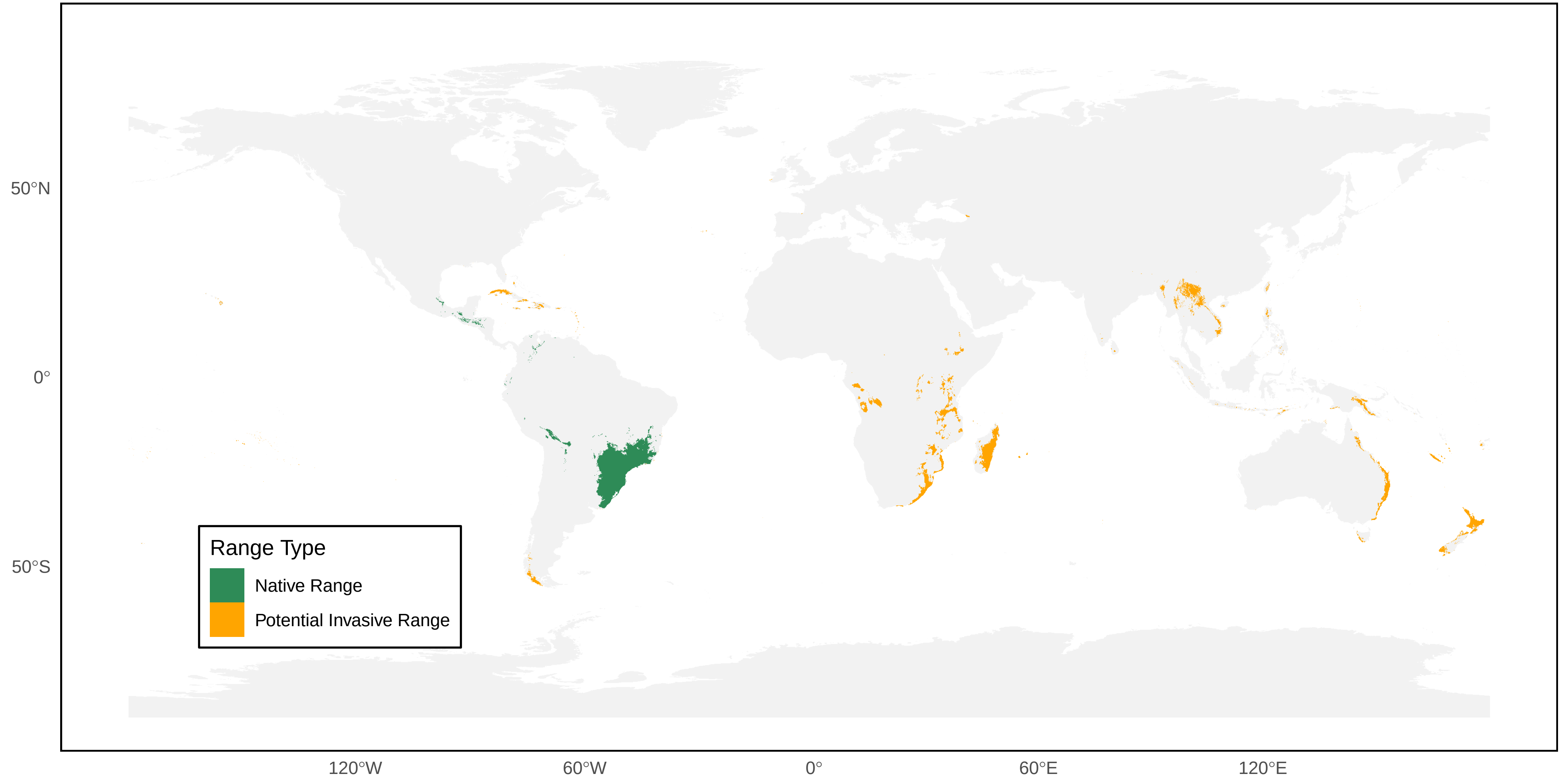

*Agelaia myrmecophila* Distribution Ranges

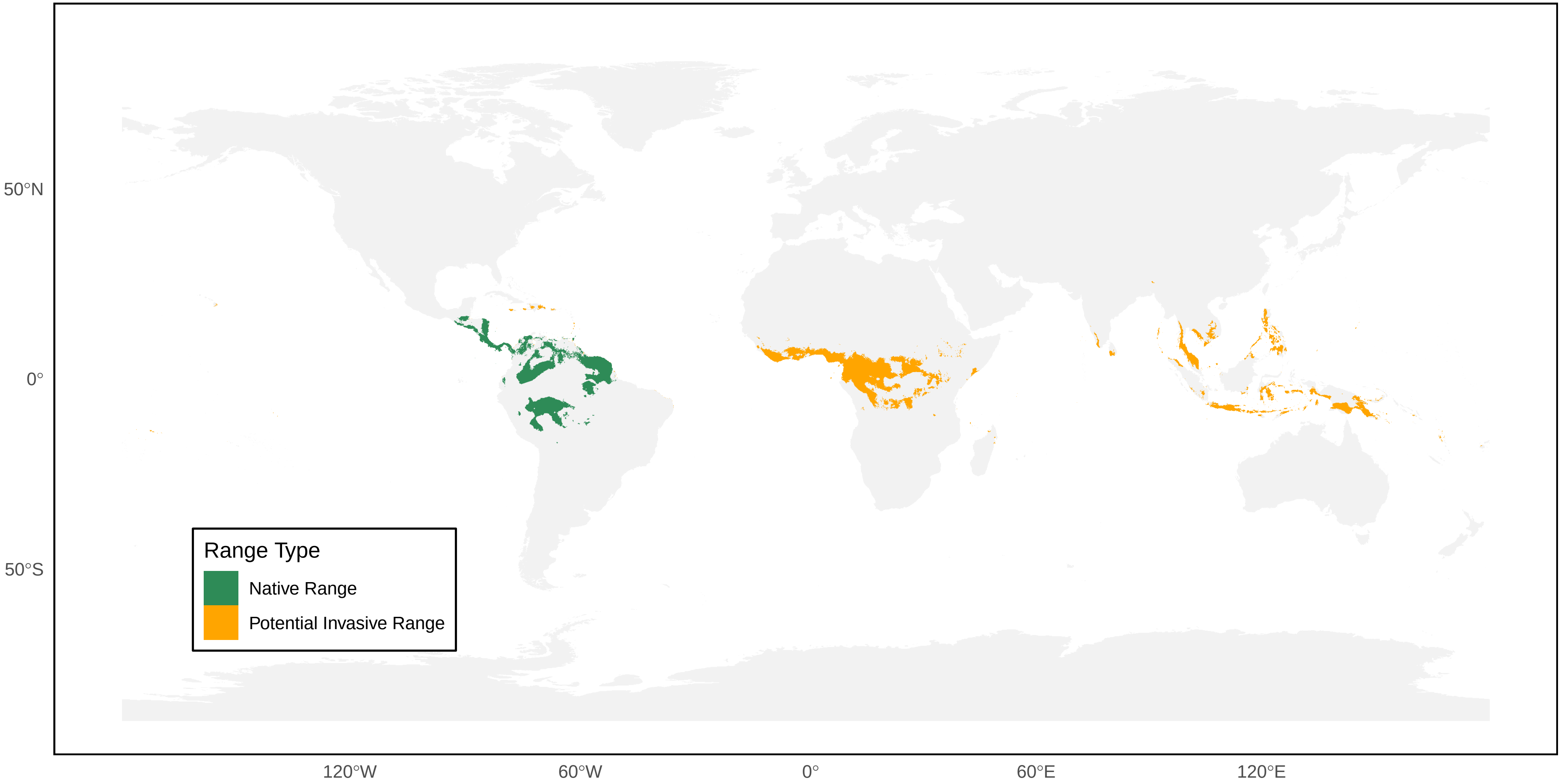

*Agelaia pallipes* Distribution Ranges

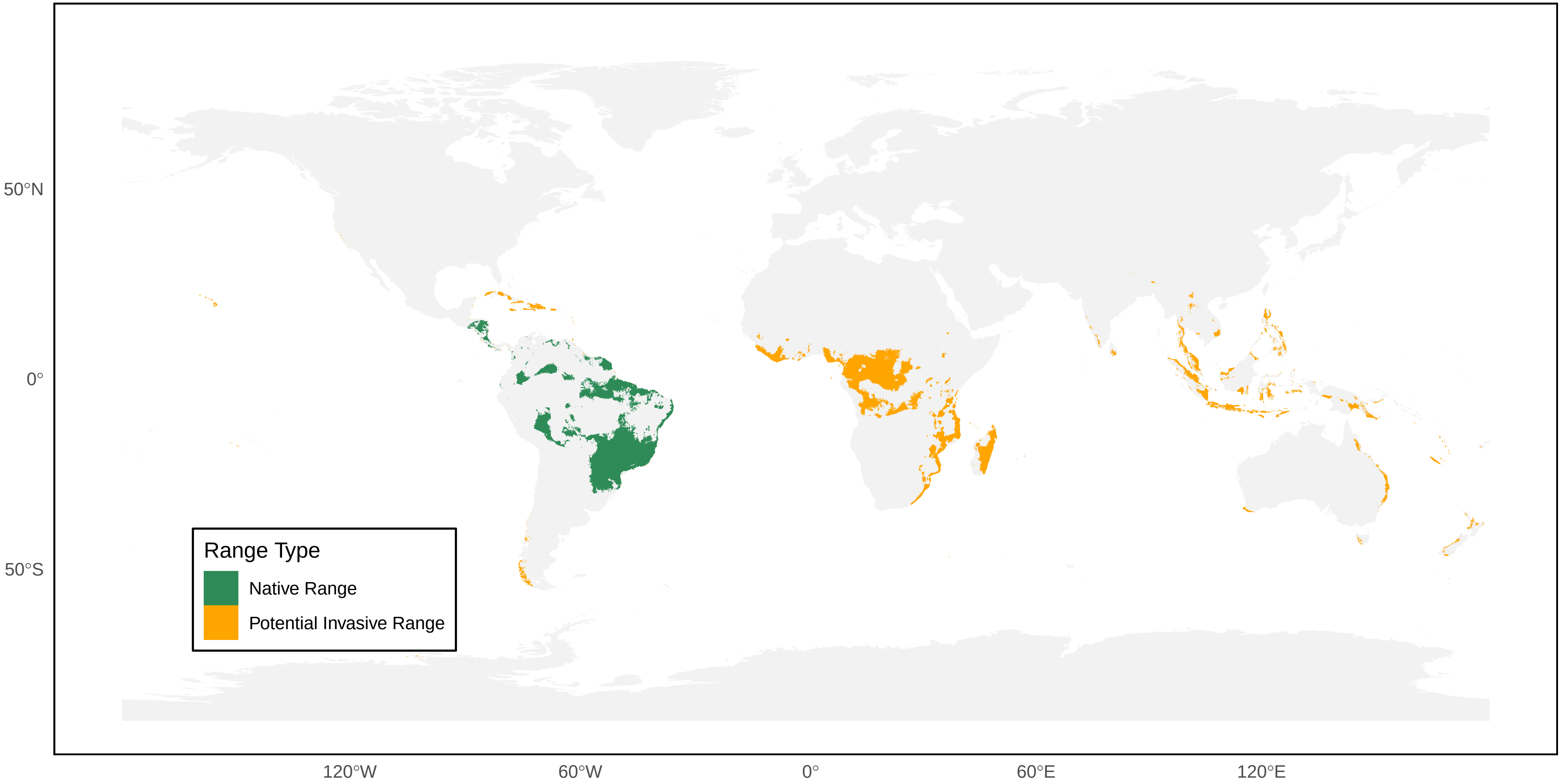

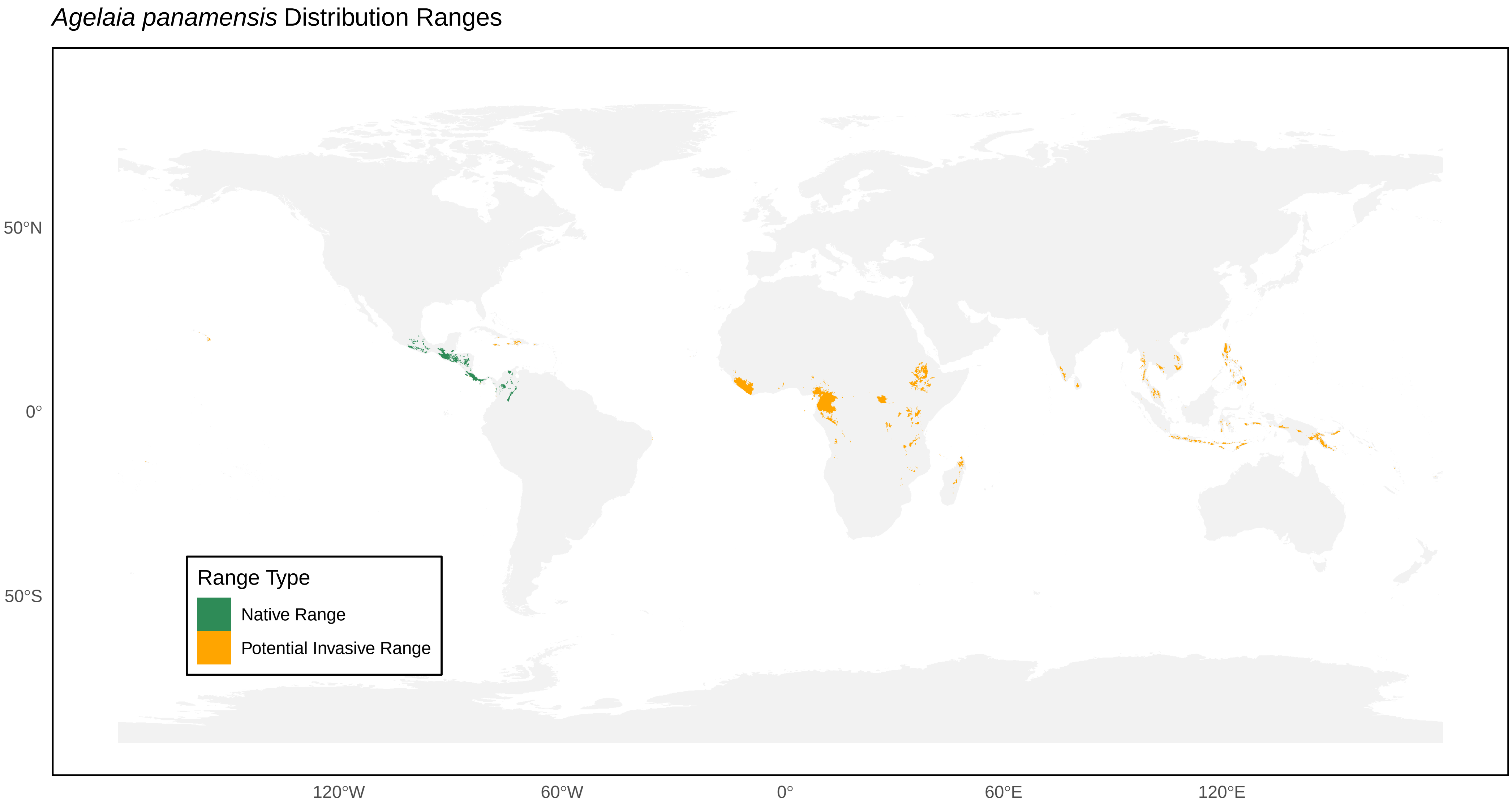

*Agelaia panamensis* Distribution Ranges
50°N
0°
50°S
120°W
60°W
0°
60°E
120°E
Range Type
Native Range
Potential Invasive Range

*Agelaia testacea* Distribution Ranges

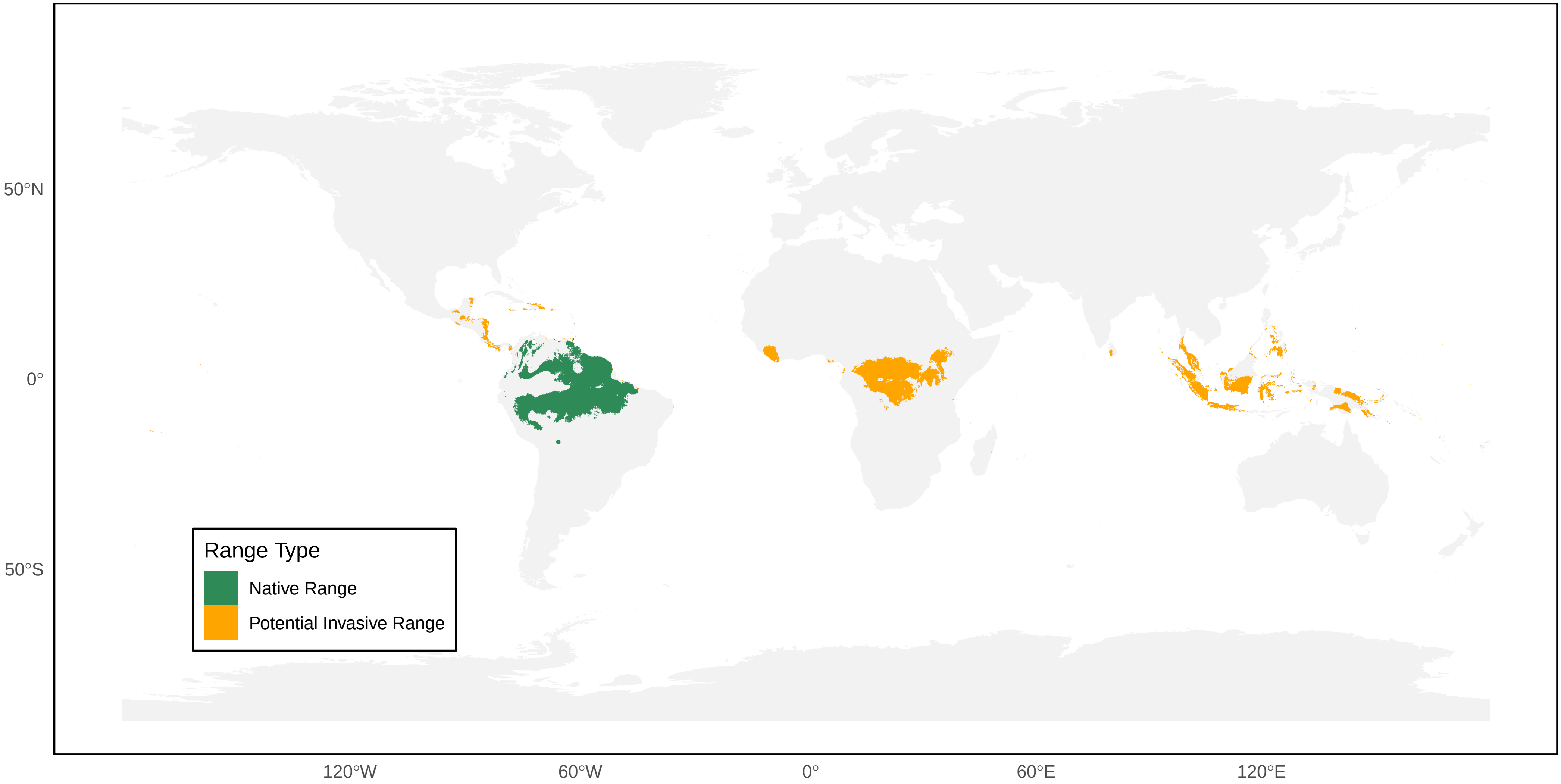

*Agelaia vicina* Distribution Ranges

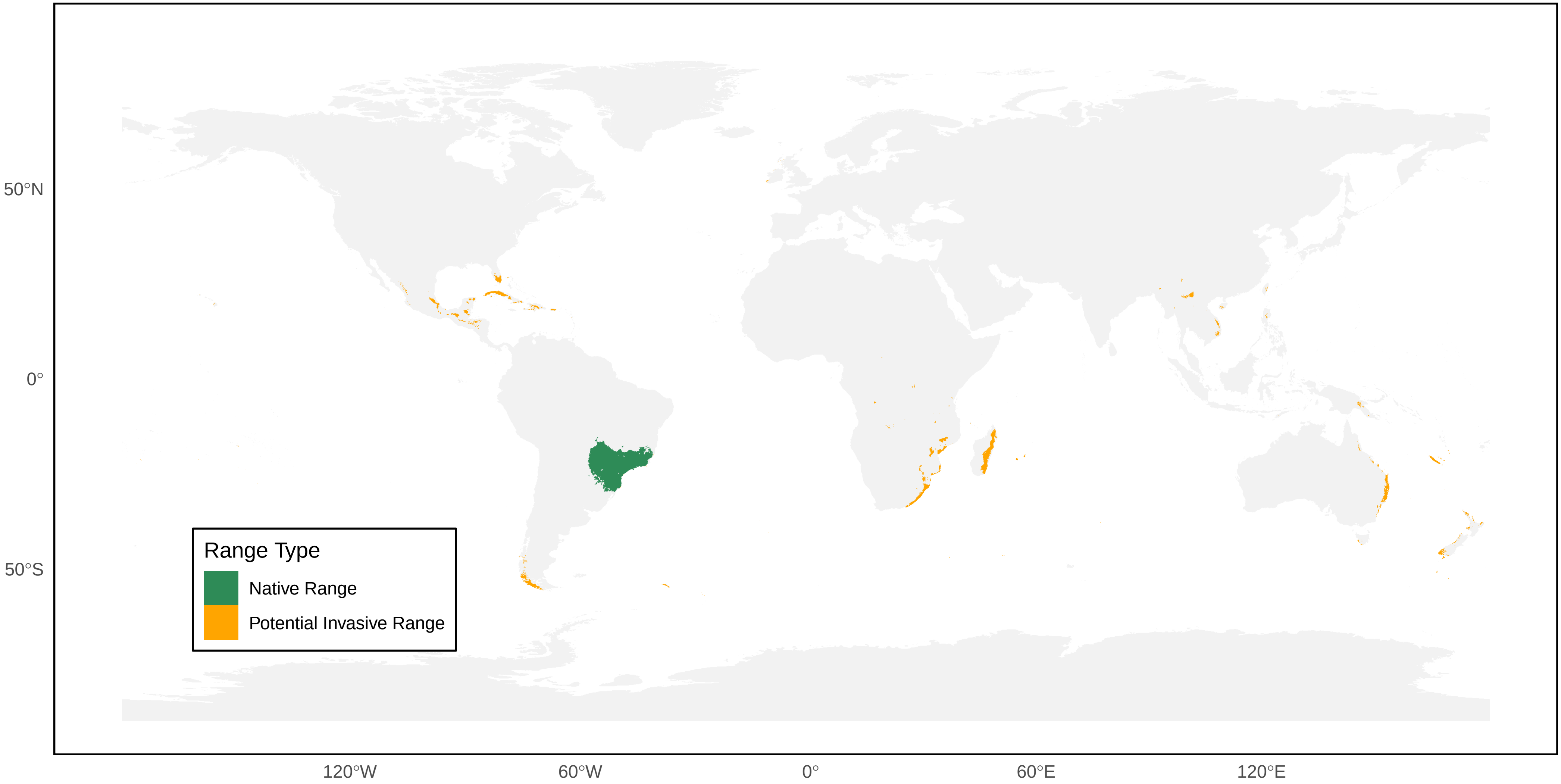

*Agelaia xanthopus* Distribution Ranges

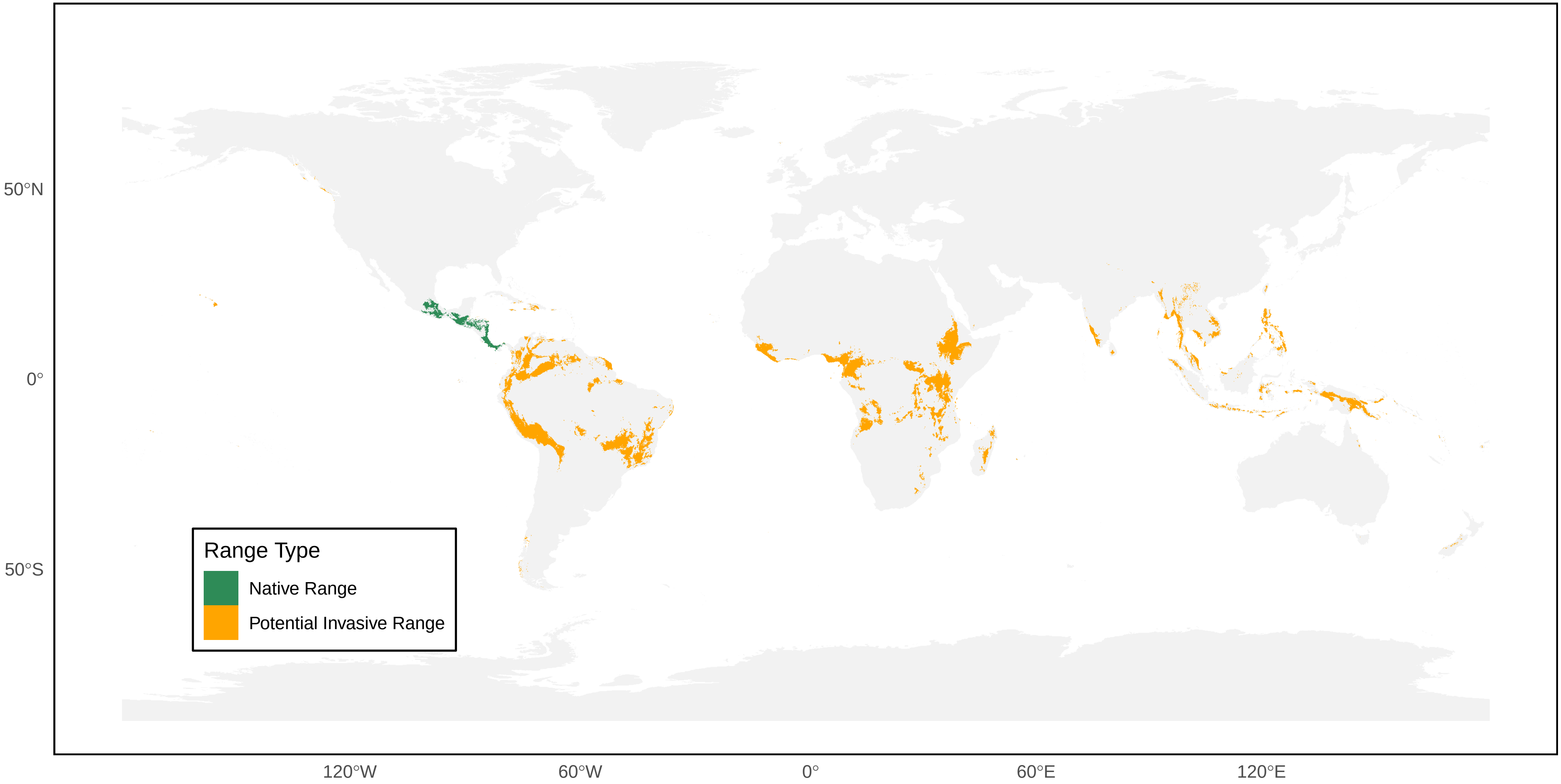

*Agelaia yepocapa* Distribution Ranges

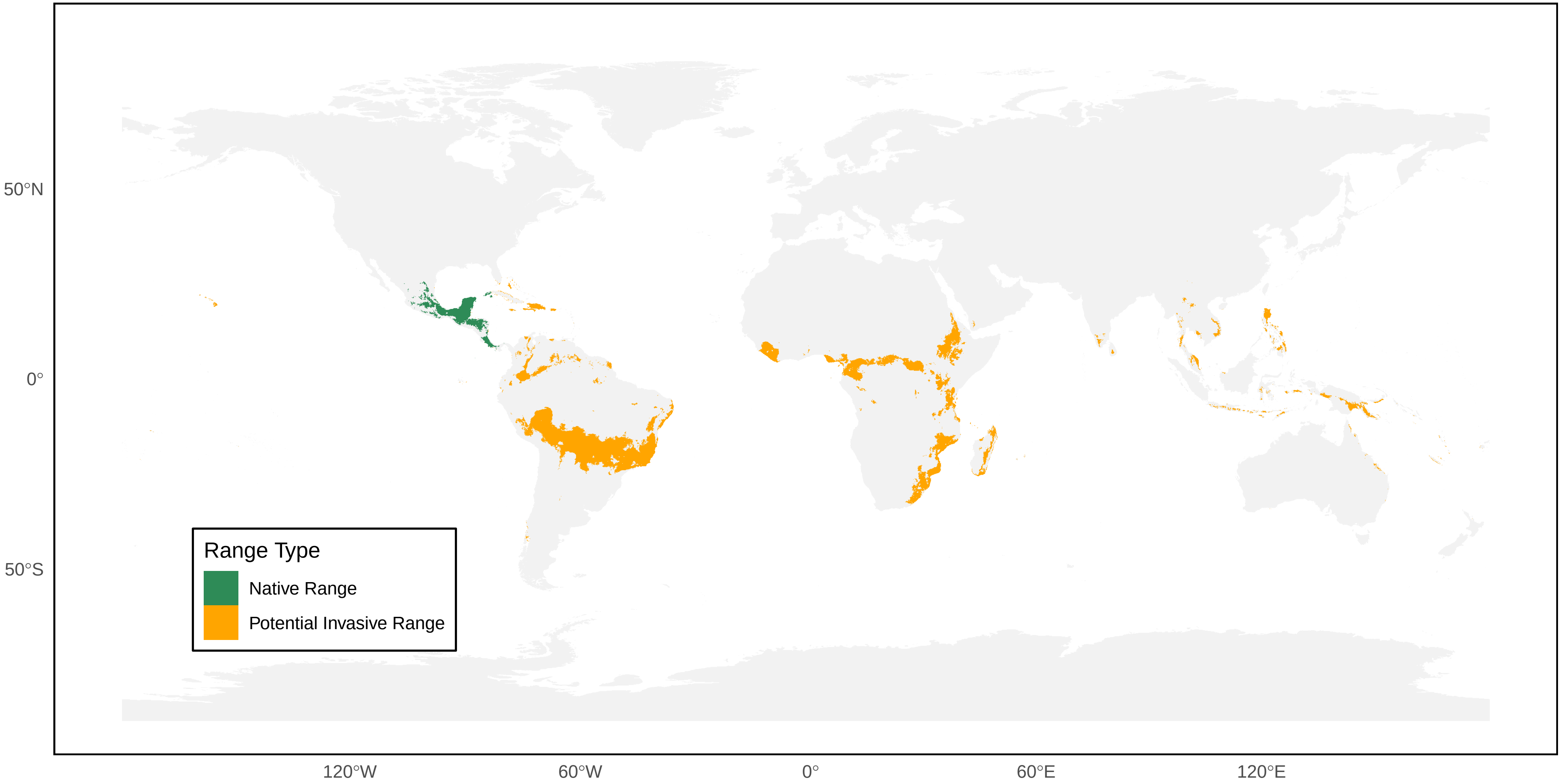

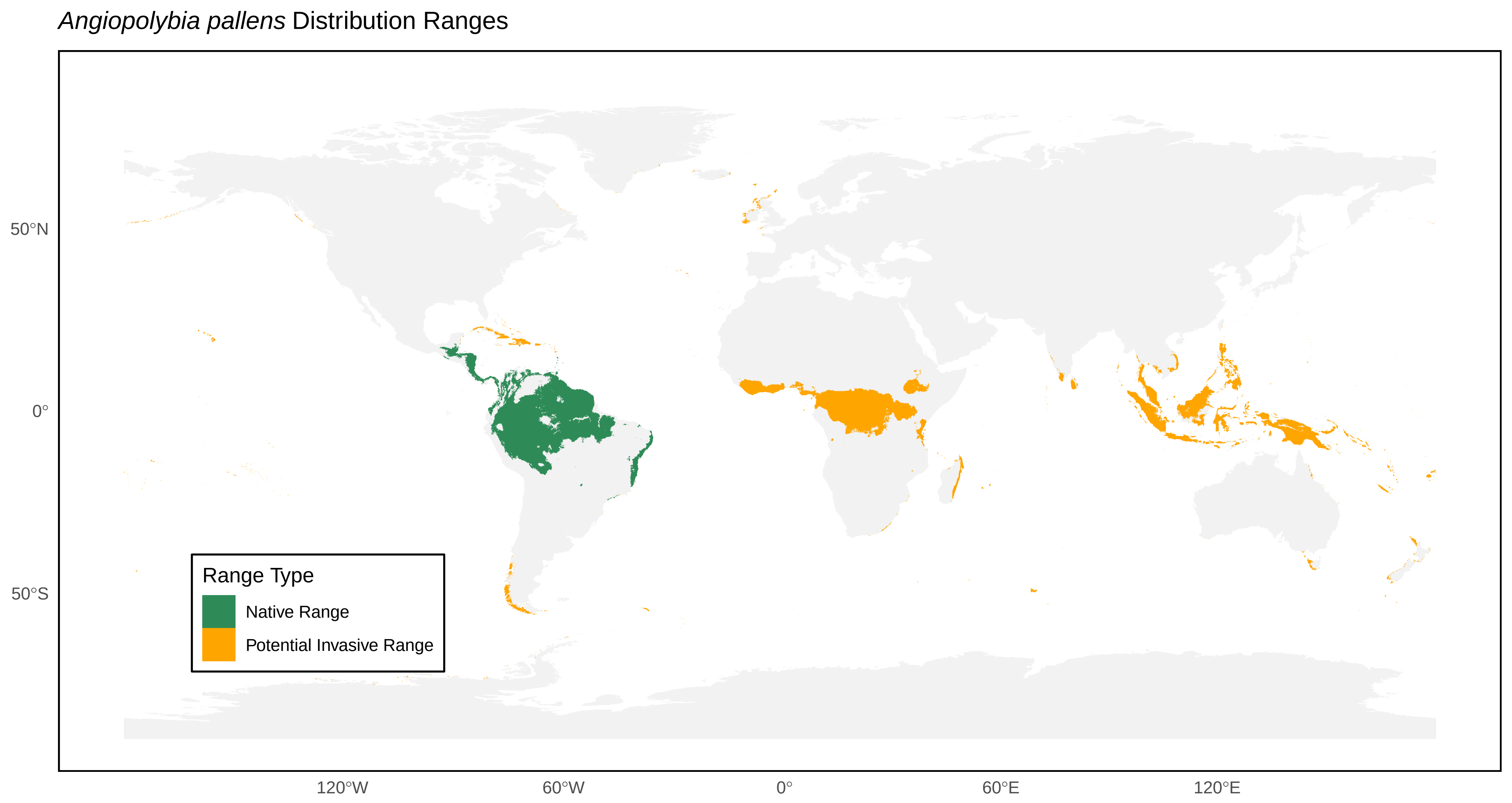
*Angiopolybia pallens* Distribution Ranges
50°N
0°
50°S
120°W
60°W
0°
60°E
120°E
Range Type
Native Range
Potential Invasive Range

*Angiopolybia paraensis* Distribution Ranges

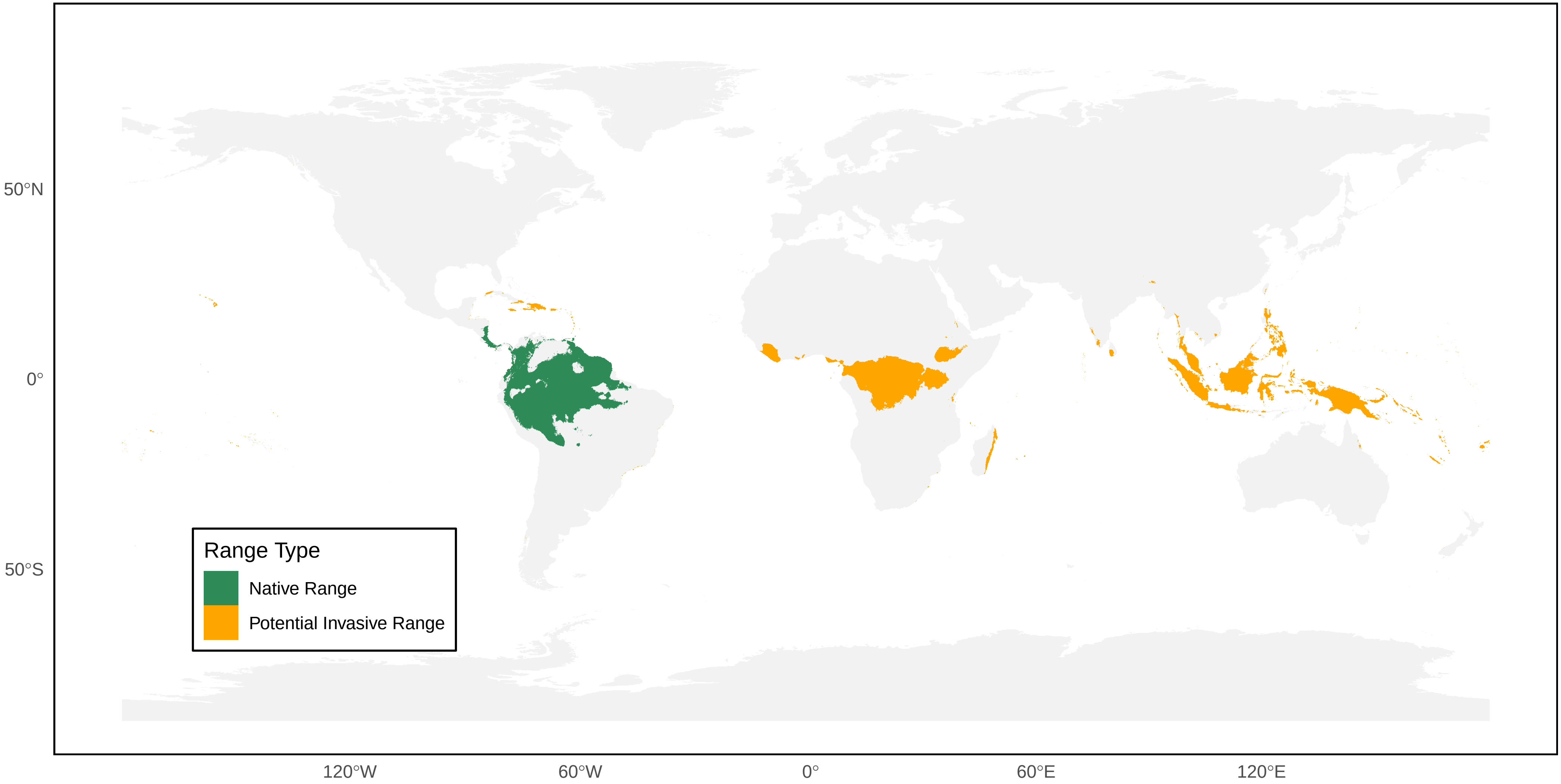

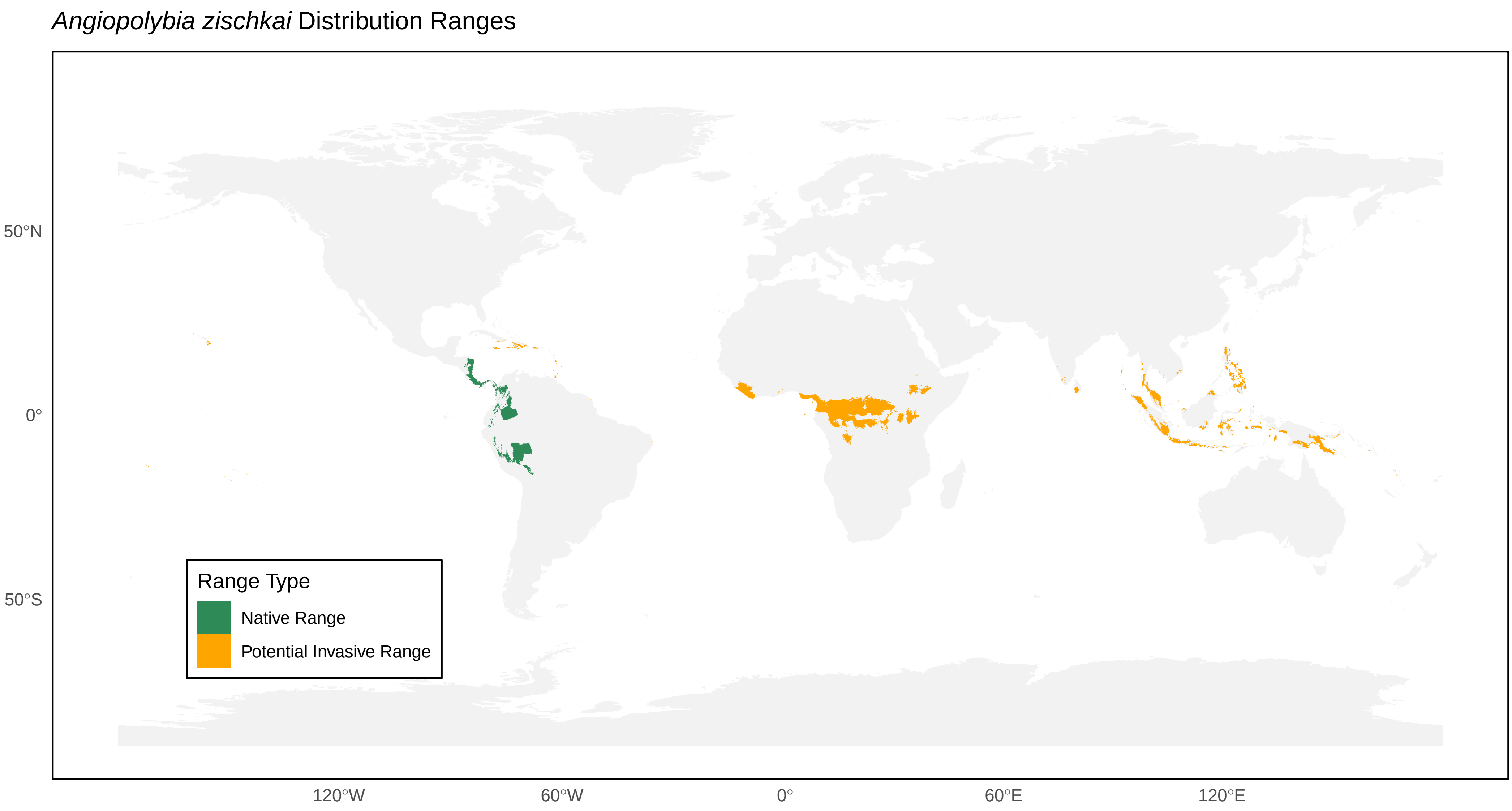
*Angiopolybia zischkai* Distribution Ranges
50°N
0°
50°S
120°W
60°W
0°
60°E
120°E
Range Type
Native Range
Potential Invasive Range

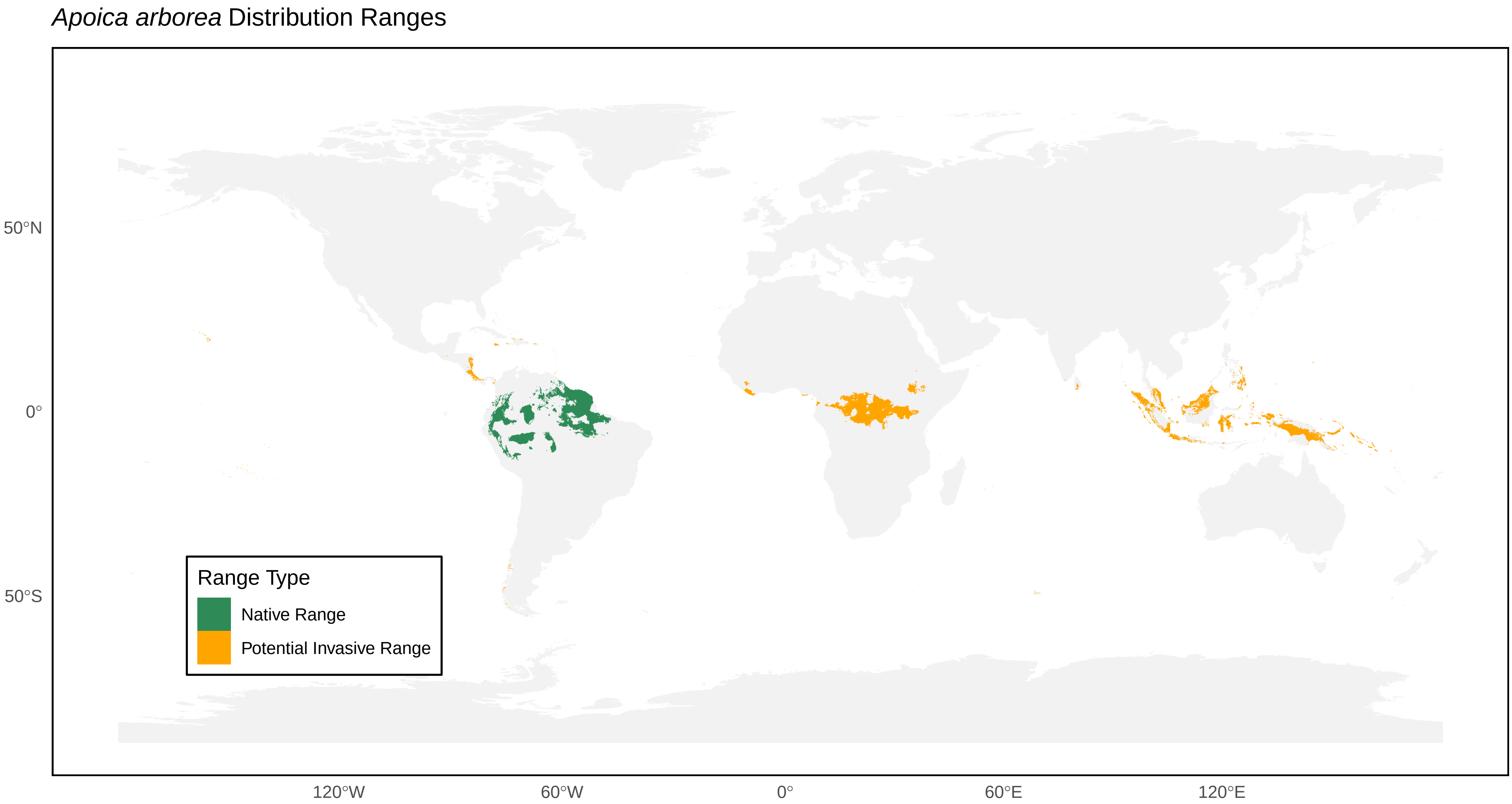
*Apoica arborea* Distribution Ranges
50°N
0°
50°S
120°W
60°W
0°
60°E
120°E
Range Type
Native Range
Potential Invasive Range

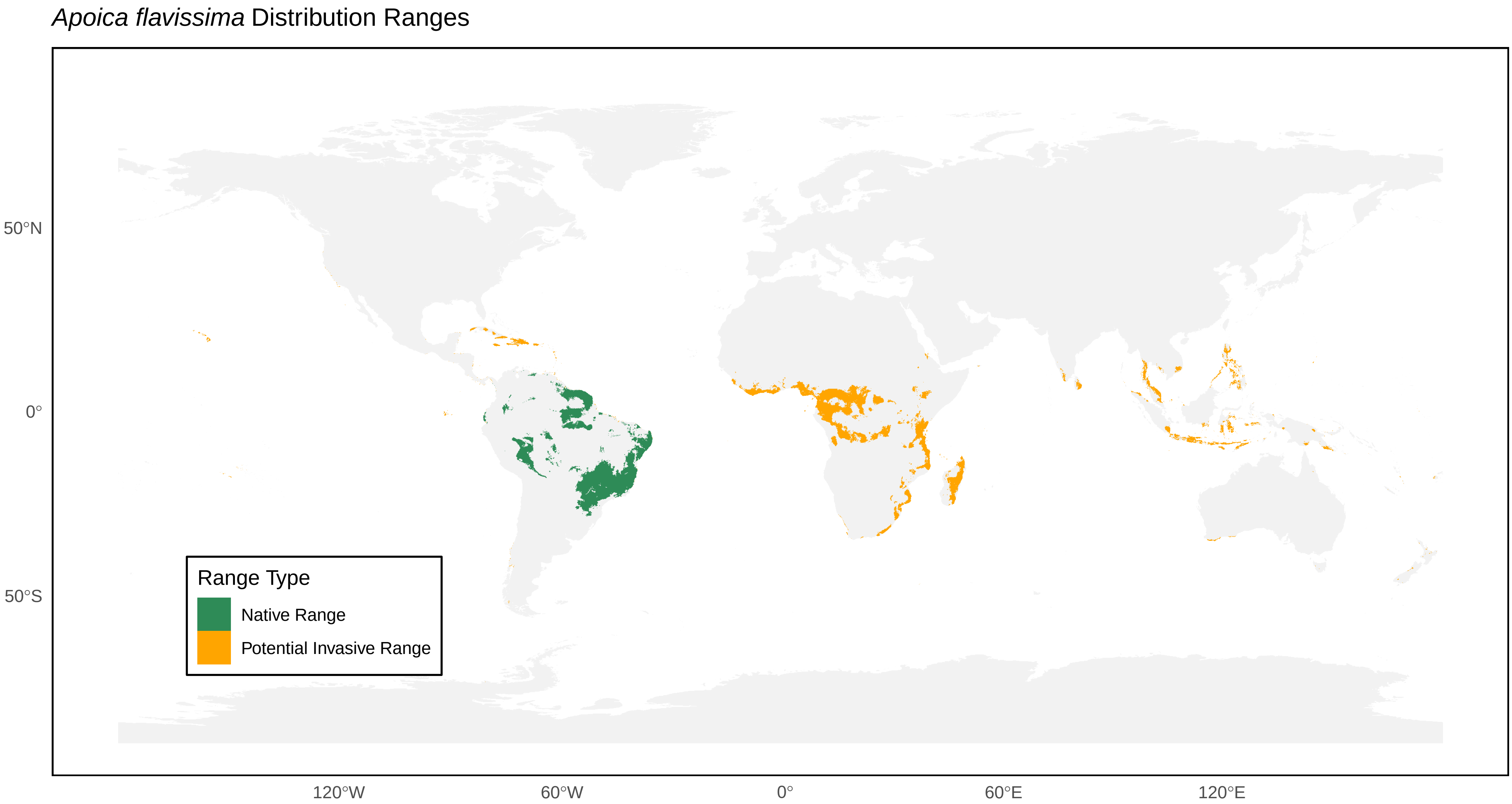

*Apoica flavissima* Distribution Ranges
50°N
0°
50°S
120°W
60°W
0°
60°E
120°E
Range Type
Native Range
Potential Invasive Range

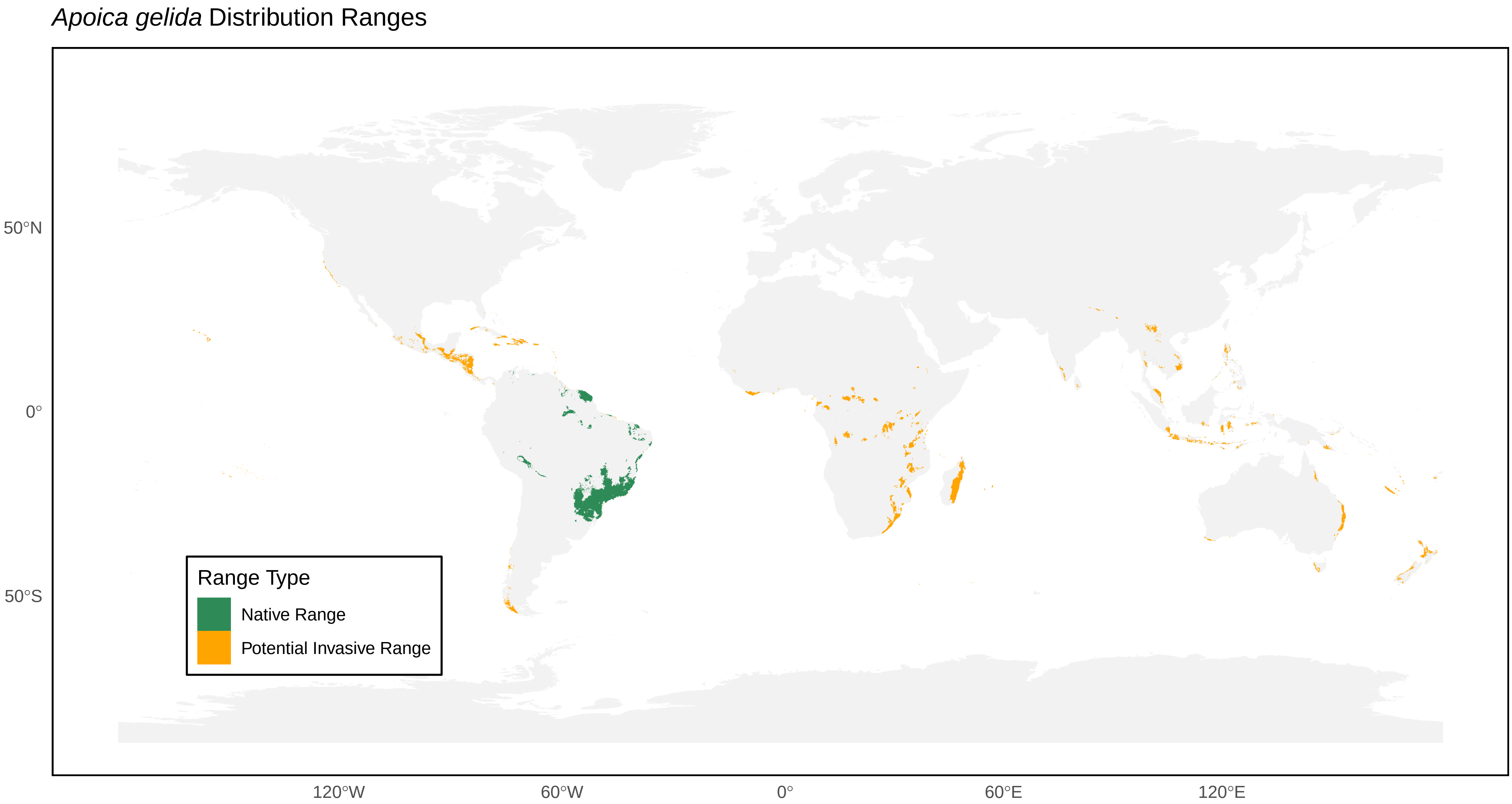
*Apoica gelida* Distribution Ranges
50°N
0°
50°S
120°W
60°W
0°
60°E
120°E
Range Type
Native Range
Potential Invasive Range

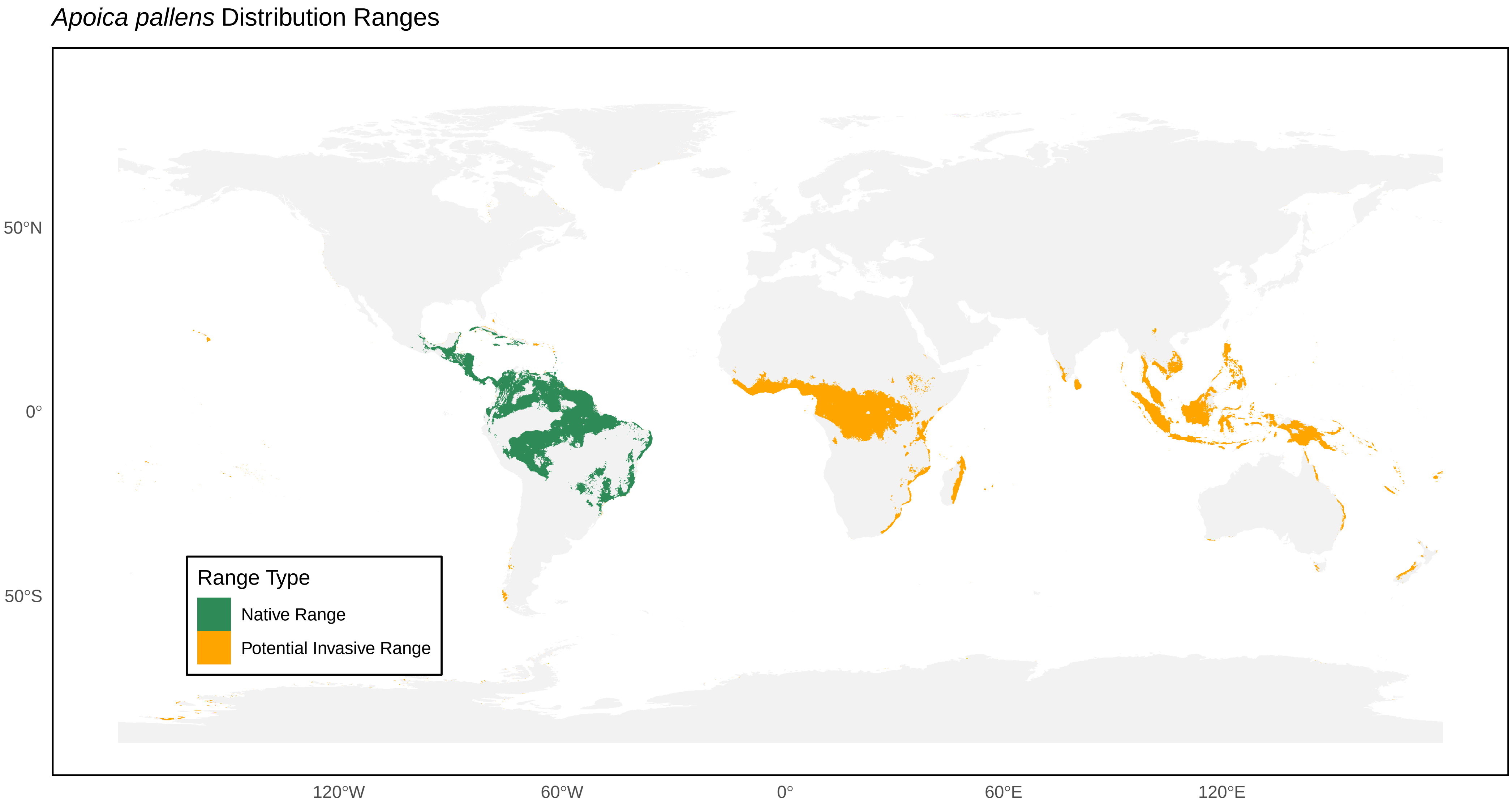
Apoica pallens Distribution Ranges
50°N
0°
50°S
120°W
60°W
0°
60°E
120°E
Range Type
Native Range
Potential Invasive Range

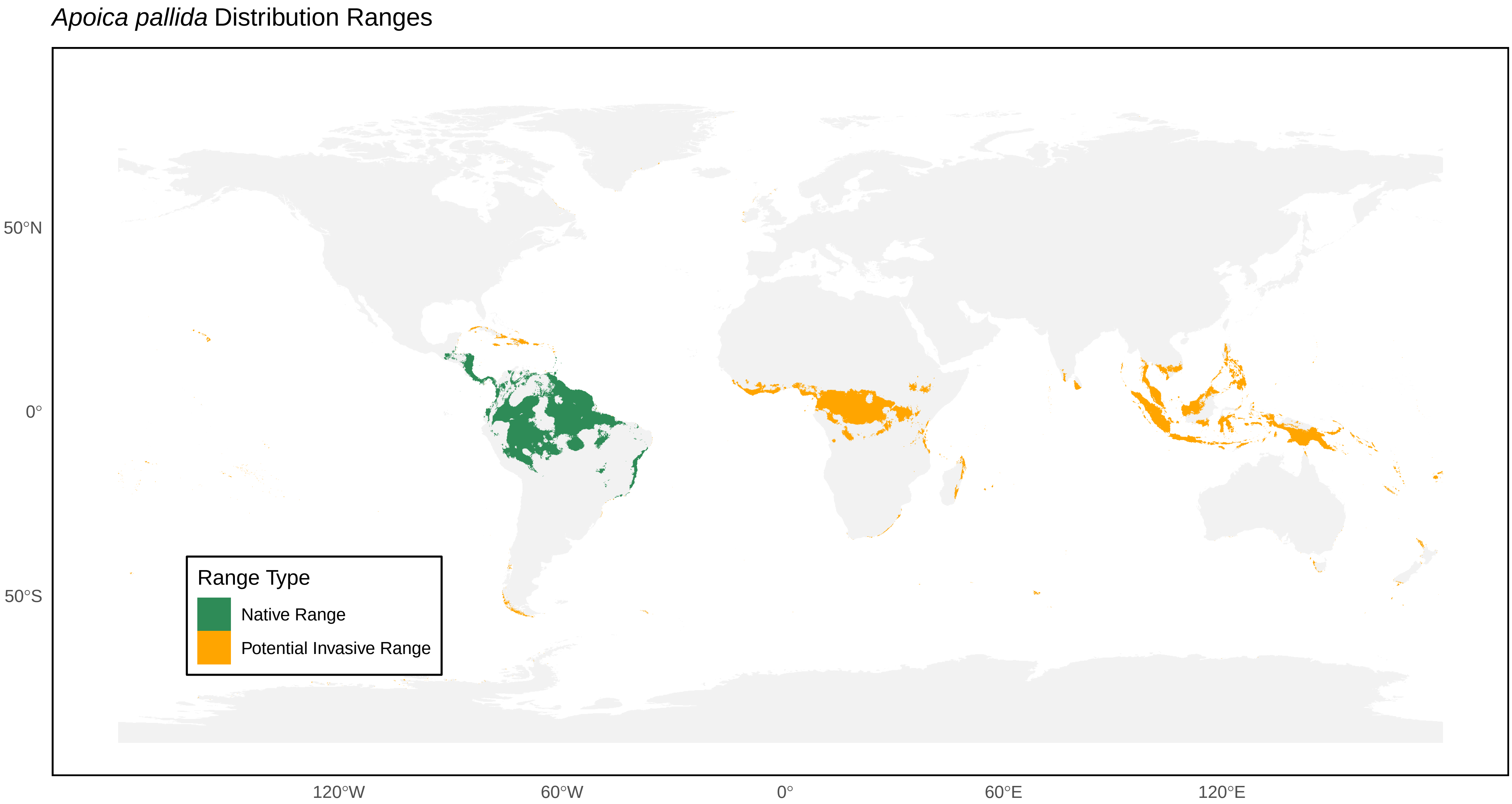
*Apoica pallida* Distribution Ranges
50°N
0°
50°S
120°W
60°W
0°
60°E
120°E
Range Type
Native Range
Potential Invasive Range

*Apoica strigata* Distribution Ranges

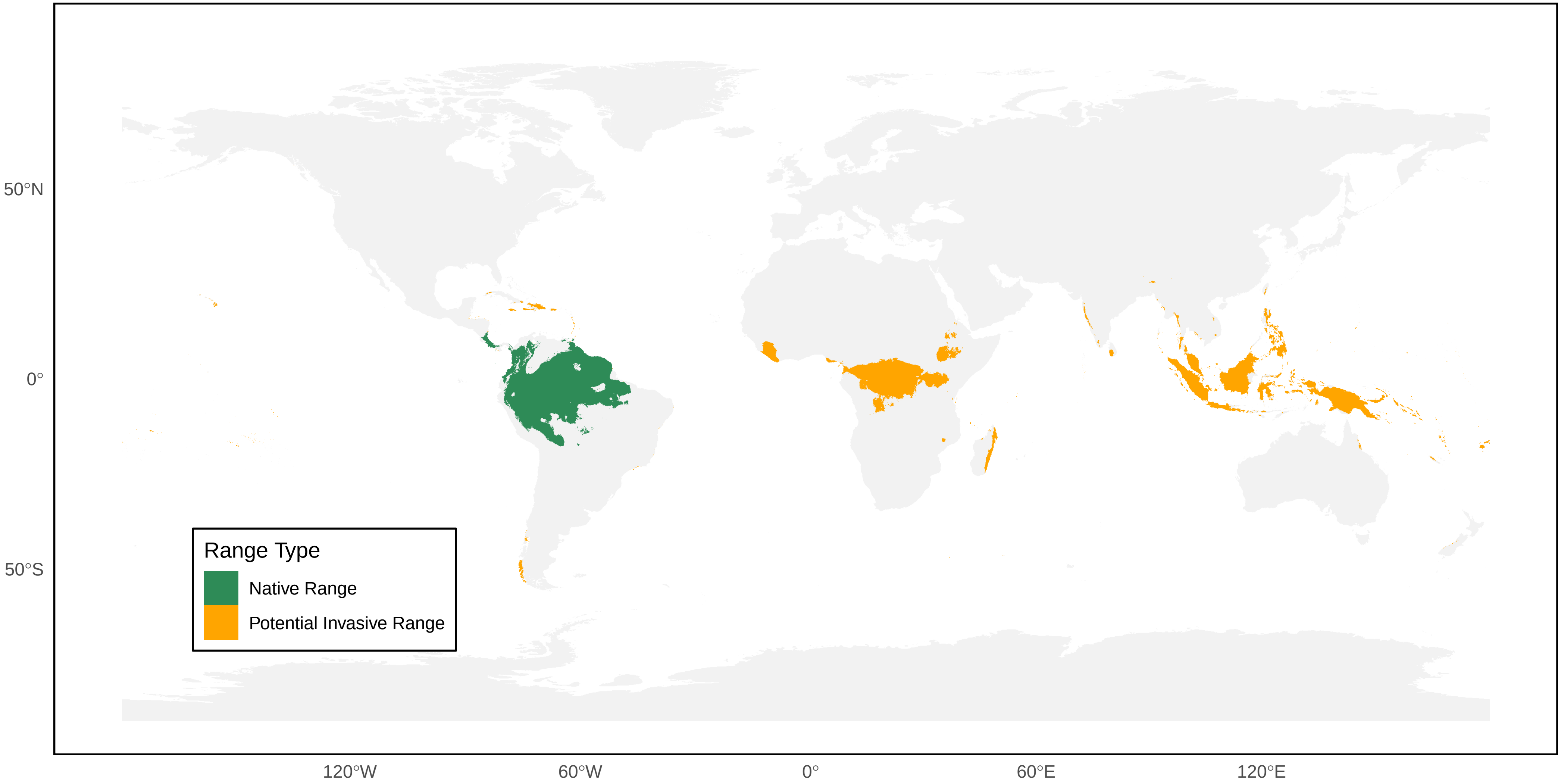

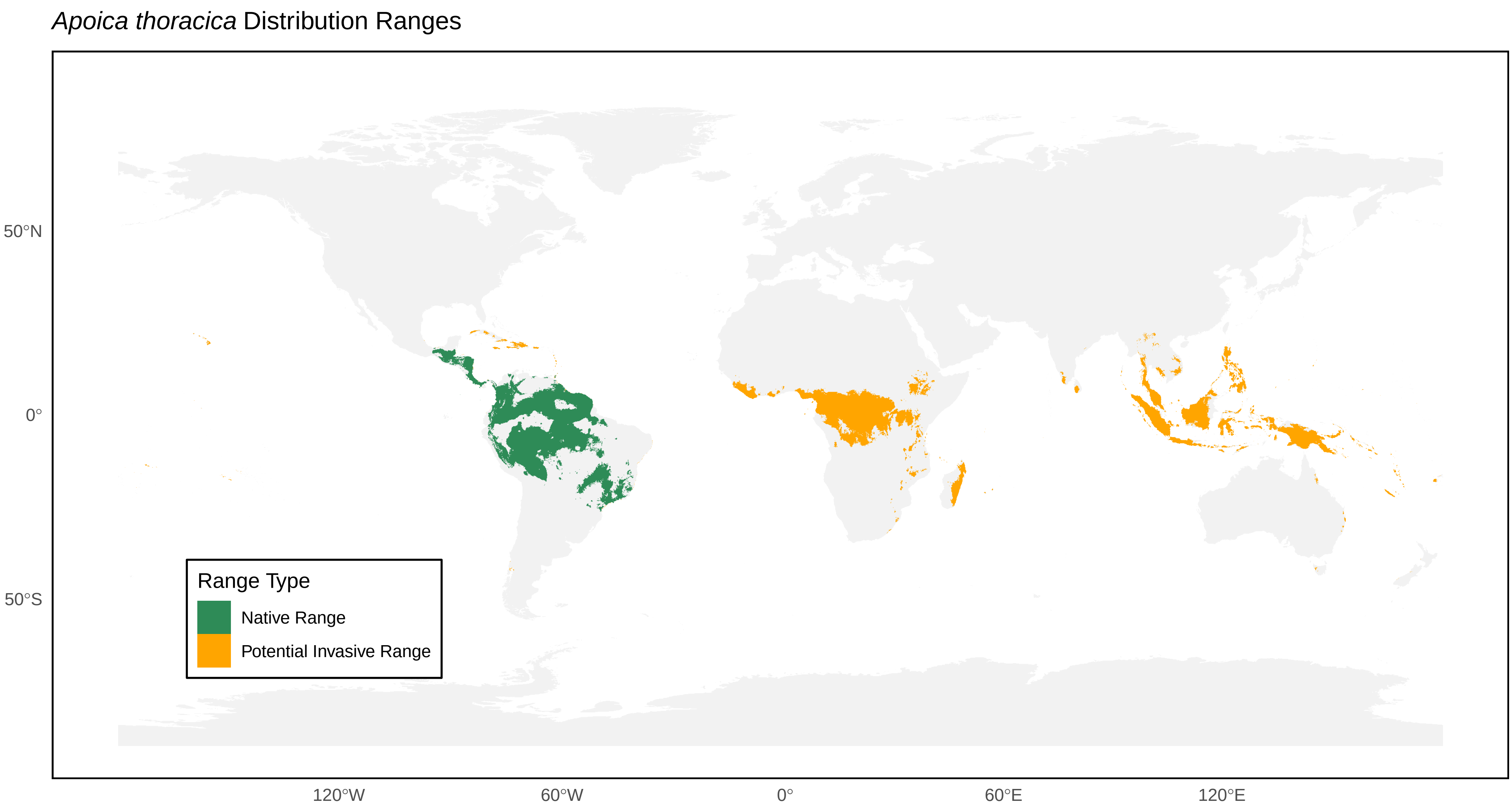
*Apoica thoracica* Distribution Ranges
50°N
0°
50°S
120°W
60°W
0°
60°E
120°E
Range Type
Native Range
Potential Invasive Range

*Belonogaster dubia* Distribution Ranges

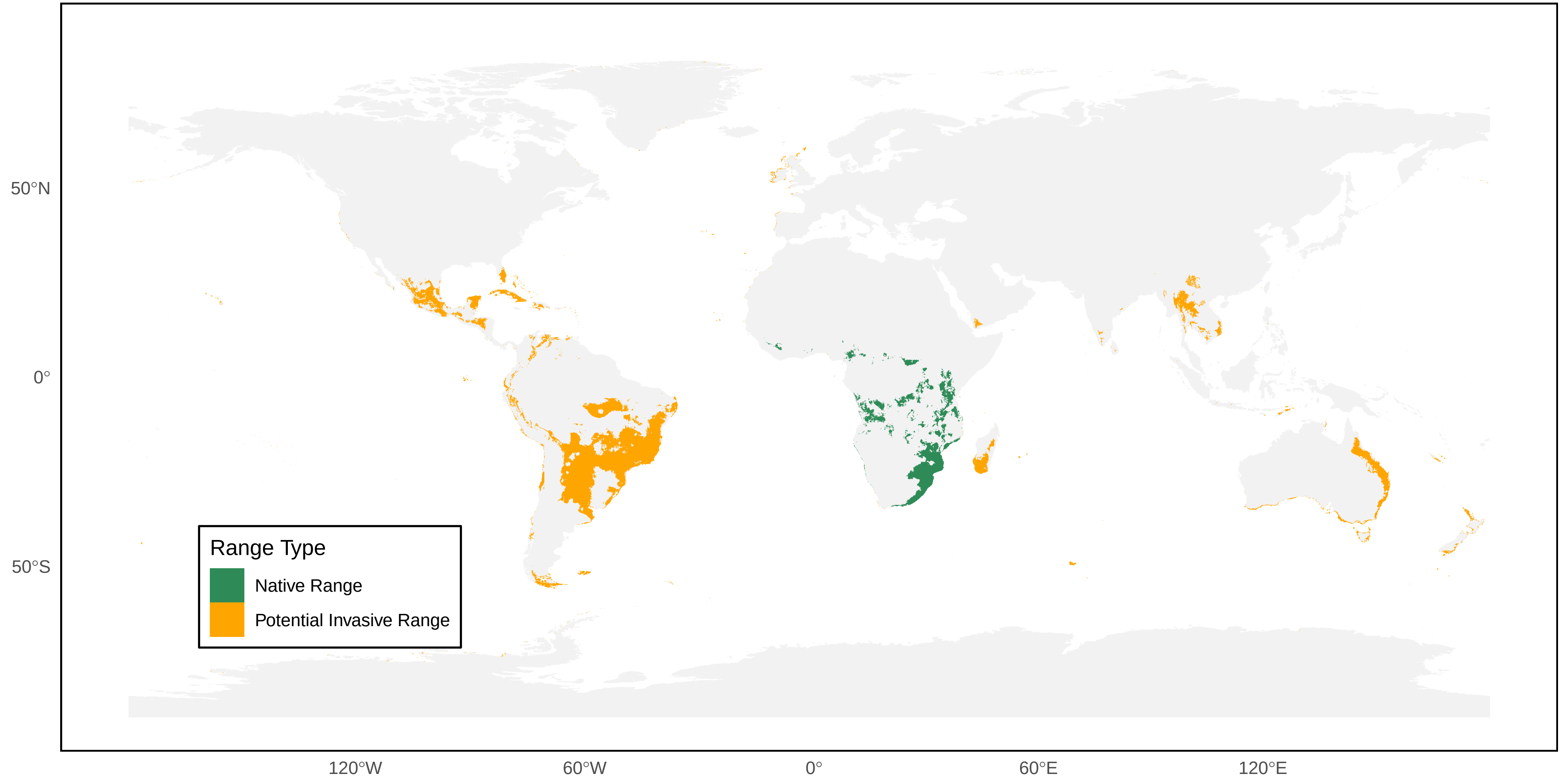

*Belonogaster filiventris* Distribution Ranges

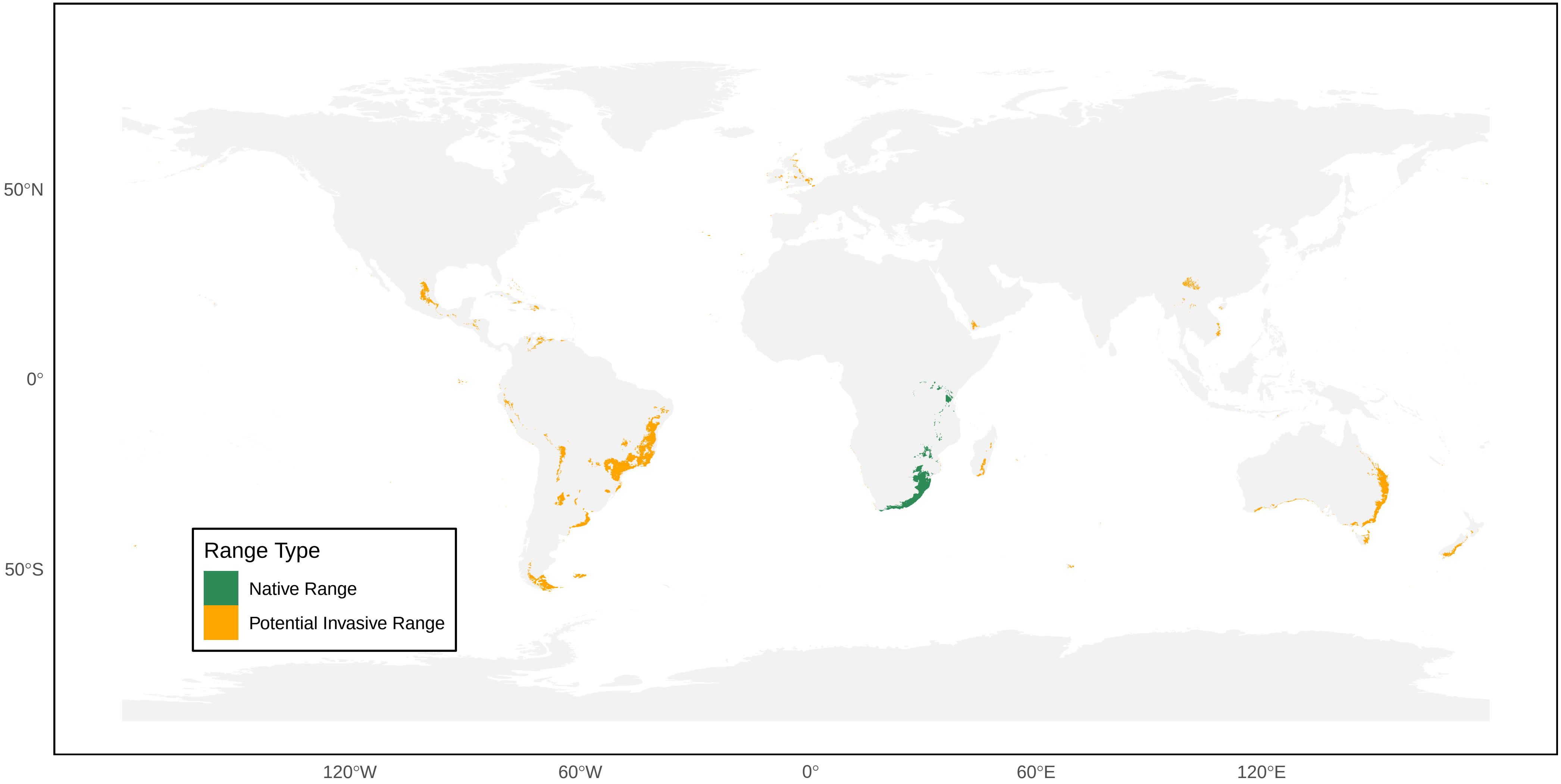

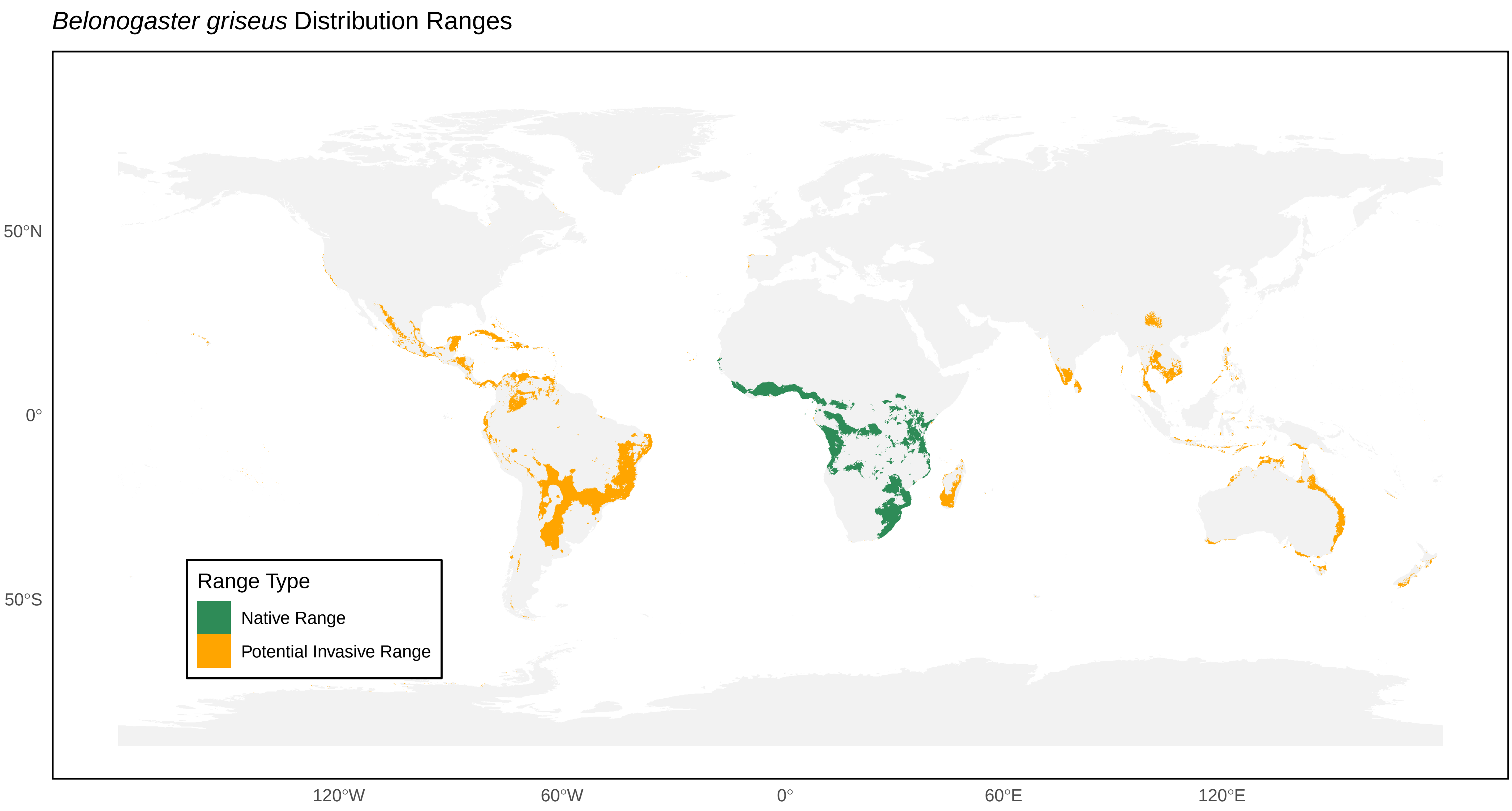
*Belonogaster griseus* Distribution Ranges
50°N
0°
50°S
120°W
60°W
0°
60°E
120°E
Range Type
Native Range
Potential Invasive Range

*Belonogaster juncea* Distribution Ranges

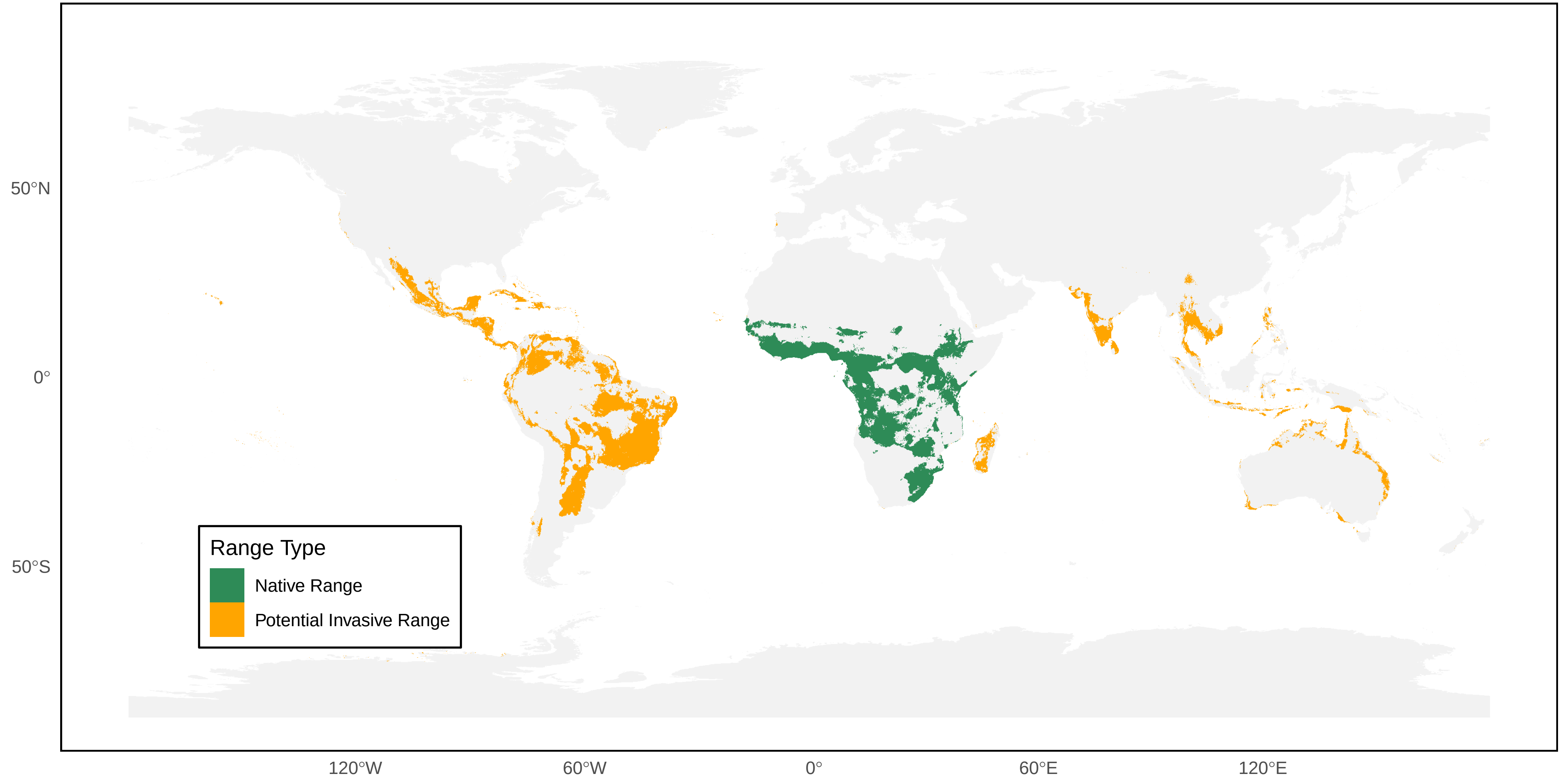

*Belonogaster lateritia* Distribution Ranges

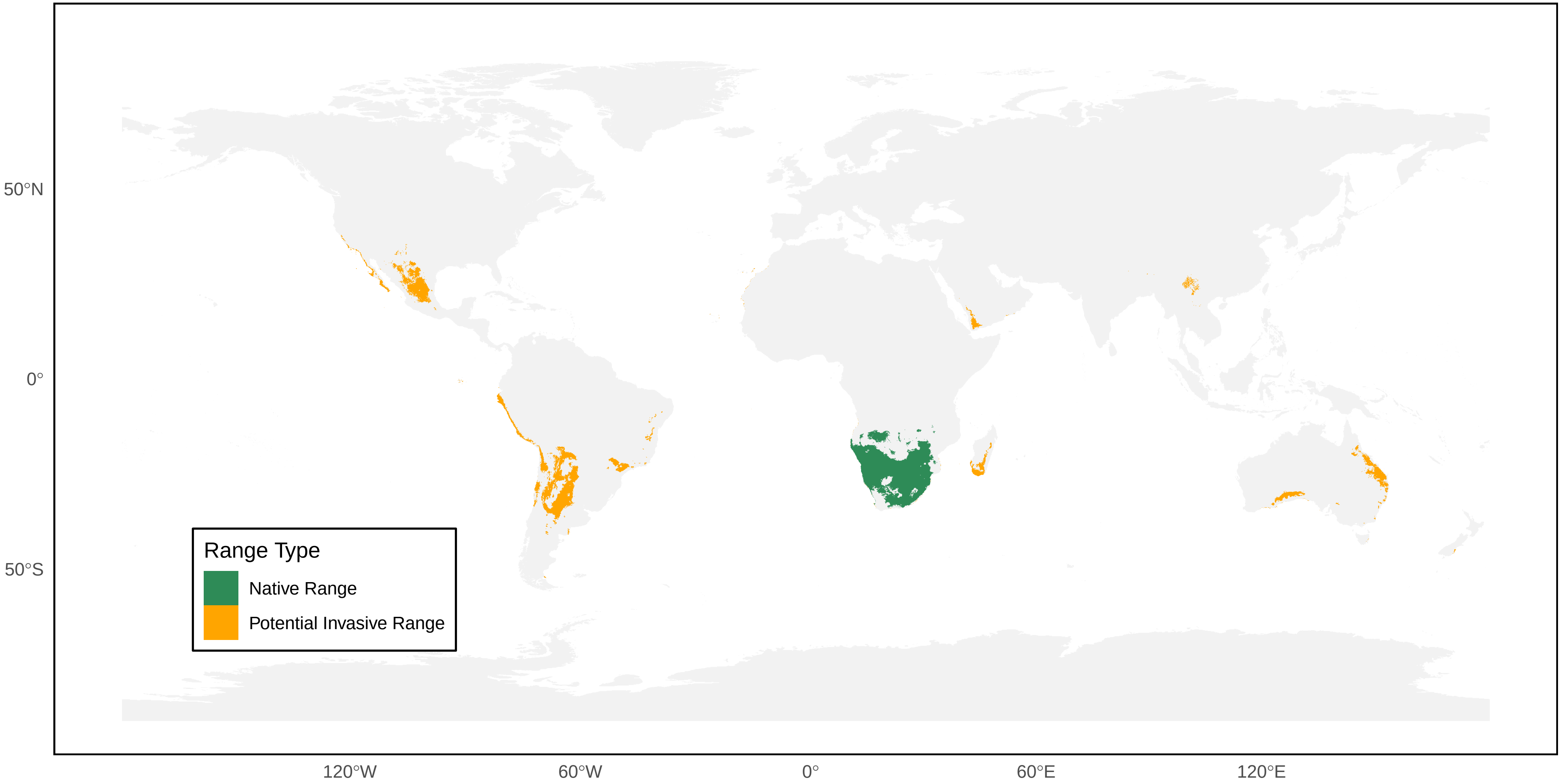

*Brachygastra augusti* Distribution Ranges

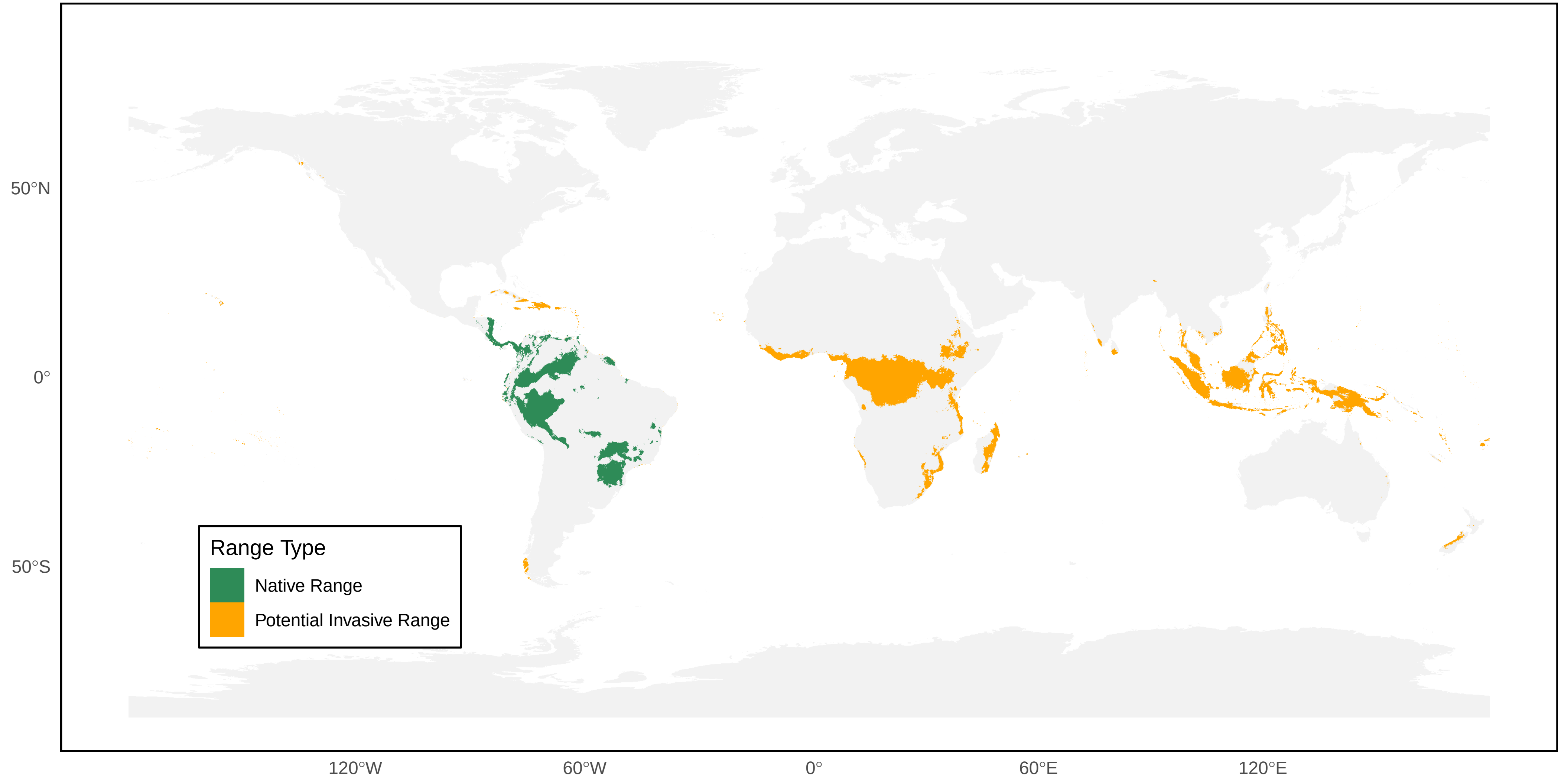

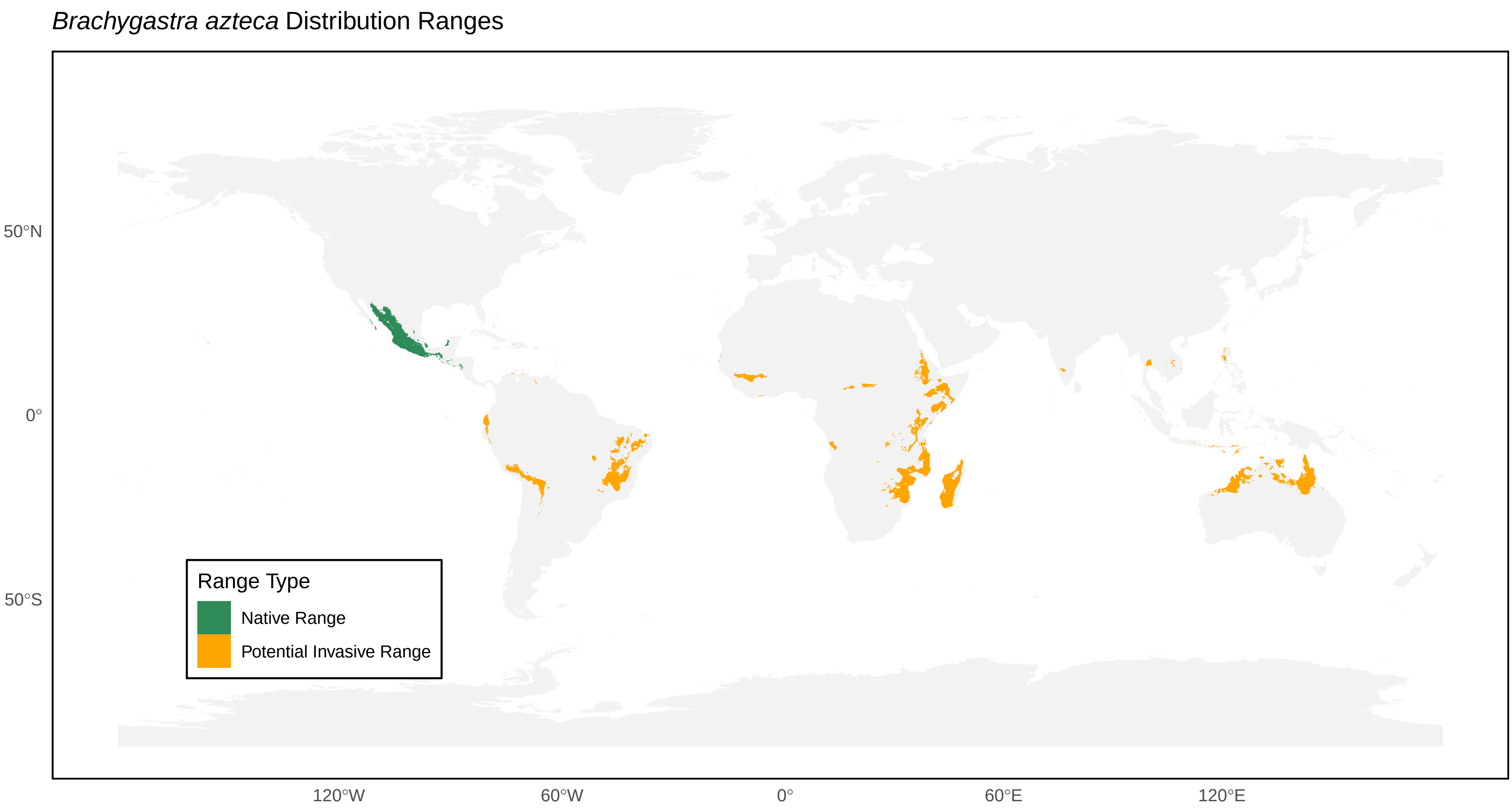

*Brachygastra azteca* Distribution Ranges
50°N
0°
50°S
120°W
60°W
0°
60°E
120°E
Range Type
Native Range
Potential Invasive Range

*Brachygastra lecheguana* Distribution Ranges

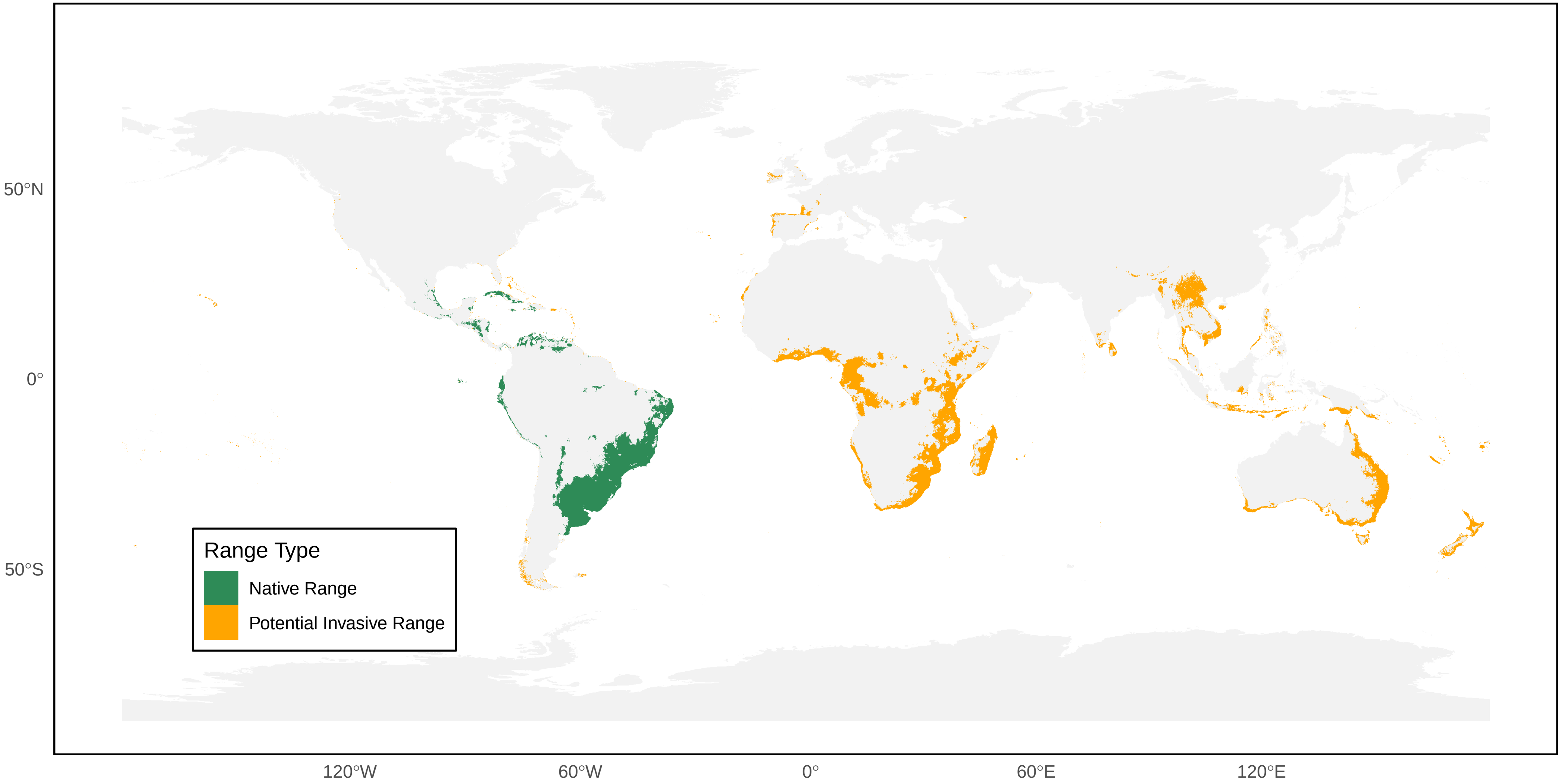

*Brachygastra mellifica* Distribution Ranges

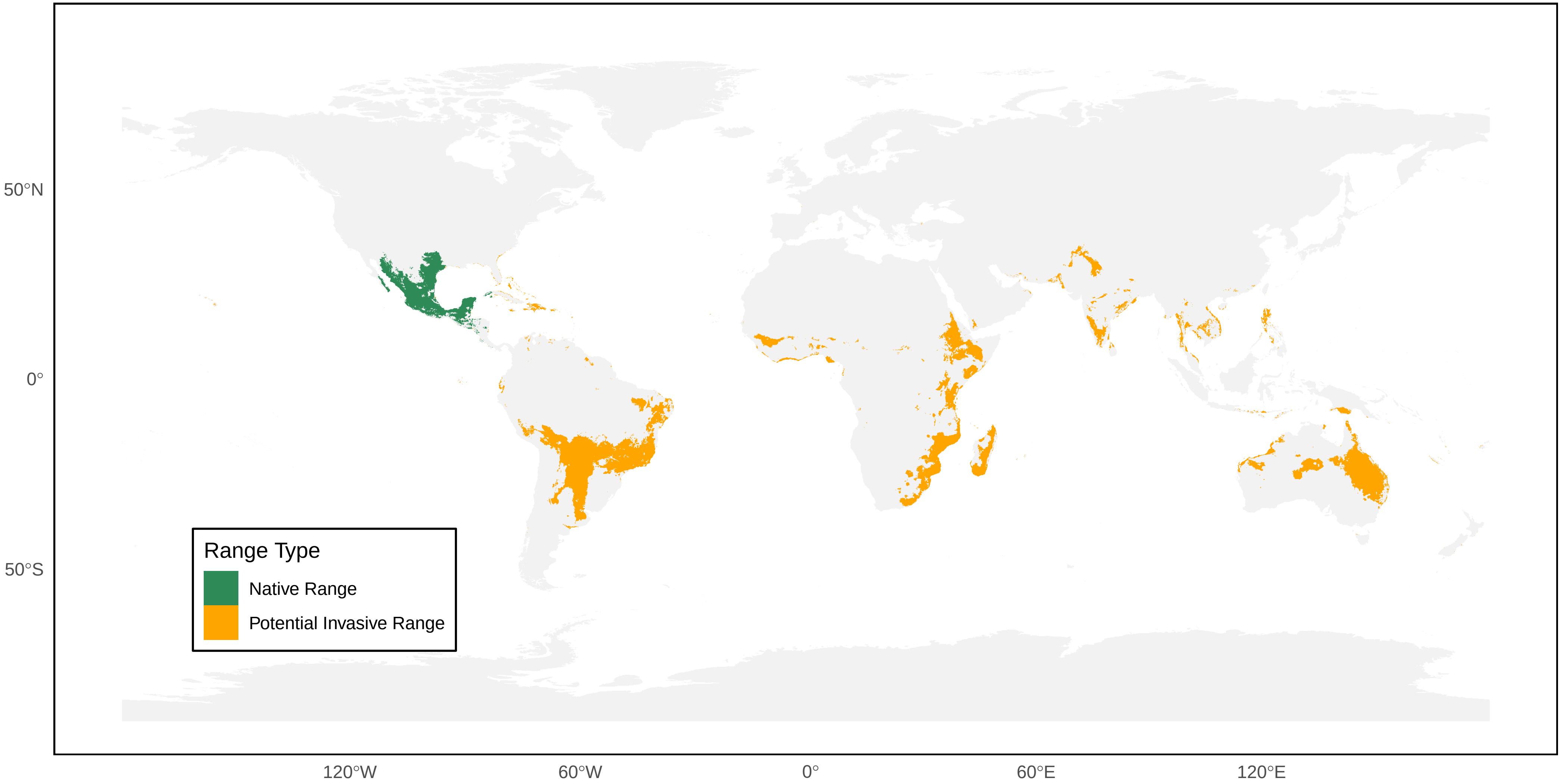

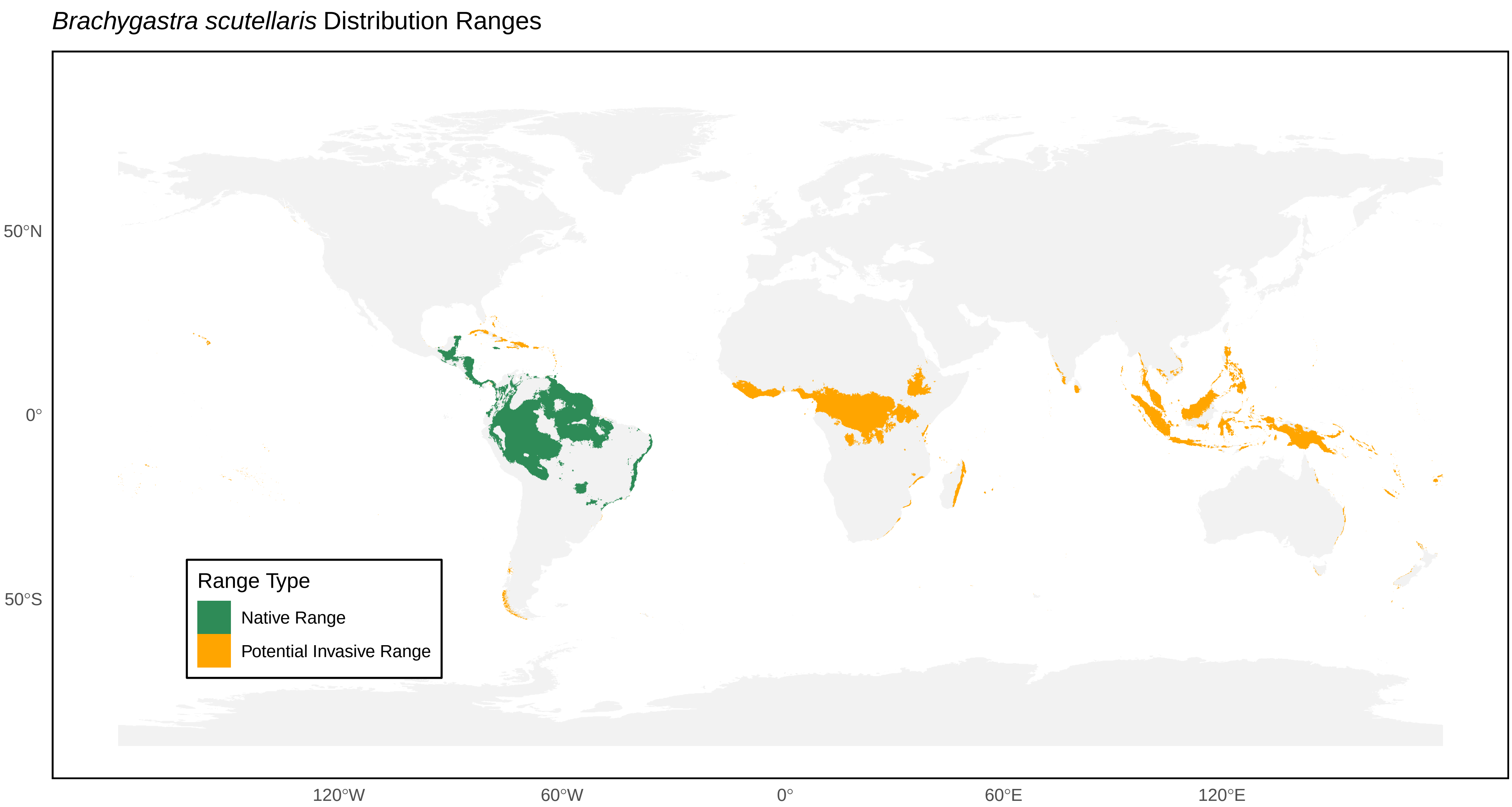

*Brachygastra scutellaris* Distribution Ranges
50°N
0°
50°S
120°W
60°W
0°
60°E
120°E
Range Type
Native Range
Potential Invasive Range

*Brachygastra smithii* Distribution Ranges

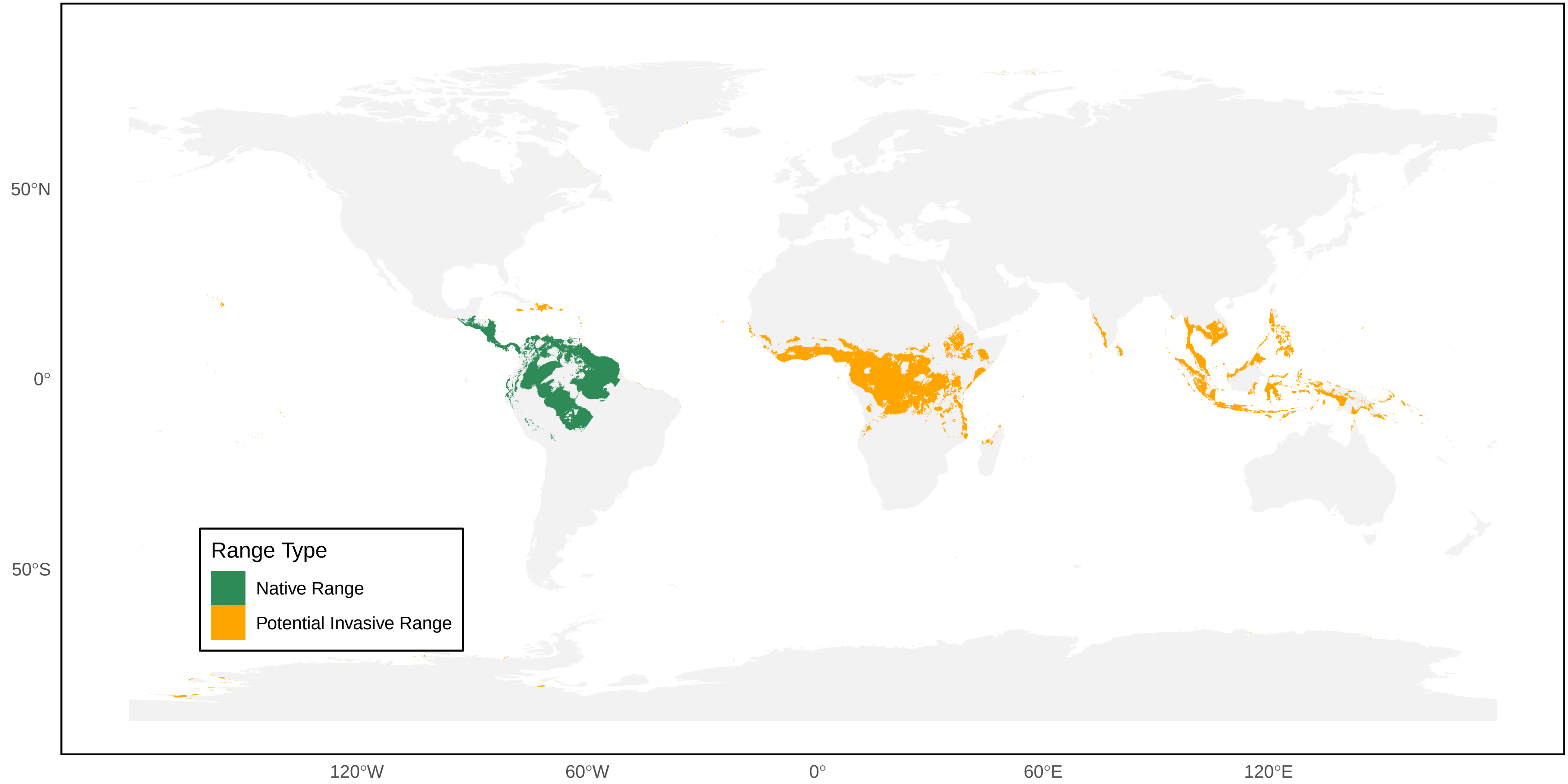

*Charterginus fulvus* Distribution Ranges

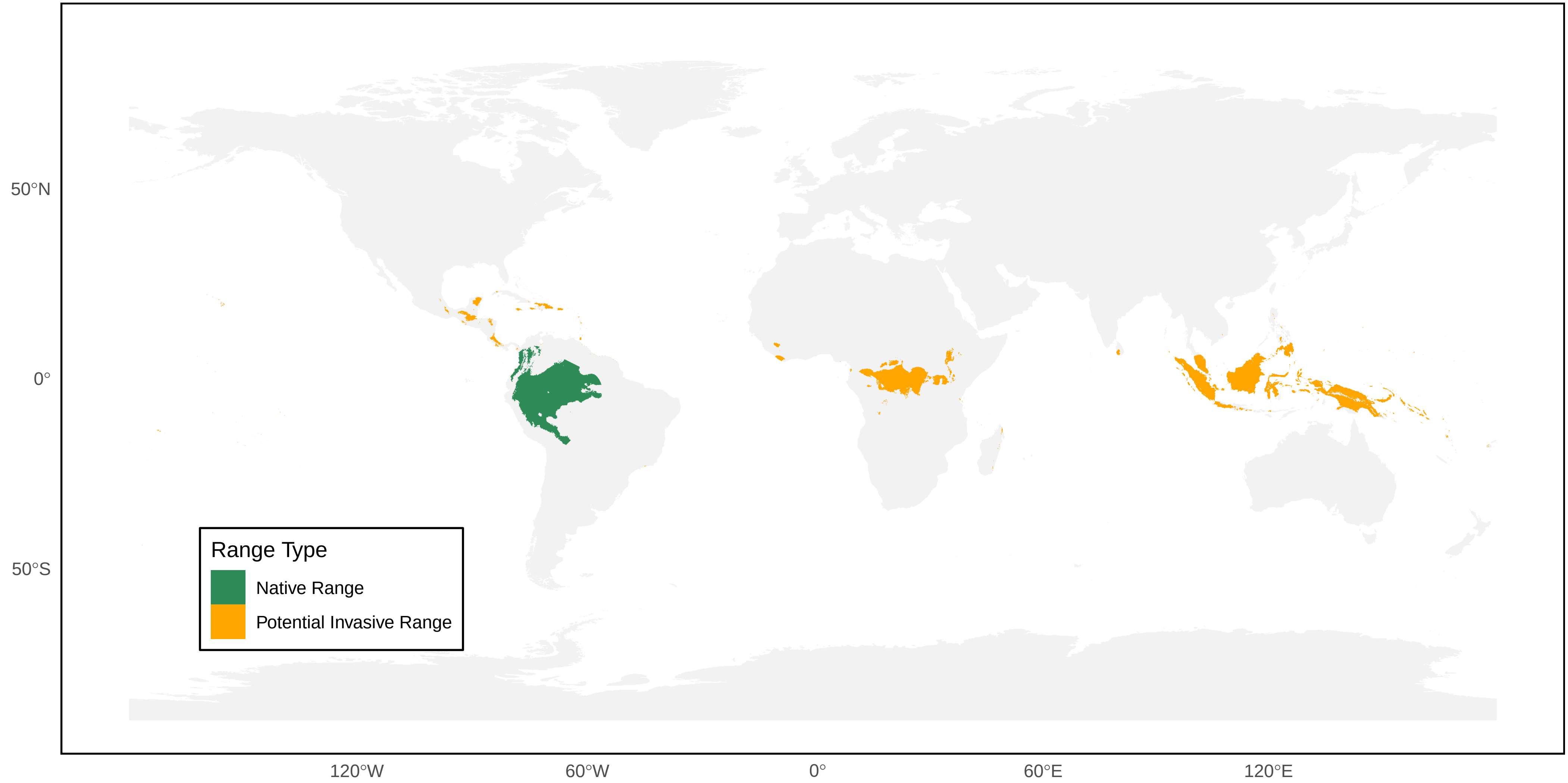

*Charterginus nevermanni* Distribution Ranges

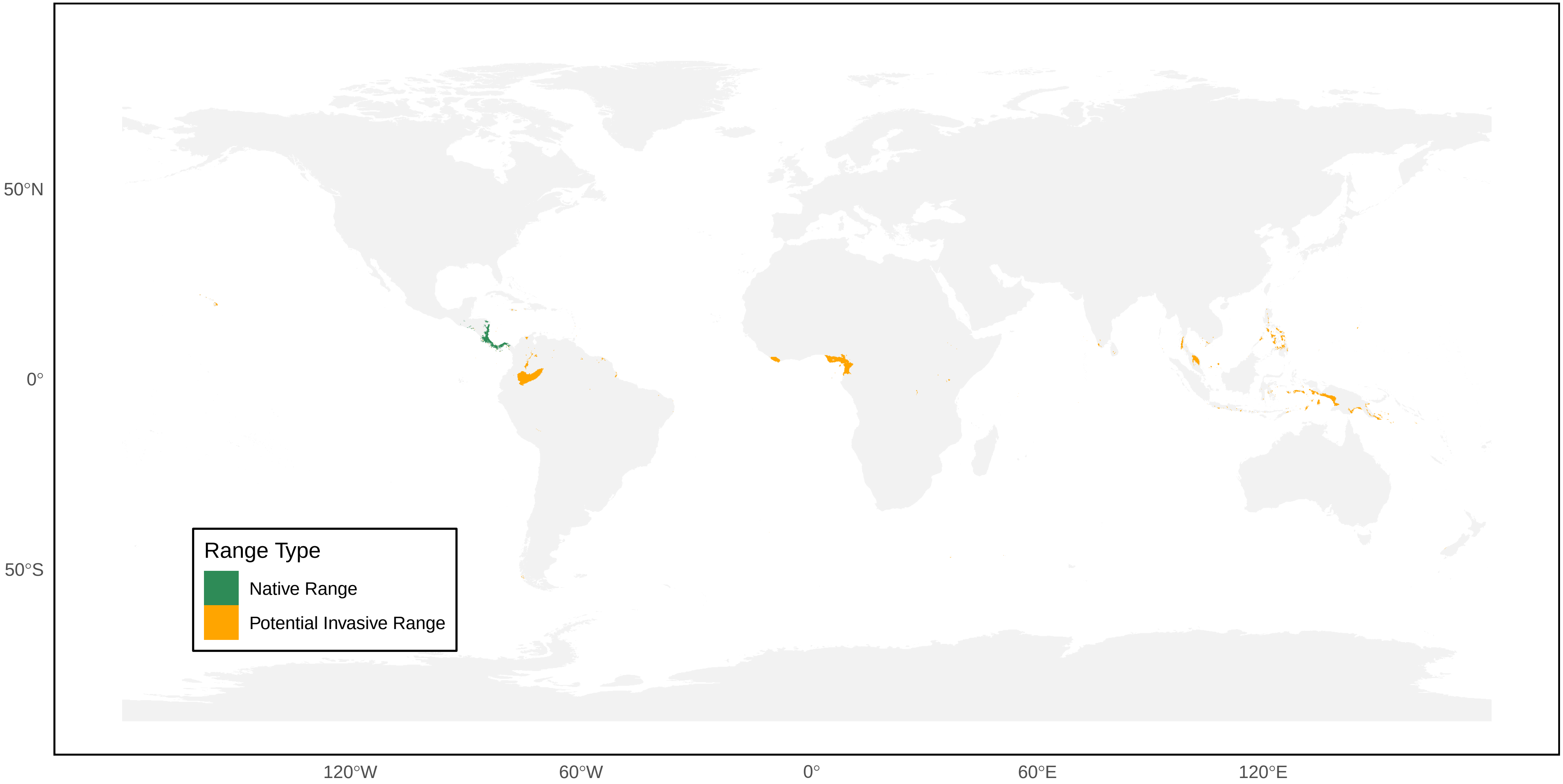

*Dolichovespula adulterina* Distribution Ranges

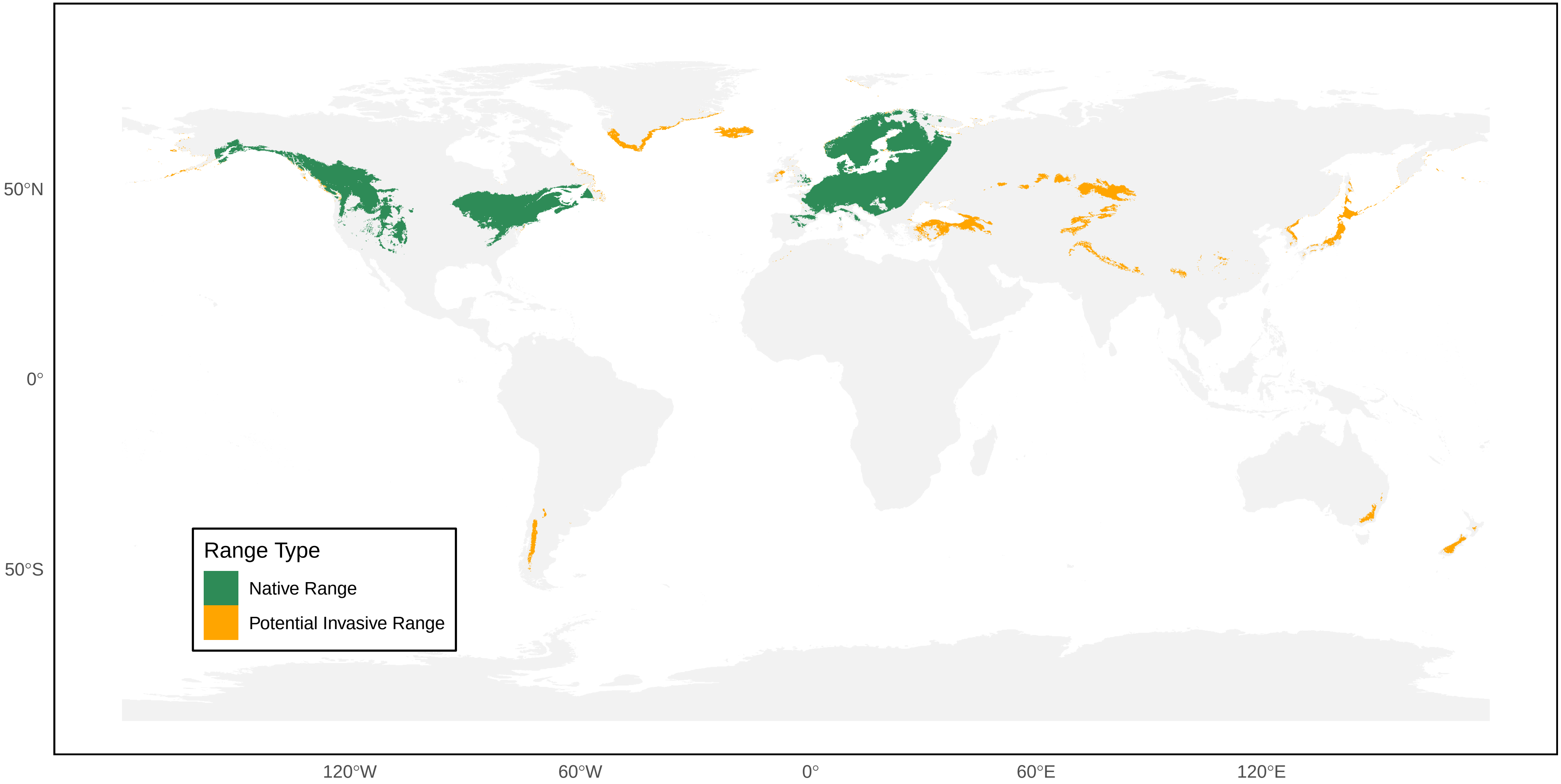

*Dolichovespula albida* Distribution Ranges

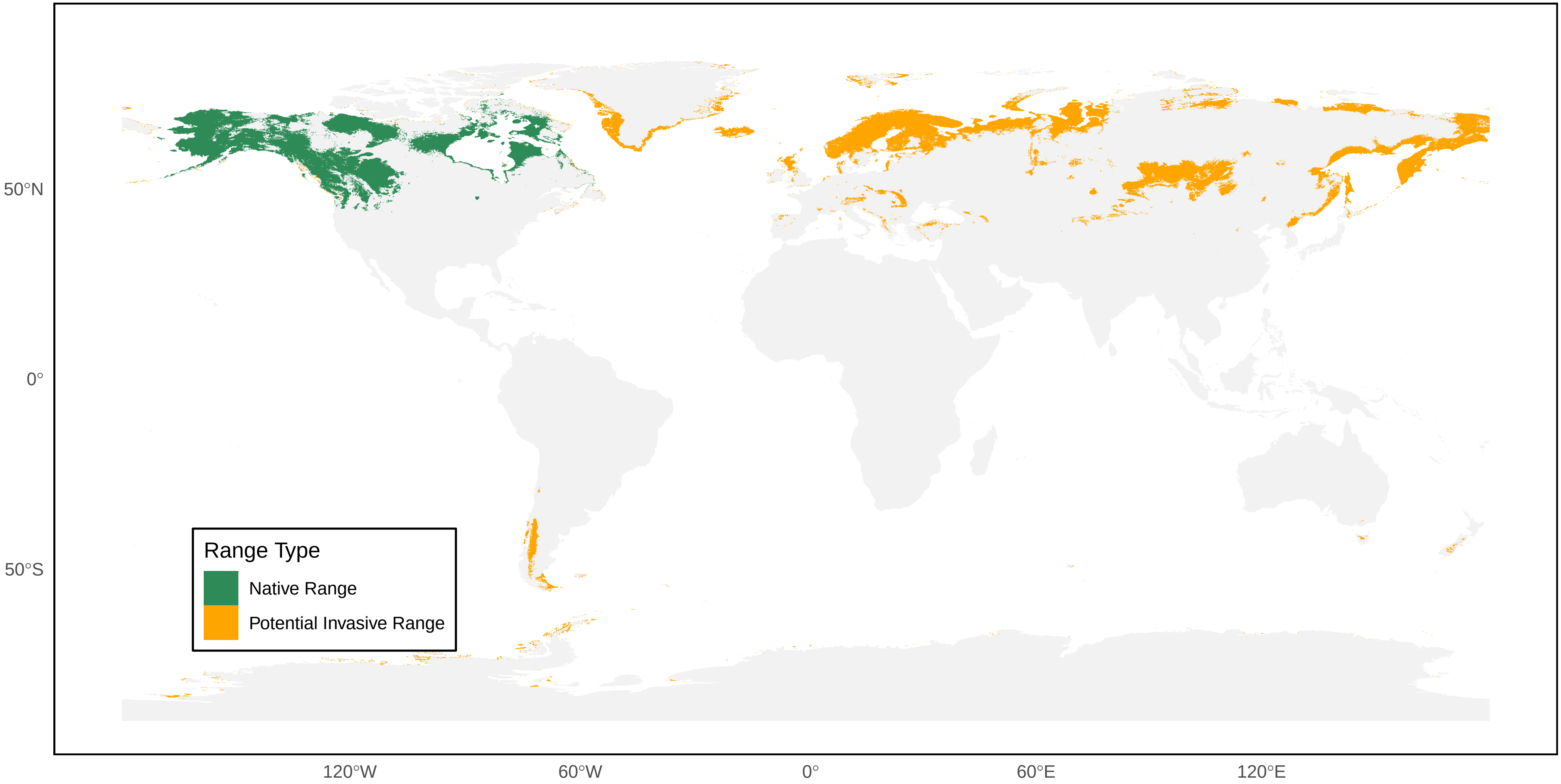

*Dolichovespula alpicola* Distribution Ranges

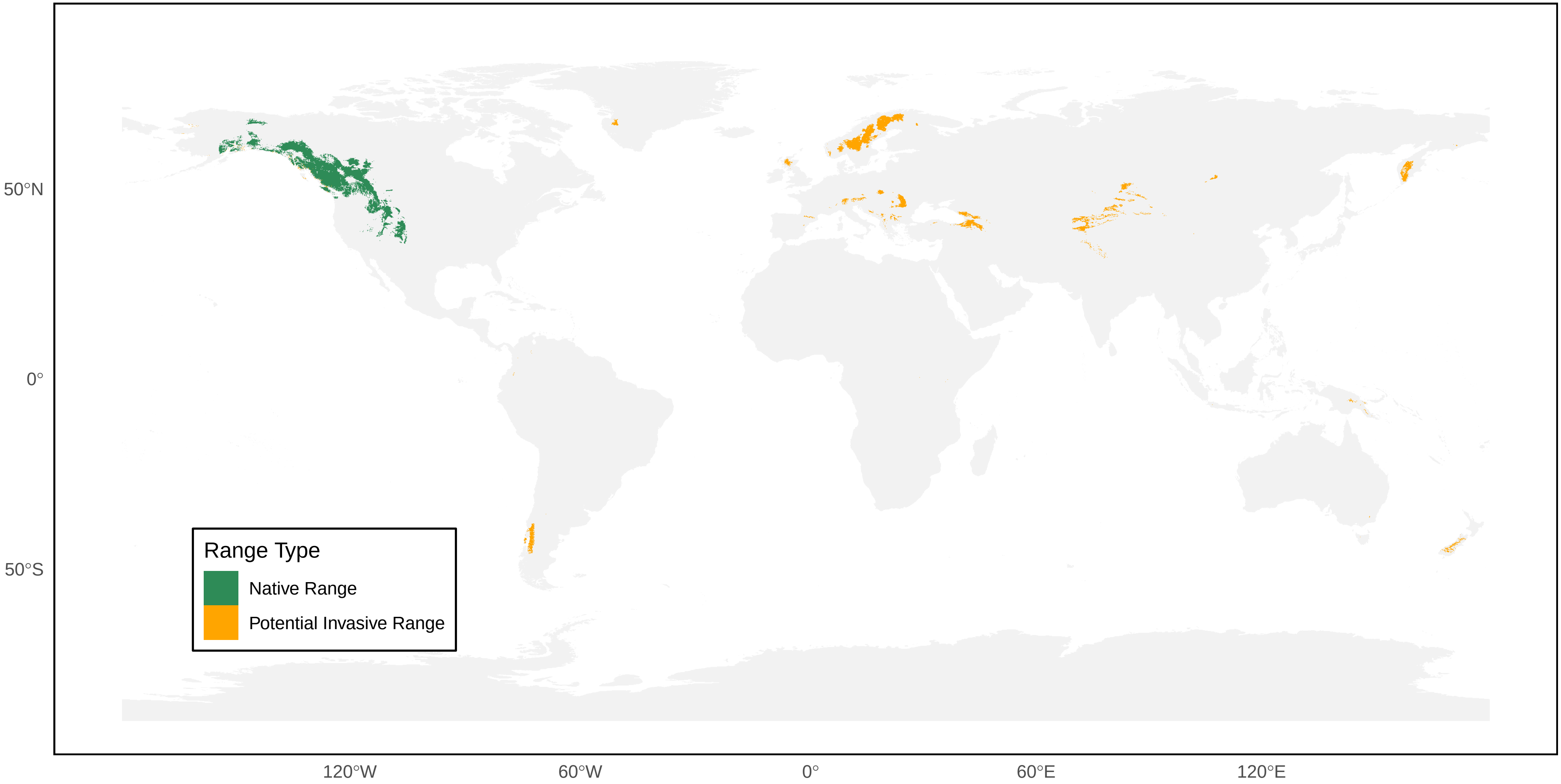

*Dolichovespula arctica* Distribution Ranges

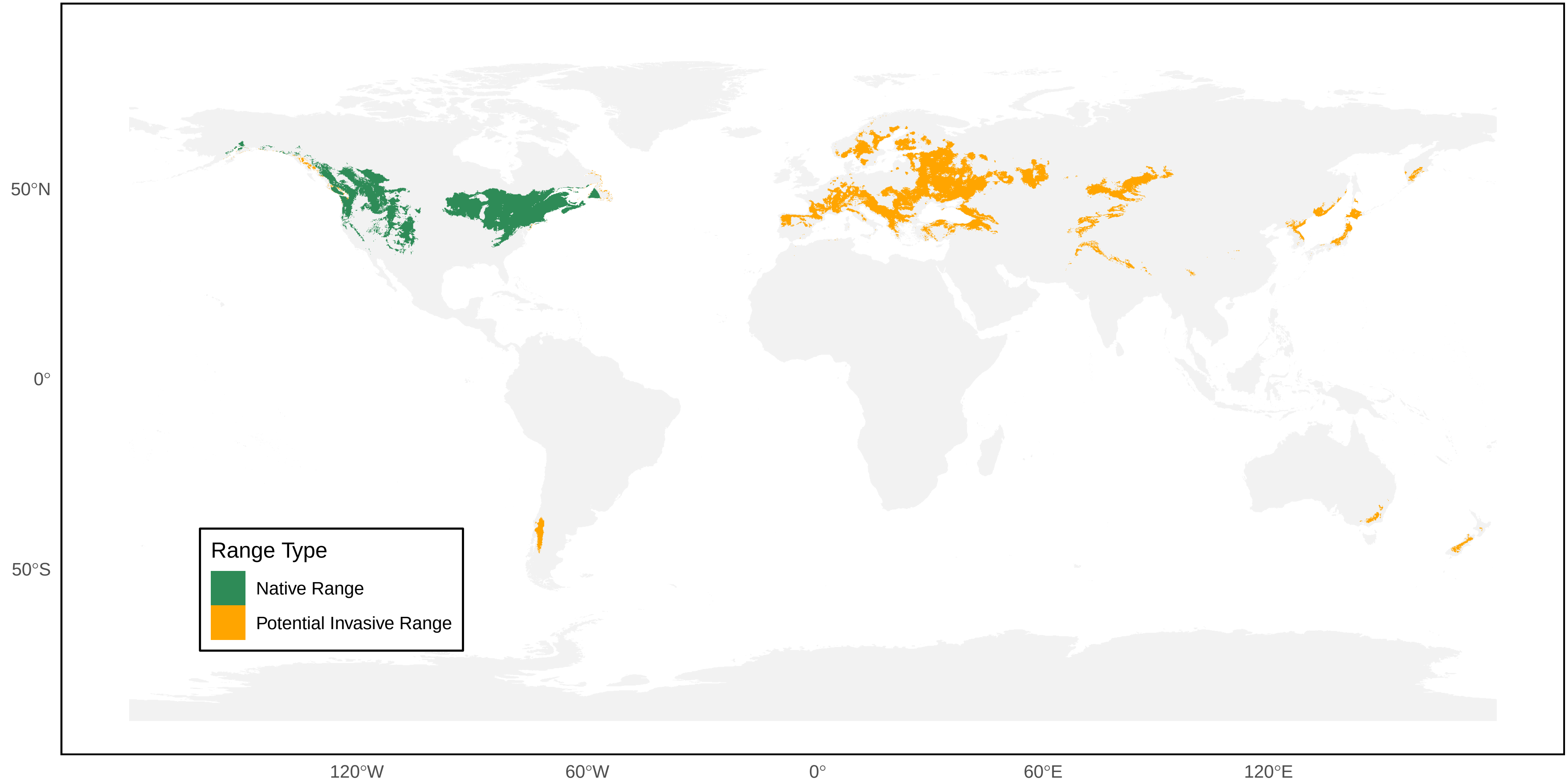

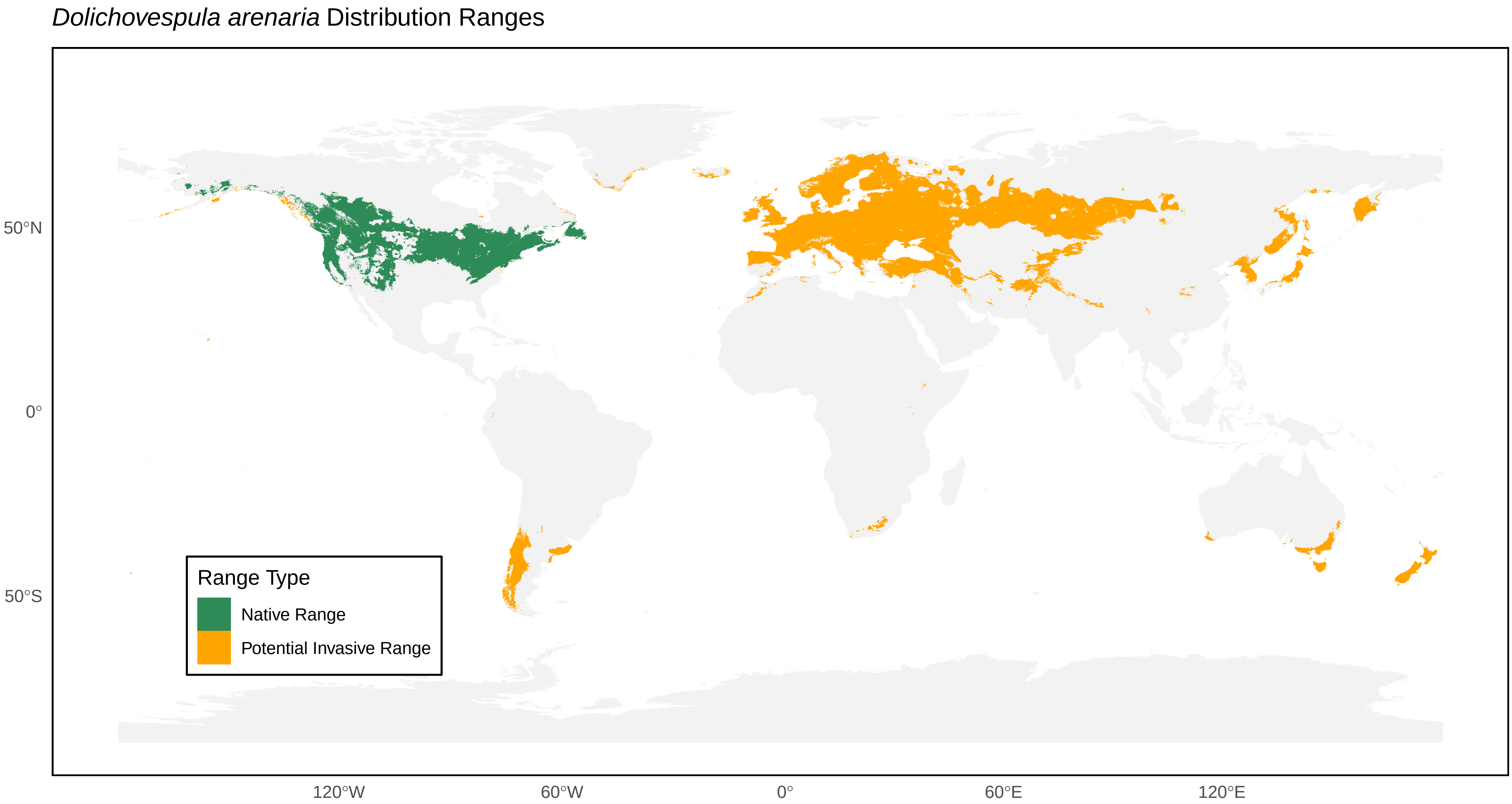

Dolichovespula arenaria Distribution Ranges
50°N
0°
50°S
120°W
60°W
0°
60°E
120°E
Range Type
Native Range
Potential Invasive Range

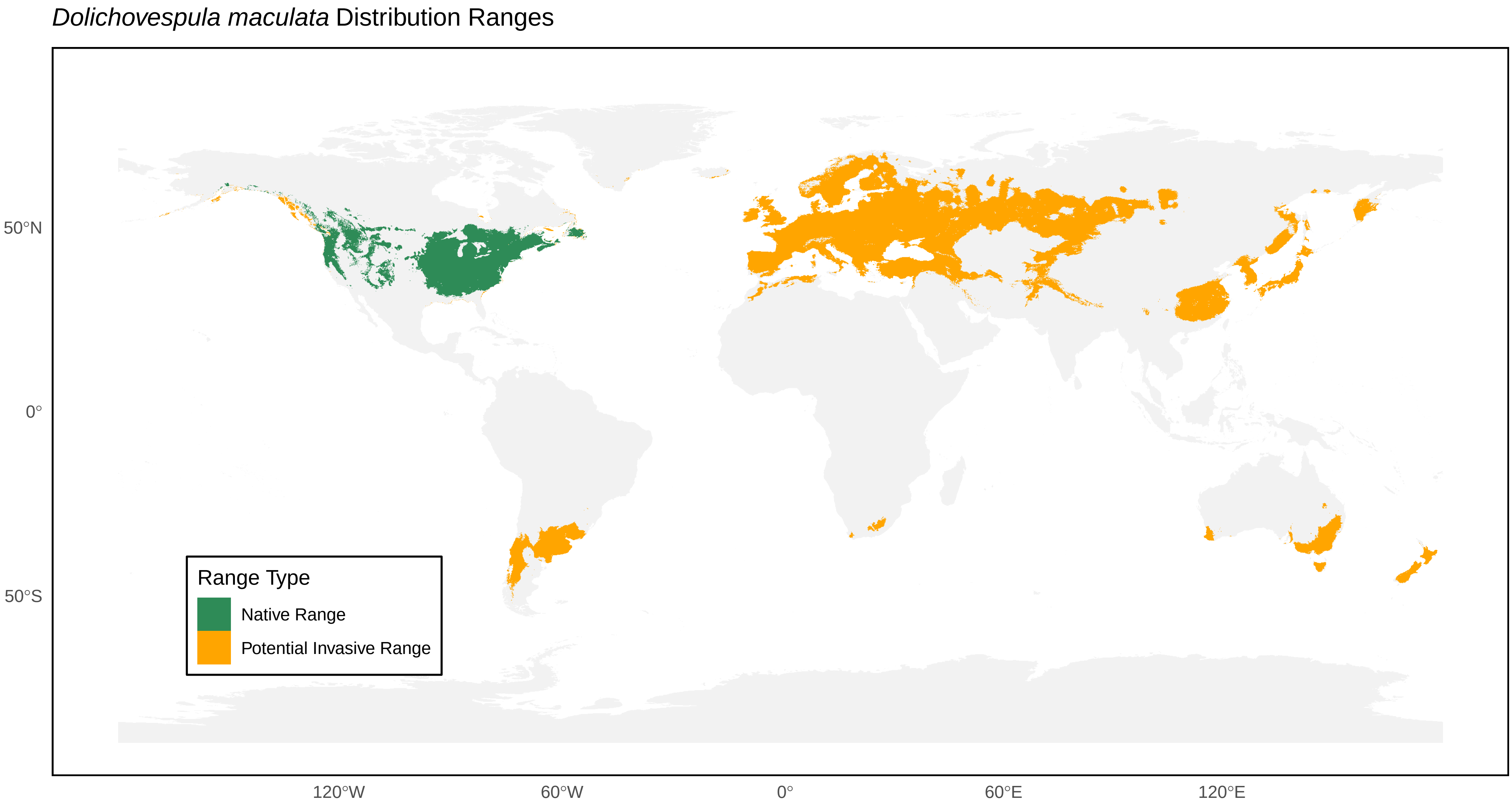

*Dolichovespula maculata* Distribution Ranges
50°N
0°
50°S
120°W
60°W
0°
60°E
120°E
Range Type
Native Range
Potential Invasive Range

*Dolichovespula media* Distribution Ranges

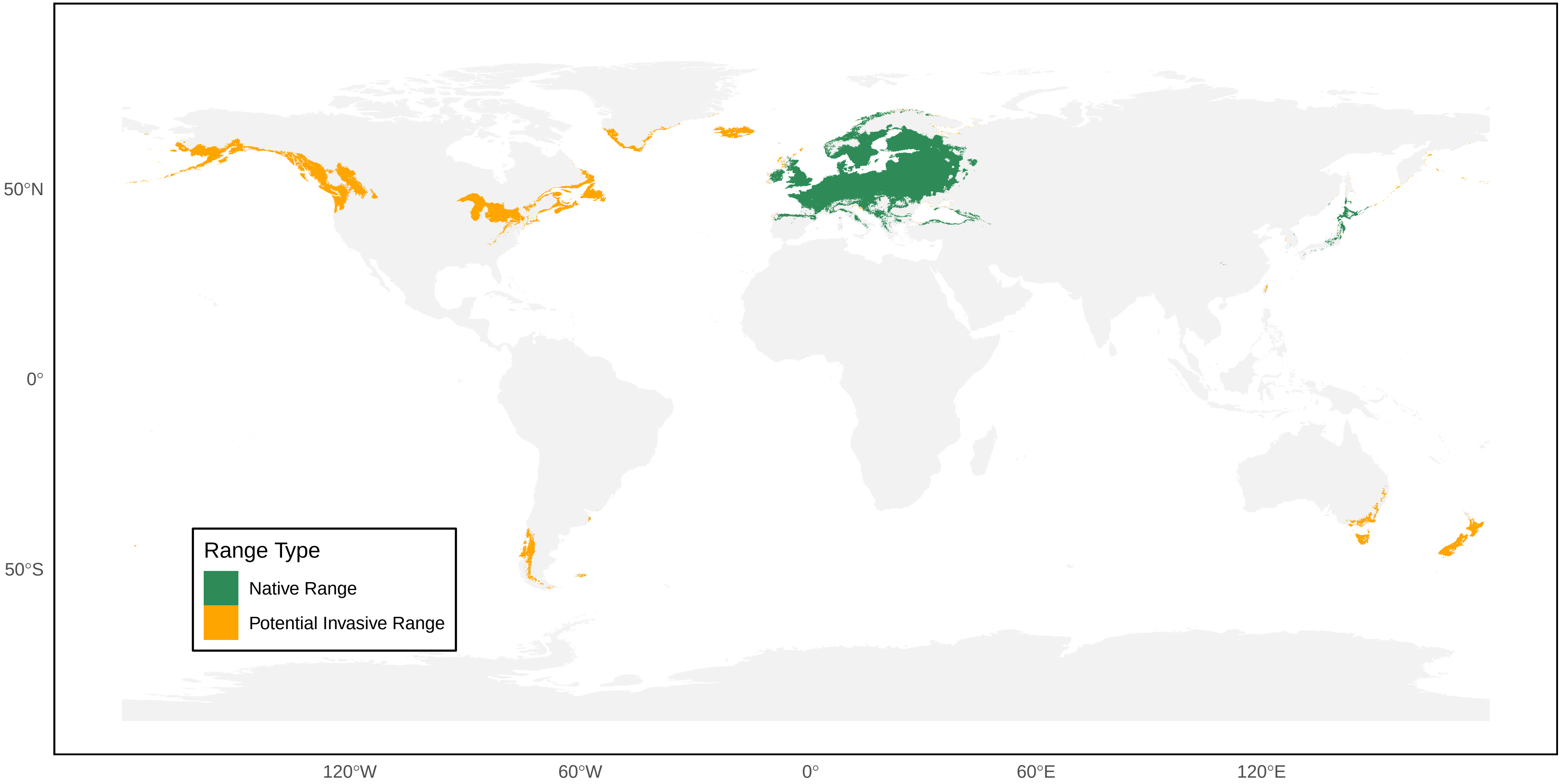

*Dolichovespula norvegicoides* Distribution Ranges

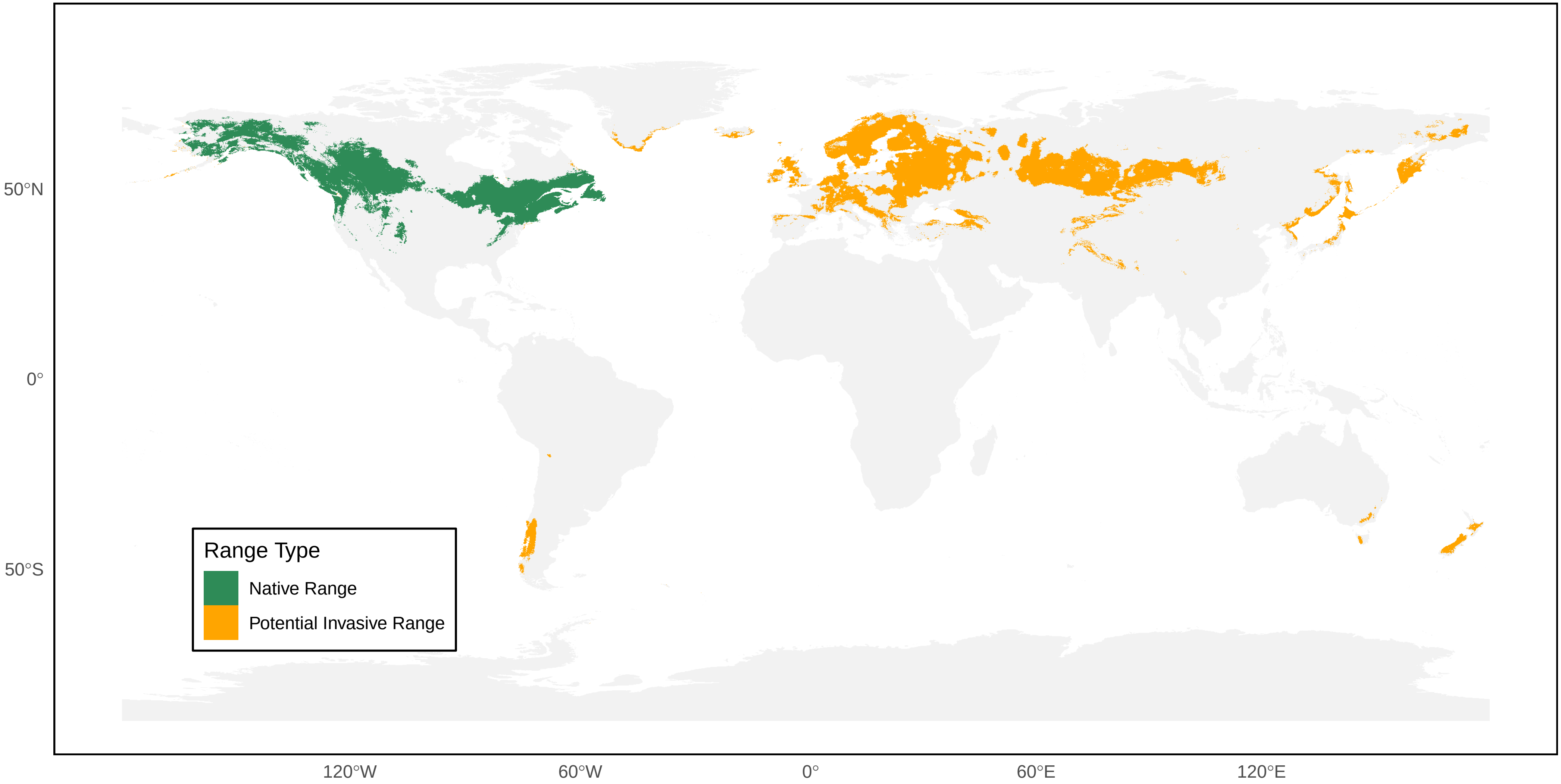

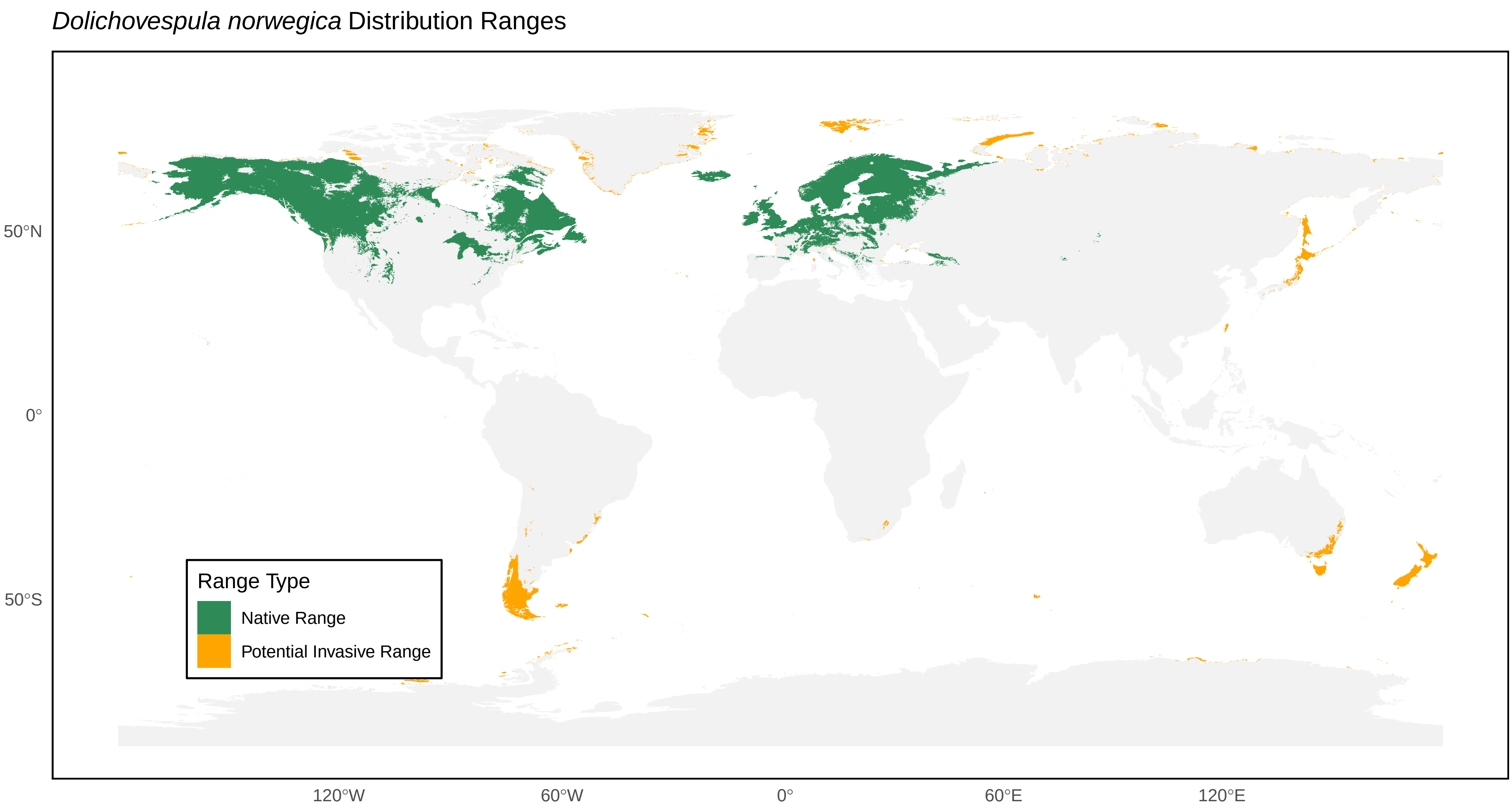
*Dolichovespula norwegica* Distribution Ranges
50°N
0°
50°S
120°W
60°W
0°
60°E
120°E
Range Type
Native Range
Potential Invasive Range

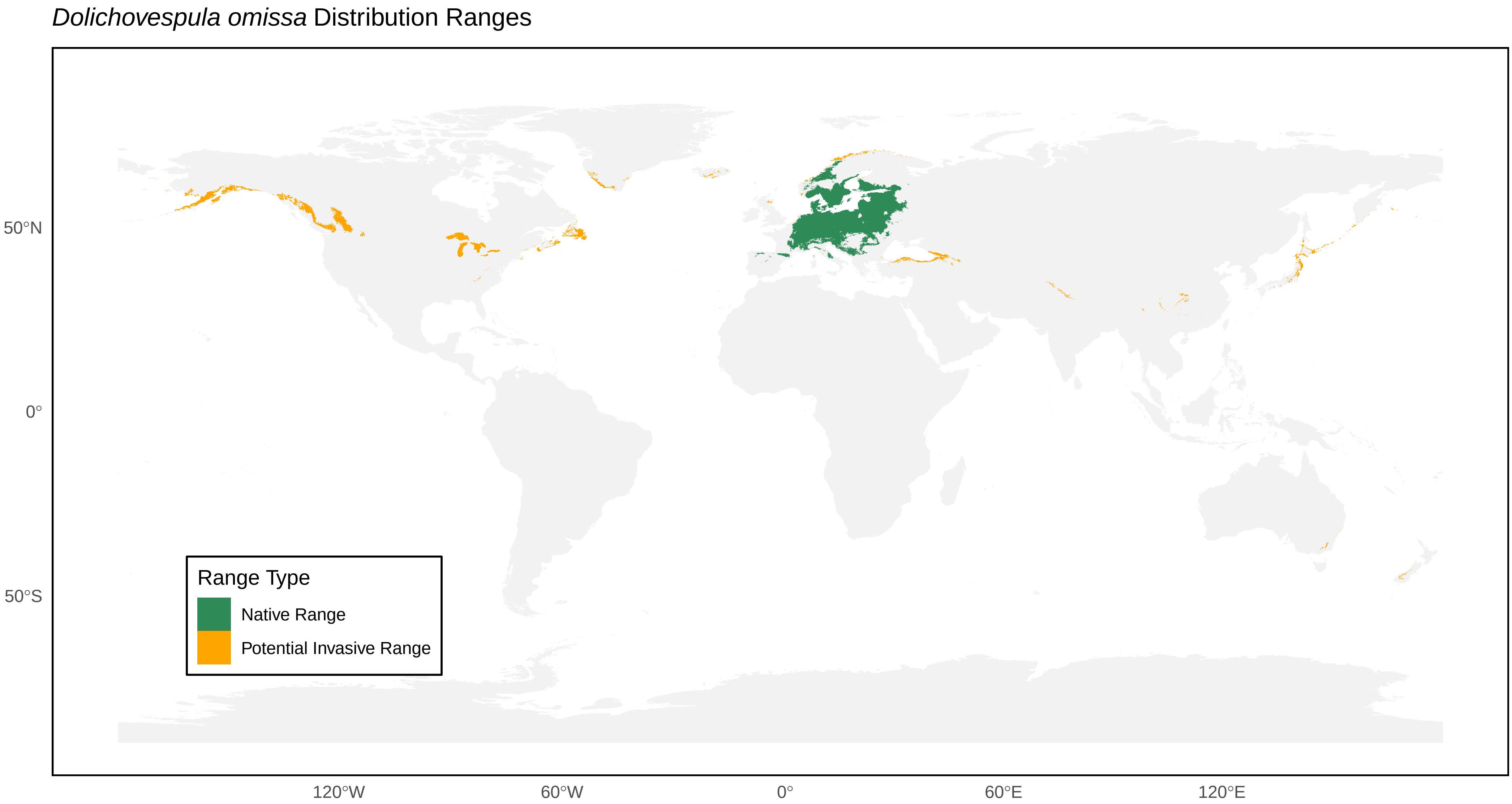

*Dolichovespula omissa* Distribution Ranges
50°N
0°
50°S
120°W
60°W
0°
60°E
120°E
Range Type
Native Range
Potential Invasive Range

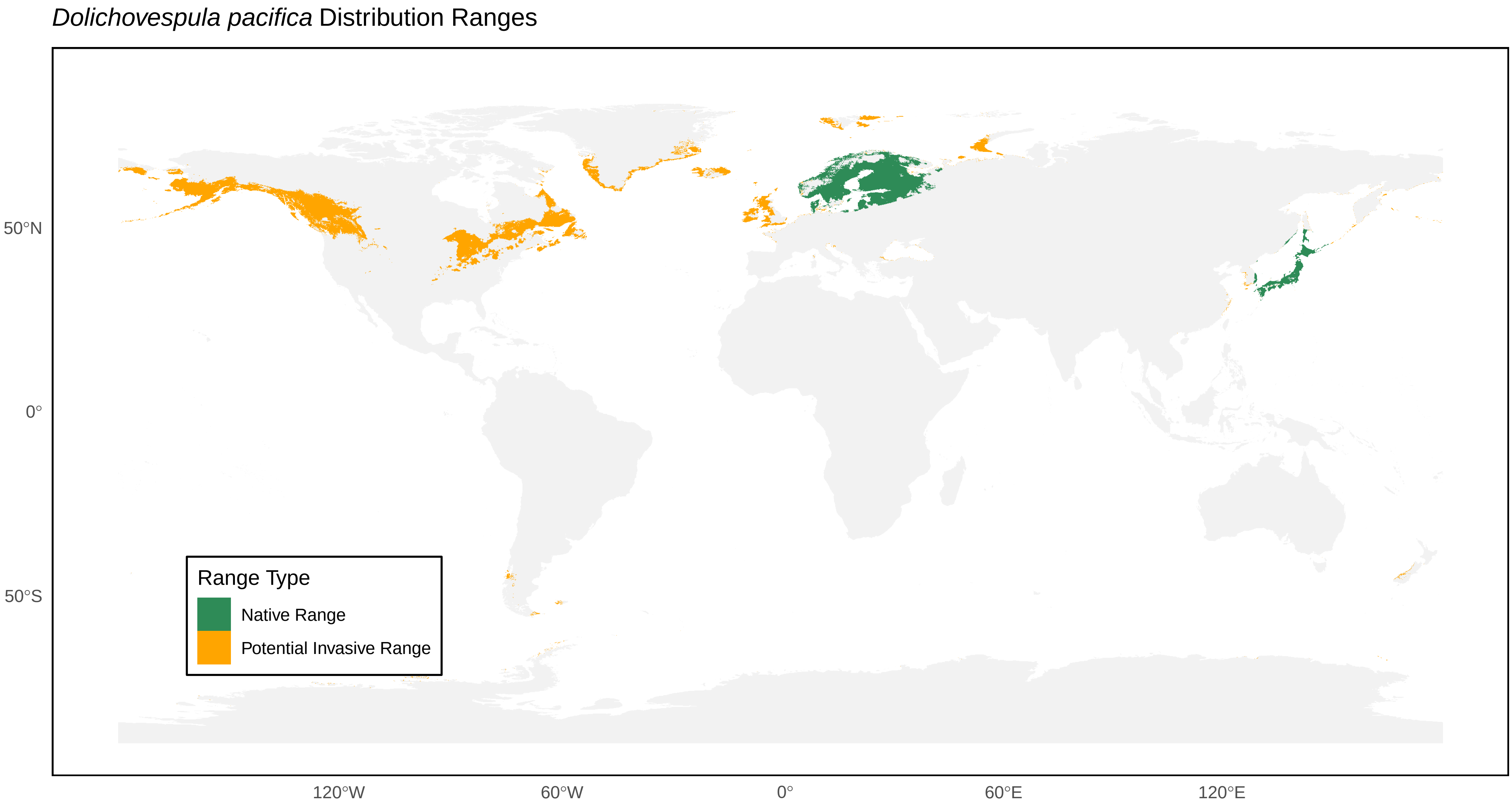

*Dolichovespula pacifica* Distribution Ranges
50°N
0°
50°S
120°W
60°W
0°
60°E
120°E
Range Type
Native Range
Potential Invasive Range

*Dolichovespula saxonica* Distribution Ranges

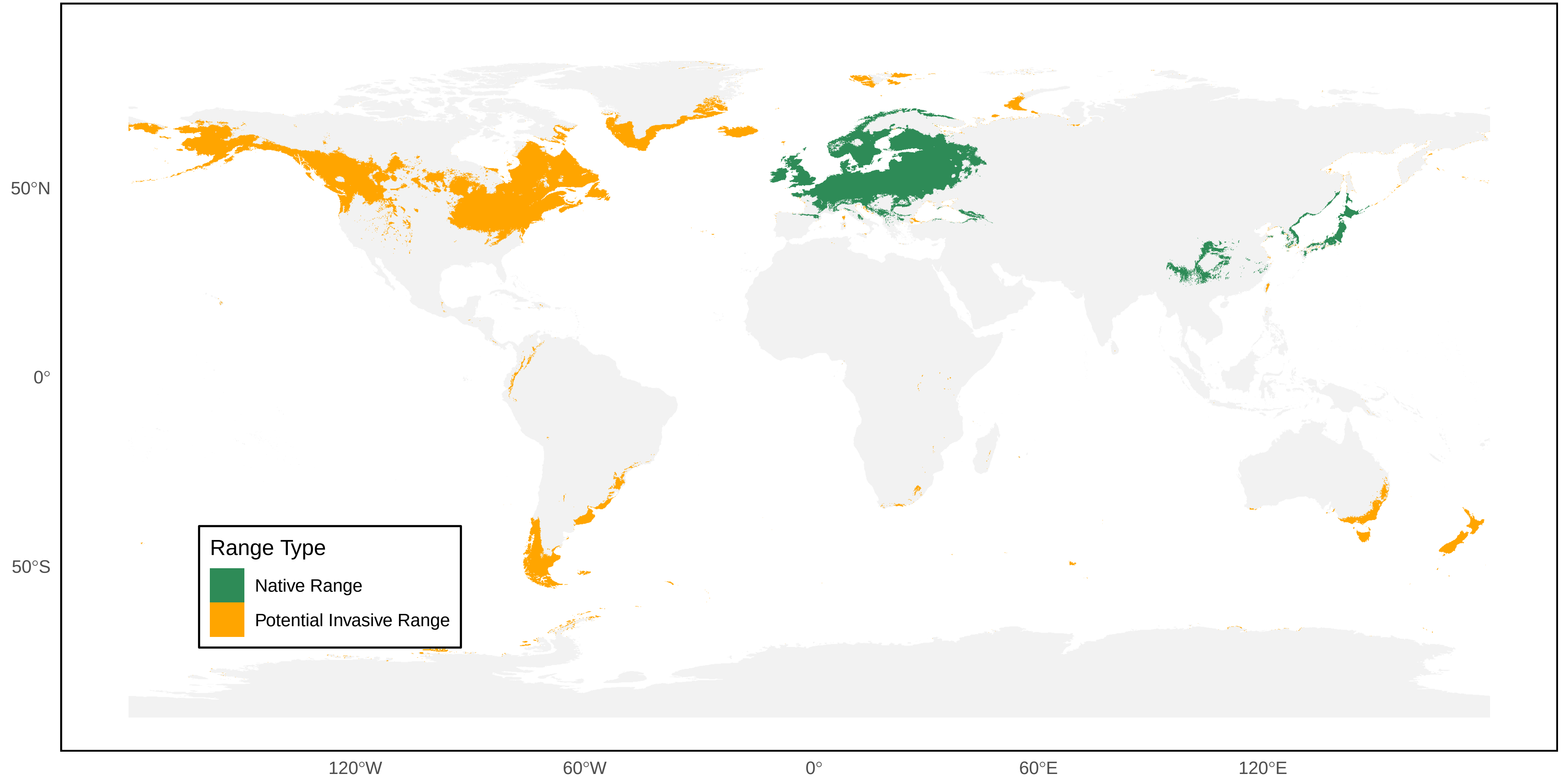

*Dolichovespula sylvestris* Distribution Ranges

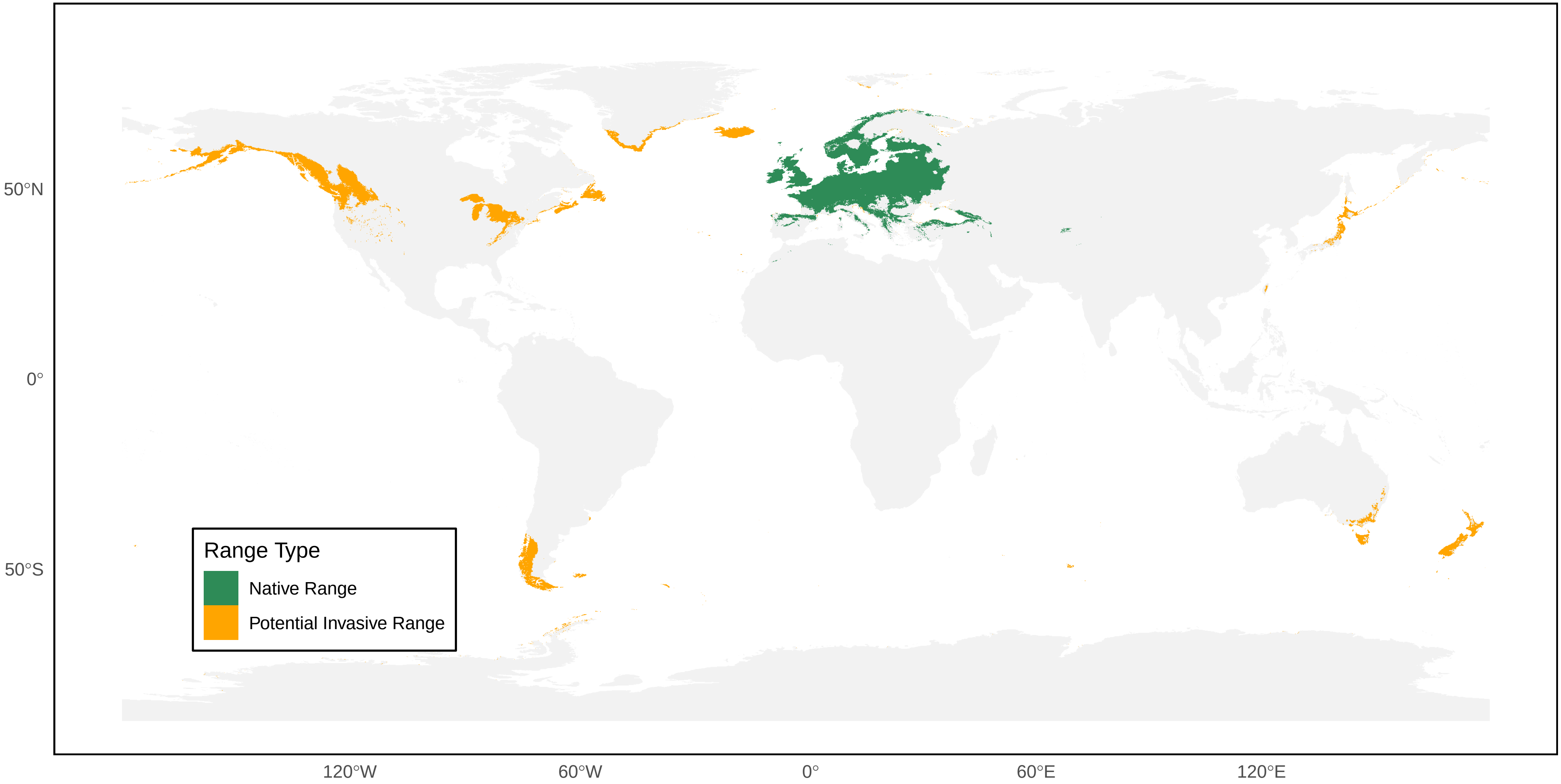

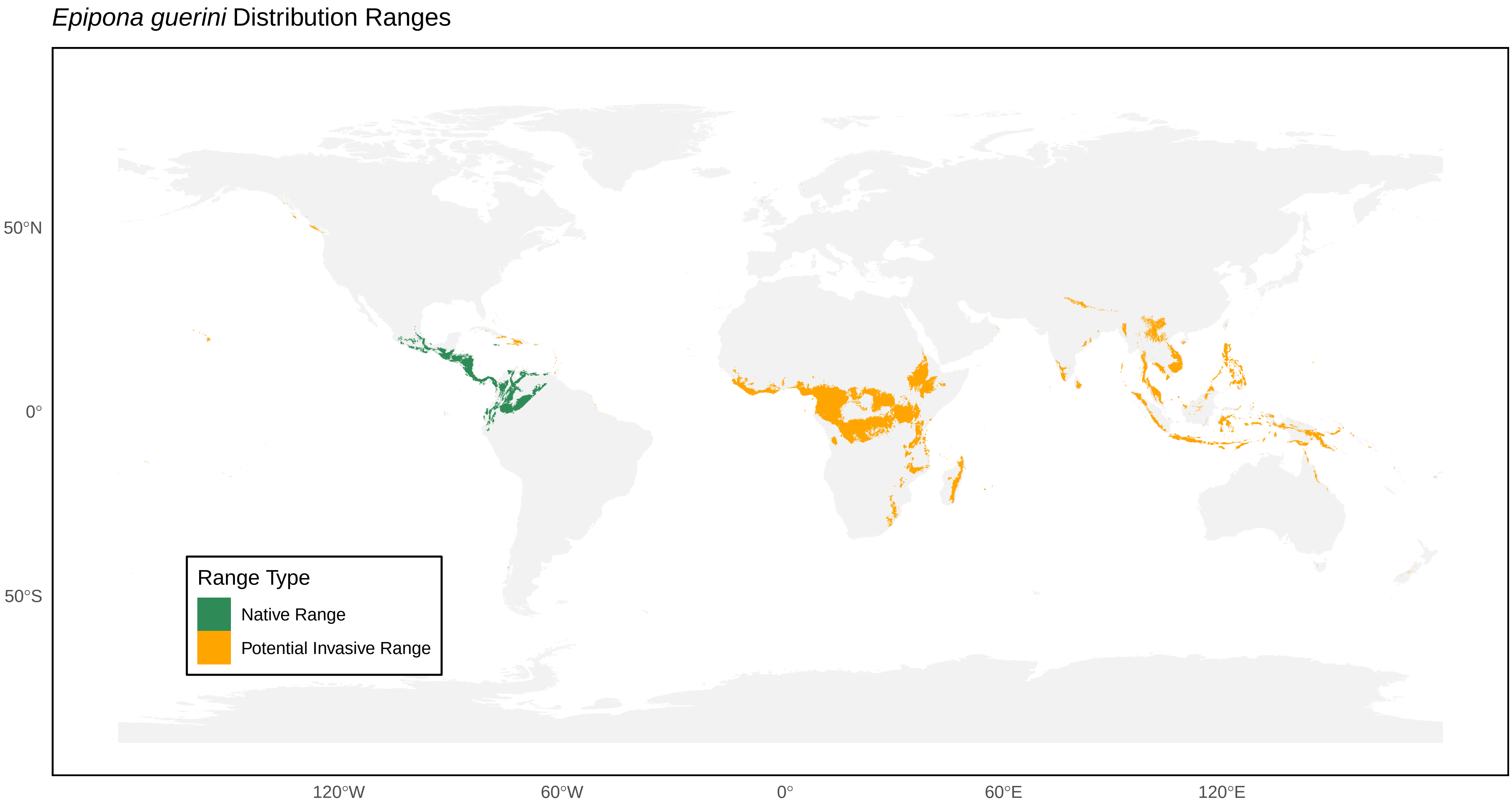
*Epipona guerini* Distribution Ranges
50°N
0°
50°S
120°W
60°W
0°
60°E
120°E
Range Type
Native Range
Potential Invasive Range

*Epipona niger* Distribution Ranges

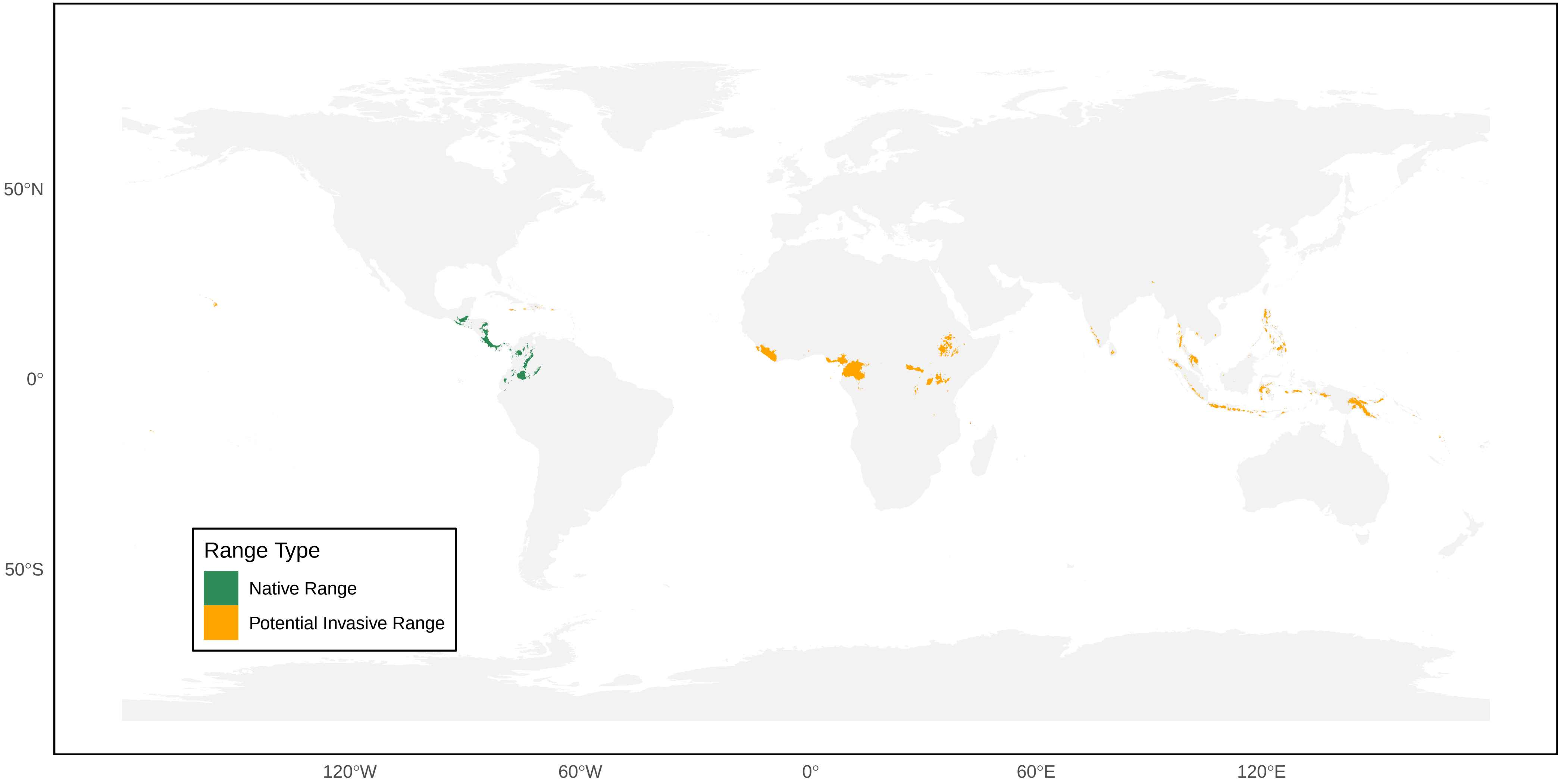

*Eustenogaster micans* Distribution Ranges

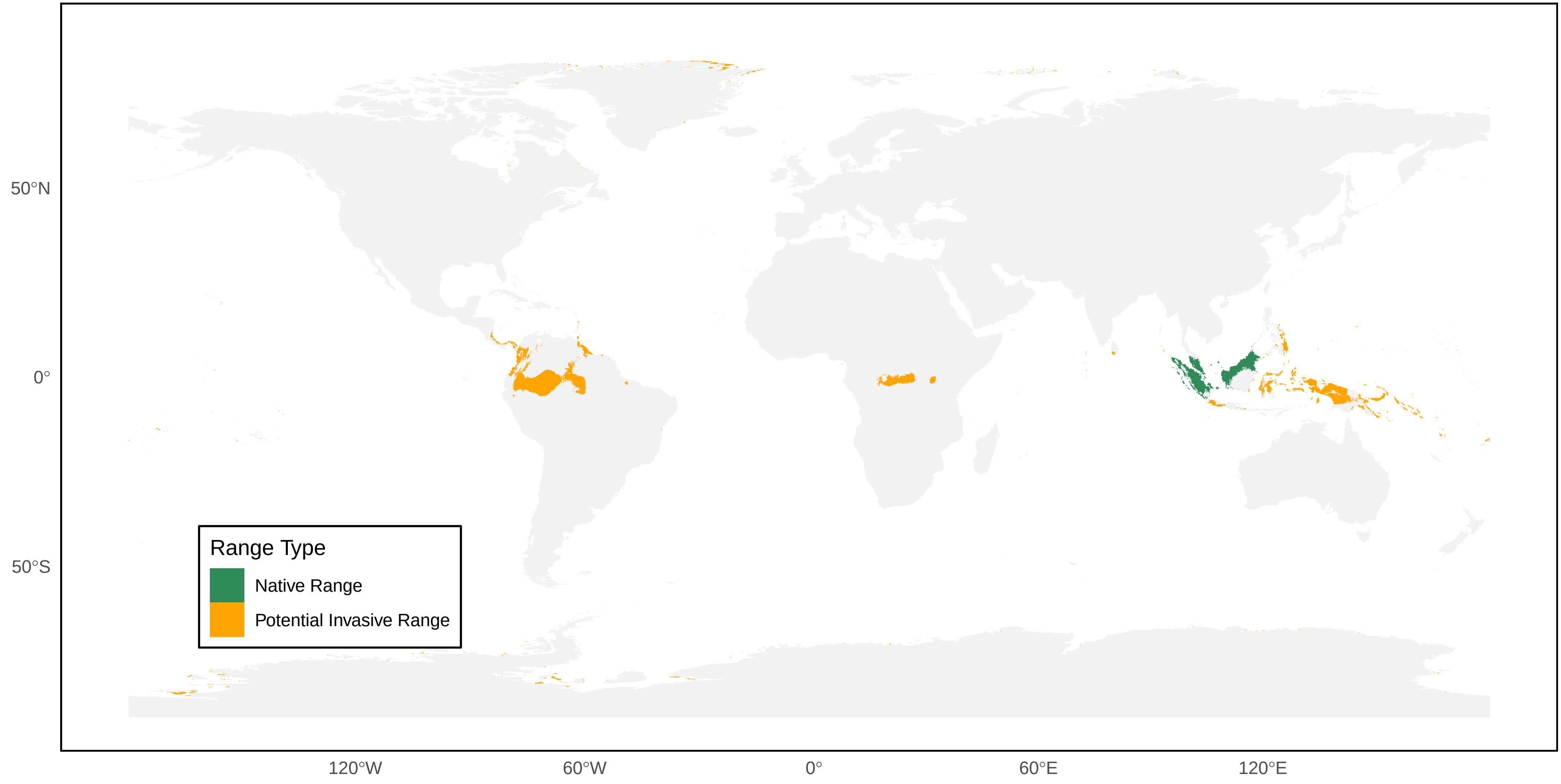

*Eustenogaster nigra* Distribution Ranges

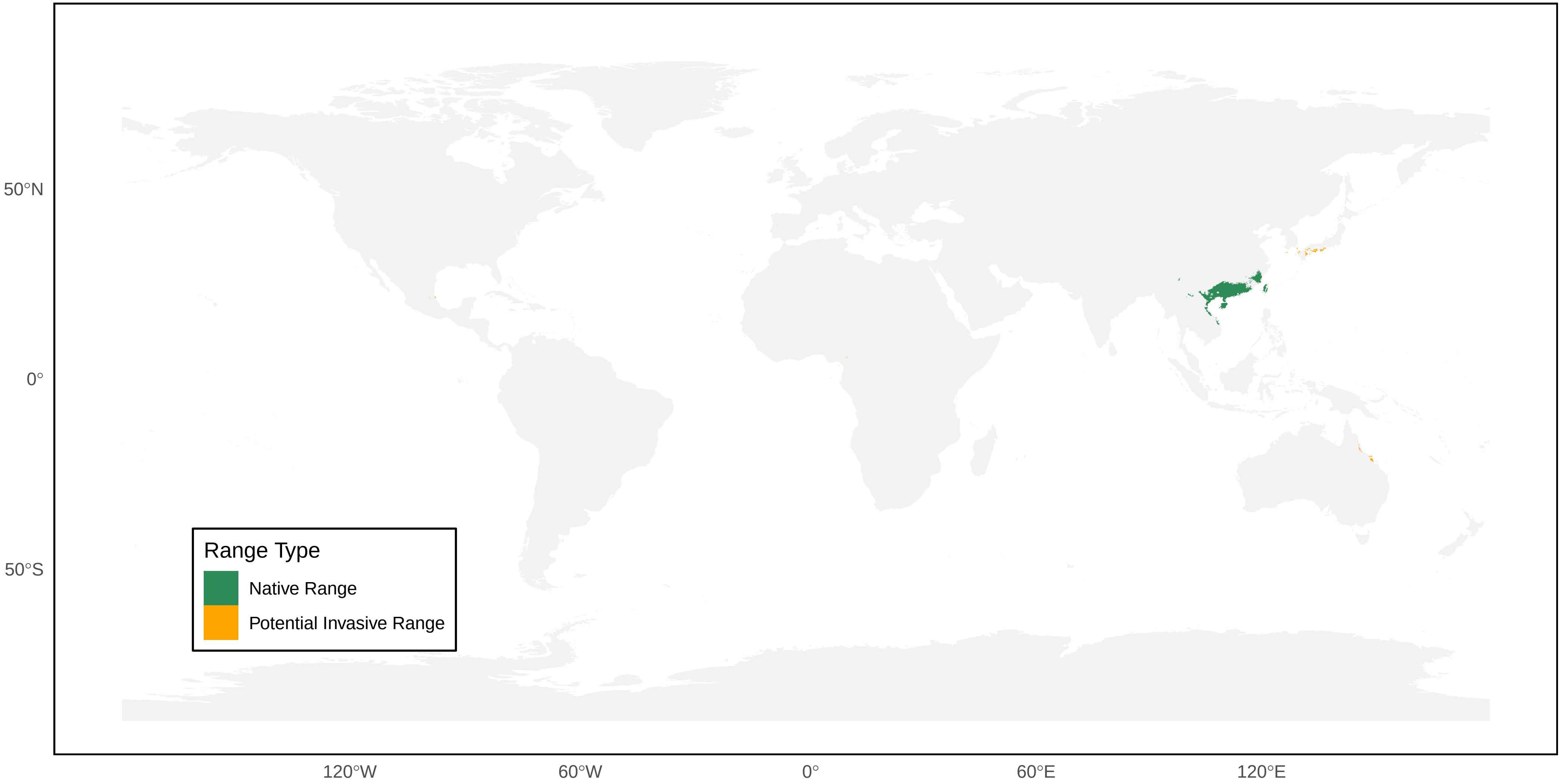

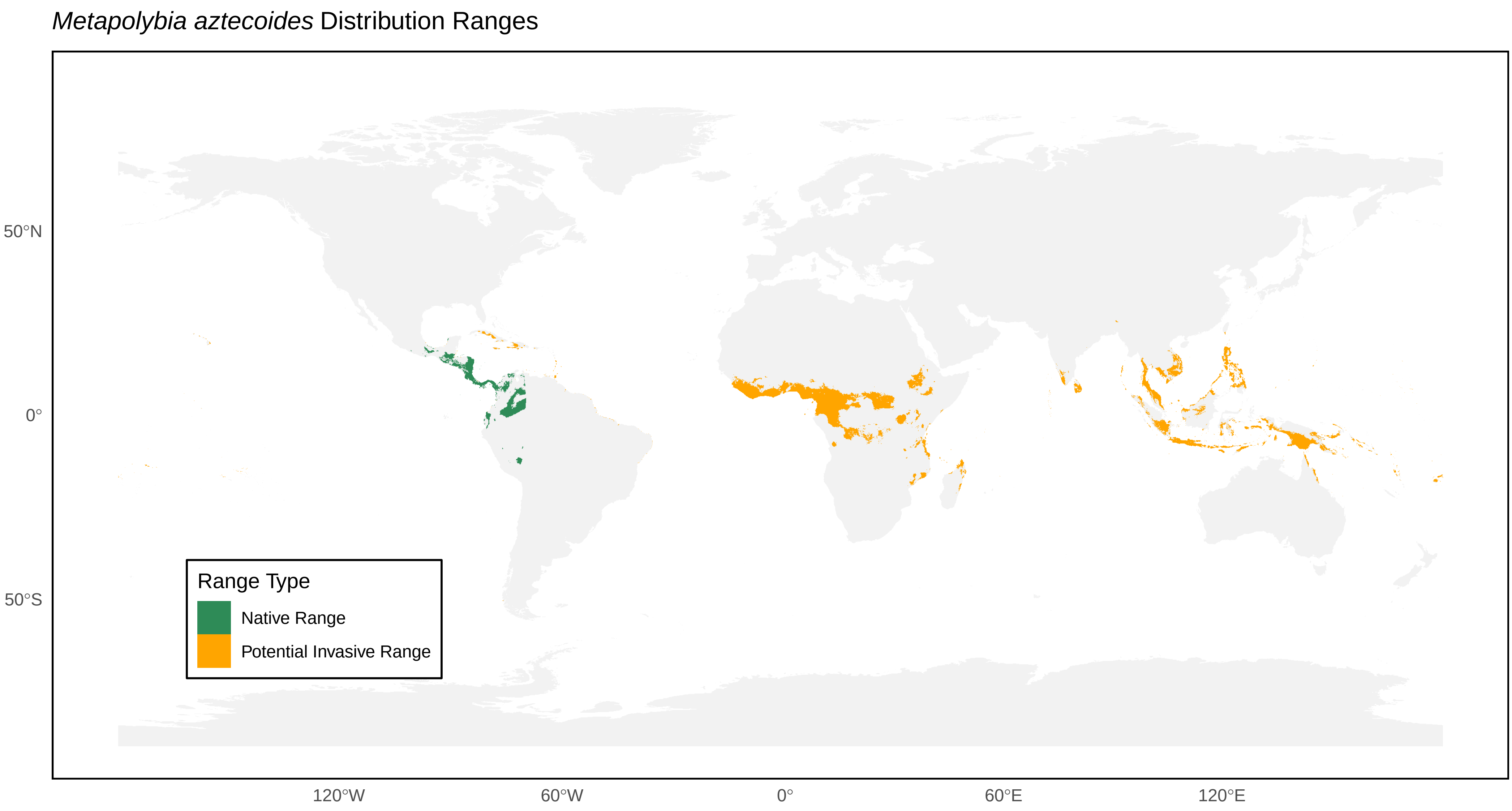
*Metapolybia aztecoides* Distribution Ranges
50°N
0°
50°S
120°W
60°W
0°
60°E
120°E
Range Type
Native Range
Potential Invasive Range

*Metapolybia cingulata* Distribution Ranges

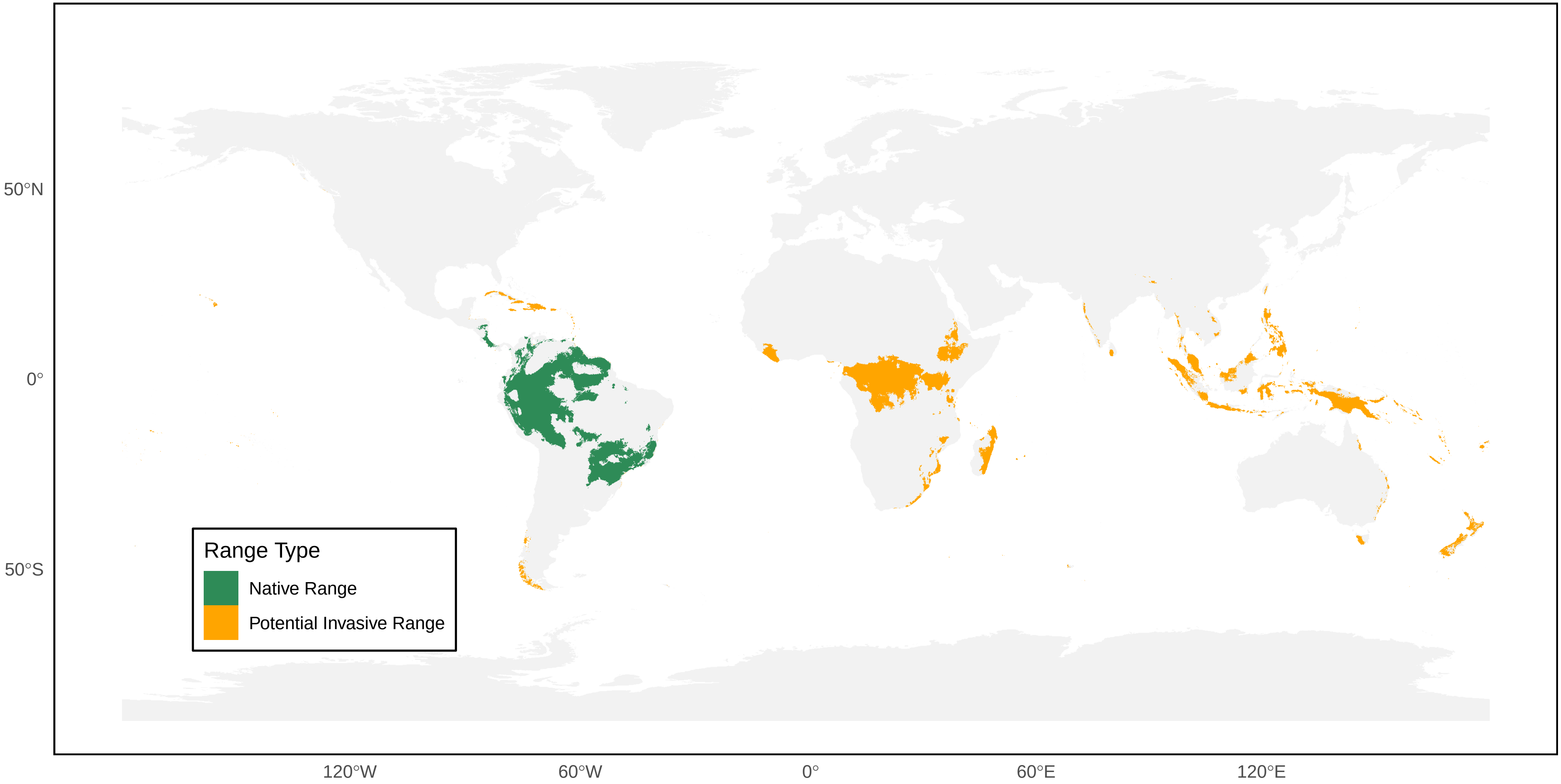

*Mischocyttarus angulatus* Distribution Ranges

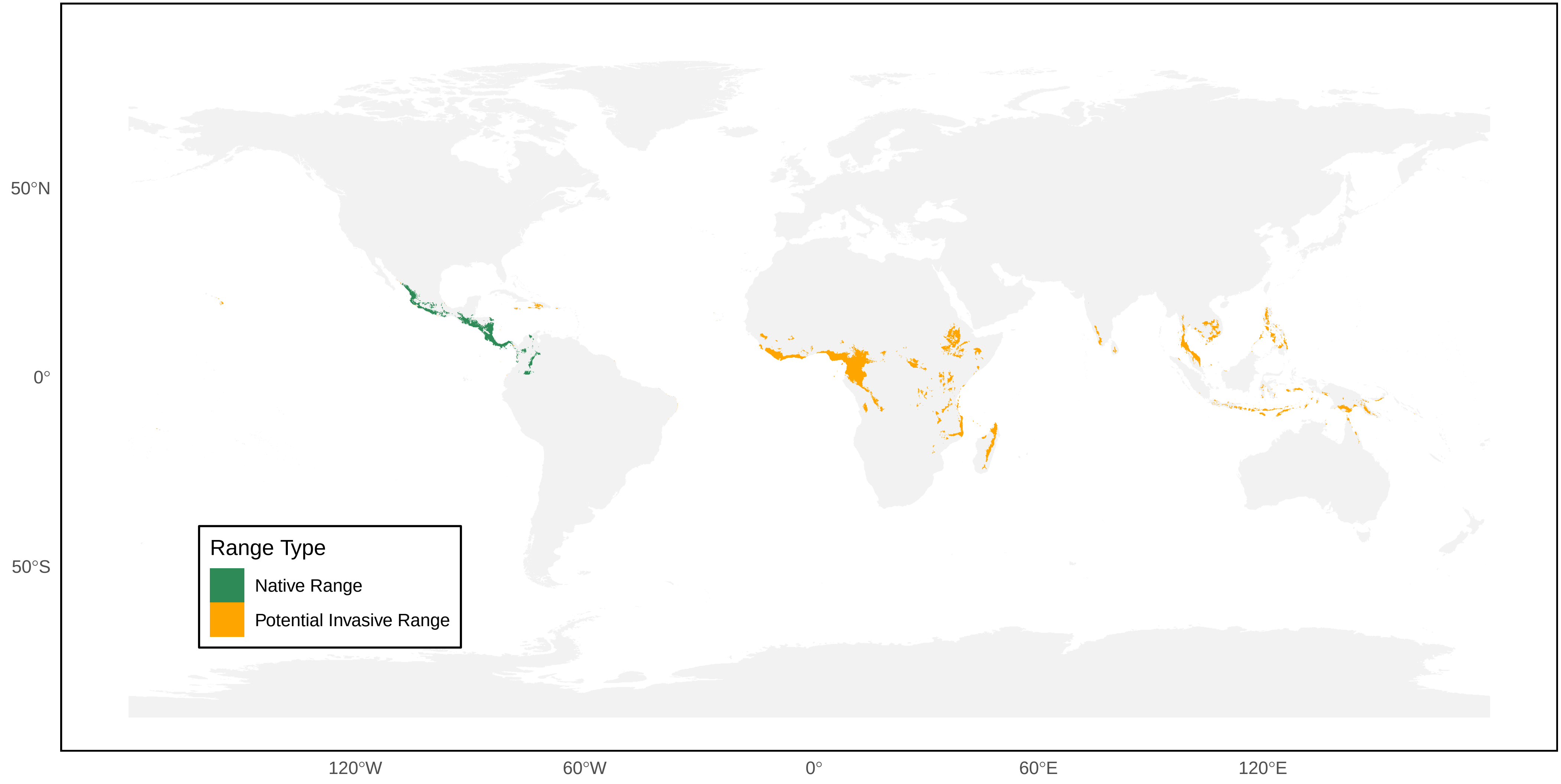

*Mischocyttarus basimacula* Distribution Ranges

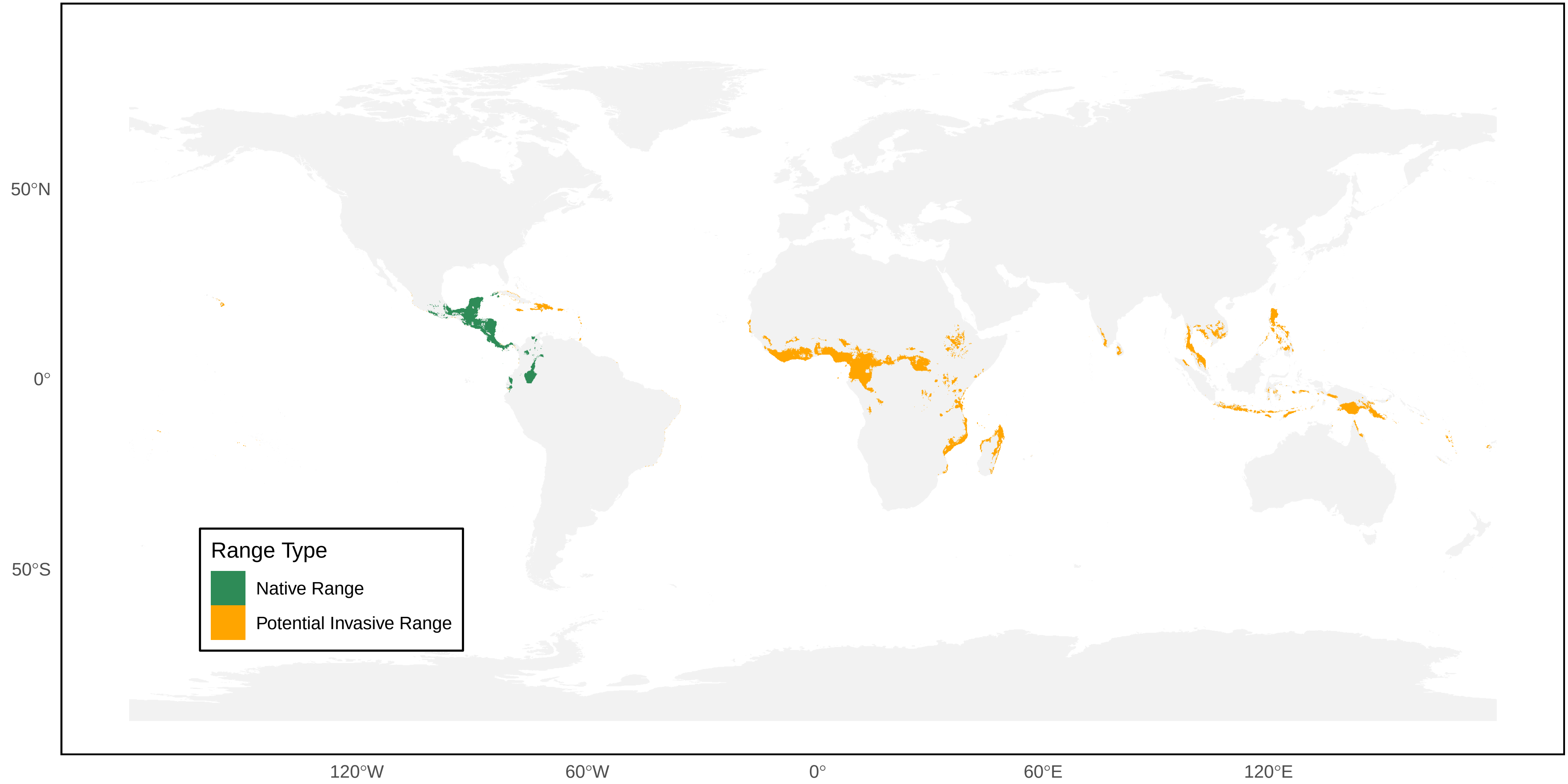

*Mischocyttarus cerberus* Distribution Ranges

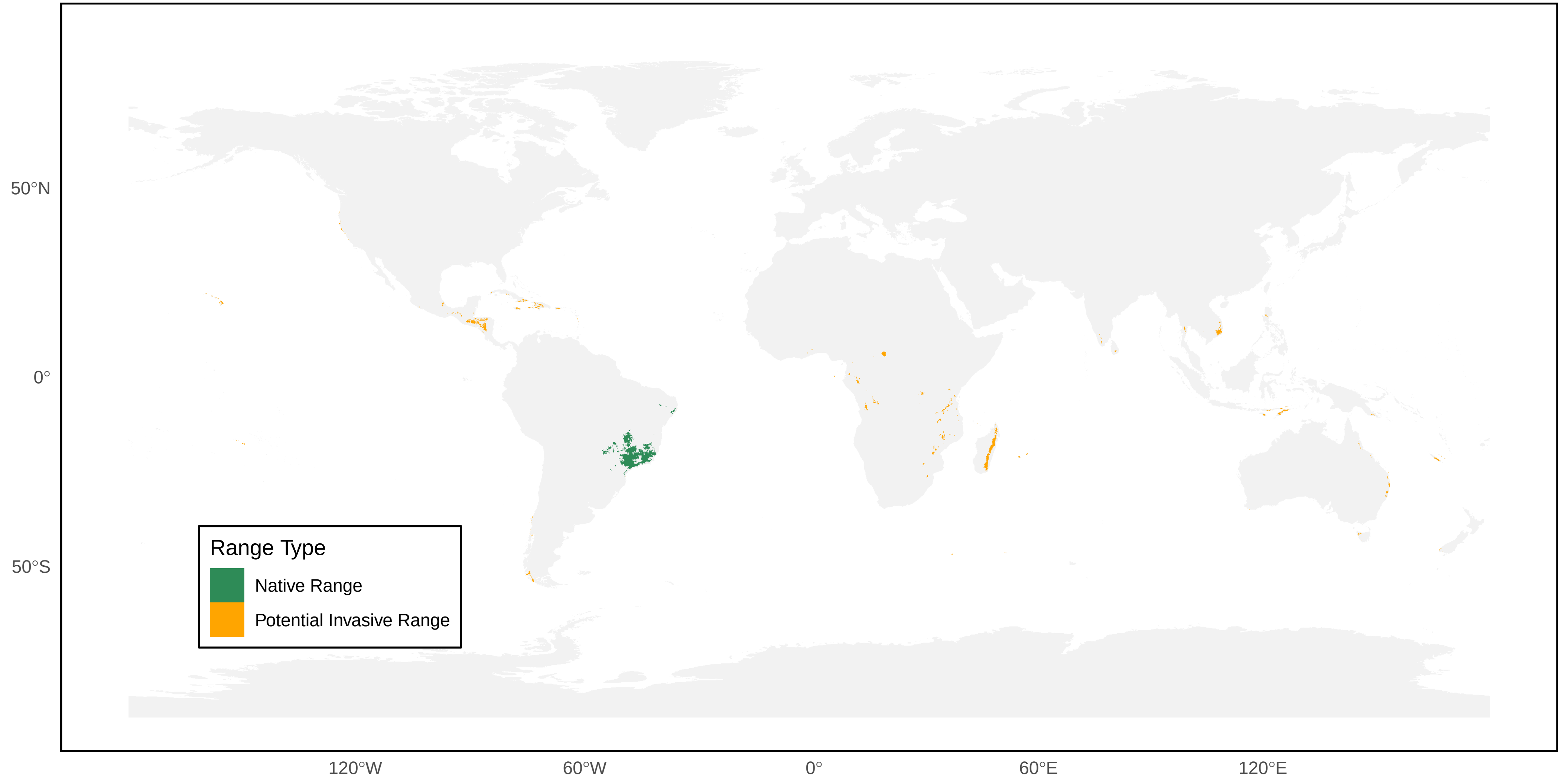

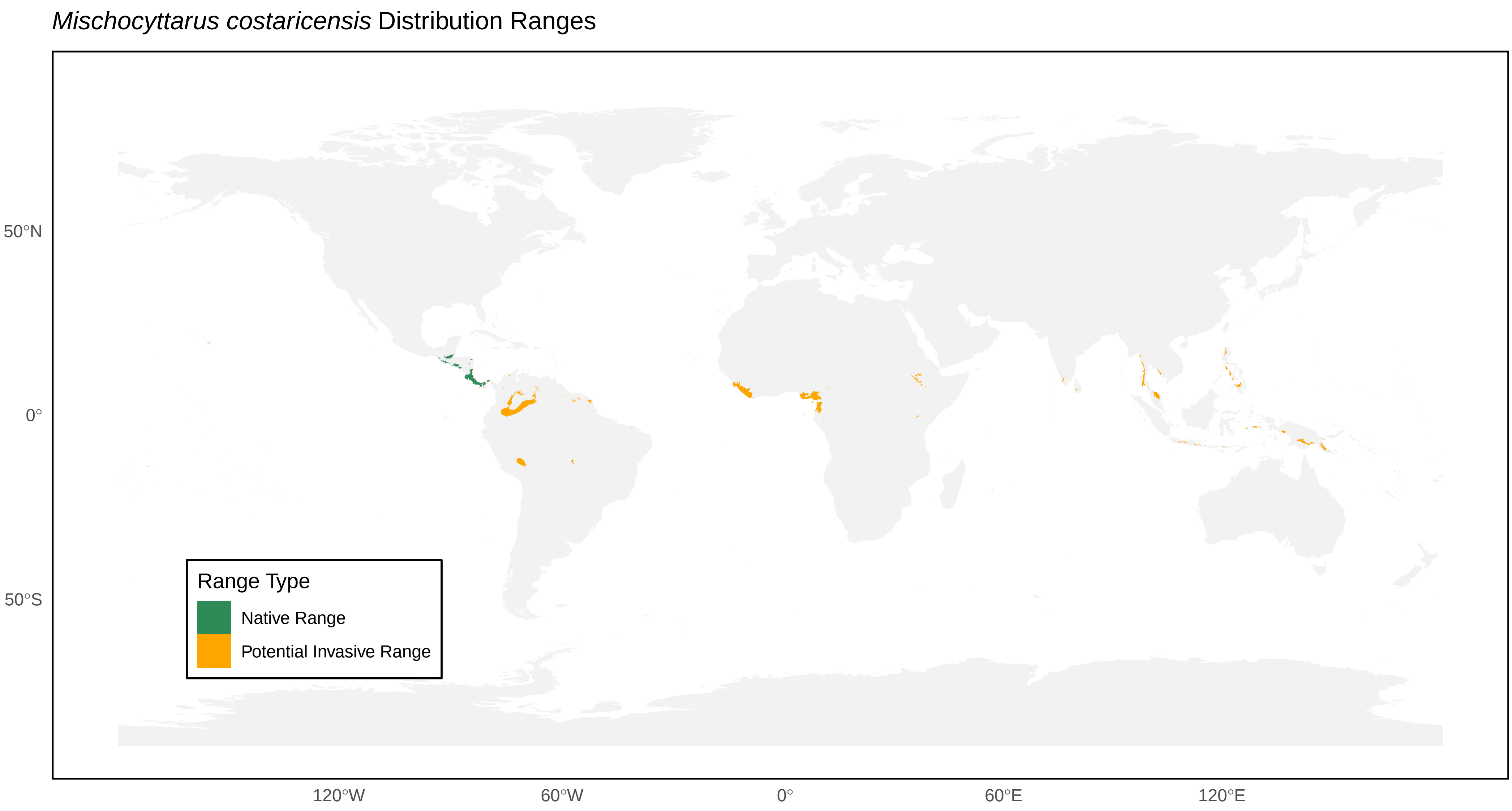

*Mischocyttarus costaricensis* Distribution Ranges
50°N
0°
50°S
120°W
60°W
0°
60°E
120°E
Range Type
Native Range
Potential Invasive Range

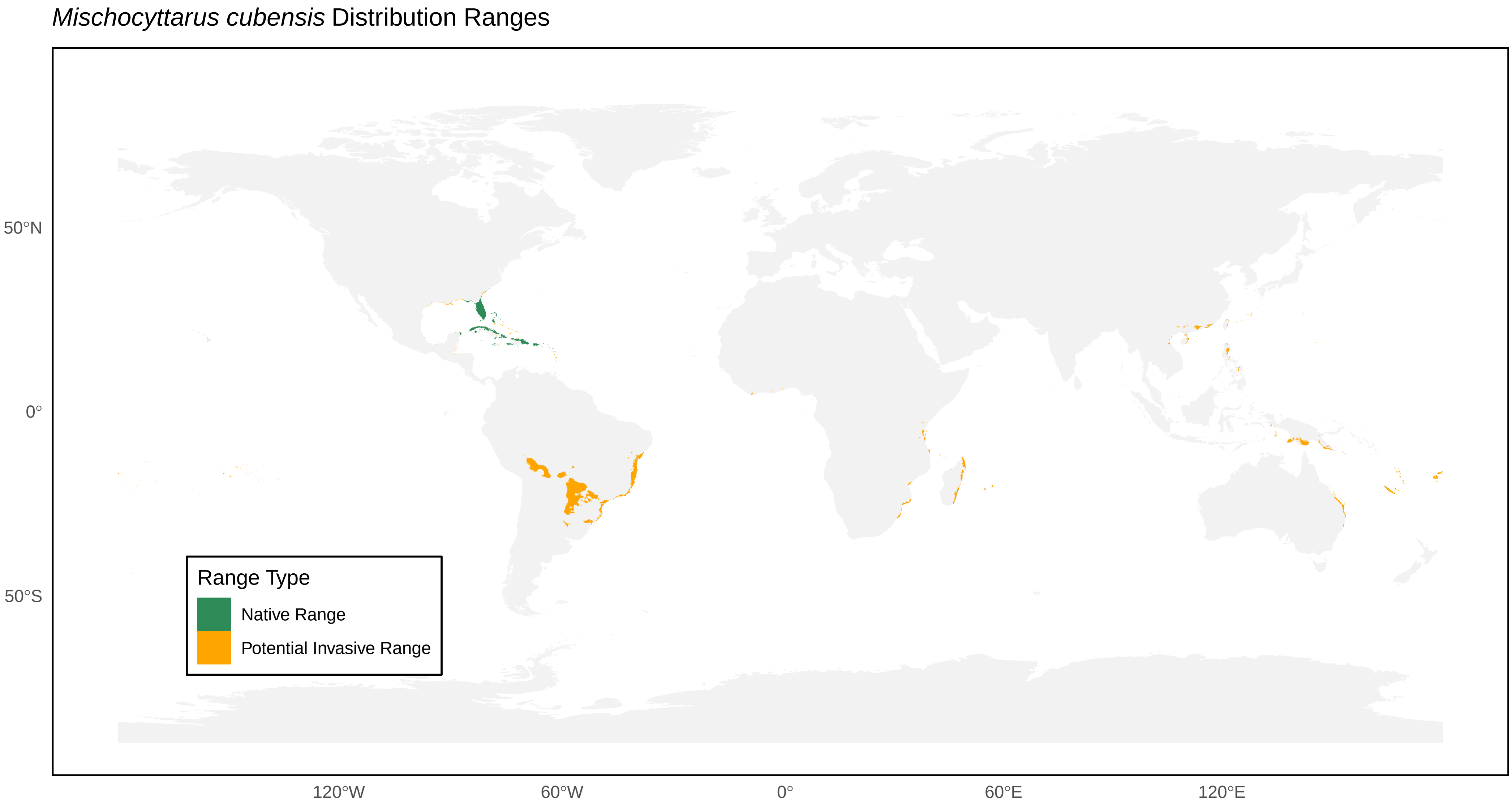
*Mischocyttarus cubensis* Distribution Ranges
50°N
0°
50°S
120°W
60°W
0°
60°E
120°E
Range Type
Native Range
Potential Invasive Range

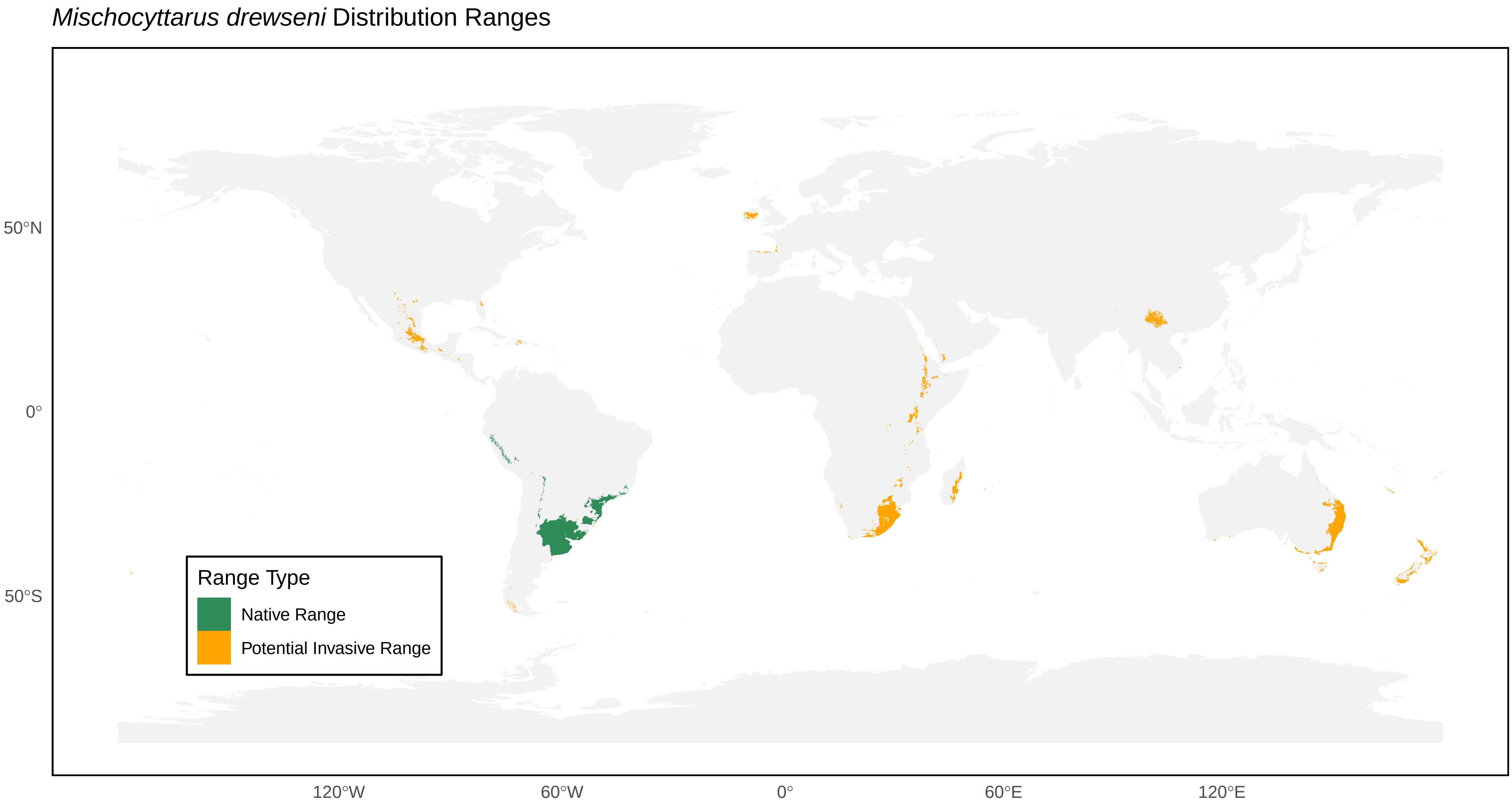
*Mischocyttarus drewseni* Distribution Ranges
50°N
0°
50°S
120°W
60°W
0°
60°E
120°E
Range Type
Native Range
Potential Invasive Range

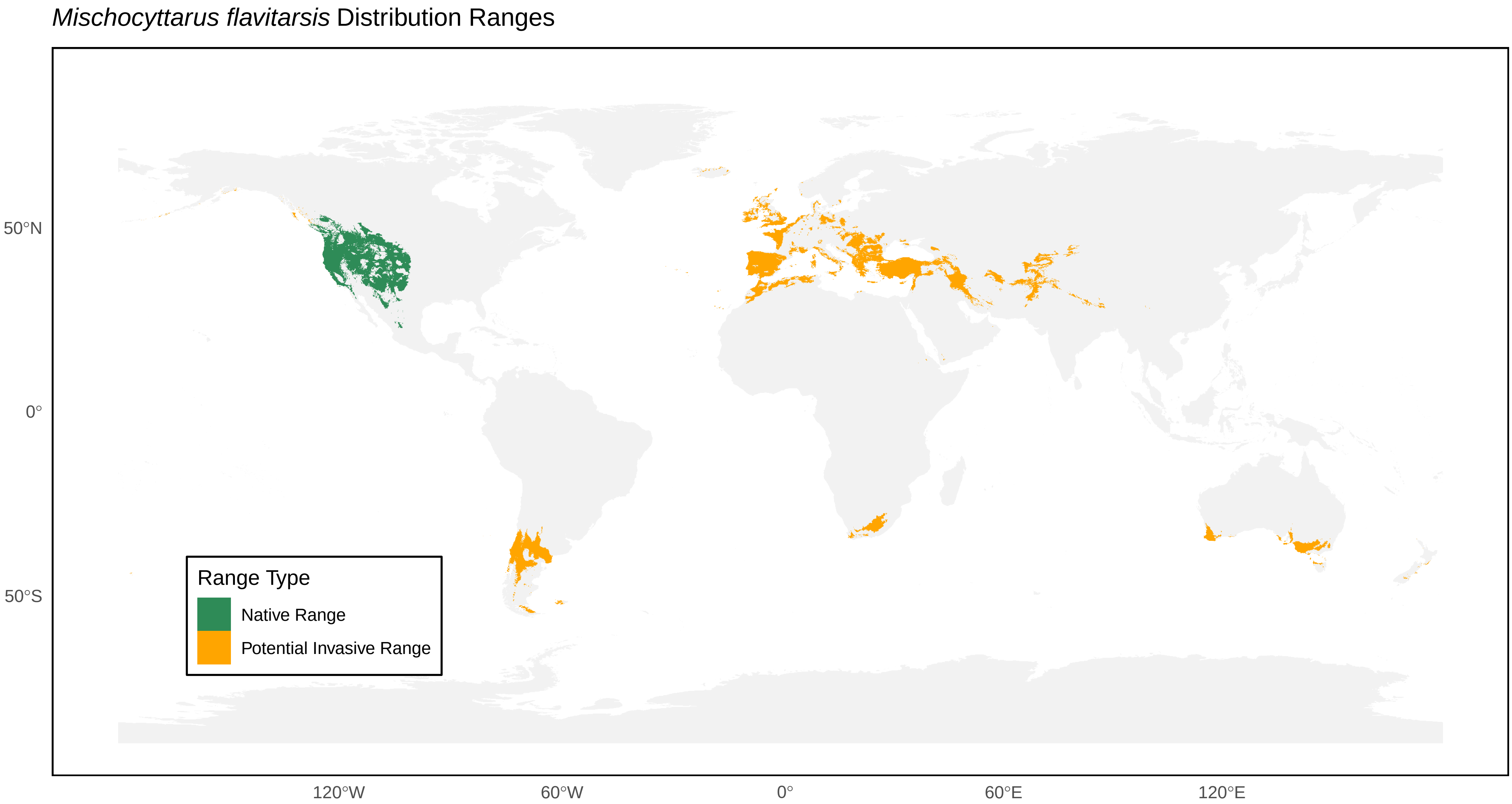

*Mischocyttarus flavitarsis* Distribution Ranges
50°N
0°
50°S
120°W
60°W
0°
60°E
120°E
Range Type
Native Range
Potential Invasive Range

*Mischocyttarus immarginatus* Distribution Ranges

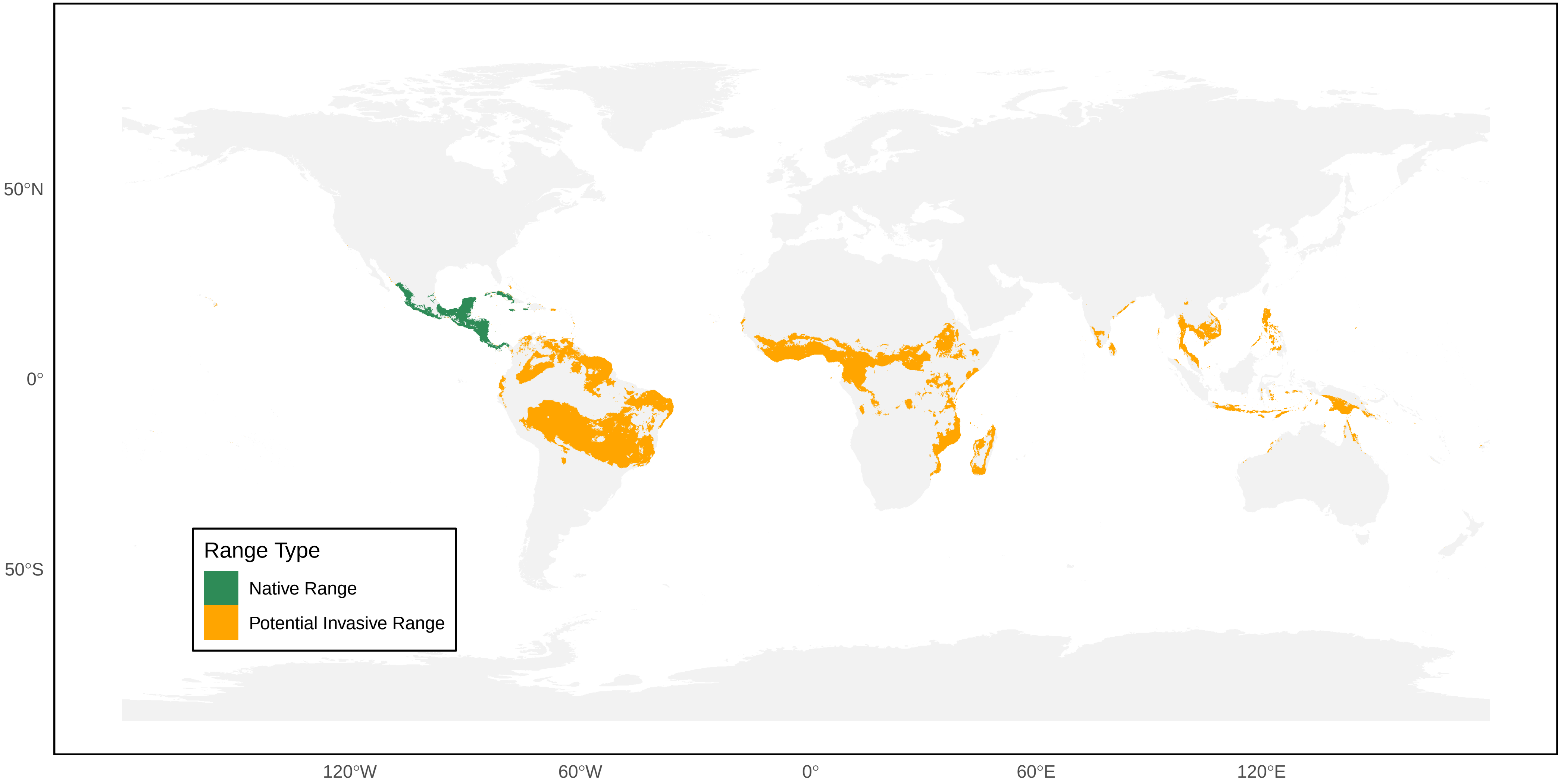

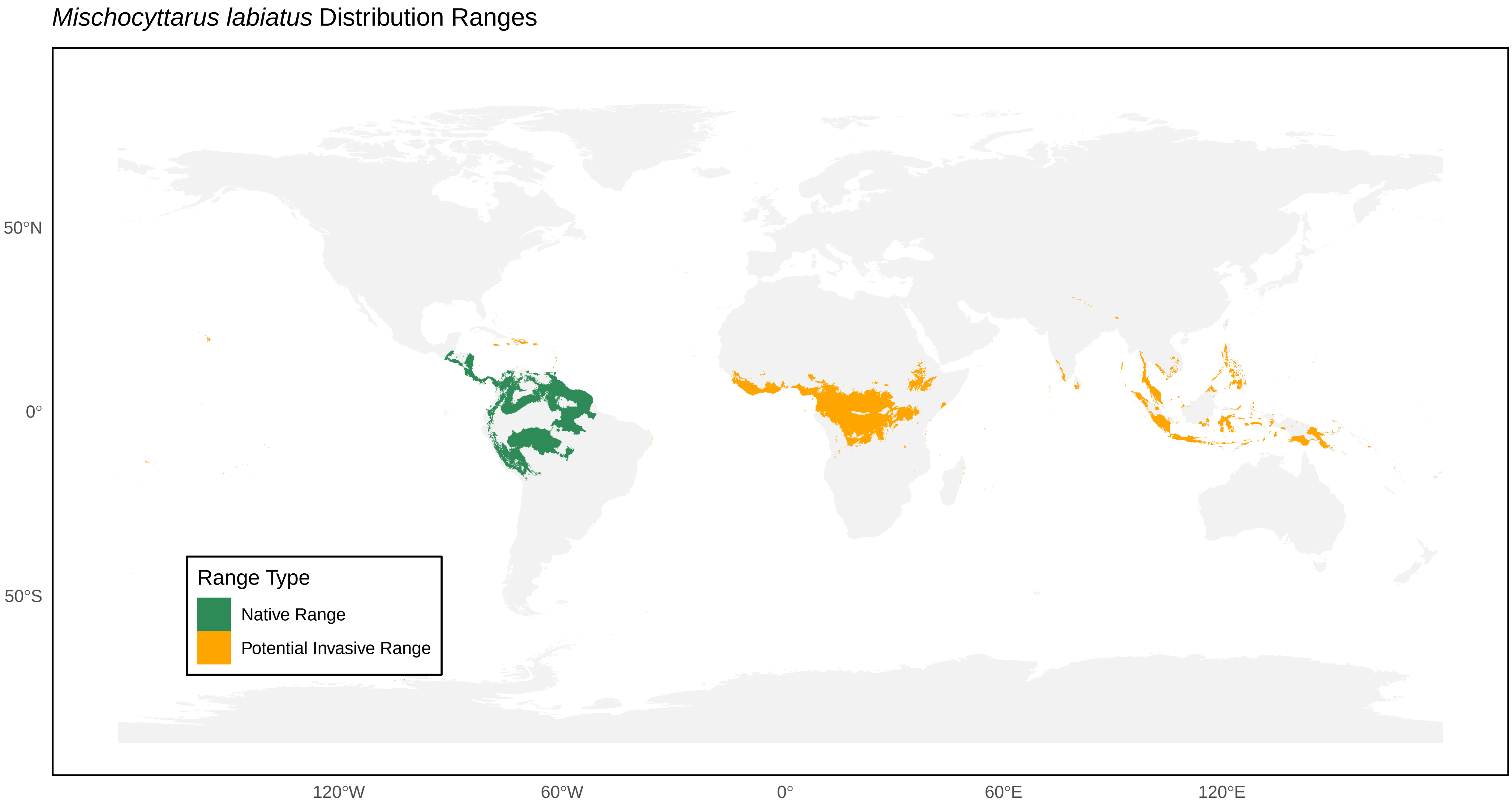
*Mischocyttarus labiatus* Distribution Ranges
50°N
0°
50°S
120°W
60°W
0°
60°E
120°E
Range Type
Native Range
Potential Invasive Range

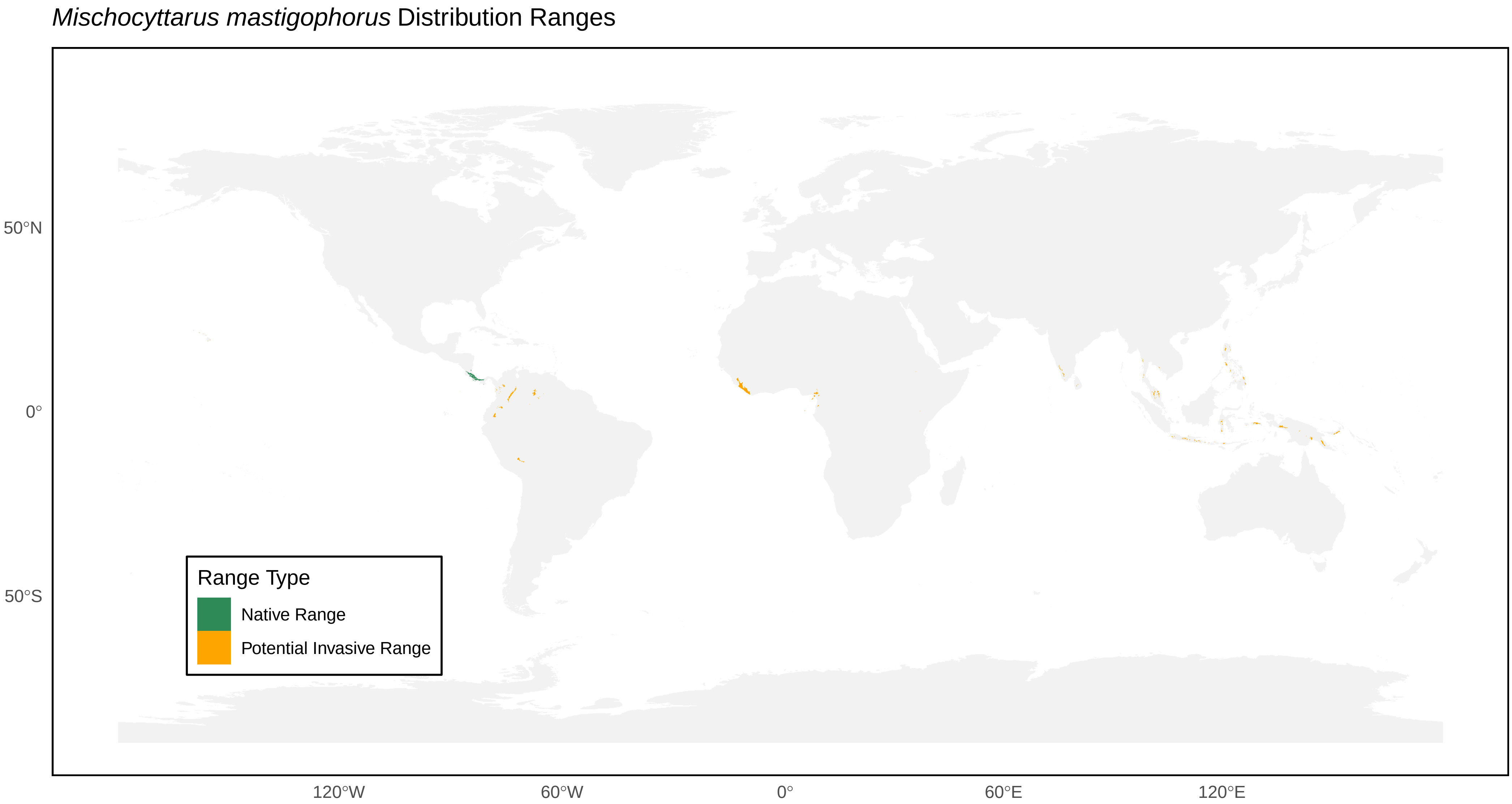

*Mischocyttarus mastigophorus* Distribution Ranges
50°N
0°
50°S
120°W
60°W
0°
60°E
120°E
Range Type
Native Range
Potential Invasive Range

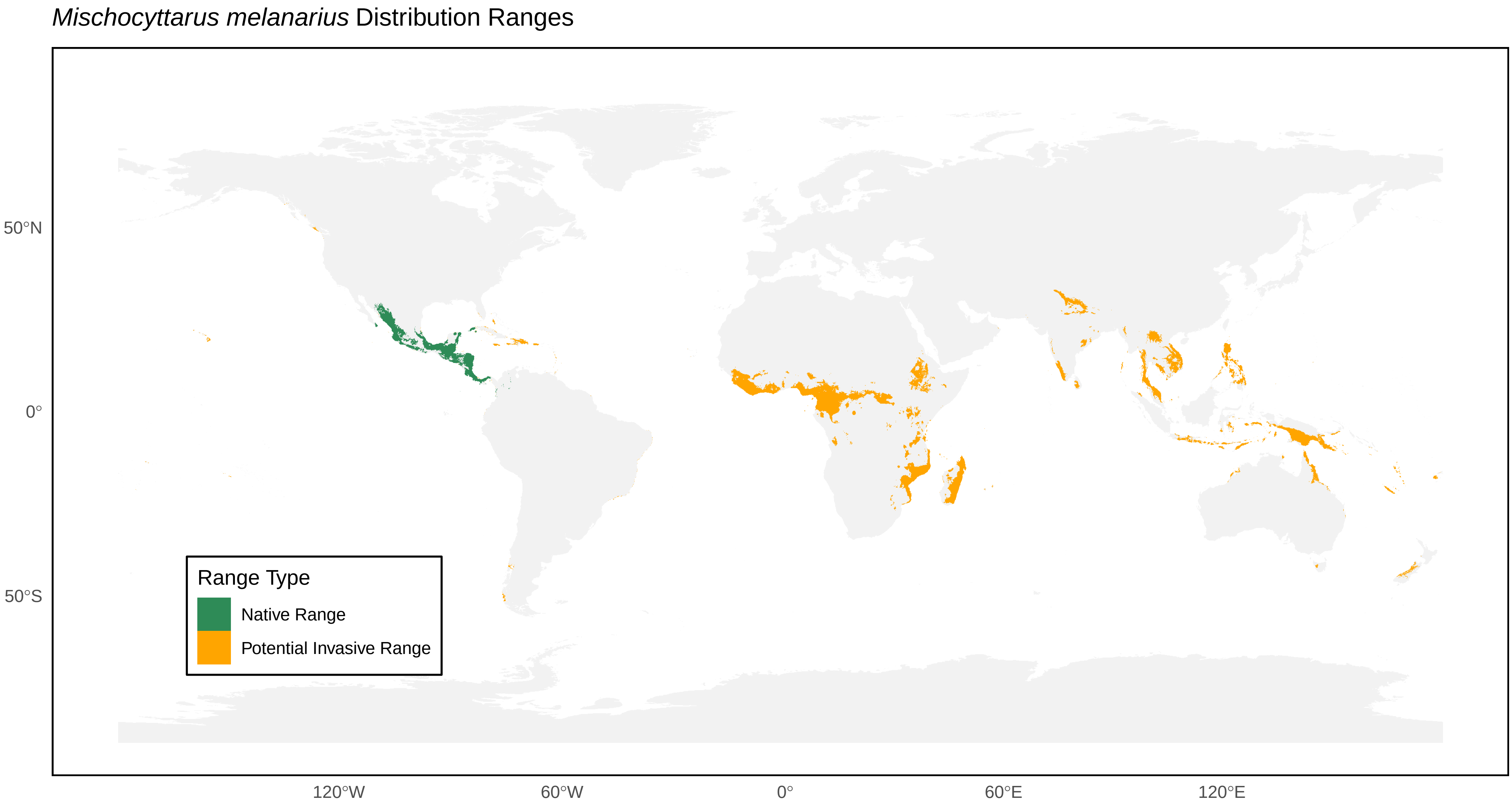
*Mischocyttarus melanarius* Distribution Ranges
50°N
0°
50°S
120°W
60°W
0°
60°E
120°E
Range Type
Native Range
Potential Invasive Range

*Mischocyttarus mexicanus* Distribution Ranges

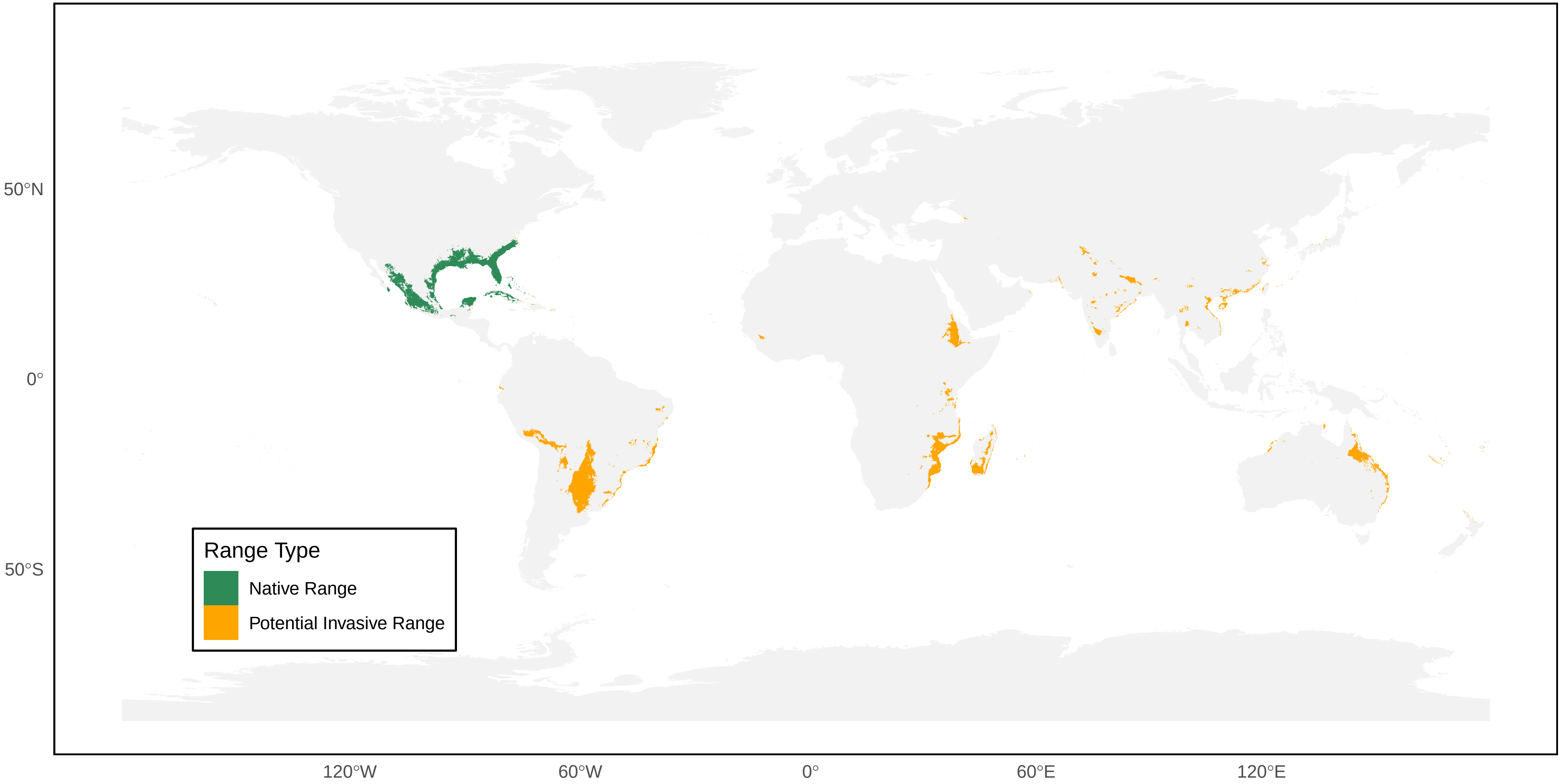

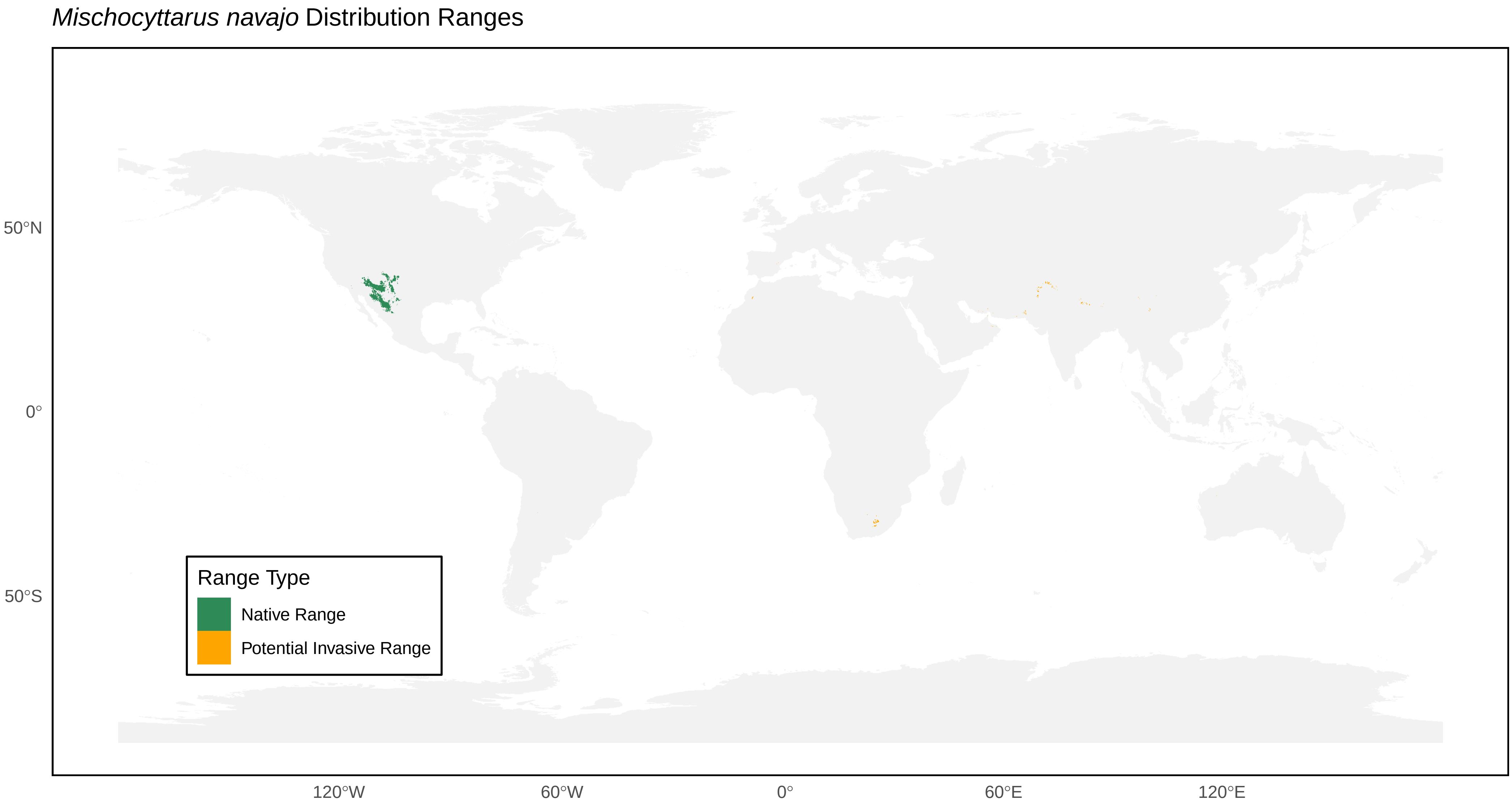
*Mischocyttarus navajo* Distribution Ranges
50°N
0°
50°S
120°W
60°W
0°
60°E
120°E
Range Type
Native Range
Potential Invasive Range

*Mischocyttarus pallidipectus* Distribution Ranges

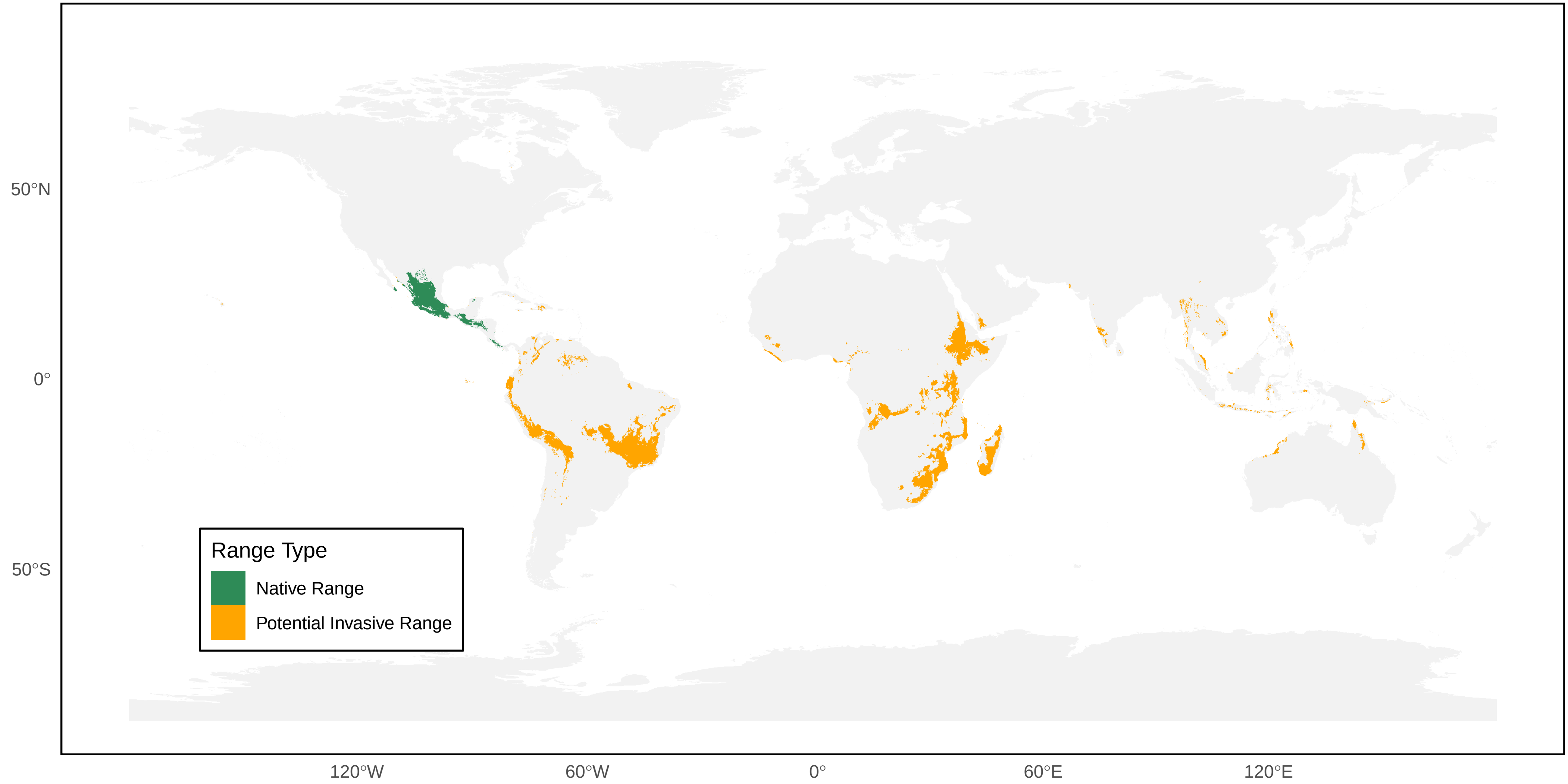

*Mischocyttarus phthisicus* Distribution Ranges

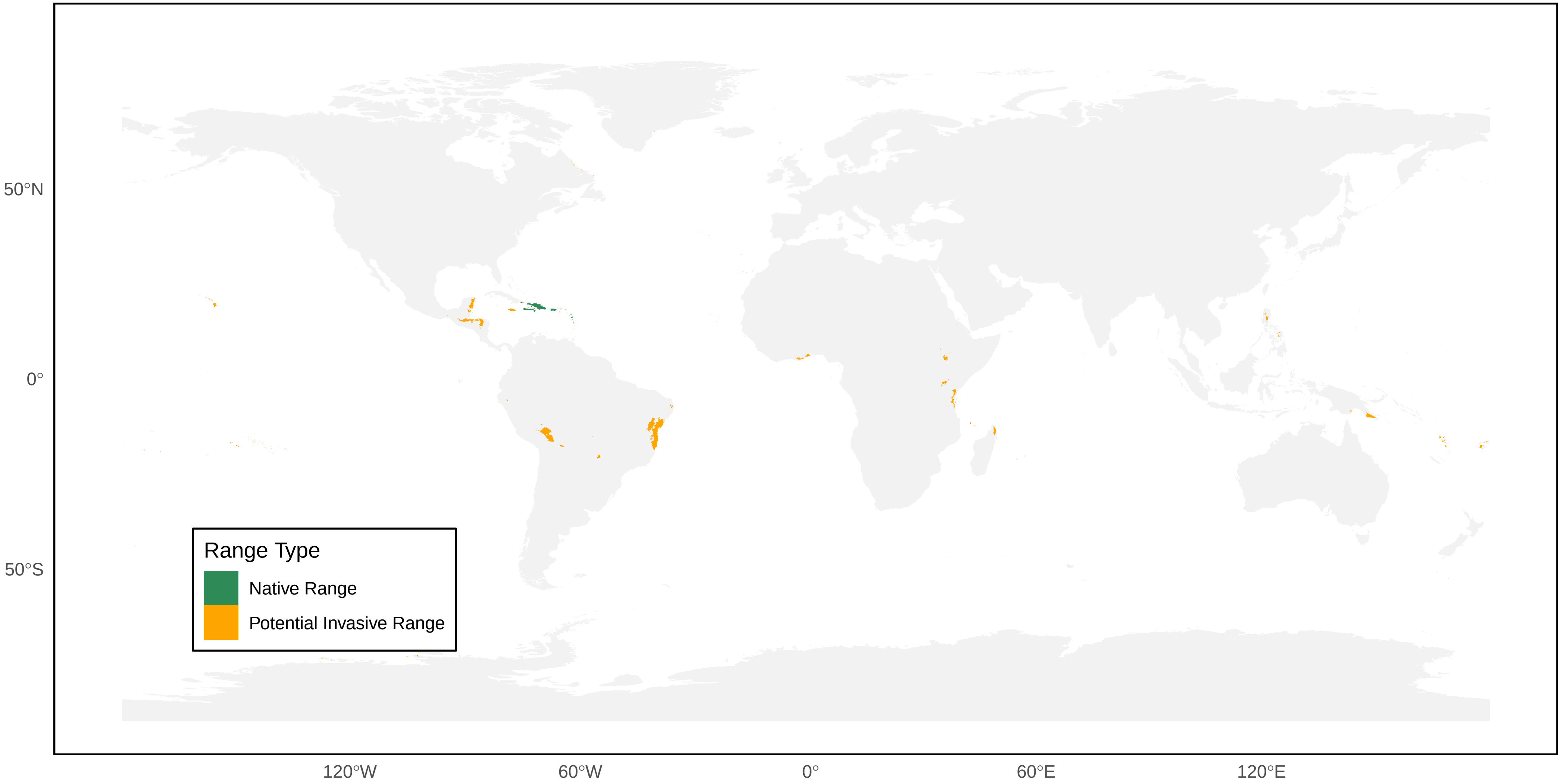

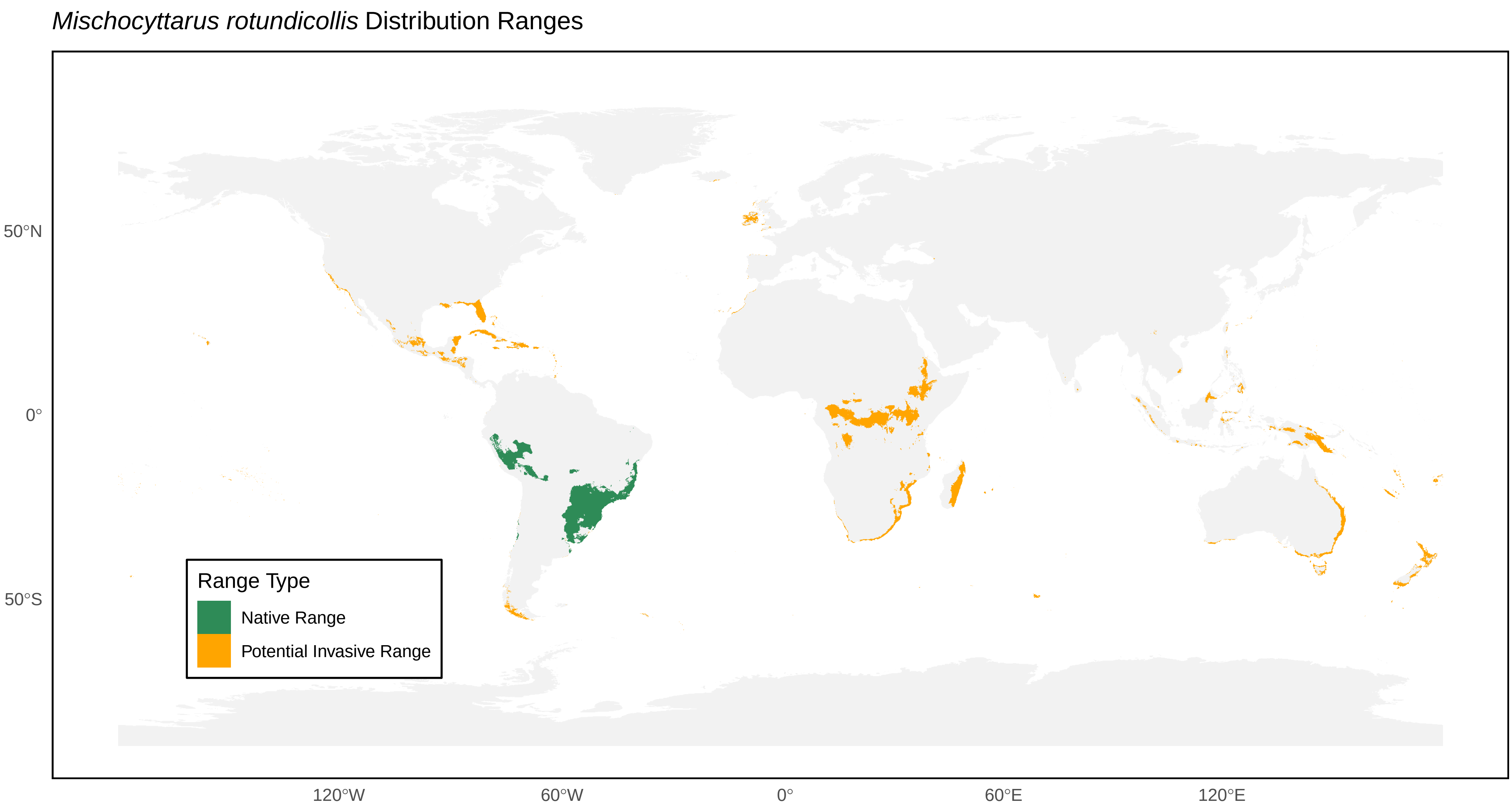

*Mischocyttarus rotundicollis* Distribution Ranges
50°N
0°
50°S
120°W
60°W
0°
60°E
120°E
Range Type
Native Range
Potential Invasive Range

*Mischocyttarus rufidens* Distribution Ranges

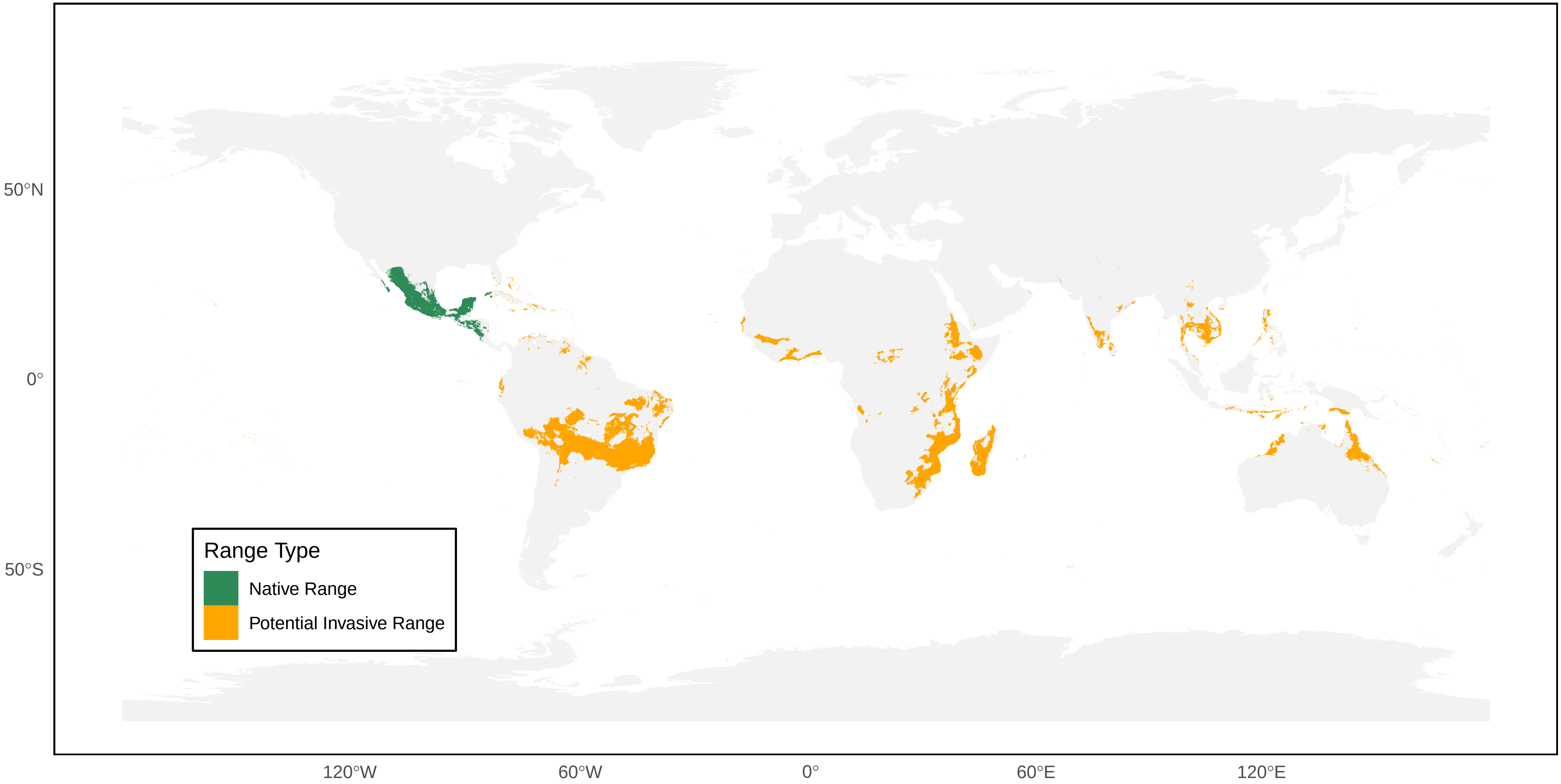

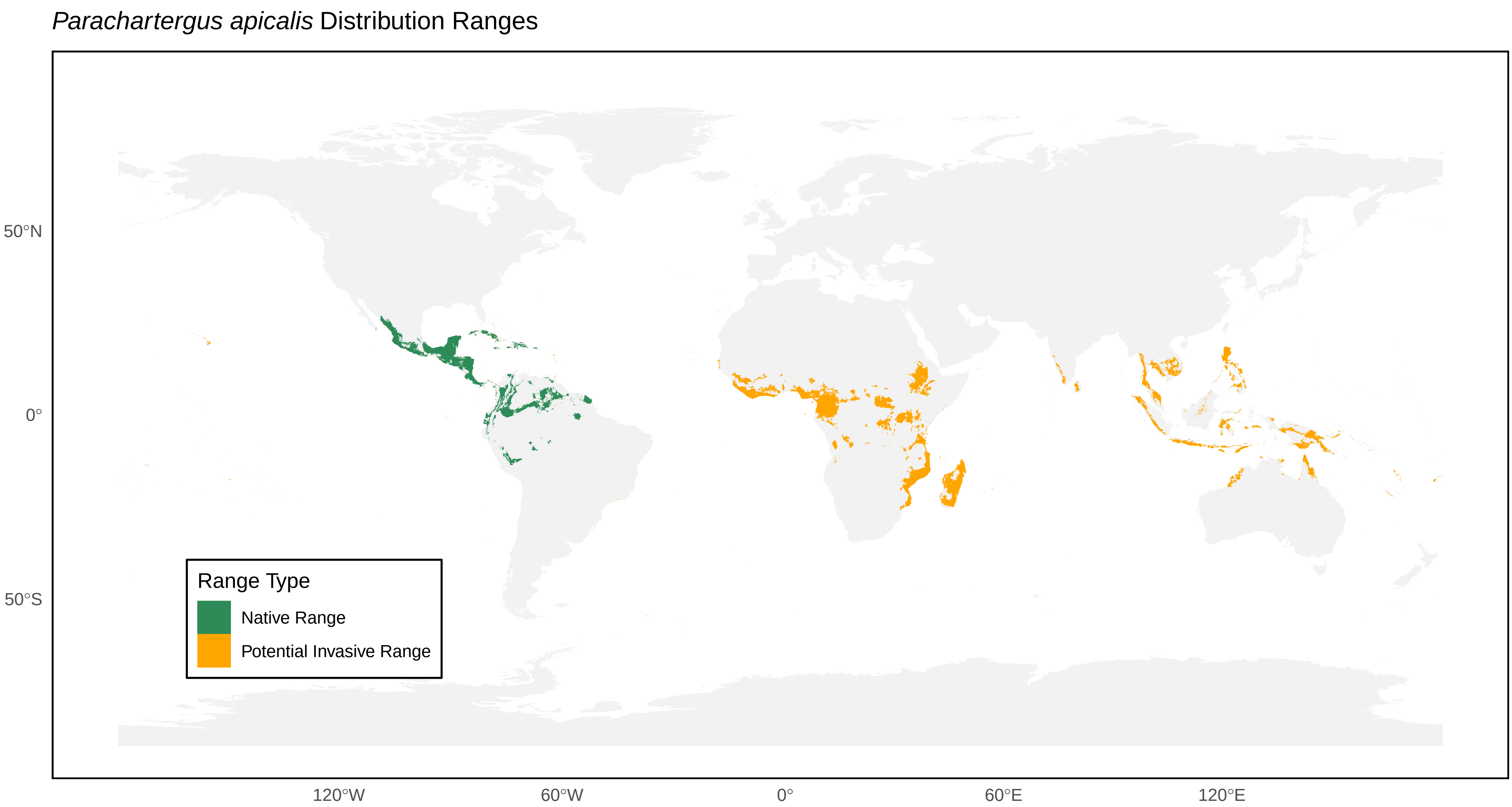
*Parachartergus apicalis* Distribution Ranges
50°N
0°
50°S
120°W
60°W
0°
60°E
120°E
Range Type
Native Range
Potential Invasive Range

*Parachartergus colobopterus* Distribution Ranges

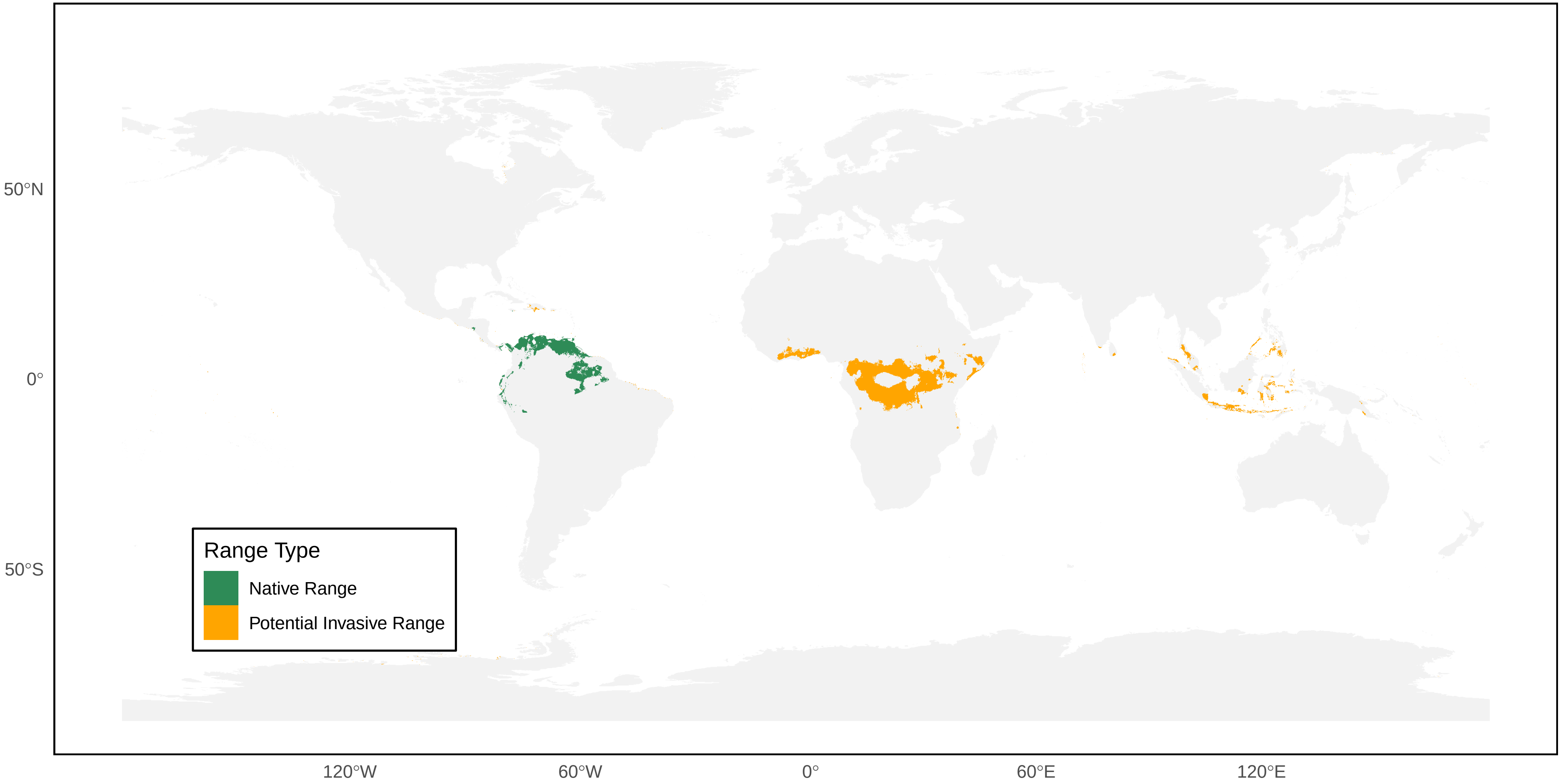

*Parachartergus compressus* Distribution Ranges

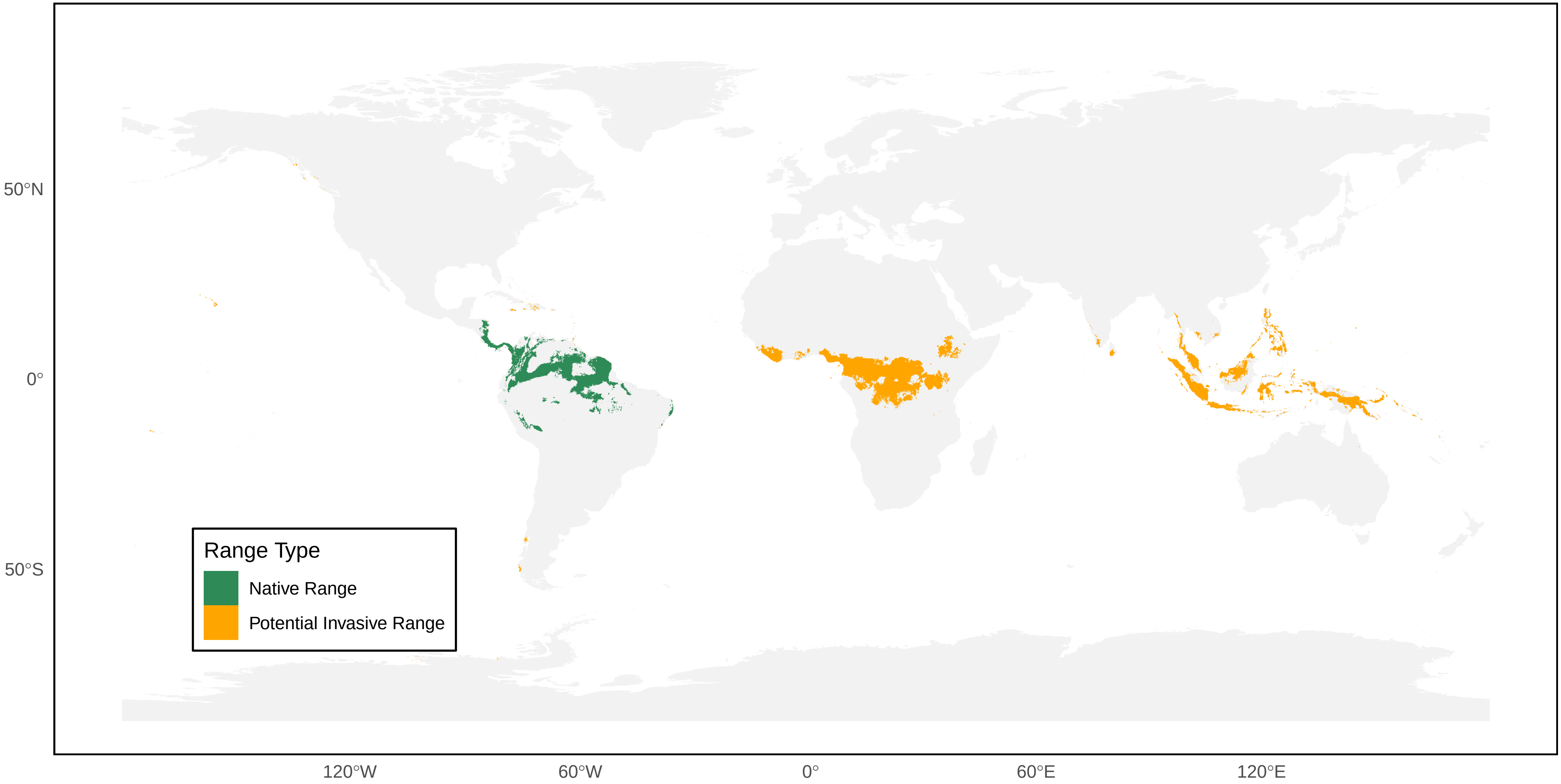

*Parachartergus fraternus* Distribution Ranges

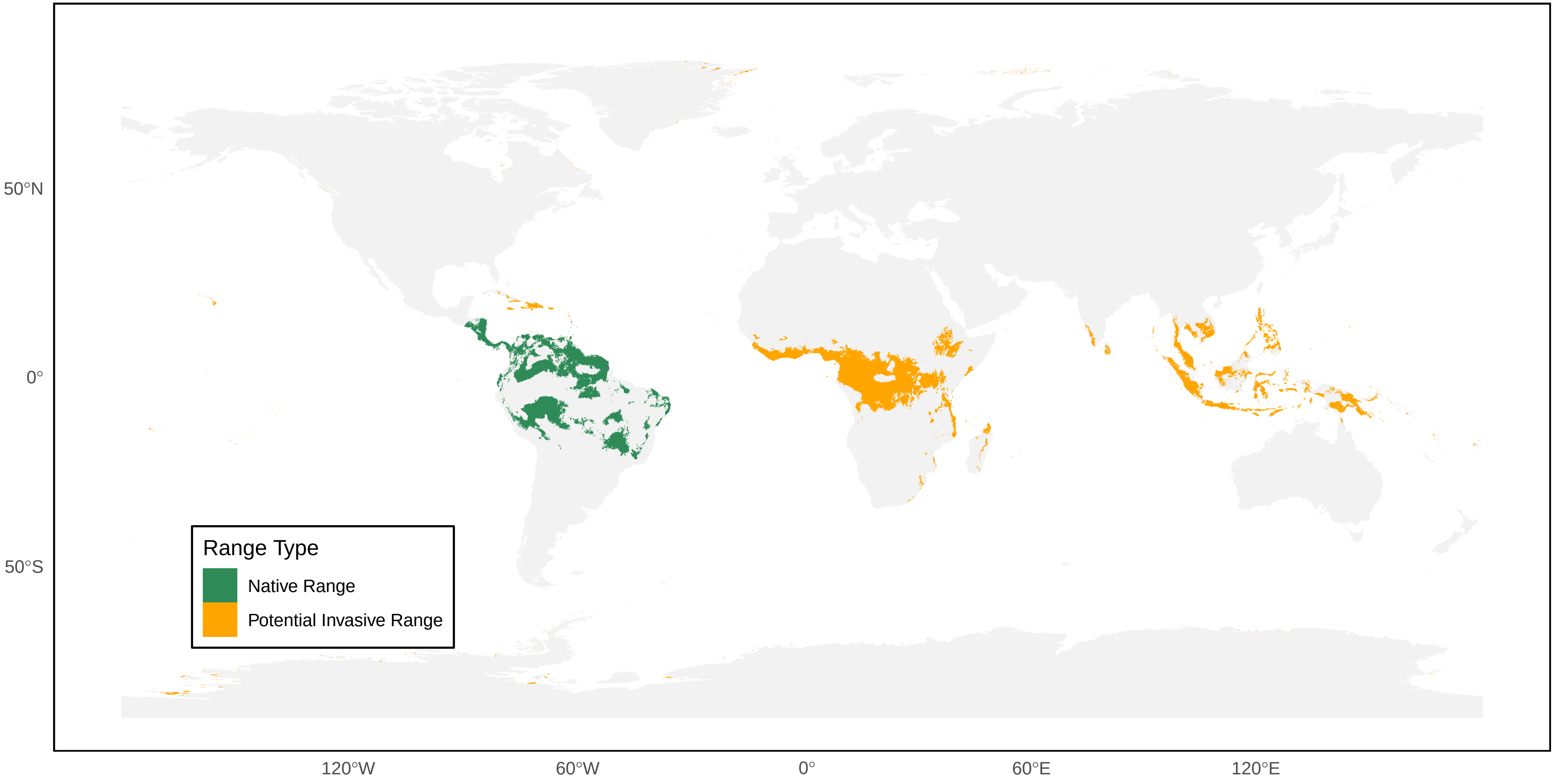

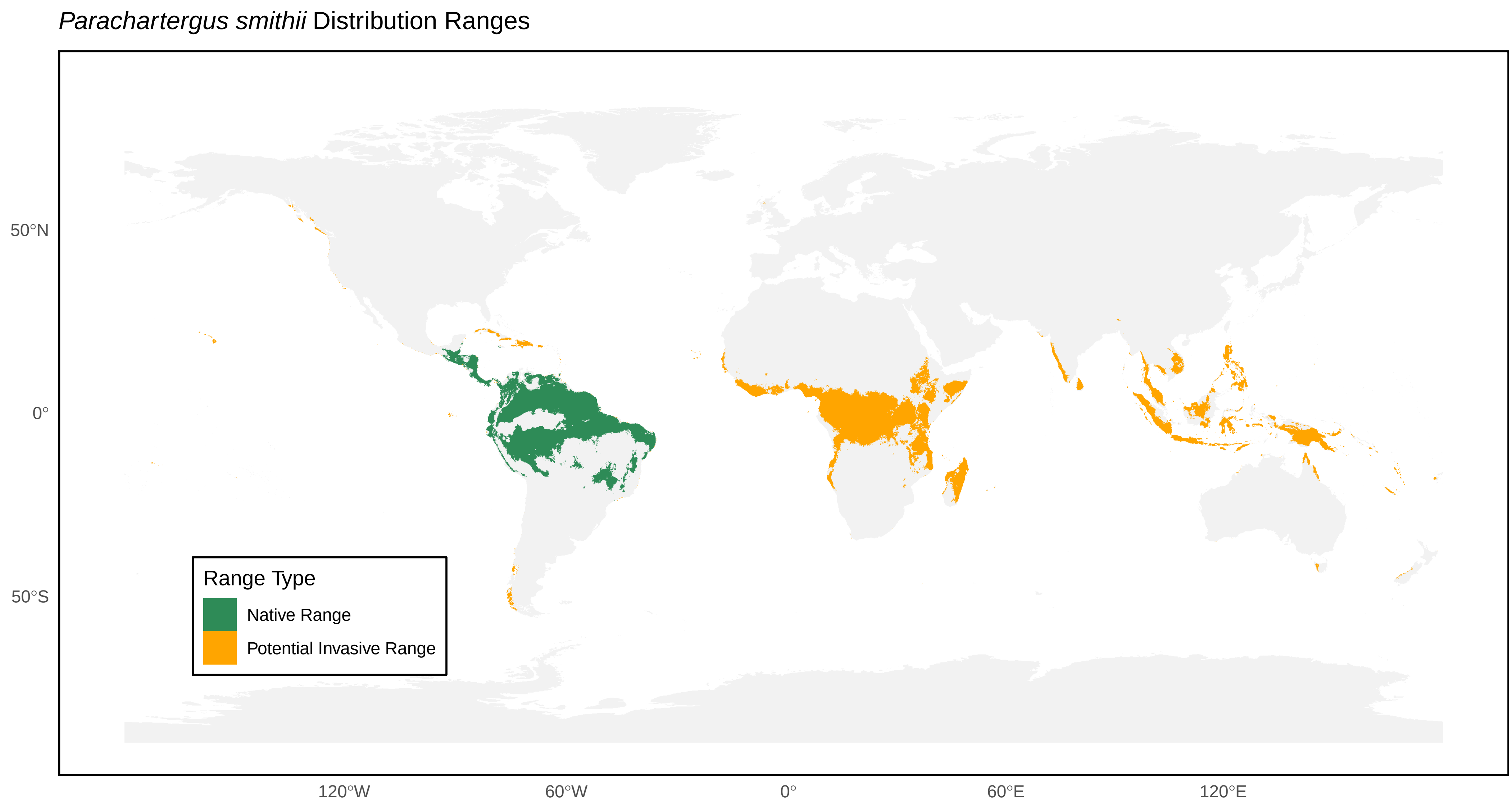
*Parachartergus smithii* Distribution Ranges
Range Type
Native Range
Potential Invasive Range
50°N
0°
50°S
120°W
60°W
0°
60°E
120°E

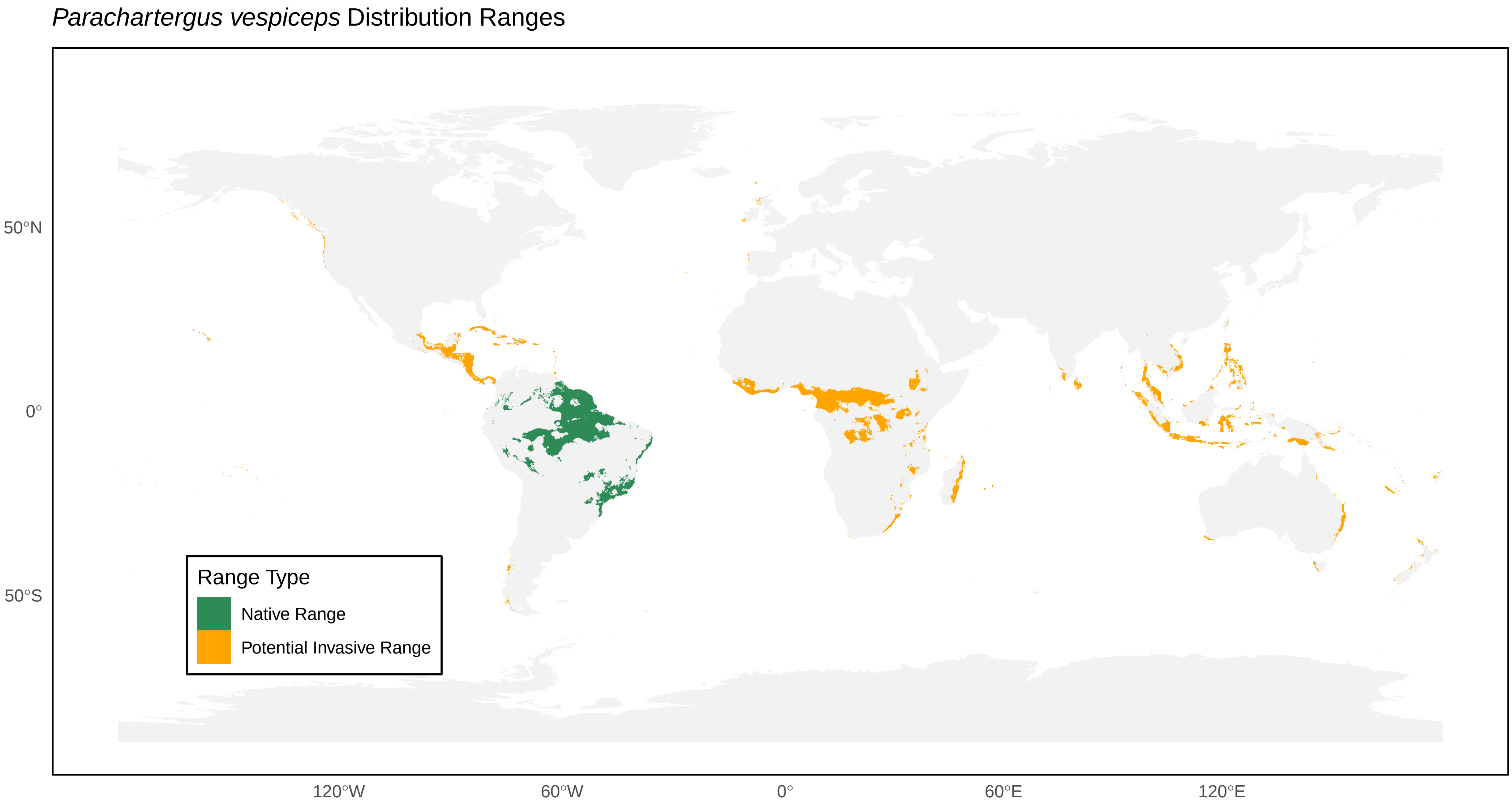

*Parachartergus vespiceps* Distribution Ranges
50°N
0°
50°S
120°W
60°W
0°
60°E
120°E
Range Type
Native Range
Potential Invasive Range

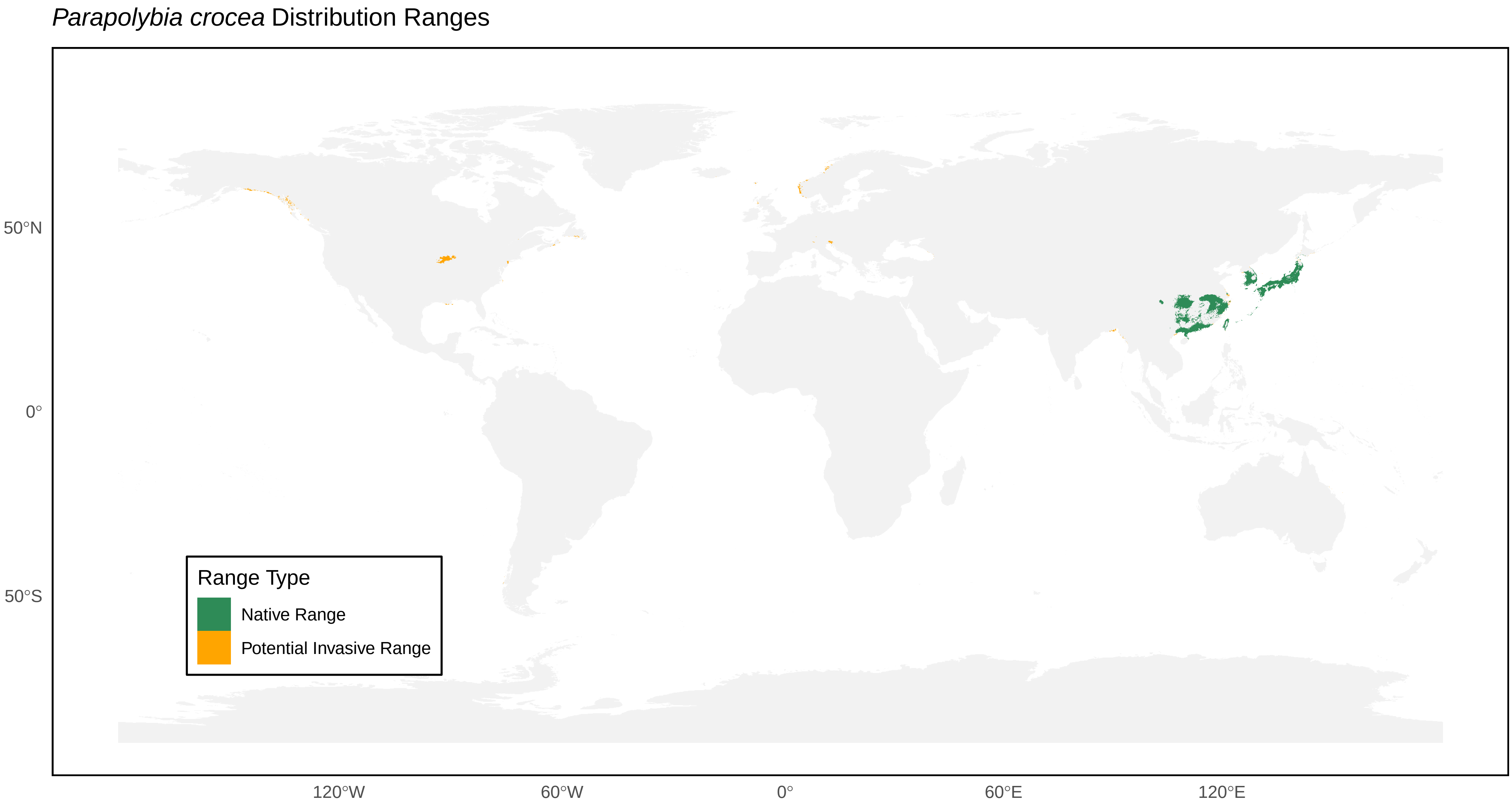

*Parapolybia crocea* Distribution Ranges
50°N
0°
50°S
120°W
60°W
0°
60°E
120°E
Range Type
Native Range
Potential Invasive Range

*Parapolybia indica* Distribution Ranges

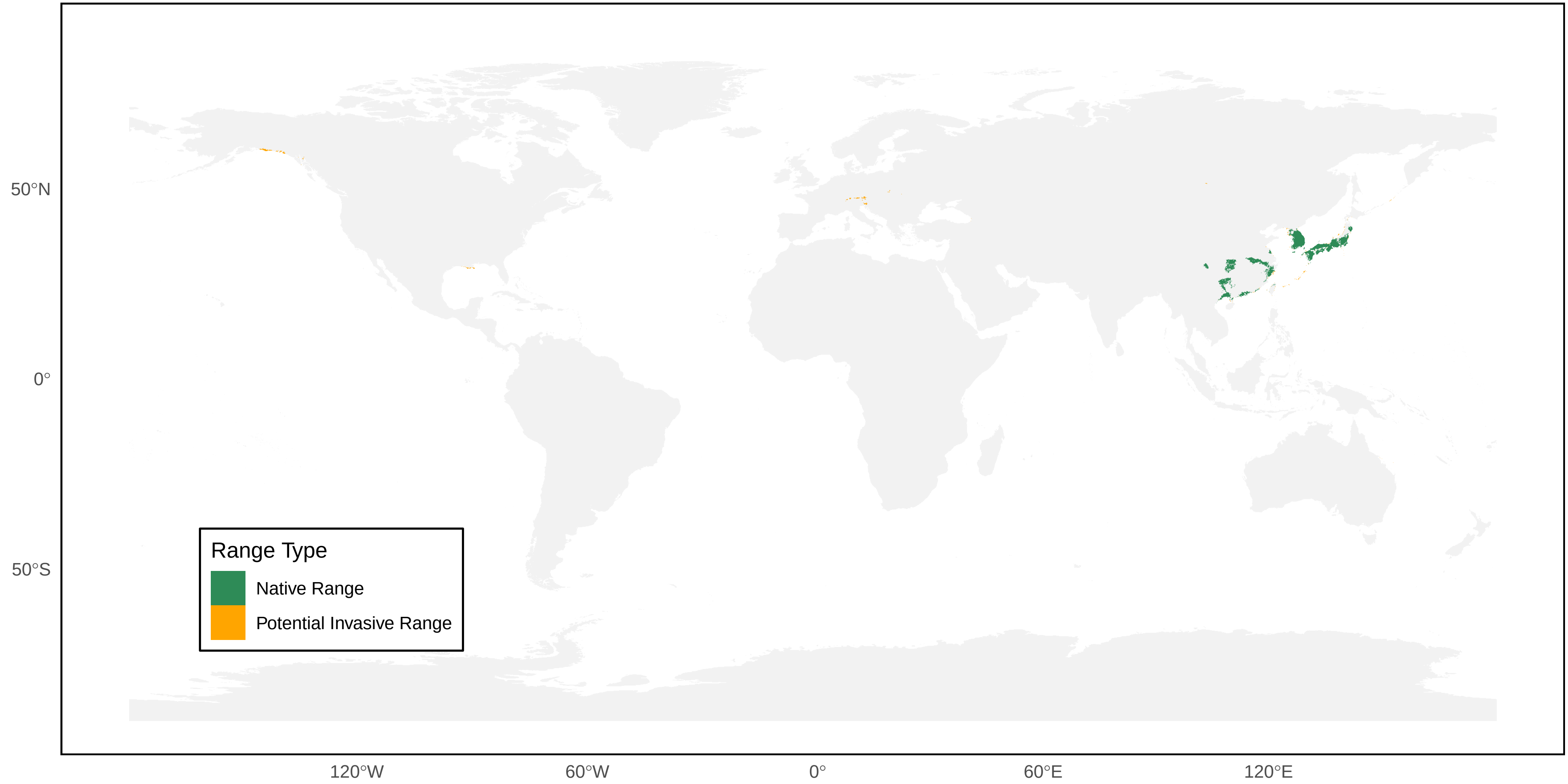

*Parapolybia nodosa* Distribution Ranges

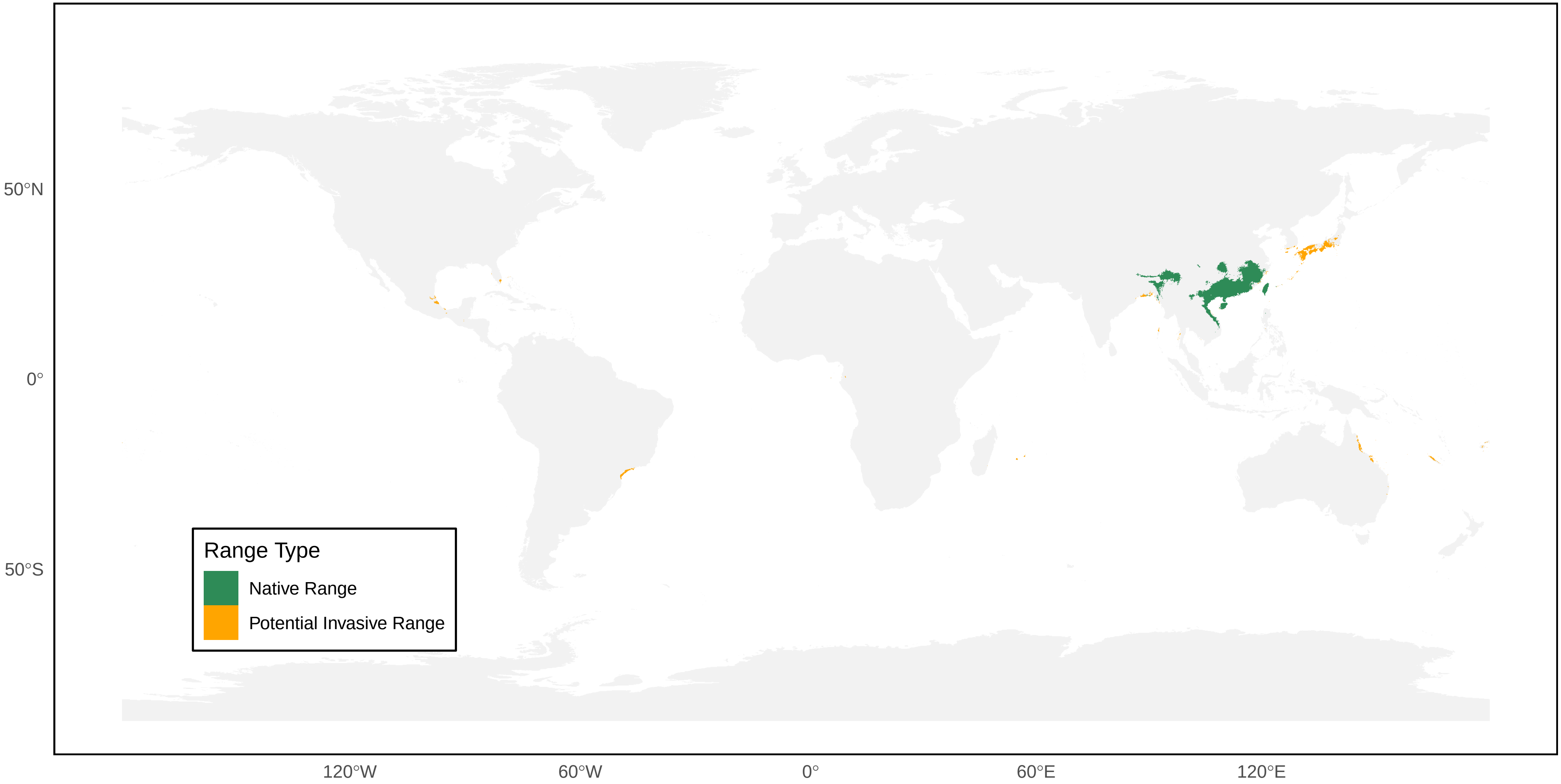

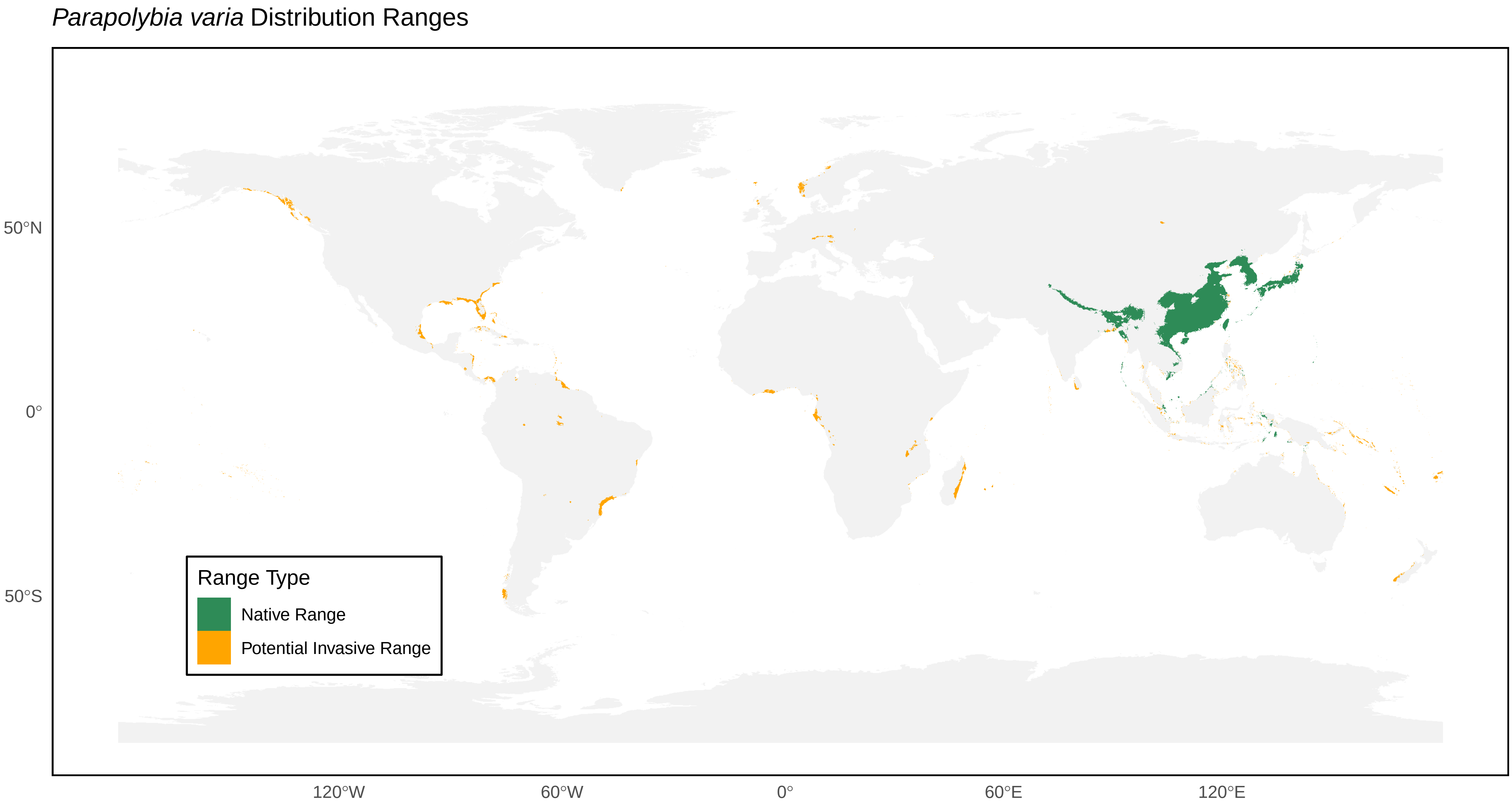

Parapolybia varia Distribution Ranges
50°N
0°
50°S
120°W
60°W
0°
60°E
120°E
Range Type
Native Range
Potential Invasive Range

*Parischnogaster mellyi* Distribution Ranges

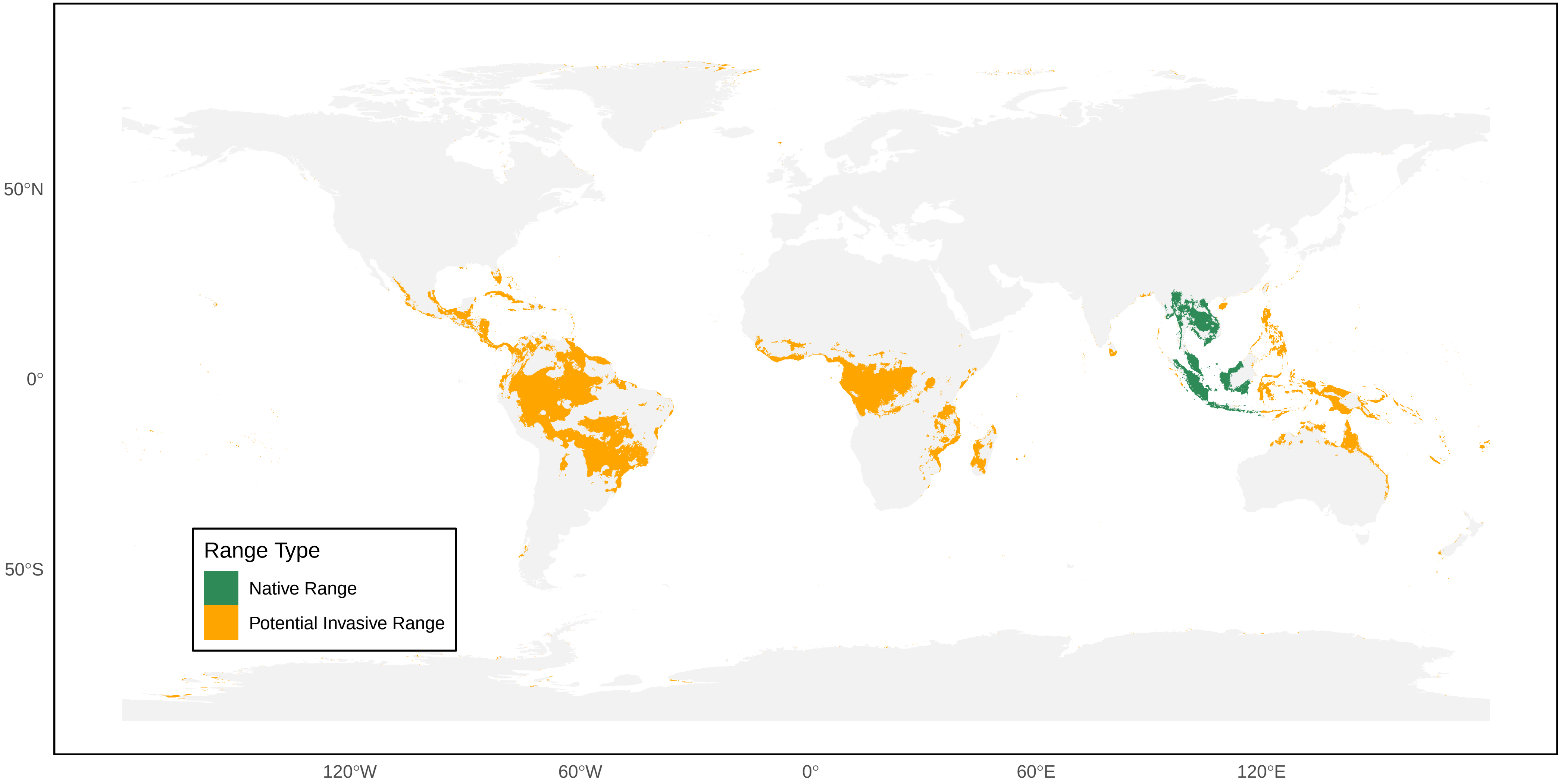

*Polistes actaeon* Distribution Ranges

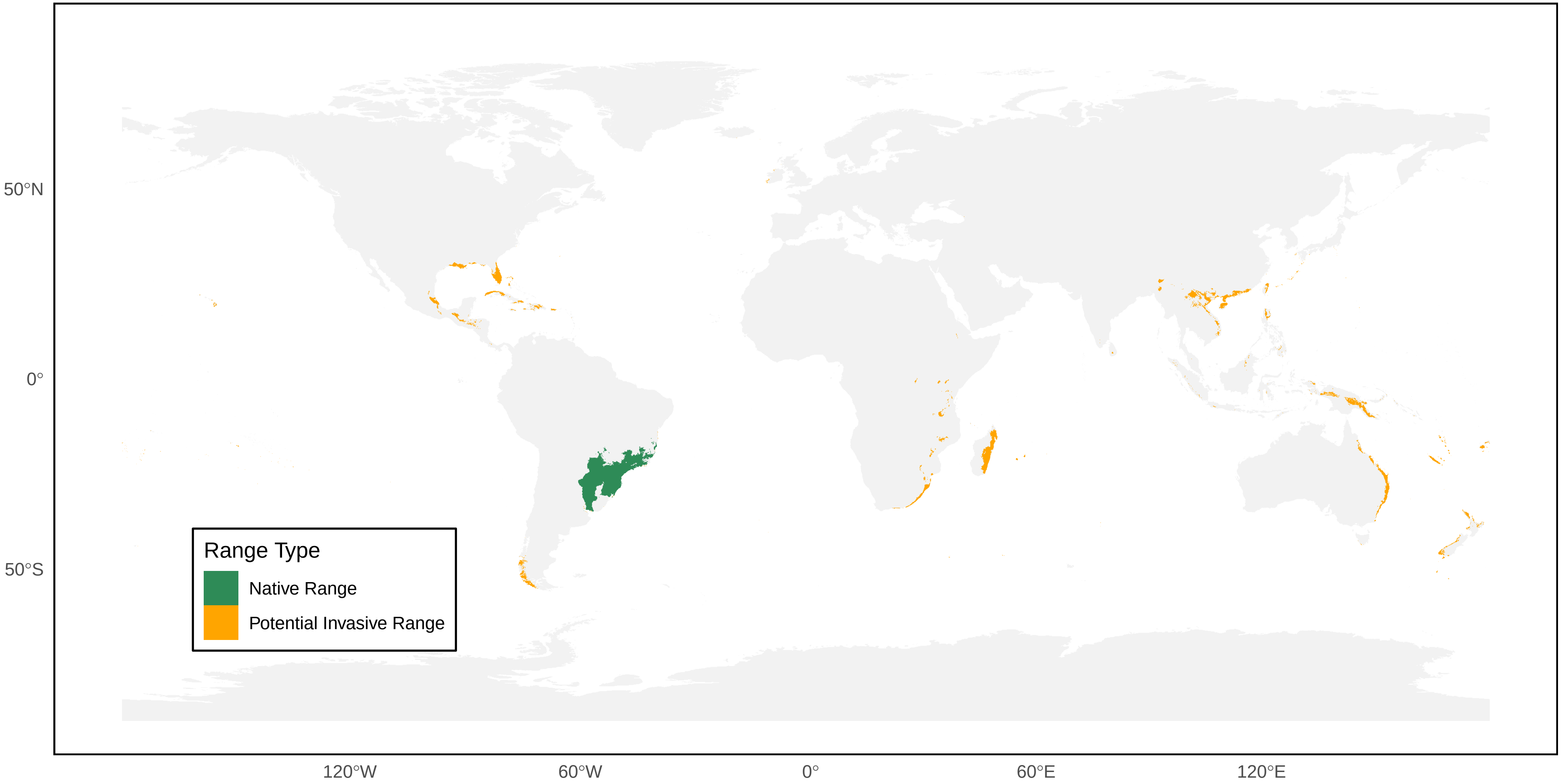

*Polistes africanus* Distribution Ranges

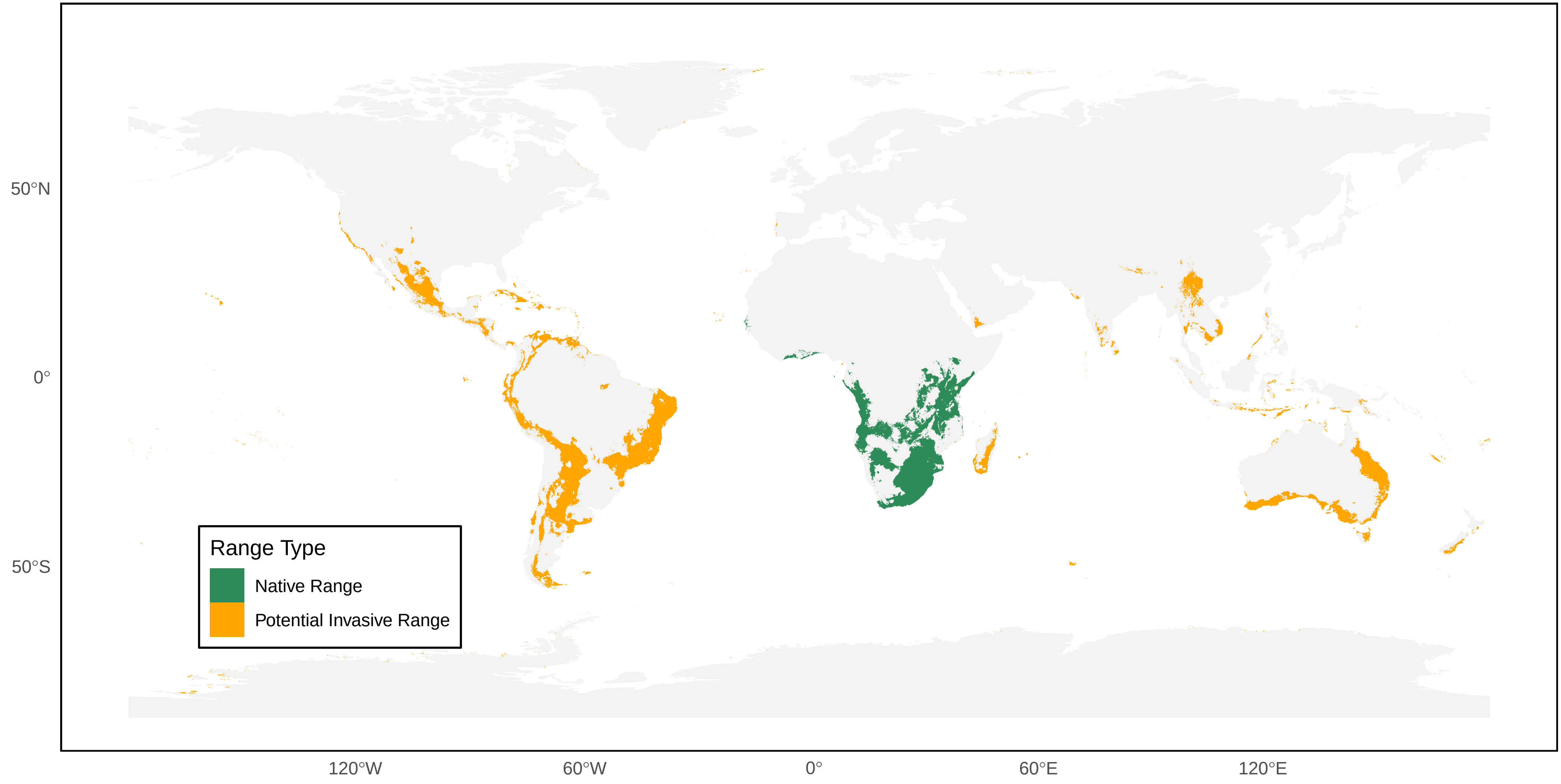

*Polistes annularis* Distribution Ranges

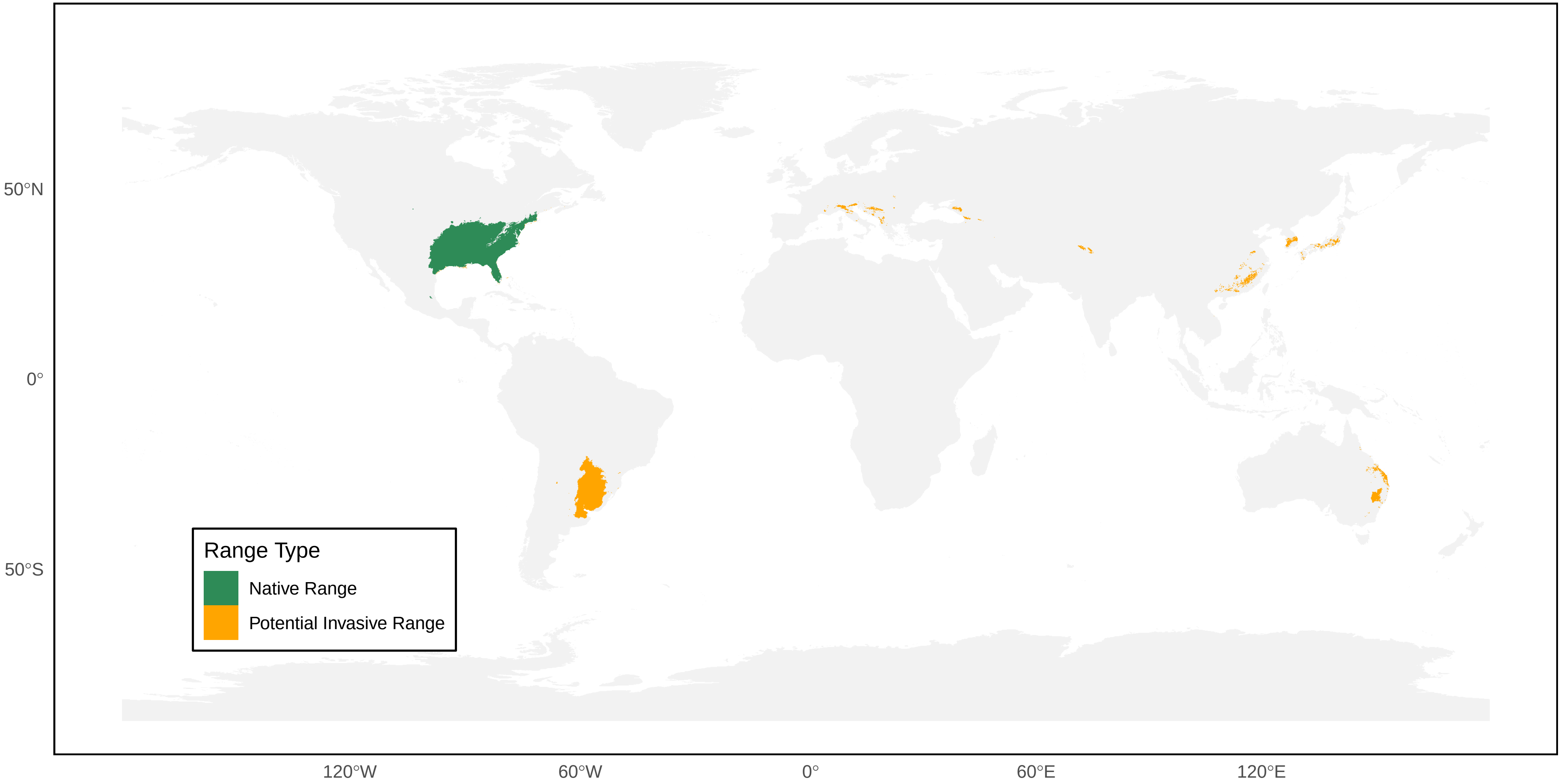

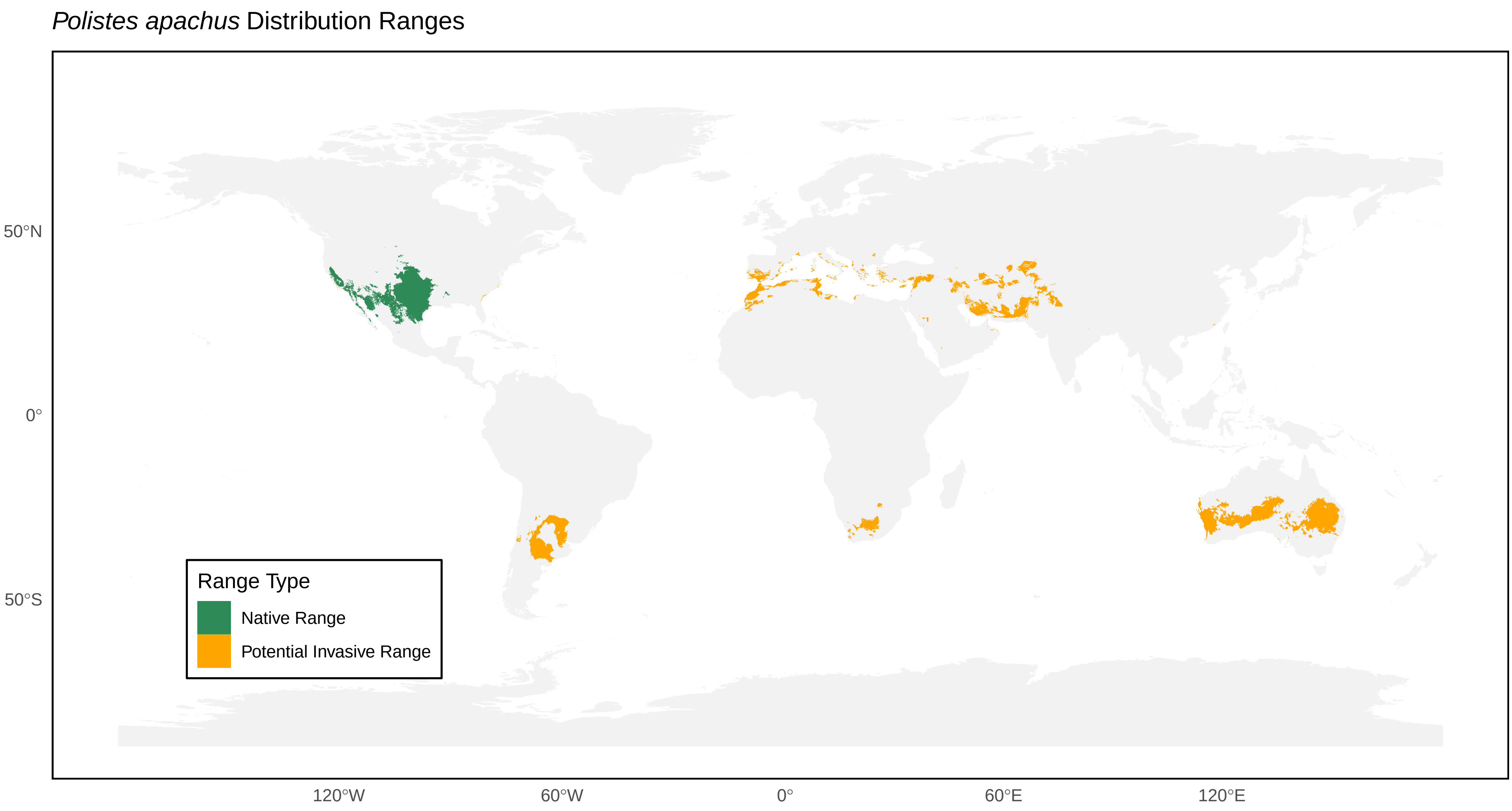
*Polistes apachus* Distribution Ranges
50°N
0°
50°S
120°W
60°W
0°
60°E
120°E
Range Type
Native Range
Potential Invasive Range

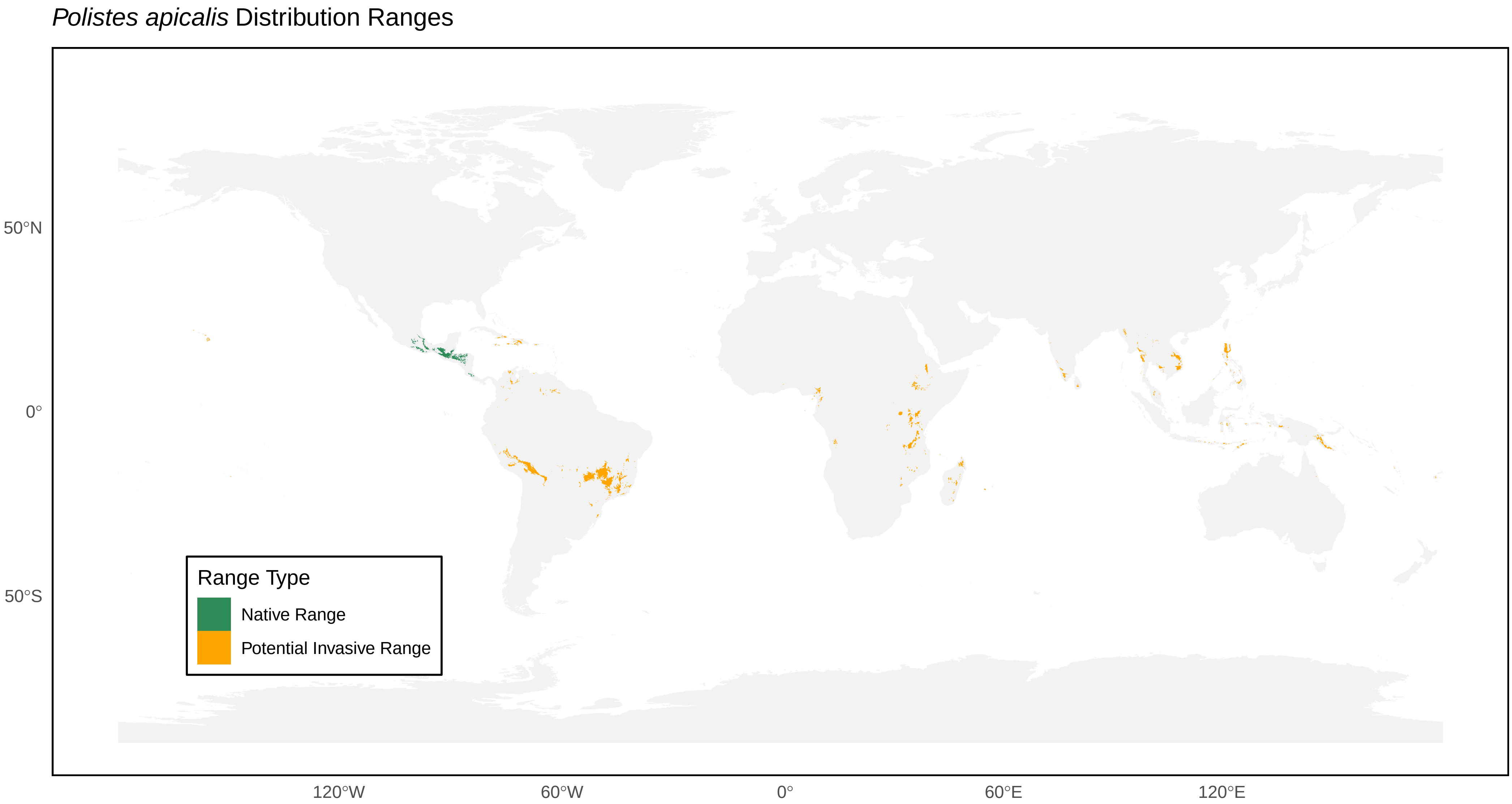
*Polistes apicalis* Distribution Ranges
50°N
0°
50°S
120°W
60°W
0°
60°E
120°E
Range Type
Native Range
Potential Invasive Range

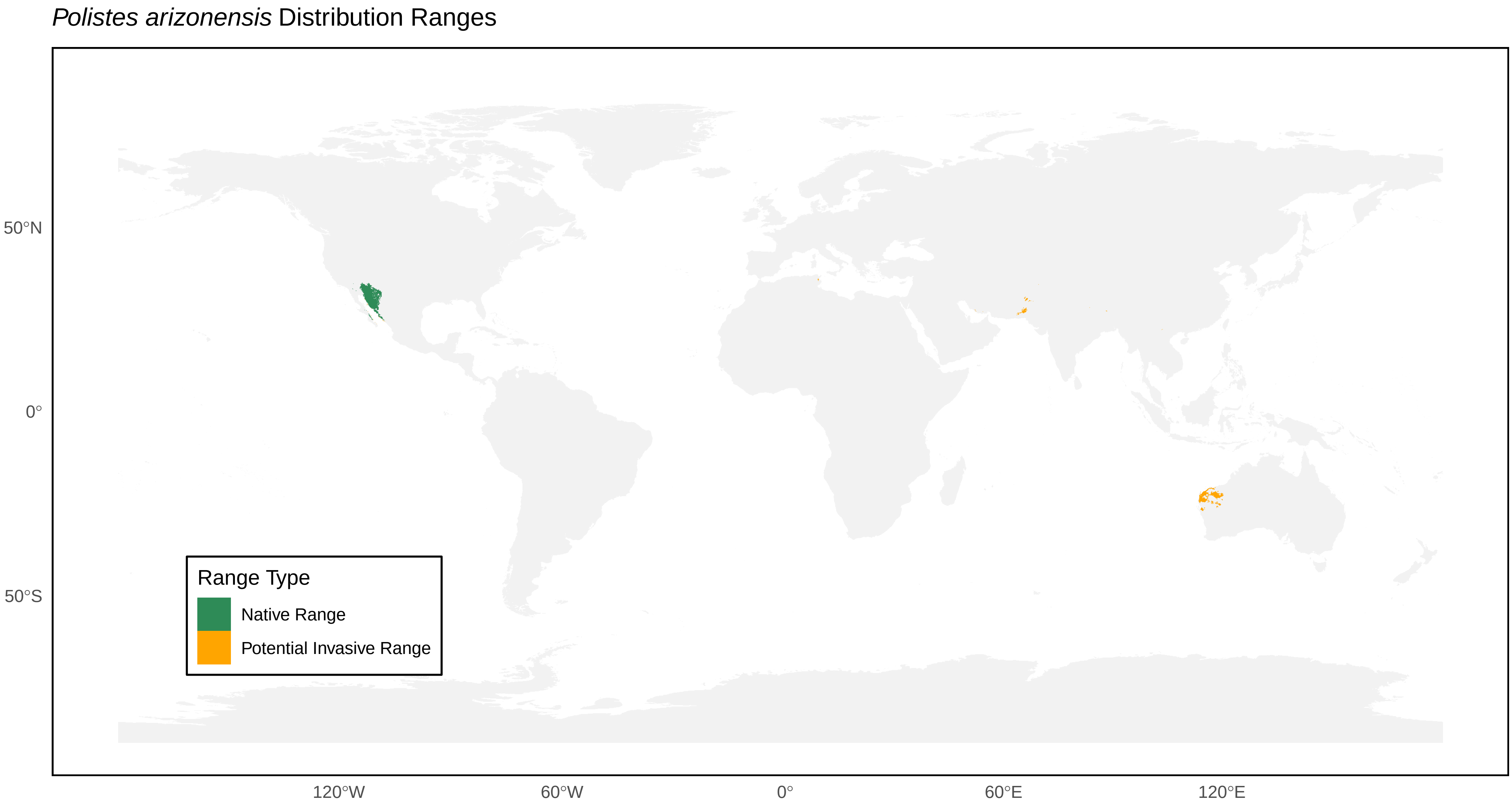

*Polistes arizonensis* Distribution Ranges
50°N
0°
50°S
120°W
60°W
0°
60°E
120°E
Range Type
Native Range
Potential Invasive Range

*Polistes associus* Distribution Ranges

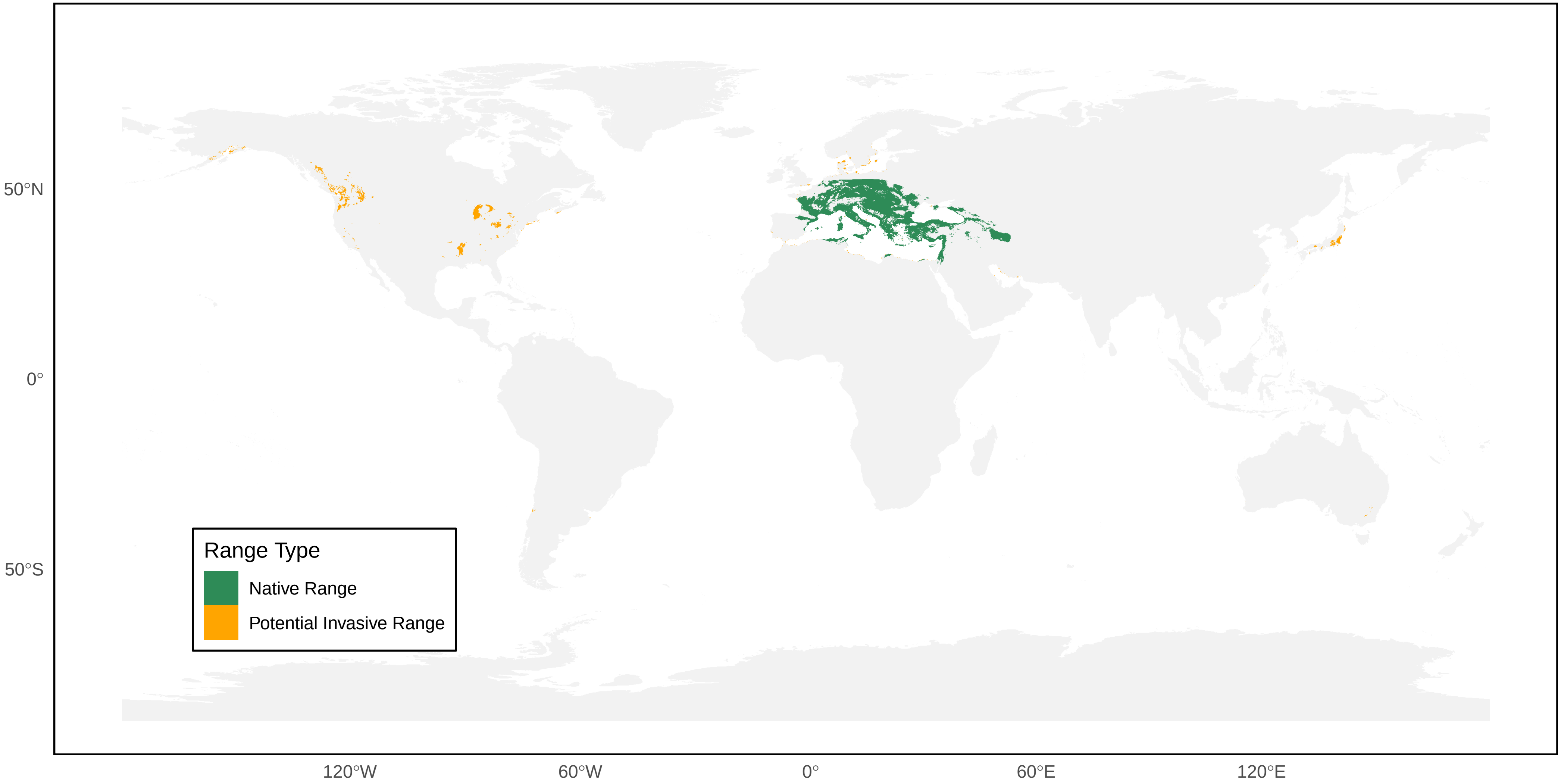

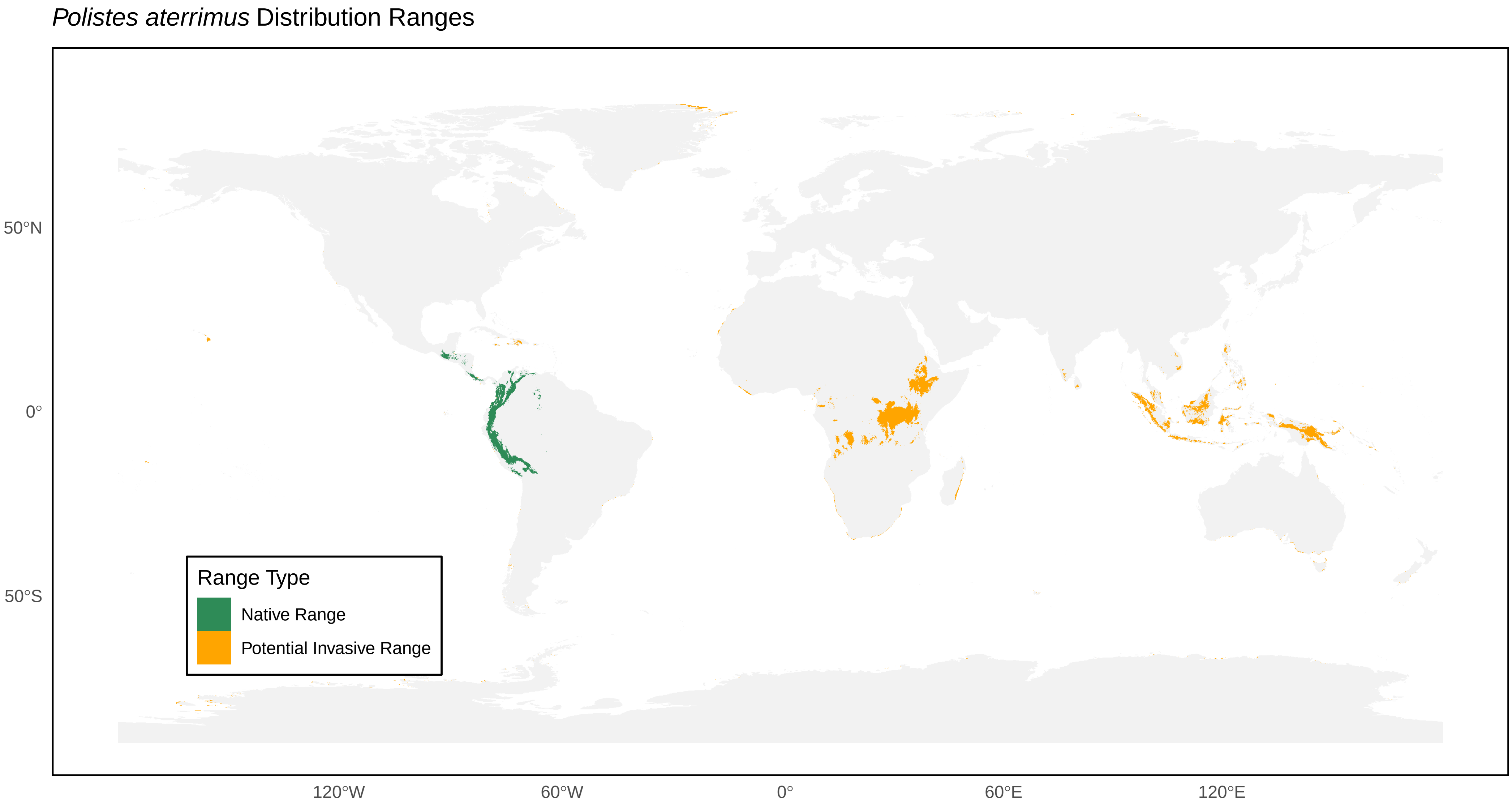

*Polistes aterrimus* Distribution Ranges
50°N
0°
50°S
120°W
60°W
0°
60°E
120°E
Range Type
Native Range
Potential Invasive Range

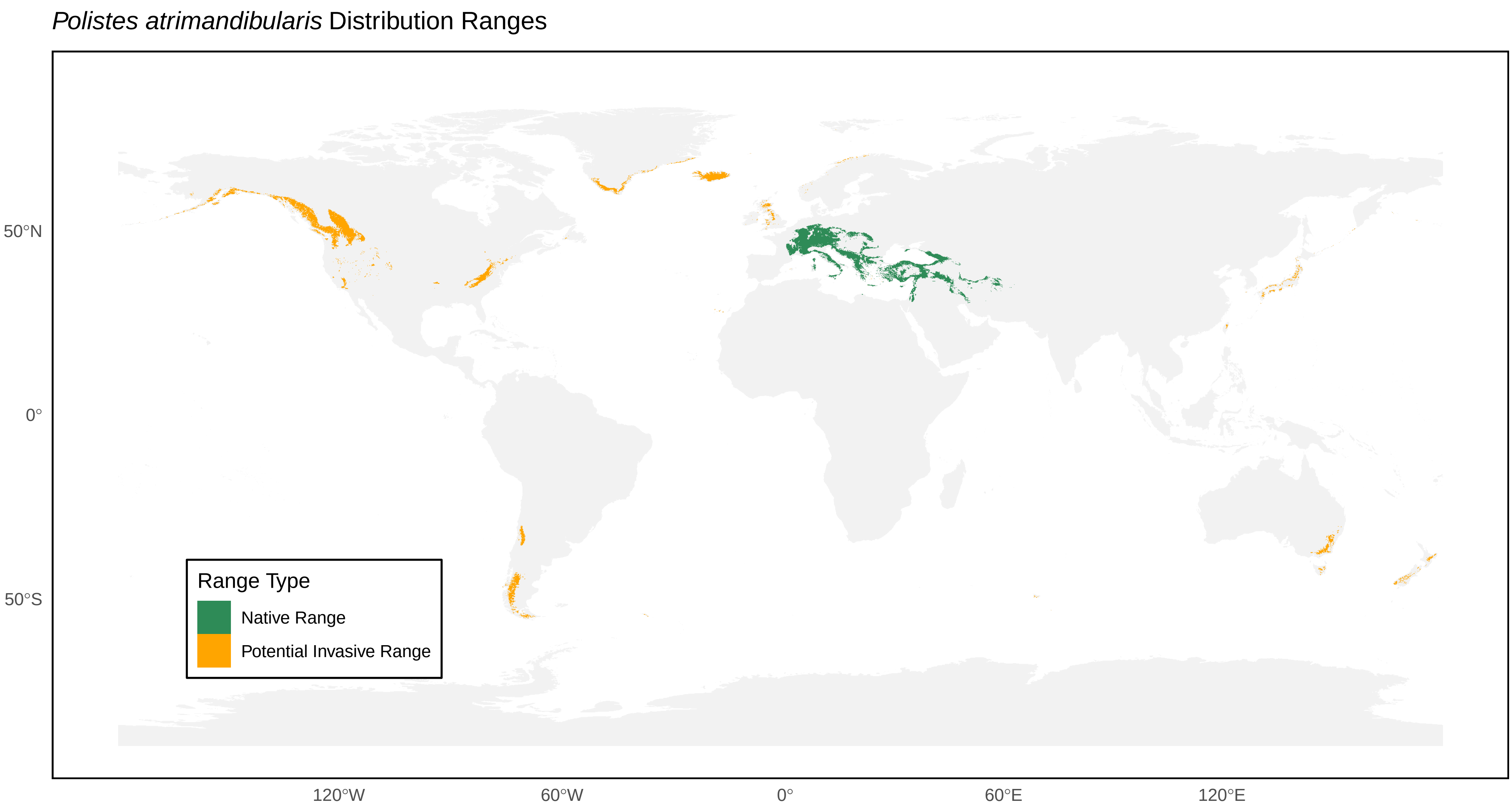

*Polistes atrimandibularis* Distribution Ranges
50°N
0°
50°S
120°W
60°W
0°
60°E
120°E
Range Type
Native Range
Potential Invasive Range

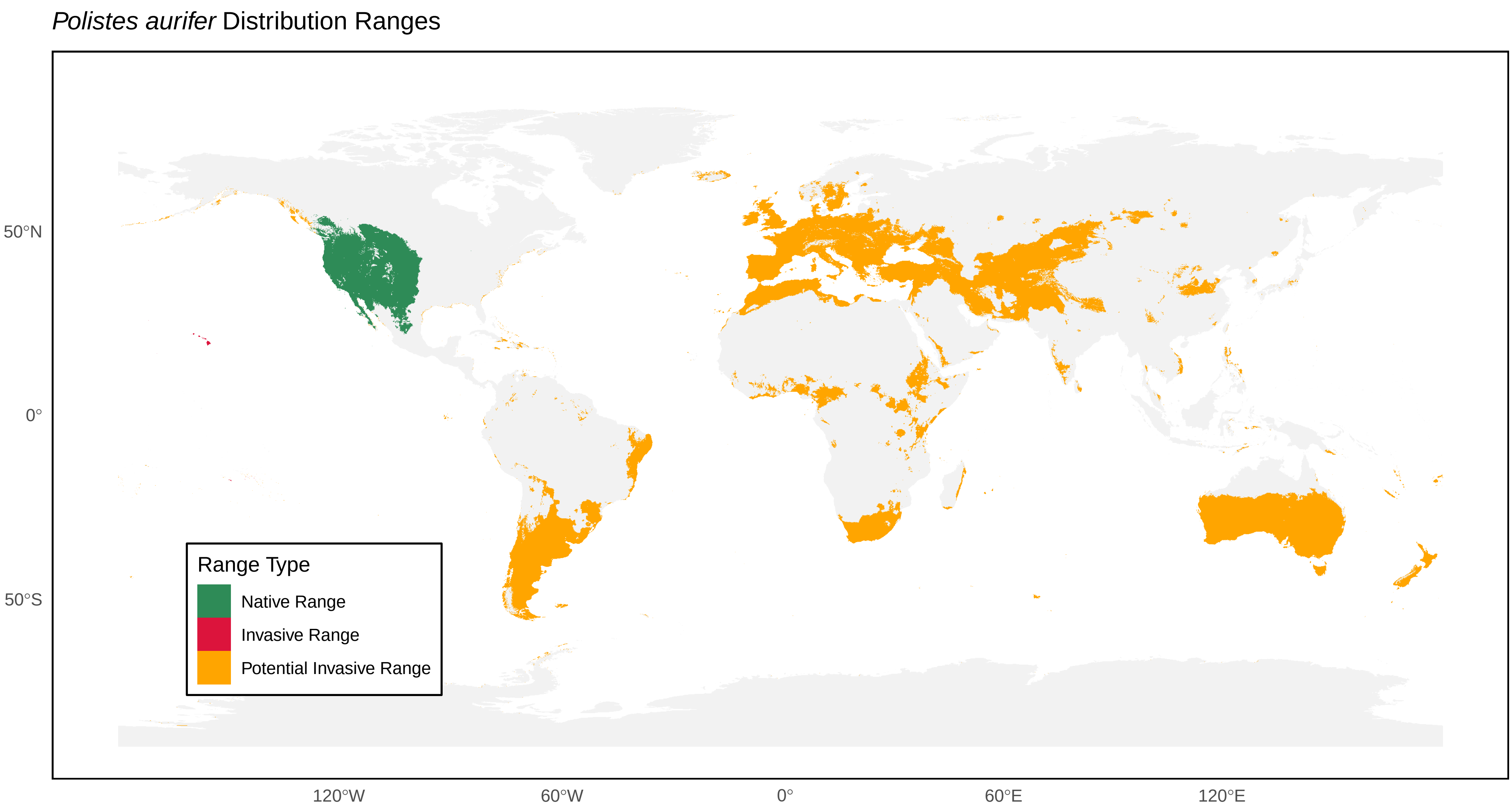
*Polistes aurifer* Distribution Ranges
50°N
0°
50°S
120°W
60°W
0°
60°E
120°E
Range Type
Native Range
Invasive Range
Potential Invasive Range

*Polistes austroccidentalis* Distribution Ranges

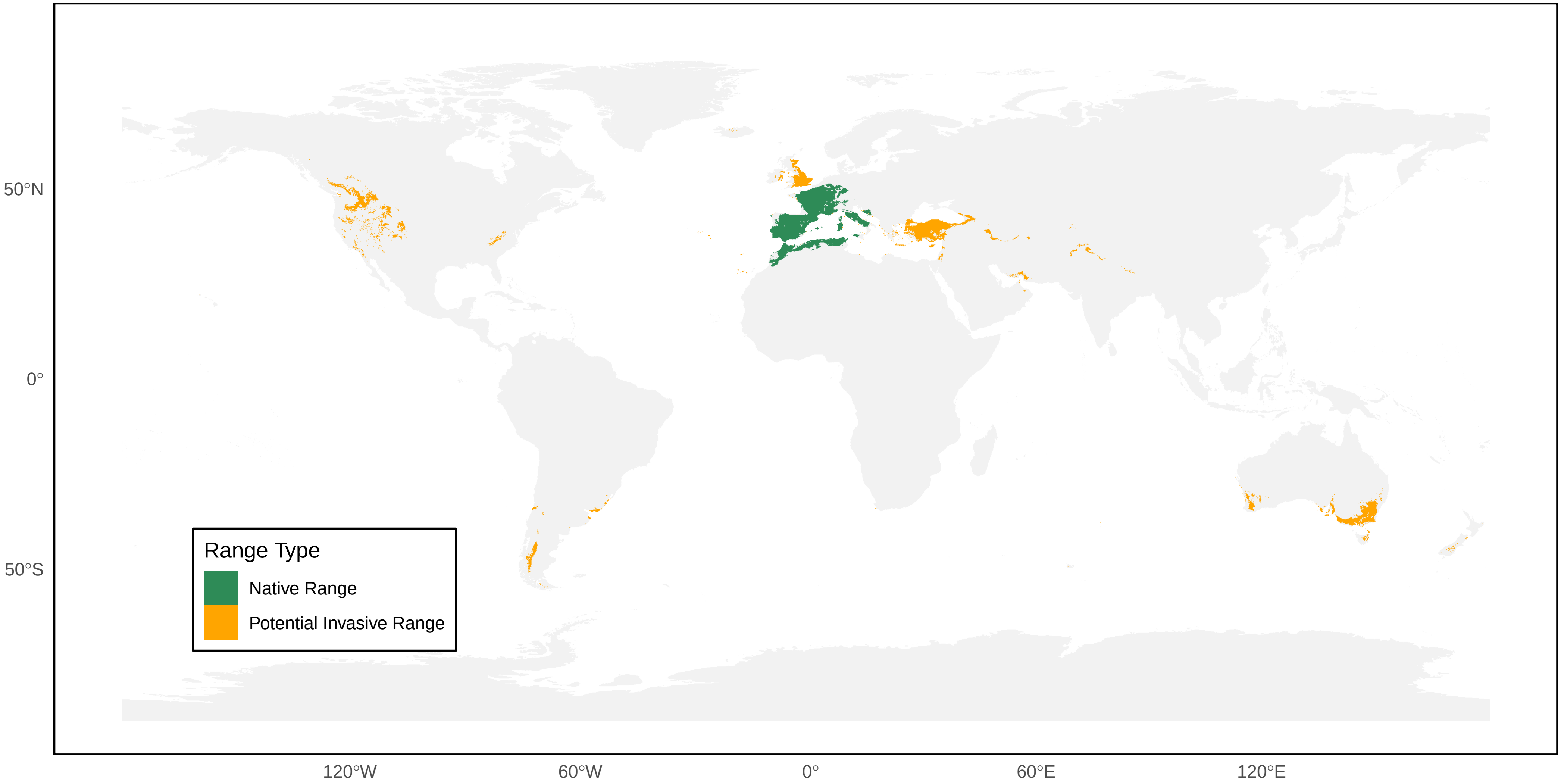

*Polistes badius* Distribution Ranges

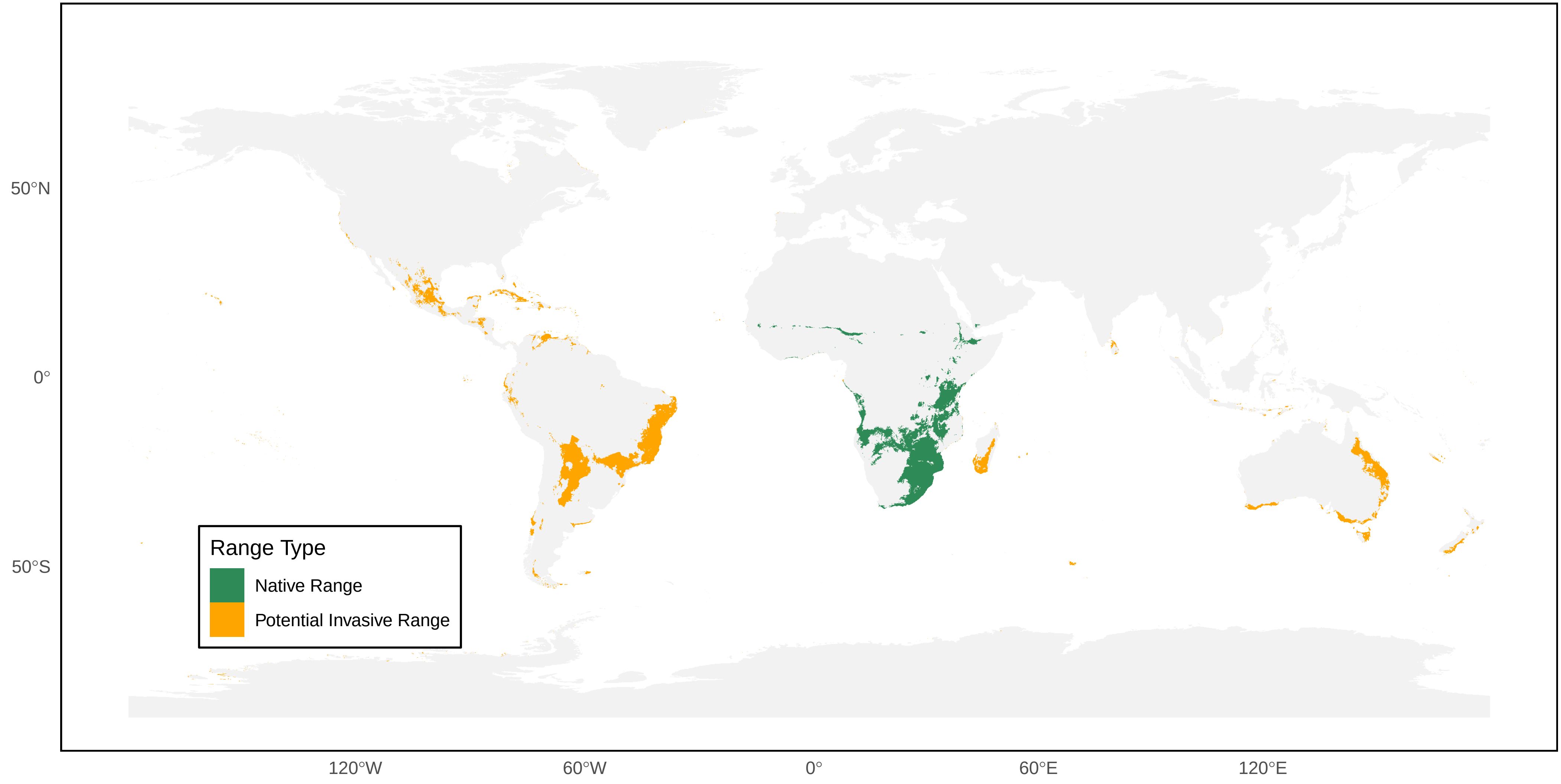

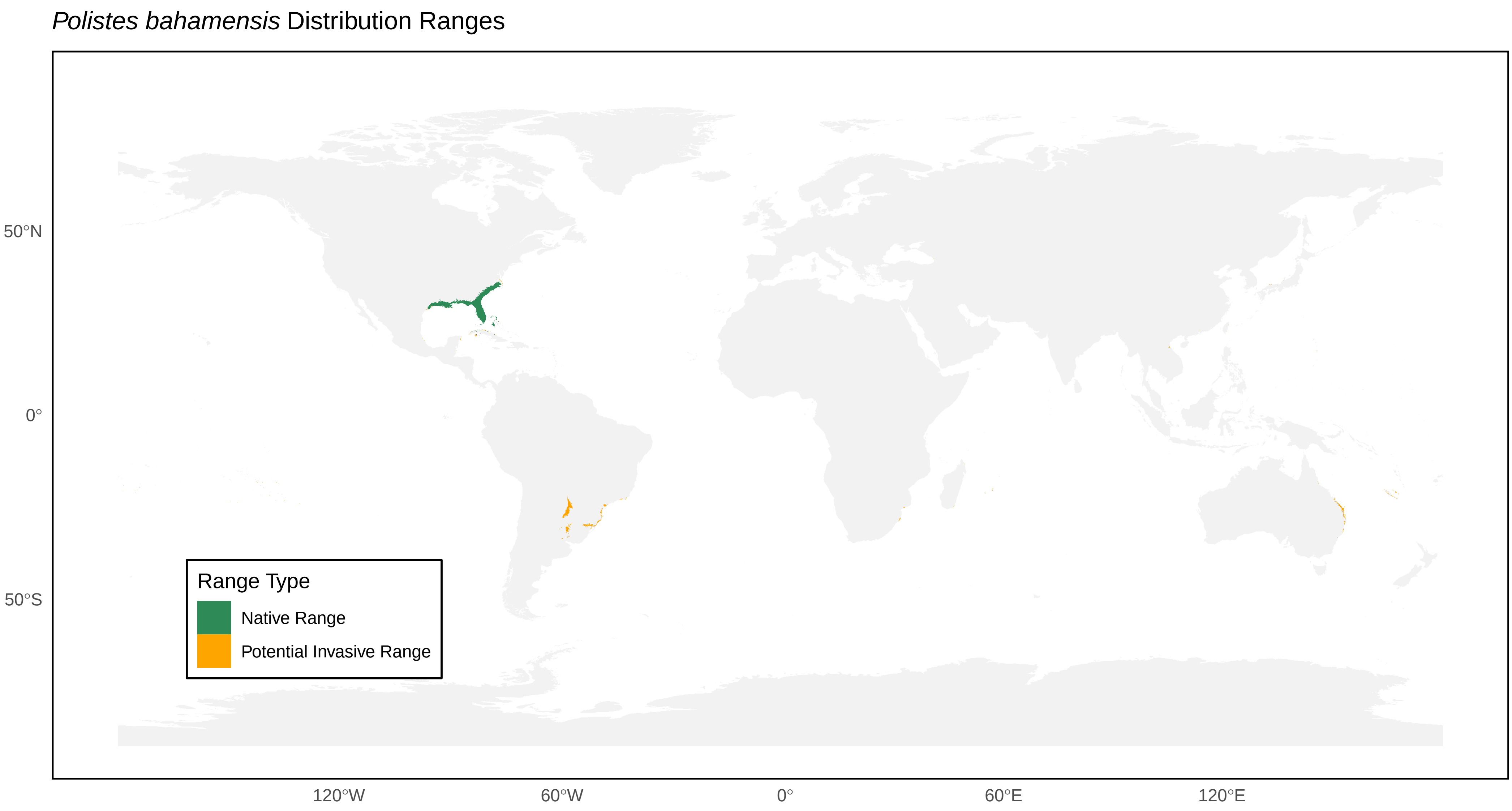
*Polistes bahamensis* Distribution Ranges
50°N
0°
50°S
120°W
60°W
0°
60°E
120°E
Range Type
Native Range
Potential Invasive Range

*Polistes bellicosus* Distribution Ranges

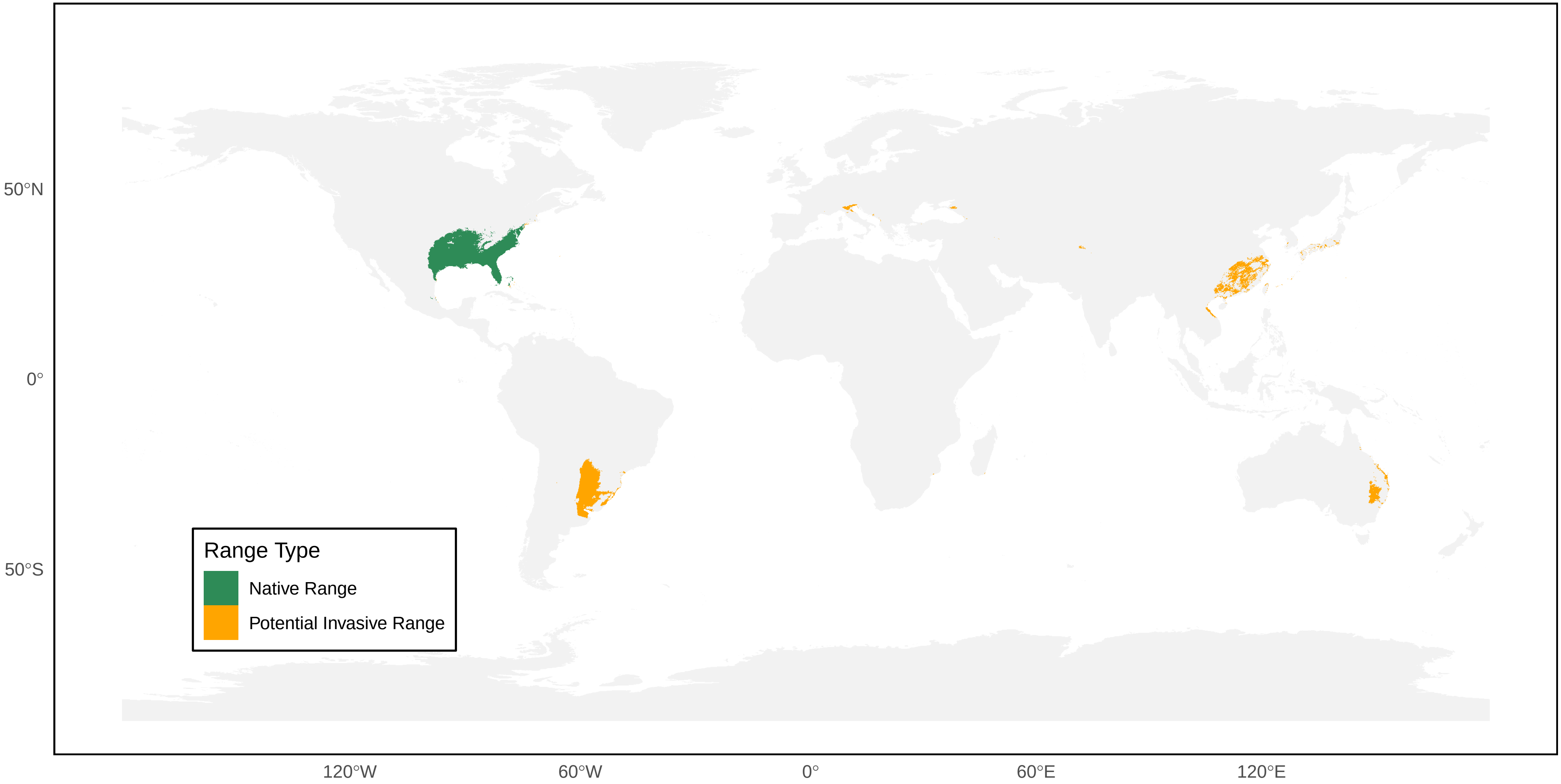

*Polistes bequaertellus* Distribution Ranges

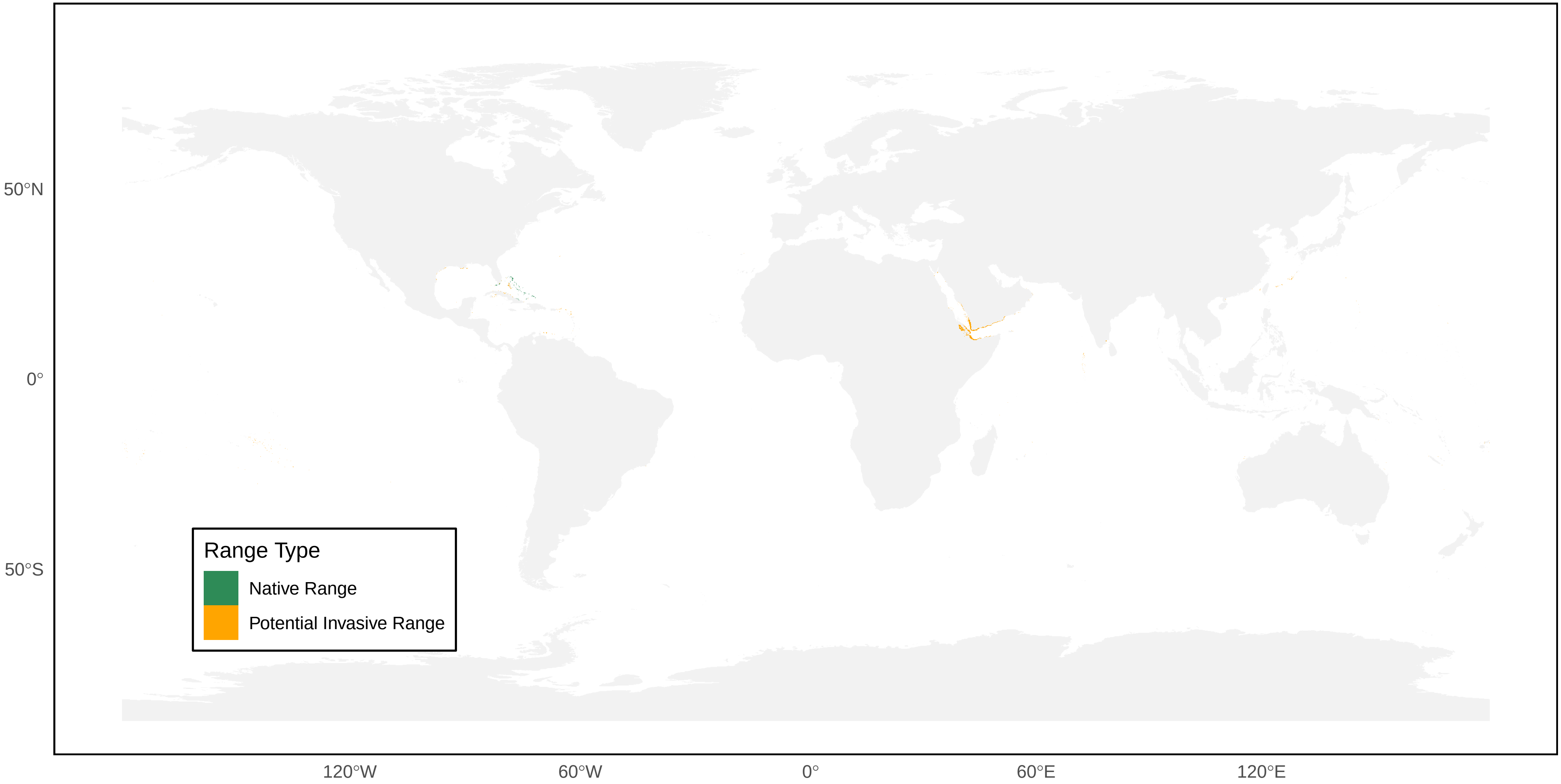

*Polistes bicolor* Distribution Ranges

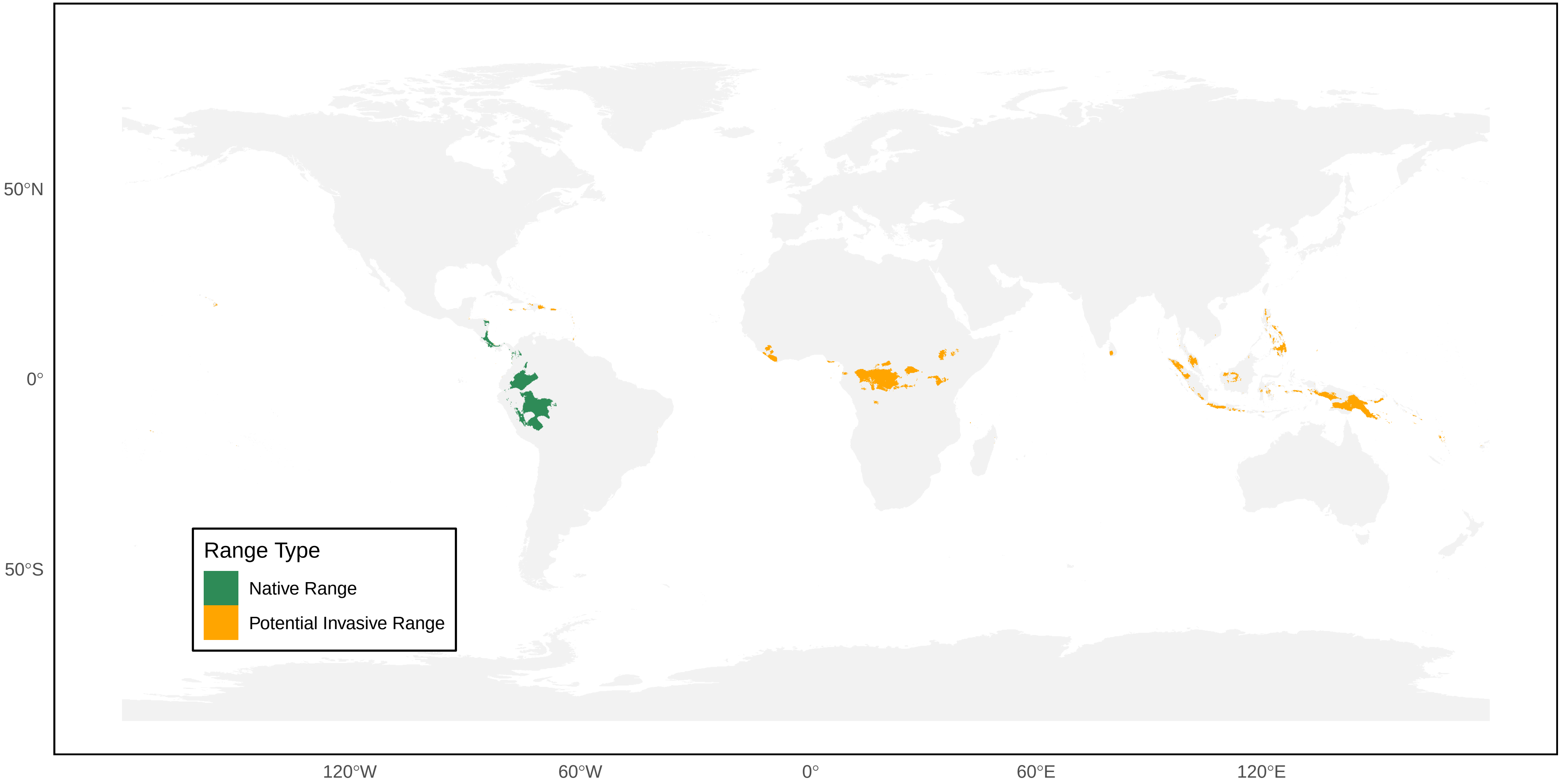

*Polistes biglumis* Distribution Ranges

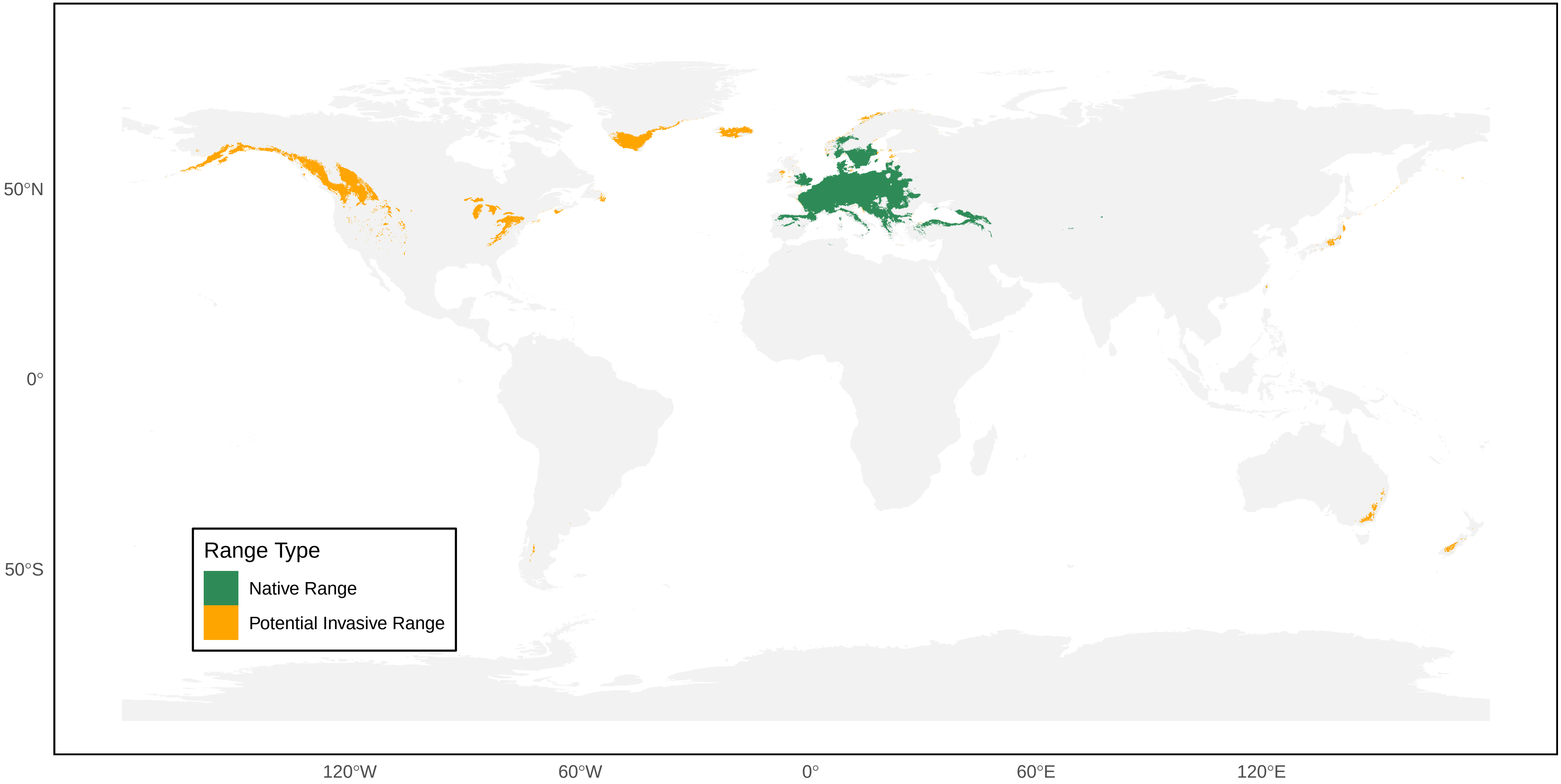

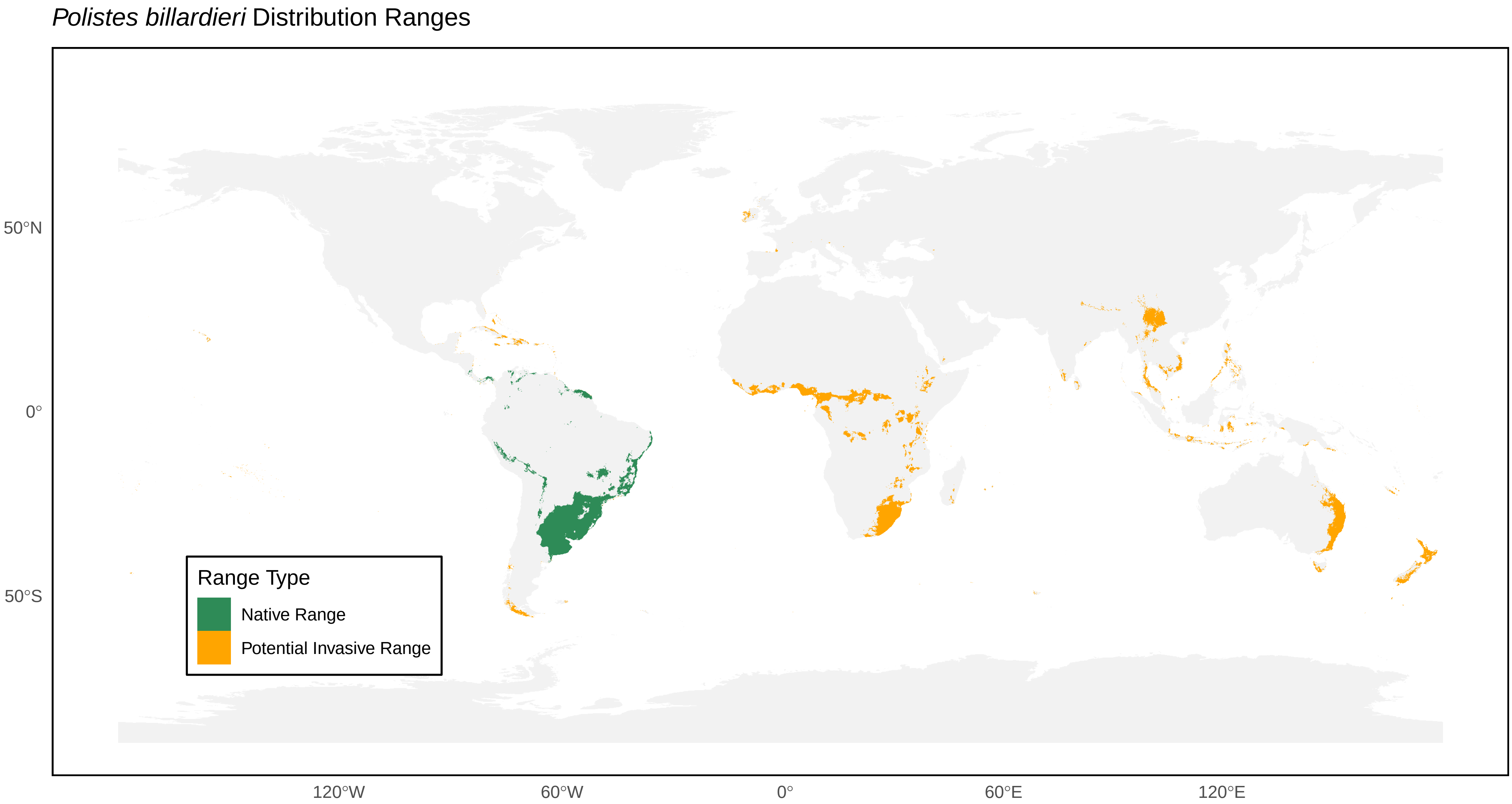

*Polistes billardieri* Distribution Ranges
50°N
0°
50°S
120°W
60°W
0°
60°E
120°E
Range Type
Native Range
Potential Invasive Range

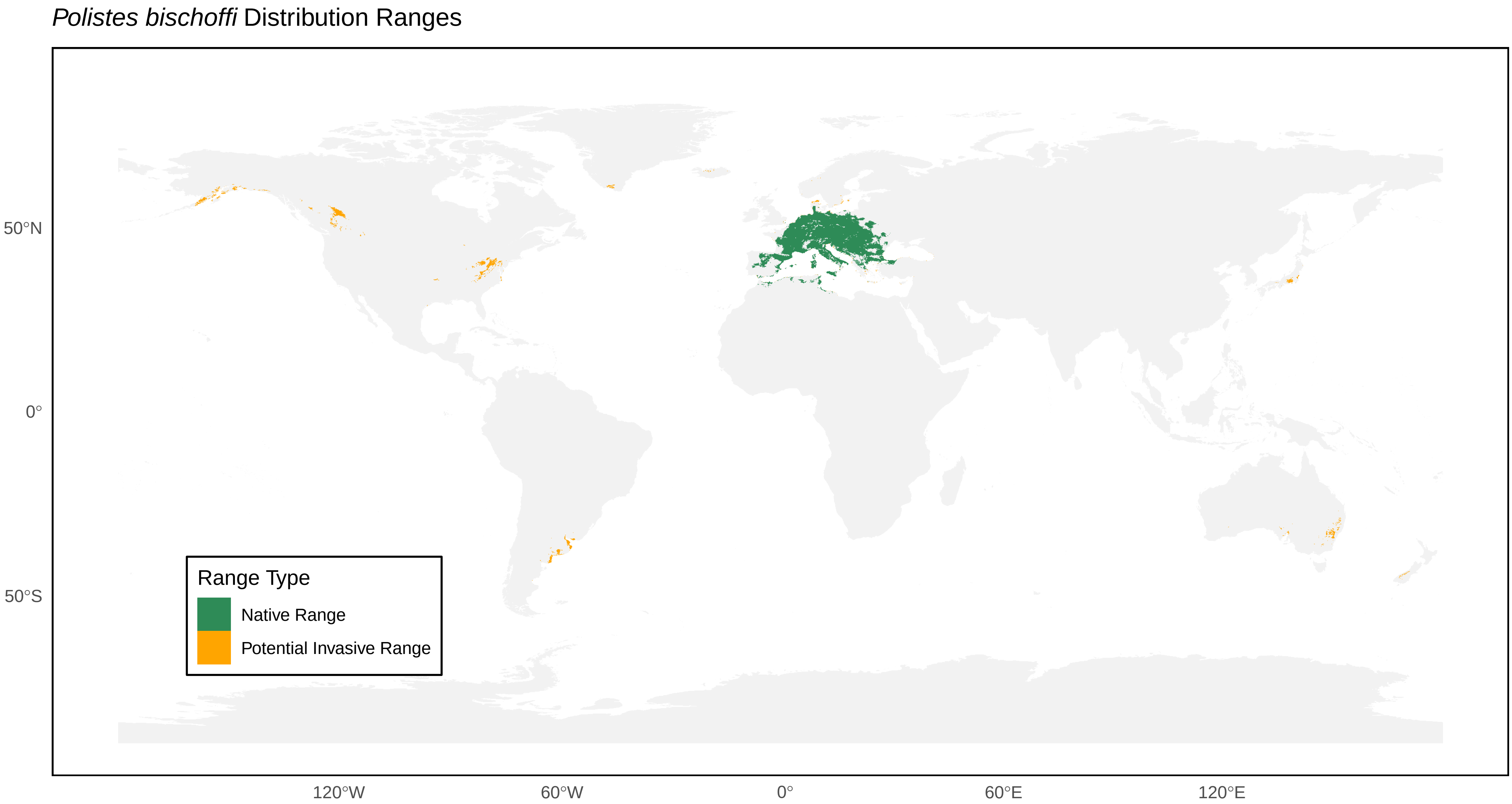
*Polistes bischoffi* Distribution Ranges
50°N
0°
50°S
120°W
60°W
0°
60°E
120°E
Range Type
Native Range
Potential Invasive Range

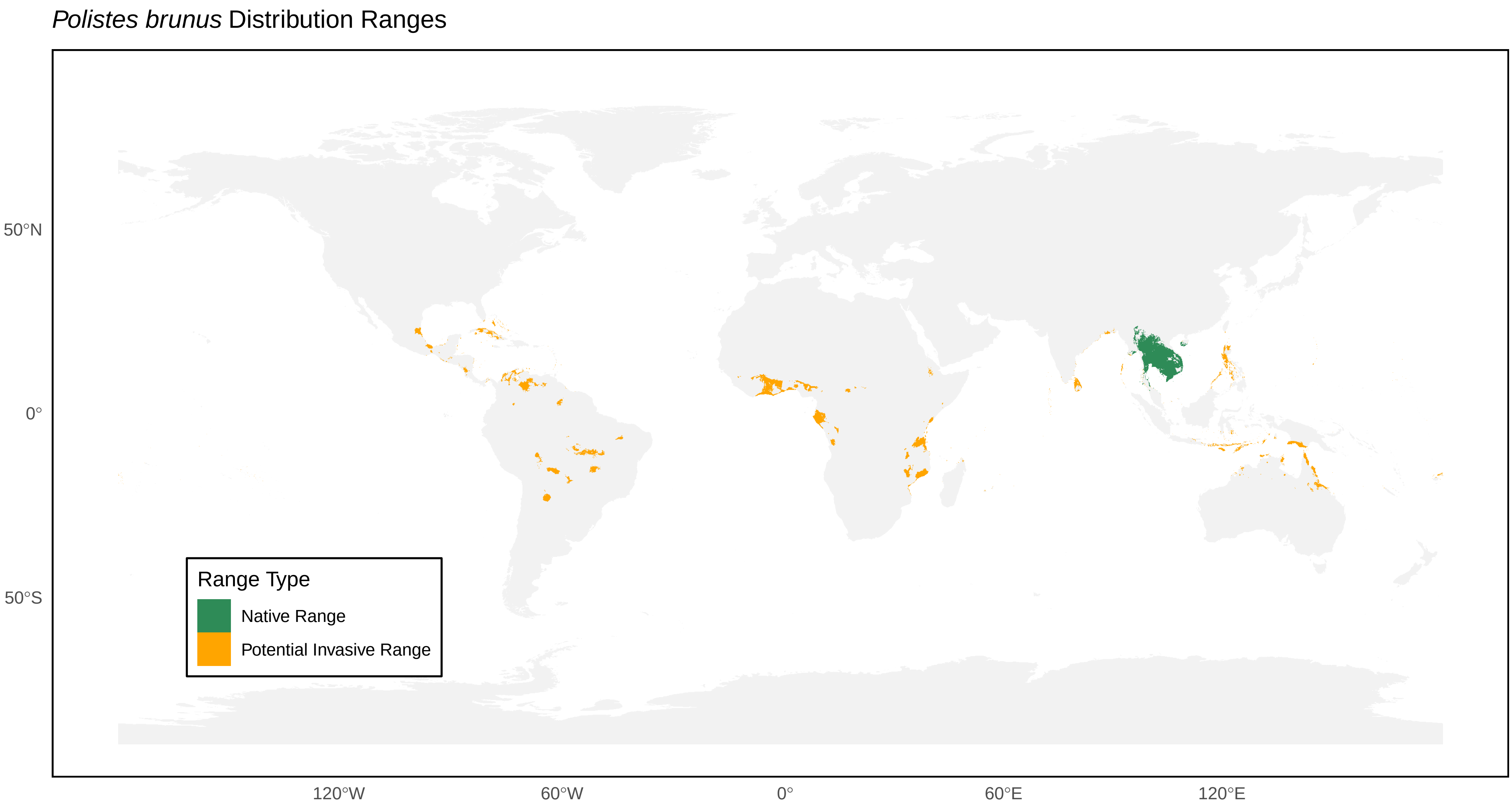
*Polistes brunus* Distribution Ranges
50°N
0°
50°S
120°W
60°W
0°
60°E
120°E
Range Type
Native Range
Potential Invasive Range

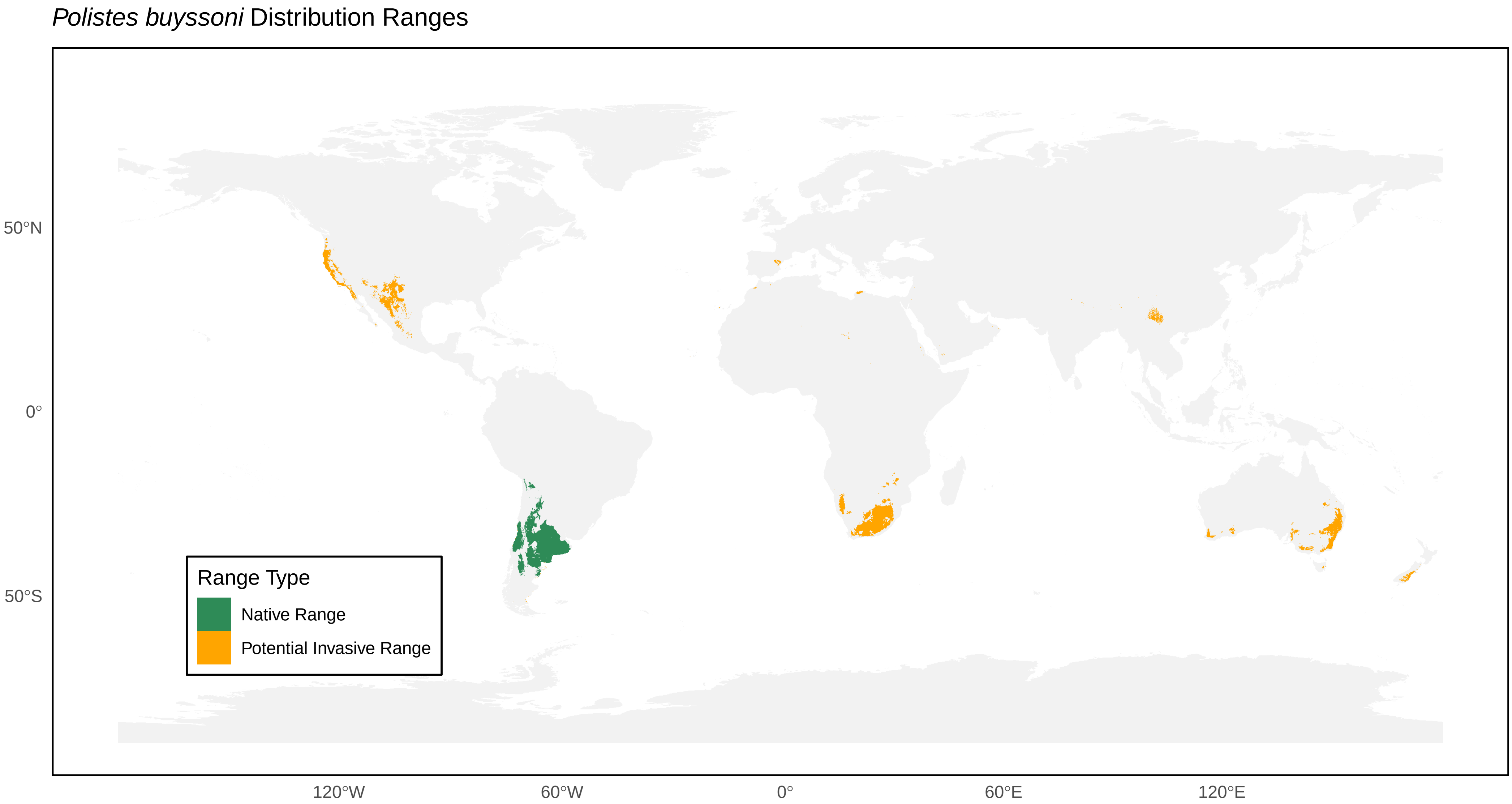
*Polistes buyssoni* Distribution Ranges
50°N
0°
50°S
120°W
60°W
0°
60°E
120°E
Range Type
Native Range
Potential Invasive Range

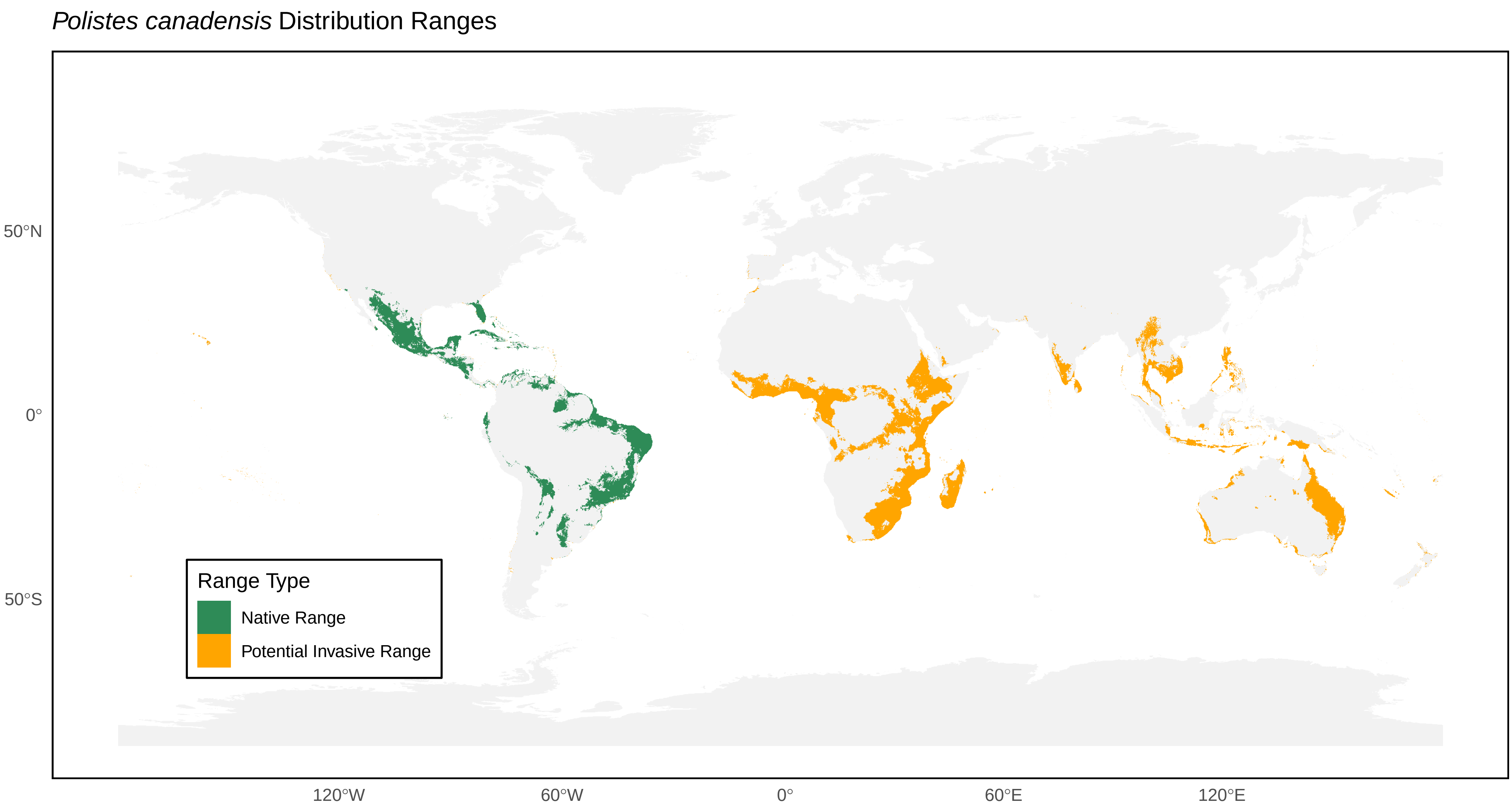
Polistes canadensis Distribution Ranges
50°N
0°
50°S
120°W
60°W
0°
60°E
120°E
Range Type
Native Range
Potential Invasive Range

#### *Polistes carnifex* Distribution Ranges

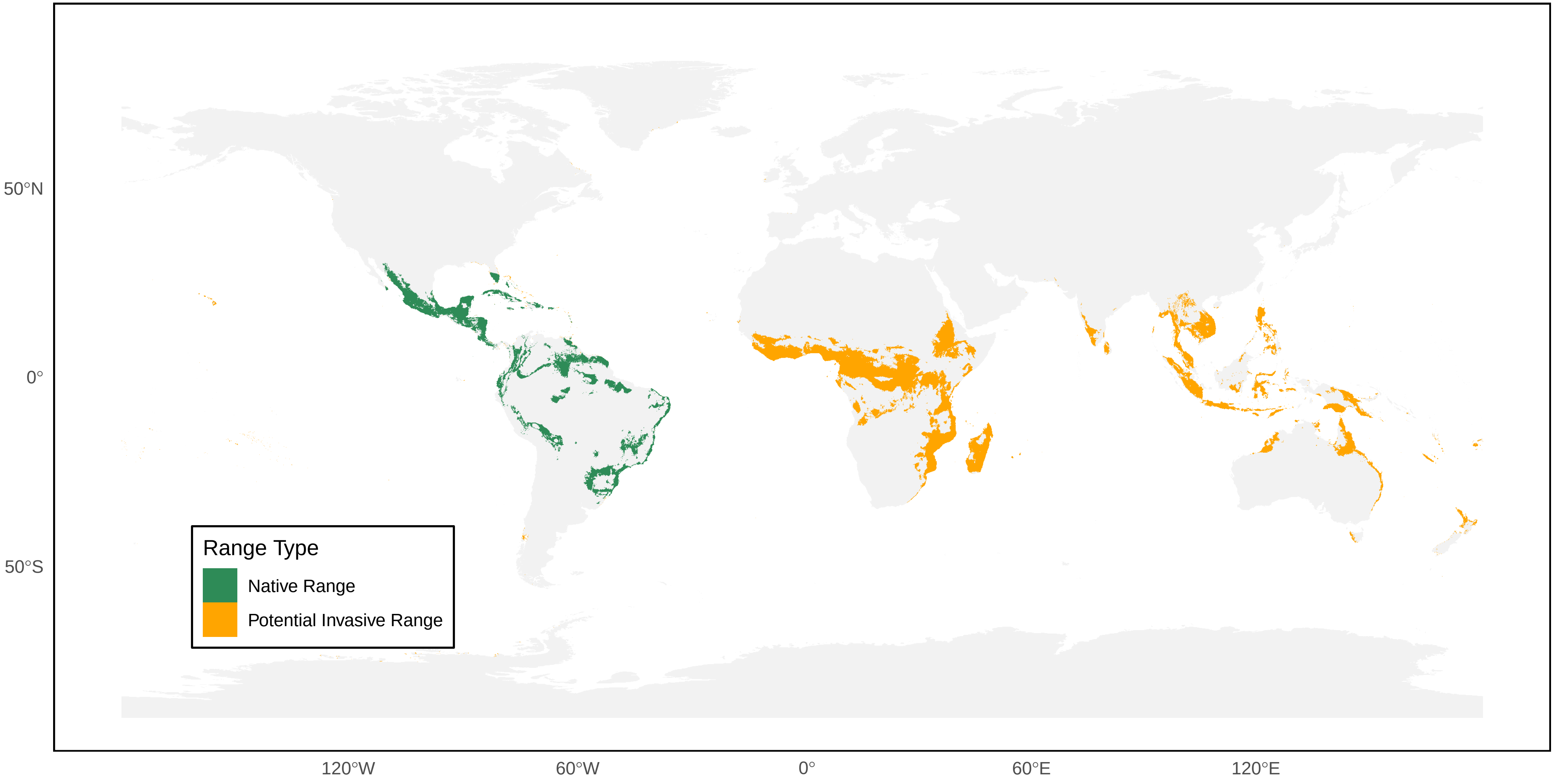

*Polistes carolina* Distribution Ranges

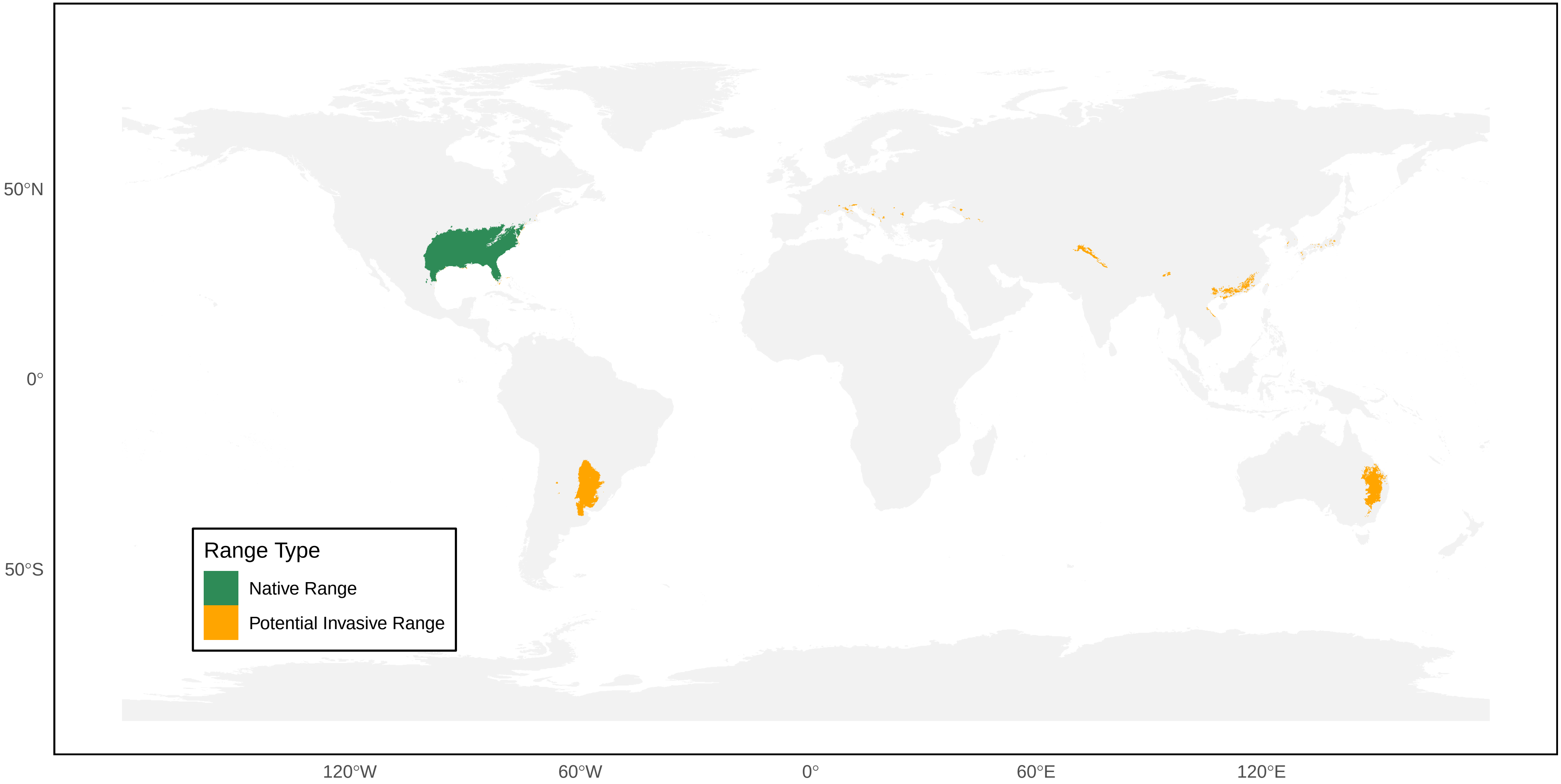

*Polistes cavapyta* Distribution Ranges

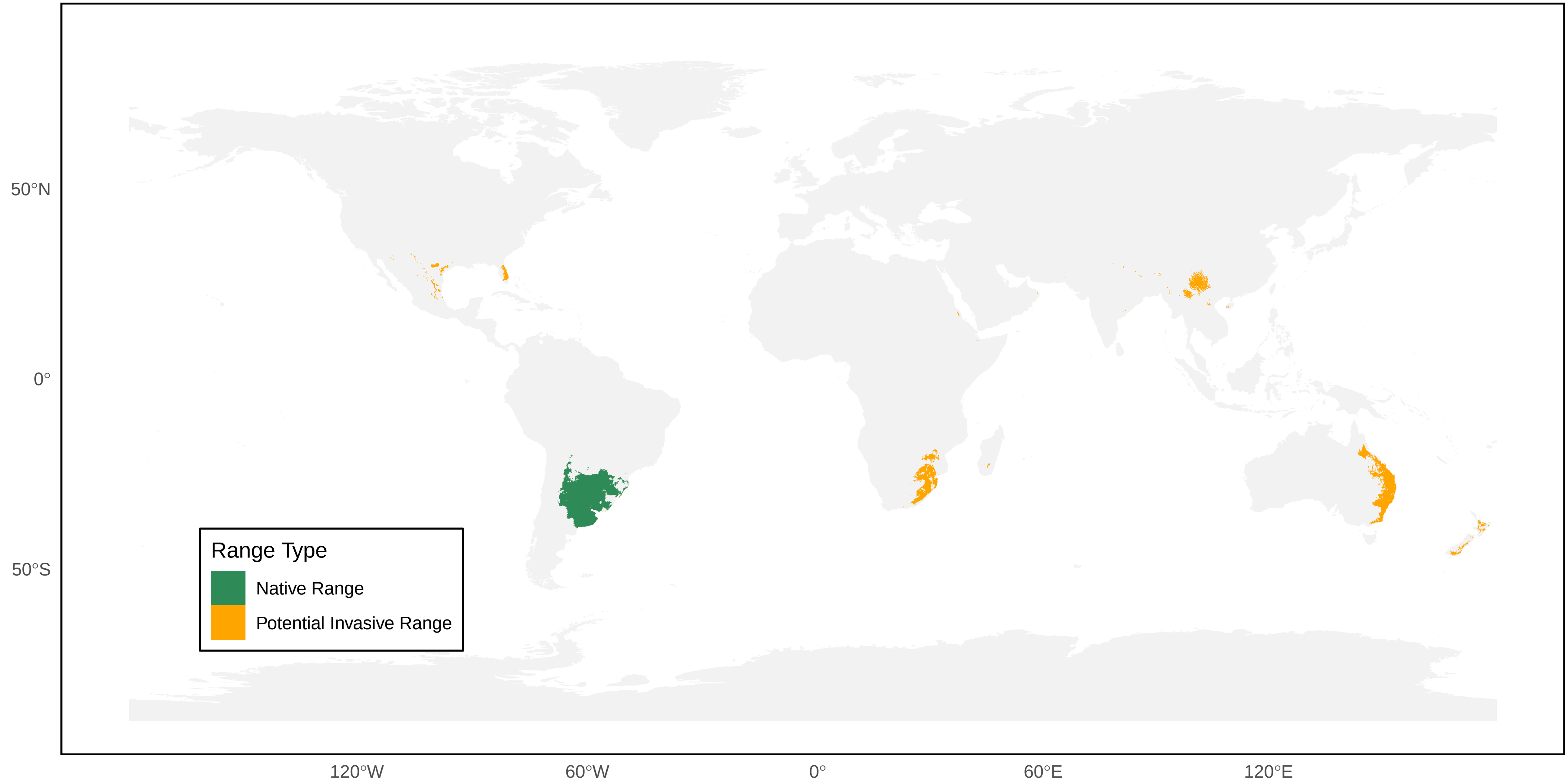

*Polistes cavapytiformis* Distribution Ranges

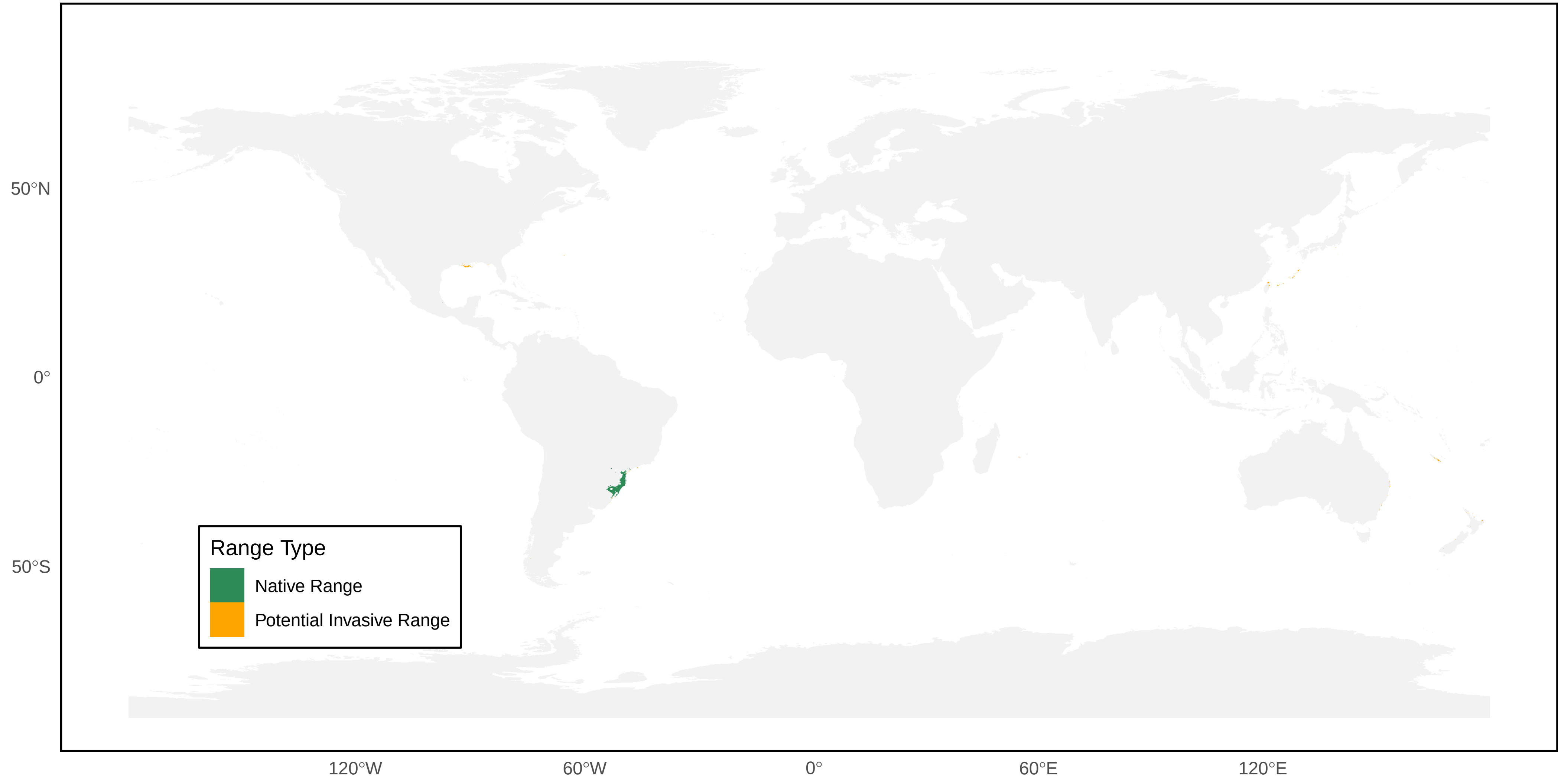

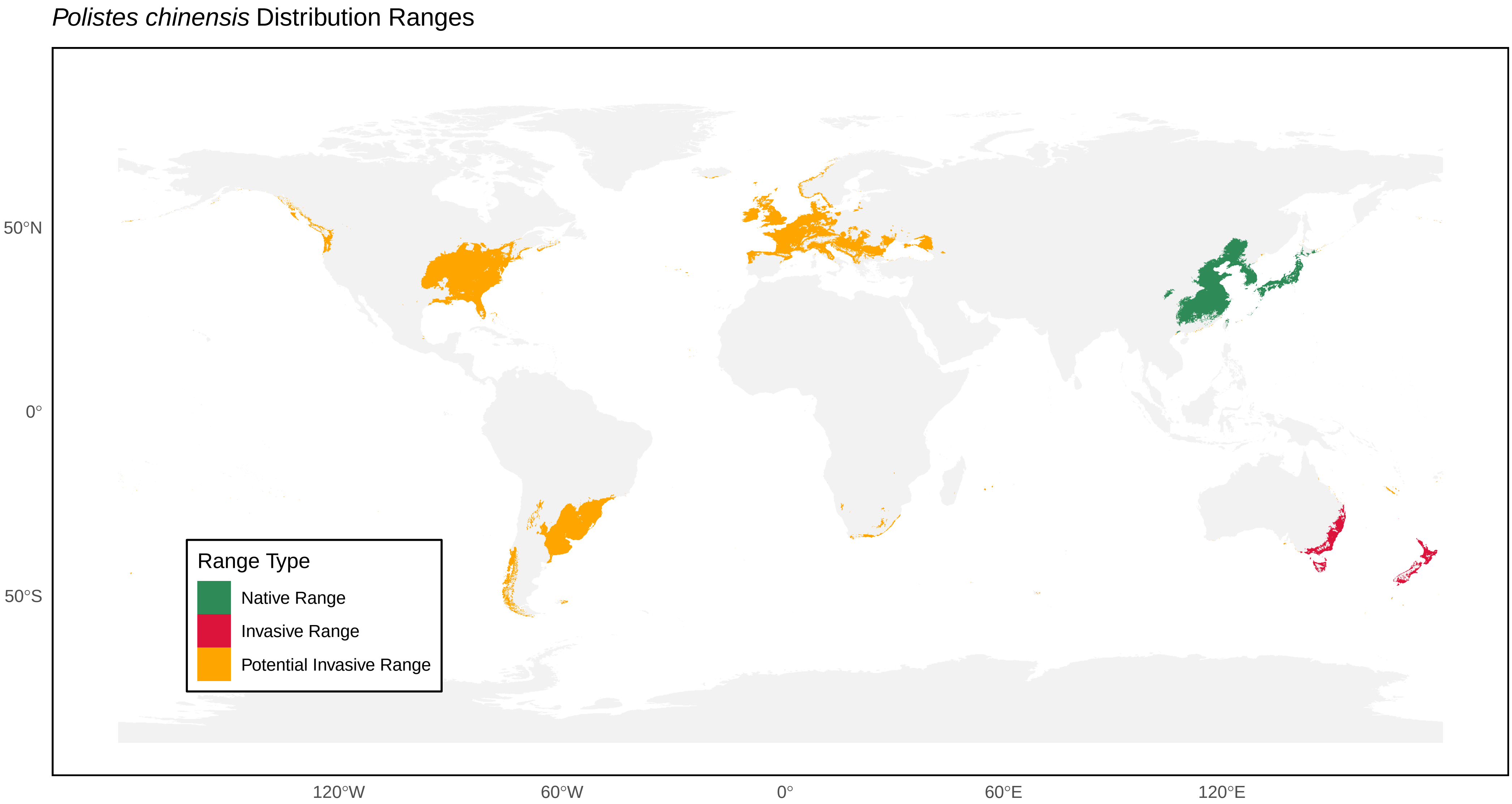
Polistes chinensis Distribution Ranges
Range Type
Native Range
Invasive Range
Potential Invasive Range
50°N
0°
50°S
120°W
60°W
0°
60°E
120°E

### *Polistes cinerascens* Distribution Ranges

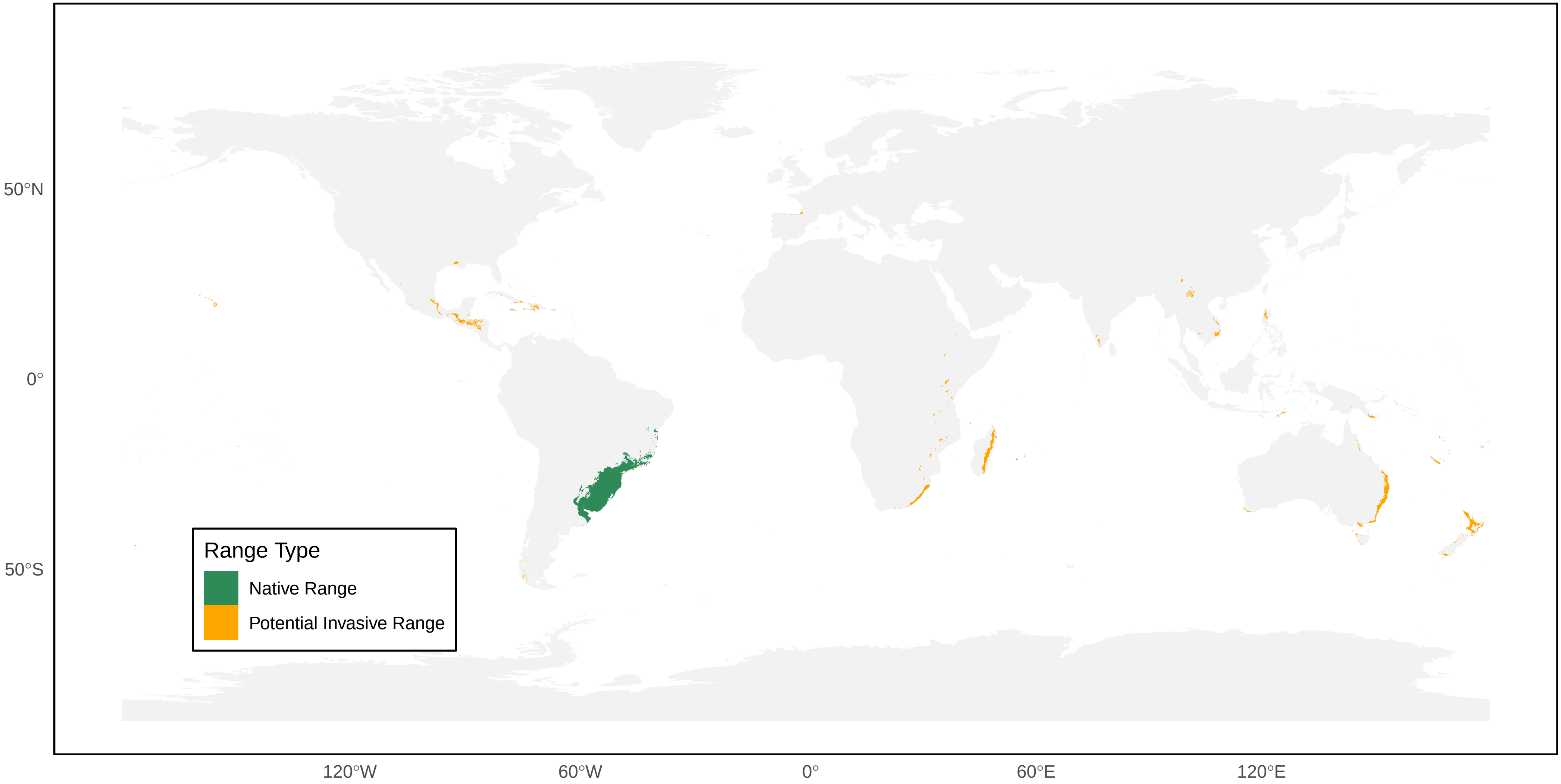

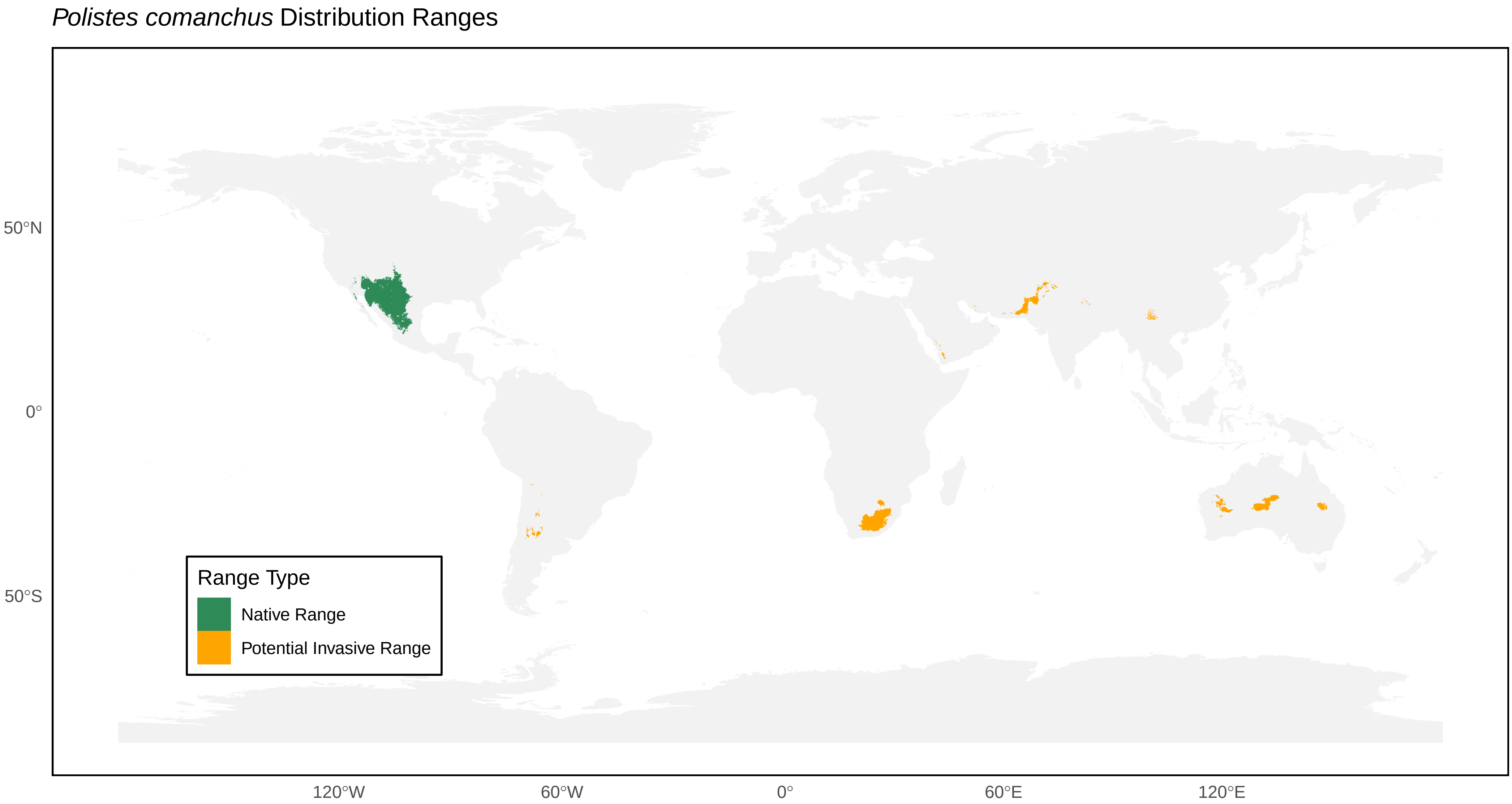
*Polistes comanchus* Distribution Ranges
50°N
0°
50°S
120°W
60°W
0°
60°E
120°E
Range Type
Native Range
Potential Invasive Range

*Polistes crinitus* Distribution Ranges

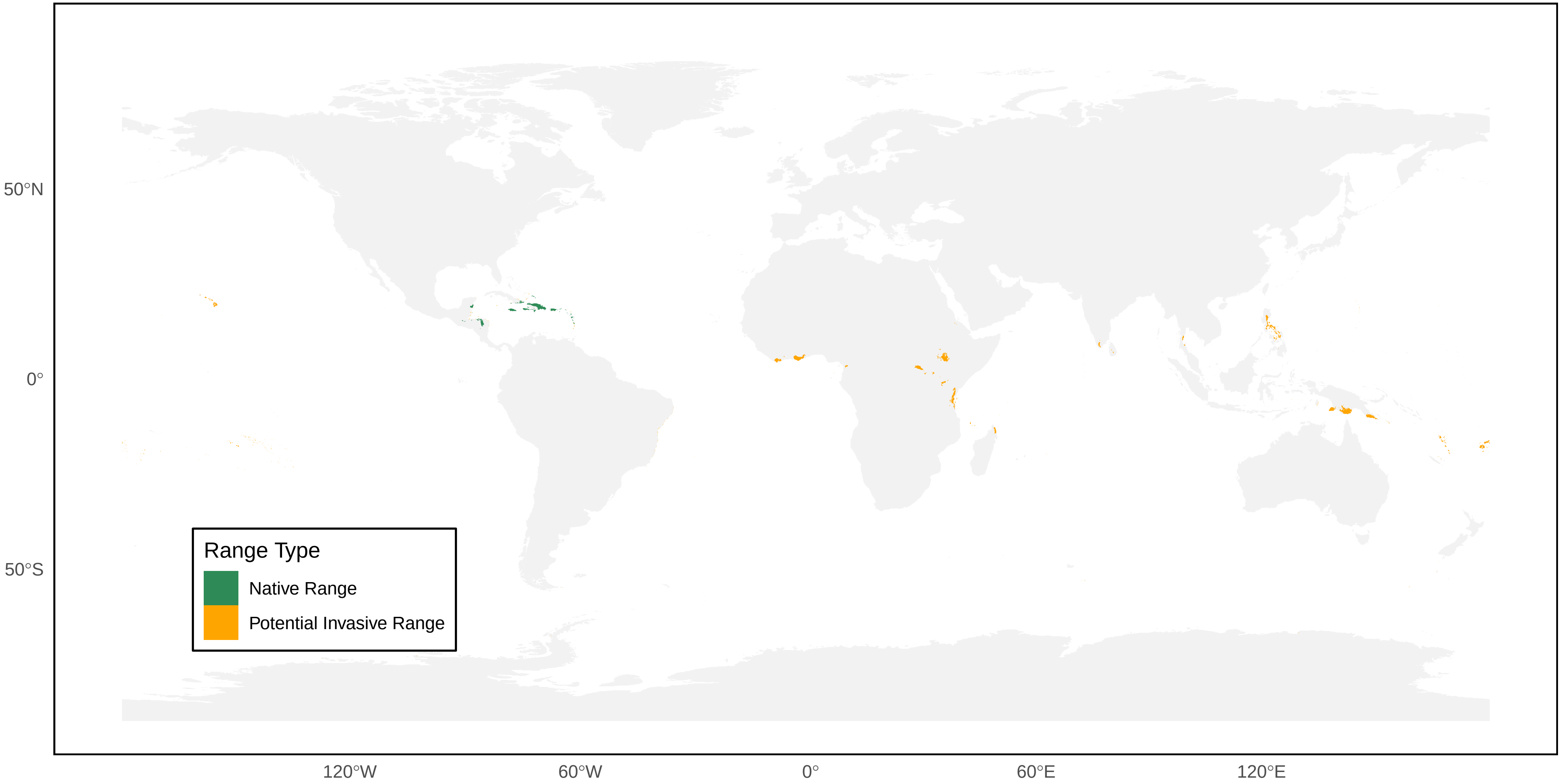

*Polistes cubensis* Distribution Ranges

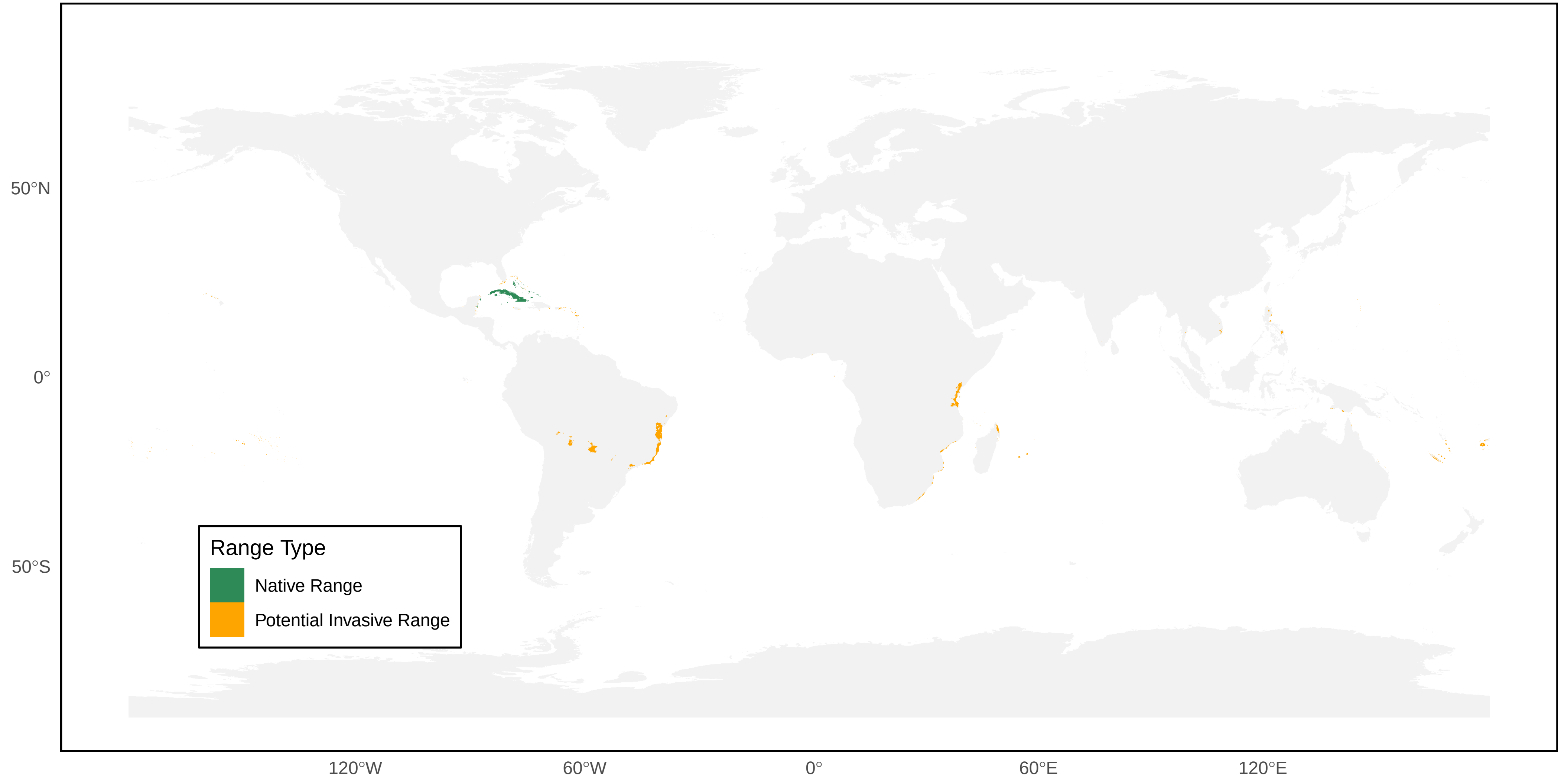

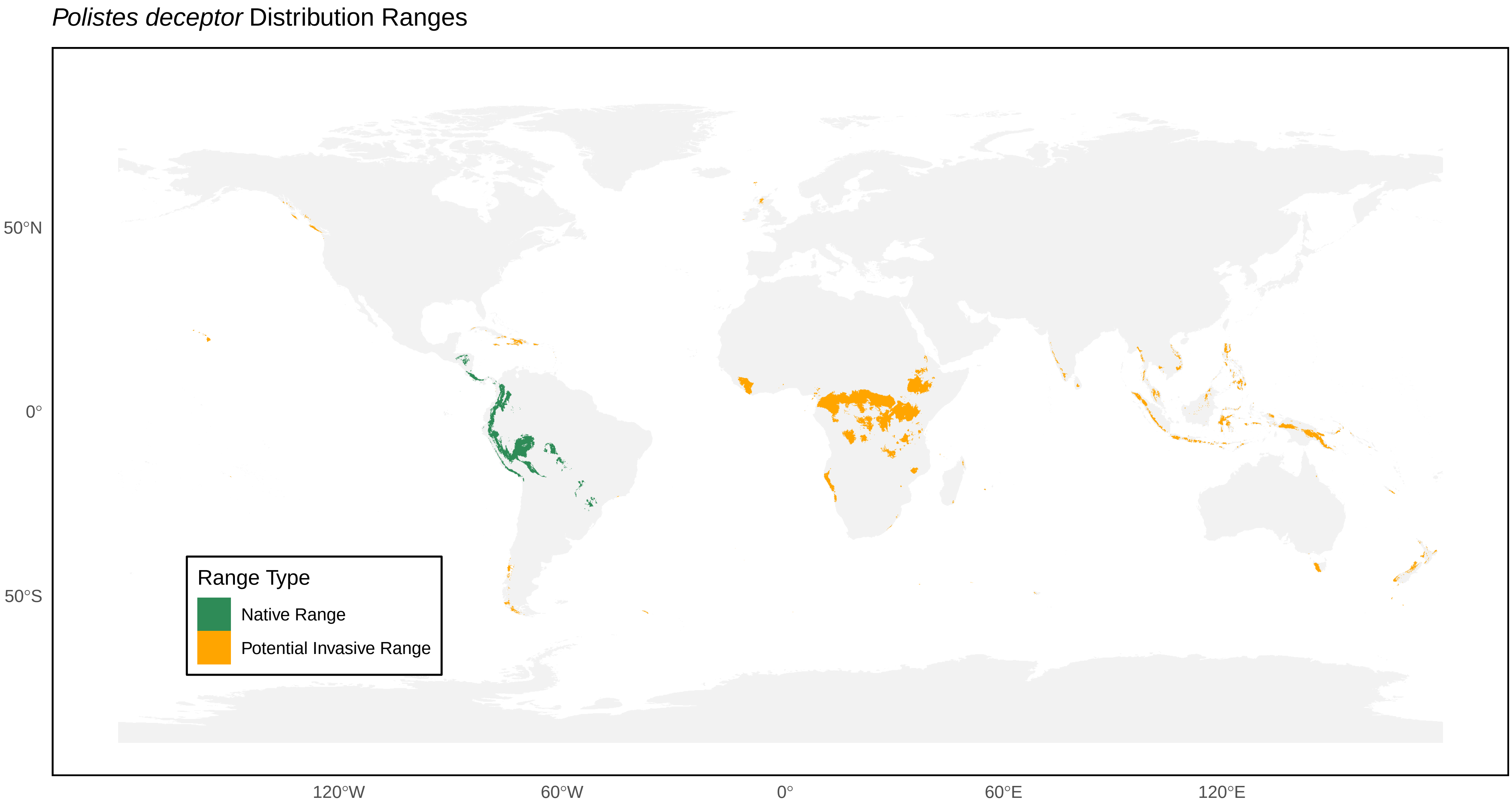

*Polistes deceptor* Distribution Ranges
50°N
0°
50°S
120°W
60°W
0°
60°E
120°E
Range Type
Native Range
Potential Invasive Range

*Polistes diabolicus* Distribution Ranges

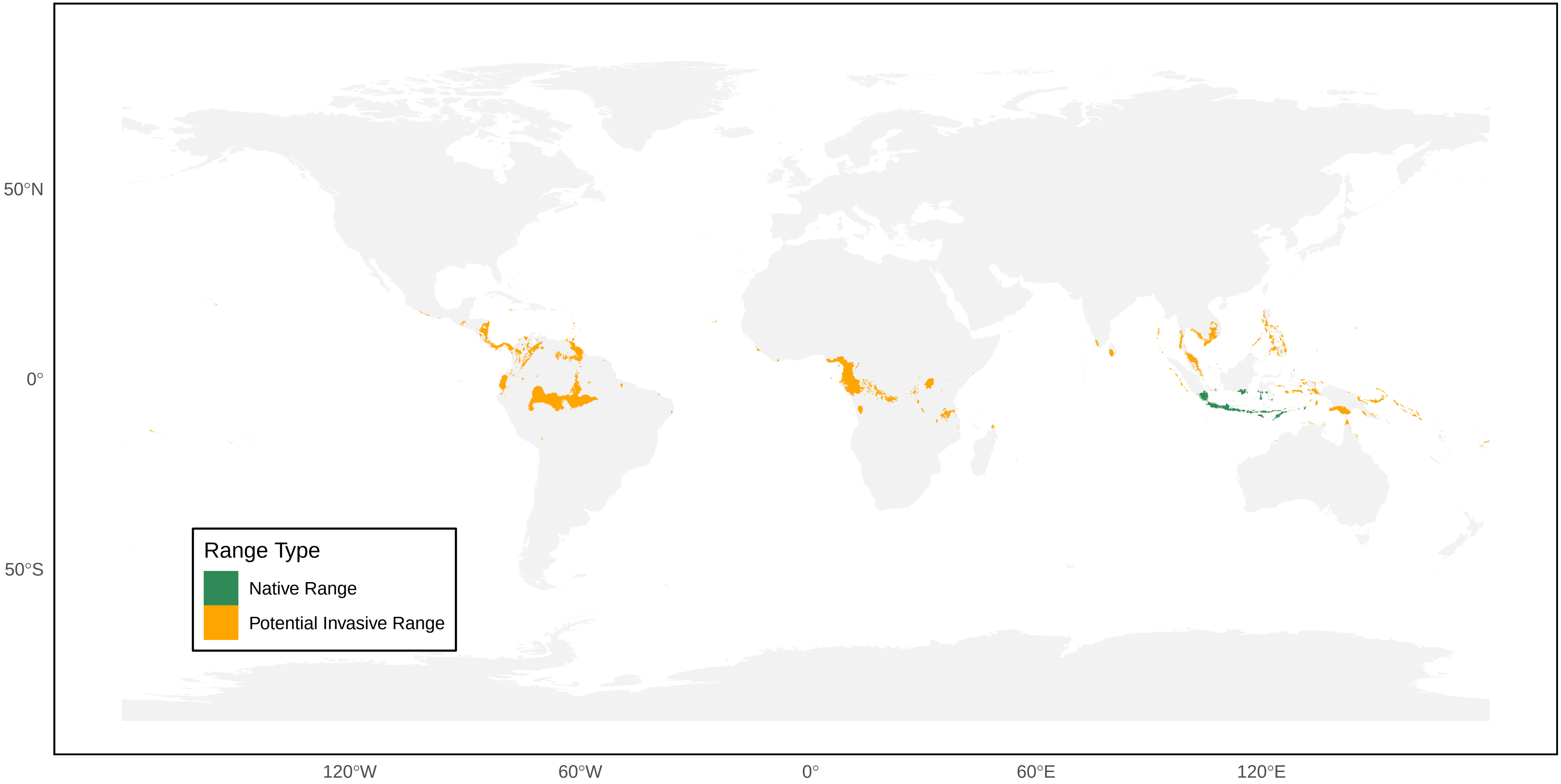

### *Polistes dominicus* Distribution Ranges

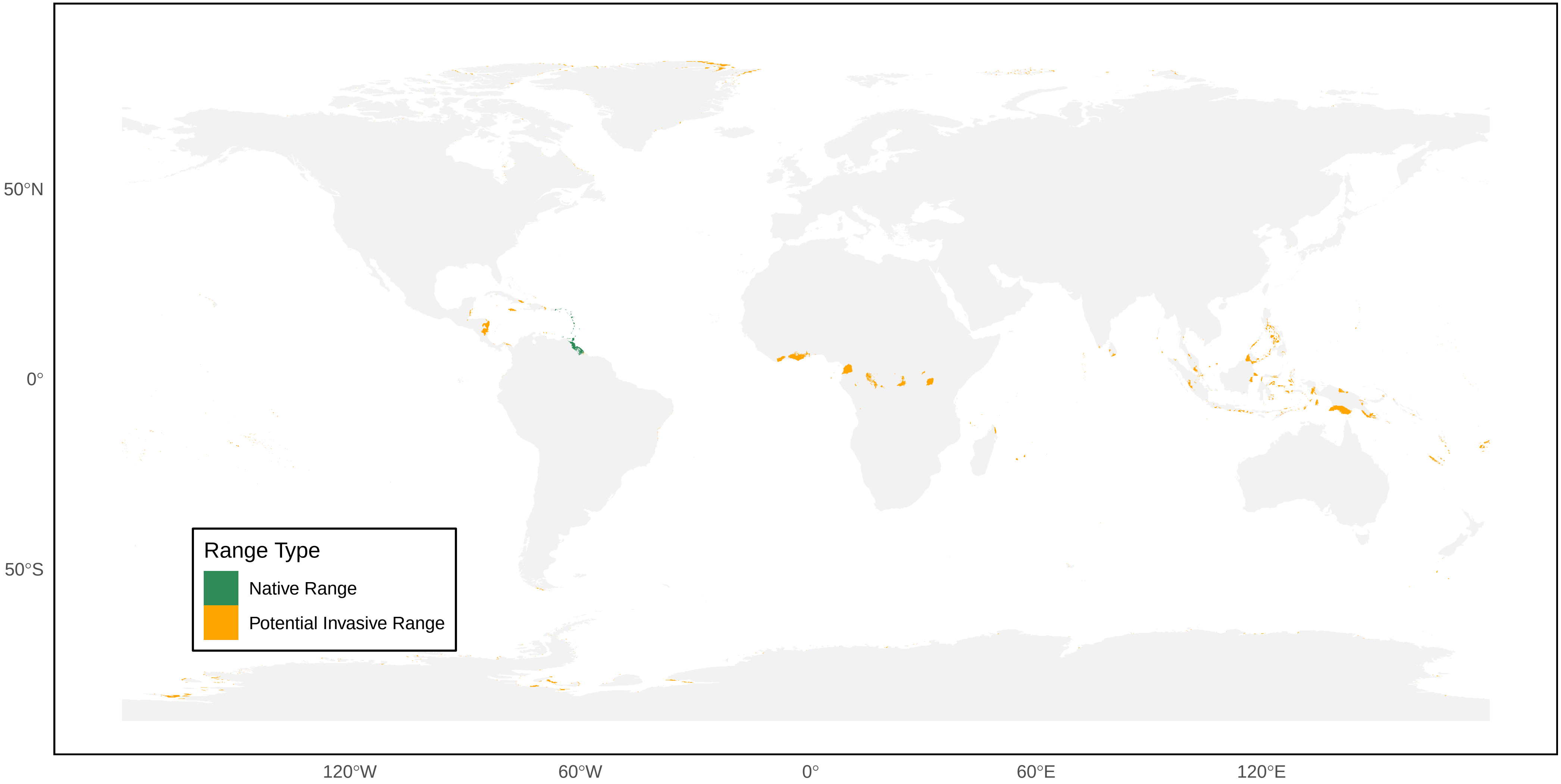

*Polistes dominula* Distribution Ranges

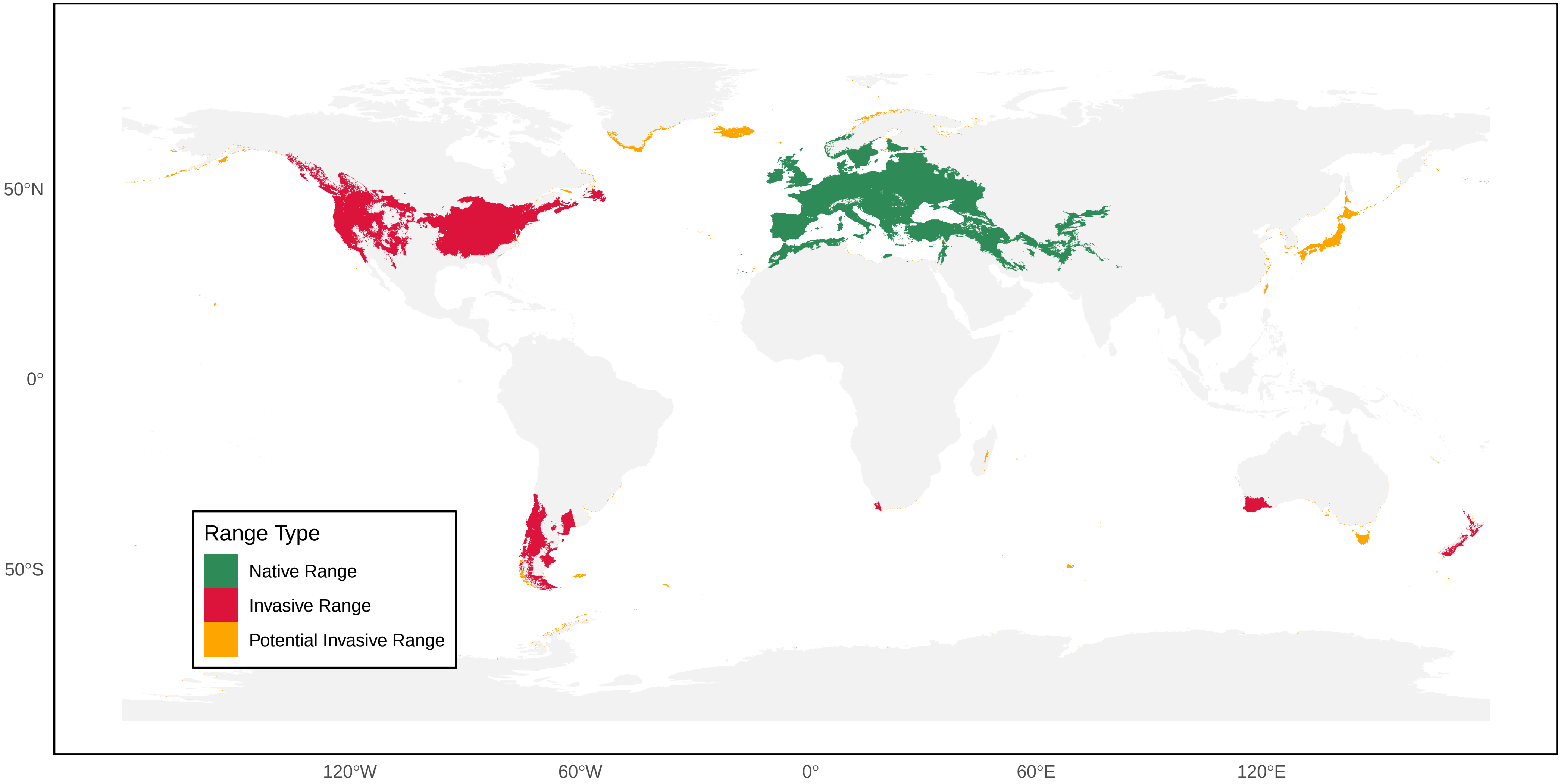

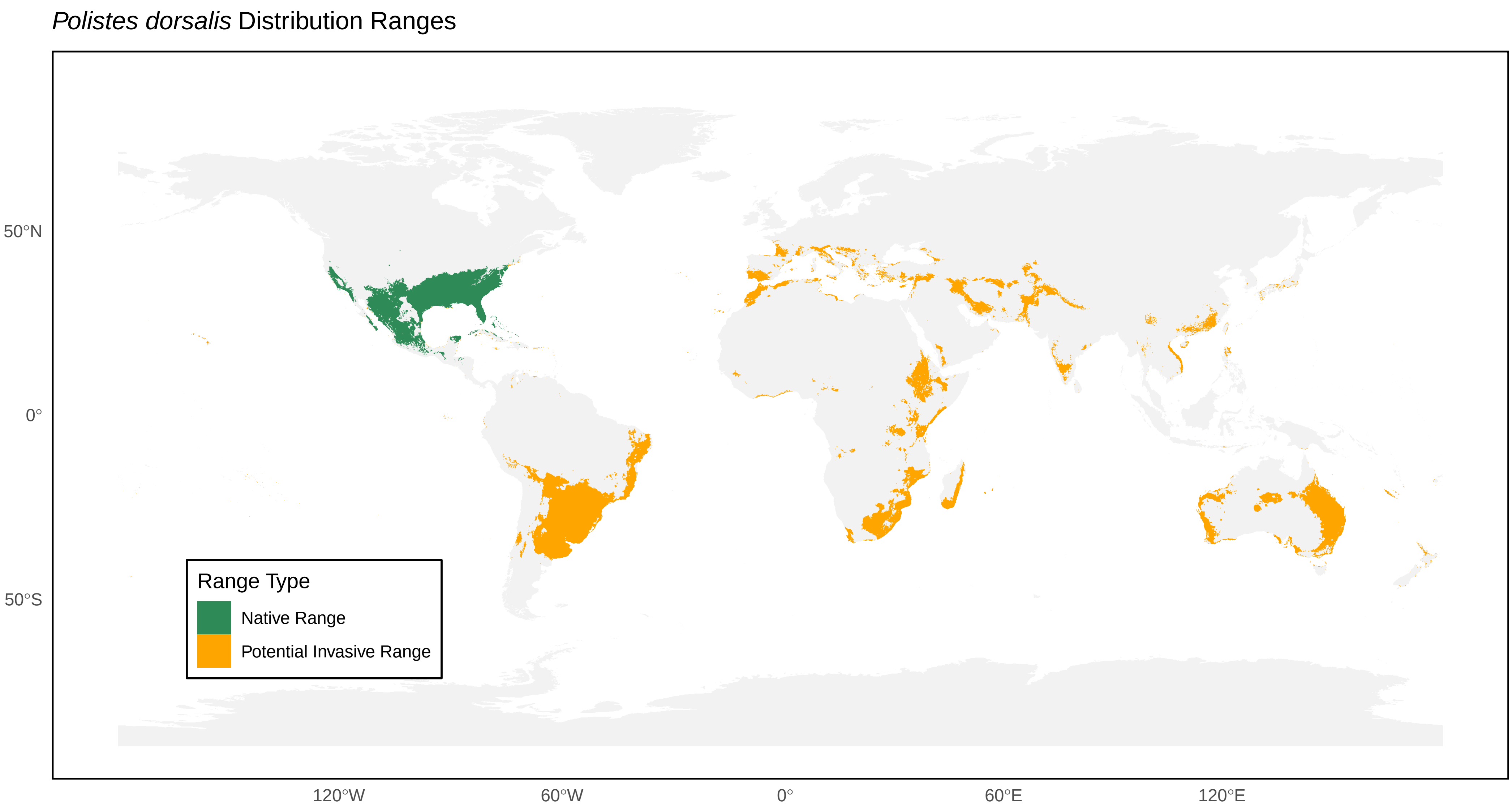
Polistes dorsalis Distribution Ranges
50°N
0°
50°S
120°W
60°W
0°
60°E
120°E
Range Type
Native Range
Potential Invasive Range

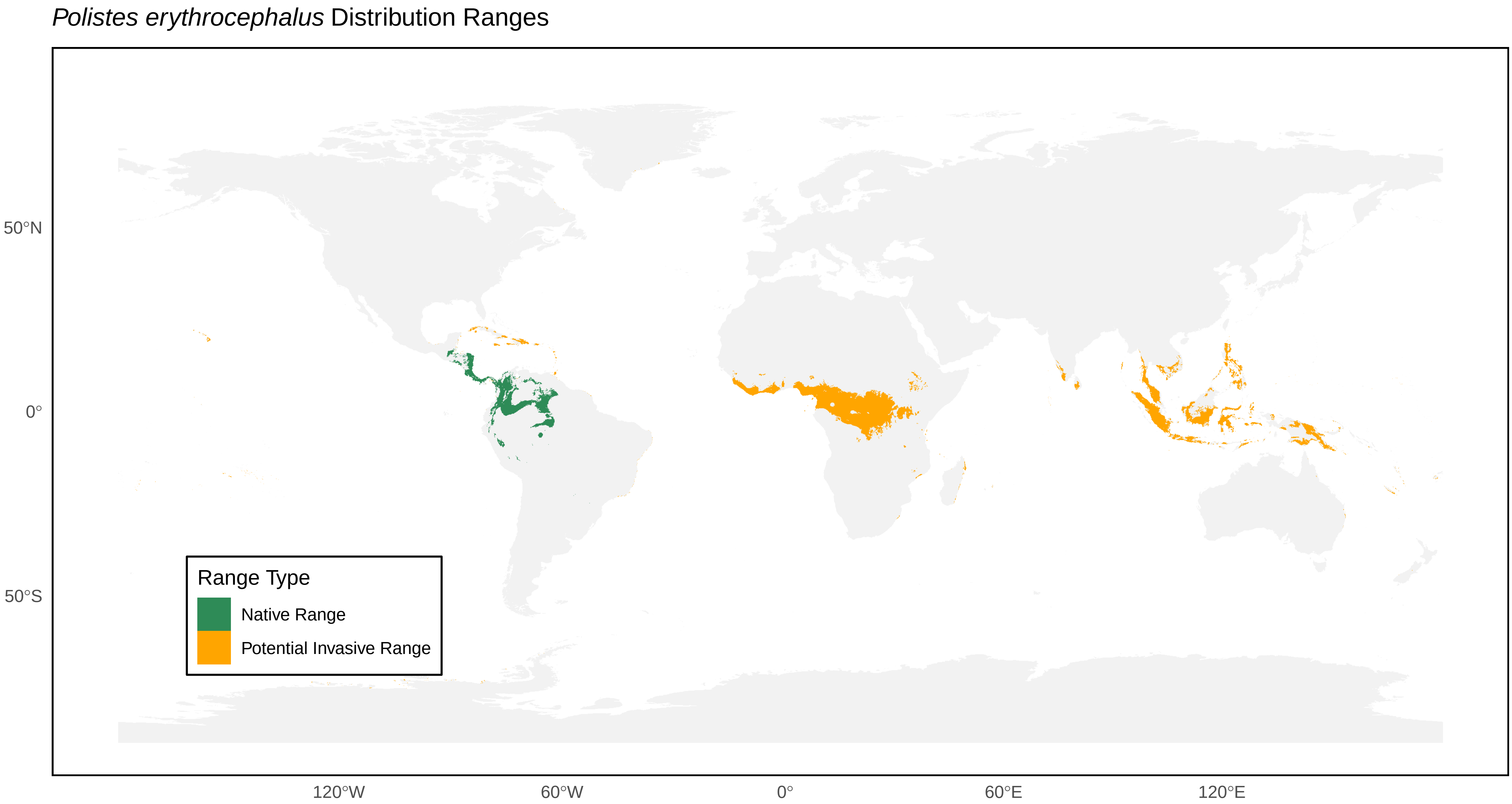

*Polistes erythrocephalus* Distribution Ranges
50°N
0°
50°S
120°W
60°W
0°
60°E
120°E
Range Type
Native Range
Potential Invasive Range

*Polistes exclamans* Distribution Ranges

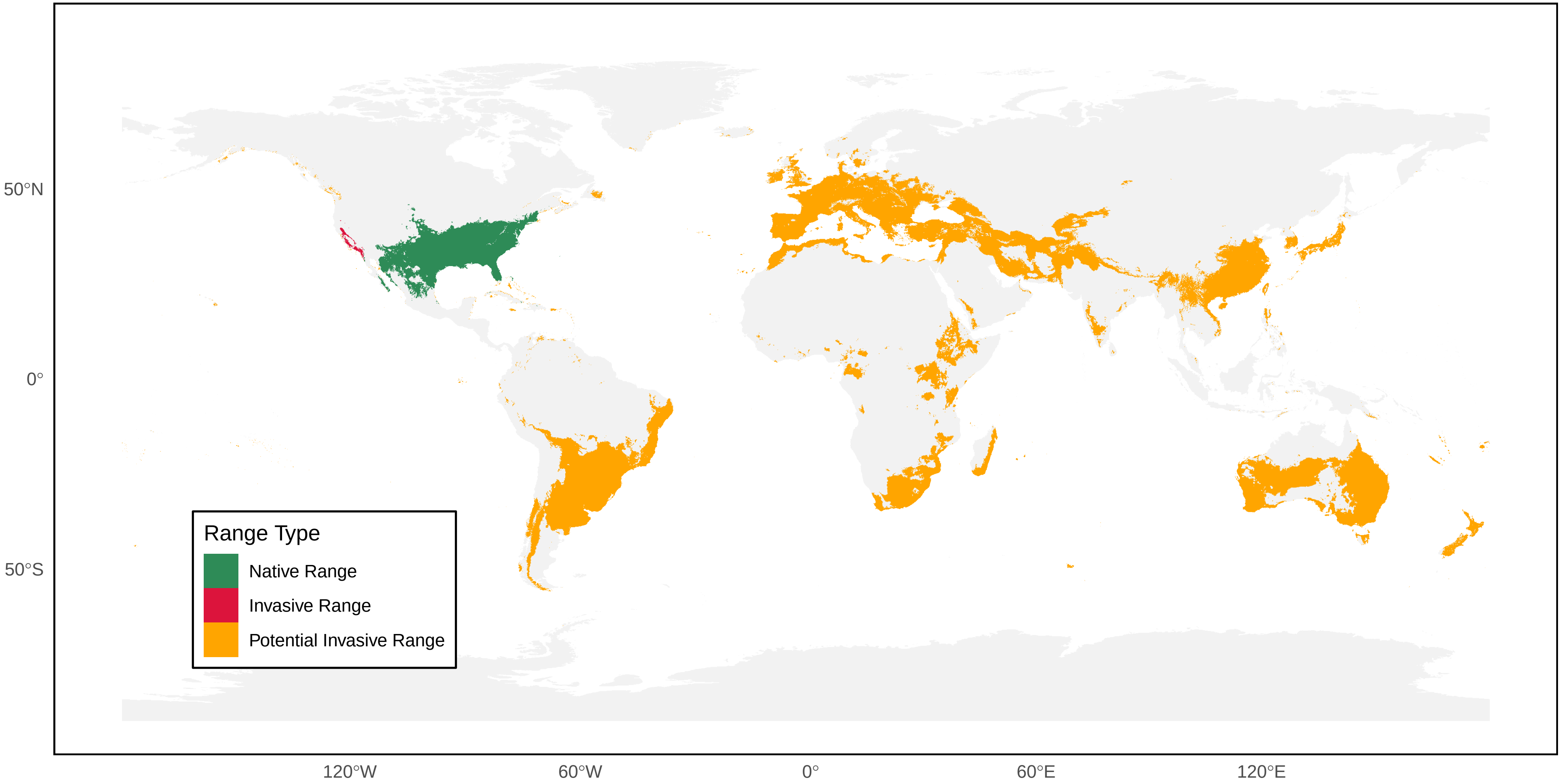

*Polistes fastidiosus* Distribution Ranges

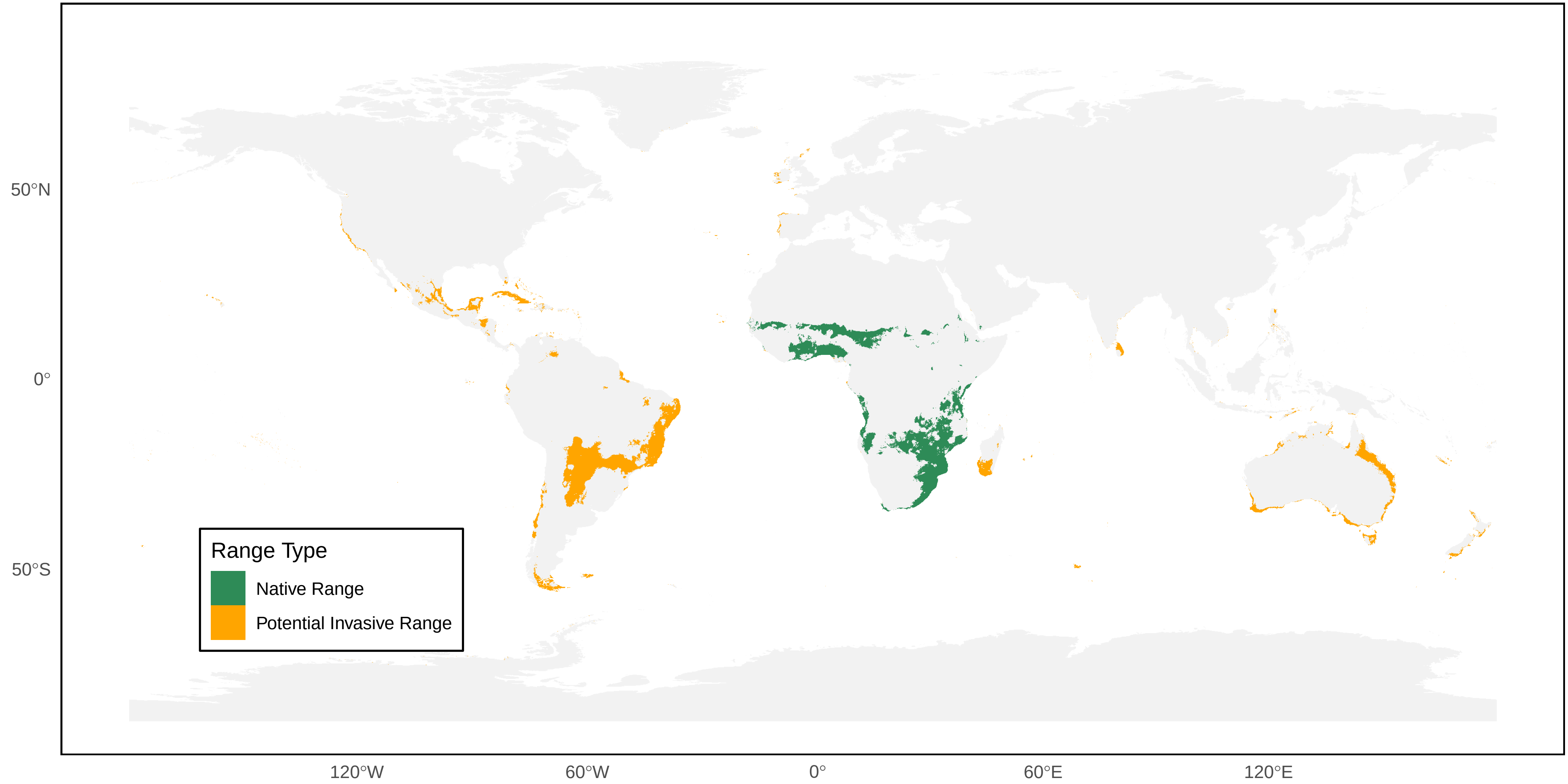

#### *Polistes ferreri* Distribution Ranges

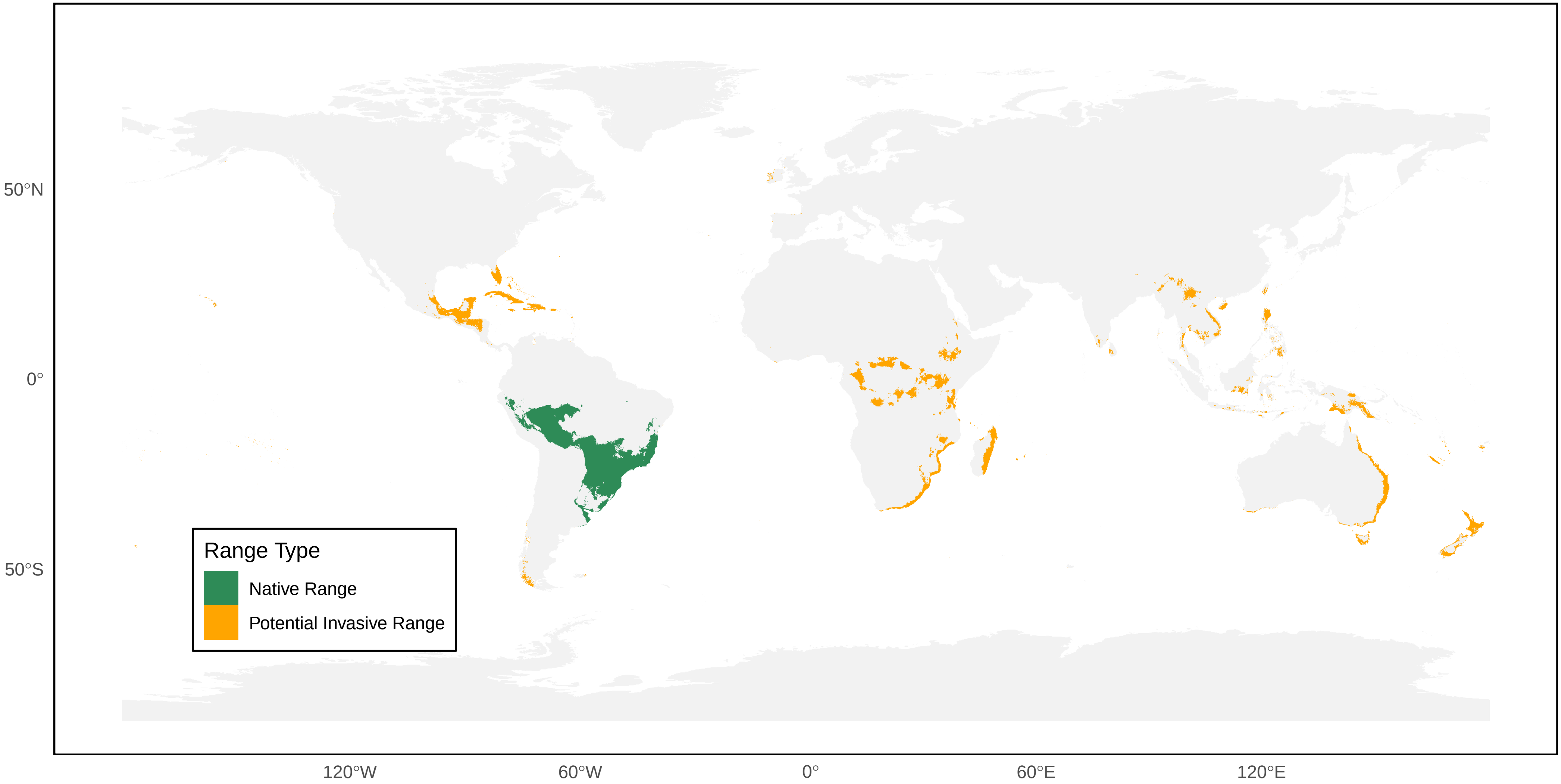

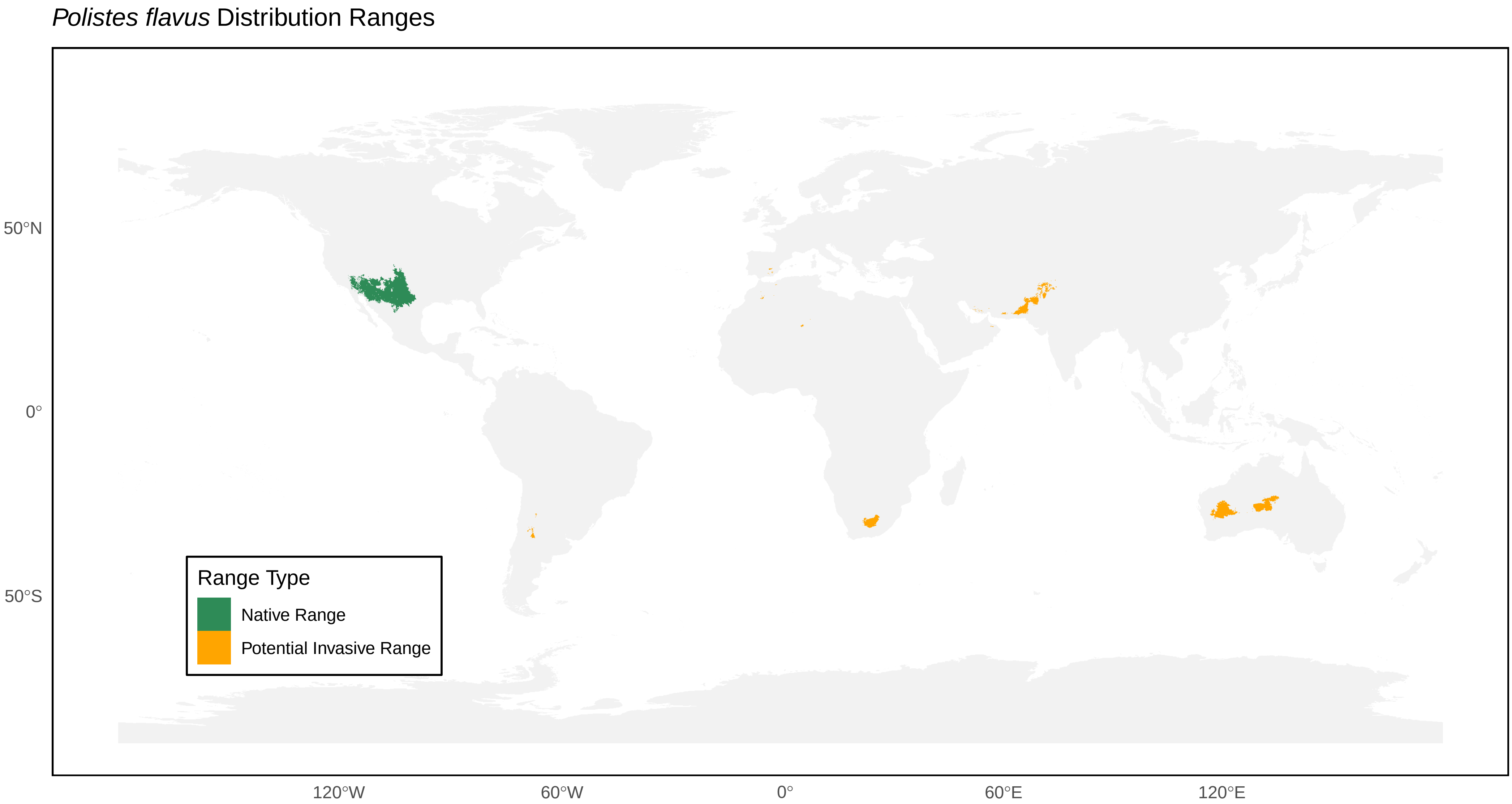
*Polistes flavus* Distribution Ranges
50°N
0°
50°S
120°W
60°W
0°
60°E
120°E
Range Type
Native Range
Potential Invasive Range

*Polistes fuscatus* Distribution Ranges

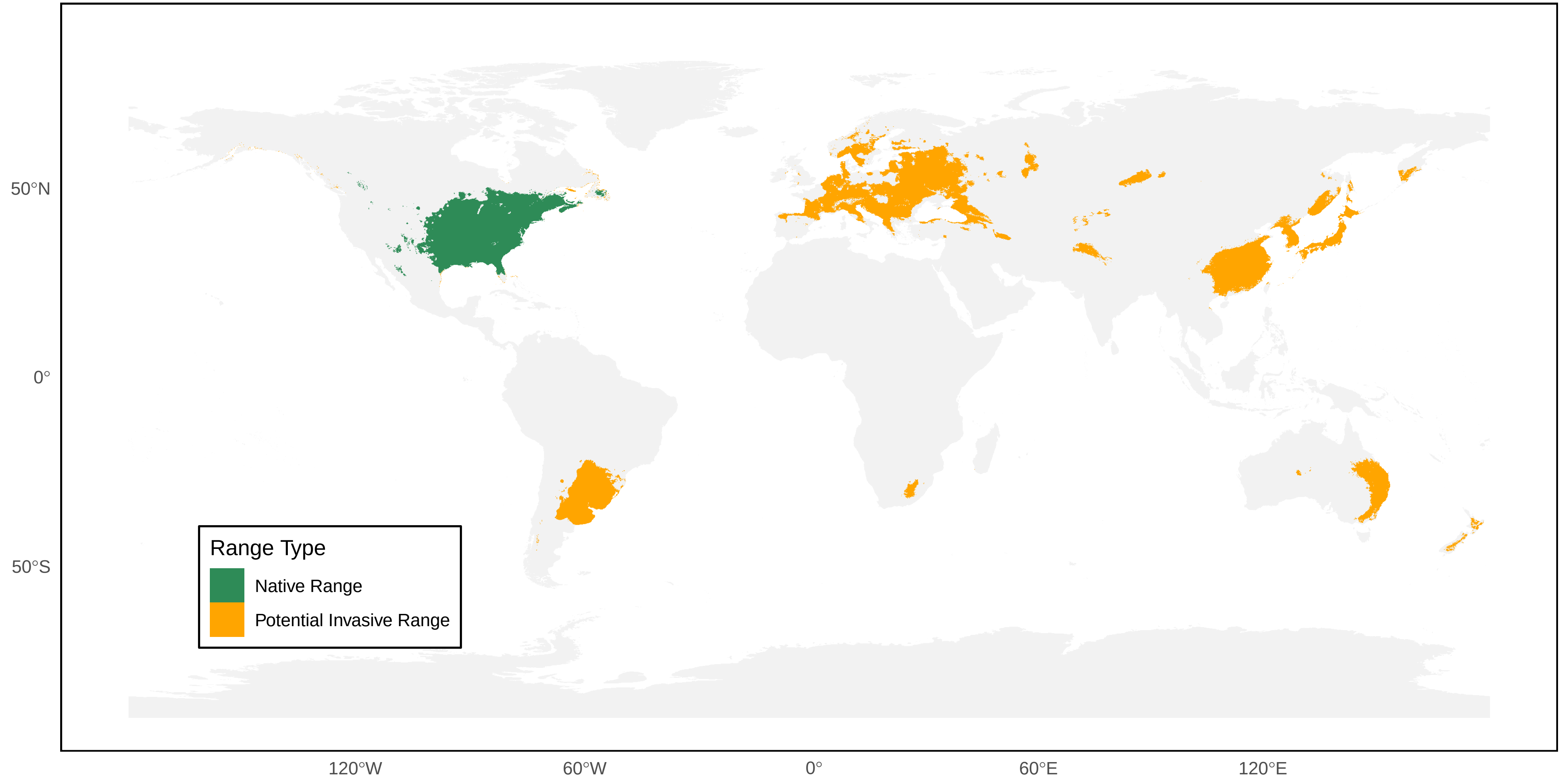

### *Polistes gallicus* Distribution Ranges

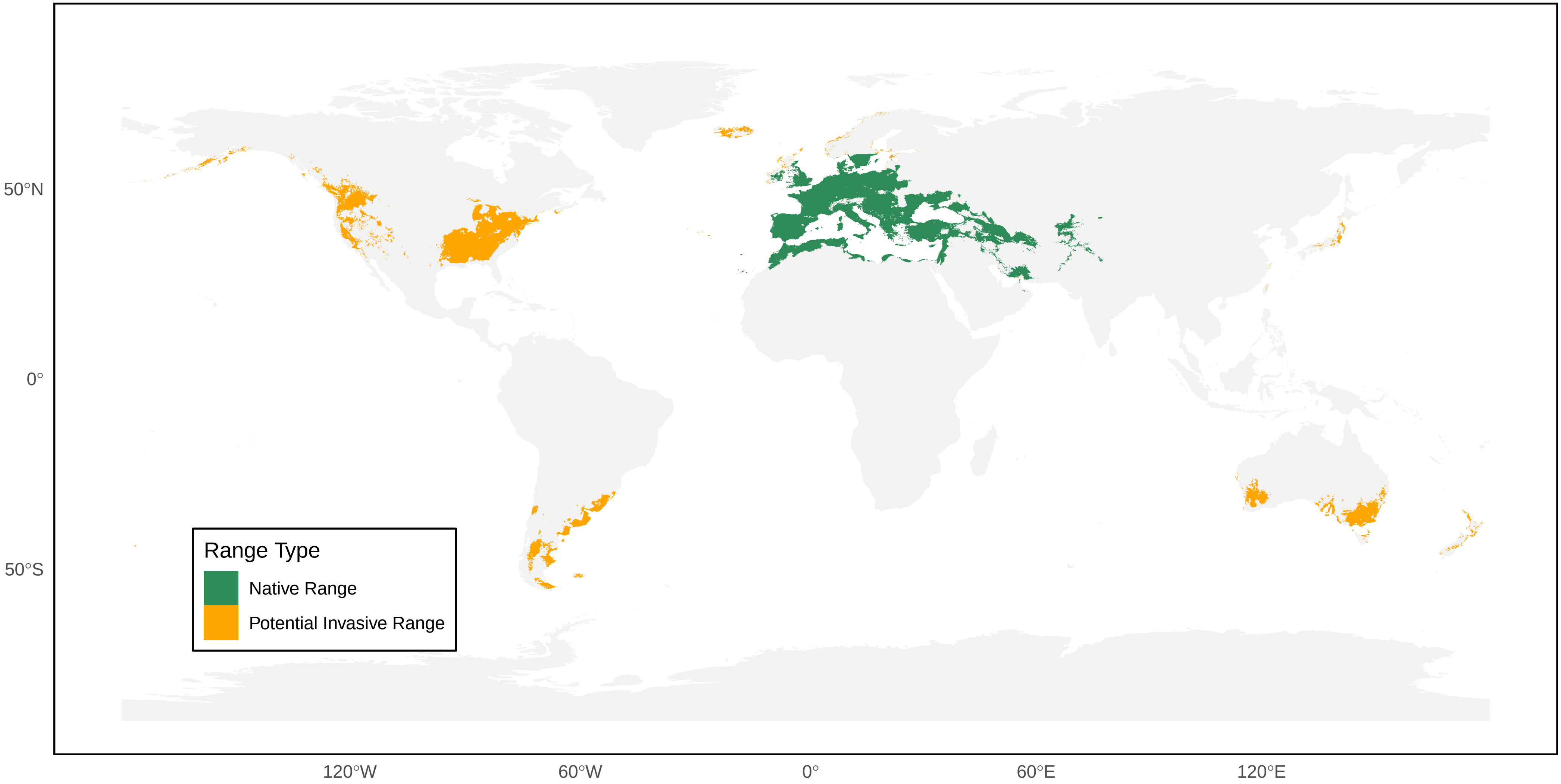

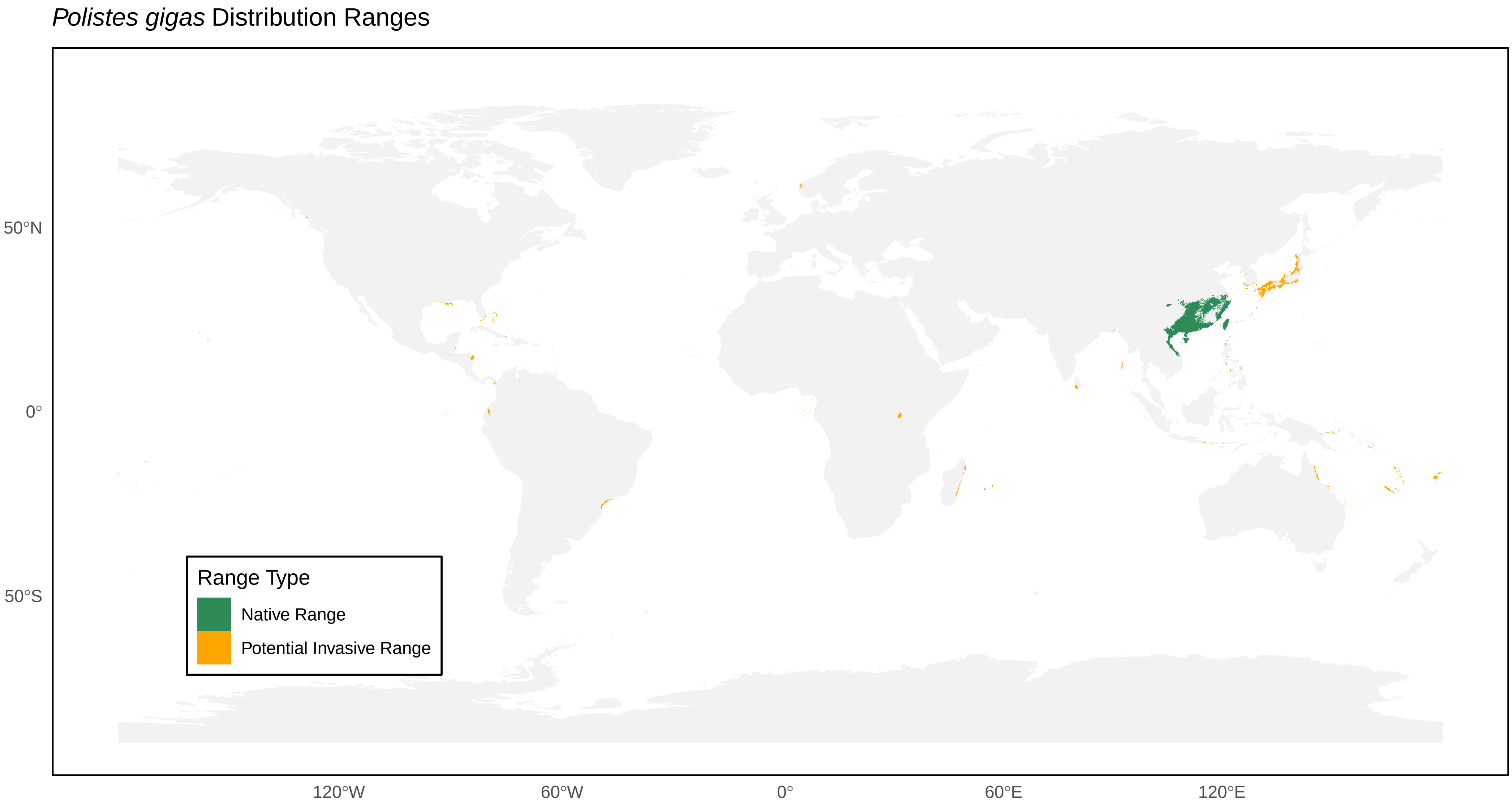
*Polistes gigas* Distribution Ranges
50°N
0°
50°S
120°W
60°W
0°
60°E
120°E
Range Type
Native Range
Potential Invasive Range

*Polistes goeldii* Distribution Ranges

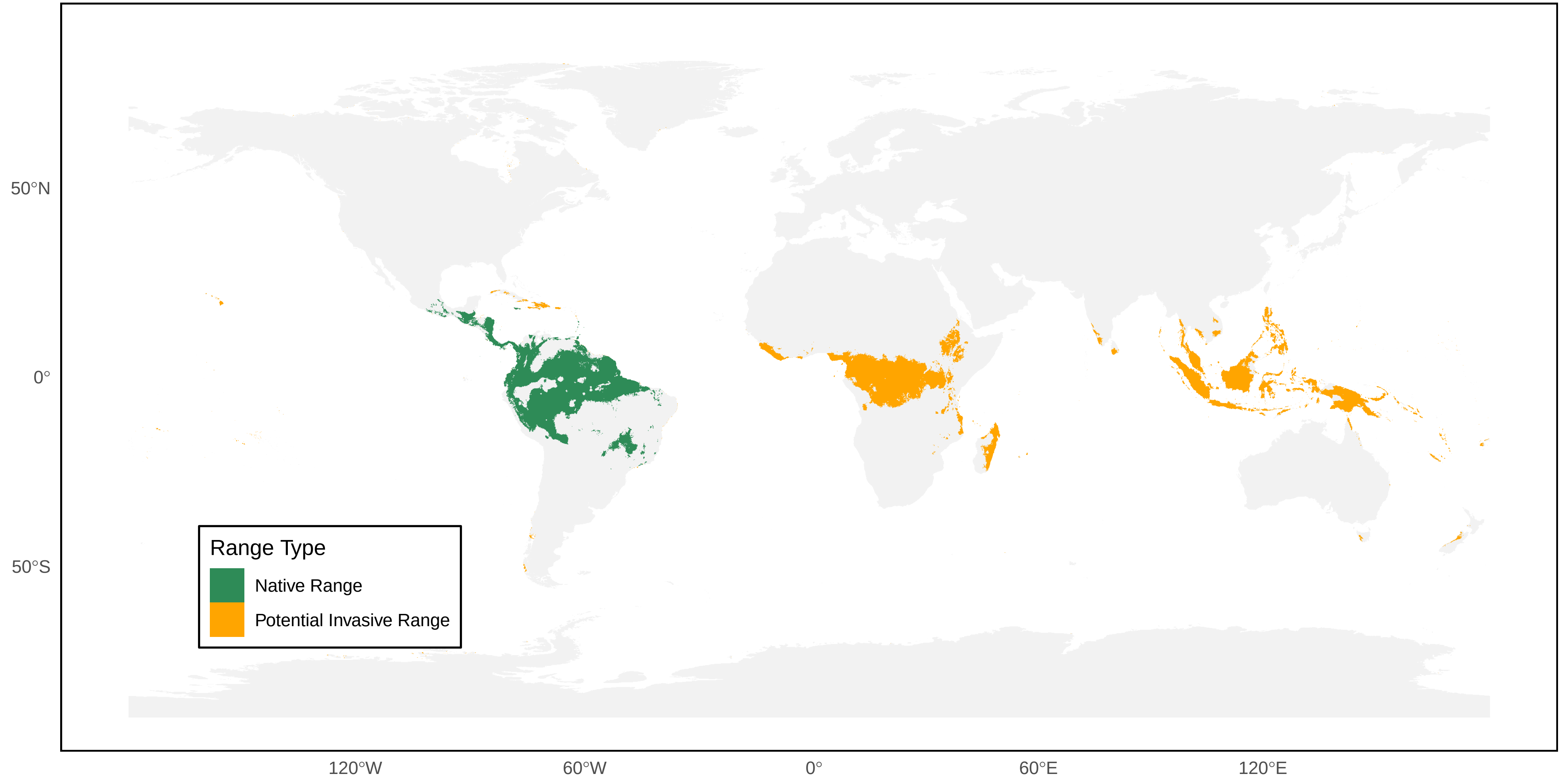

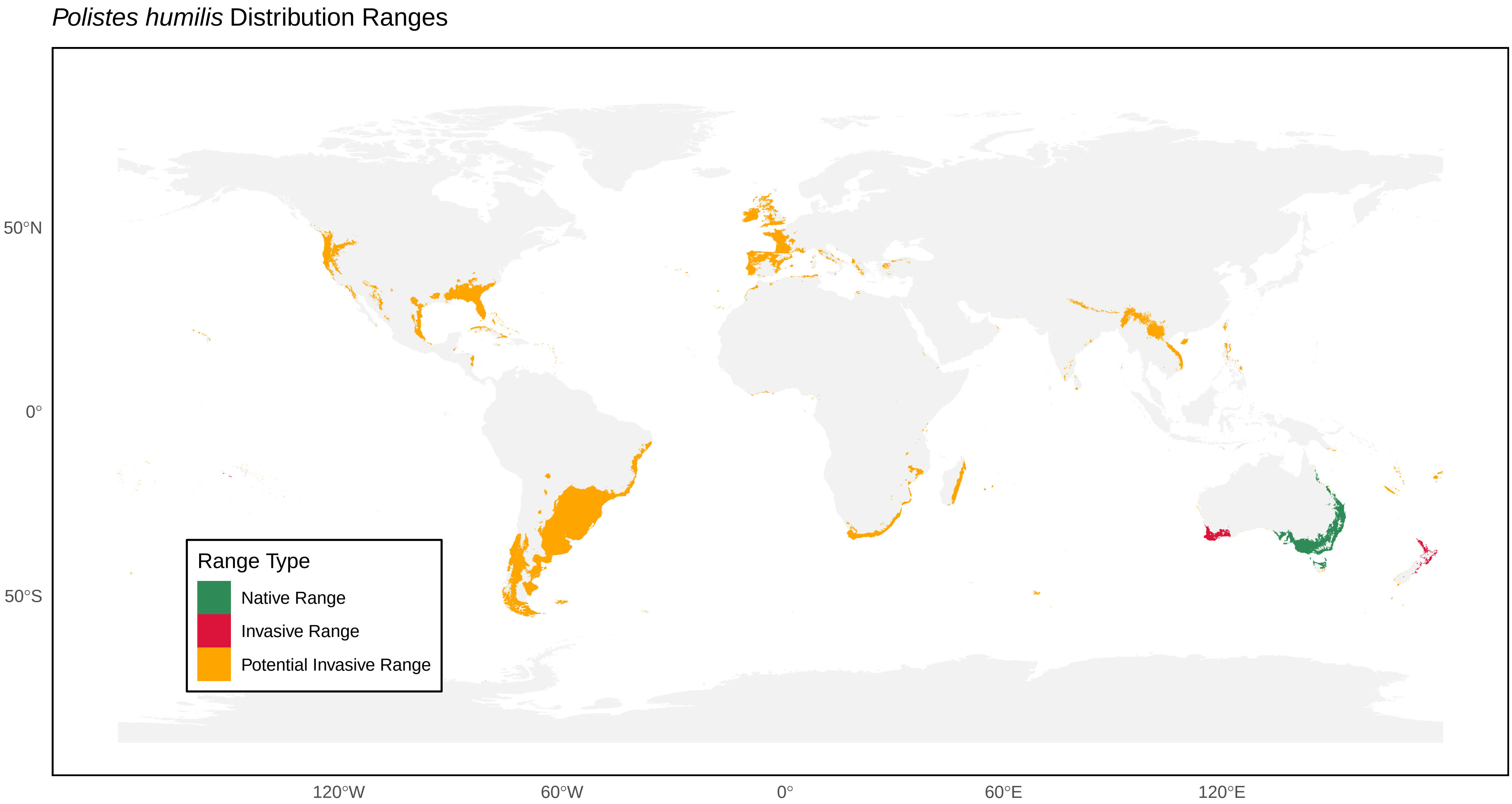

*Polistes humilis* Distribution Ranges
Range Type
Native Range
Invasive Range
Potential Invasive Range
50°N
0°
50°S
120°W
60°W
0°
60°E
120°E

*Polistes incertus* Distribution Ranges

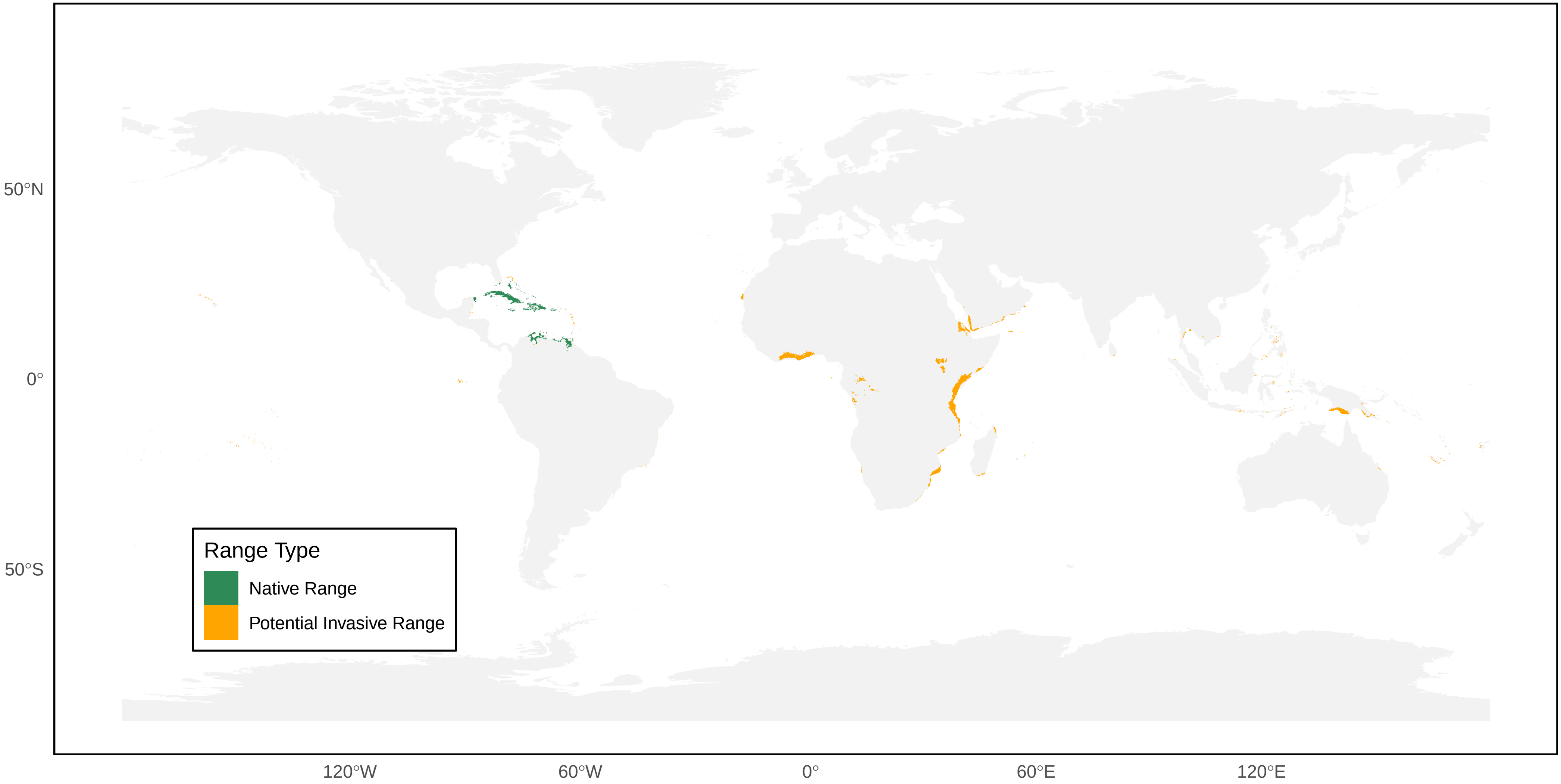

*Polistes infuscatus* Distribution Ranges

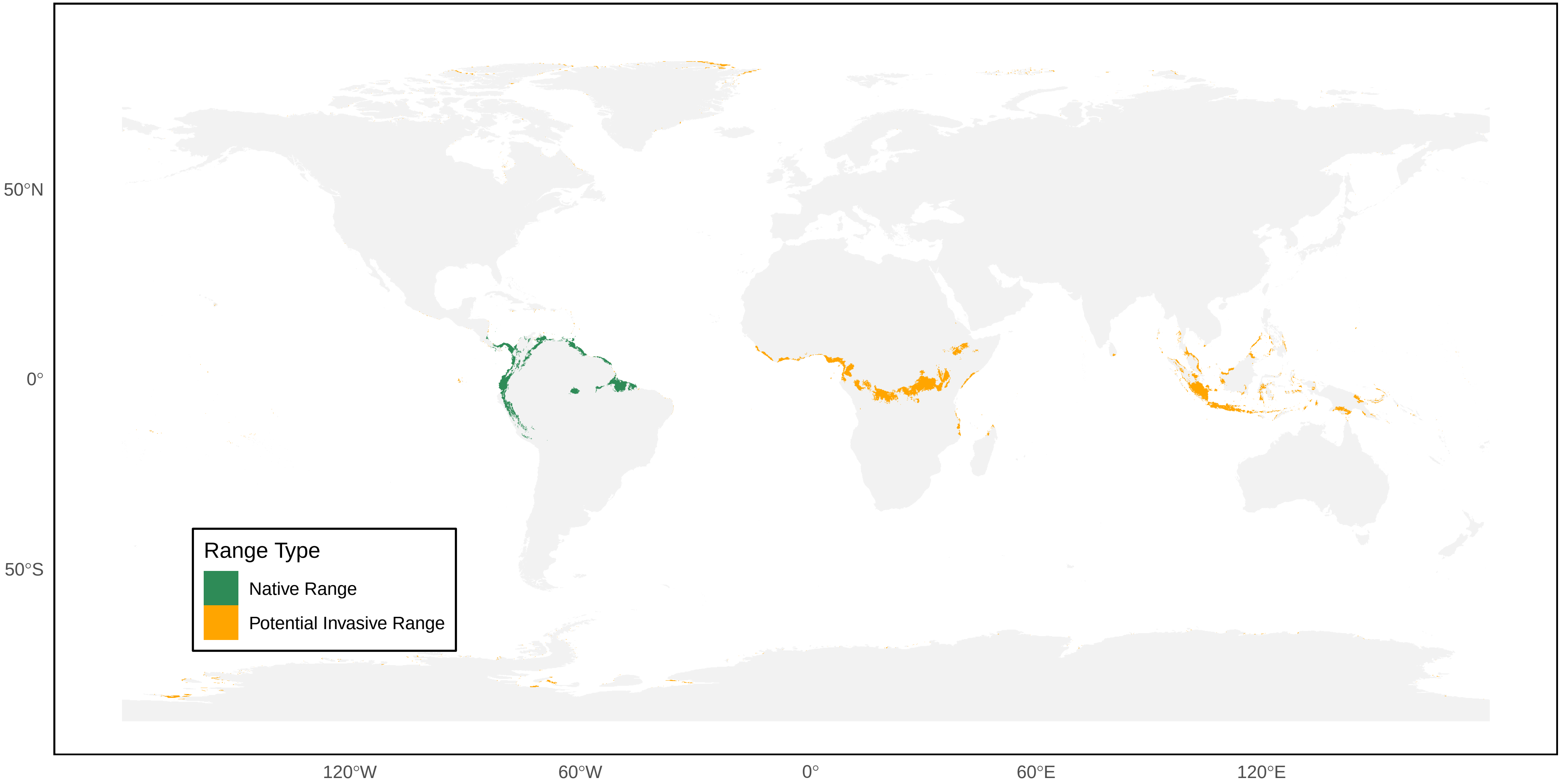

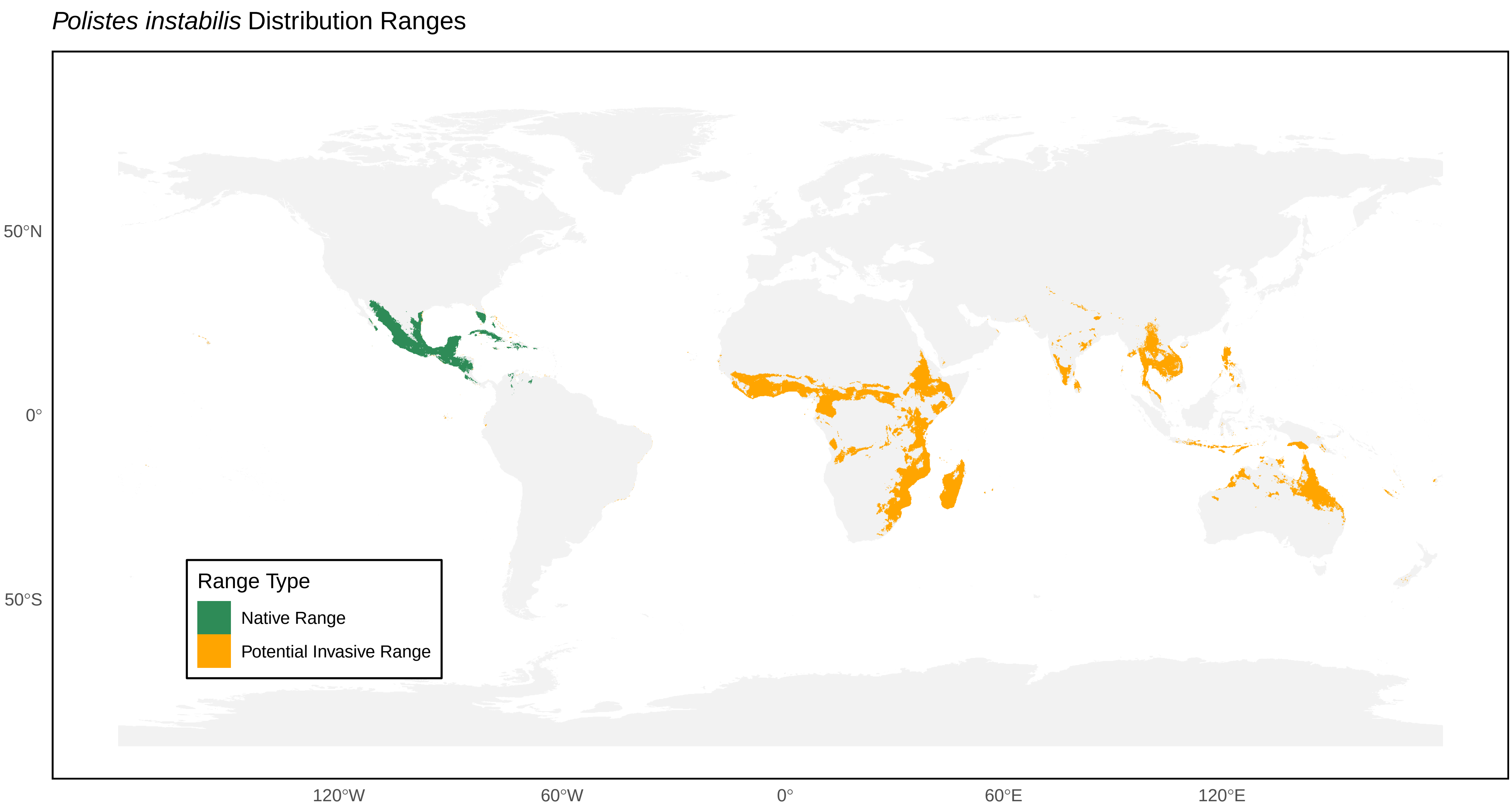

*Polistes instabilis* Distribution Ranges
50°N
0°
50°S
120°W
60°W
0°
60°E
120°E
Range Type
Native Range
Potential Invasive Range

*Polistes japonicus* Distribution Ranges

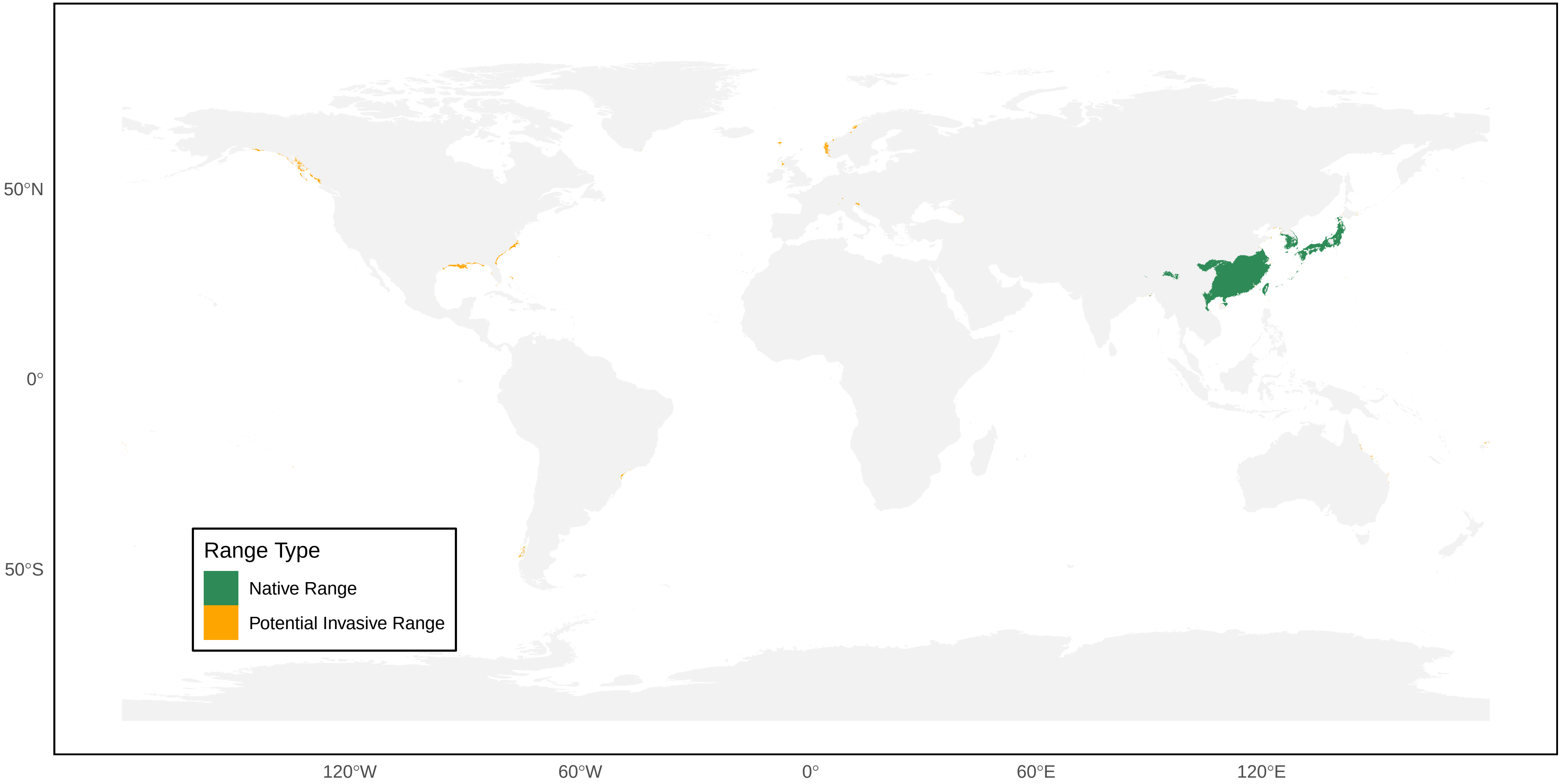

*Polistes jokahamae* Distribution Ranges

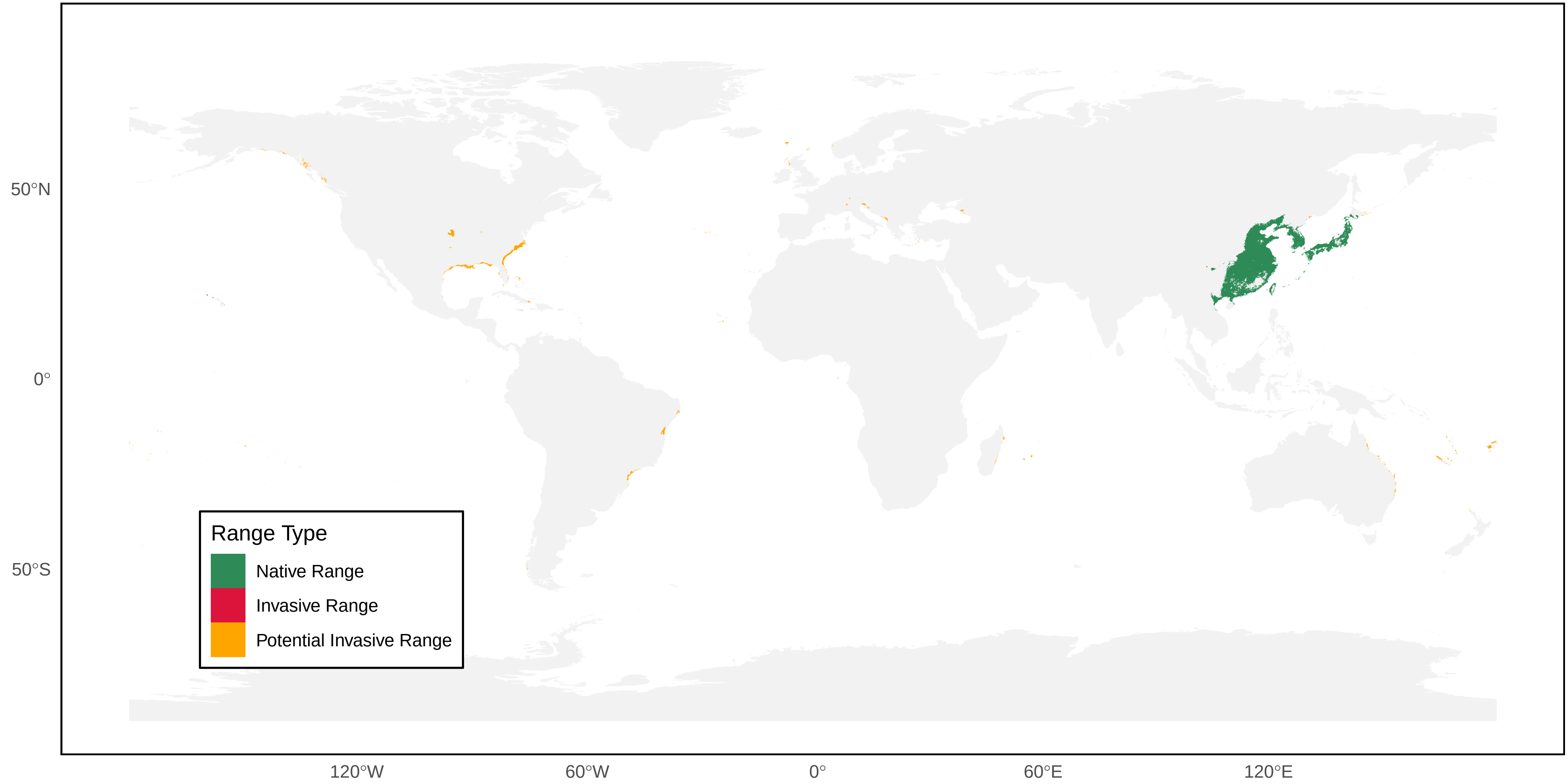

*Polistes kaibabensis* Distribution Ranges

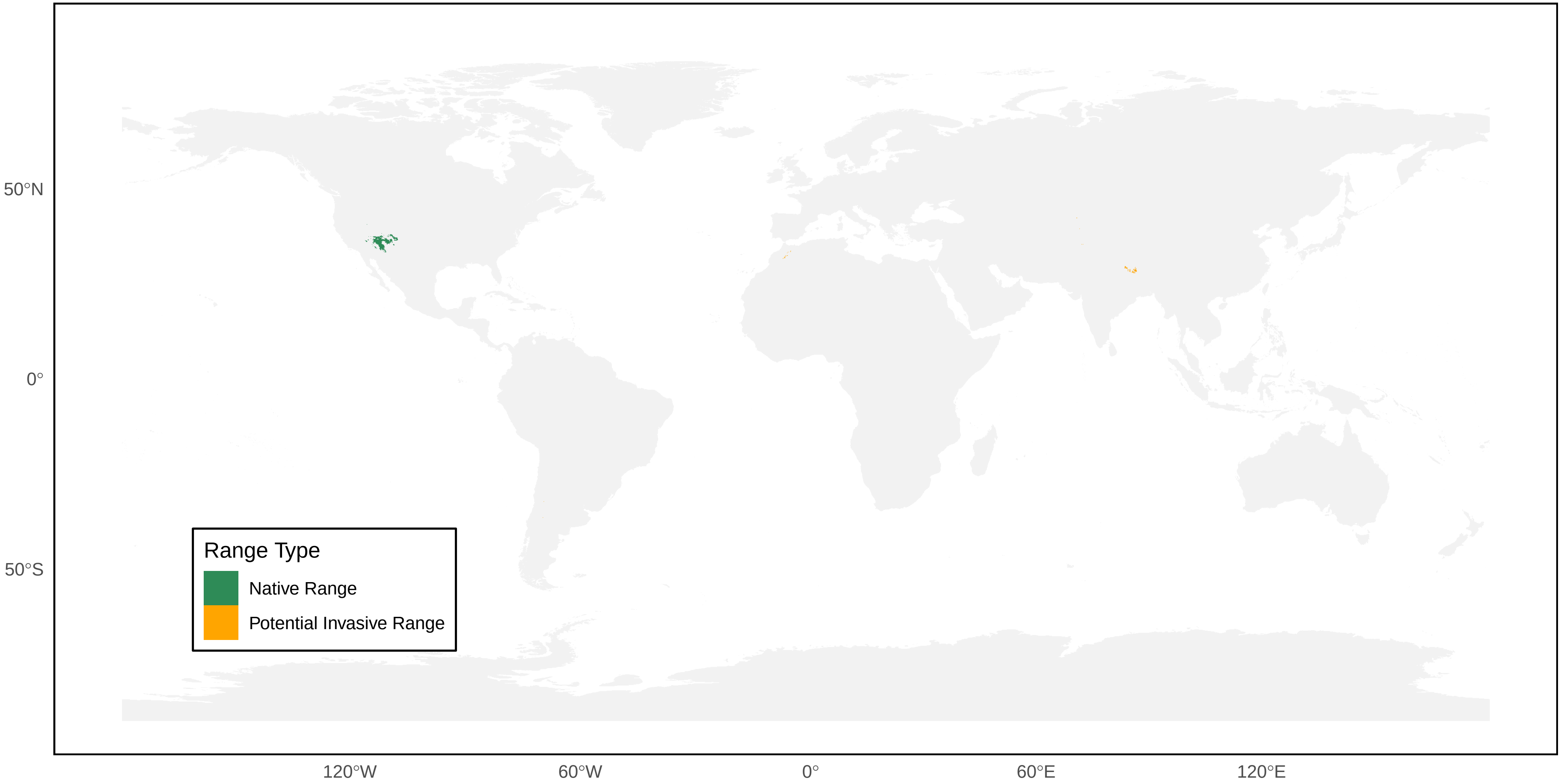

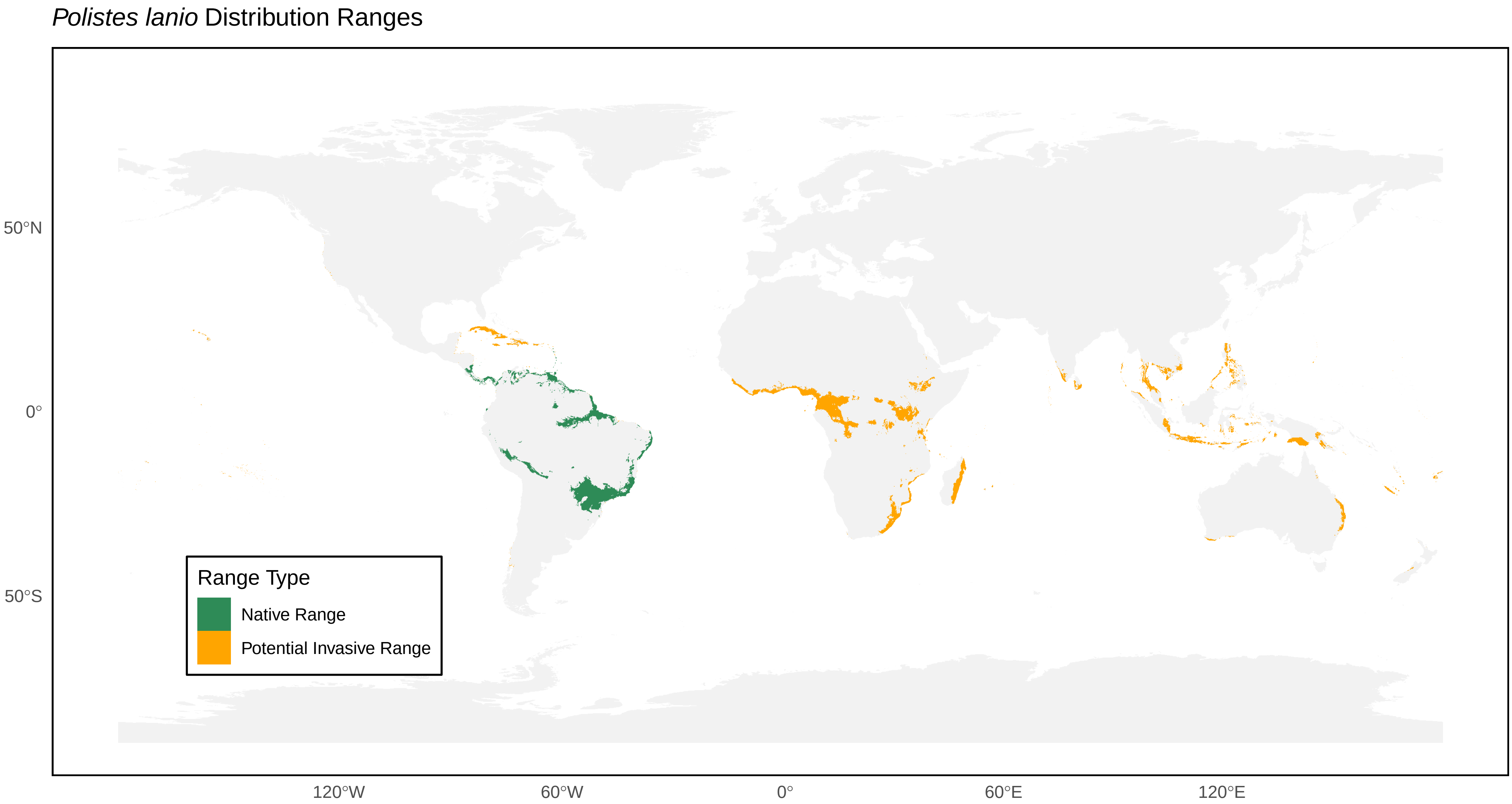
*Polistes lanio* Distribution Ranges
50°N
0°
50°S
120°W
60°W
0°
60°E
120°E
Range Type
Native Range
Potential Invasive Range

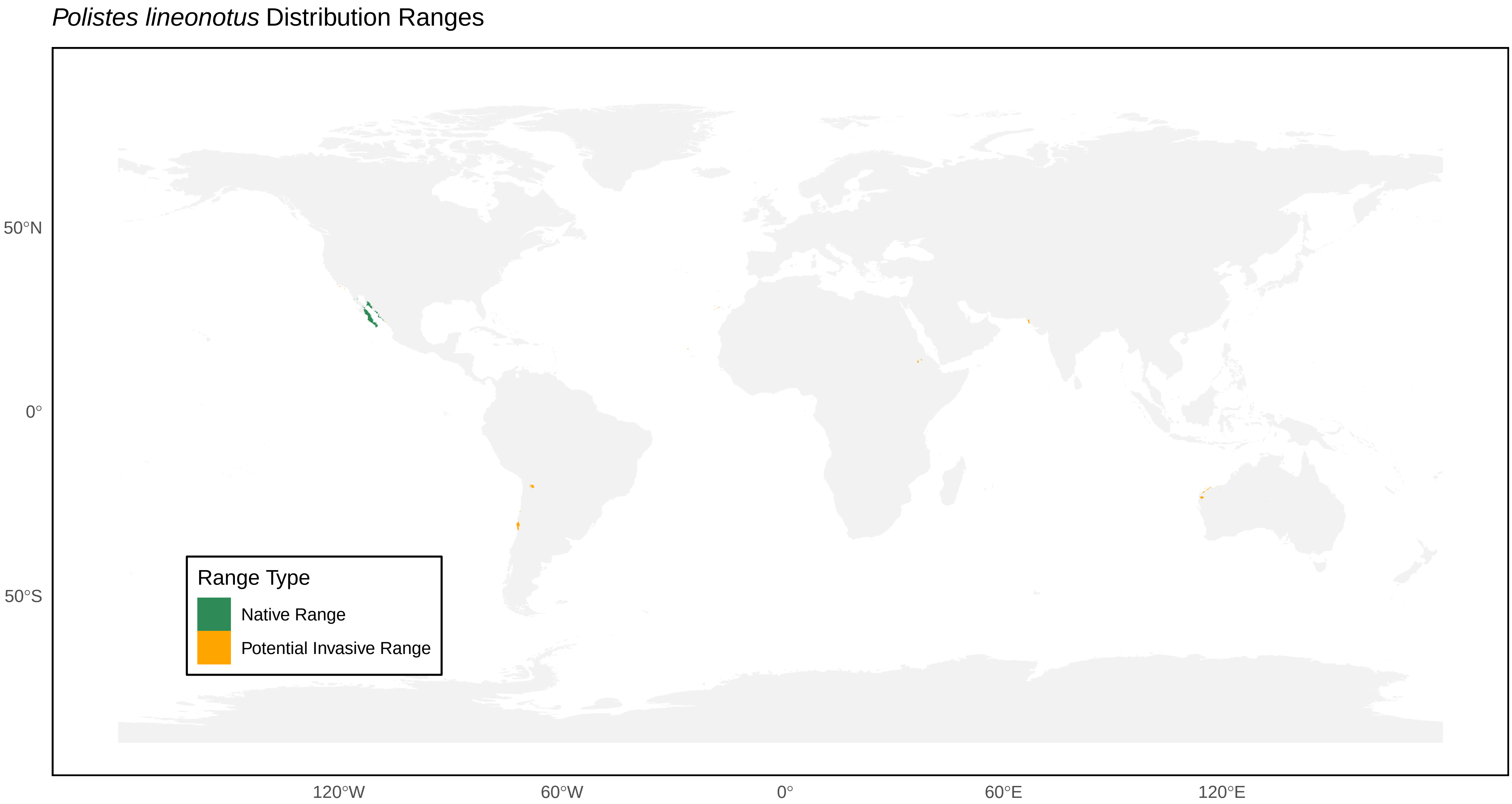

Polistes lineonotus Distribution Ranges
50°N
0°
50°S
120°W
60°W
0°
60°E
120°E
Range Type
Native Range
Potential Invasive Range

*Polistes major* Distribution Ranges

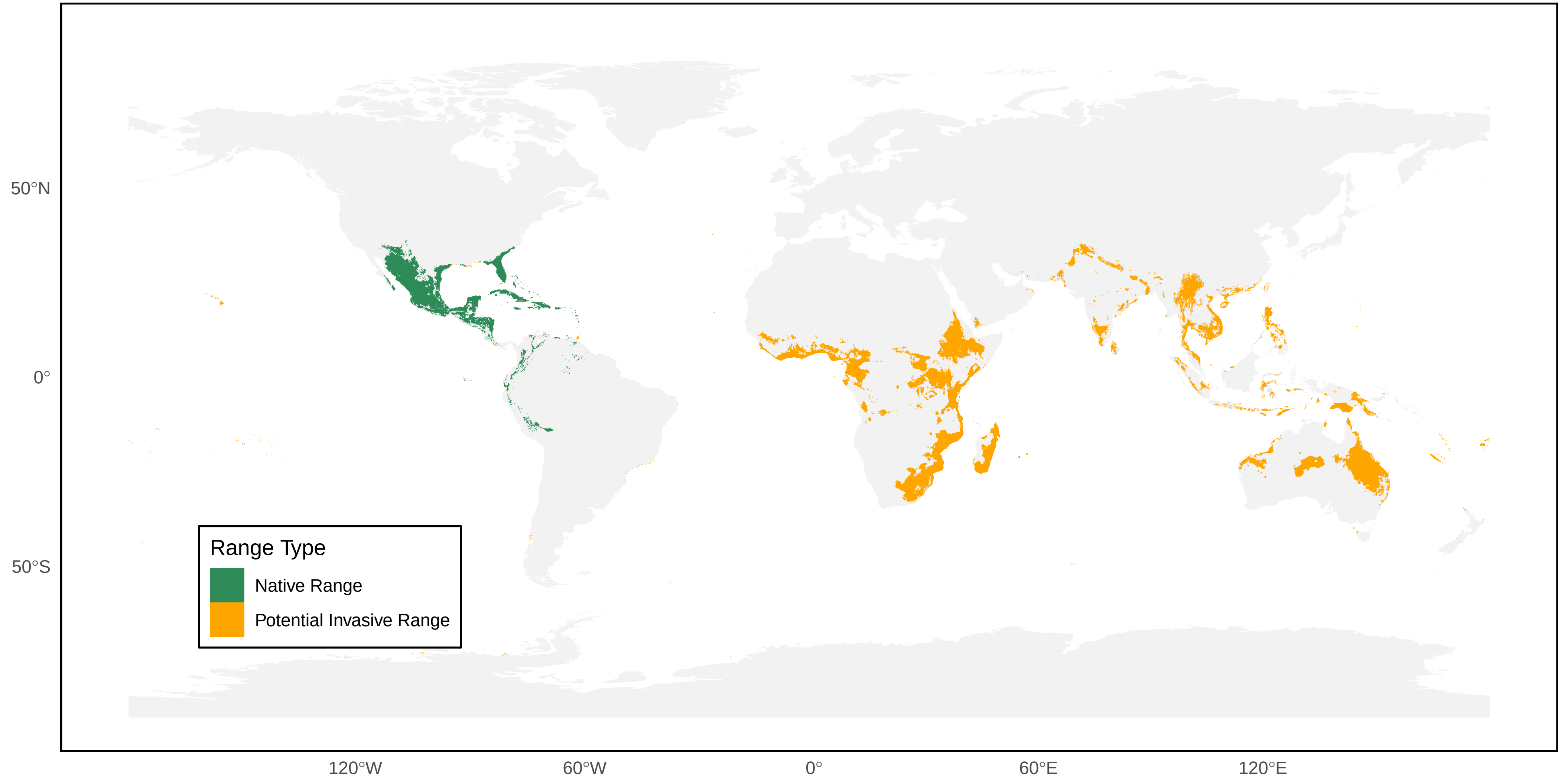

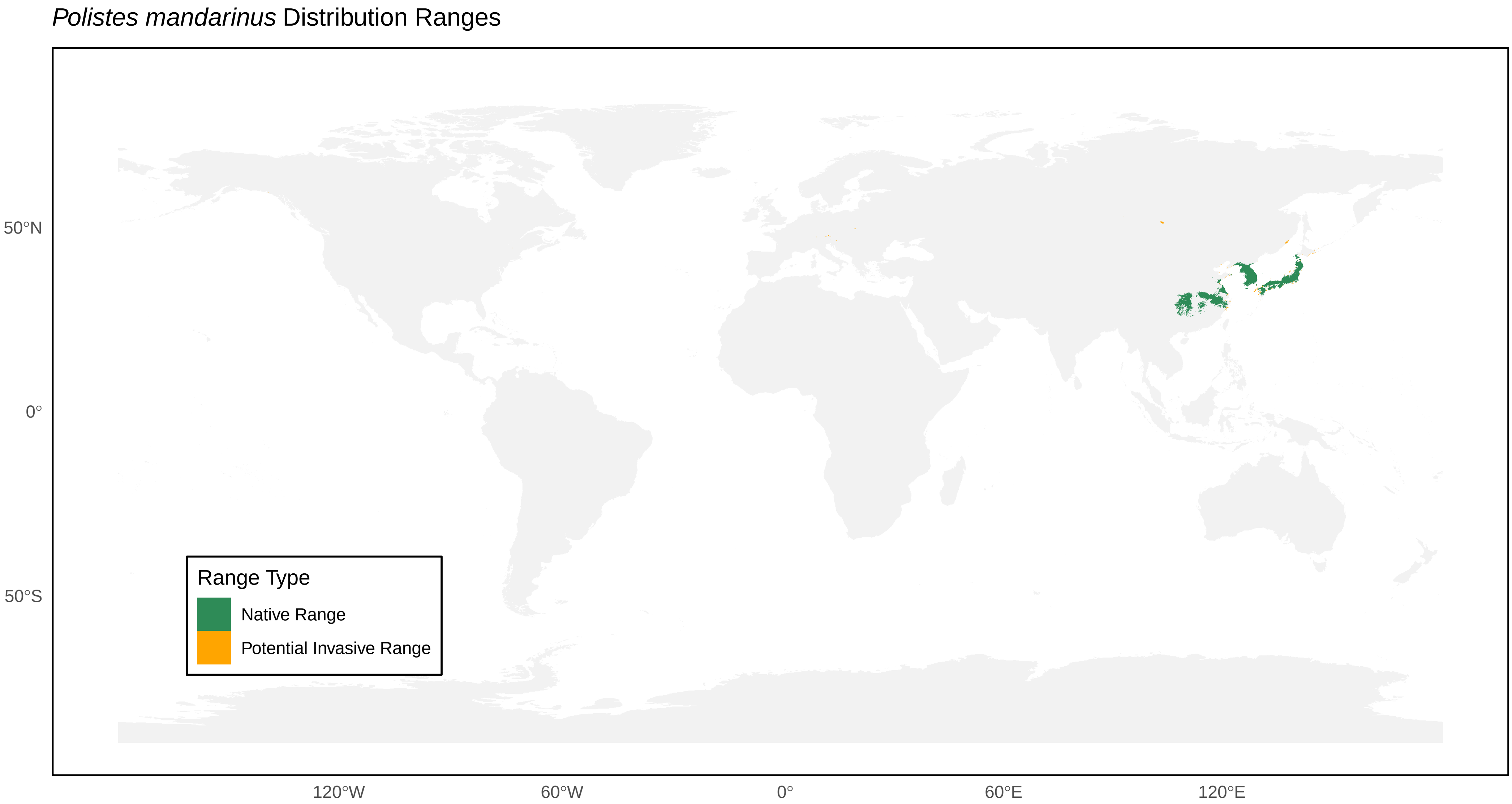
*Polistes mandarinus* Distribution Ranges
50°N
0°
50°S
120°W
60°W
0°
60°E
120°E
Range Type
Native Range
Potential Invasive Range

*Polistes marginalis* Distribution Ranges

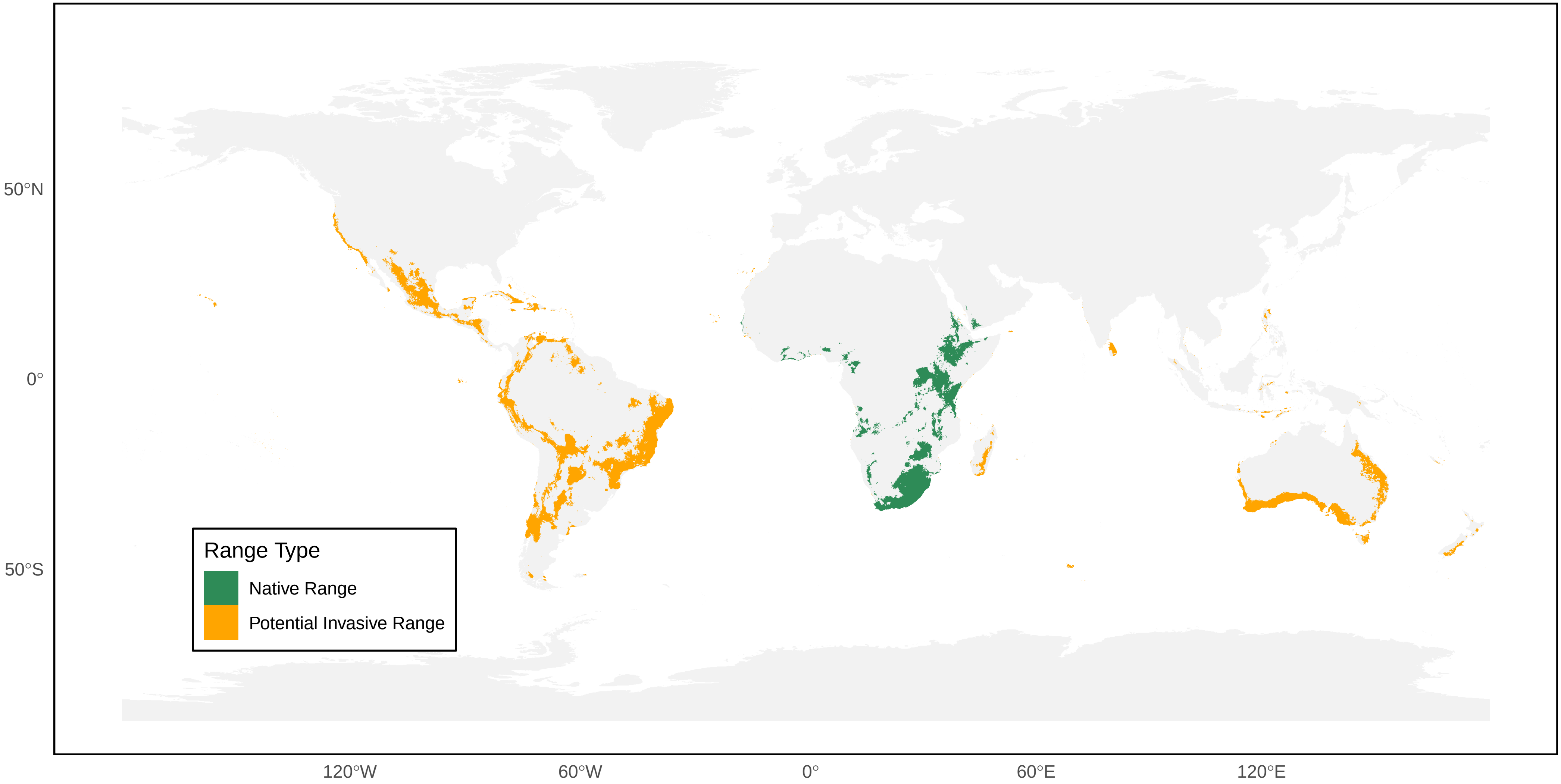

*Polistes metricus* Distribution Ranges

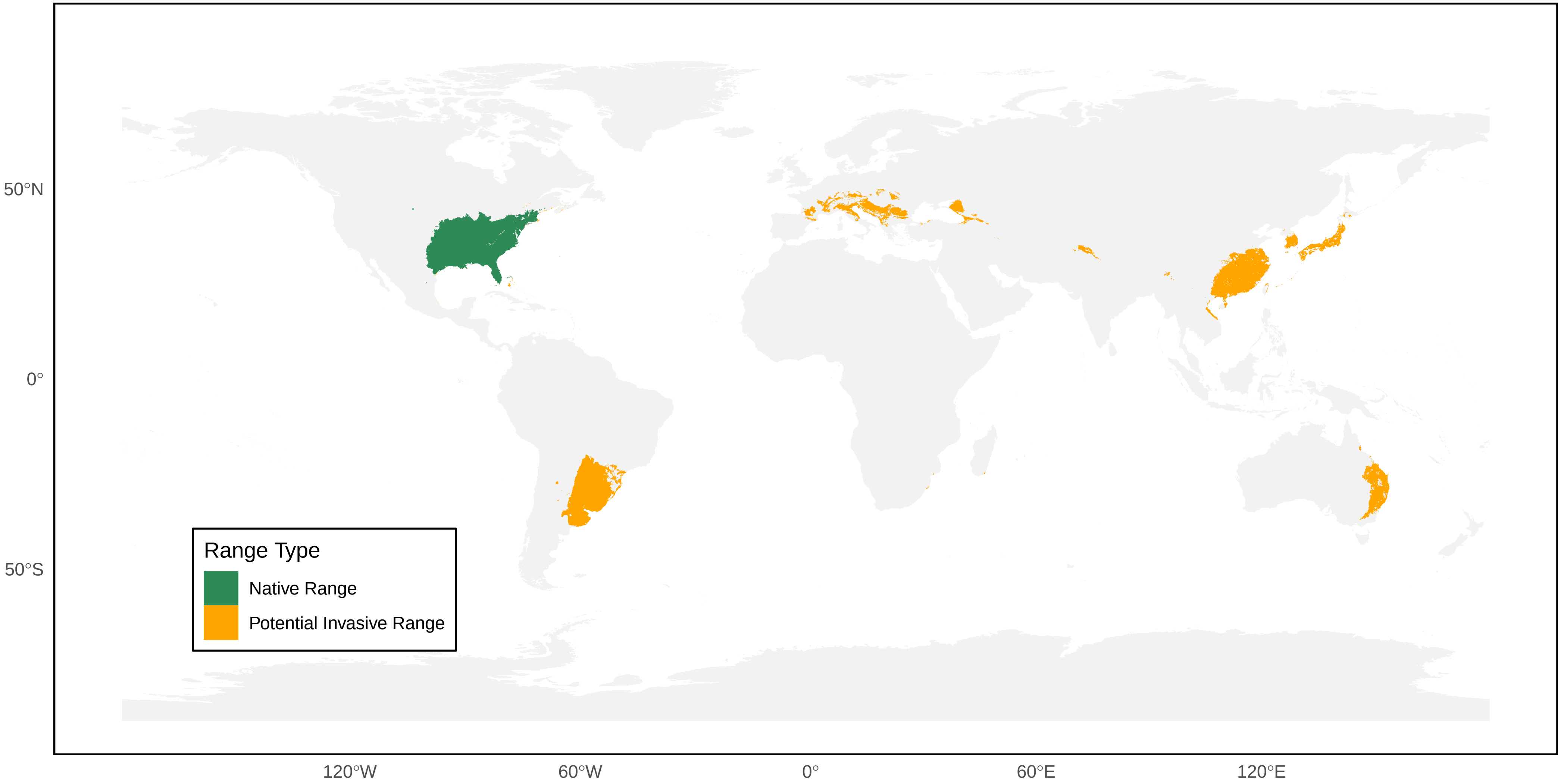

*Polistes mexicanus* Distribution Ranges

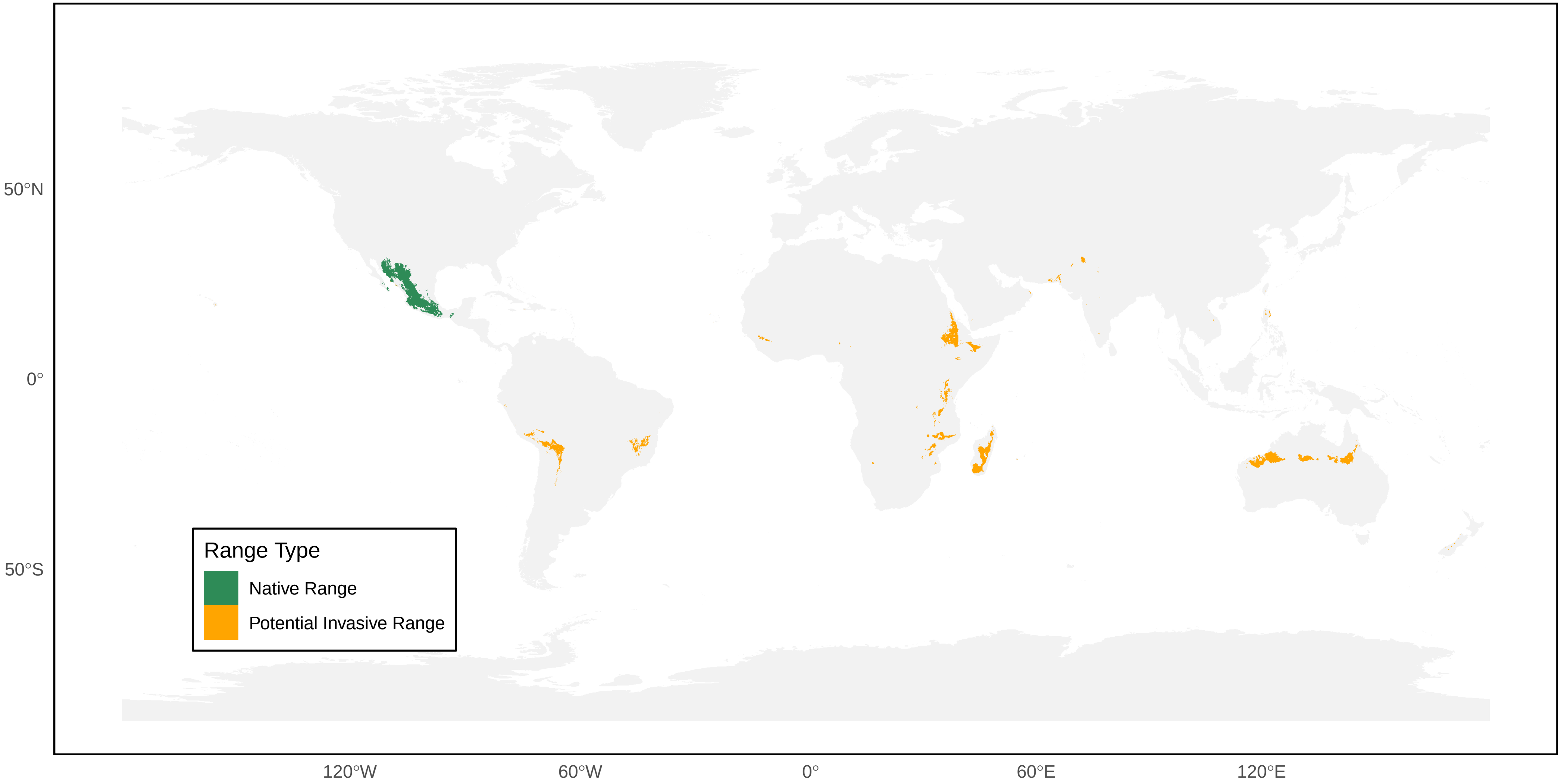

*Polistes minor* Distribution Ranges

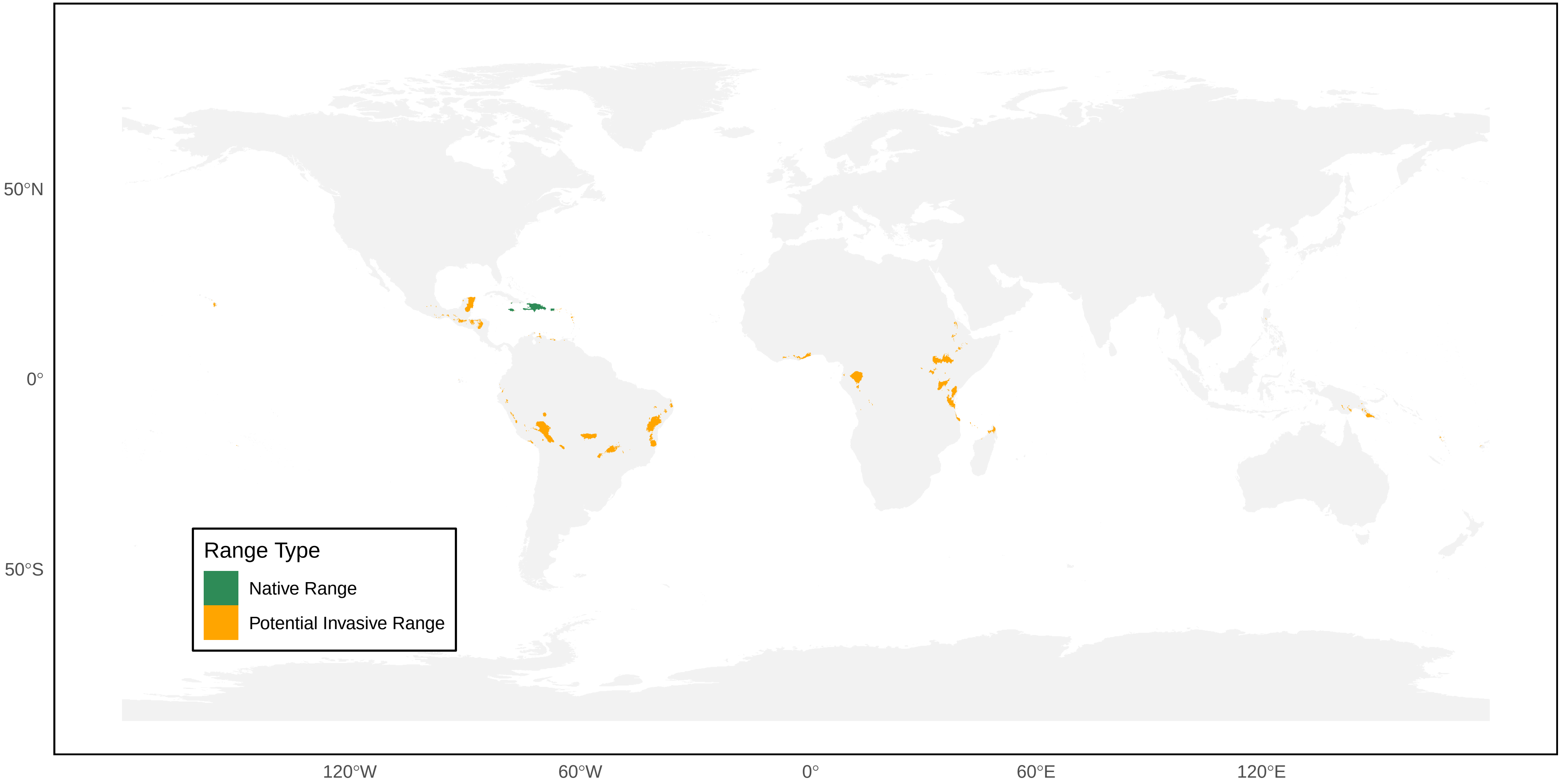

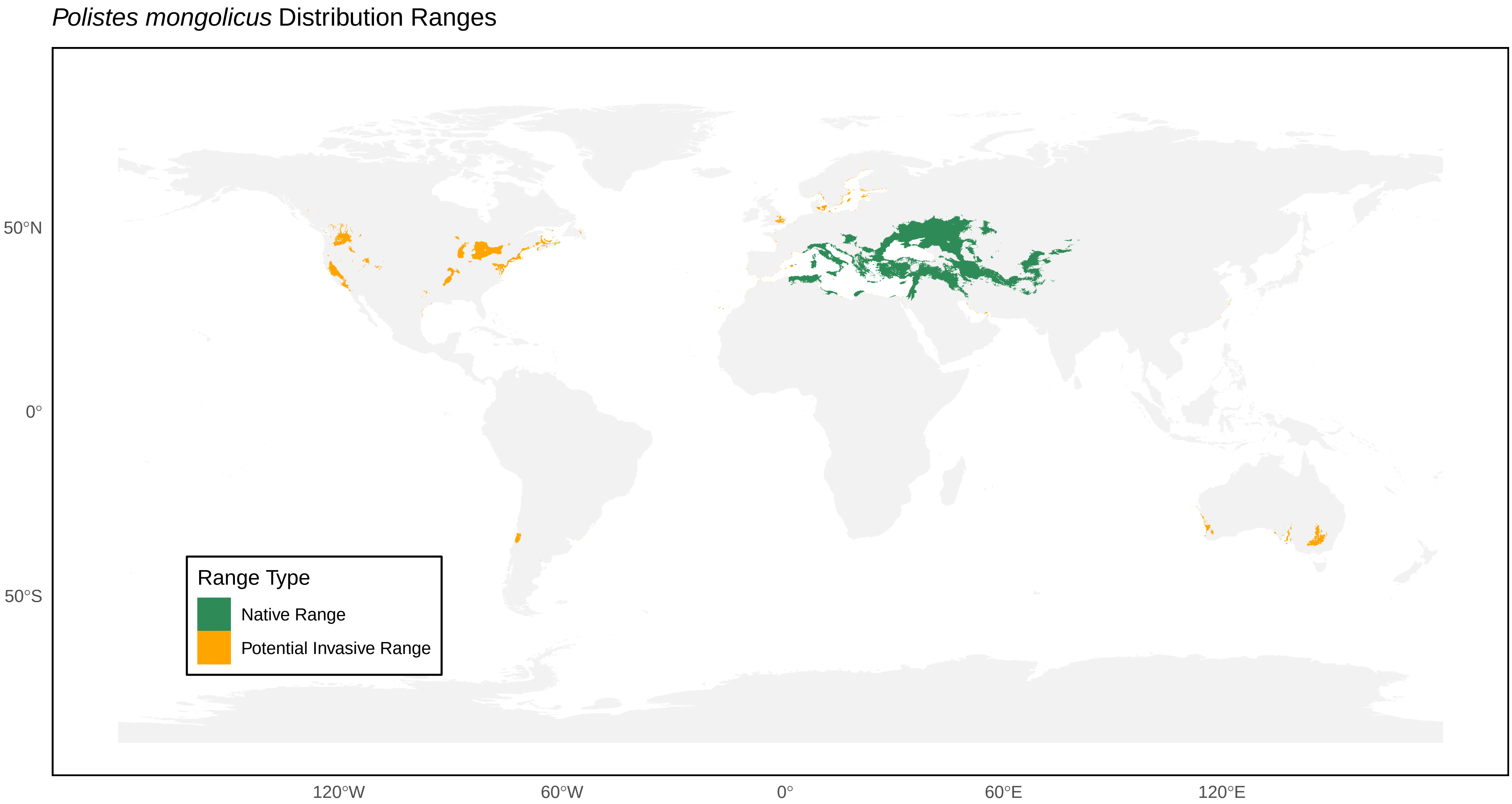

*Polistes mongolicus* Distribution Ranges
50°N
0°
50°S
120°W
60°W
0°
60°E
120°E
Range Type
Native Range
Potential Invasive Range

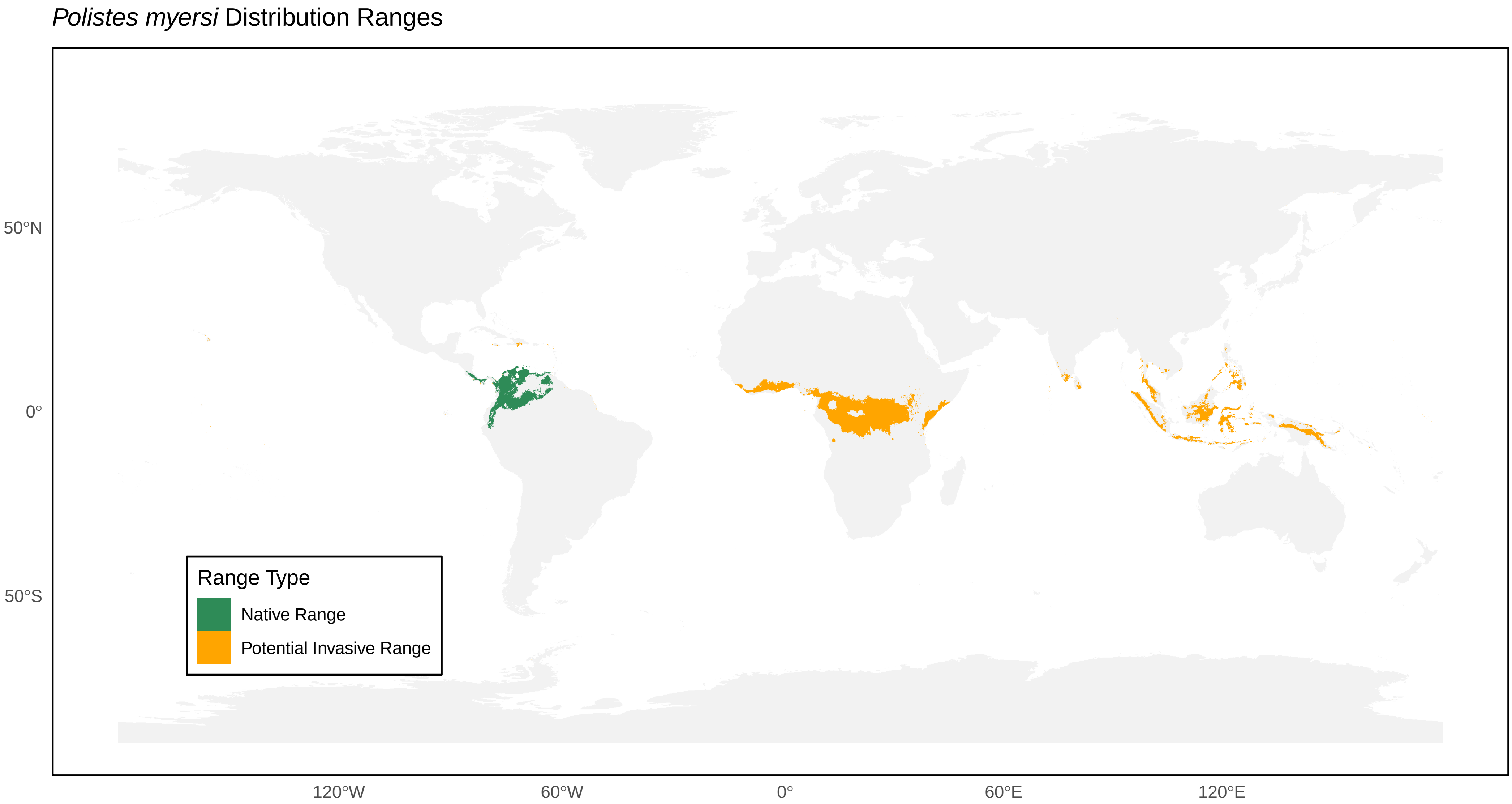

*Polistes myersi* Distribution Ranges
50°N
0°
50°S
120°W
60°W
0°
60°E
120°E
Range Type
Native Range
Potential Invasive Range

*Polistes nimpha* Distribution Ranges

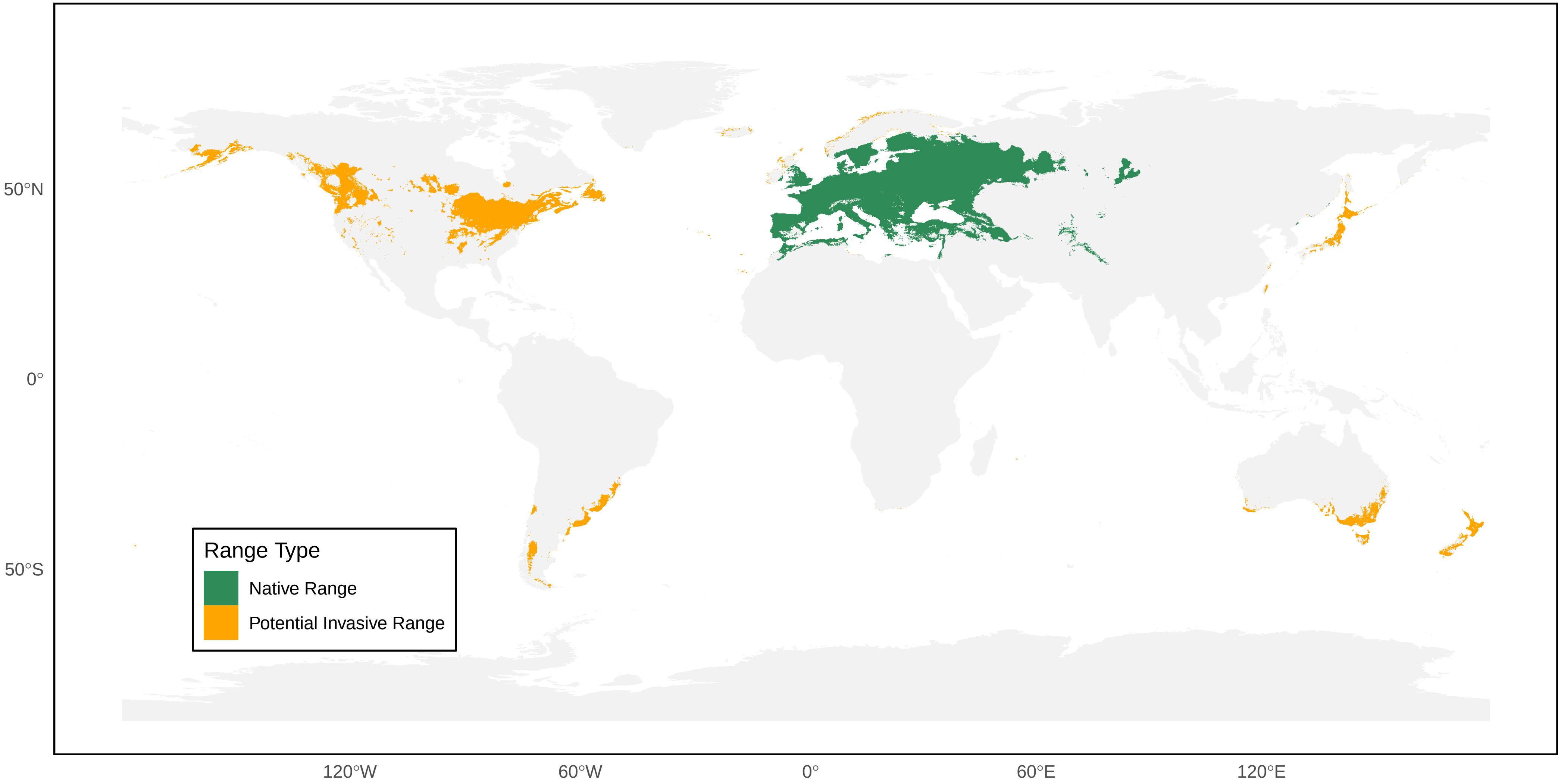

*Polistes nipponensis* Distribution Ranges

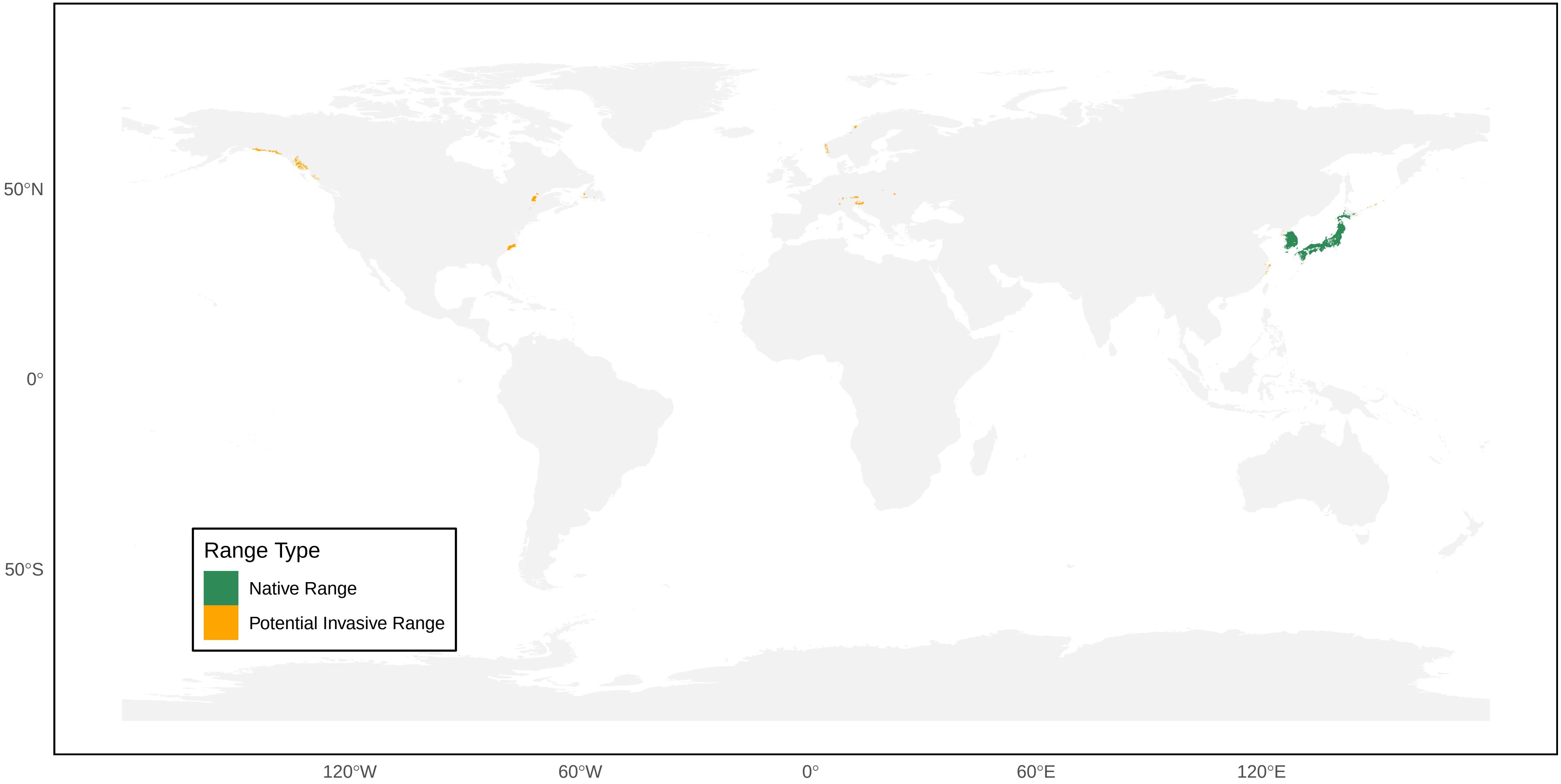

*Polistes occipitalis* Distribution Ranges

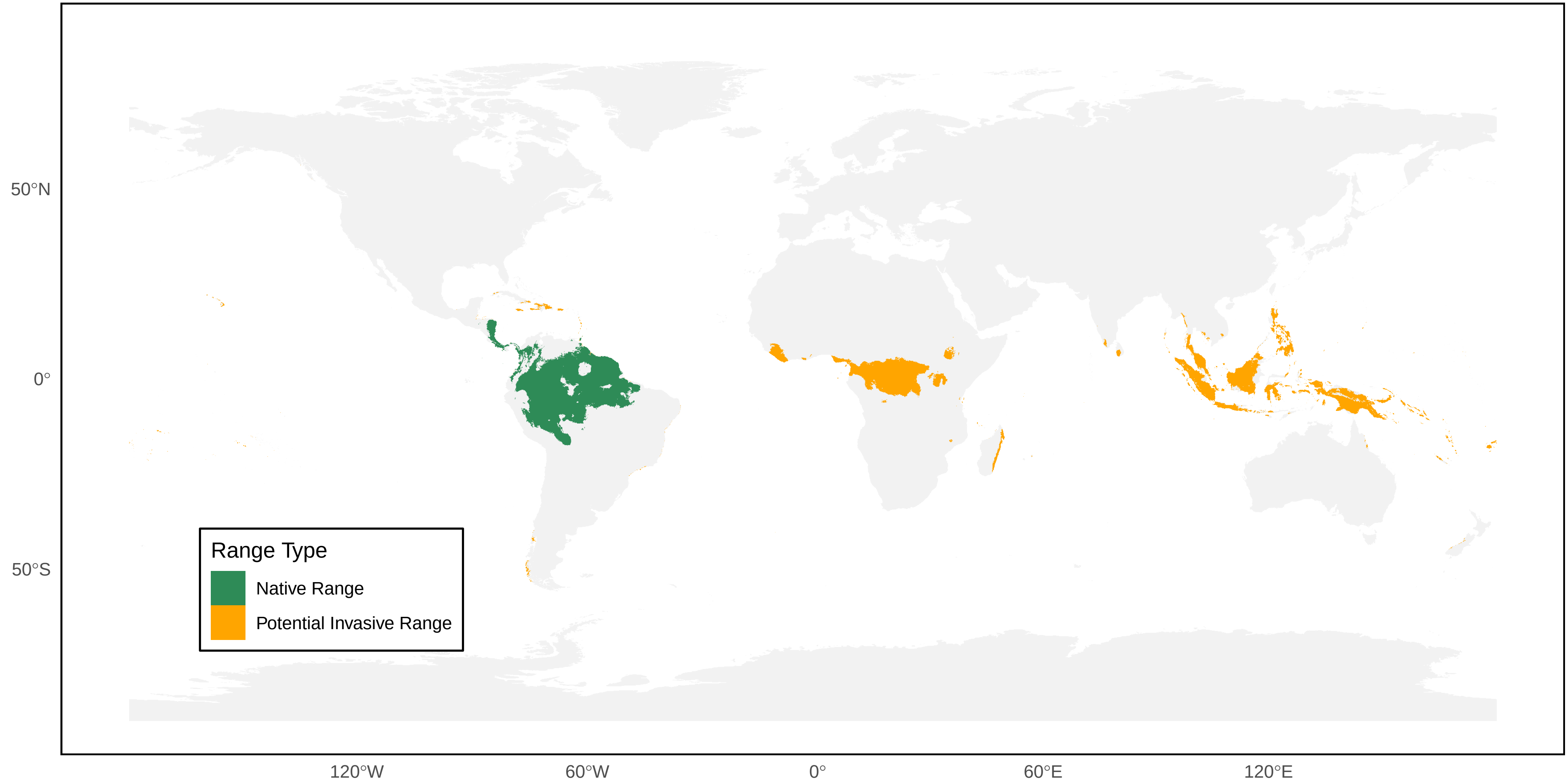

*Polistes oculatus* Distribution Ranges

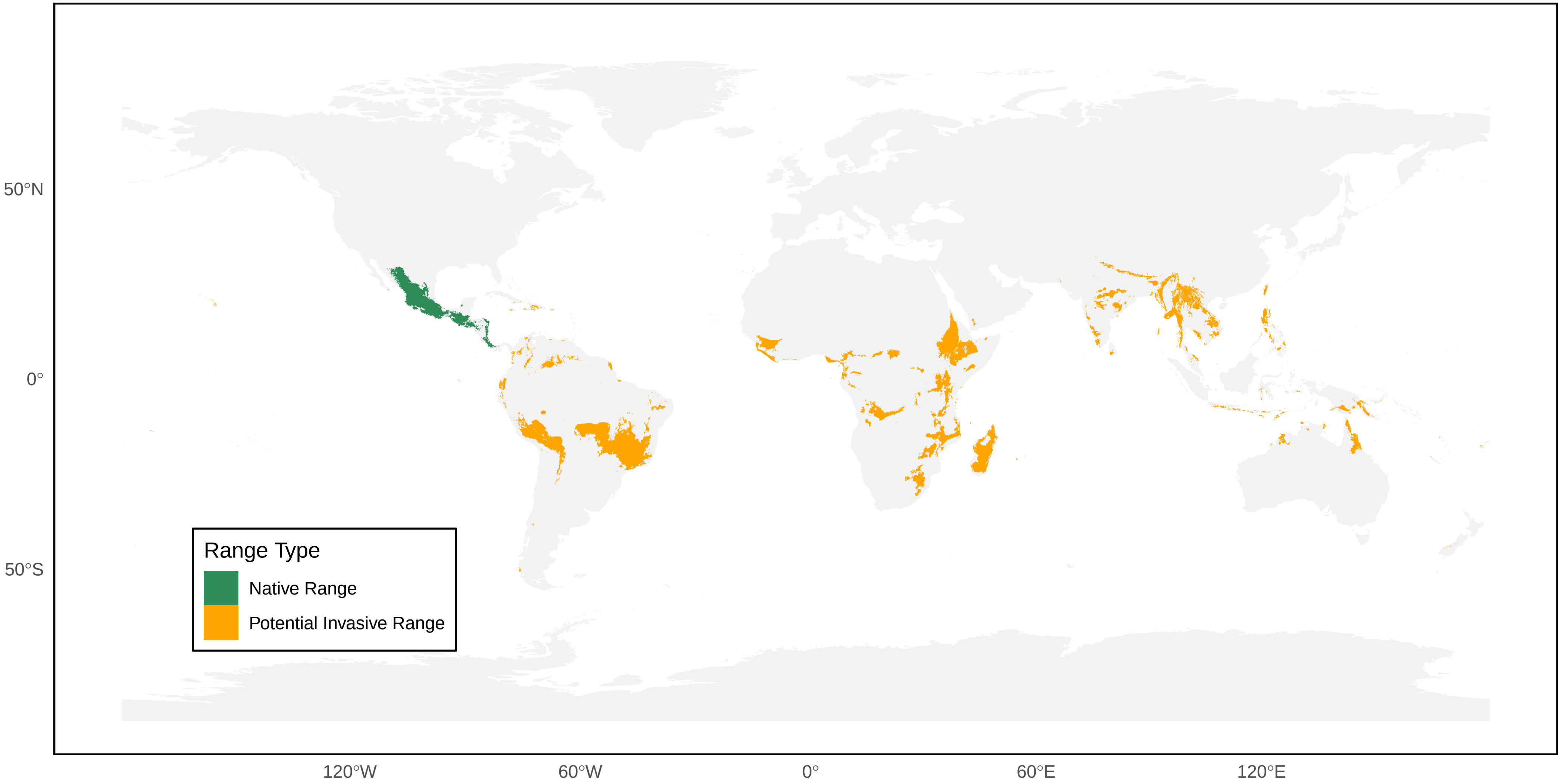

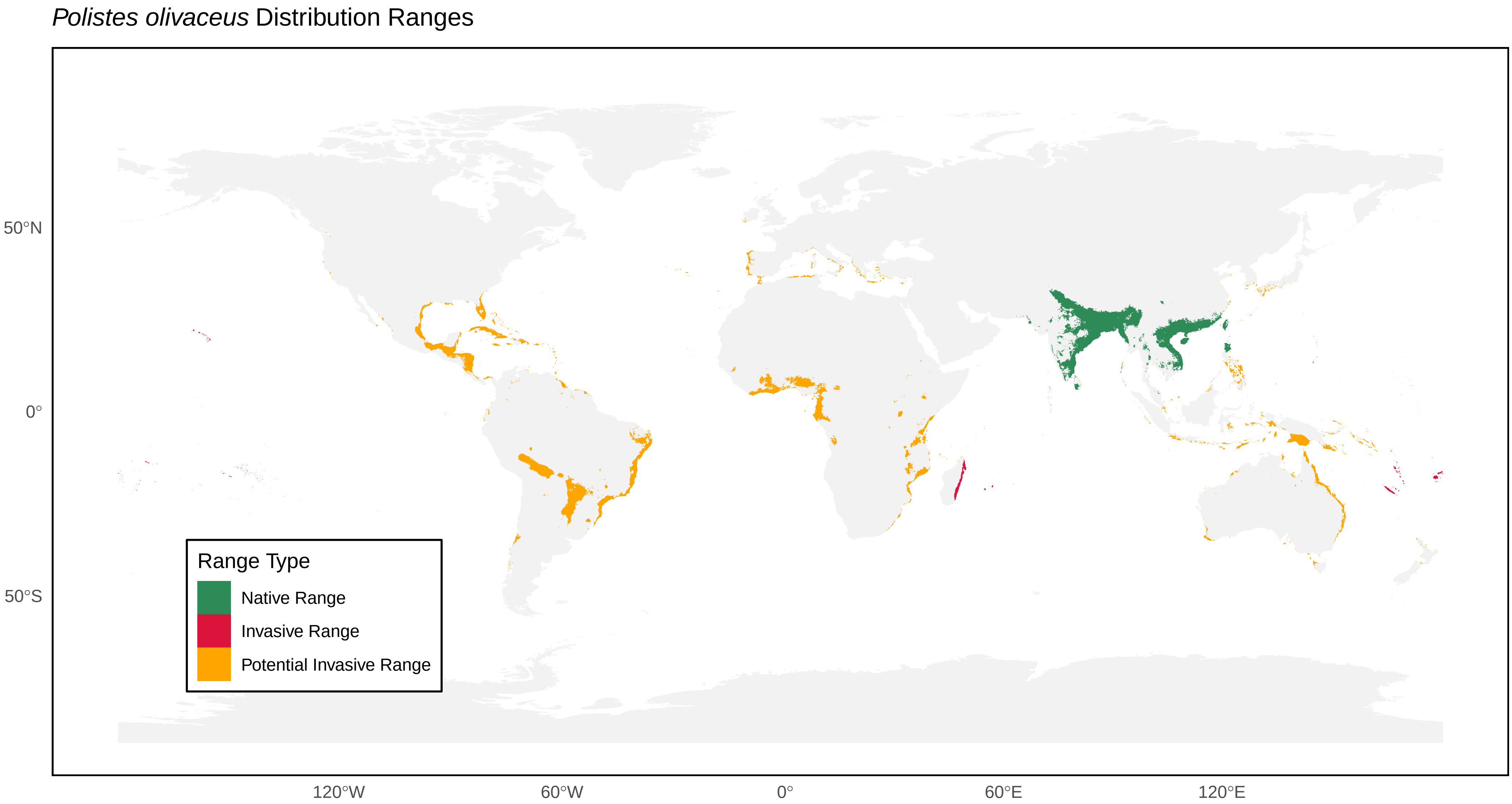
*Polistes olivaceus* Distribution Ranges
50°N
0°
50°S
120°W
60°W
0°
60°E
120°E
Range Type
Native Range
Invasive Range
Potential Invasive Range

*Polistes pacificus* Distribution Ranges

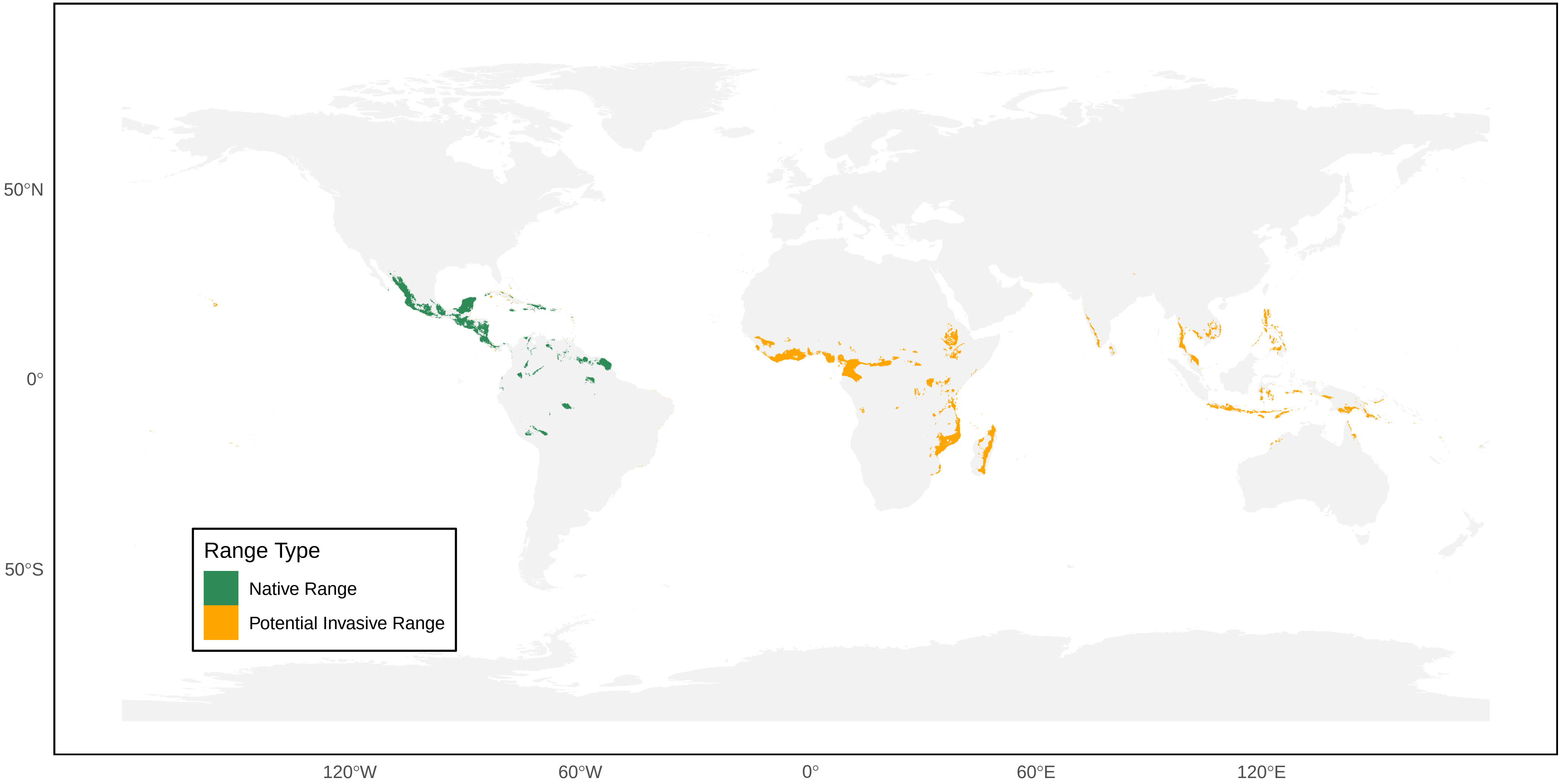

*Polistes palmarum* Distribution Ranges

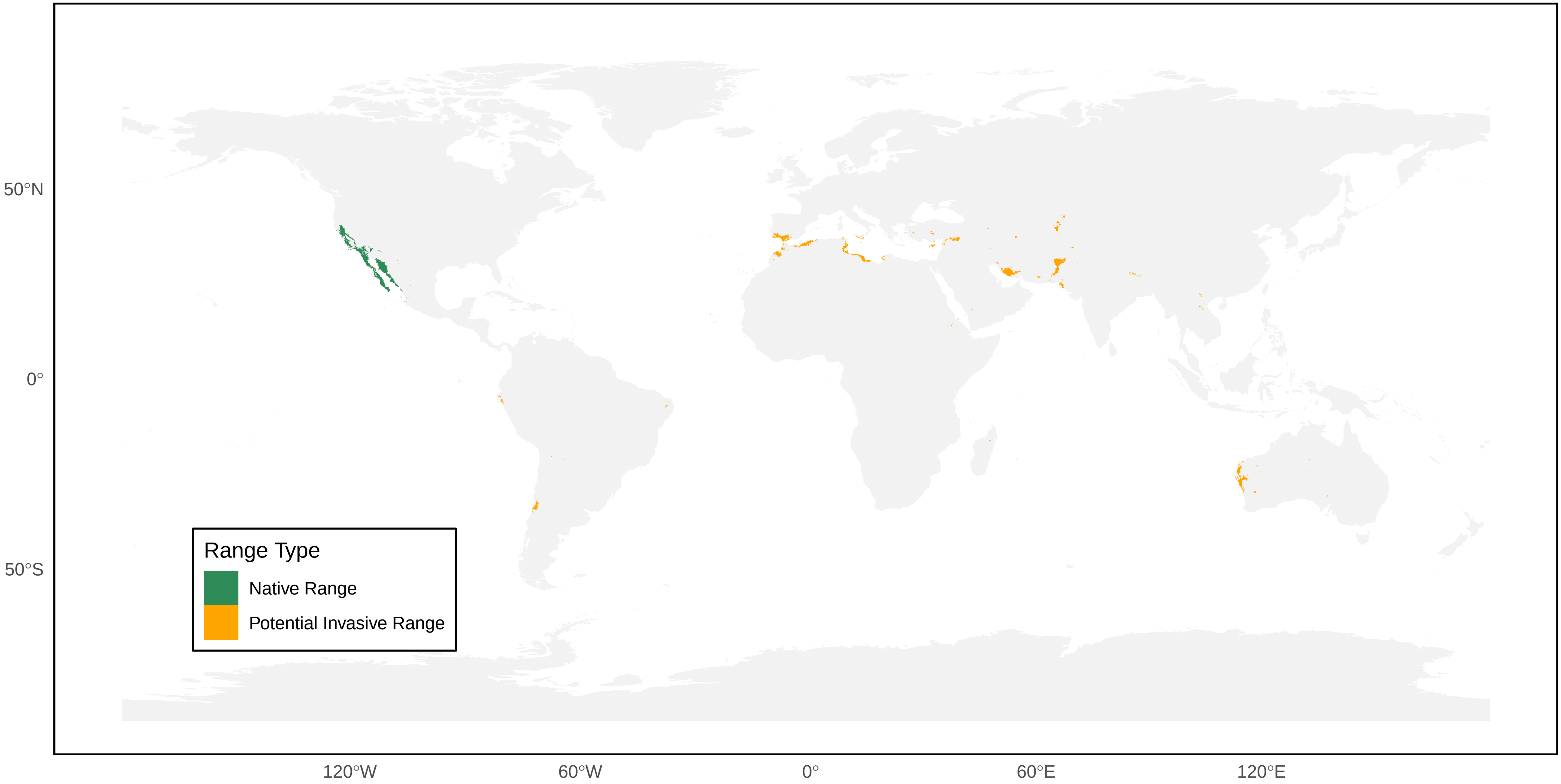

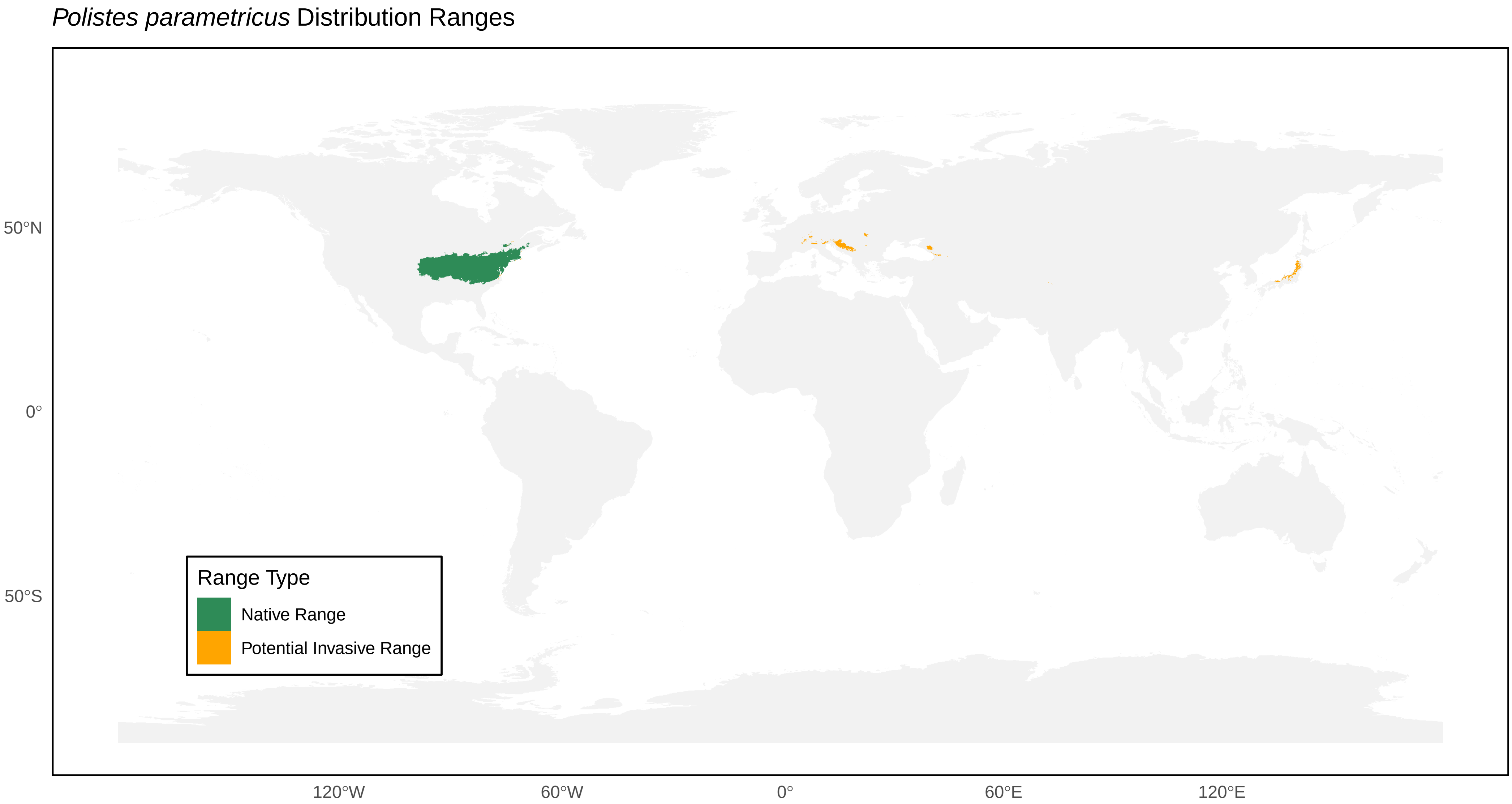

*Polistes parametricus* Distribution Ranges
50°N
0°
50°S
120°W
60°W
0°
60°E
120°E
Range Type
Native Range
Potential Invasive Range

*Polistes peruvianus* Distribution Ranges

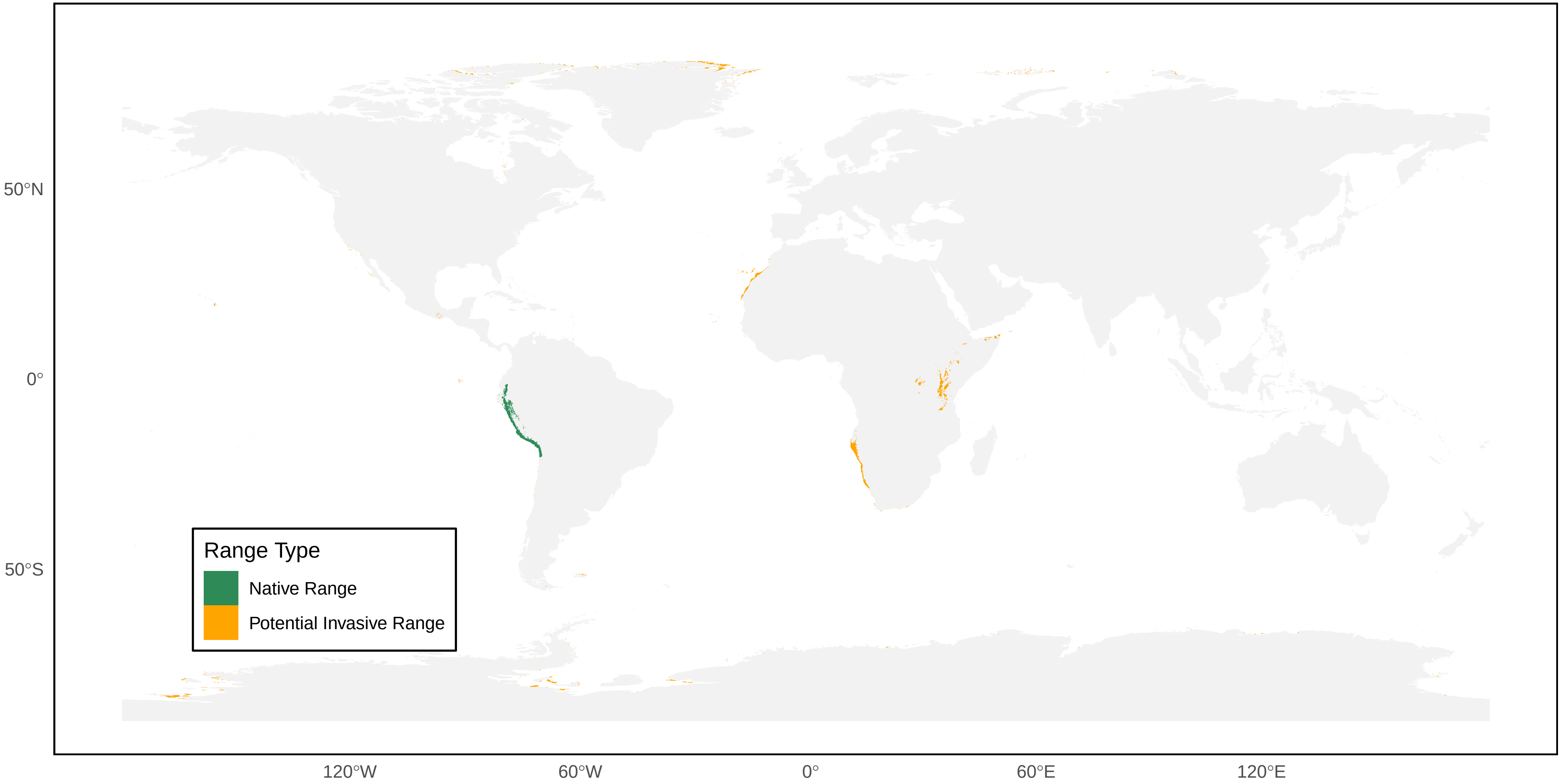

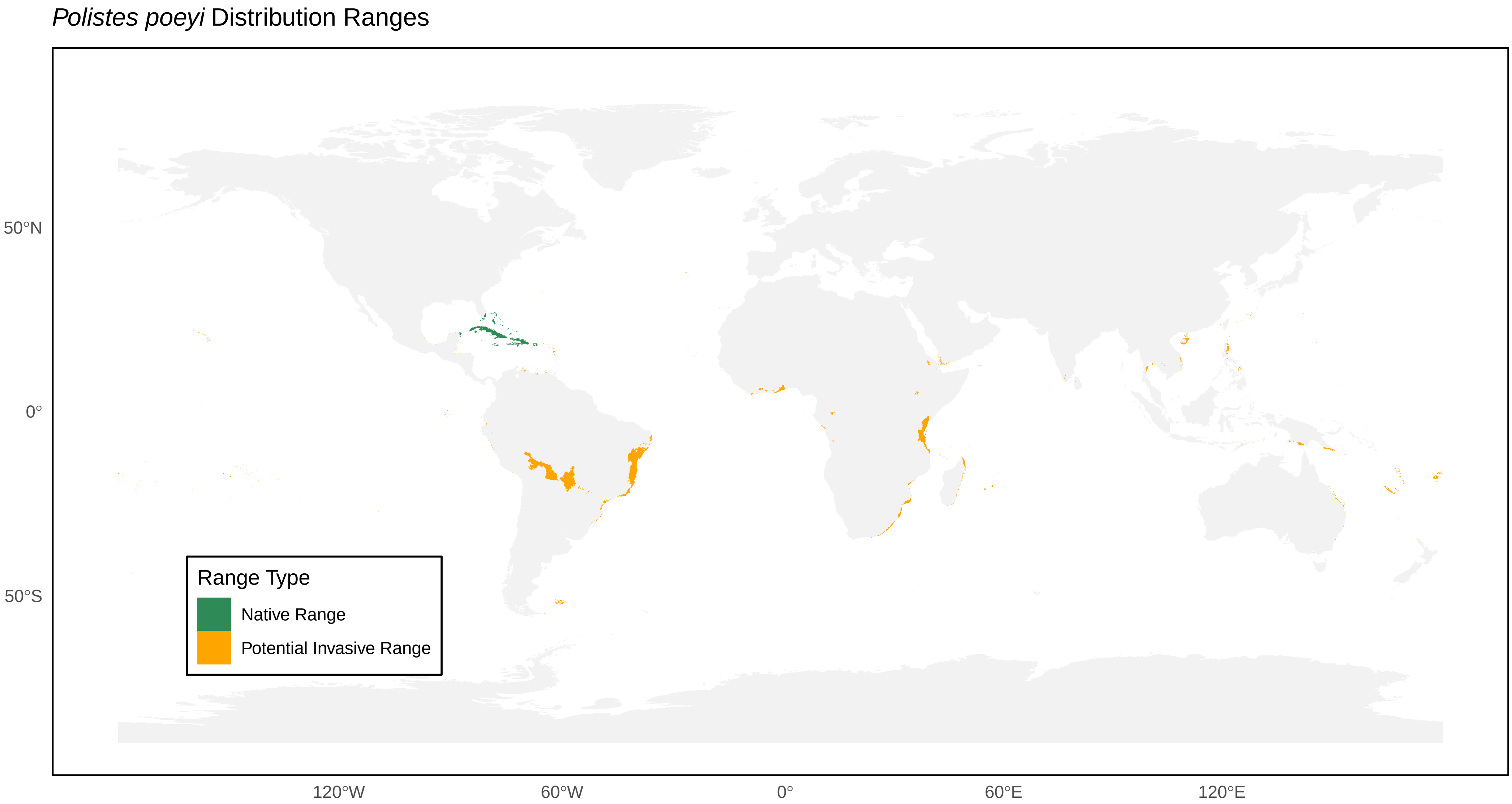

Polistes poeyi Distribution Ranges
50°N
0°
50°S
120°W
60°W
0°
60°E
120°E
Range Type
Native Range
Potential Invasive Range

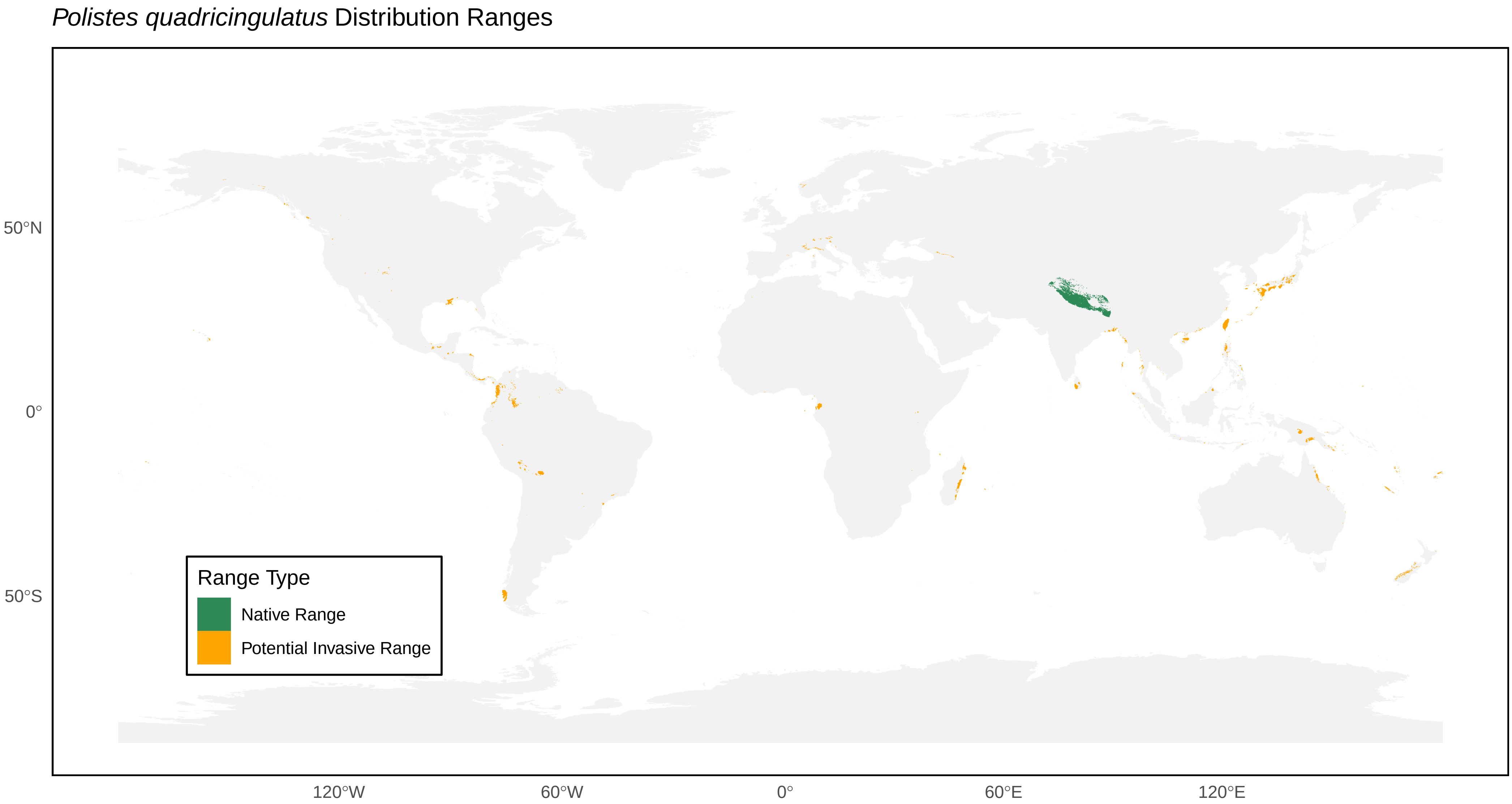
*Polistes quadricingulatus* Distribution Ranges
50°N
0°
50°S
120°W
60°W
0°
60°E
120°E
Range Type
Native Range
Potential Invasive Range

*Polistes riparius* Distribution Ranges

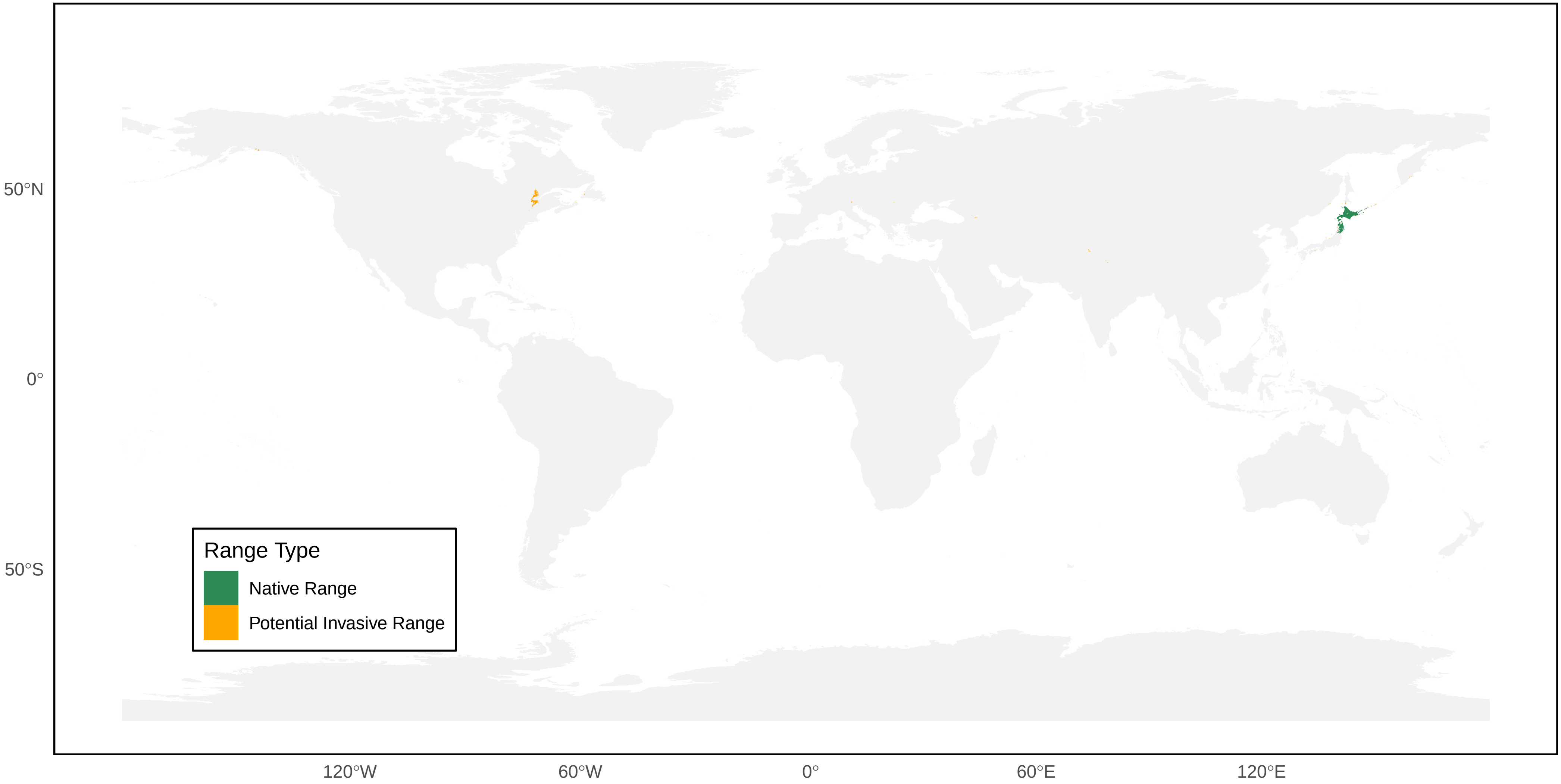

*Polistes rothneyi* Distribution Ranges

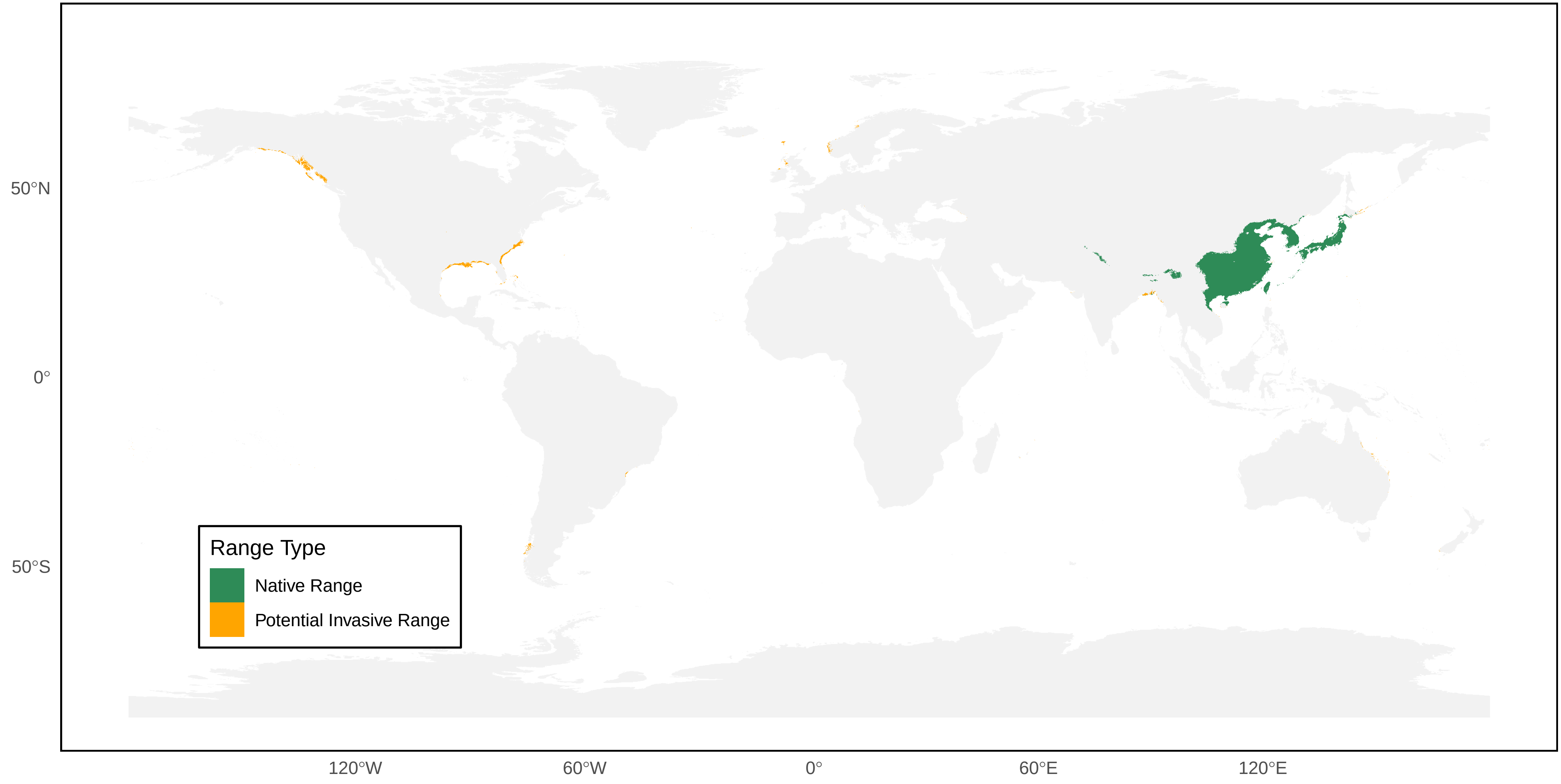

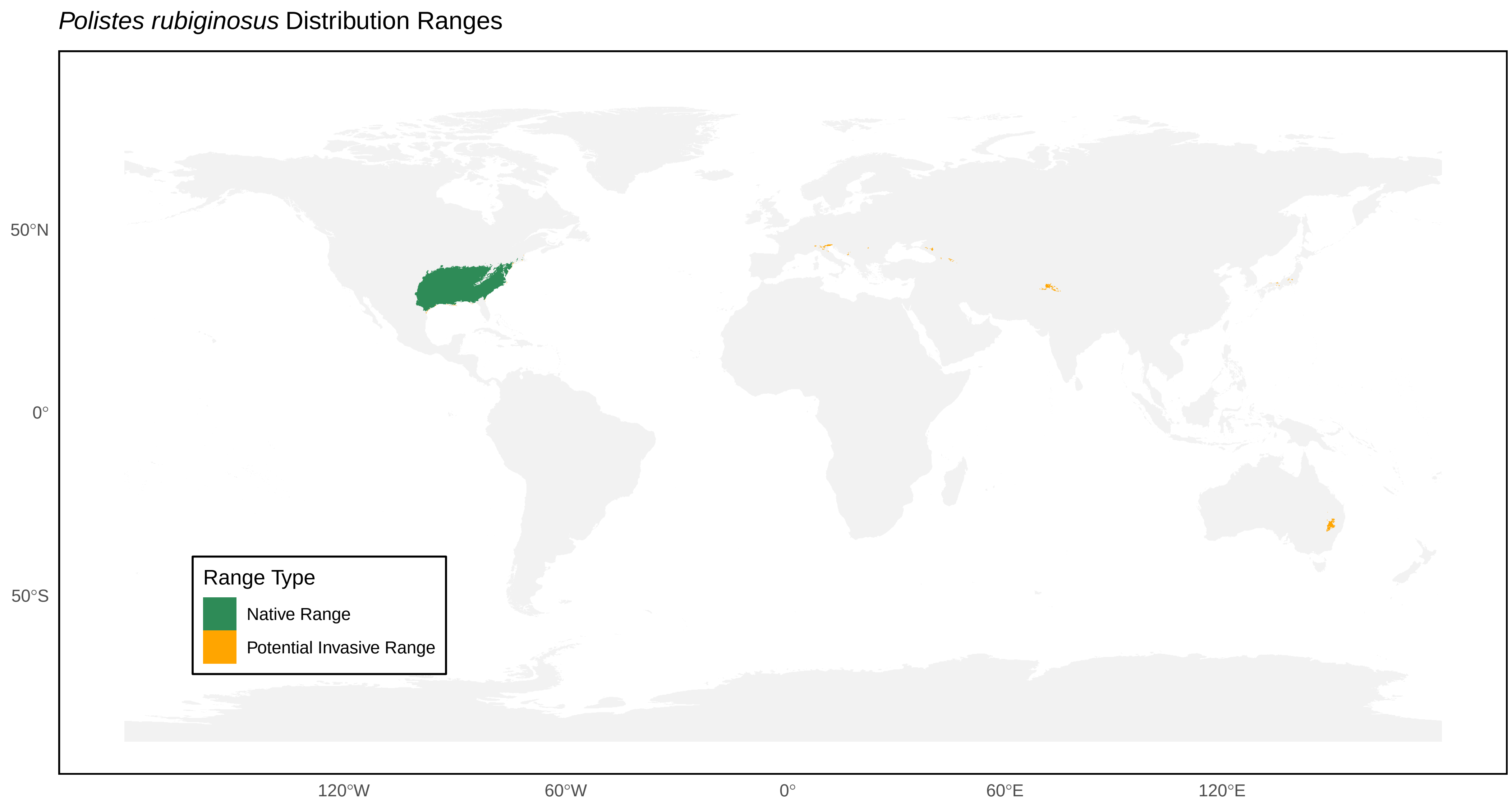

*Polistes rubiginosus* Distribution Ranges
50°N
0°
50°S
Range Type
Native Range
Potential Invasive Range
120°W
60°W
0°
60°E
120°E

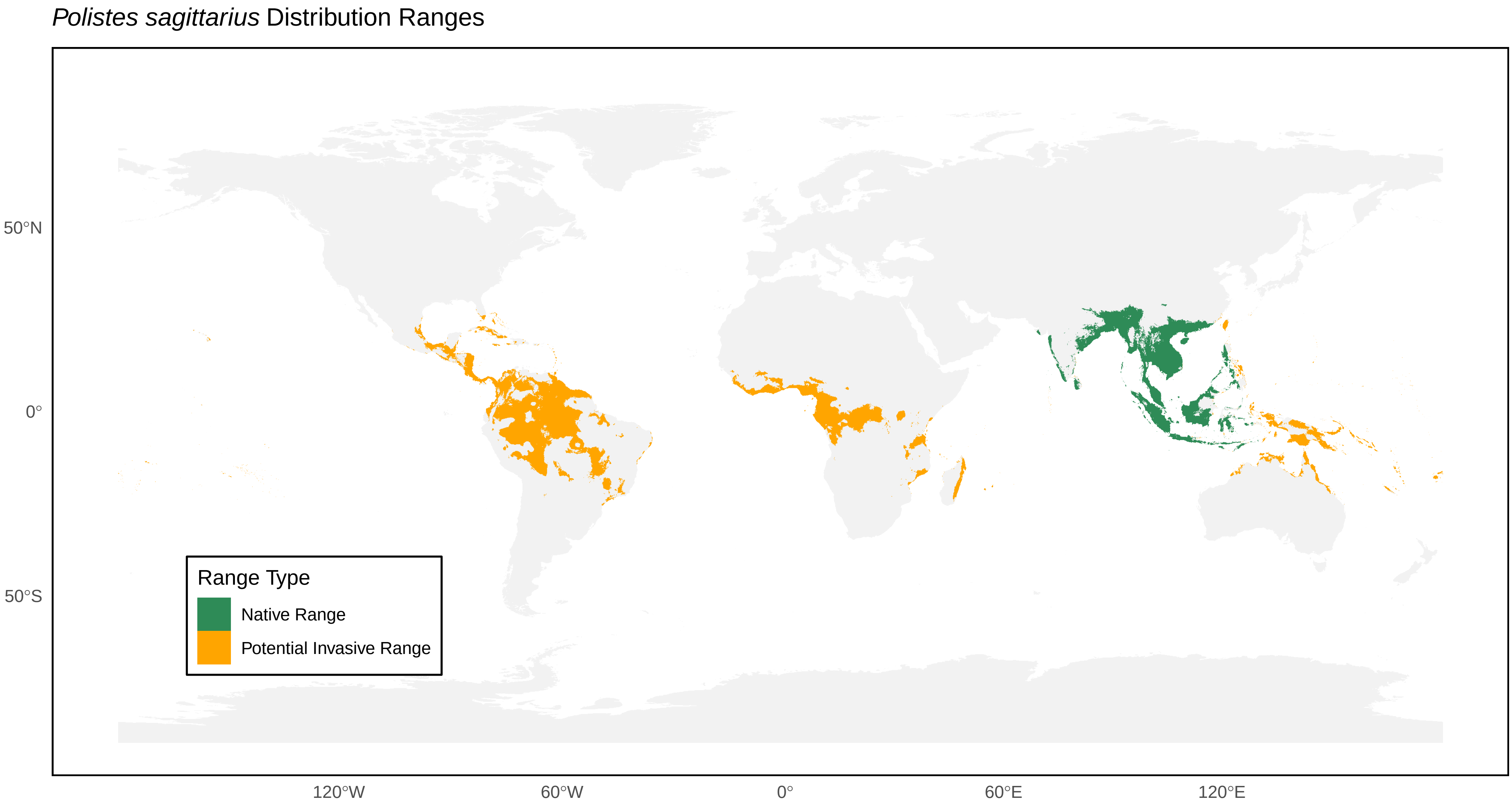

*Polistes sagittarius* Distribution Ranges
50°N
0°
50°S
120°W
60°W
0°
60°E
120°E
Range Type
Native Range
Potential Invasive Range

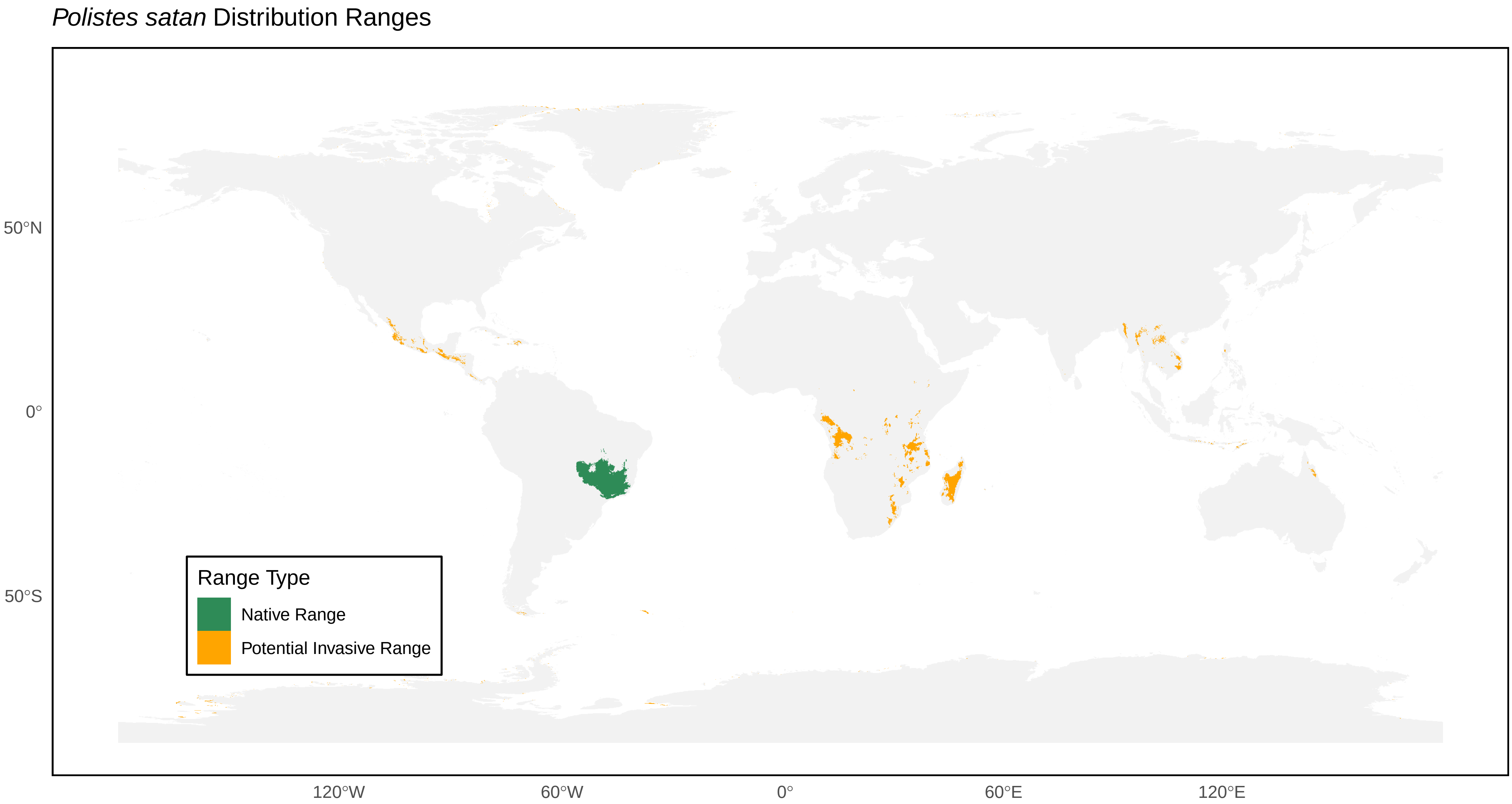

*Polistes satan* Distribution Ranges
50°N
0°
50°S
120°W
60°W
0°
60°E
120°E
Range Type
Native Range
Potential Invasive Range

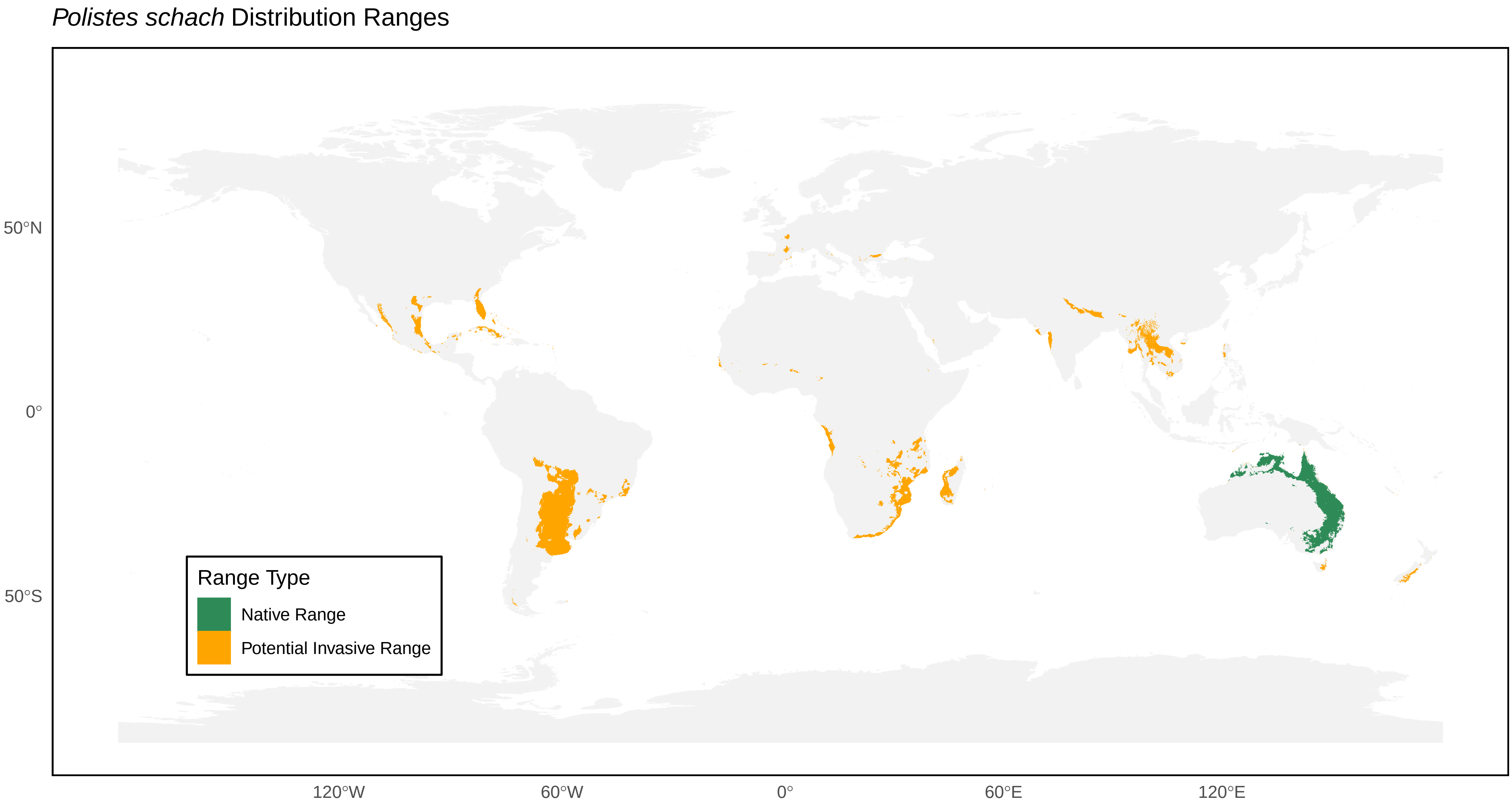
*Polistes schach* Distribution Ranges
50°N
0°
50°S
120°W
60°W
0°
60°E
120°E
Range Type
Native Range
Potential Invasive Range

*Polistes semenowi* Distribution Ranges

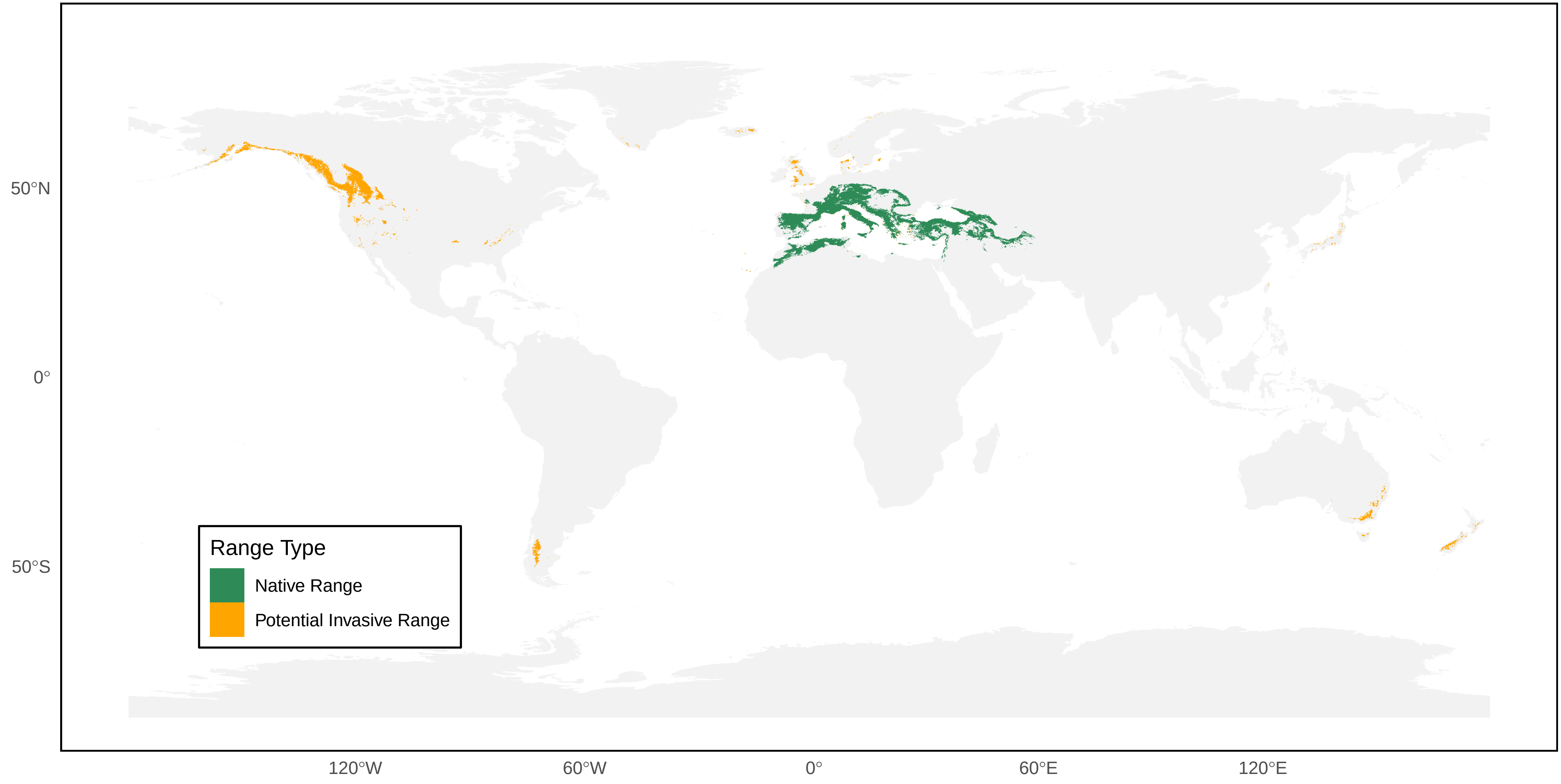

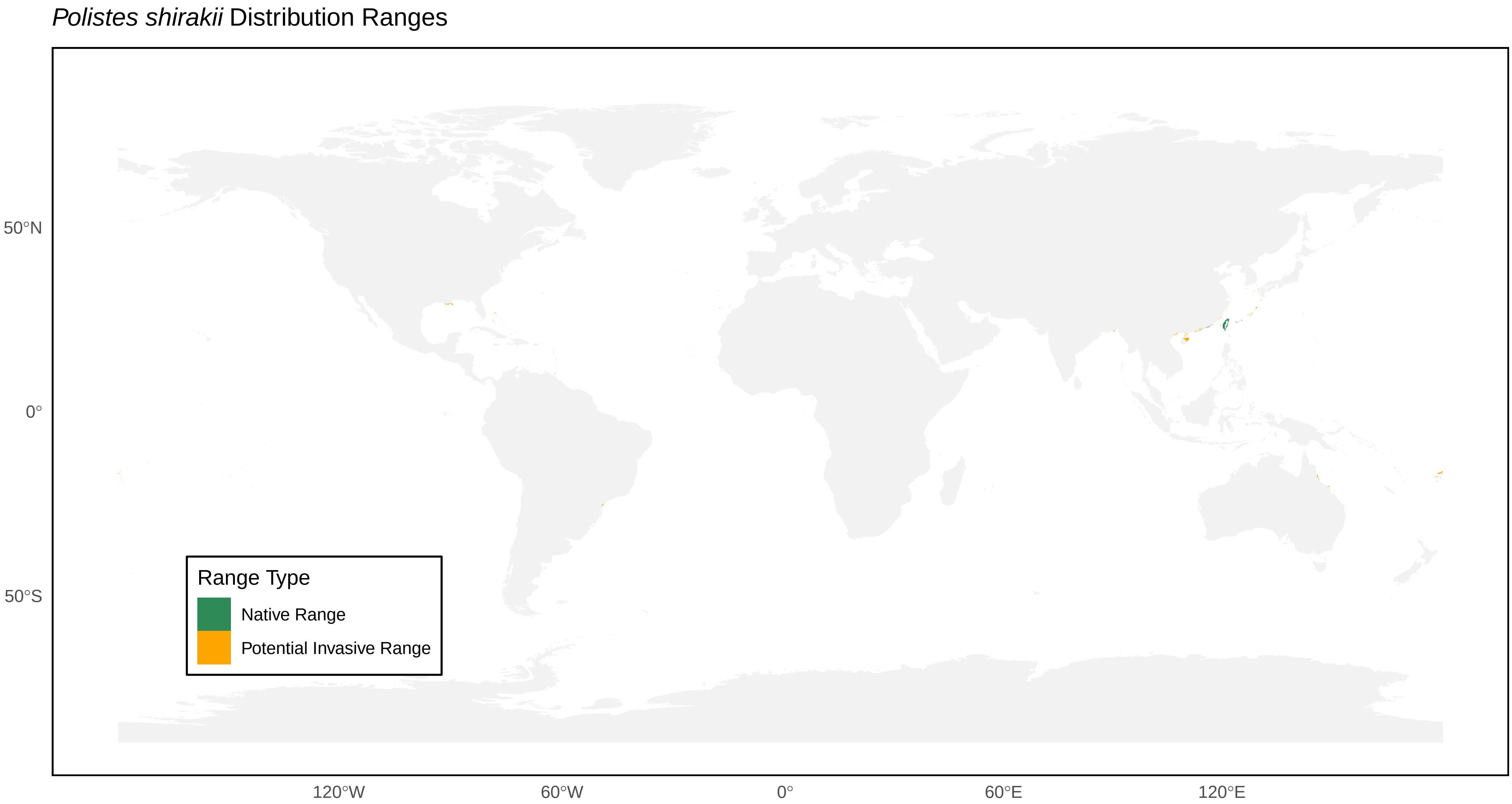

*Polistes shirakii* Distribution Ranges
50°N
0°
50°S
120°W
60°W
0°
60°E
120°E
Range Type
Native Range
Potential Invasive Range

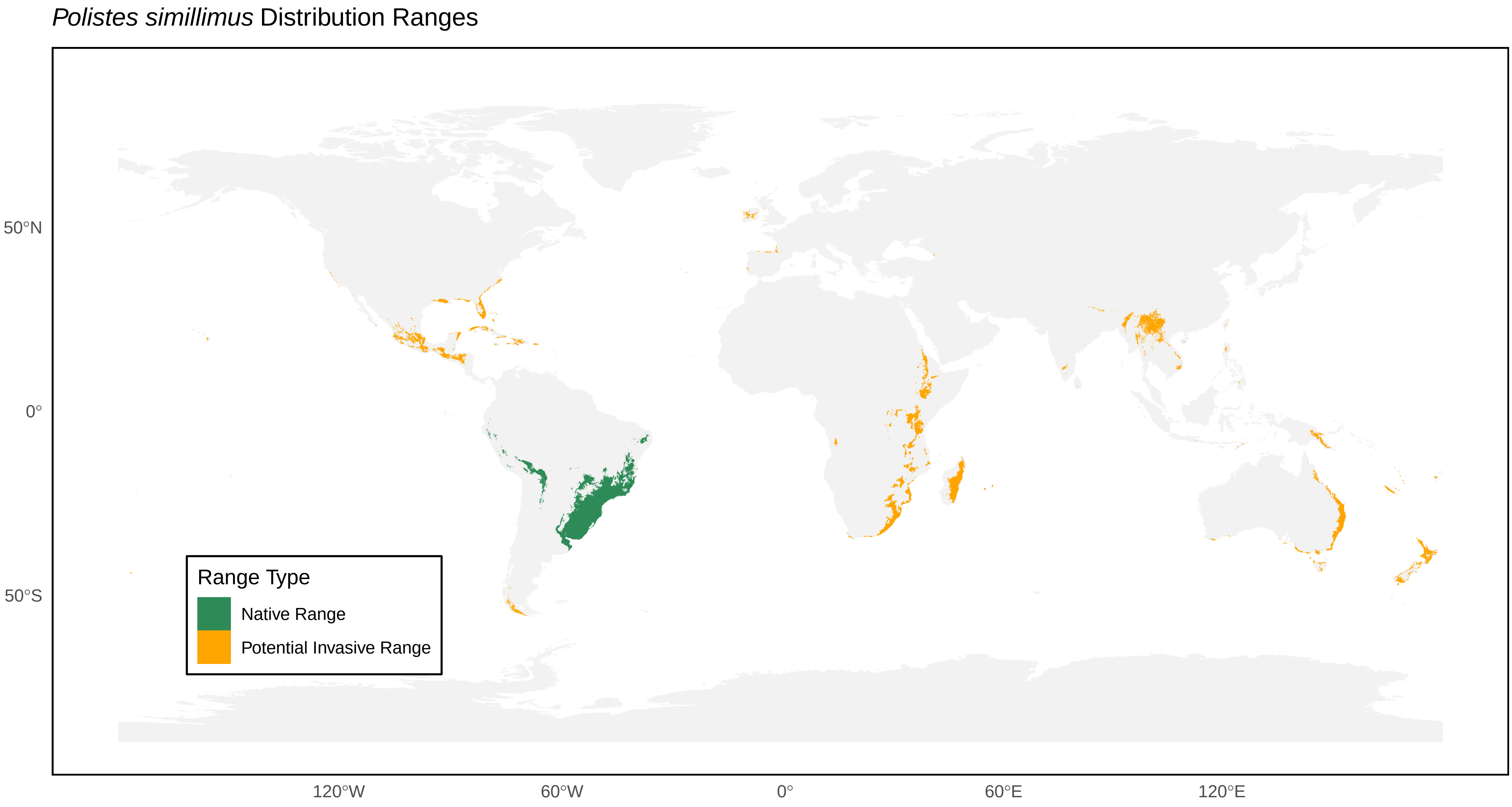

*Polistes simillimus* Distribution Ranges
50°N
0°
50°S
120°W
60°W
0°
60°E
120°E
Range Type
Native Range
Potential Invasive Range

*Polistes smithii* Distribution Ranges

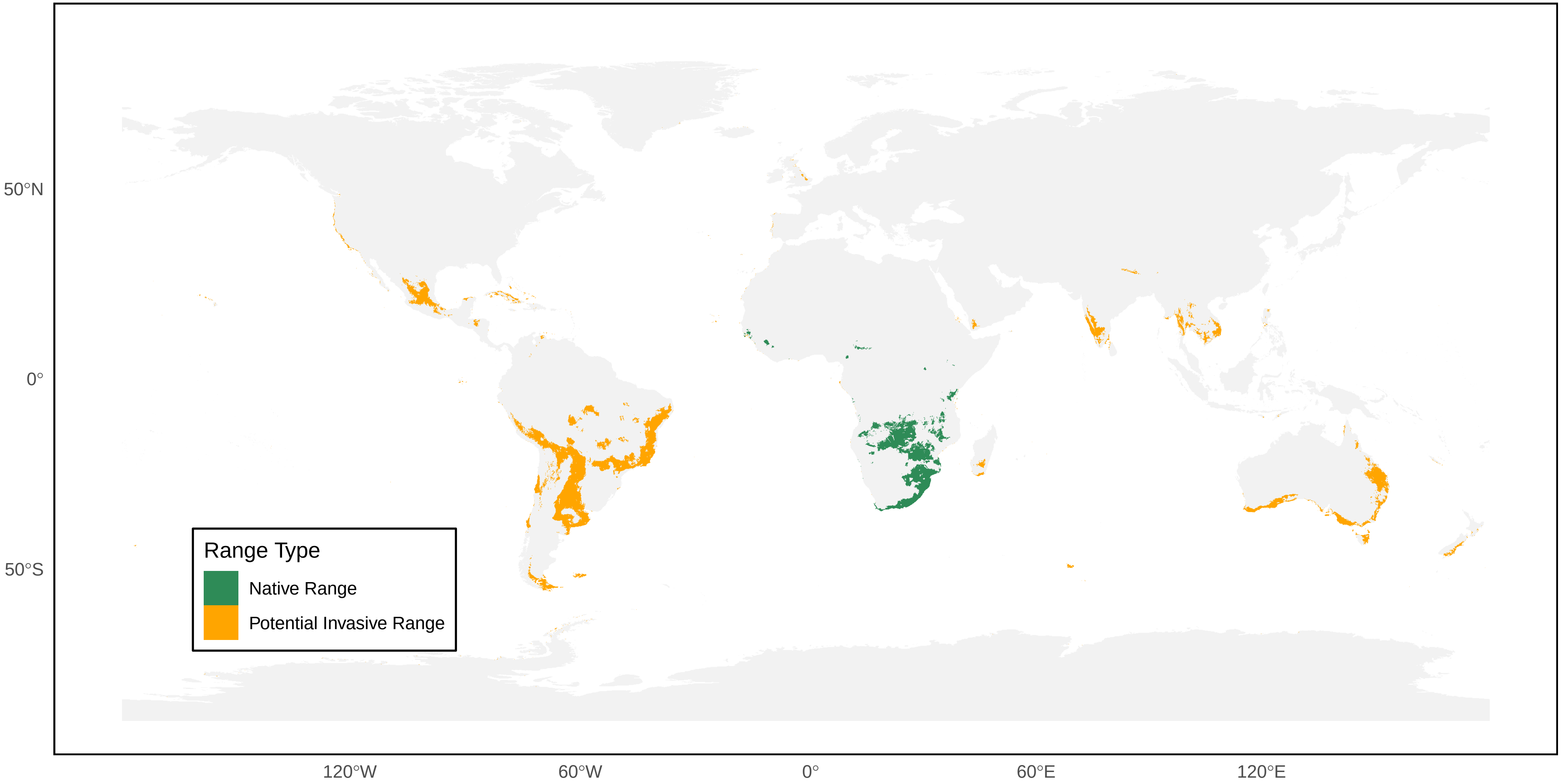

#### *Polistes snelleni* Distribution Ranges

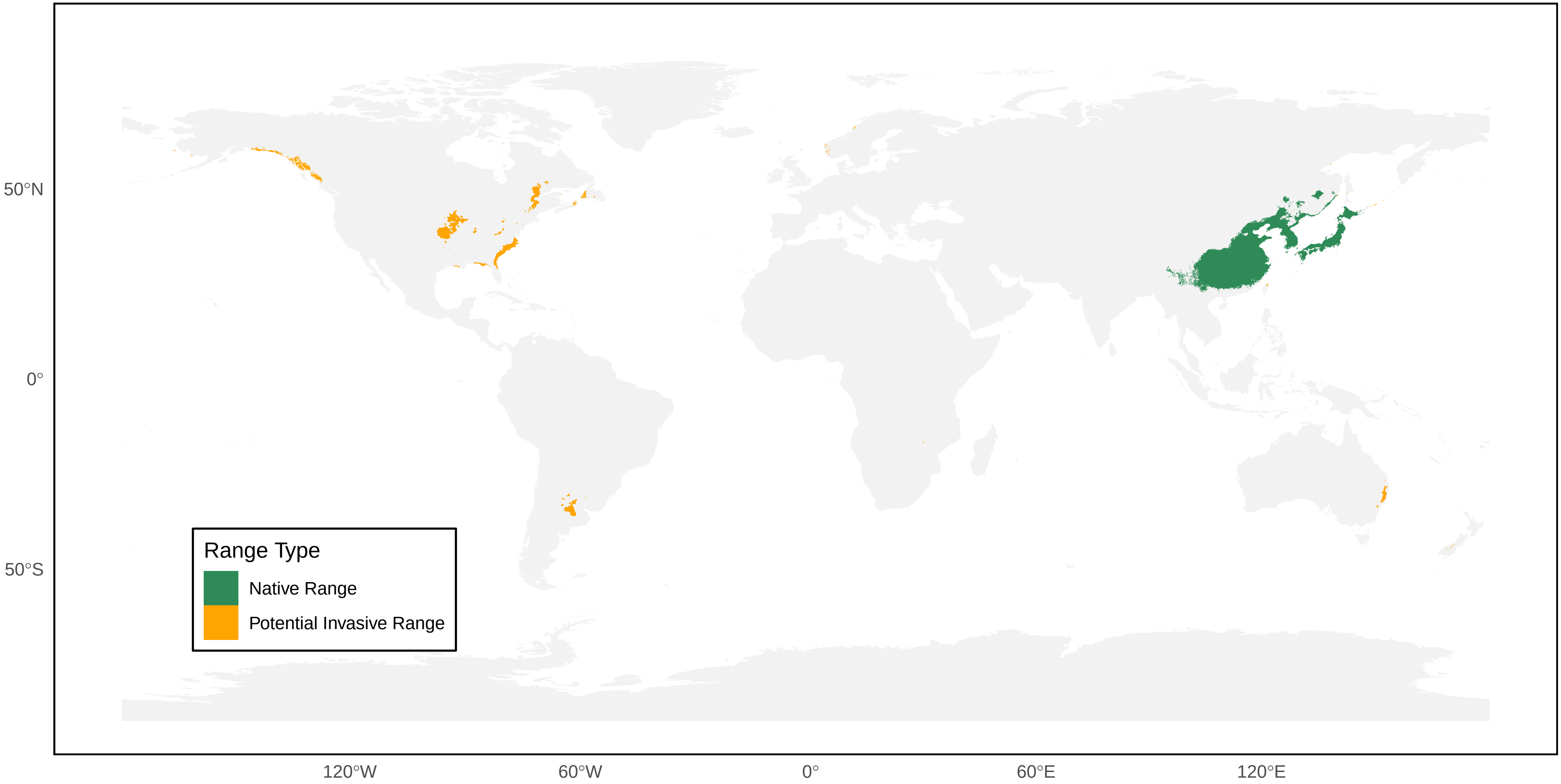

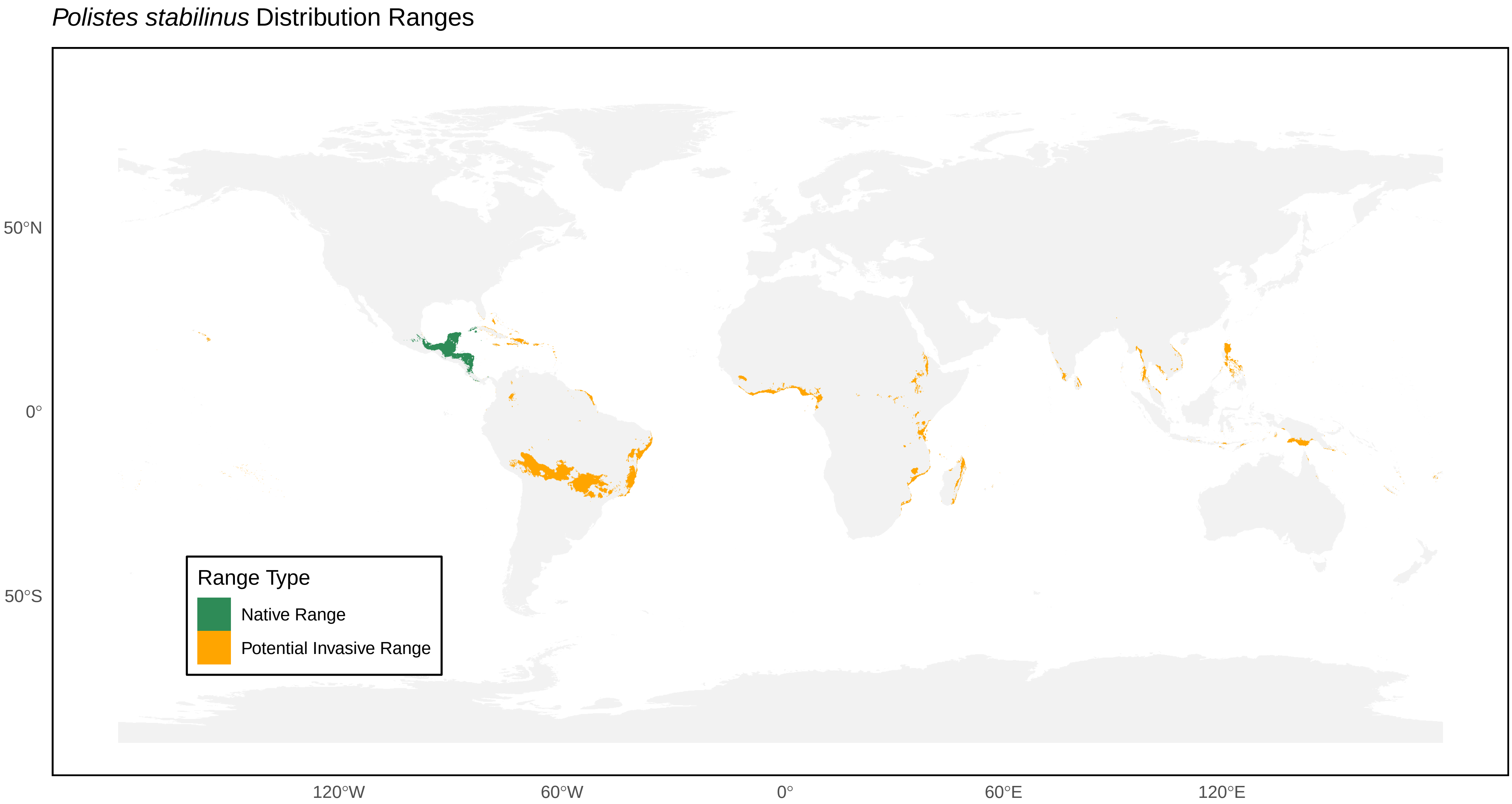

Polistes stabilinus Distribution Ranges
50°N
0°
50°S
120°W
60°W
0°
60°E
120°E
Range Type
Native Range
Potential Invasive Range

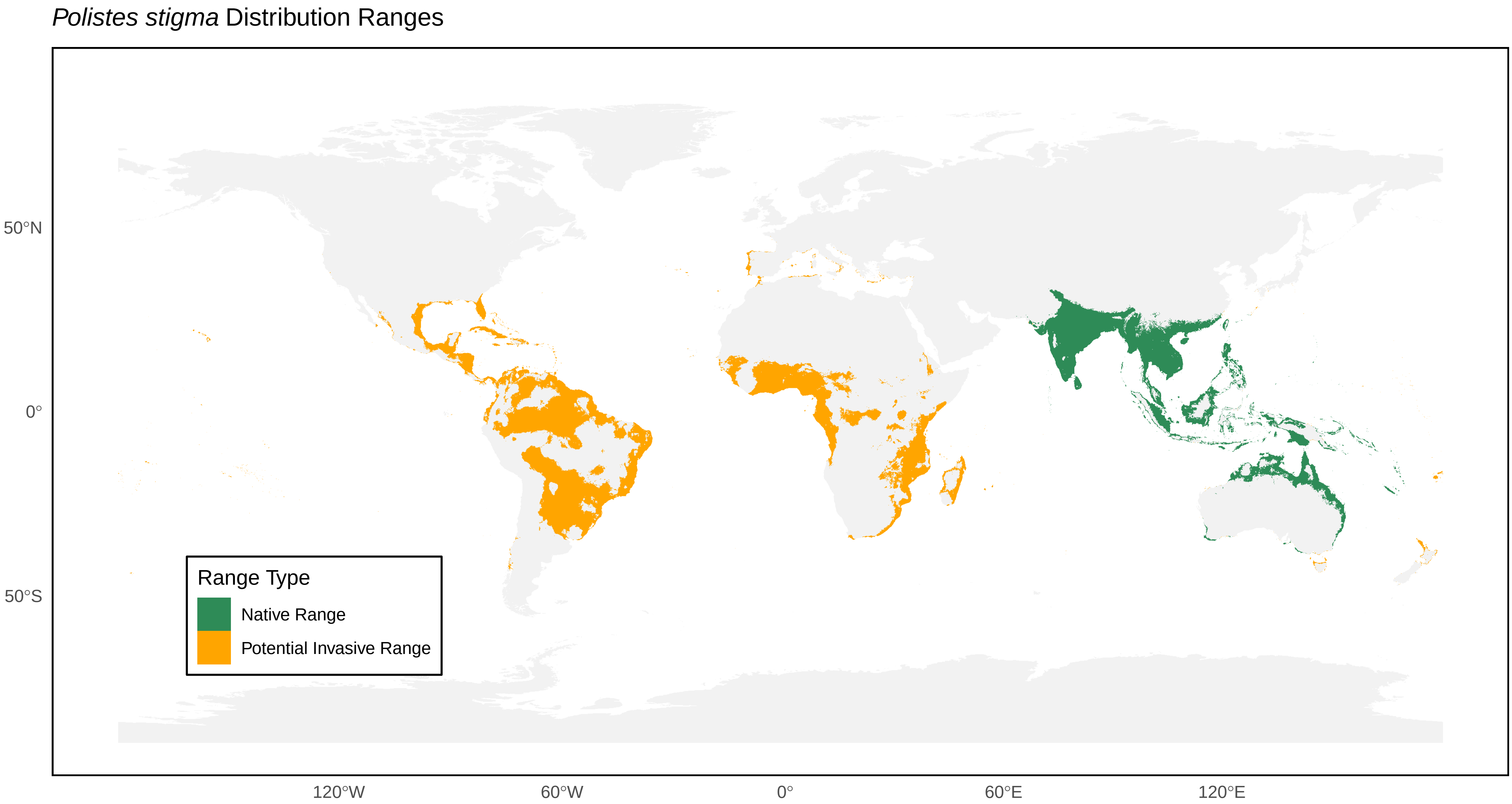
*Polistes stigma* Distribution Ranges
50°N
0°
50°S
120°W
60°W
0°
60°E
120°E
Range Type
Native Range
Potential Invasive Range

*Polistes strigosus* Distribution Ranges

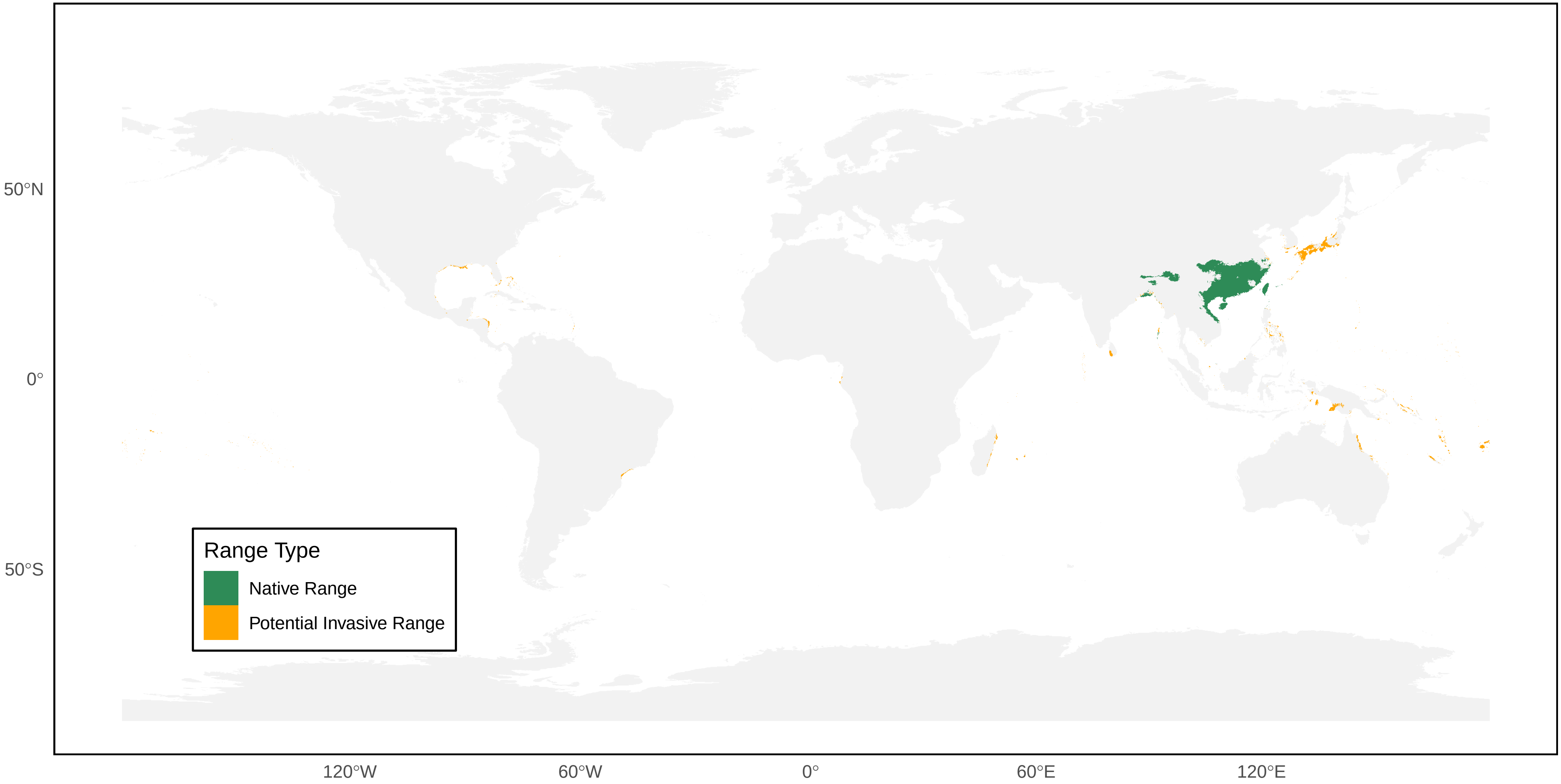

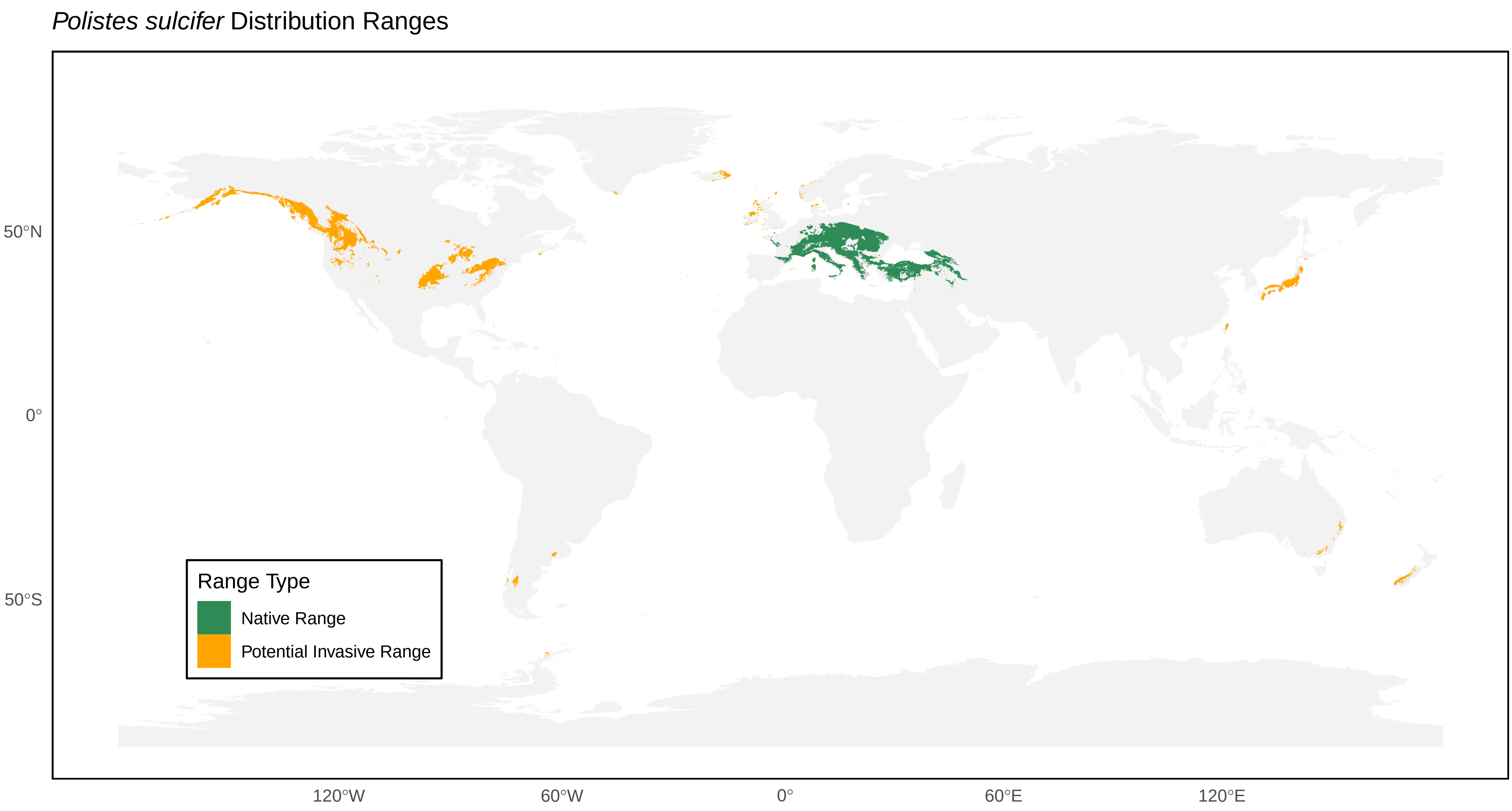

*Polistes sulcifer* Distribution Ranges
50°N
0°
50°S
120°W
60°W
0°
60°E
120°E
Range Type
Native Range
Potential Invasive Range

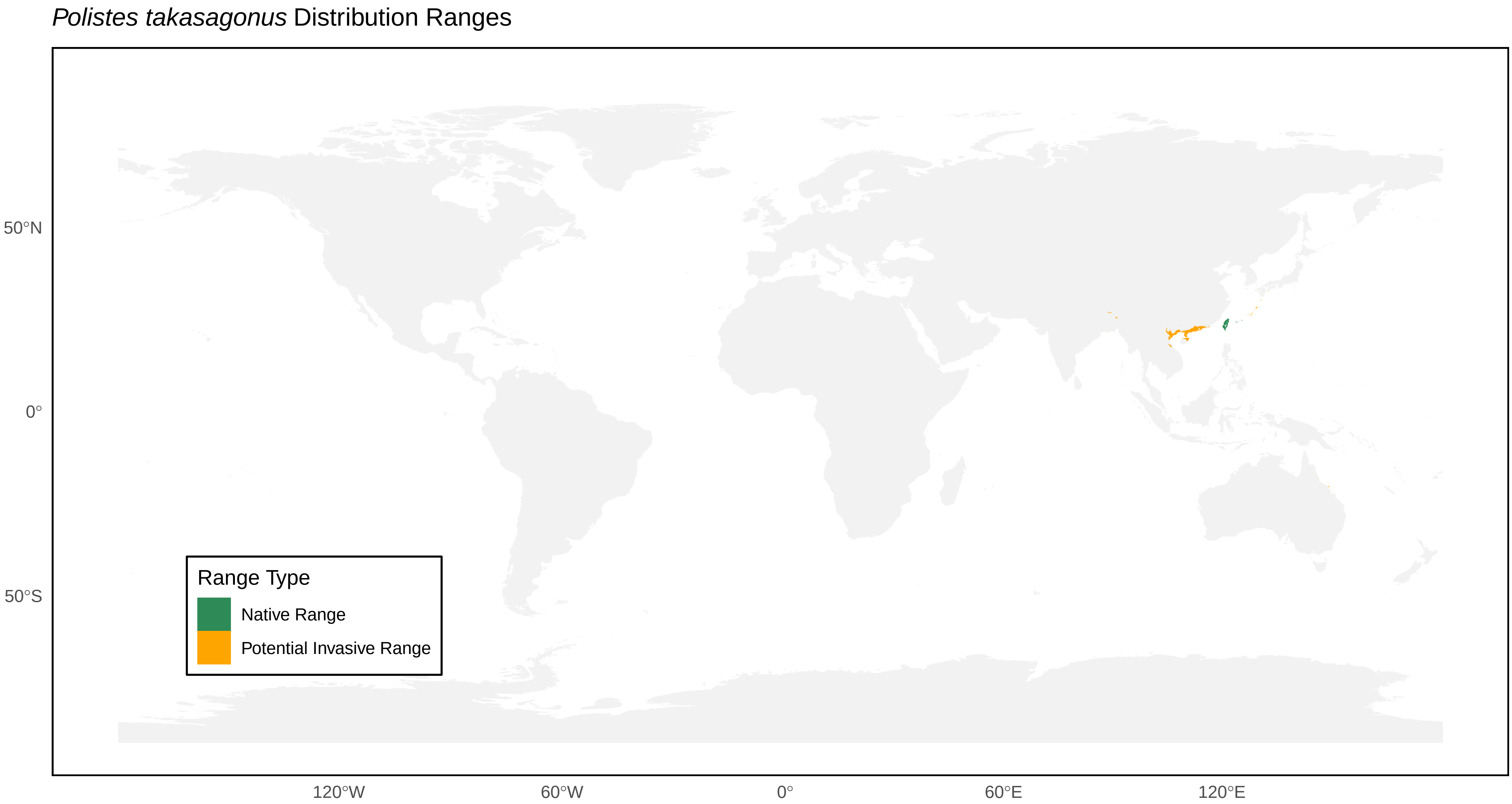

*Polistes takasagonus* Distribution Ranges
50°N
0°
50°S
120°W
60°W
0°
60°E
120°E
Range Type
Native Range
Potential Invasive Range

*Polistes tenebricosus* Distribution Ranges

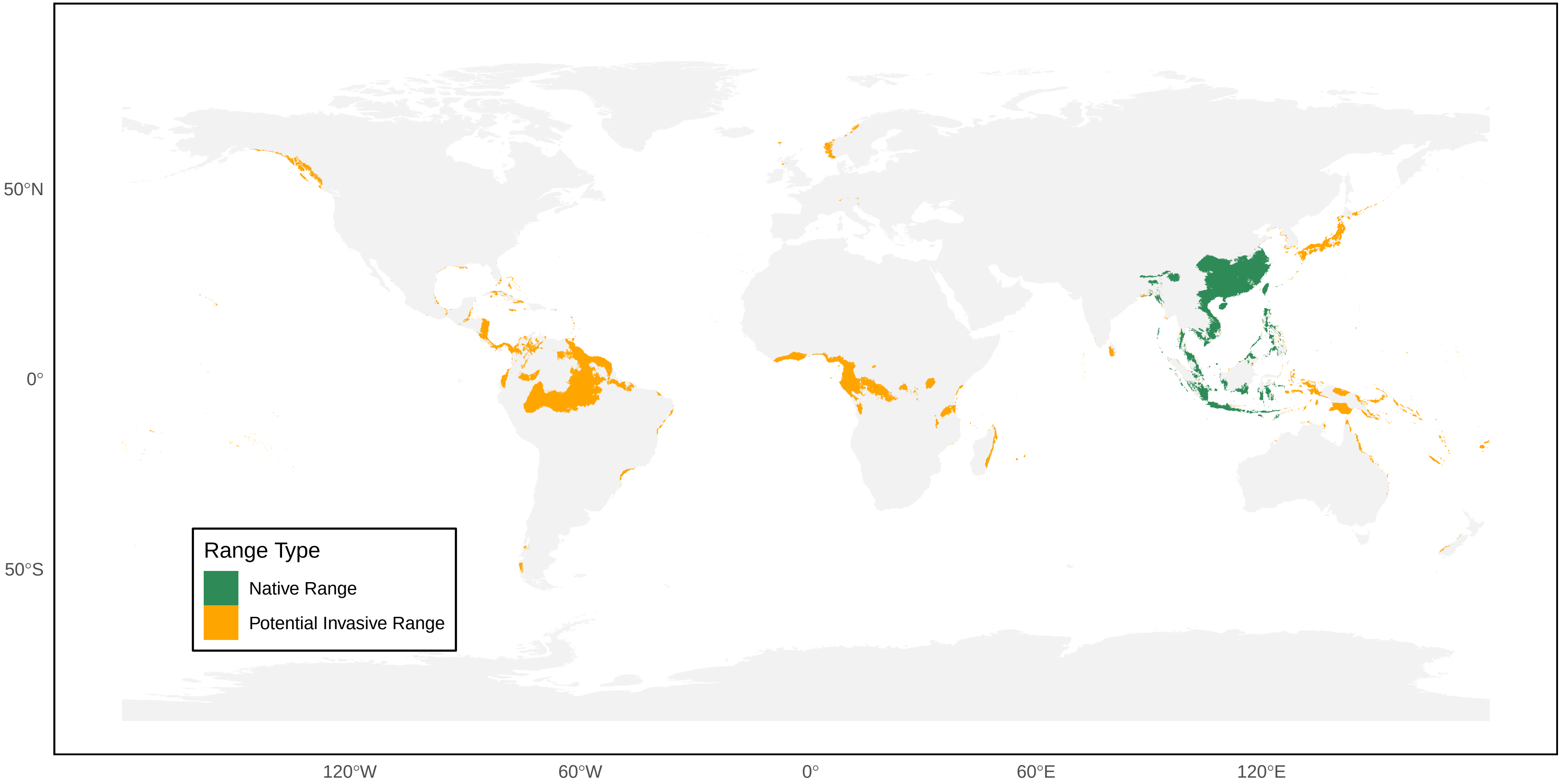

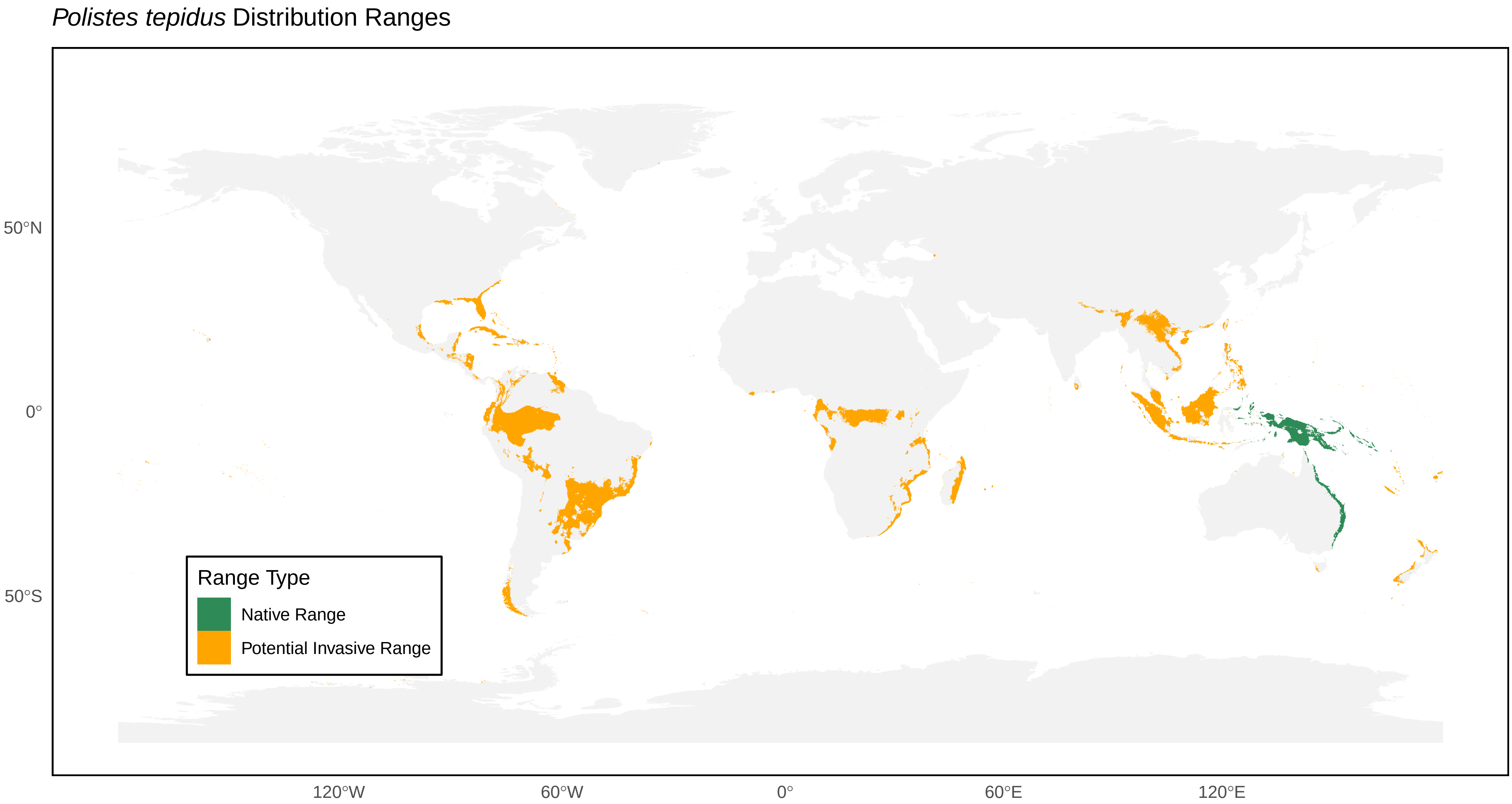
*Polistes tepidus* Distribution Ranges
50°N
0°
50°S
120°W
60°W
0°
60°E
120°E
Range Type
Native Range
Potential Invasive Range

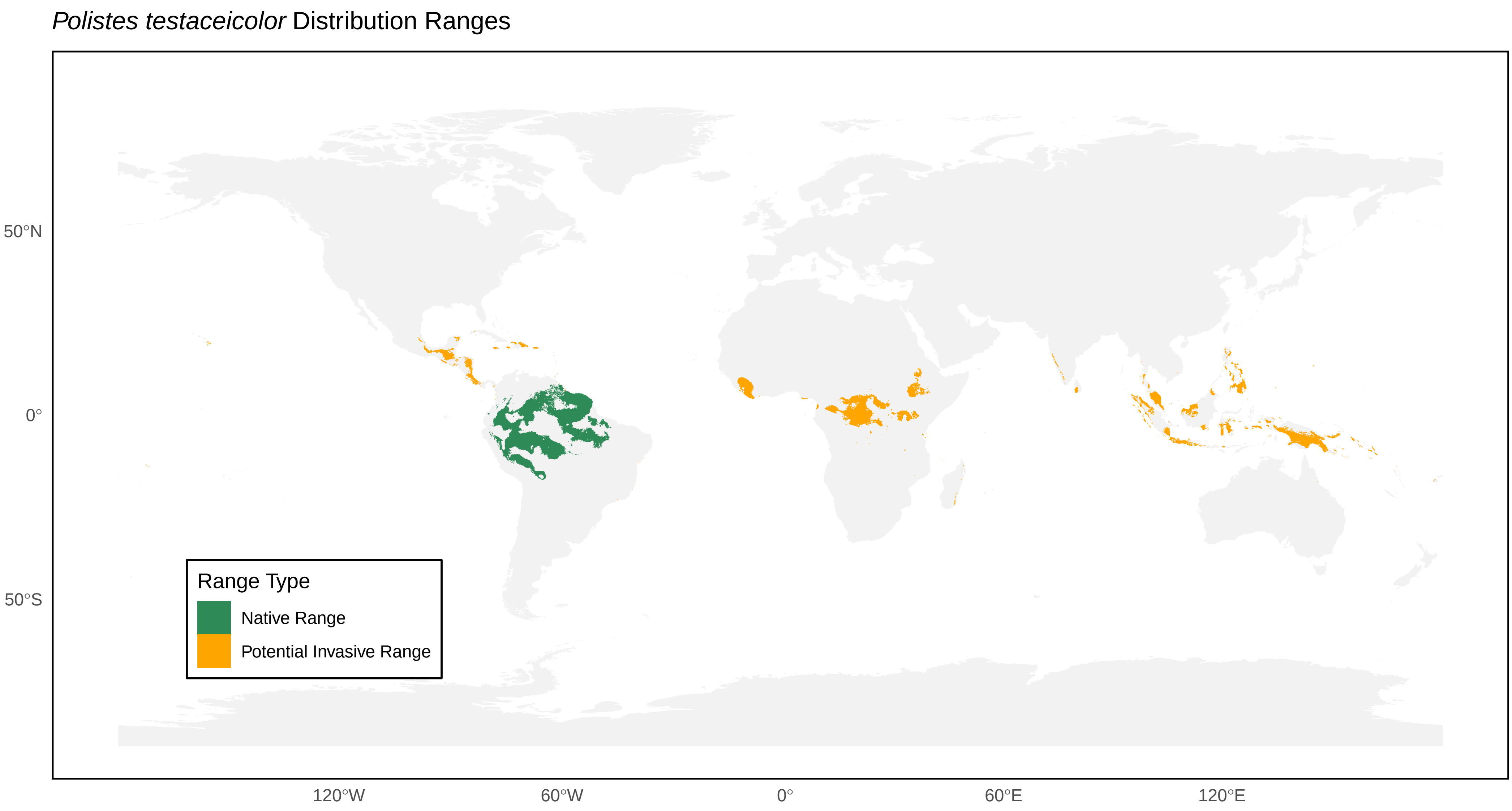
*Polistes testaceicolor* Distribution Ranges
50°N
0°
50°S
120°W
60°W
0°
60°E
120°E
Range Type
Native Range
Potential Invasive Range

*Polistes veracrucis* Distribution Ranges

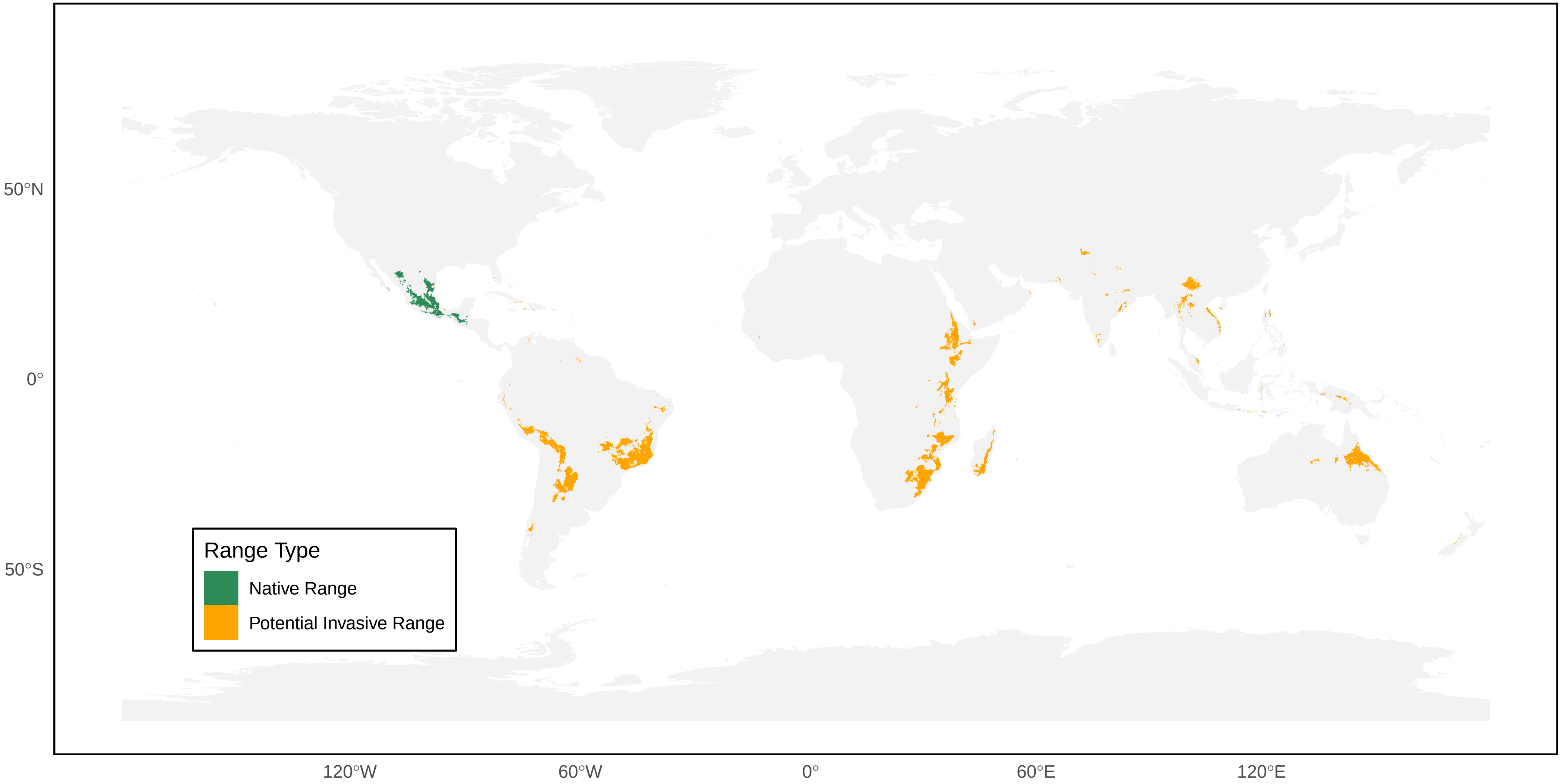

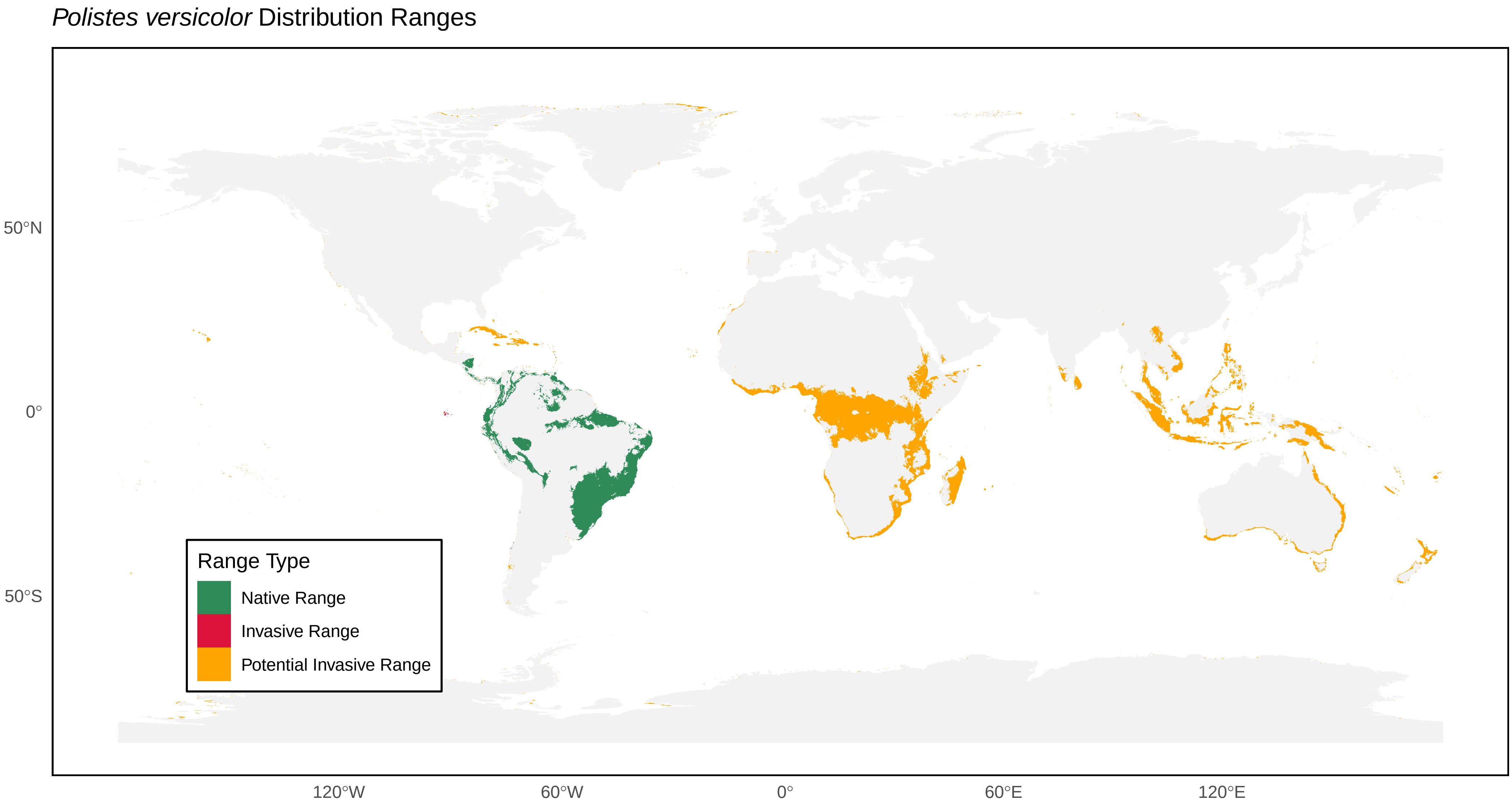
*Polistes versicolor* Distribution Ranges
50°N
0°
50°S
120°W
60°W
0°
60°E
120°E
Range Type
Native Range
Invasive Range
Potential Invasive Range

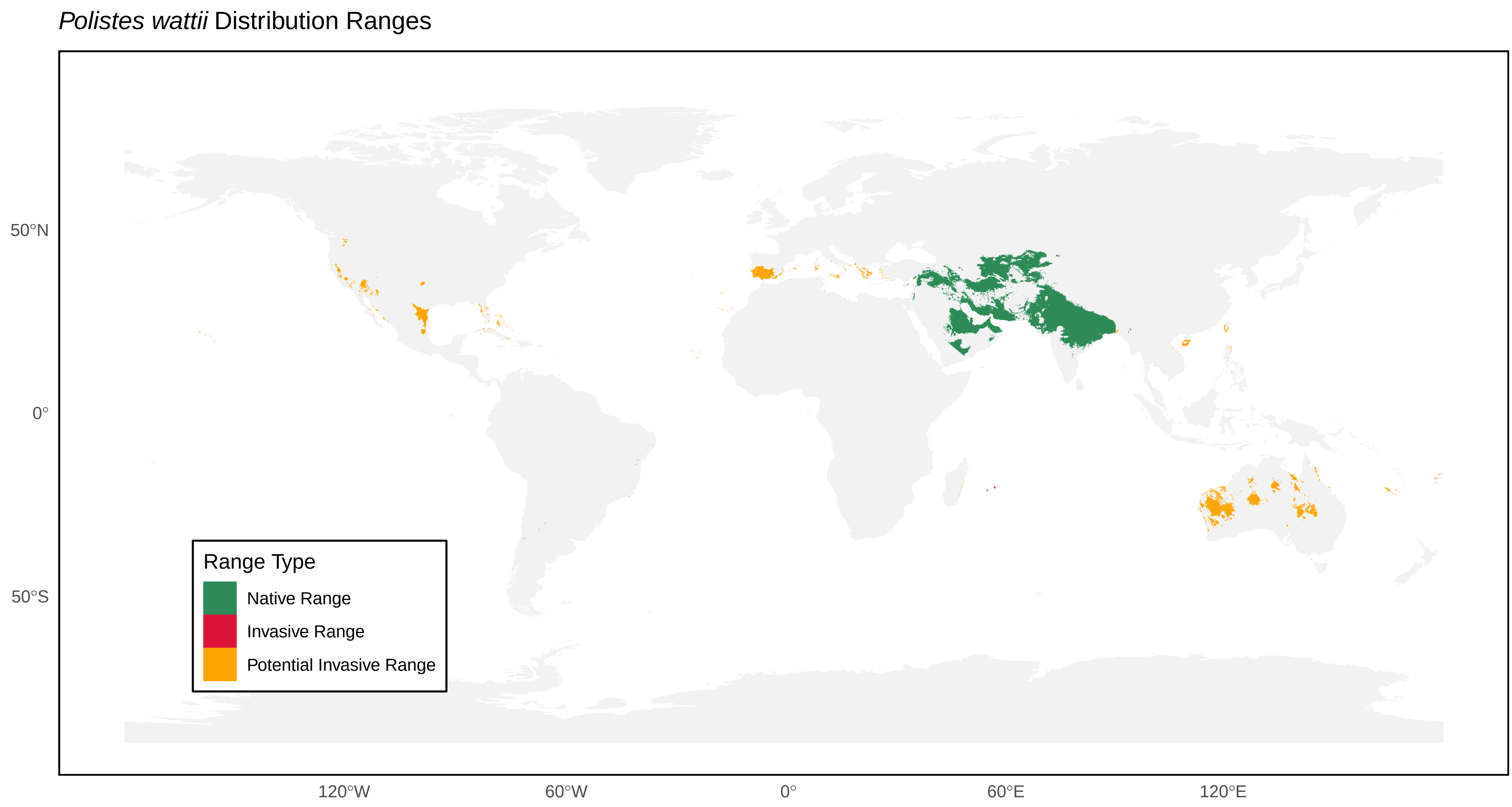

*Polistes wattii* Distribution Ranges
50°N
0°
50°S
120°W
60°W
0°
60°E
120°E
Range Type
Native Range
Invasive Range
Potential Invasive Range

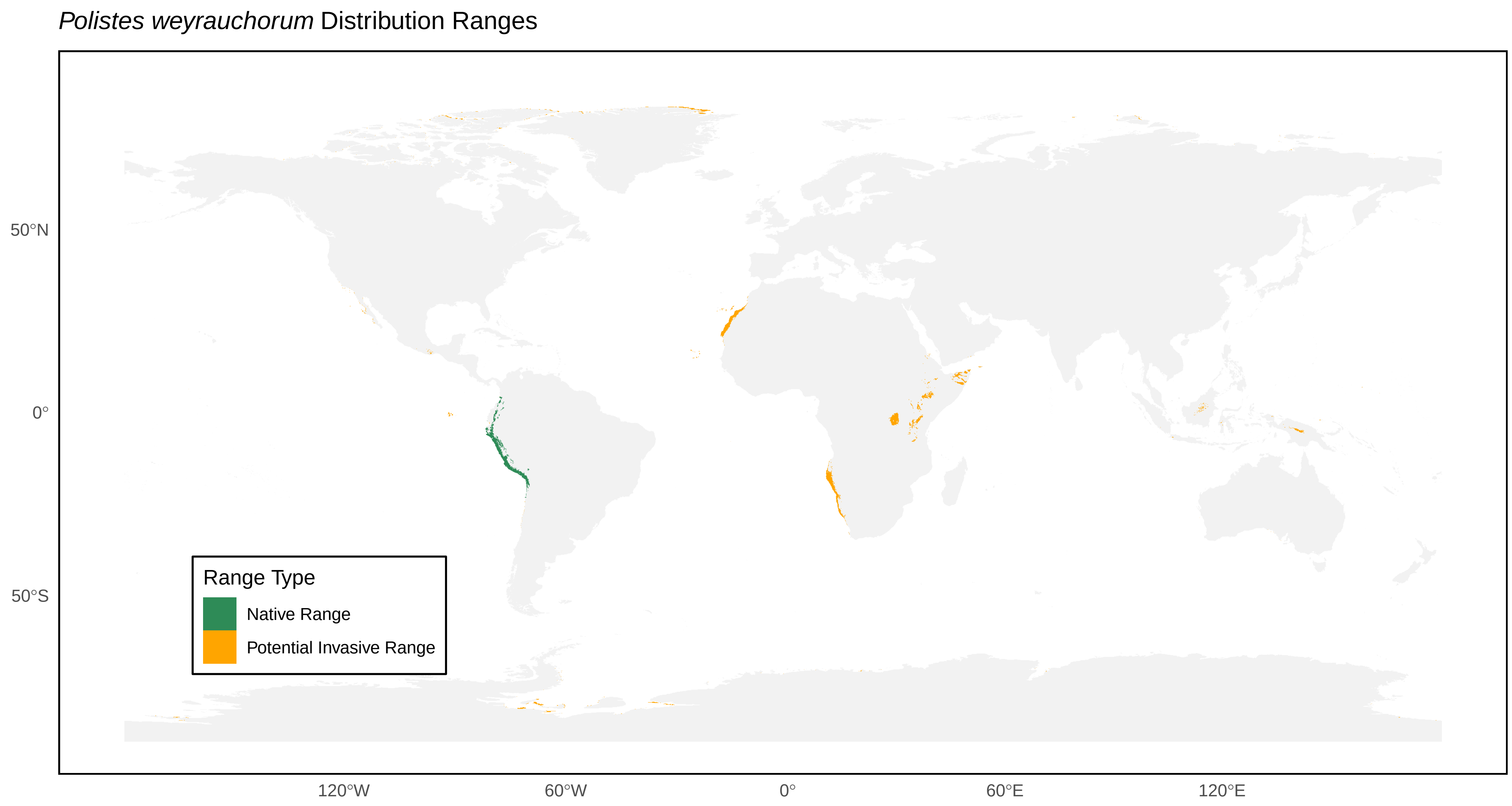

Polistes weyrauchorum Distribution Ranges
50°N
0°
50°S
120°W
60°W
0°
60°E
120°E
Range Type
Native Range
Potential Invasive Range

*Polistes xanthogaster* Distribution Ranges

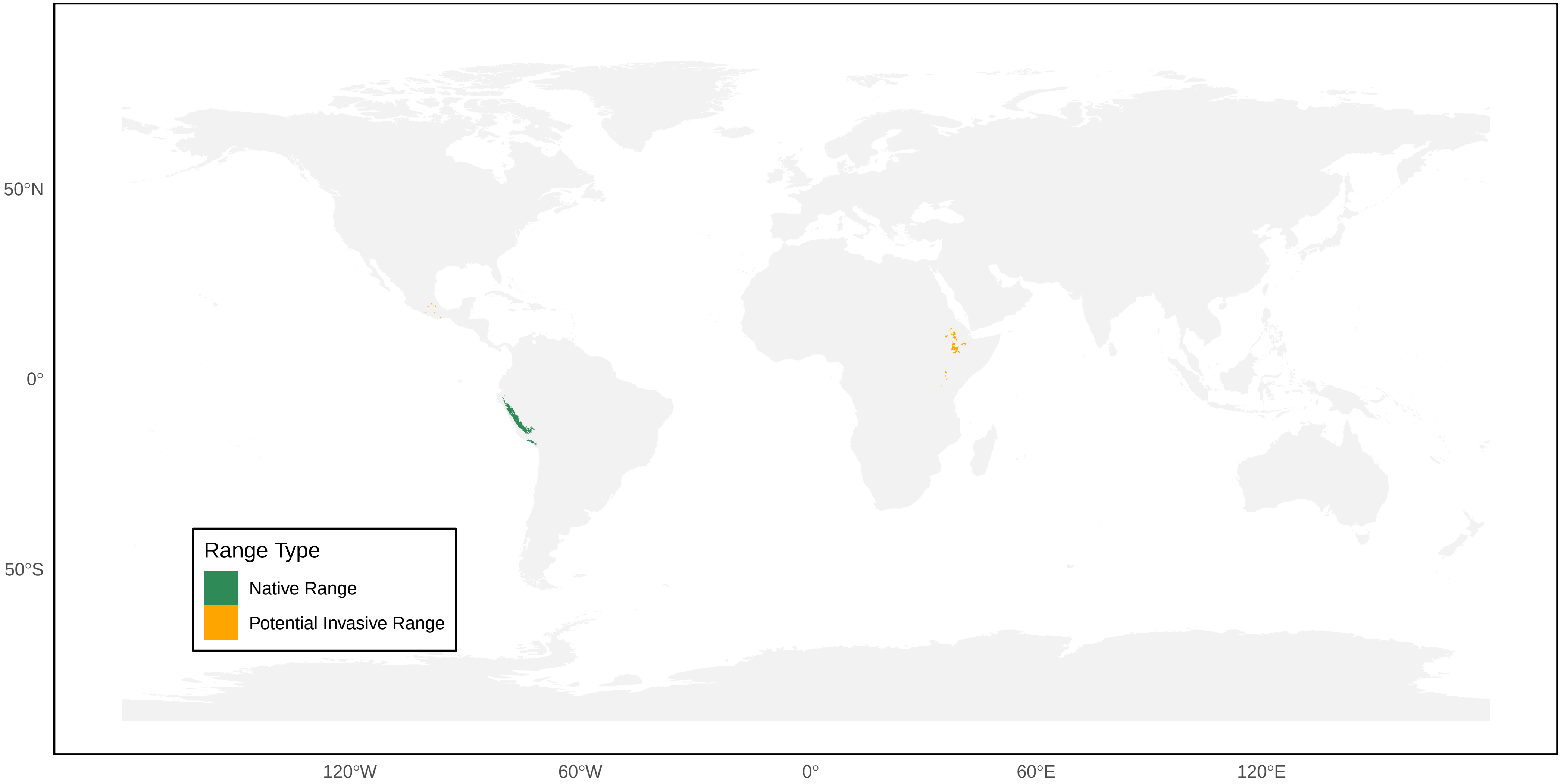

*Polybia aequatorialis* Distribution Ranges

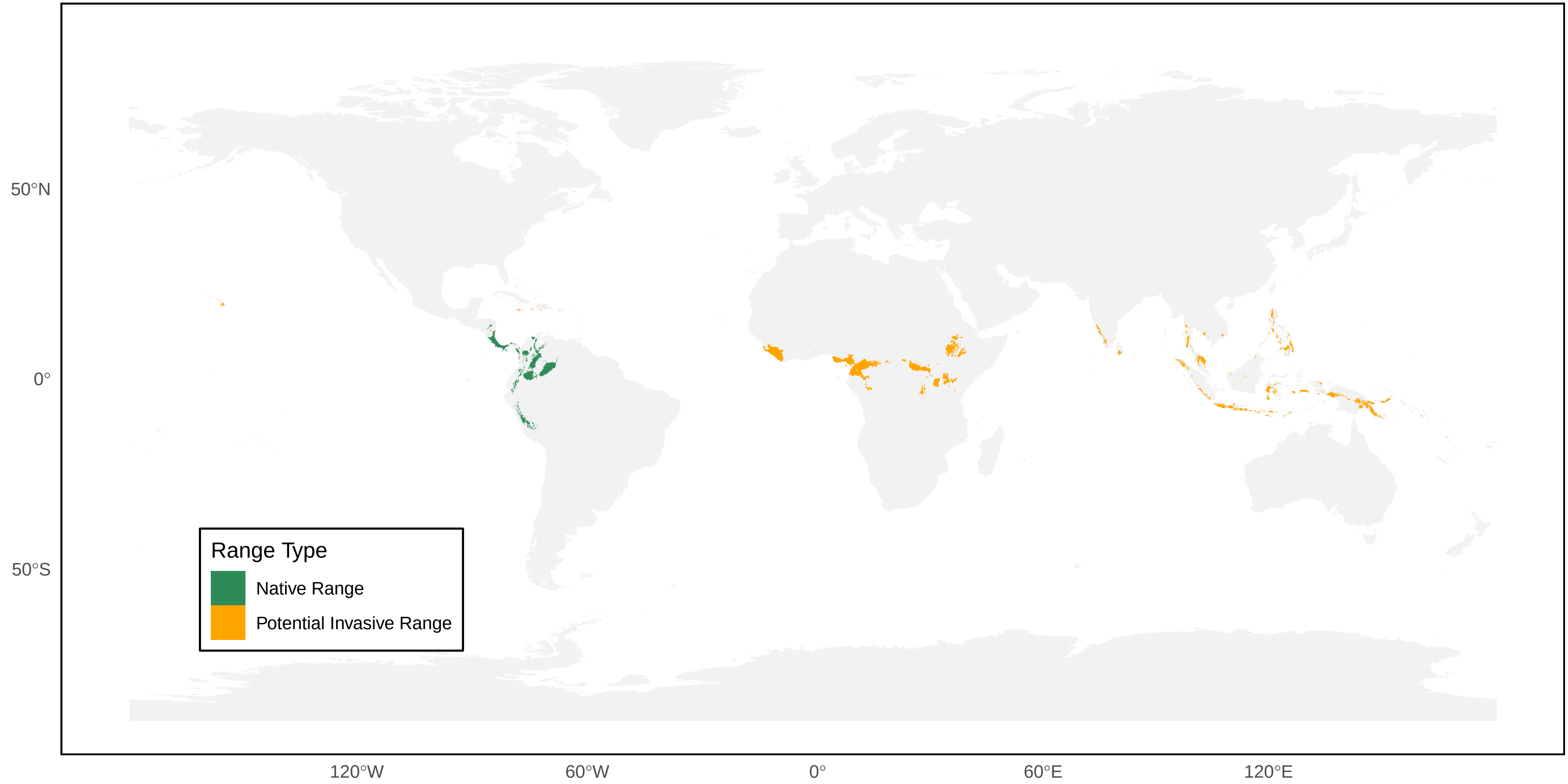

*Polybia belemensis* Distribution Ranges

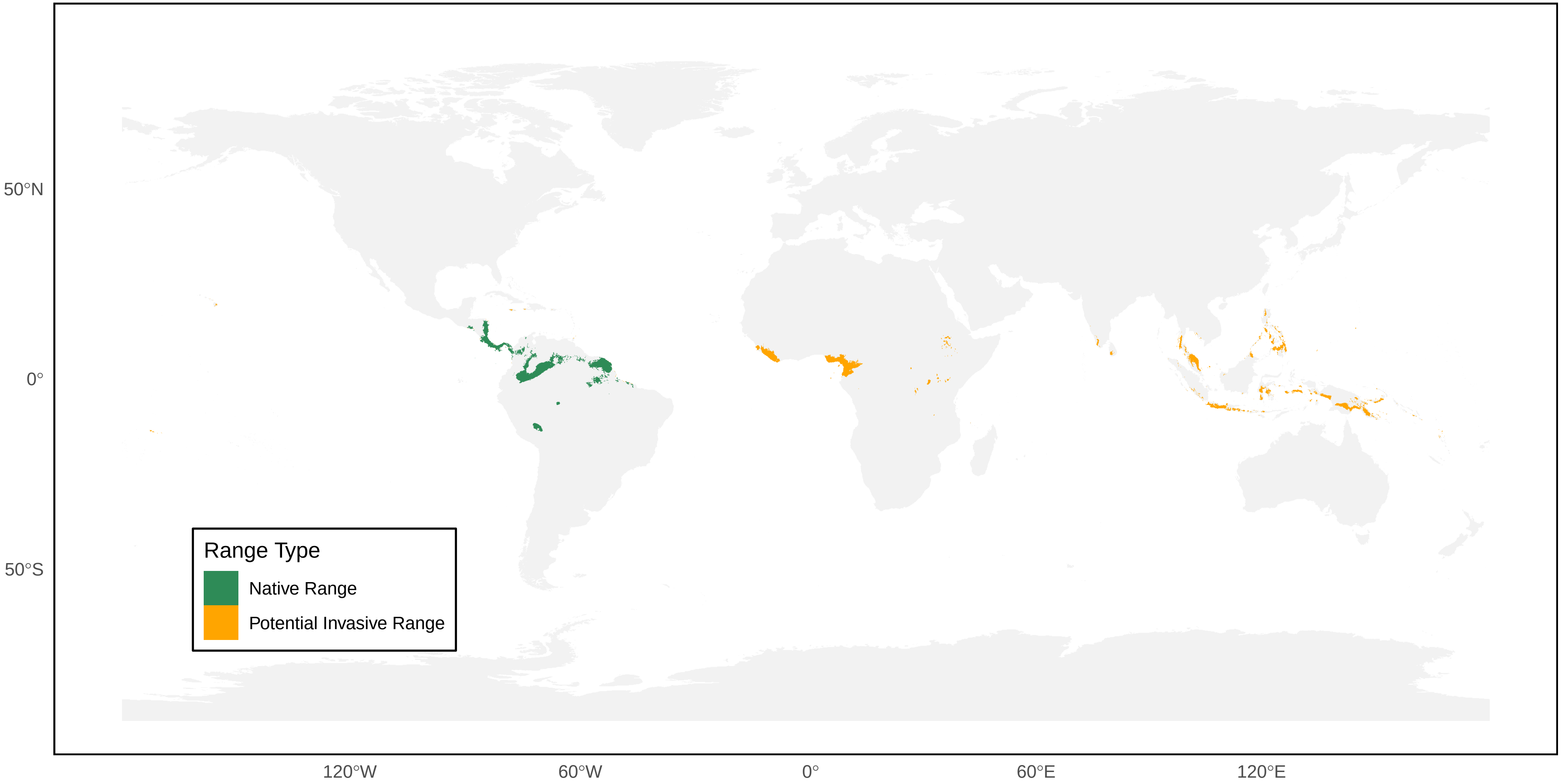

*Polybia bistriata* Distribution Ranges

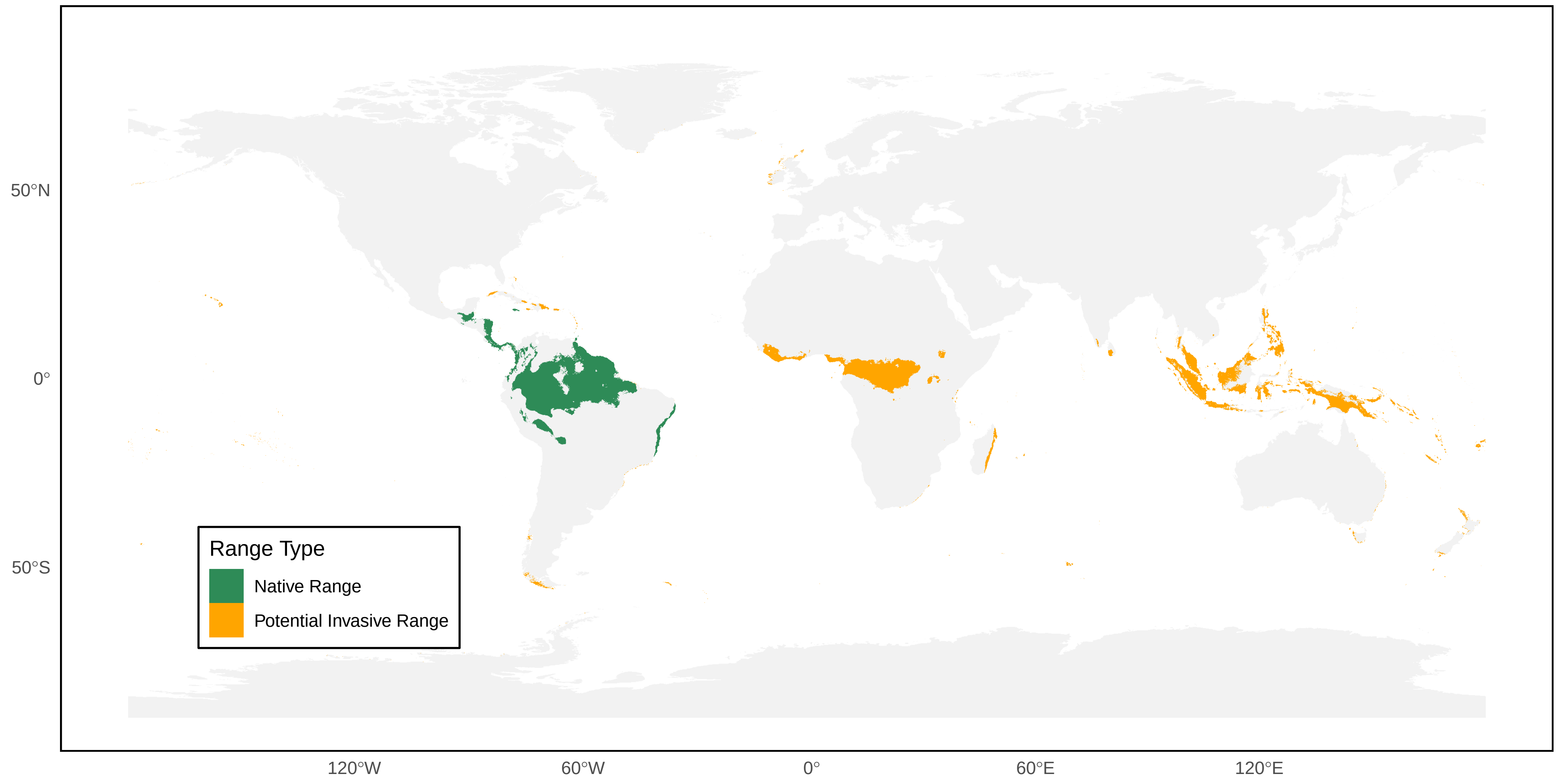

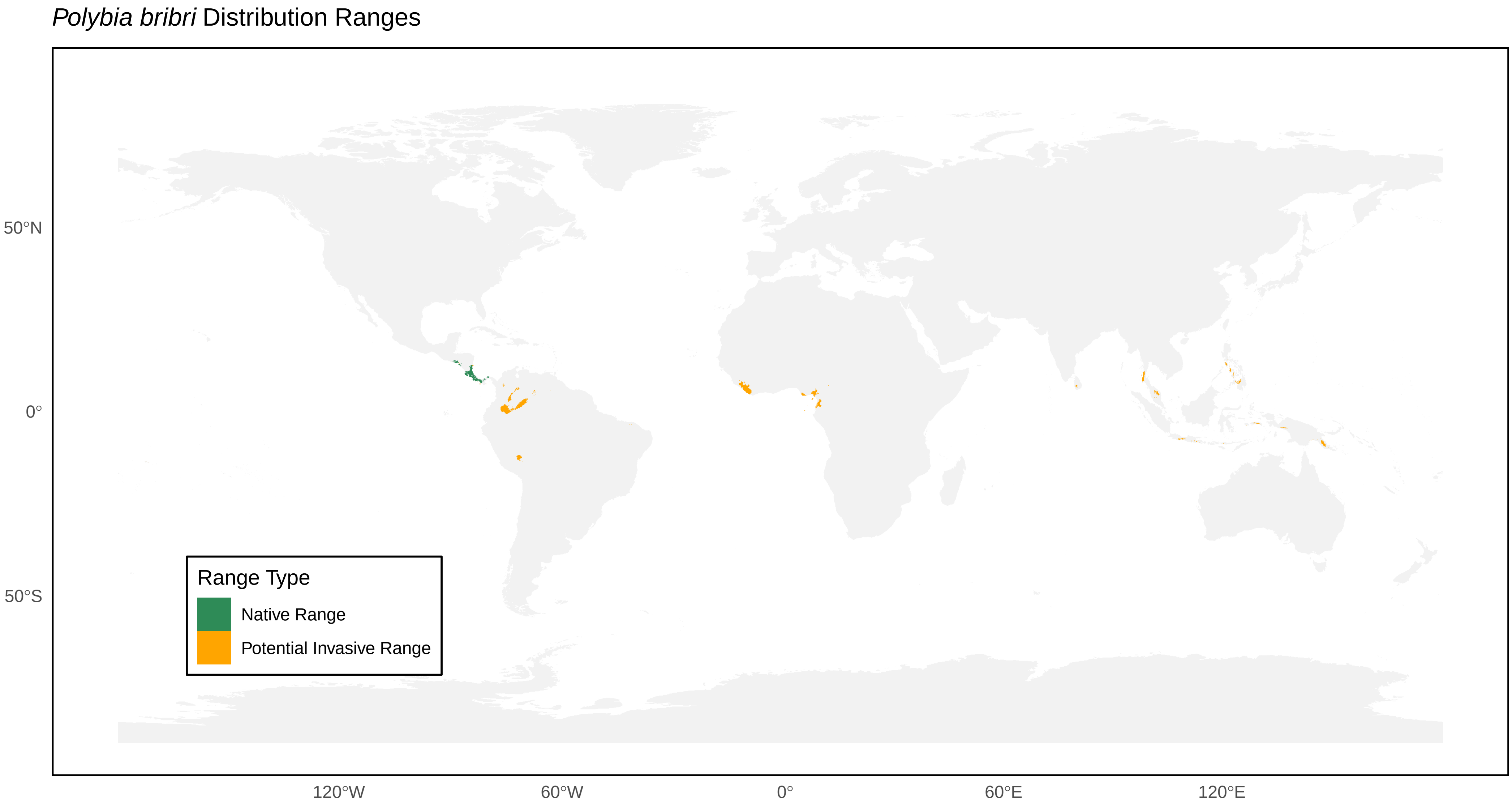

*Polybia bribri* Distribution Ranges
Range Type
Native Range
Potential Invasive Range
50°N
0°
50°S
120°W
60°W
0°
60°E
120°E

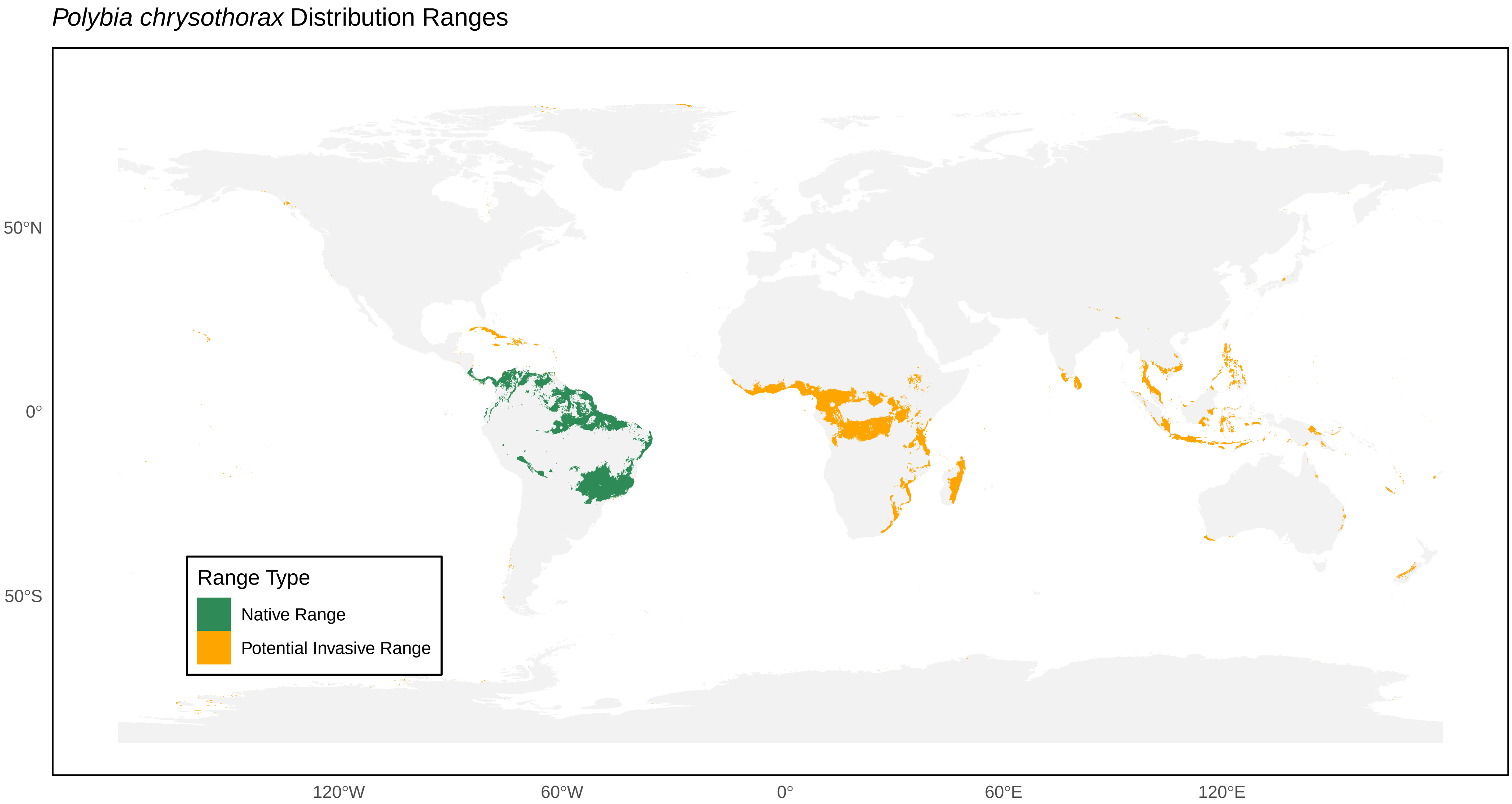
*Polybia chrysothorax* Distribution Ranges
50°N
0°
50°S
120°W
60°W
0°
60°E
120°E
Range Type
Native Range
Potential Invasive Range

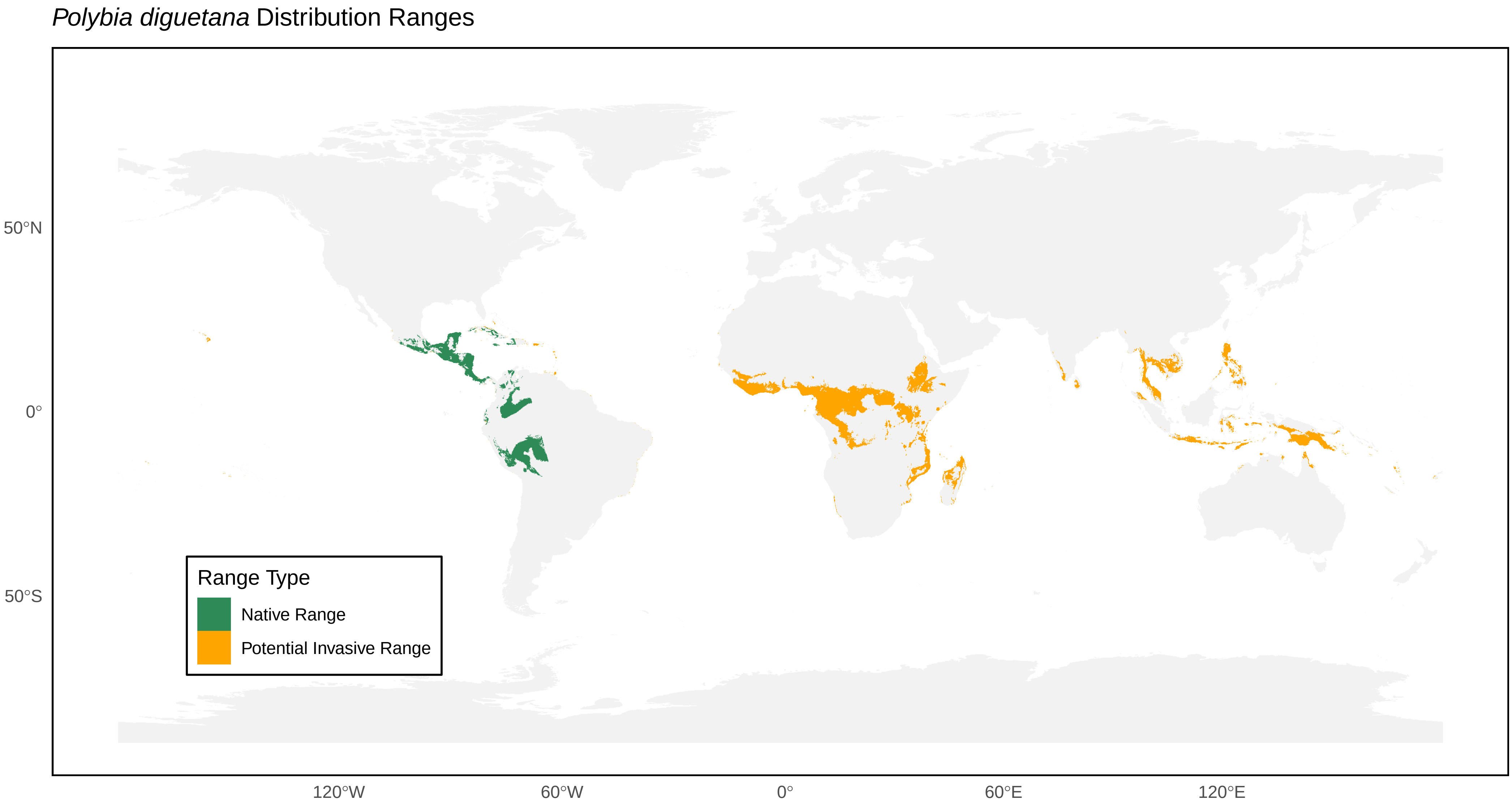
*Polybia diguetana* Distribution Ranges
50°N
0°
50°S
120°W
60°W
0°
60°E
120°E
Range Type
Native Range
Potential Invasive Range

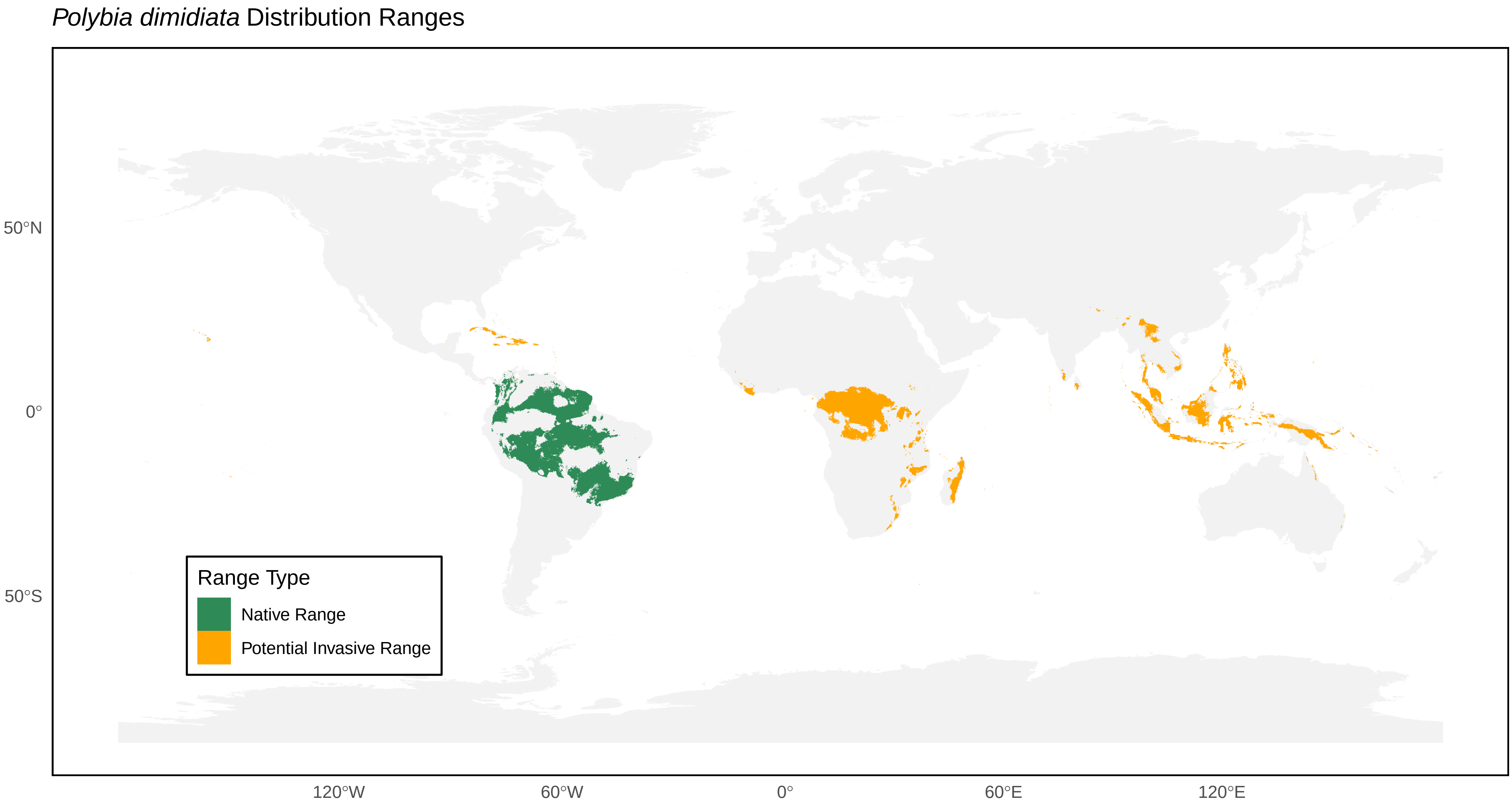

*Polybia dimidiata* Distribution Ranges
50°N
0°
50°S
120°W
60°W
0°
60°E
120°E
Range Type
Native Range
Potential Invasive Range

*Polybia emaciata* Distribution Ranges

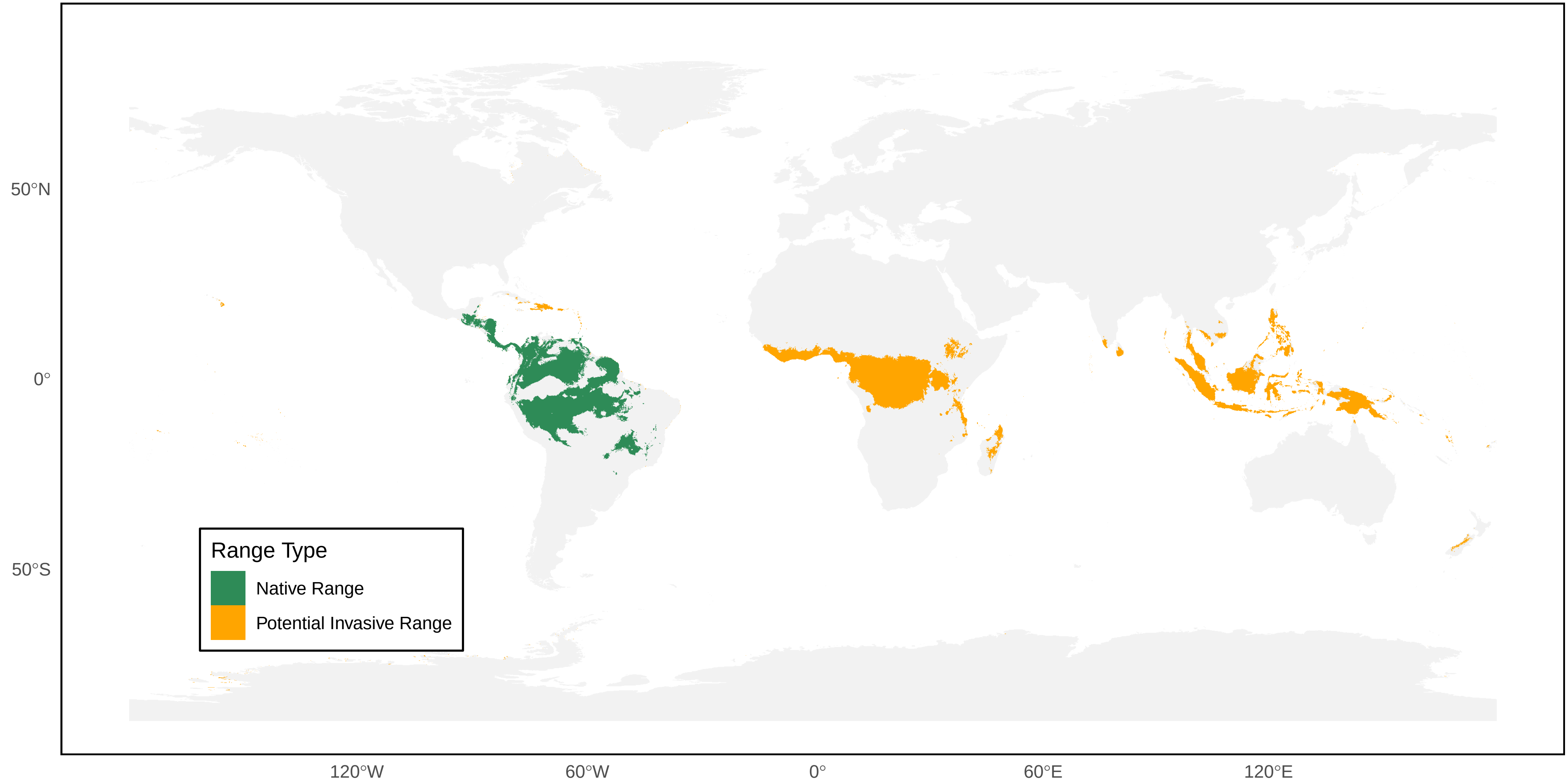

*Polybia erythrothoraxla* Distribution Ranges

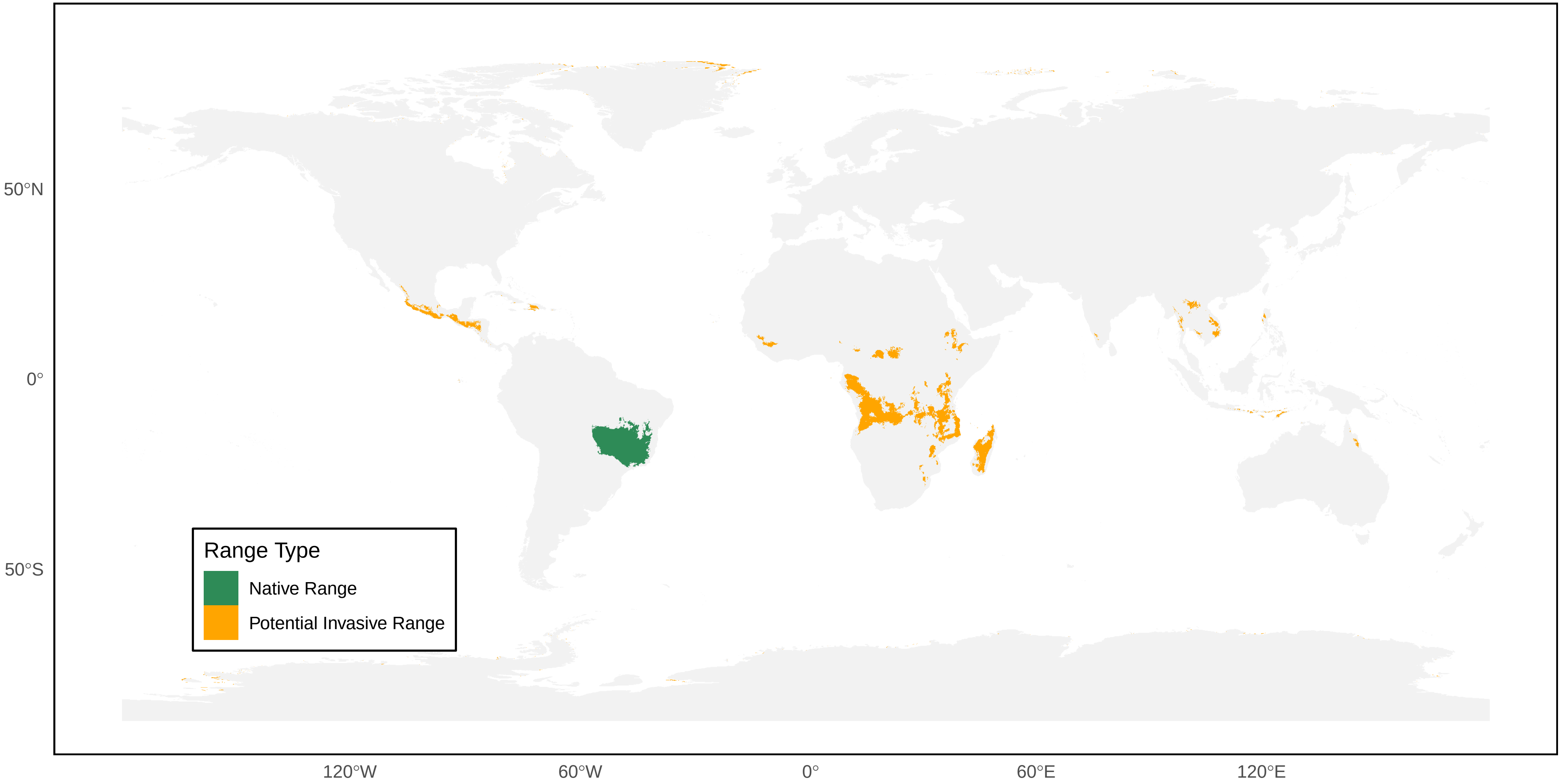

*Polybia fastidiosuscula* Distribution Ranges

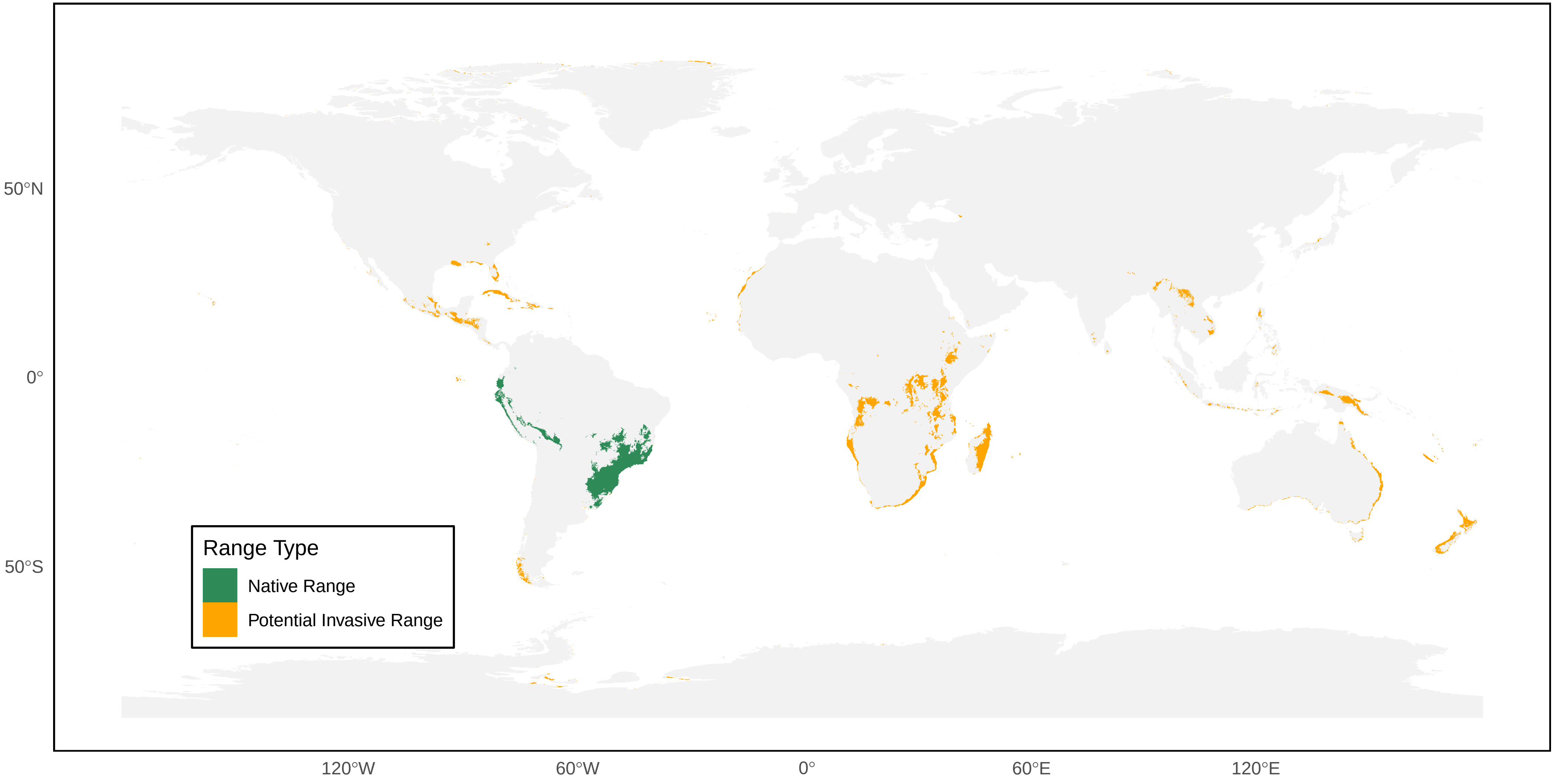

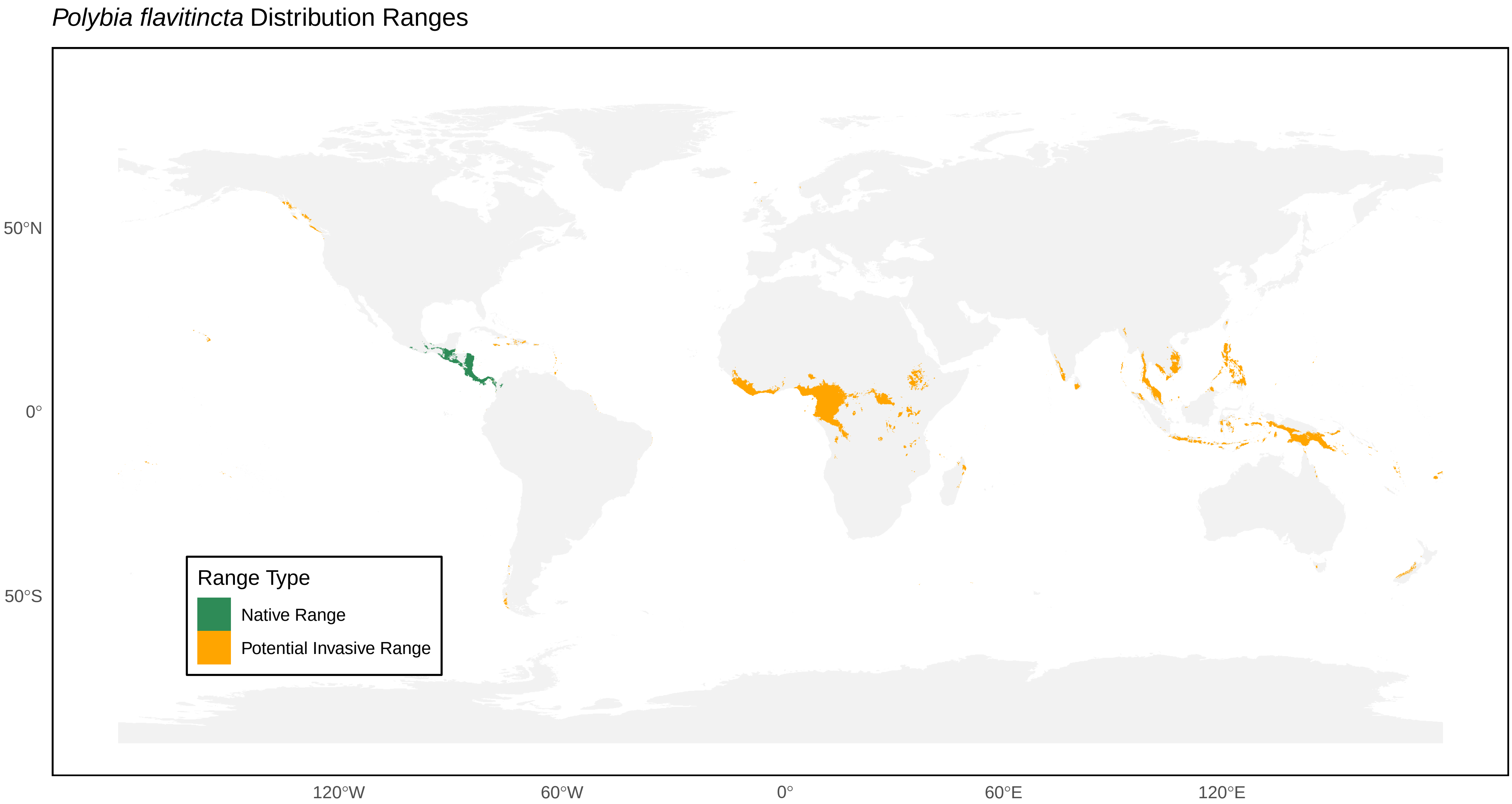

*Polybia flavitincta* Distribution Ranges
50°N
0°
50°S
120°W
60°W
0°
60°E
120°E
Range Type
Native Range
Potential Invasive Range

*Polybia ignobilis* Distribution Ranges

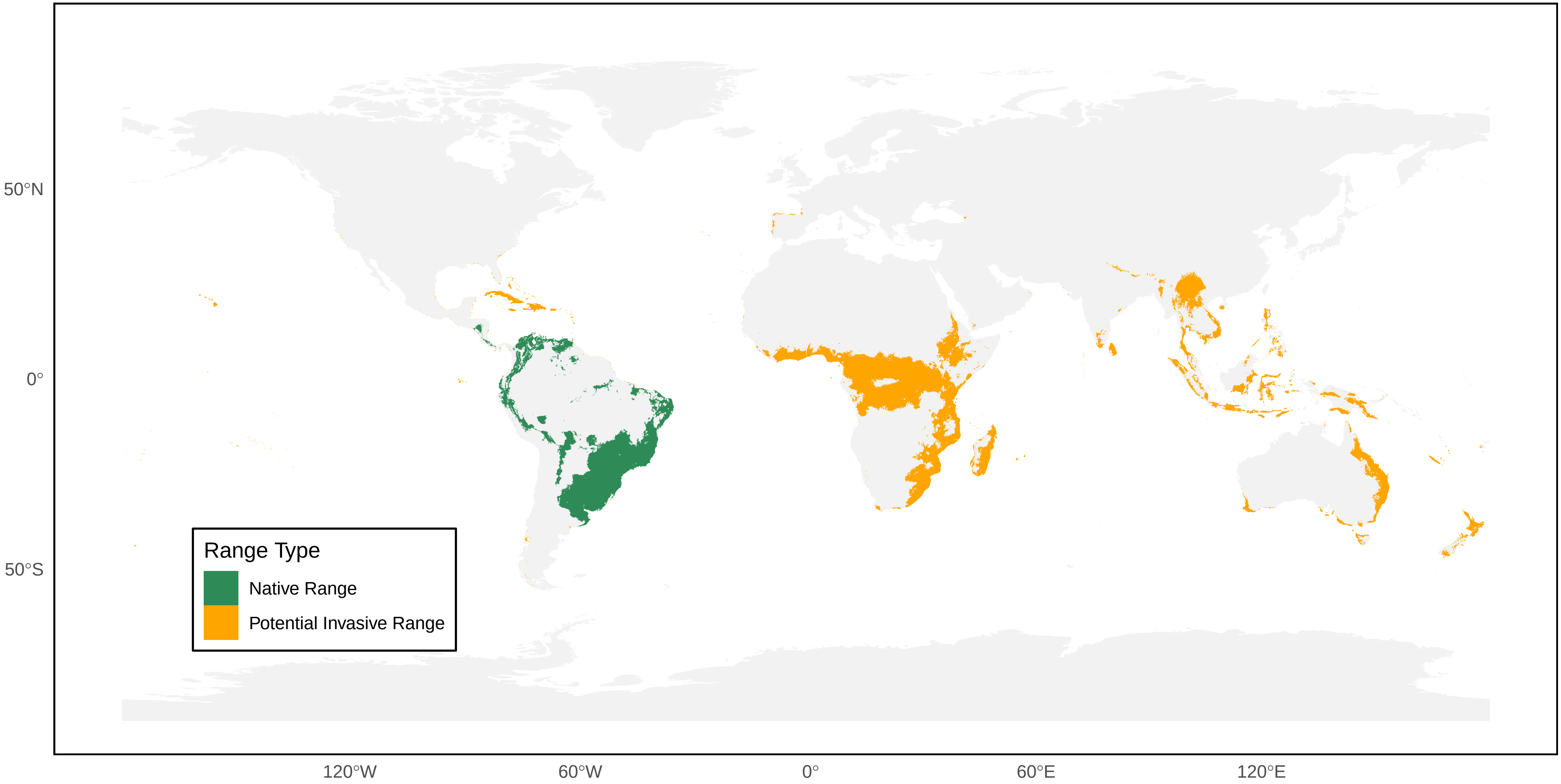

*Polybia jurinei* Distribution Ranges

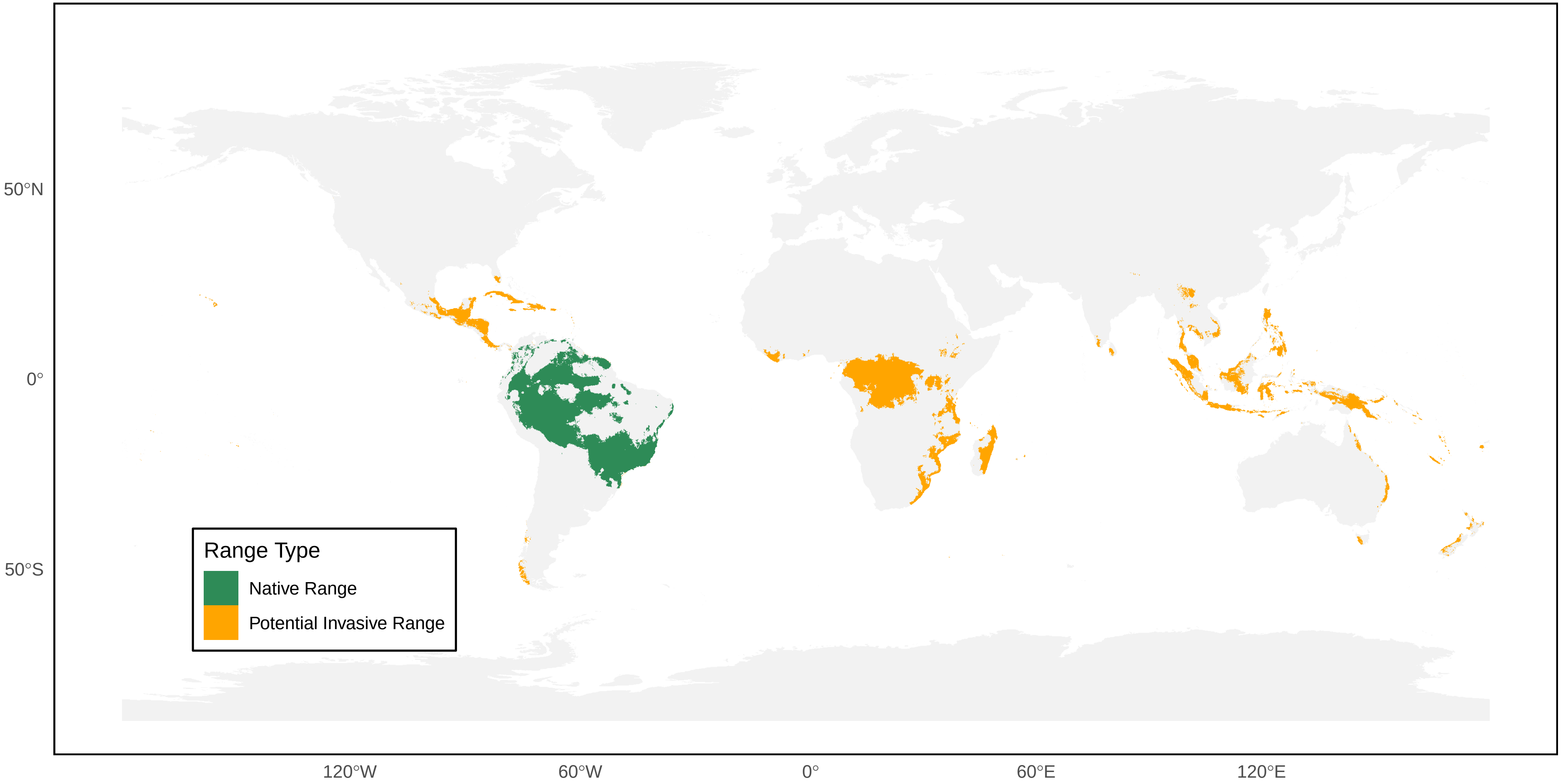

*Polybia liliacea* Distribution Ranges

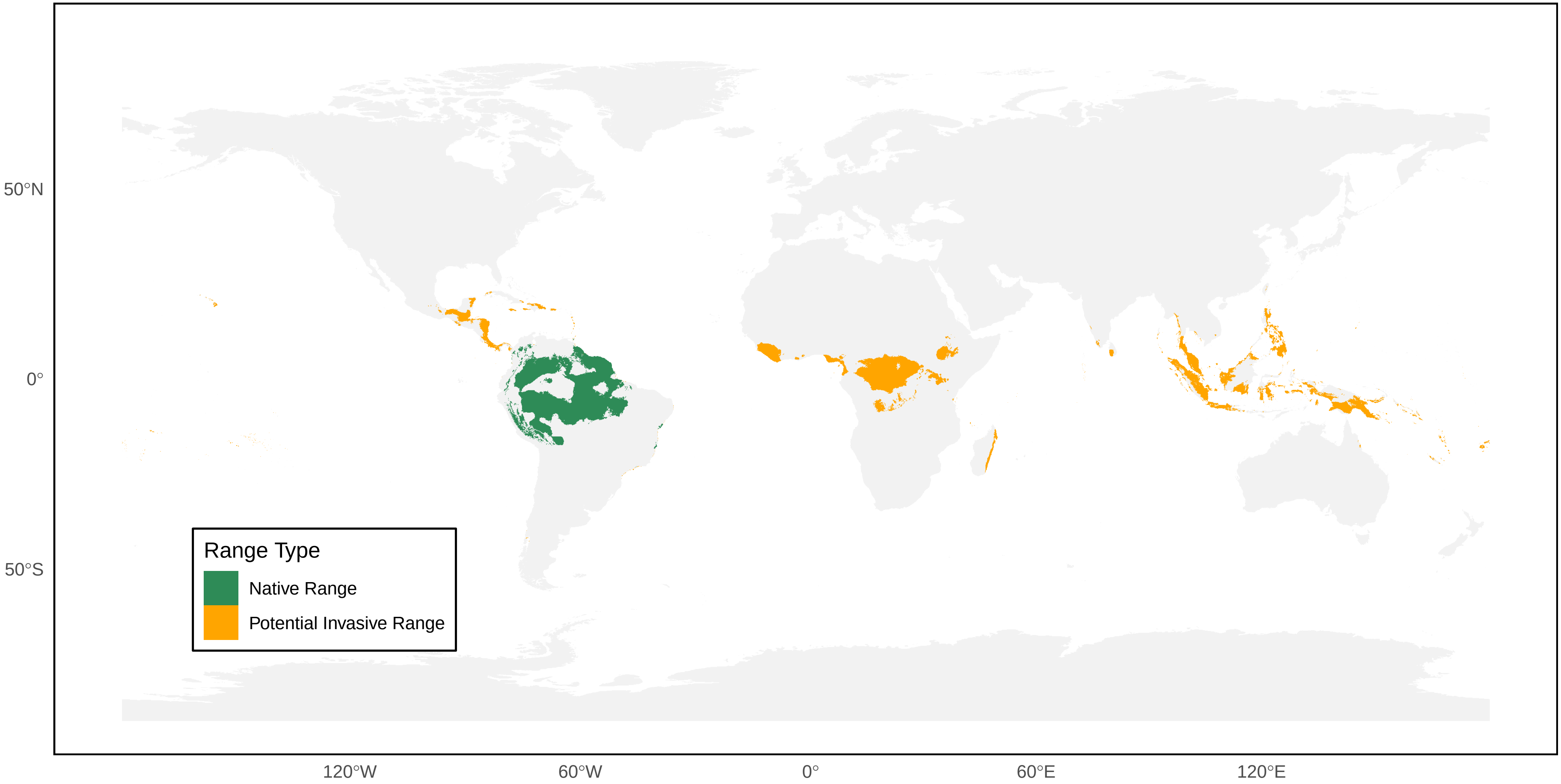

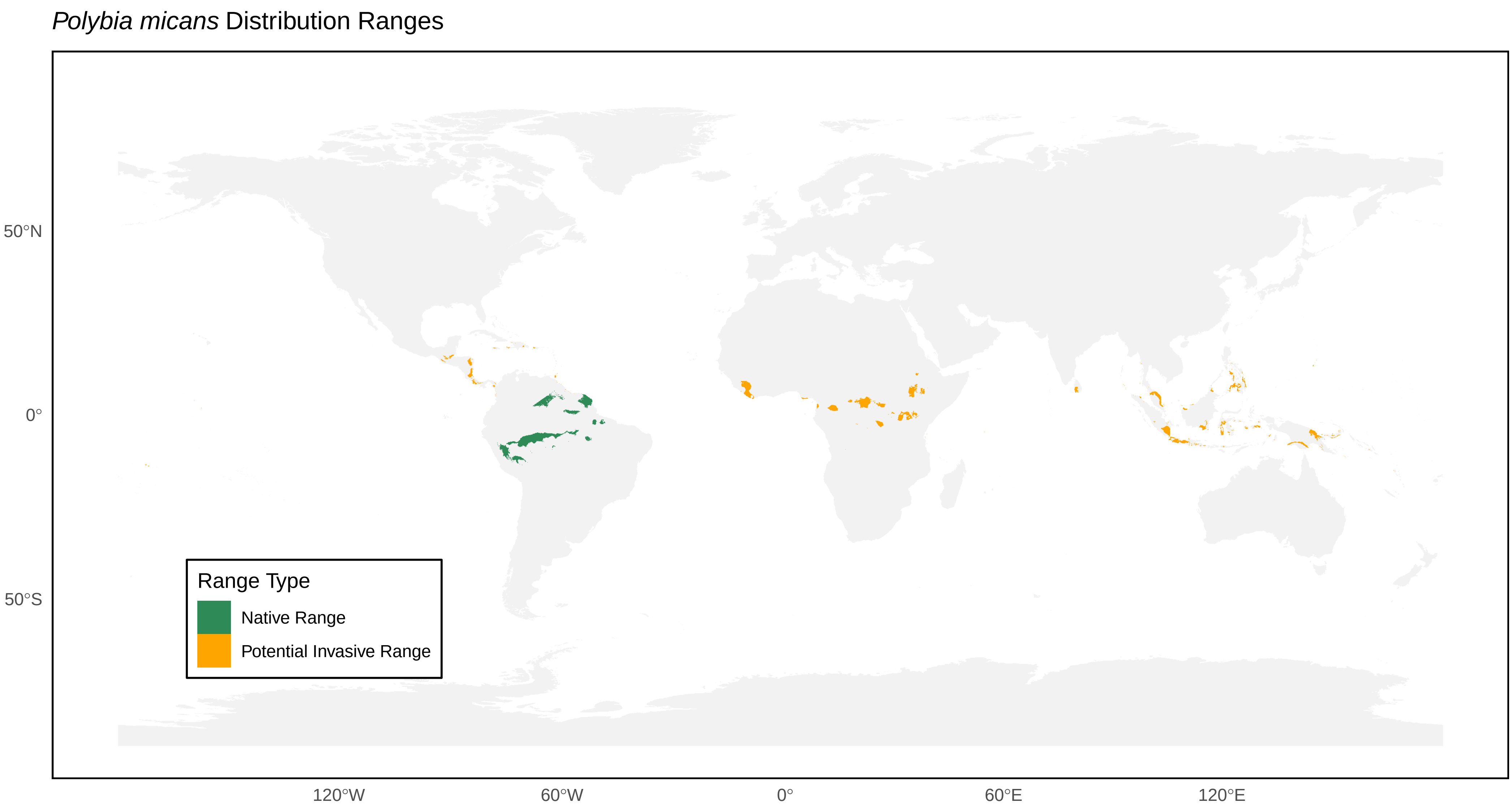

*Polybia micans* Distribution Ranges
50°N
0°
50°S
120°W
60°W
0°
60°E
120°E
Range Type
Native Range
Potential Invasive Range

*Polybia nidulatrix* Distribution Ranges

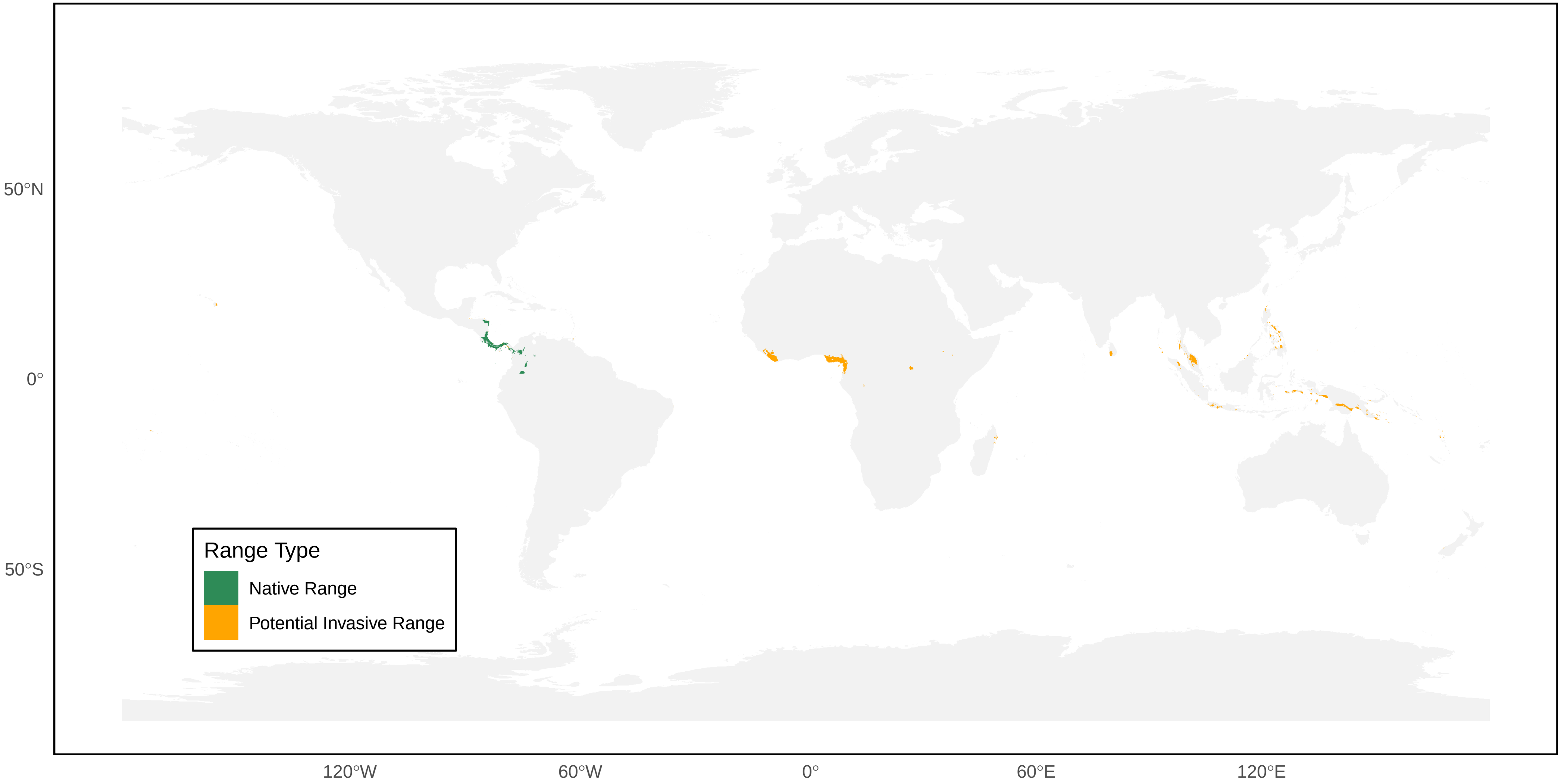

*Polybia occidentalis* Distribution Ranges

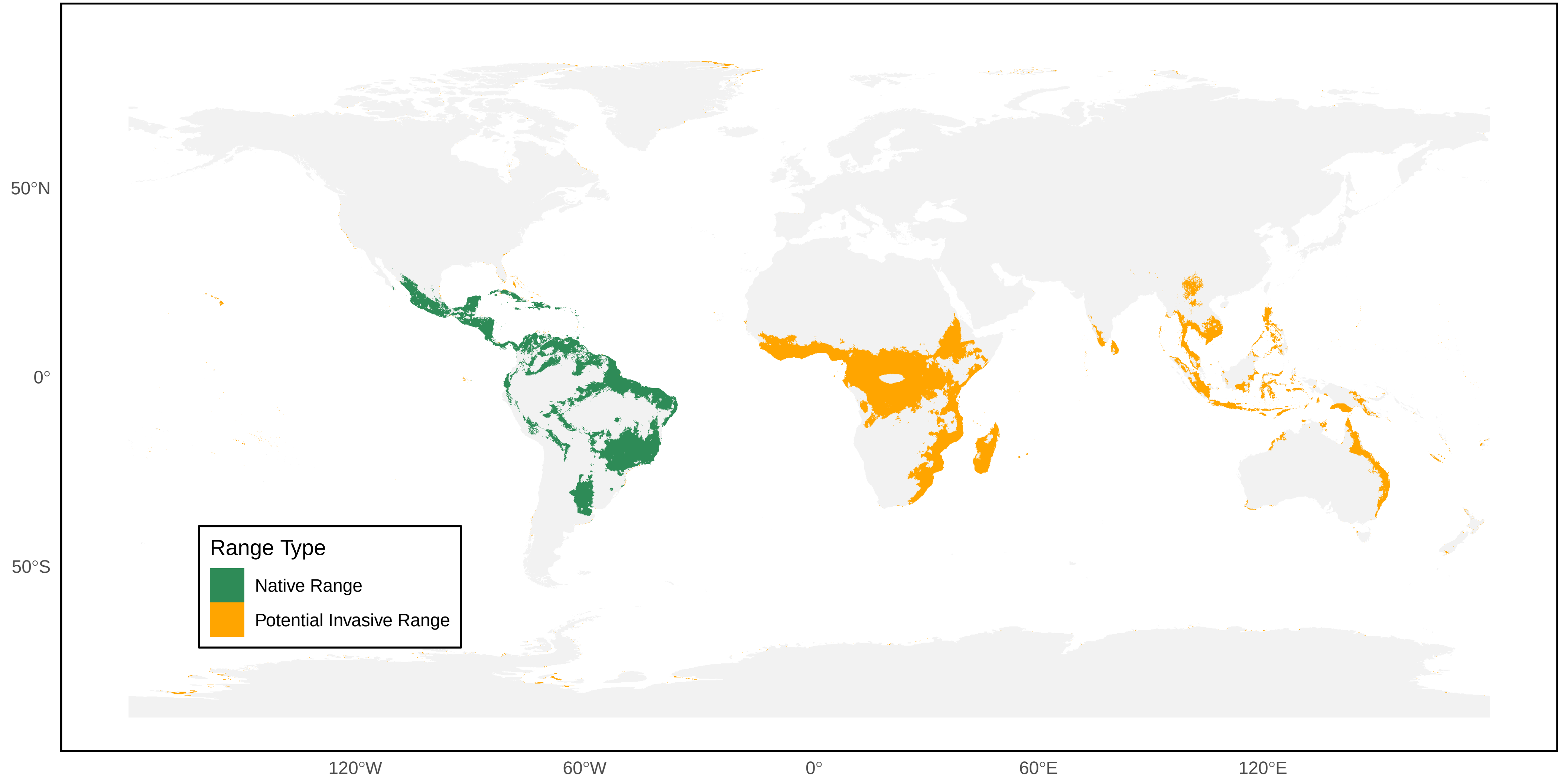

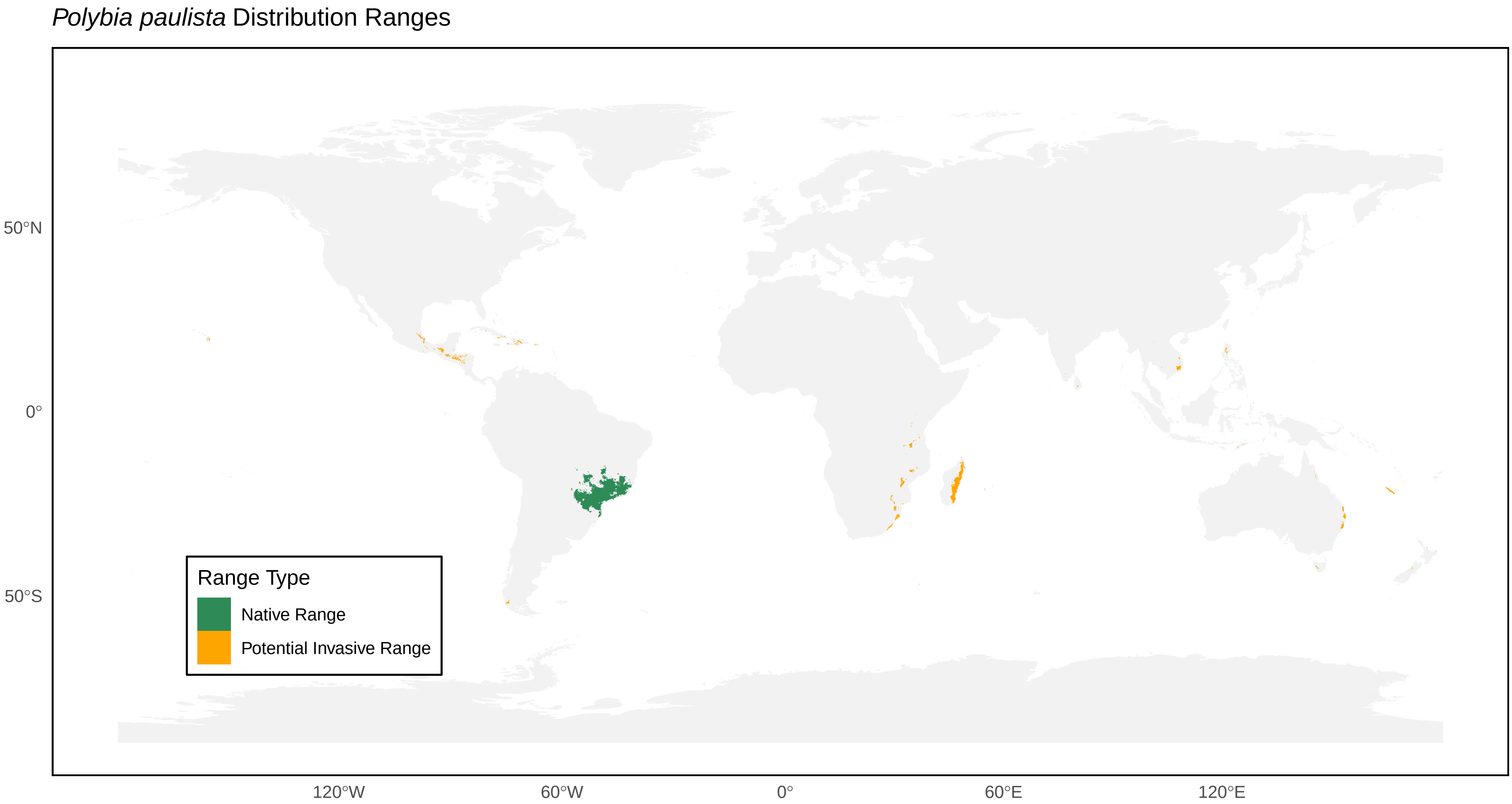
*Polybia paulista* Distribution Ranges
50°N
0°
50°S
120°W
60°W
0°
60°E
120°E
Range Type
Native Range
Potential Invasive Range

*Polybia platycephala* Distribution Ranges

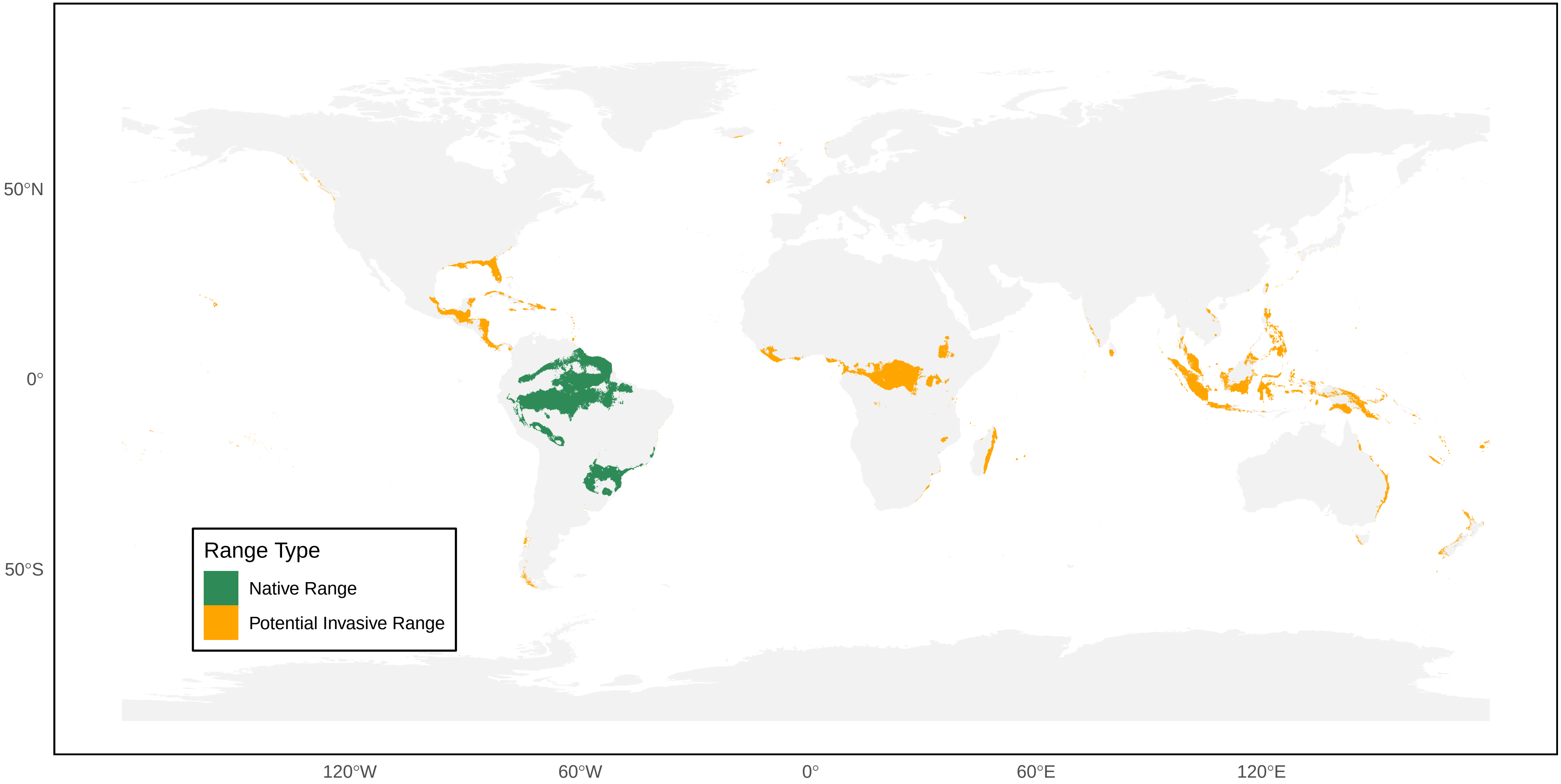

*Polybia plebeja* Distribution Ranges

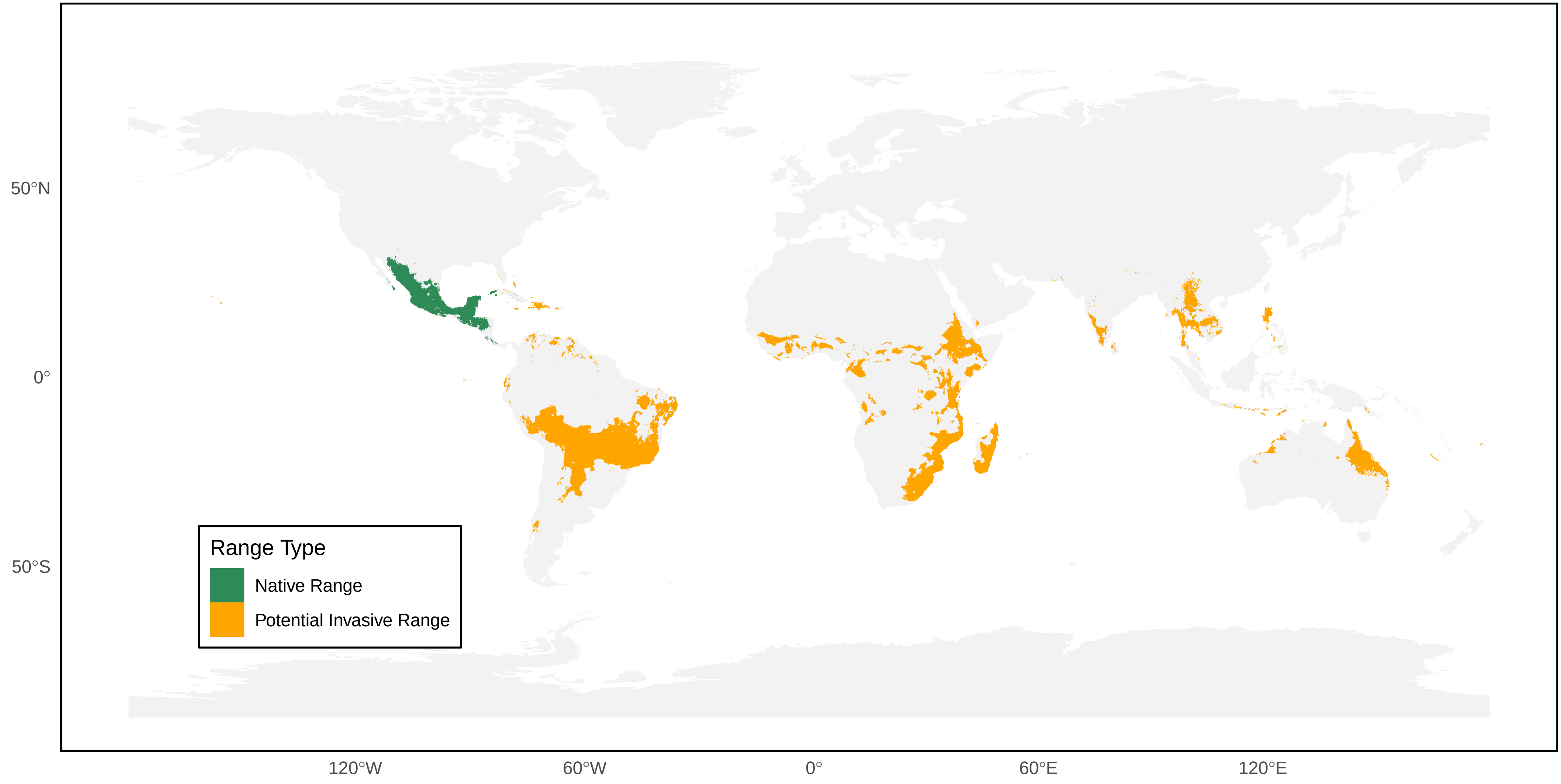

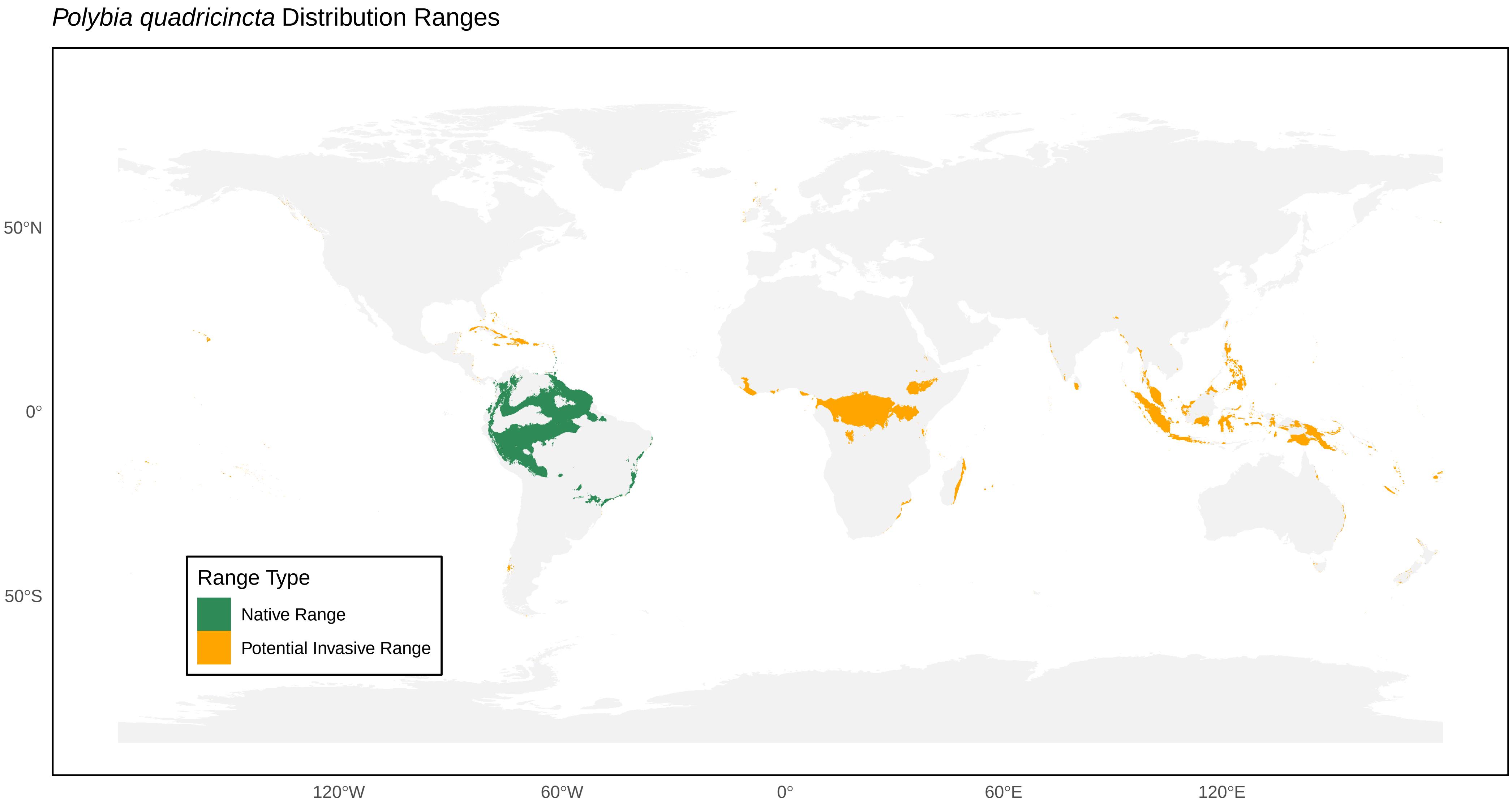
*Polybia quadricincta* Distribution Ranges
50°N
0°
50°S
120°W
60°W
0°
60°E
120°E
Range Type
Native Range
Potential Invasive Range

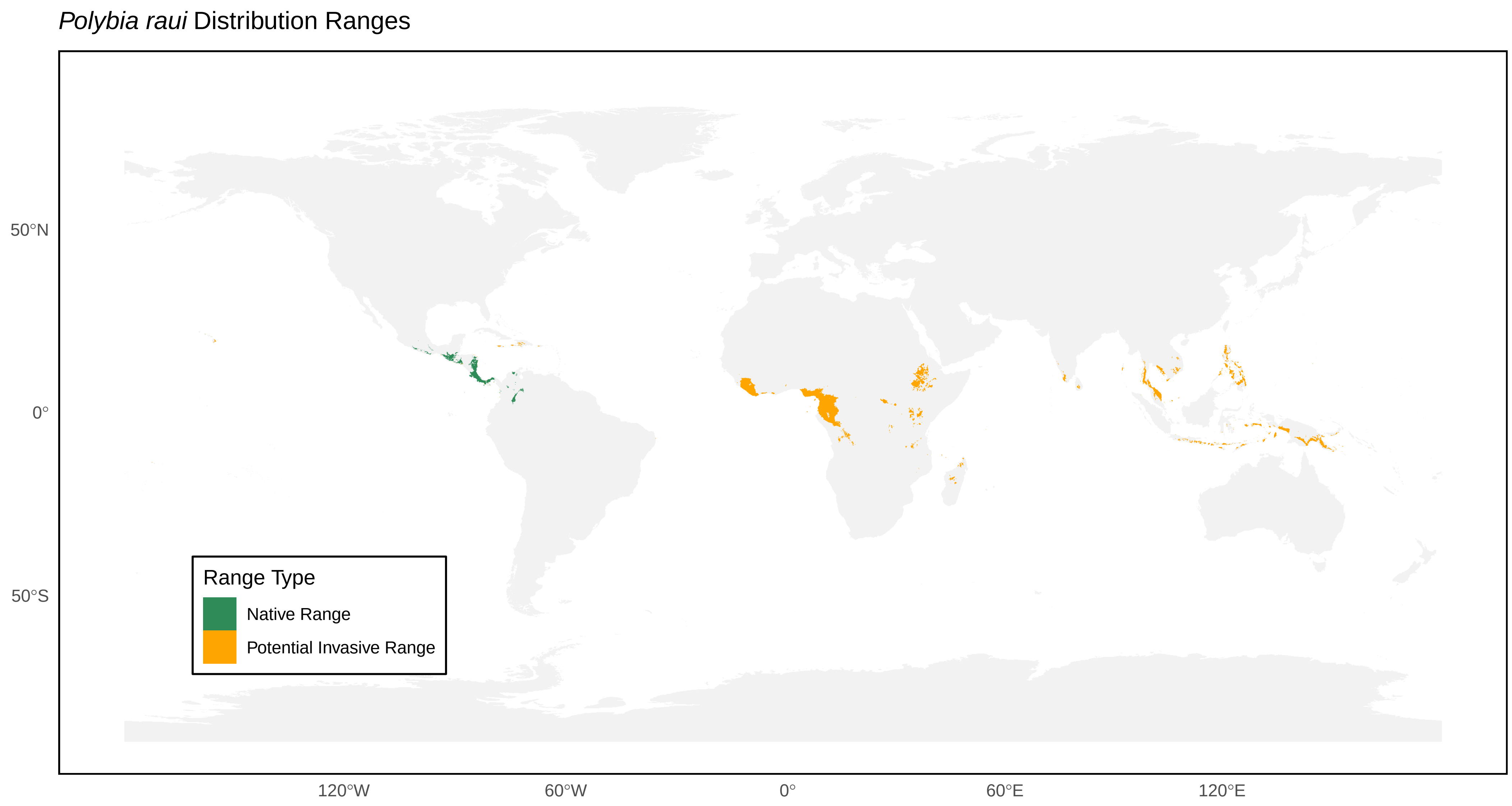

*Polybia raui* Distribution Ranges
50°N
0°
50°S
120°W
60°W
0°
60°E
120°E
Range Type
Native Range
Potential Invasive Range

*Polybia rejecta* Distribution Ranges

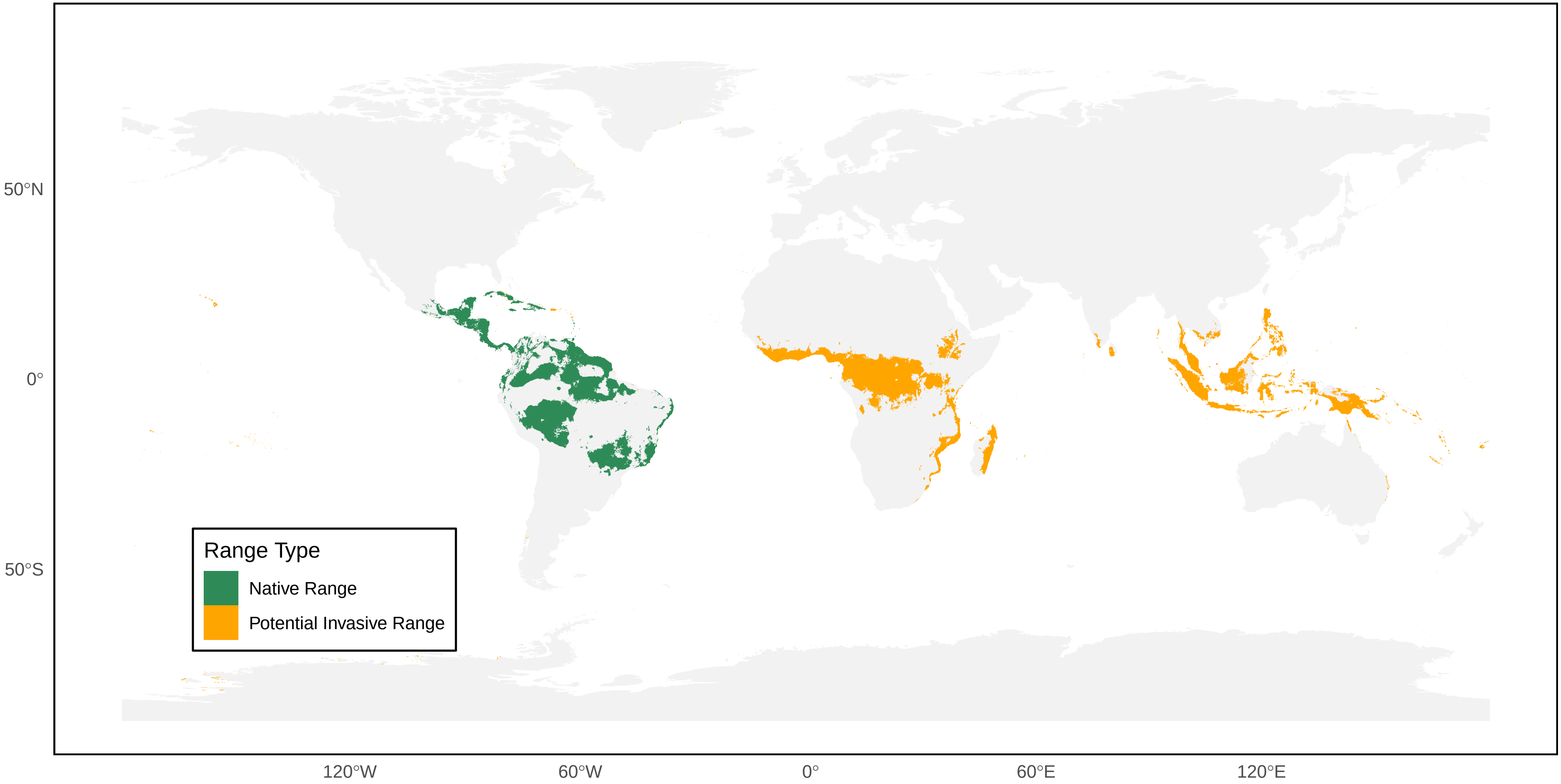

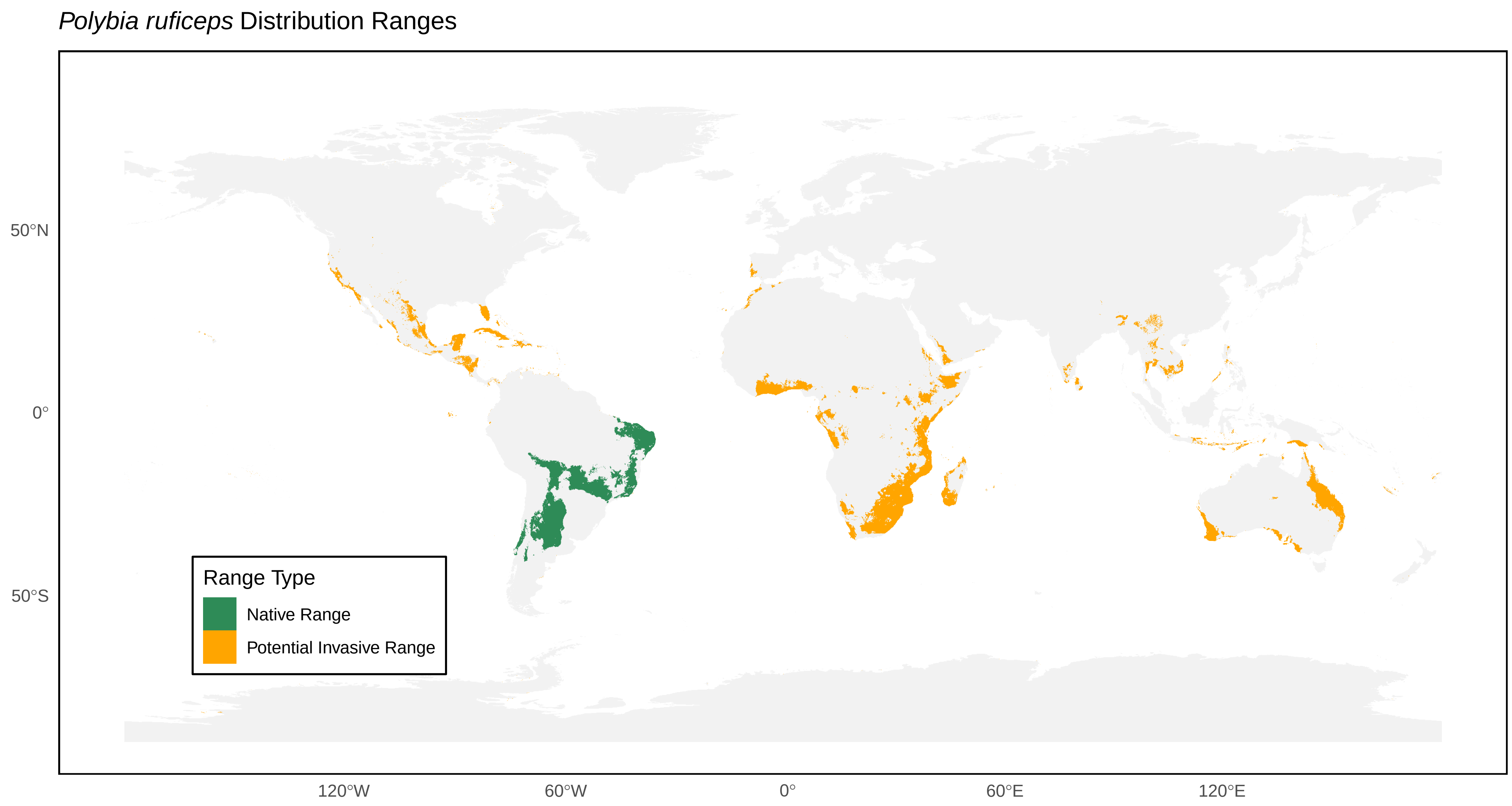

*Polybia ruficeps* Distribution Ranges
50°N
0°
50°S
120°W
60°W
0°
60°E
120°E
Range Type
Native Range
Potential Invasive Range

*Polybia scrobalis* Distribution Ranges

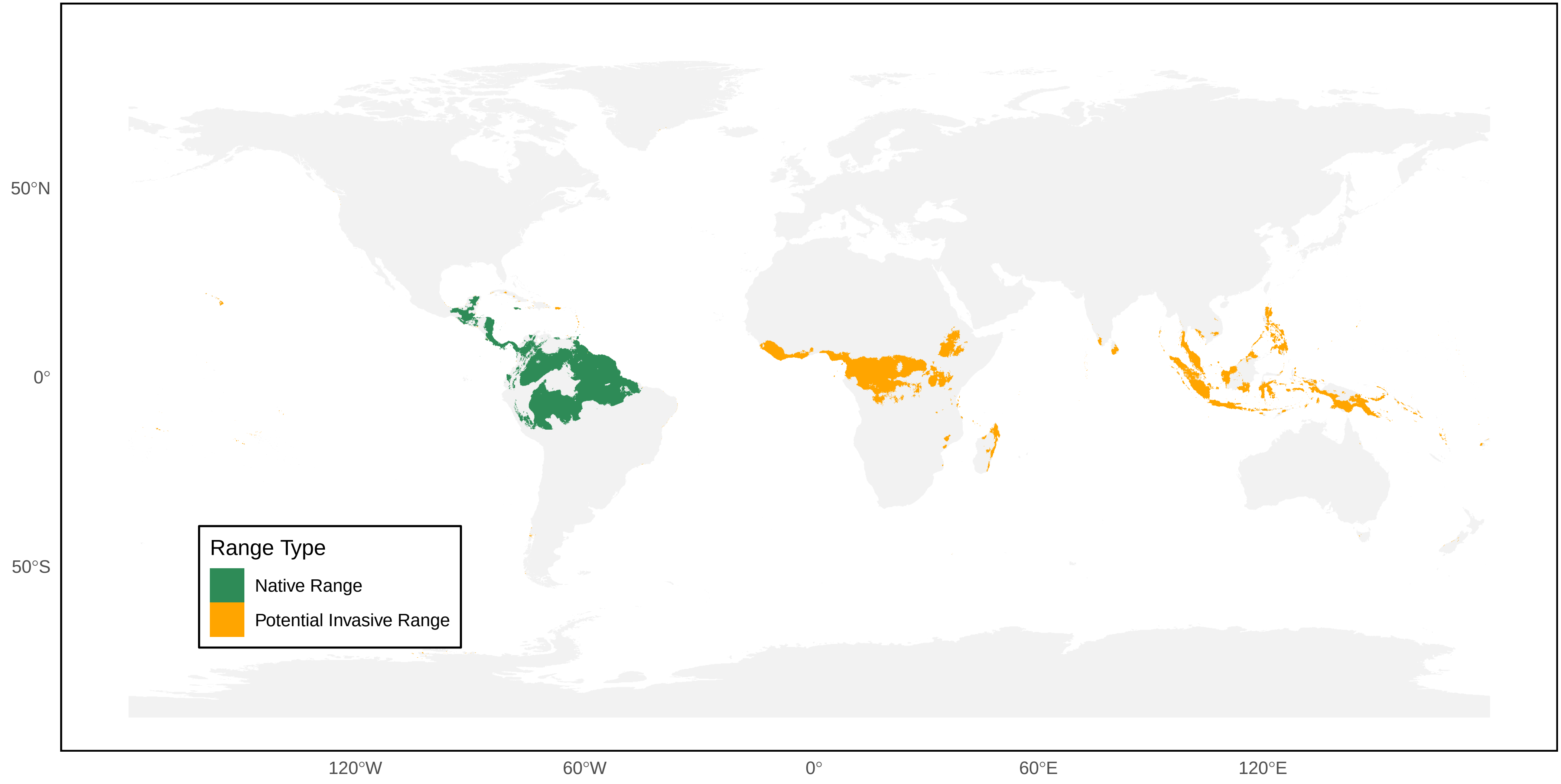

*Polybia scutellaris* Distribution Ranges

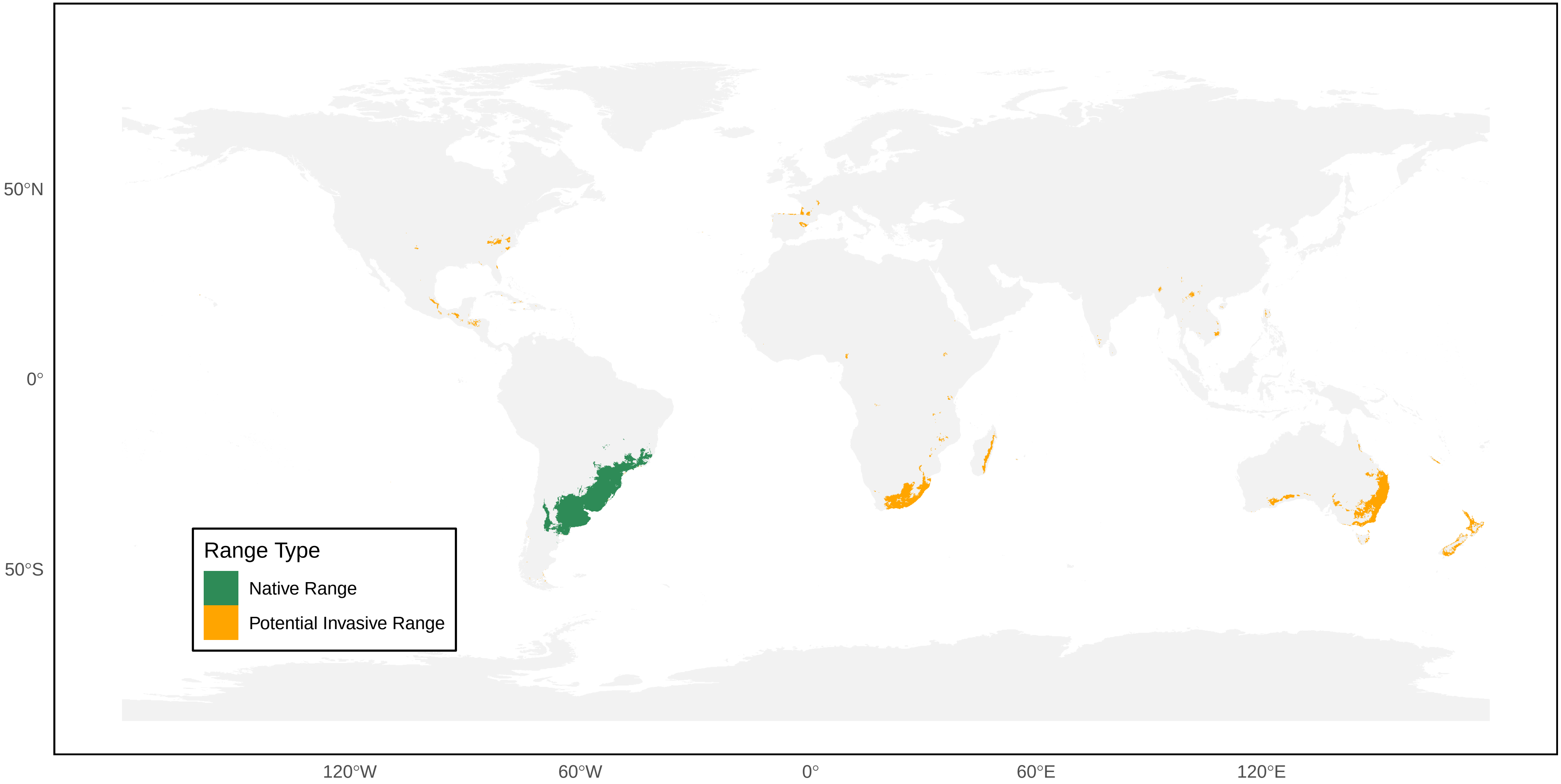

*Polybia selvana* Distribution Ranges

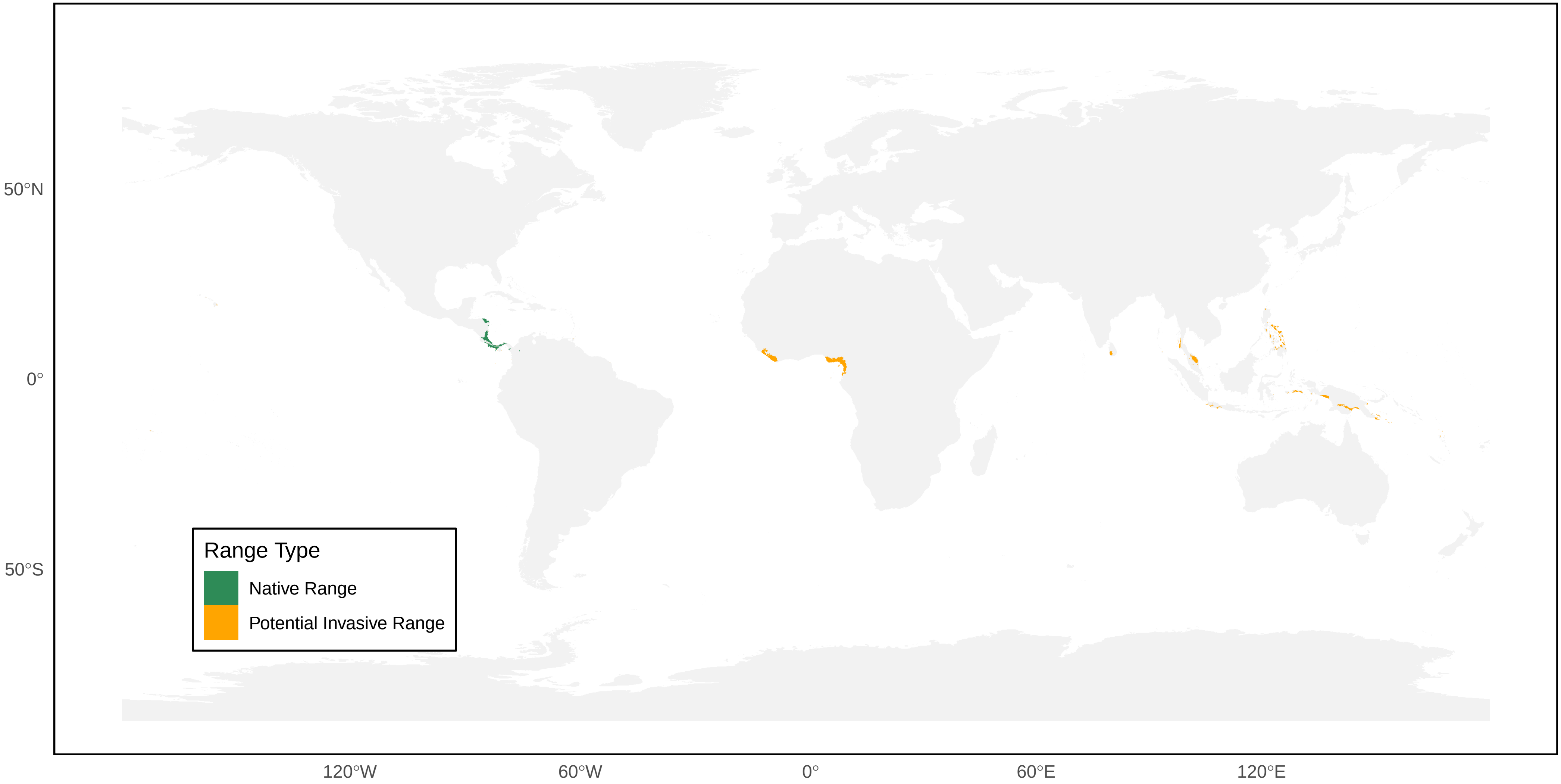

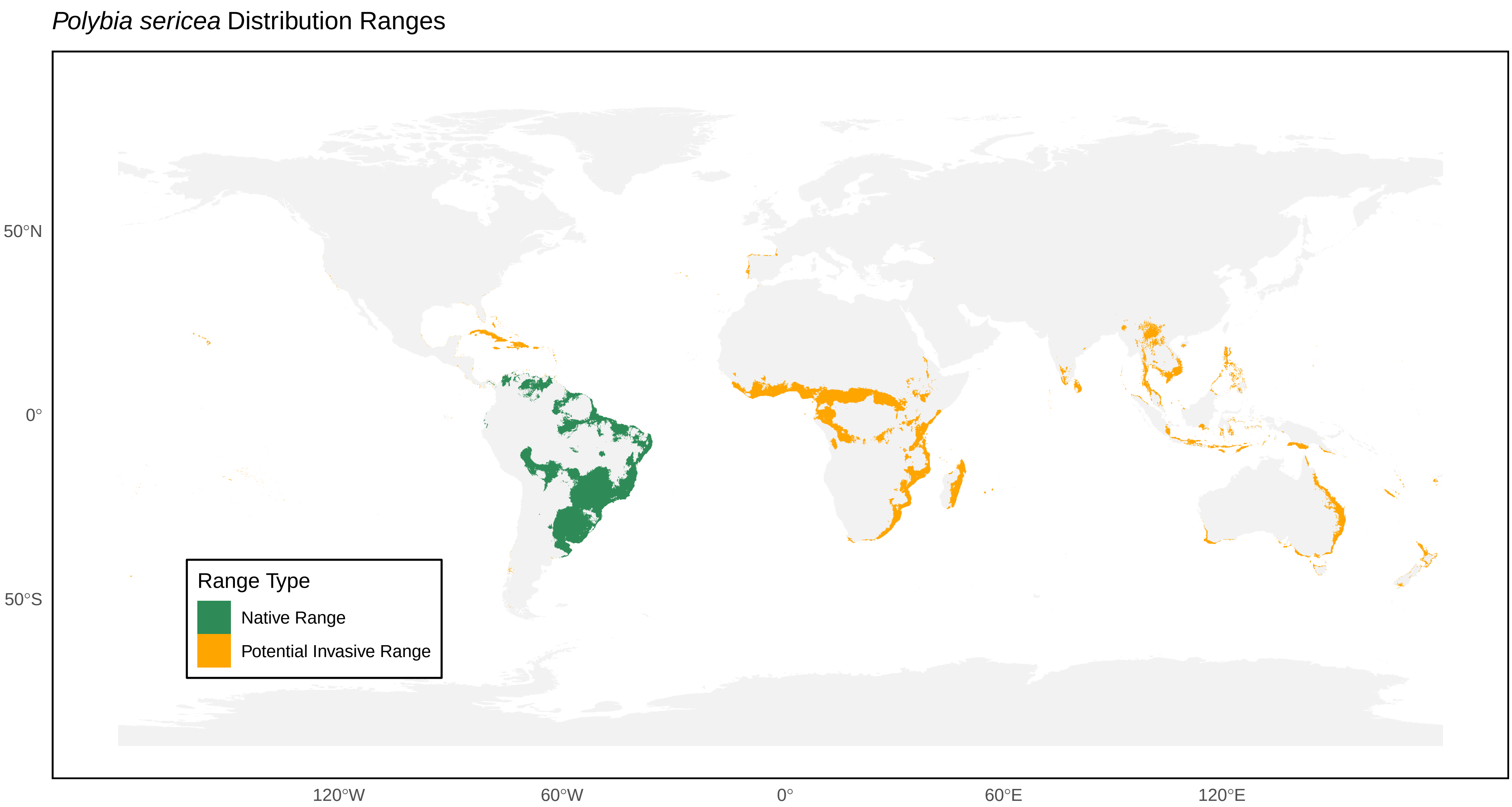
*Polybia sericea* Distribution Ranges
50°N
0°
50°S
120°W
60°W
0°
60°E
120°E
Range Type
Native Range
Potential Invasive Range

*Polybia simillima* Distribution Ranges

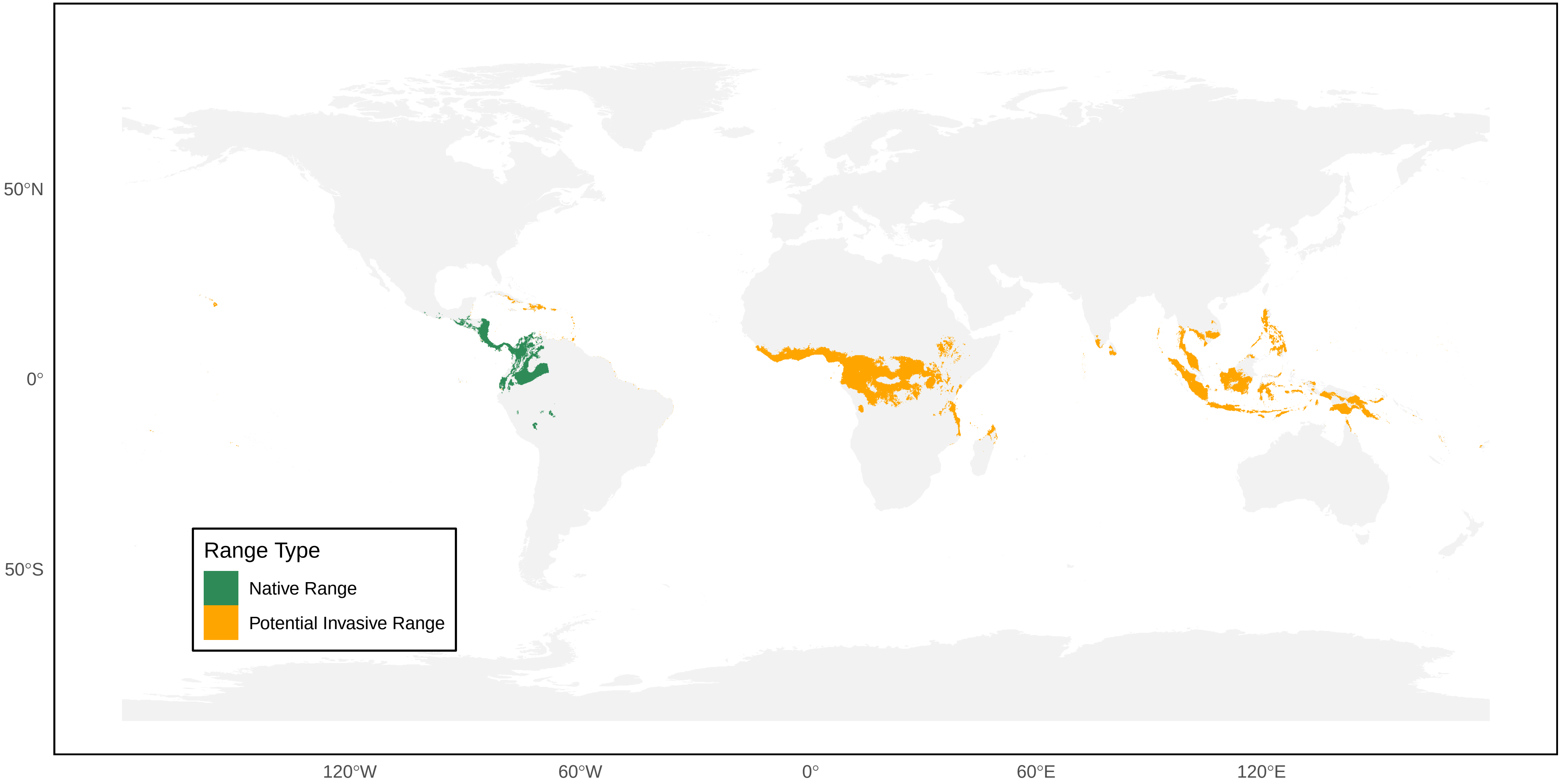

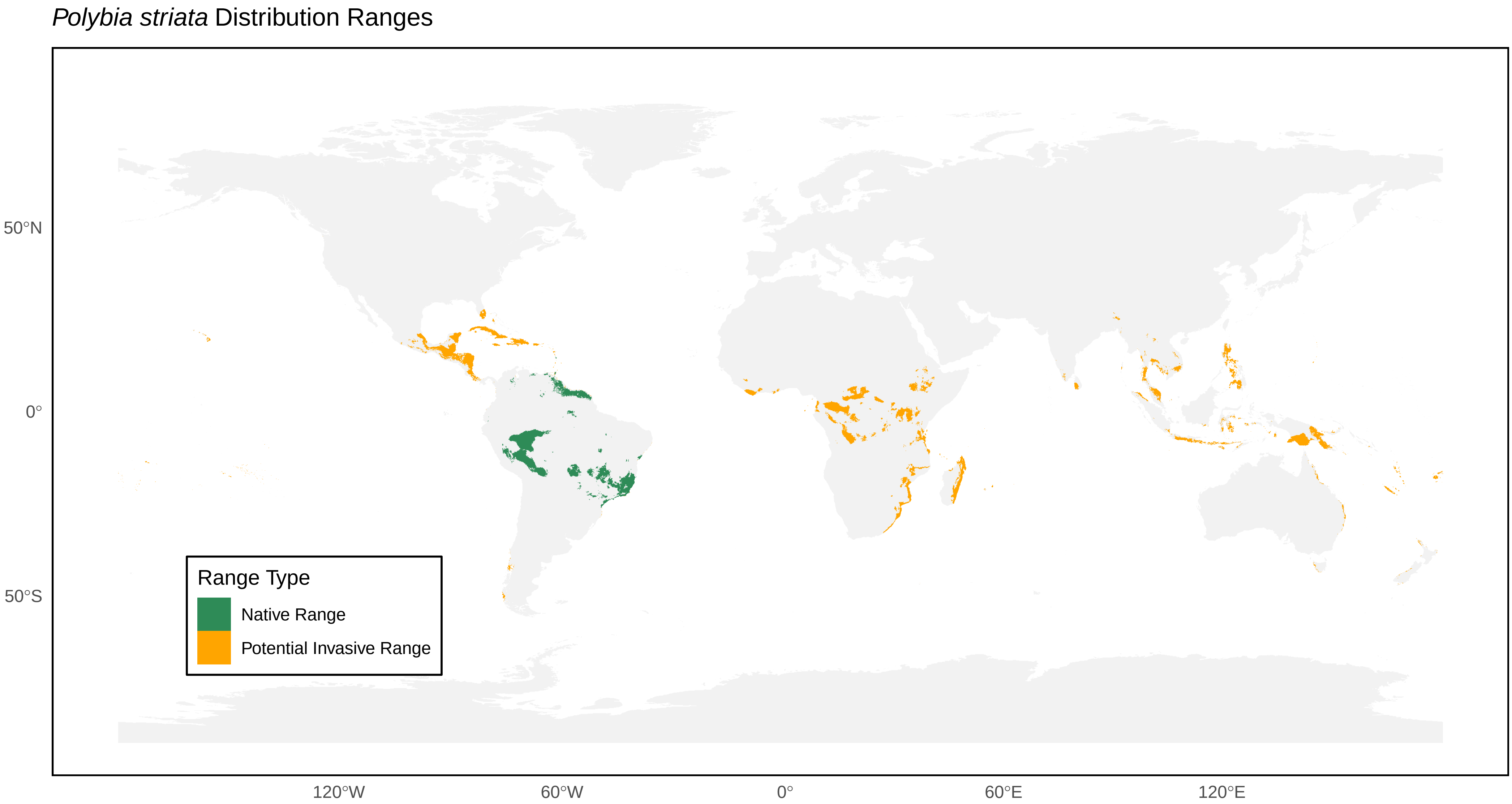
*Polybia striata* Distribution Ranges
50°N
0°
50°S
120°W
60°W
0°
60°E
120°E
Range Type
Native Range
Potential Invasive Range

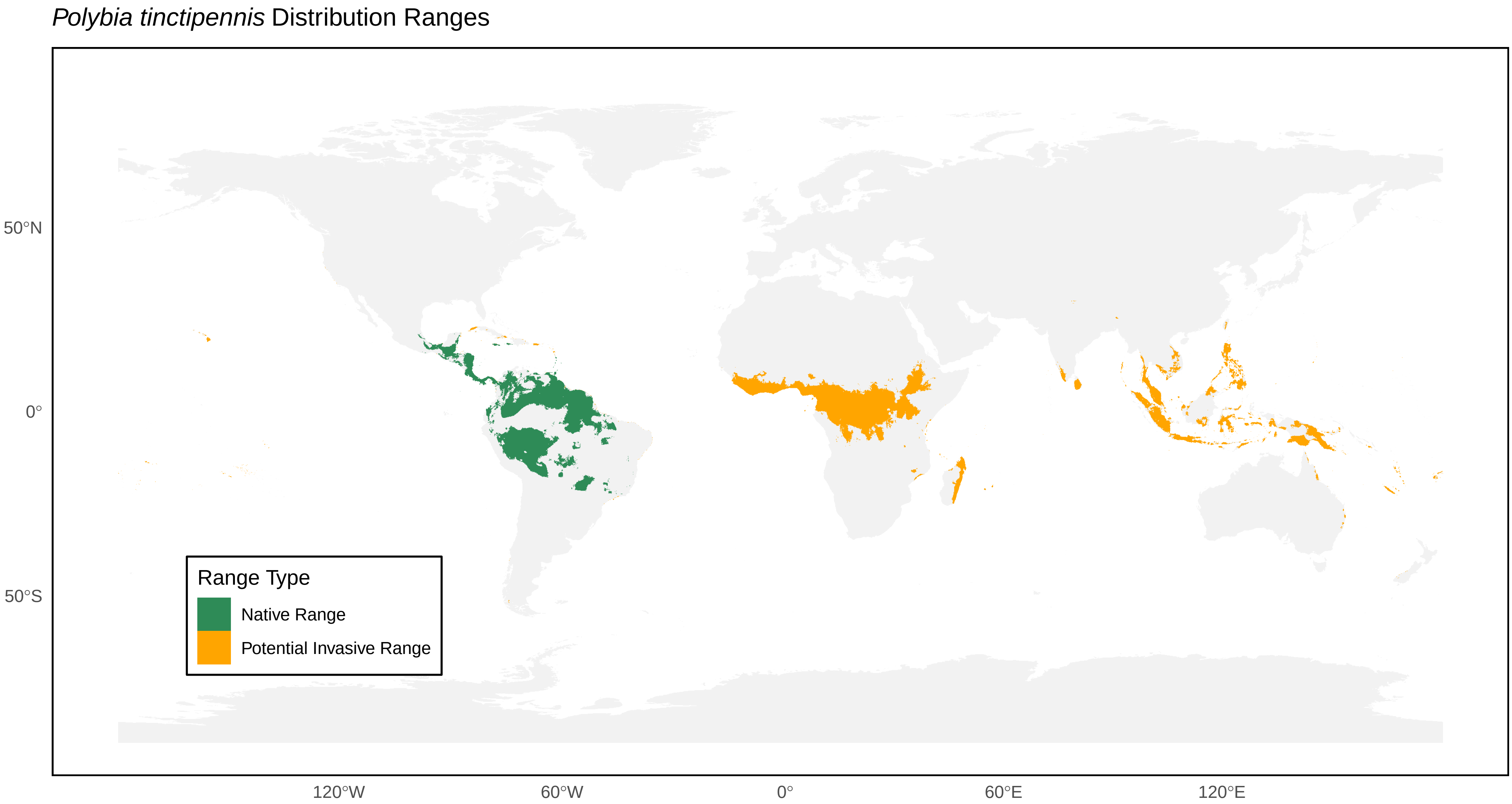

*Polybia tinctipennis* Distribution Ranges
50°N
0°
50°S
120°W
60°W
0°
60°E
120°E
Range Type
Native Range
Potential Invasive Range

*Polybioides raphigastra* Distribution Ranges

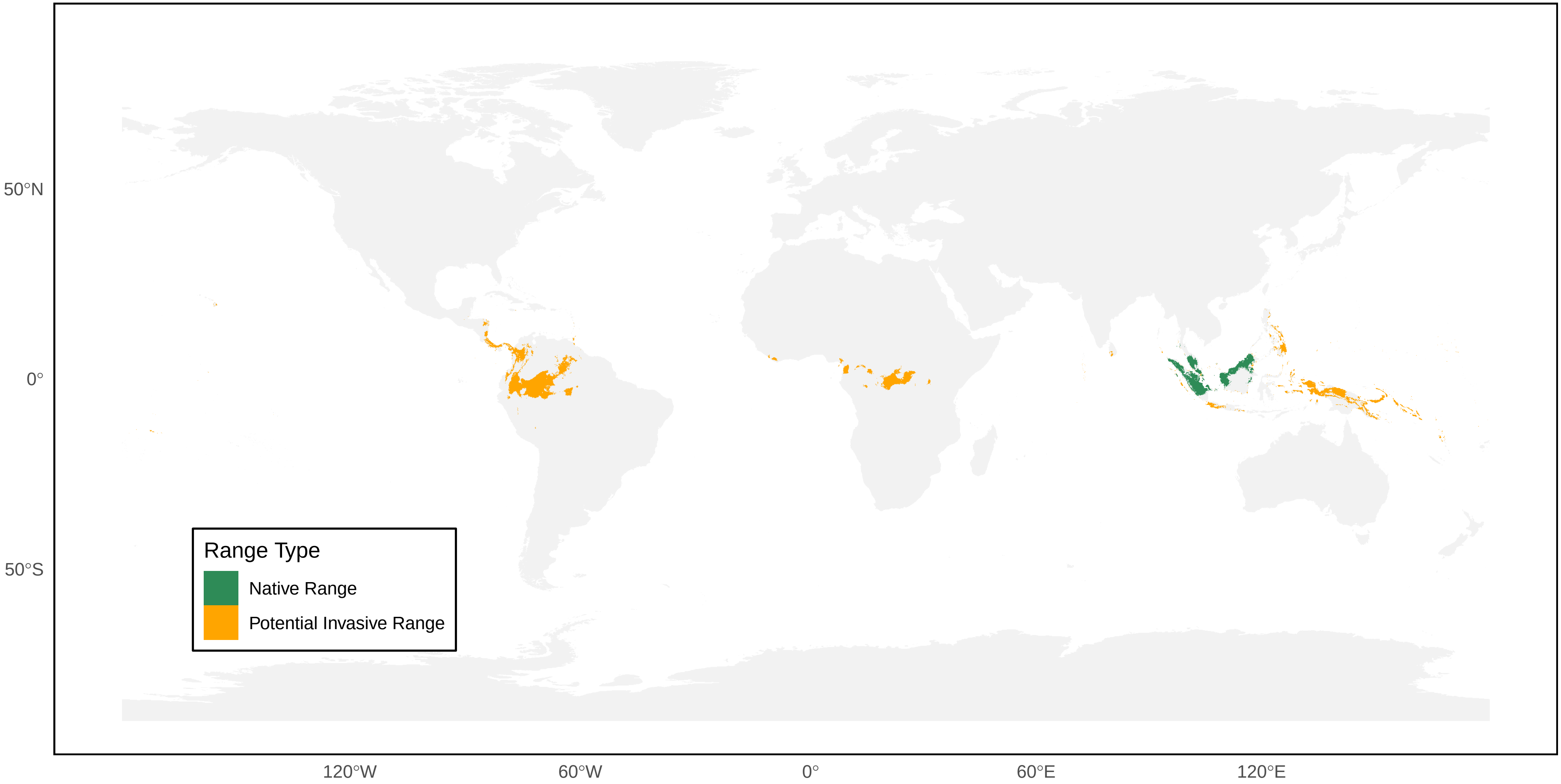

*Protonectarina sylveirae* Distribution Ranges

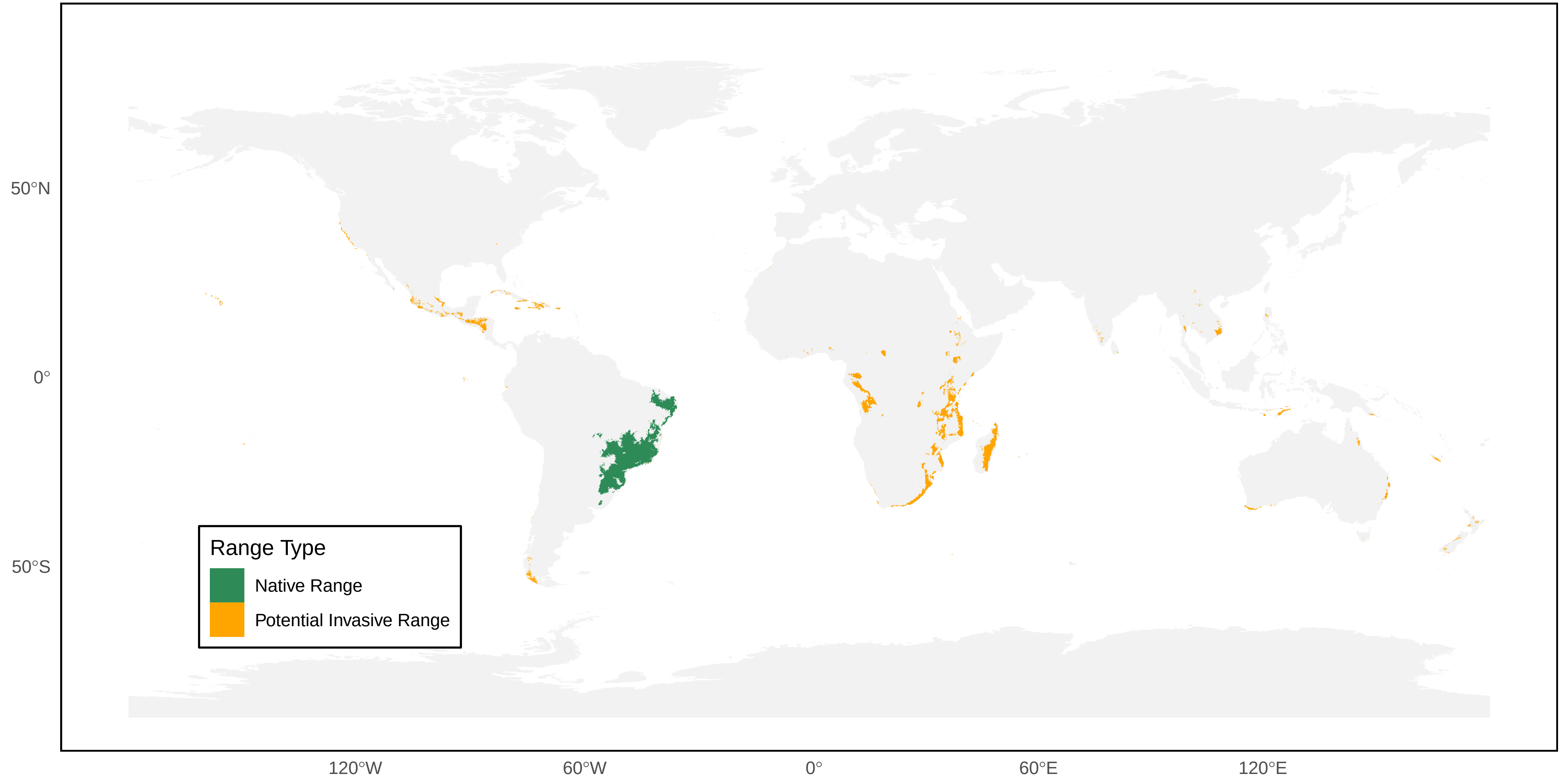

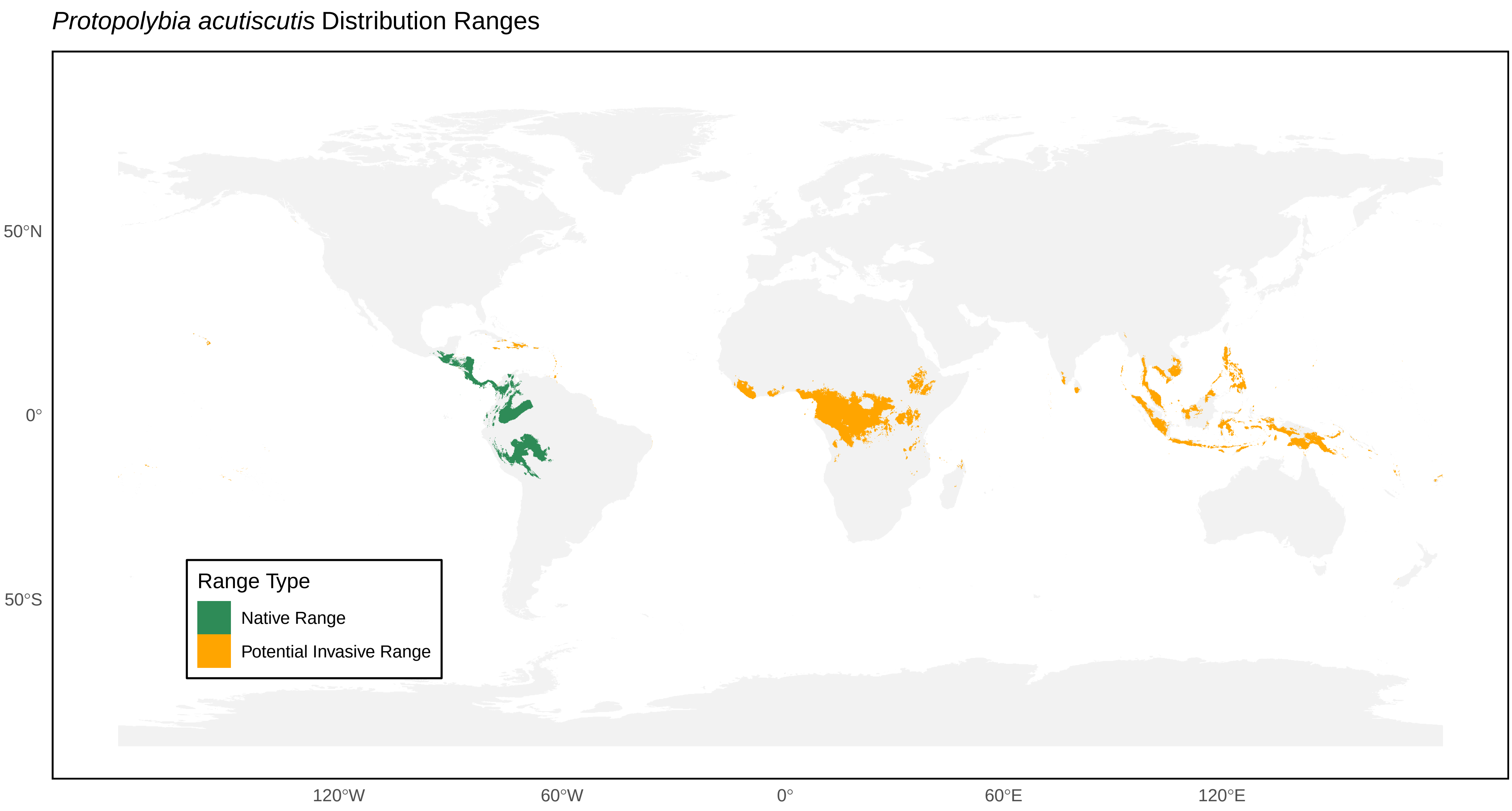
*Protopolybia acutiscutis* Distribution Ranges
50°N
0°
50°S
120°W
60°W
0°
60°E
120°E
Range Type
Native Range
Potential Invasive Range

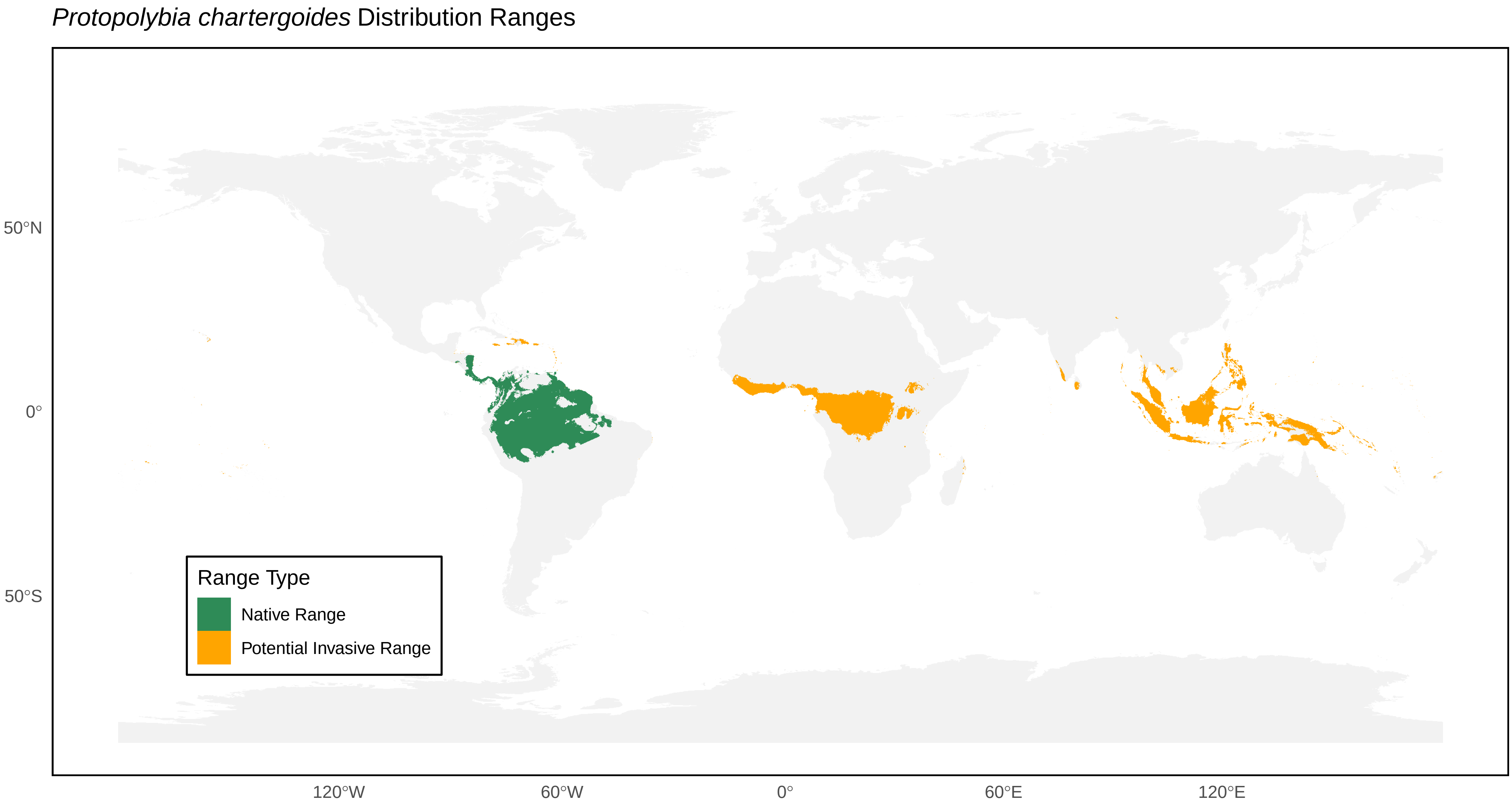

Protopolybia chartergoides Distribution Ranges
50°N
0°
50°S
120°W
60°W
0°
60°E
120°E
Range Type
Native Range
Potential Invasive Range

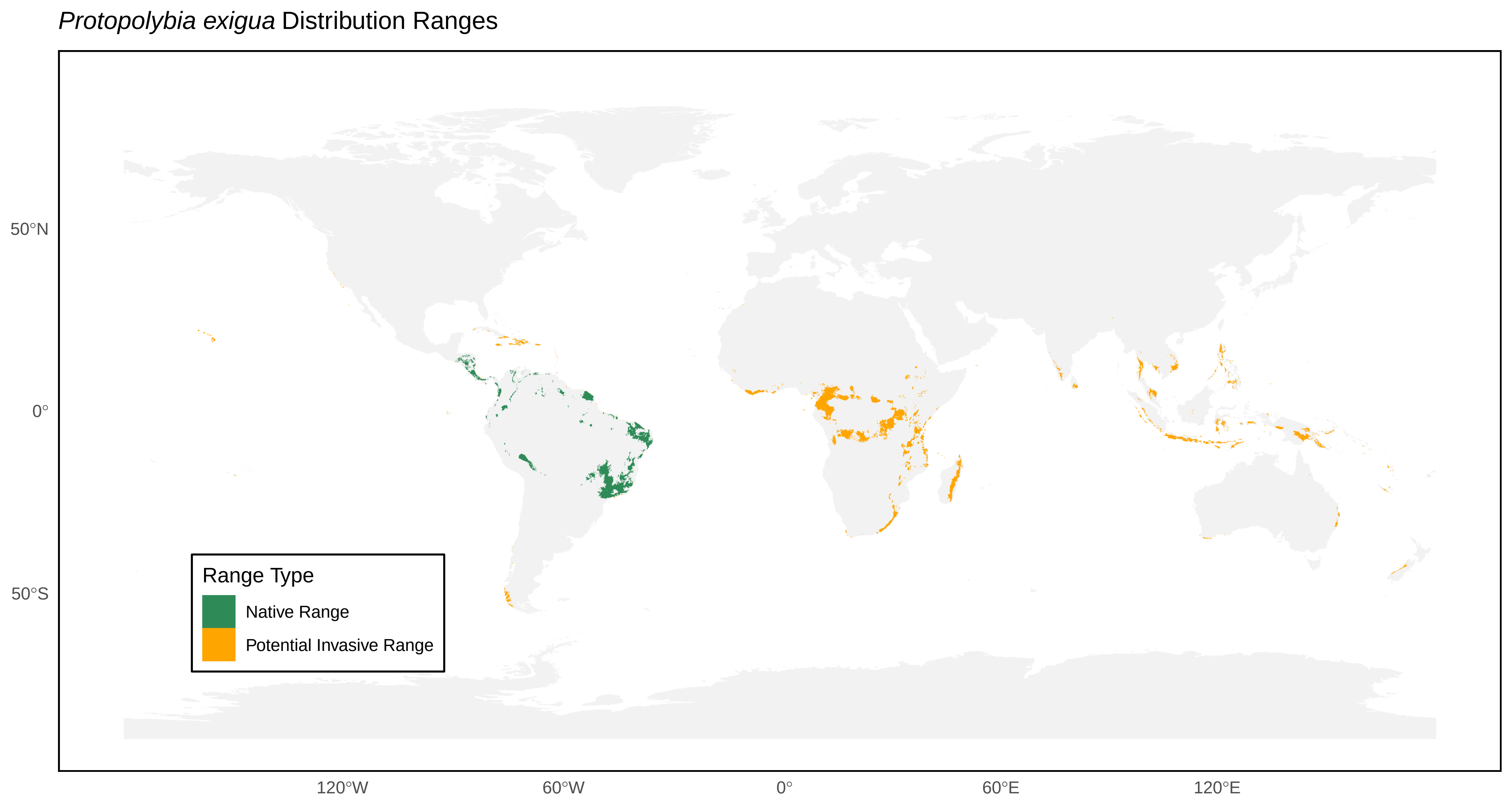

*Protopolybia exigua* Distribution Ranges
50°N
0°
50°S
120°W
60°W
0°
60°E
120°E
Range Type
Native Range
Potential Invasive Range

*Protopolybia sedula* Distribution Ranges

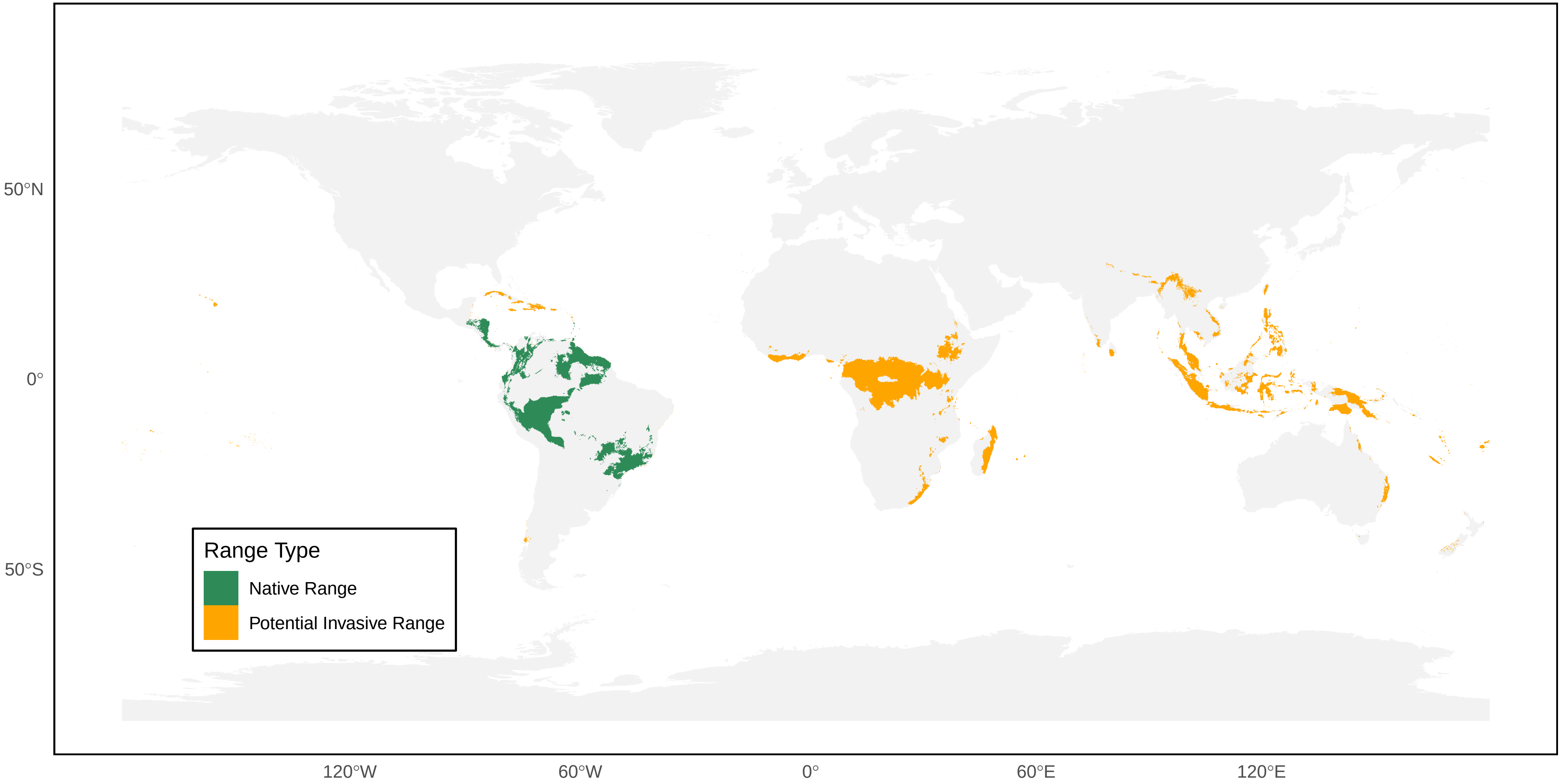

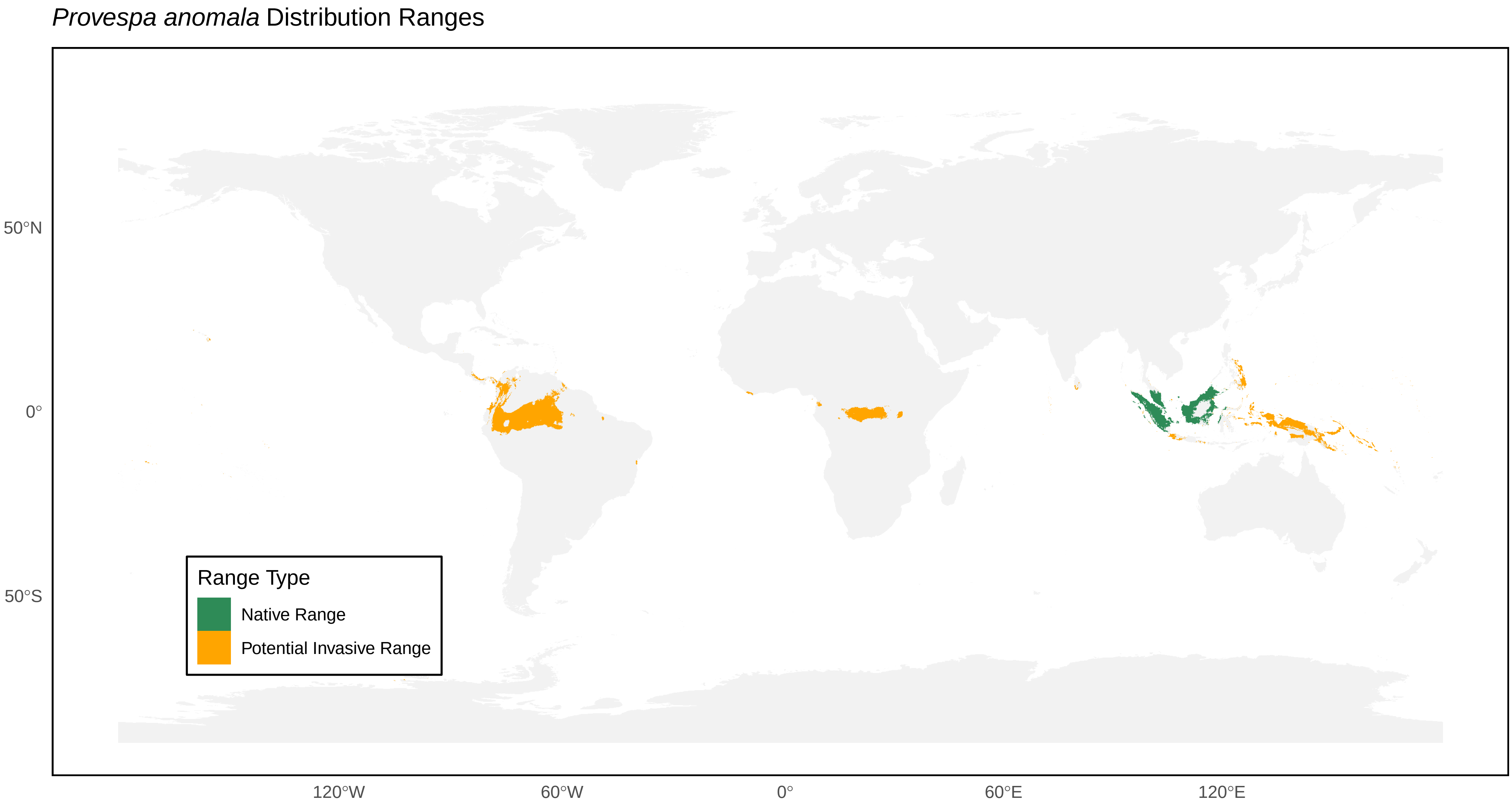

*Provespa anomala* Distribution Ranges
50°N
0°
50°S
120°W
60°W
0°
60°E
120°E
Range Type
Native Range
Potential Invasive Range

*Provespa barthelemyi* Distribution Ranges

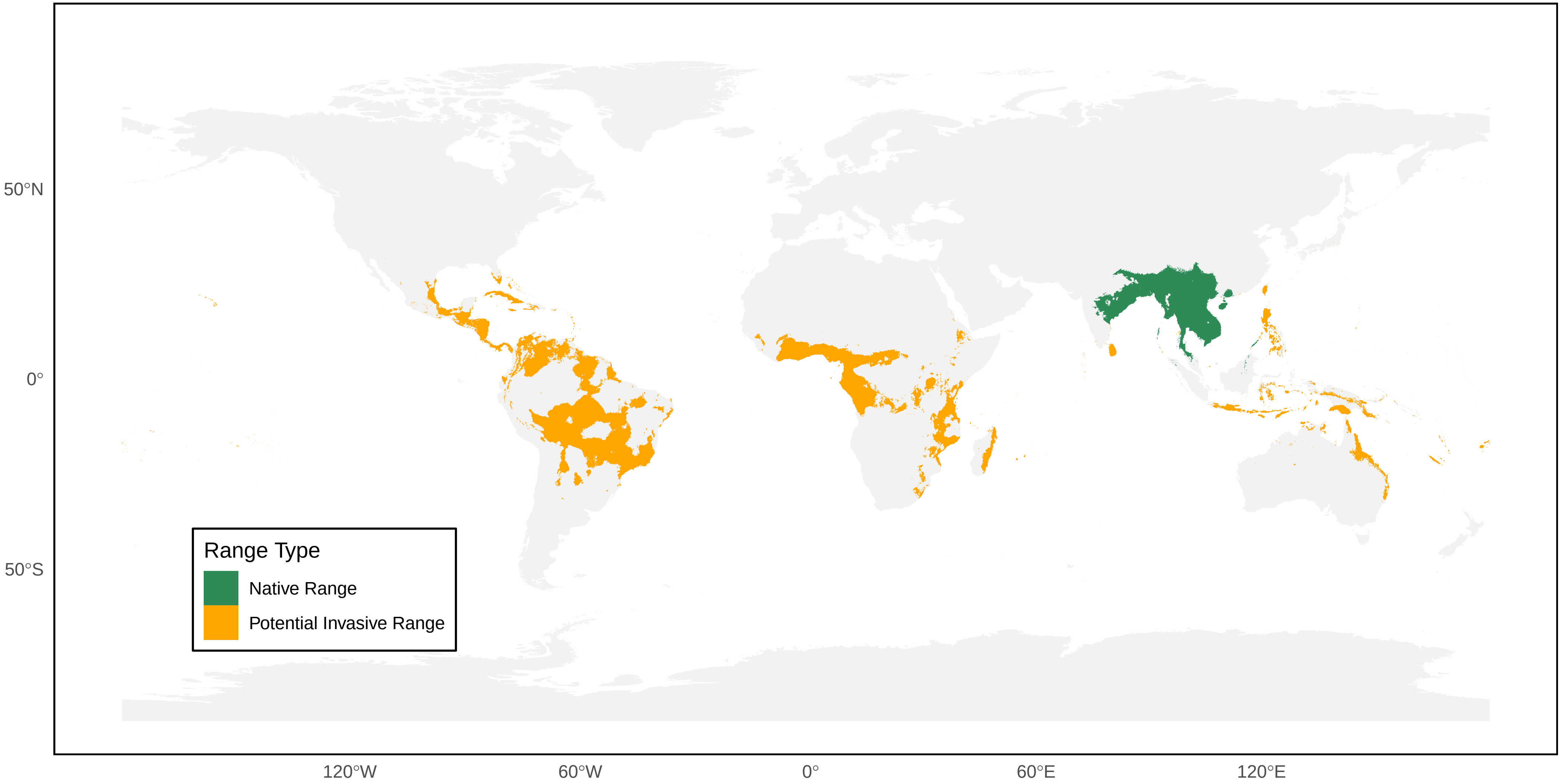

*Provespa nocturna* Distribution Ranges

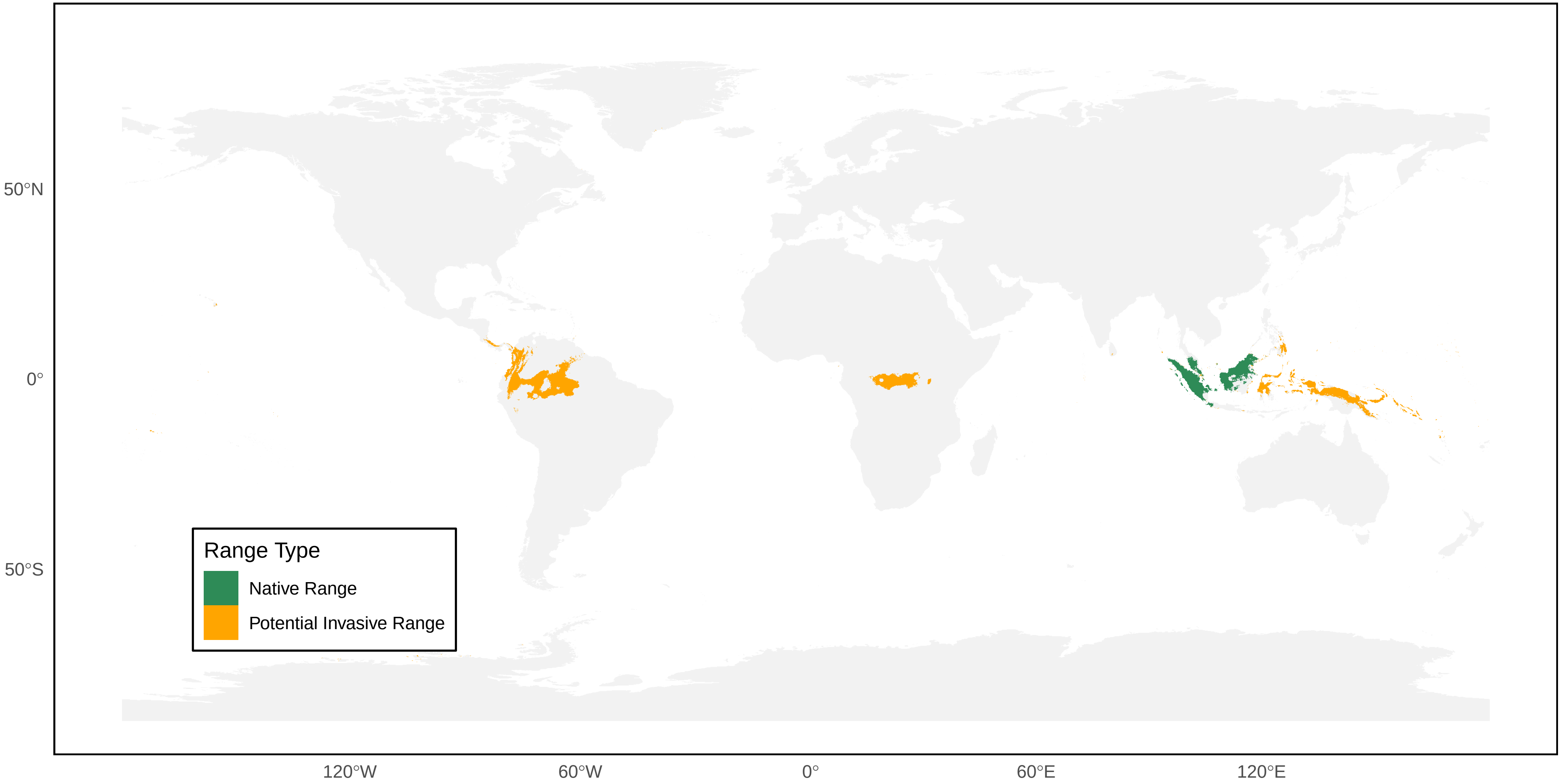

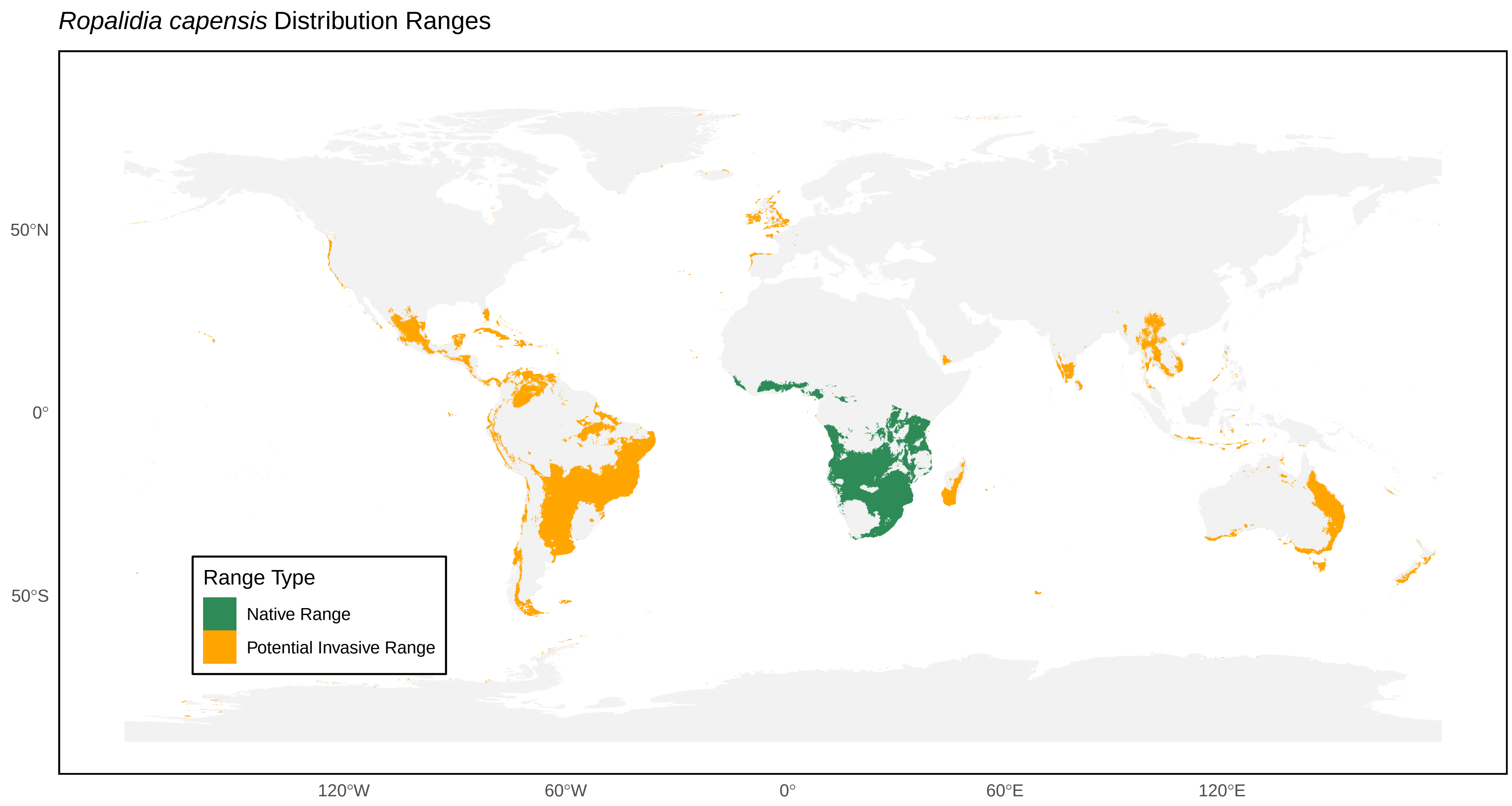

*Ropalidia capensis* Distribution Ranges
50°N
0°
50°S
120°W
60°W
0°
60°E
120°E
Range Type
Native Range
Potential Invasive Range

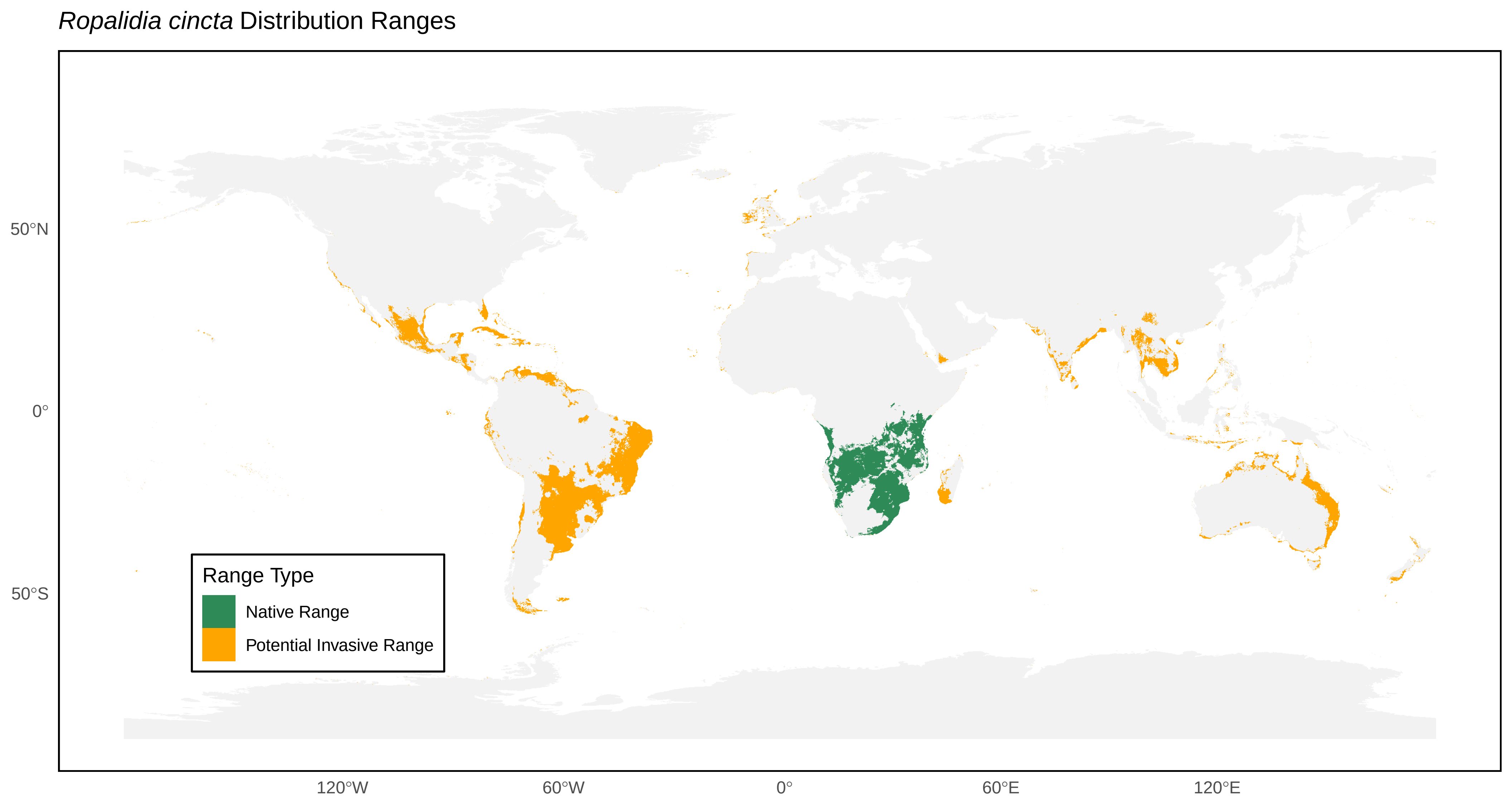
*Ropalidia cincta* Distribution Ranges
Range Type
Native Range
Potential Invasive Range
50°N
0°
50°S
120°W
60°W
0°
60°E
120°E

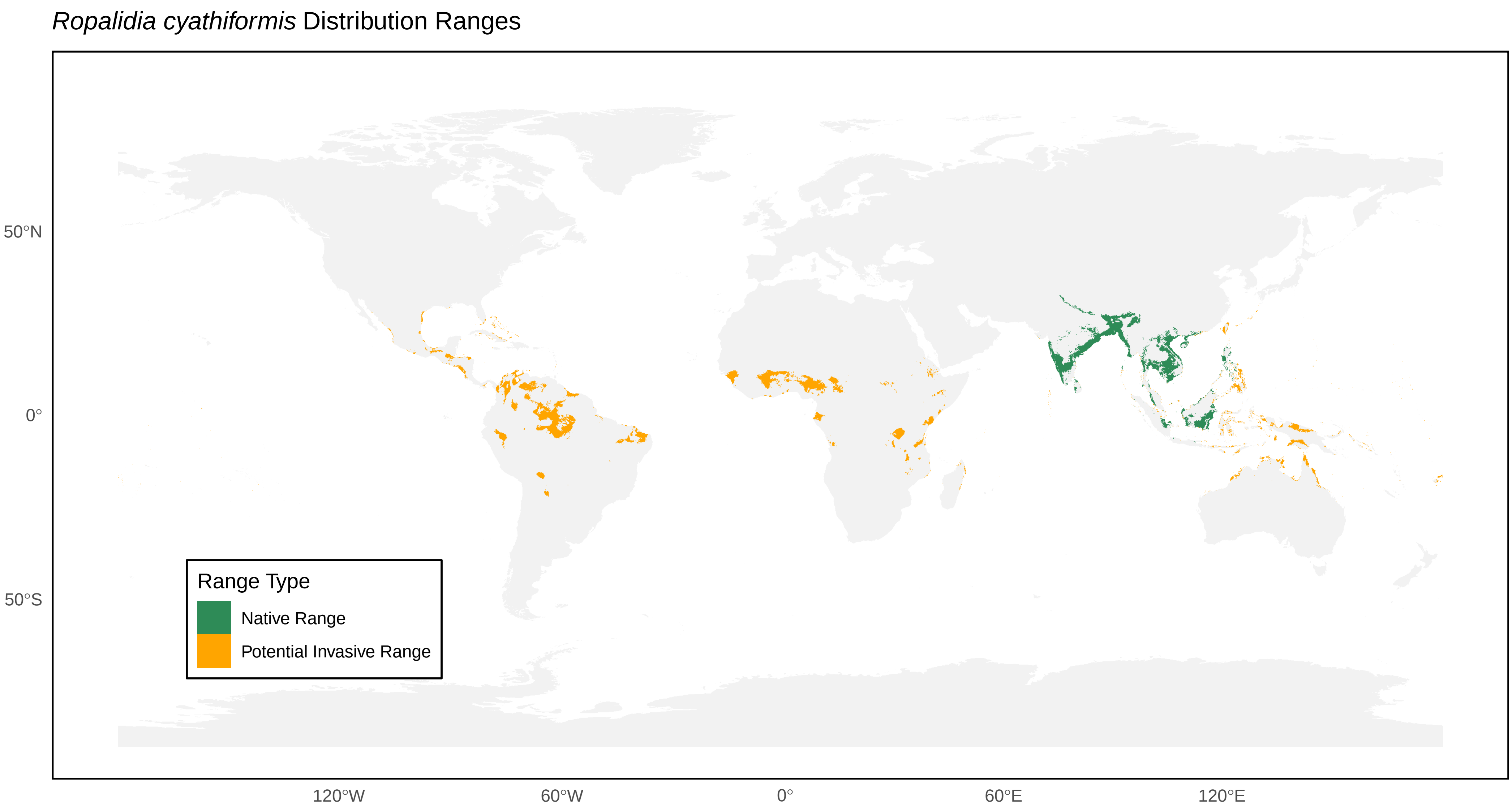
*Ropalidia cyathiformis* Distribution Ranges
50°N
0°
50°S
120°W
60°W
0°
60°E
120°E
Range Type
Native Range
Potential Invasive Range

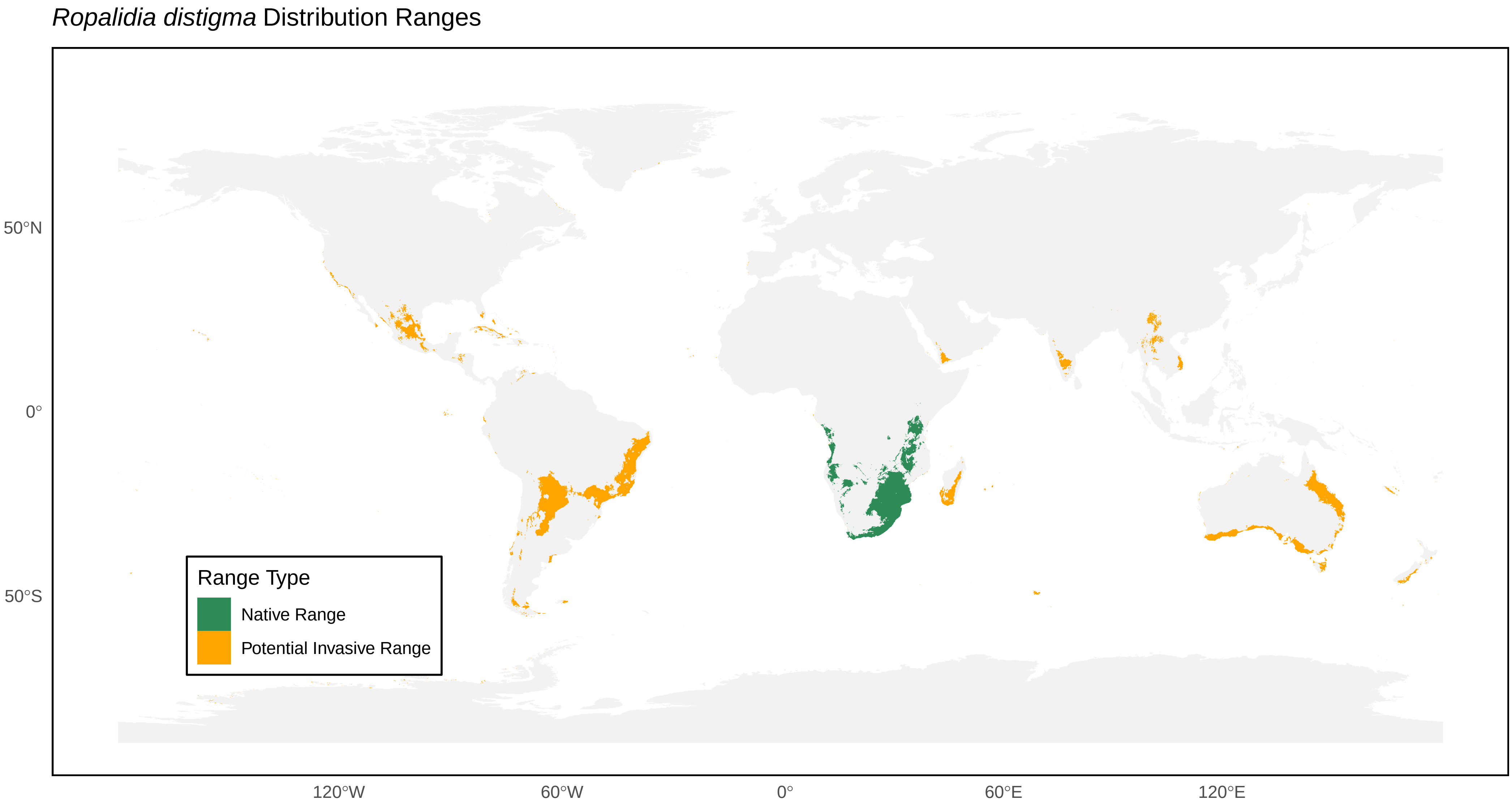
*Ropalidia distigma* Distribution Ranges
50°N
0°
50°S
120°W
60°W
0°
60°E
120°E
Range Type
Native Range
Potential Invasive Range

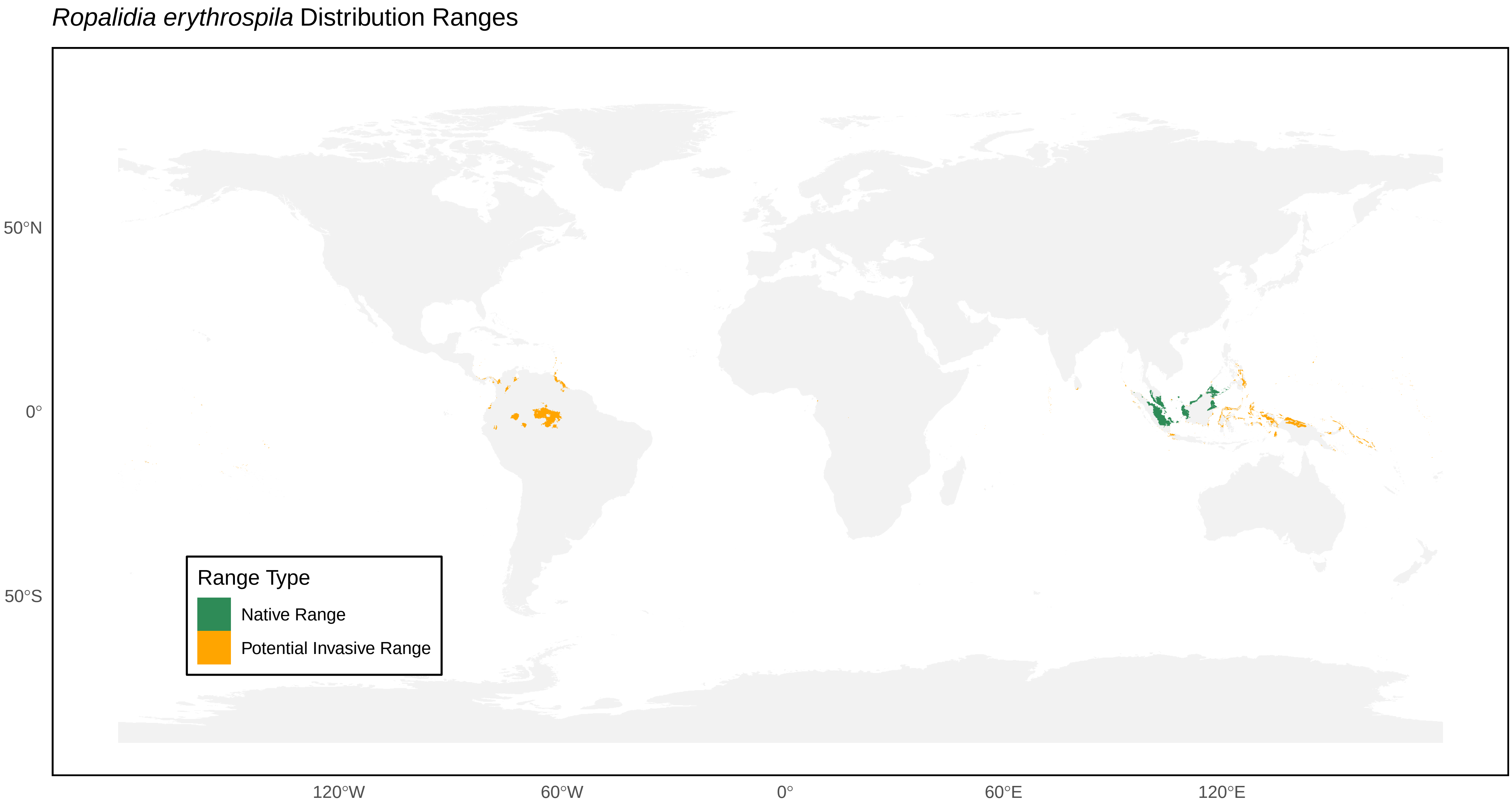
*Ropalidia erythrospila* Distribution Ranges
Range Type
Native Range
Potential Invasive Range
50°N
0°
50°S
120°W
60°W
0°
60°E
120°E

*Ropalidia fasciata* Distribution Ranges

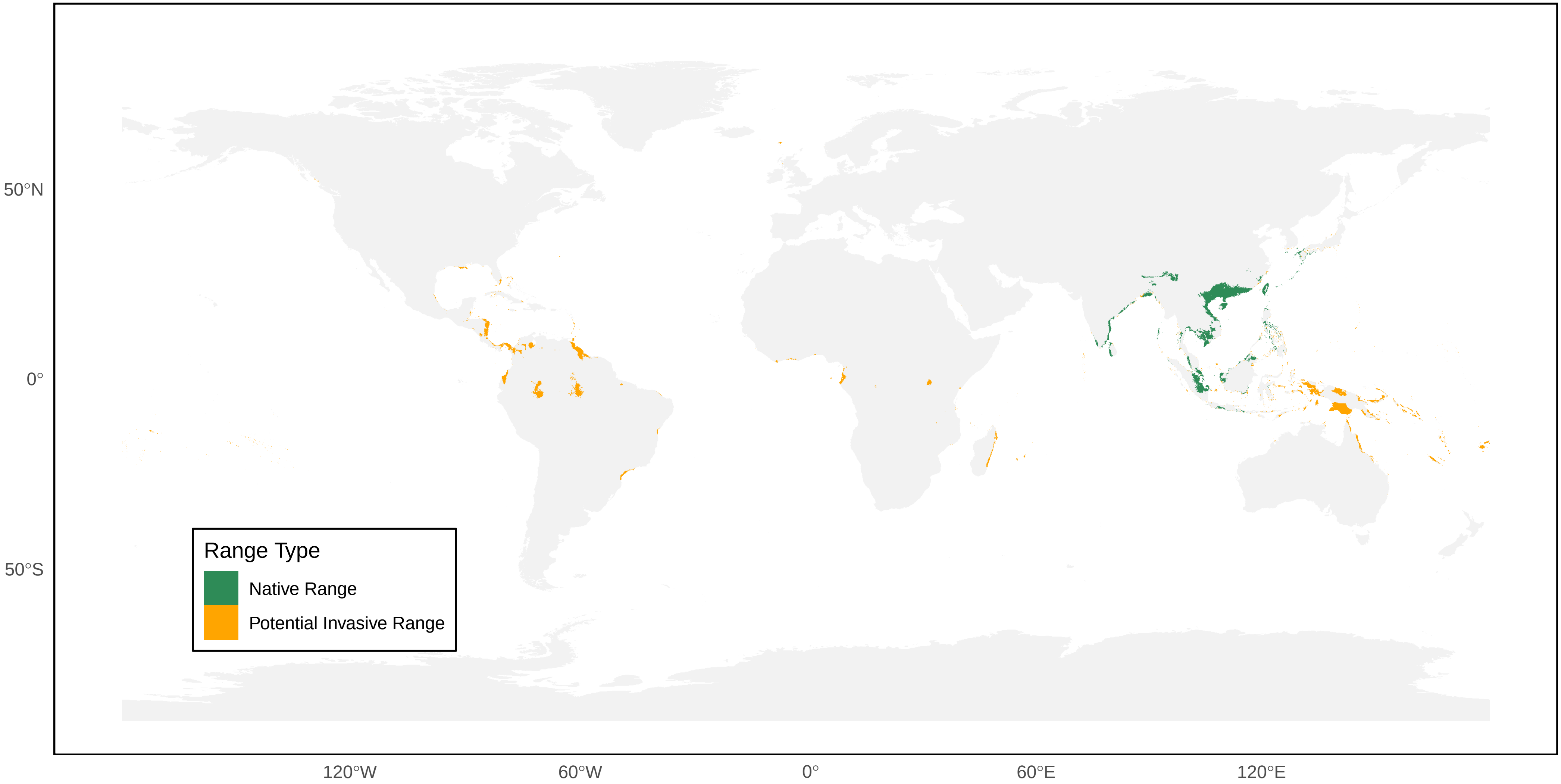

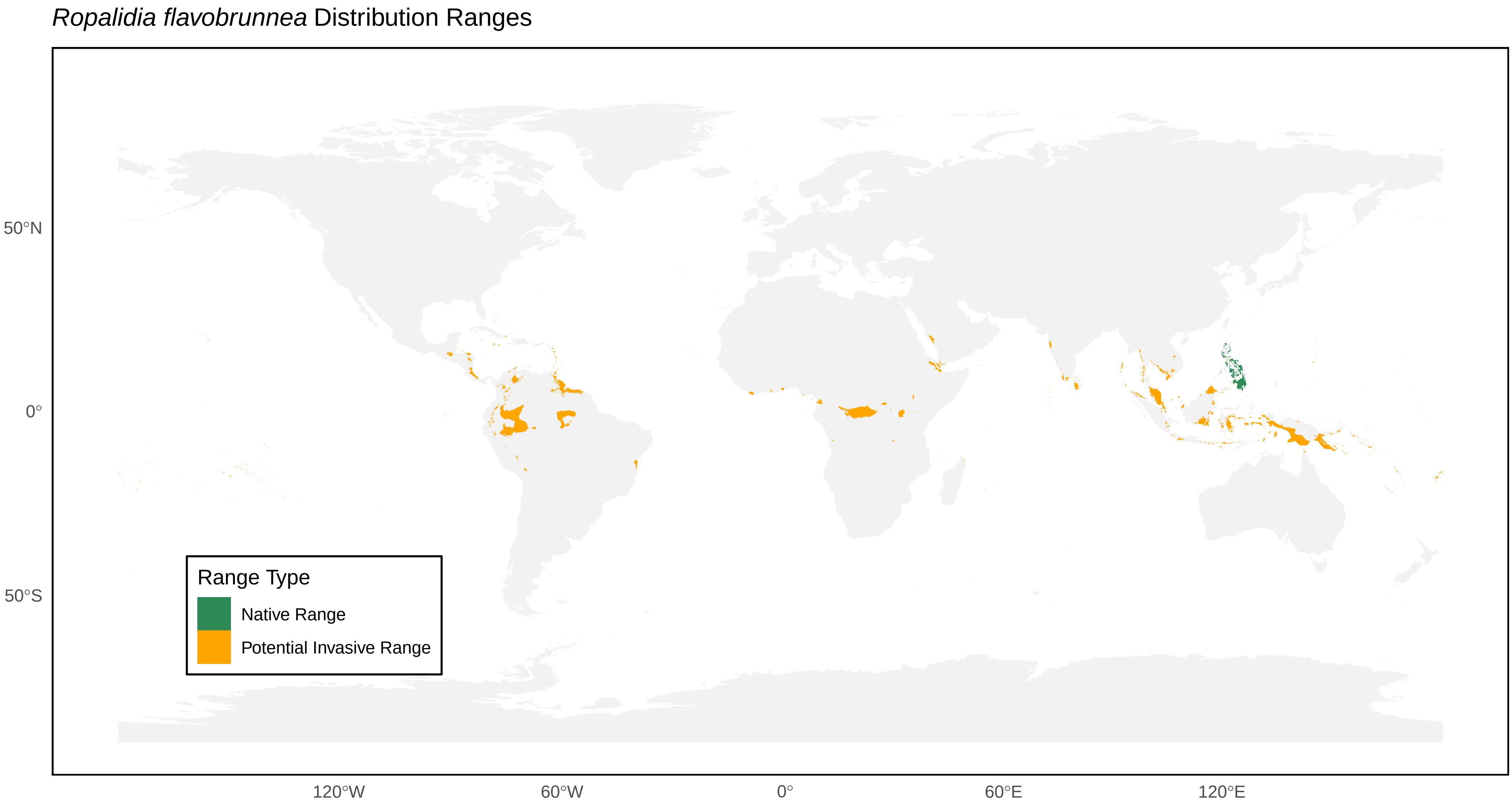

*Ropalidia flavobrunnea* Distribution Ranges
50°N
0°
50°S
120°W
60°W
0°
60°E
120°E
Range Type
Native Range
Potential Invasive Range

*Ropalidia flavopicta* Distribution Ranges

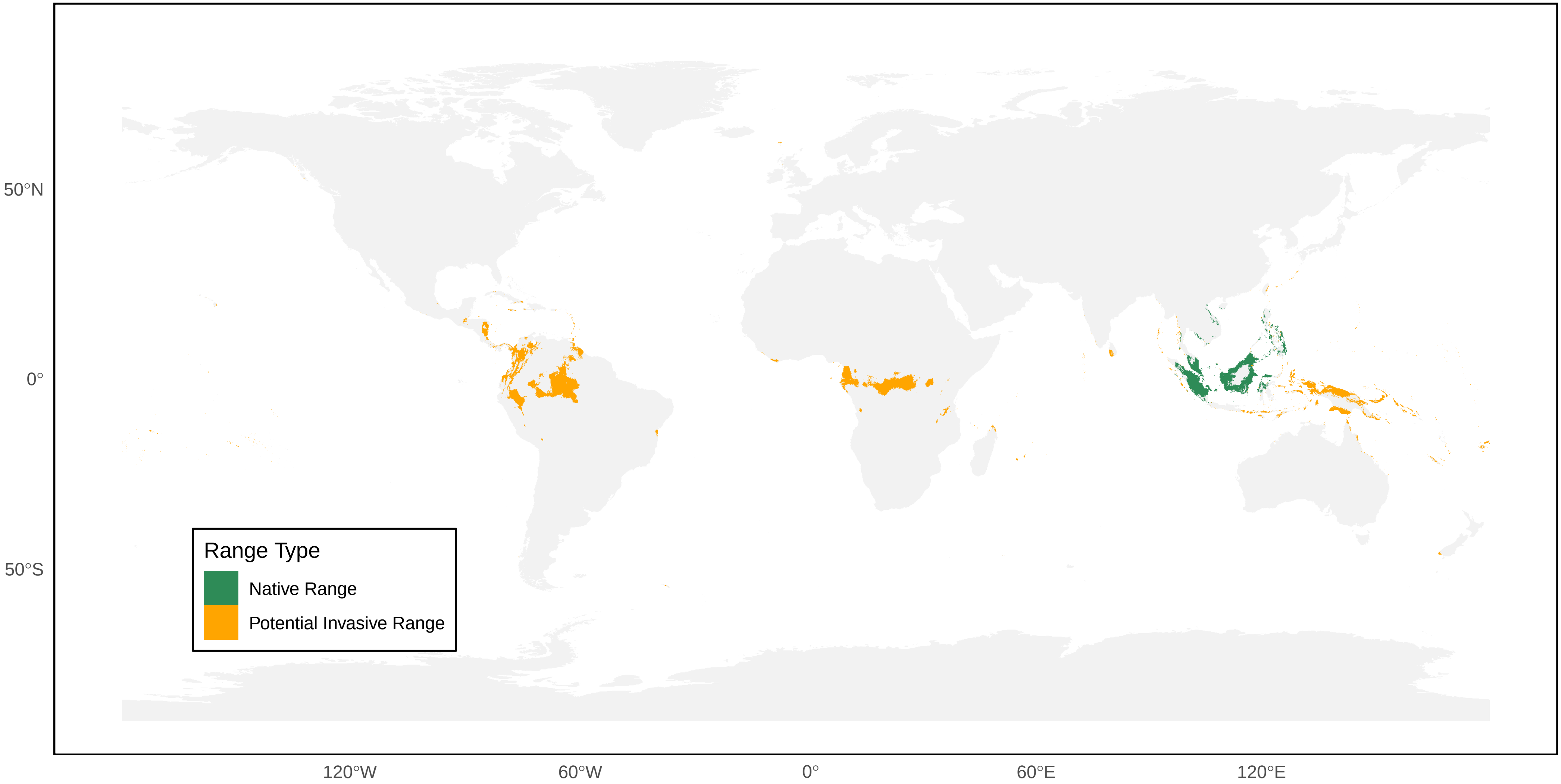

*Ropalidia galimatia* Distribution Ranges

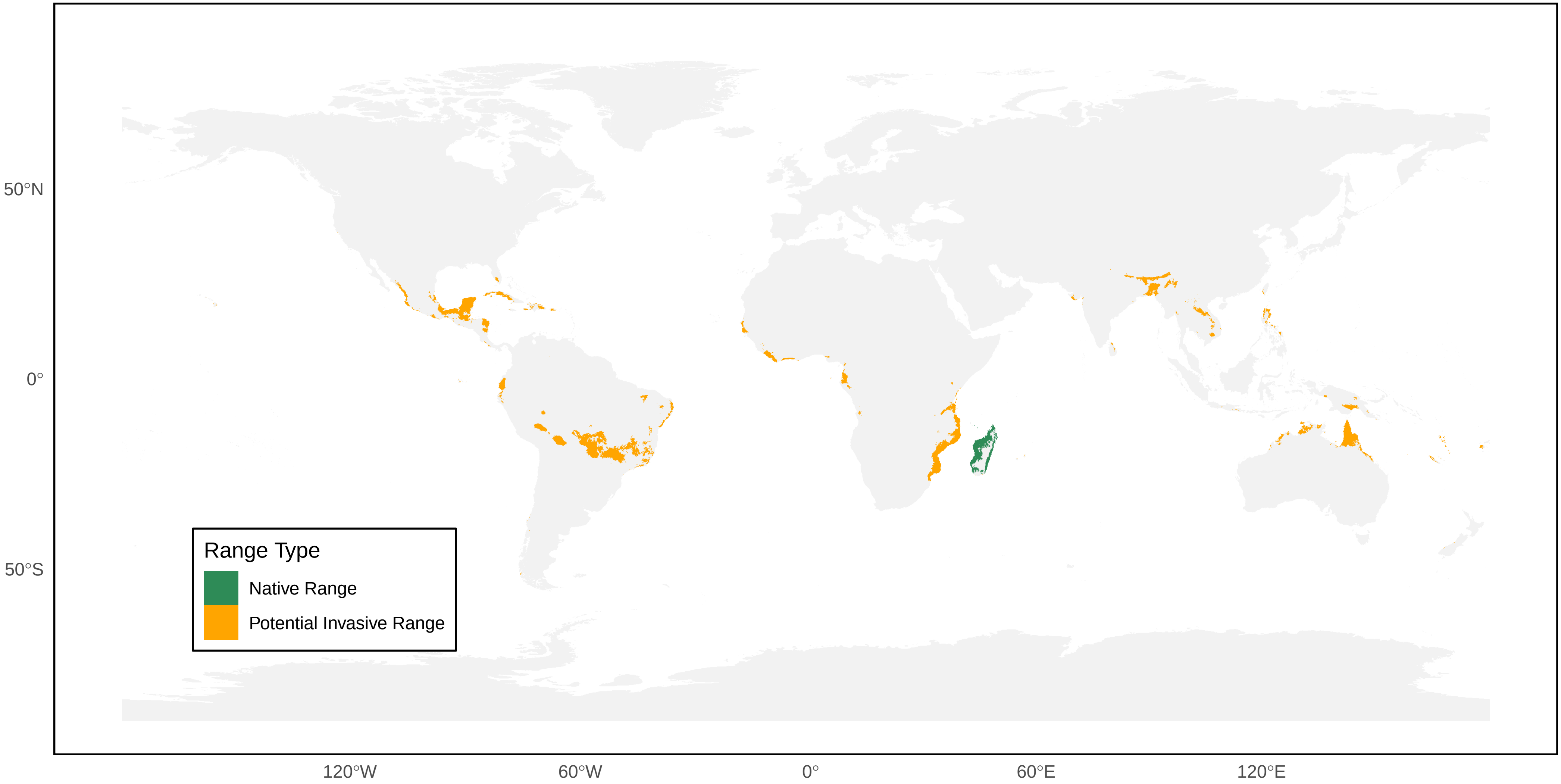

*Ropalidia gregaria* Distribution Ranges

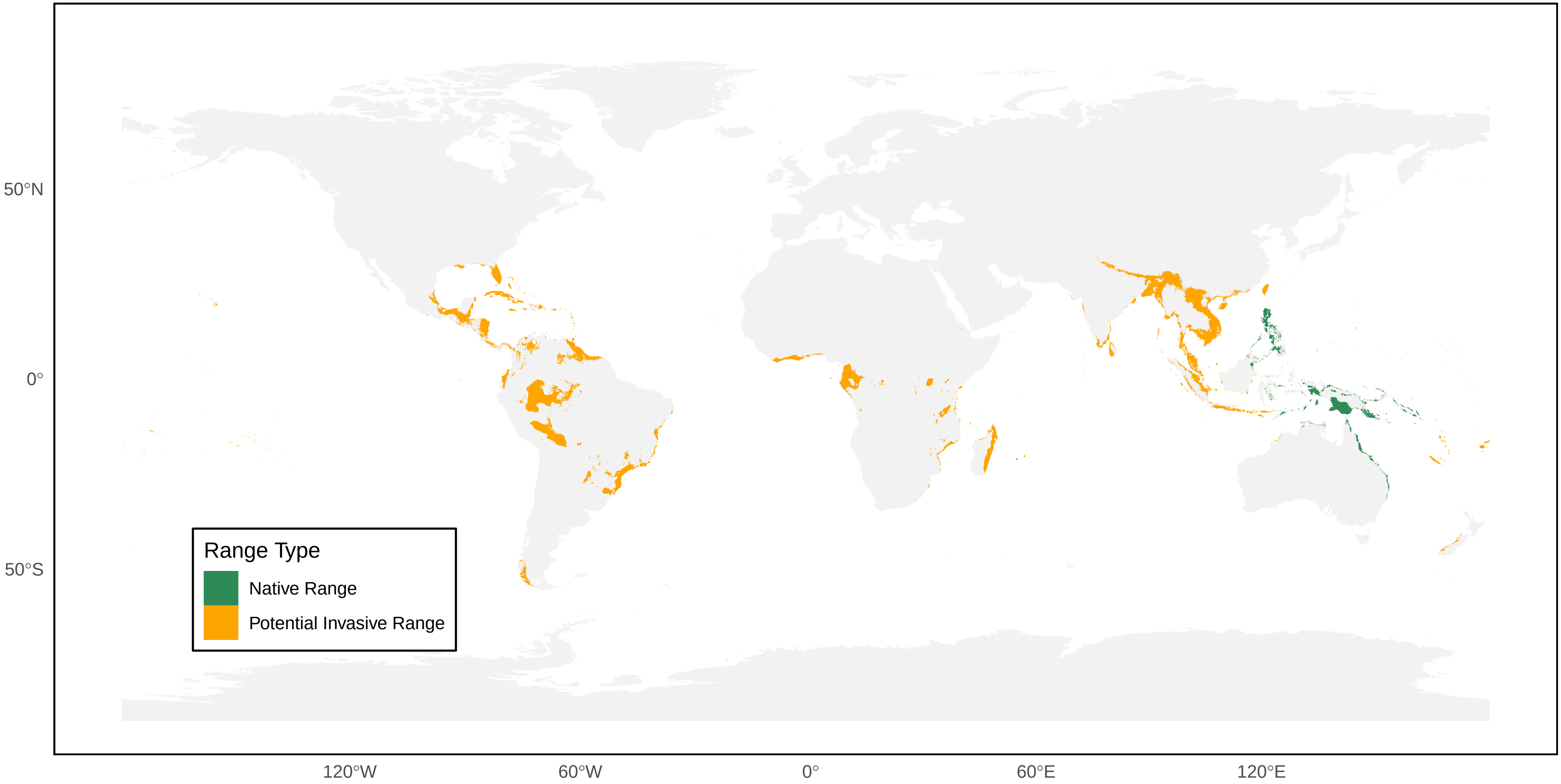

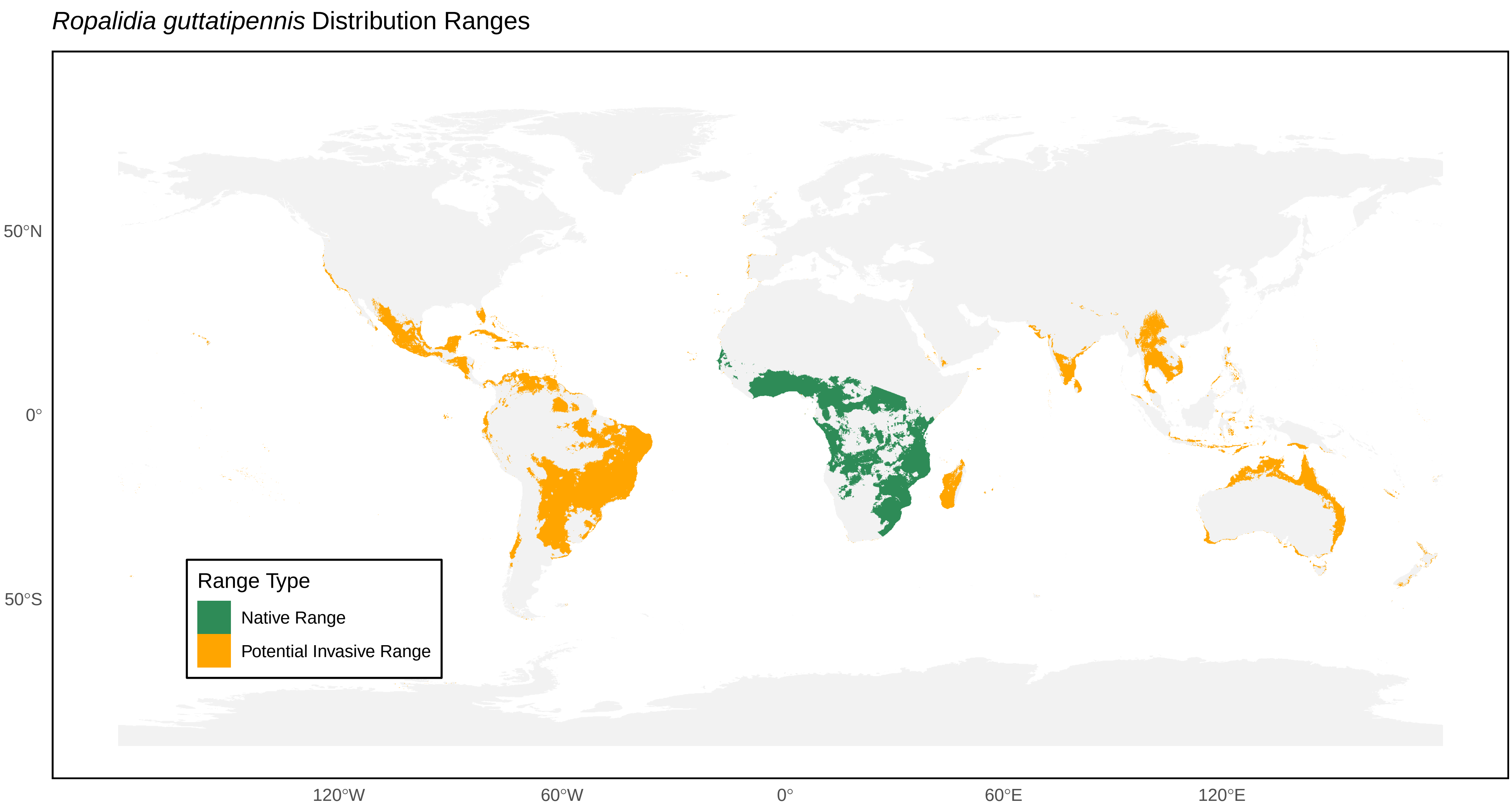

*Ropalidia guttatipennis* Distribution Ranges
50°N
0°
50°S
120°W
60°W
0°
60°E
120°E
Range Type
Native Range
Potential Invasive Range

*Ropalidia hongkongensis* Distribution Ranges

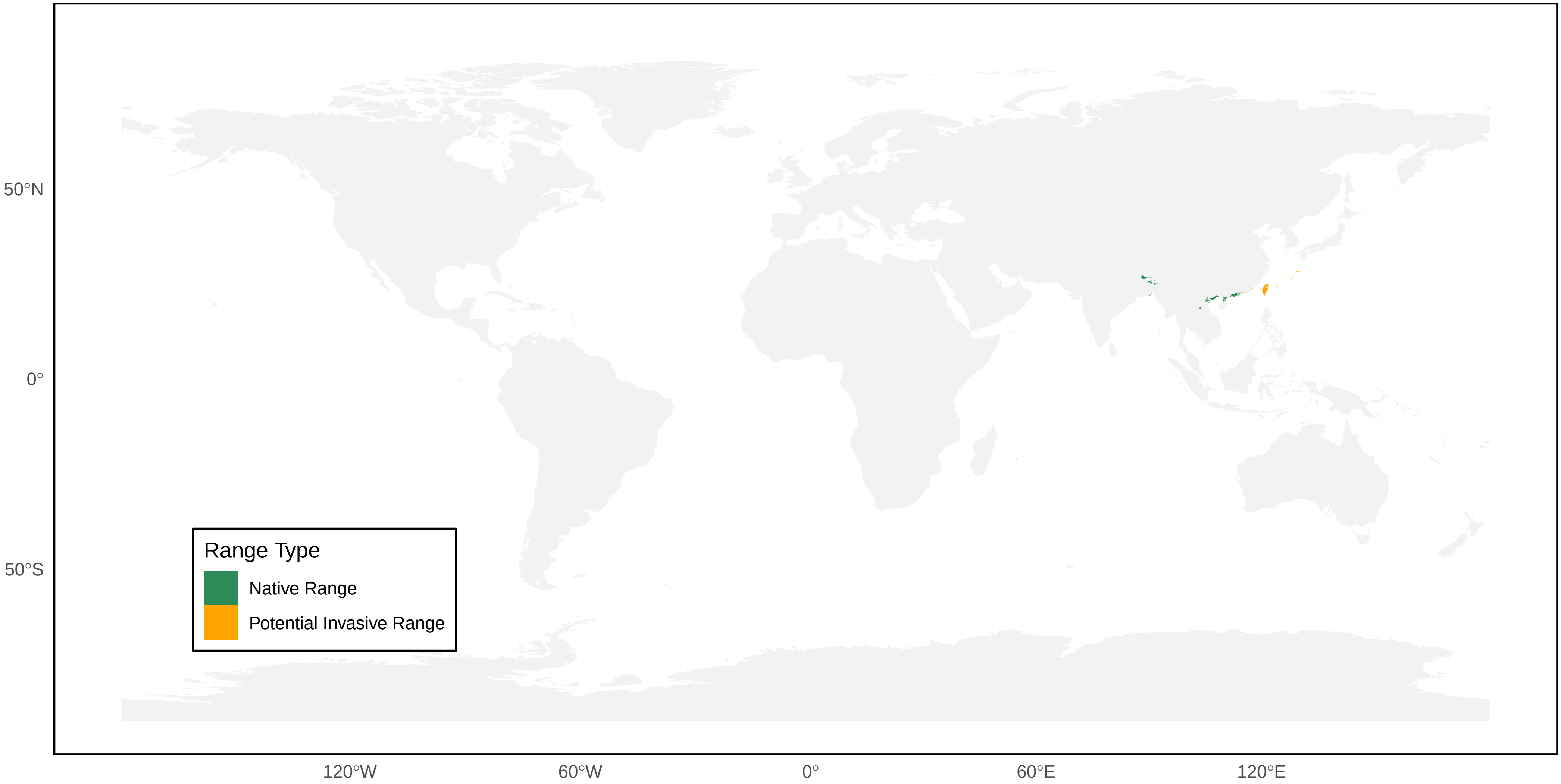

*Ropalidia horni* Distribution Ranges

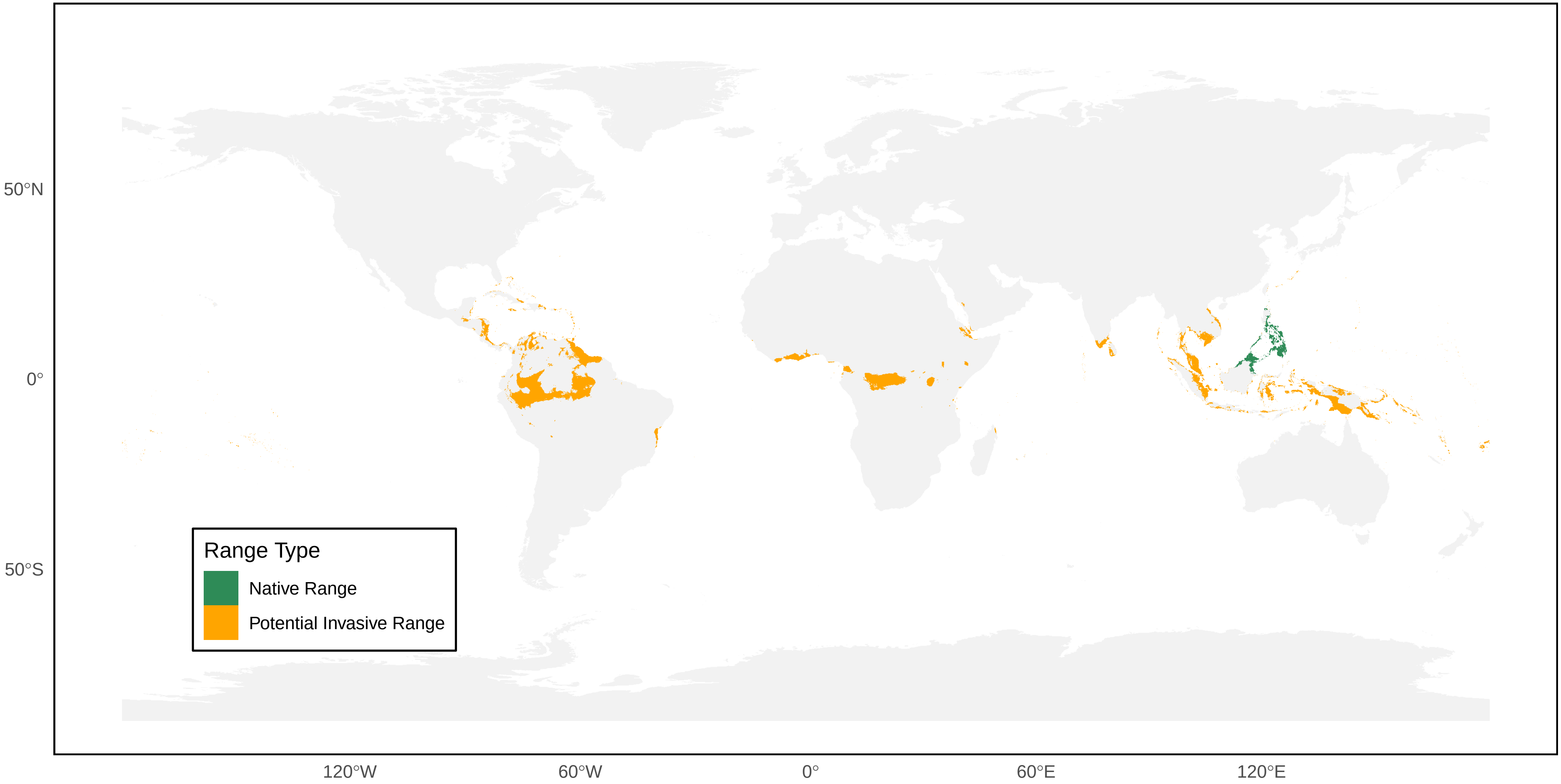

*Ropalidia impetuosa* Distribution Ranges

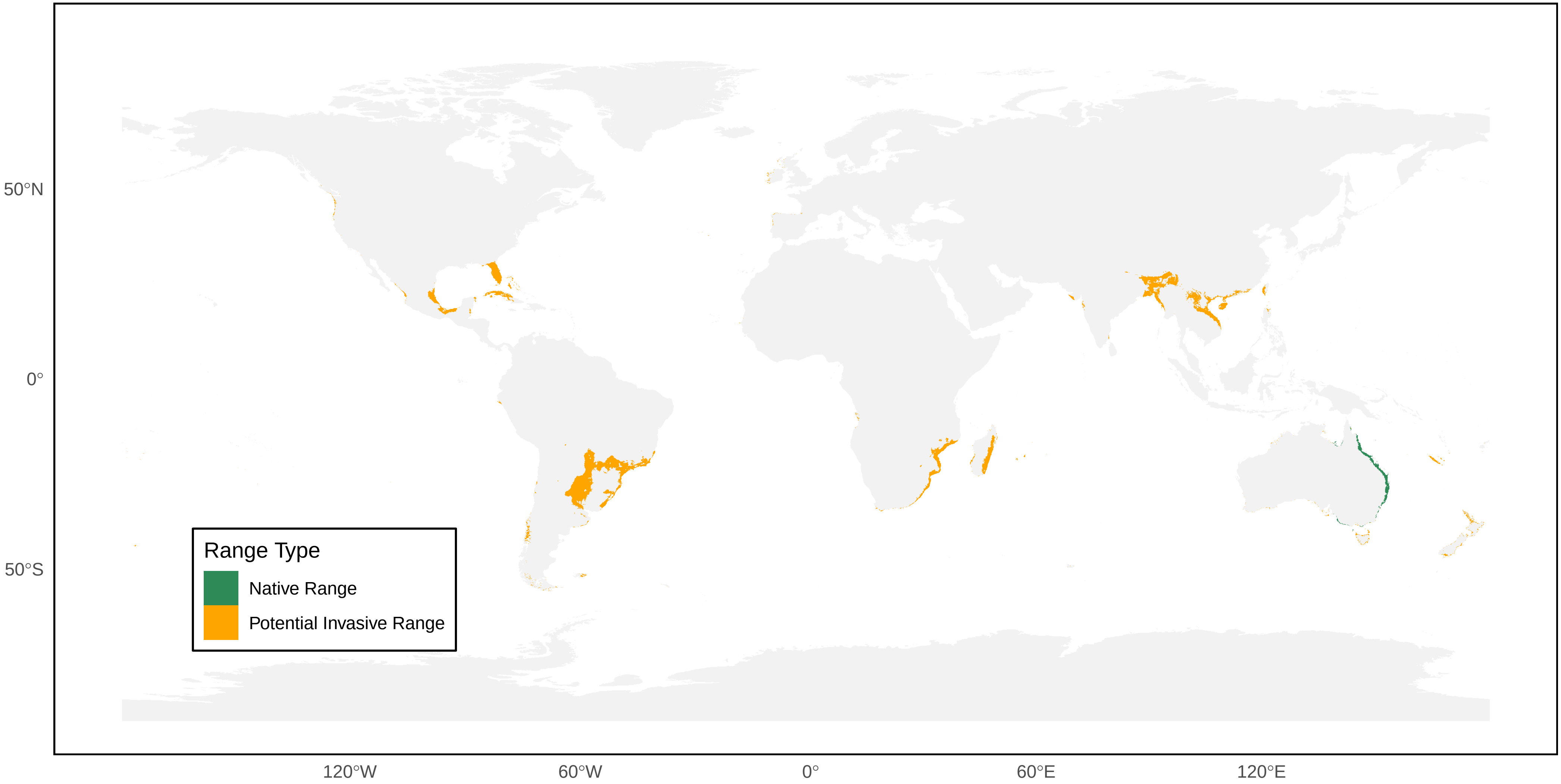

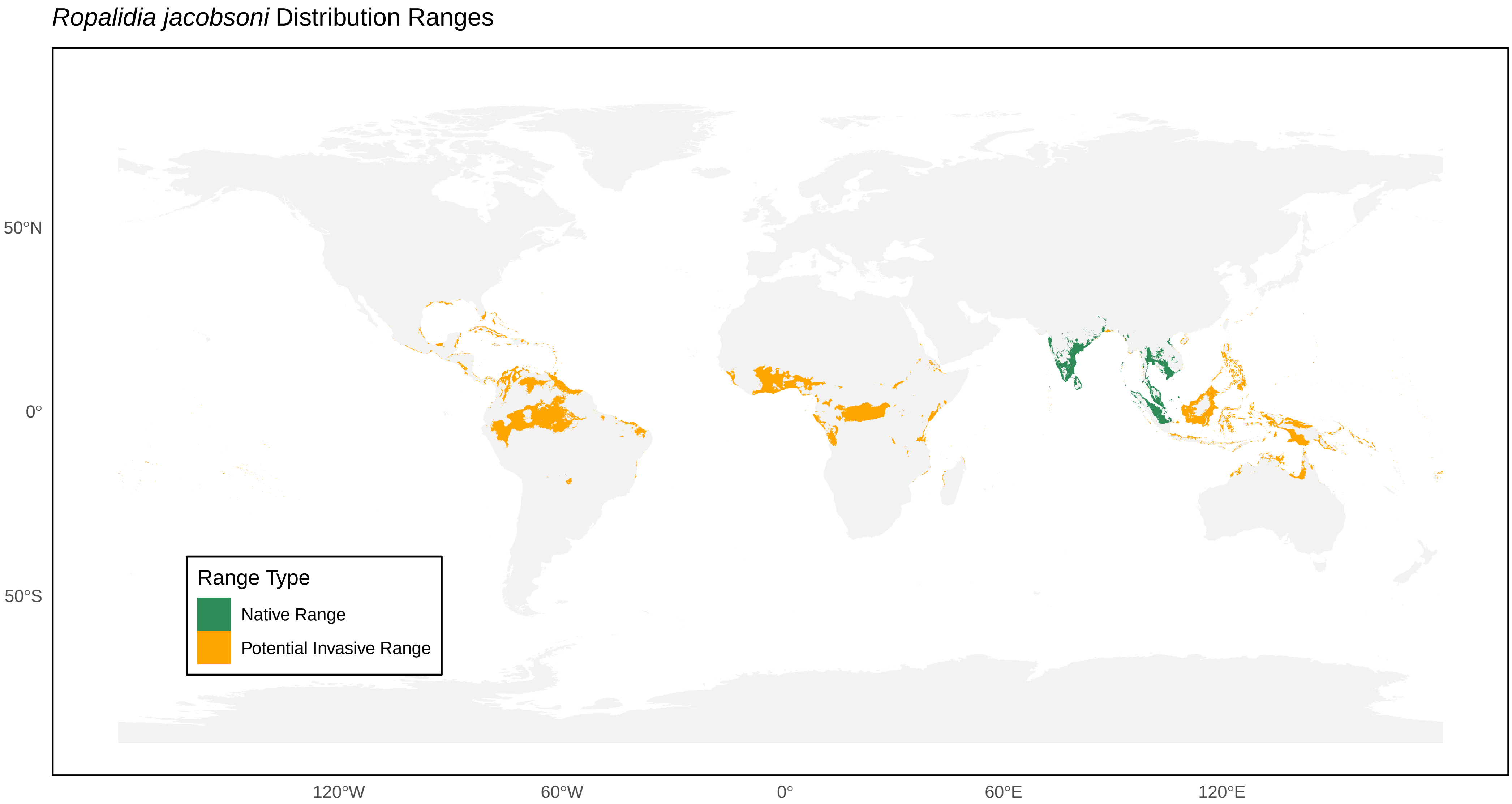
*Ropalidia jacobsoni* Distribution Ranges
Range Type
Native Range
Potential Invasive Range
50°N
0°
50°S
120°W
60°W
0°
60°E
120°E

### *Ropalidia magnanima* Distribution Ranges

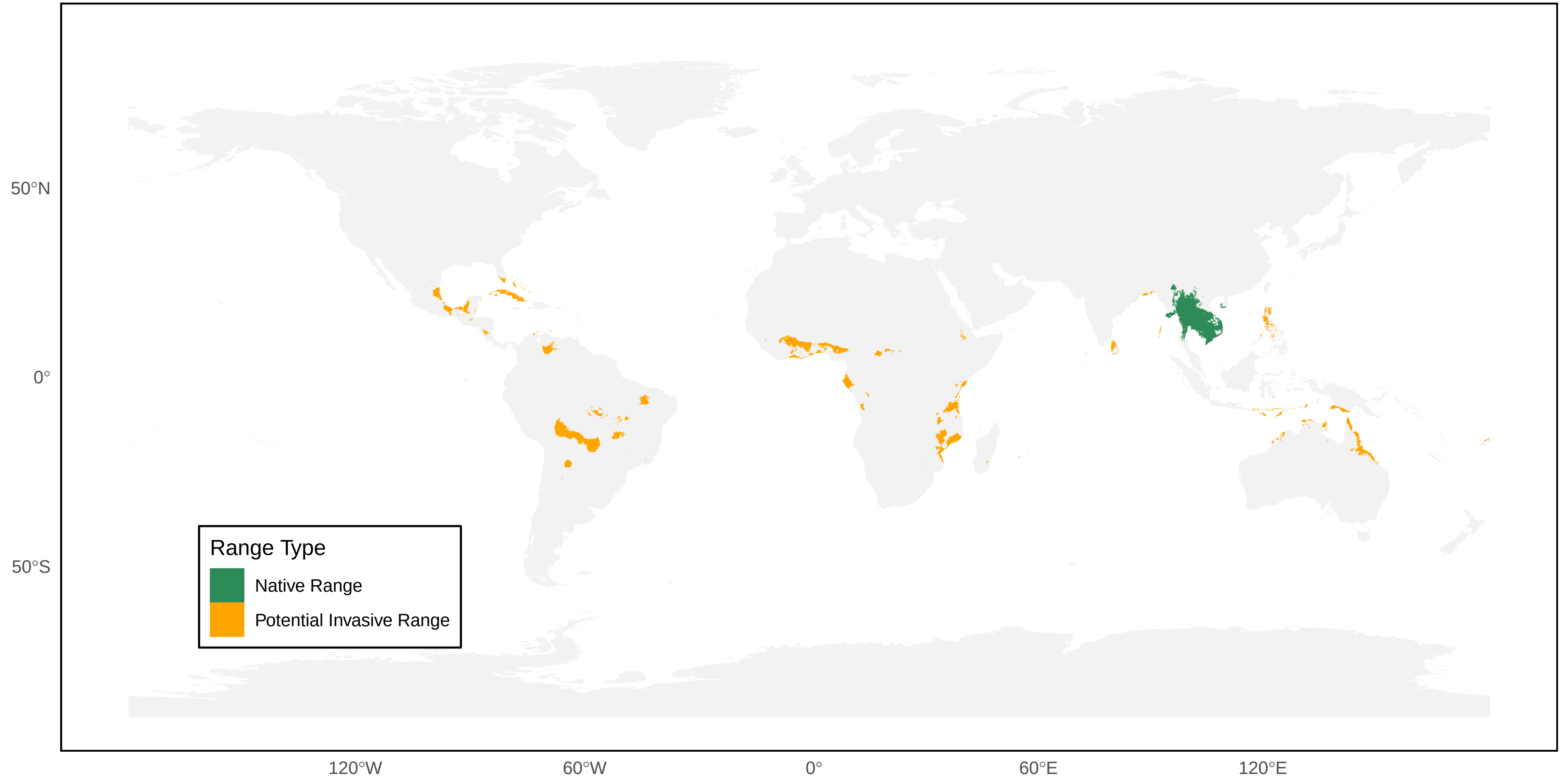

*Ropalidia marginata* Distribution Ranges

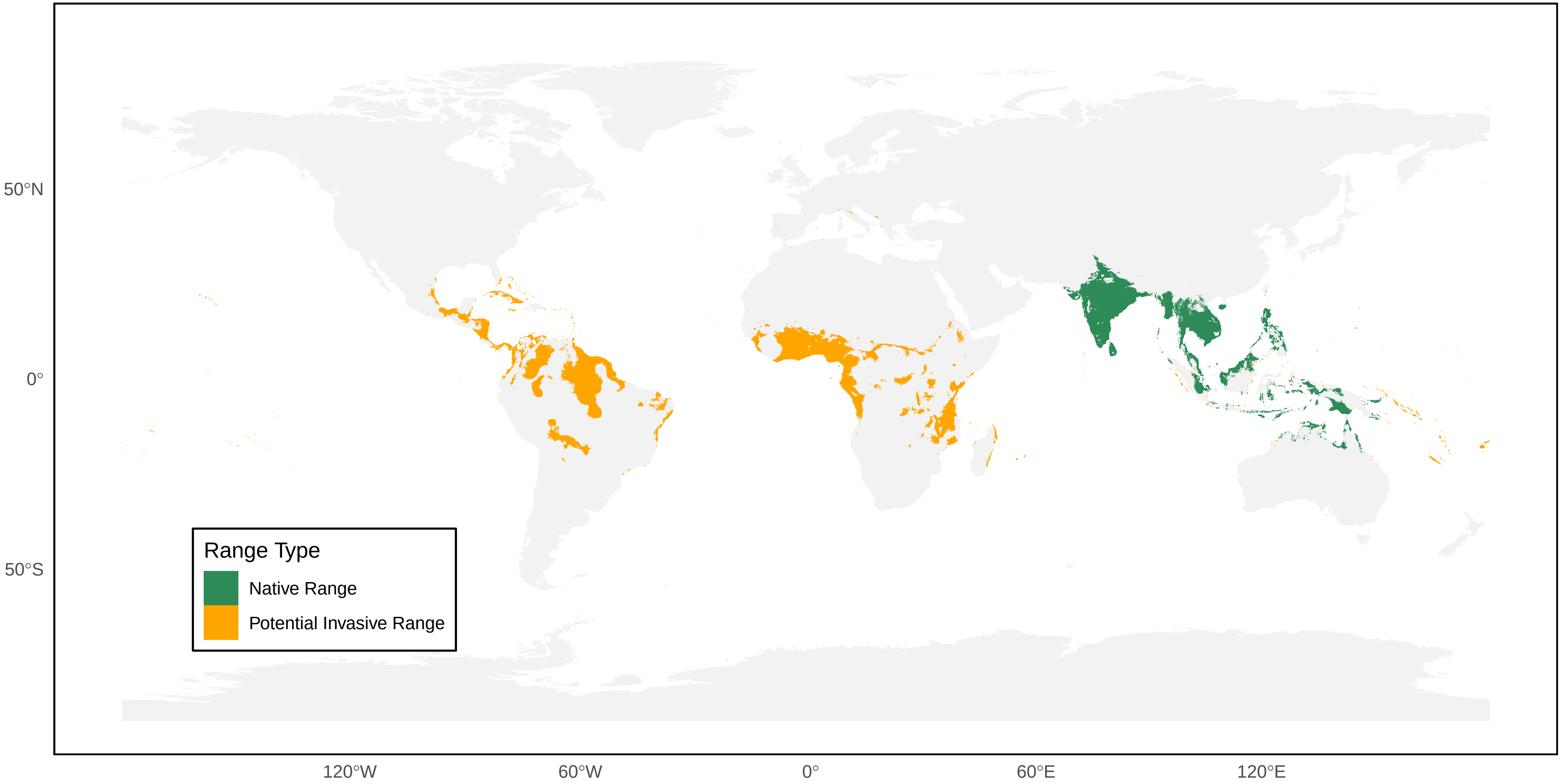

*Ropalidia merina* Distribution Ranges

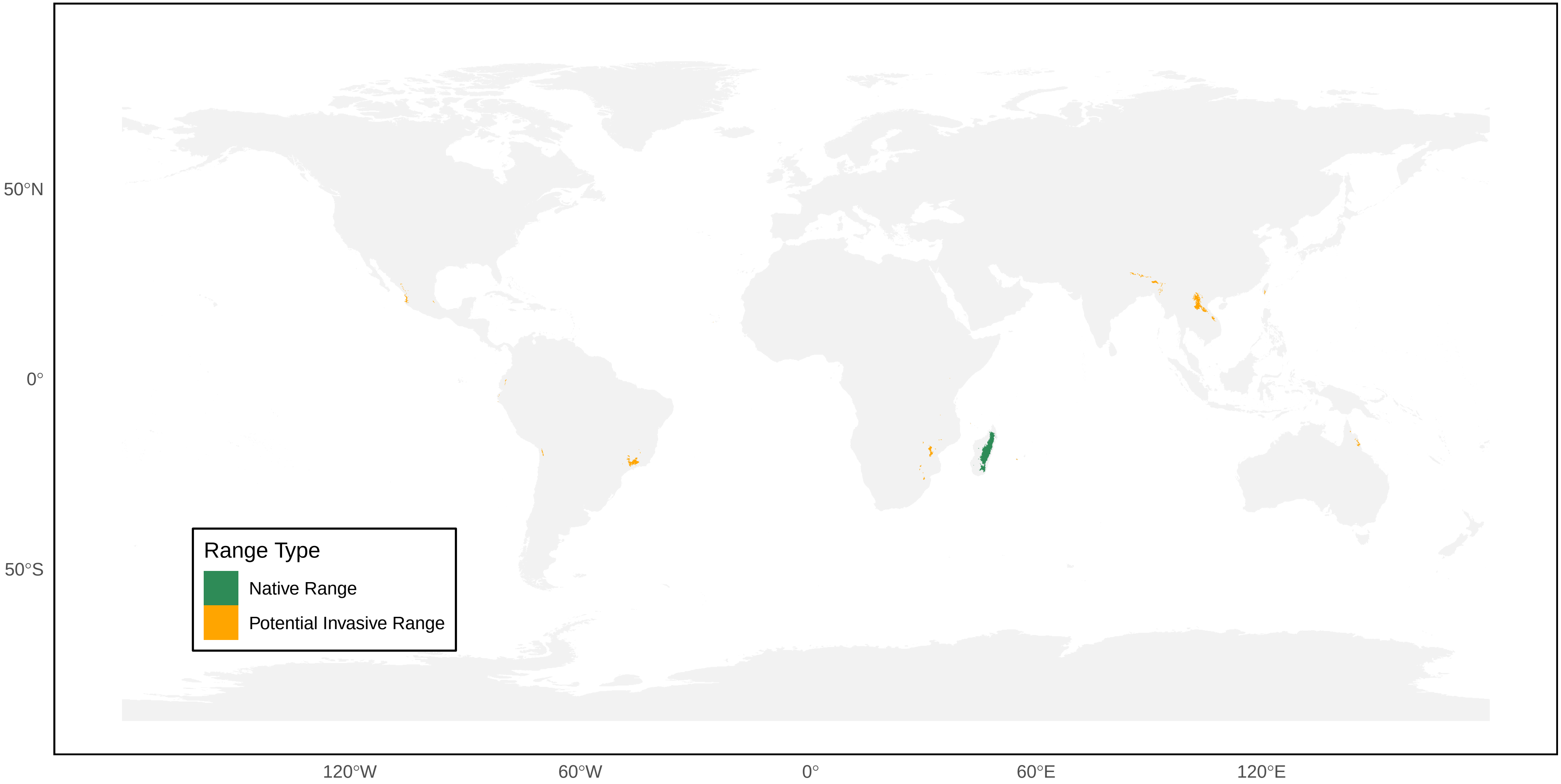

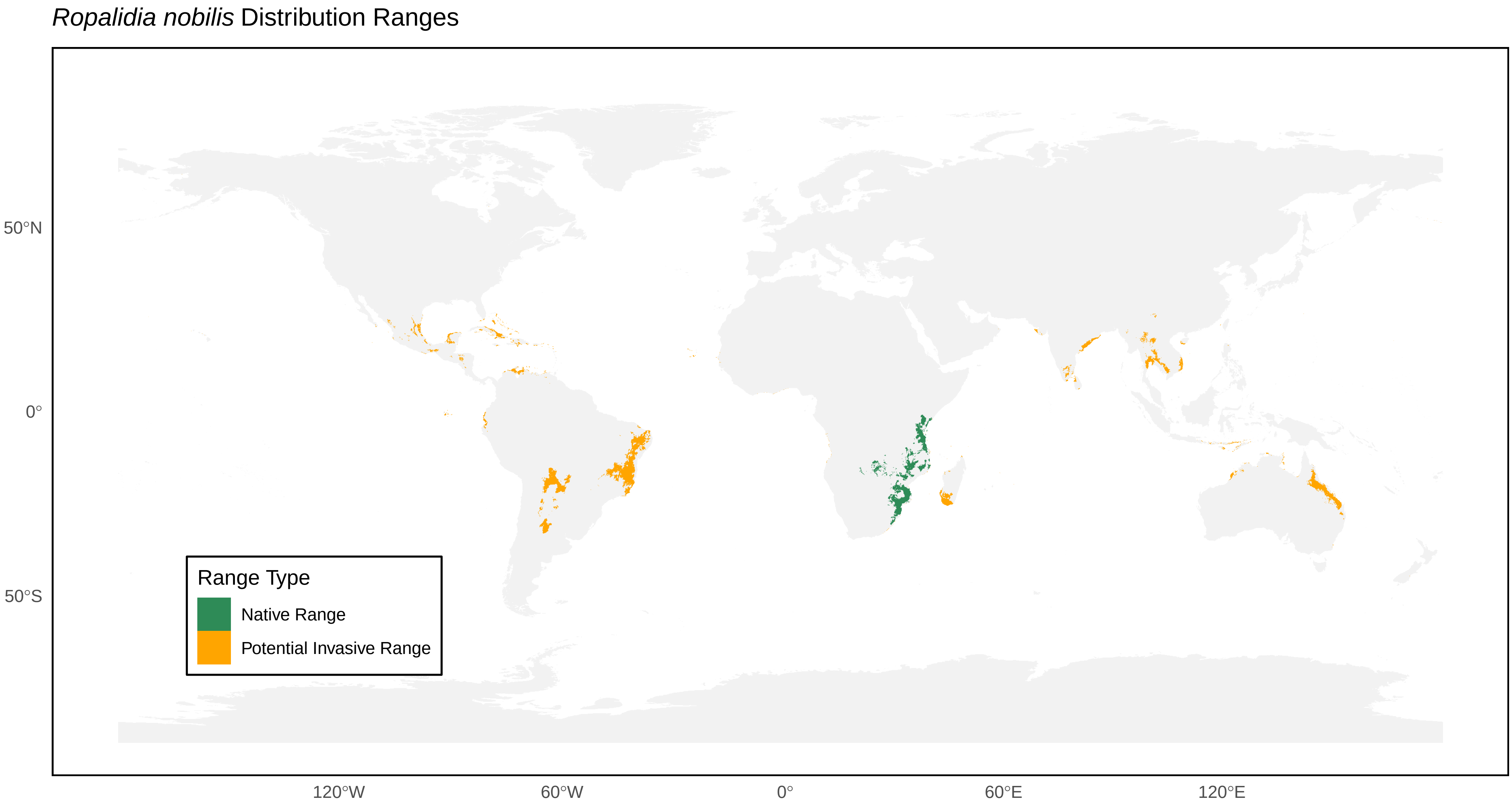

*Ropalidia nobilis* Distribution Ranges
50°N
0°
50°S
120°W
60°W
0°
60°E
120°E
Range Type
Native Range
Potential Invasive Range

*Ropalidia ornaticeps* Distribution Ranges

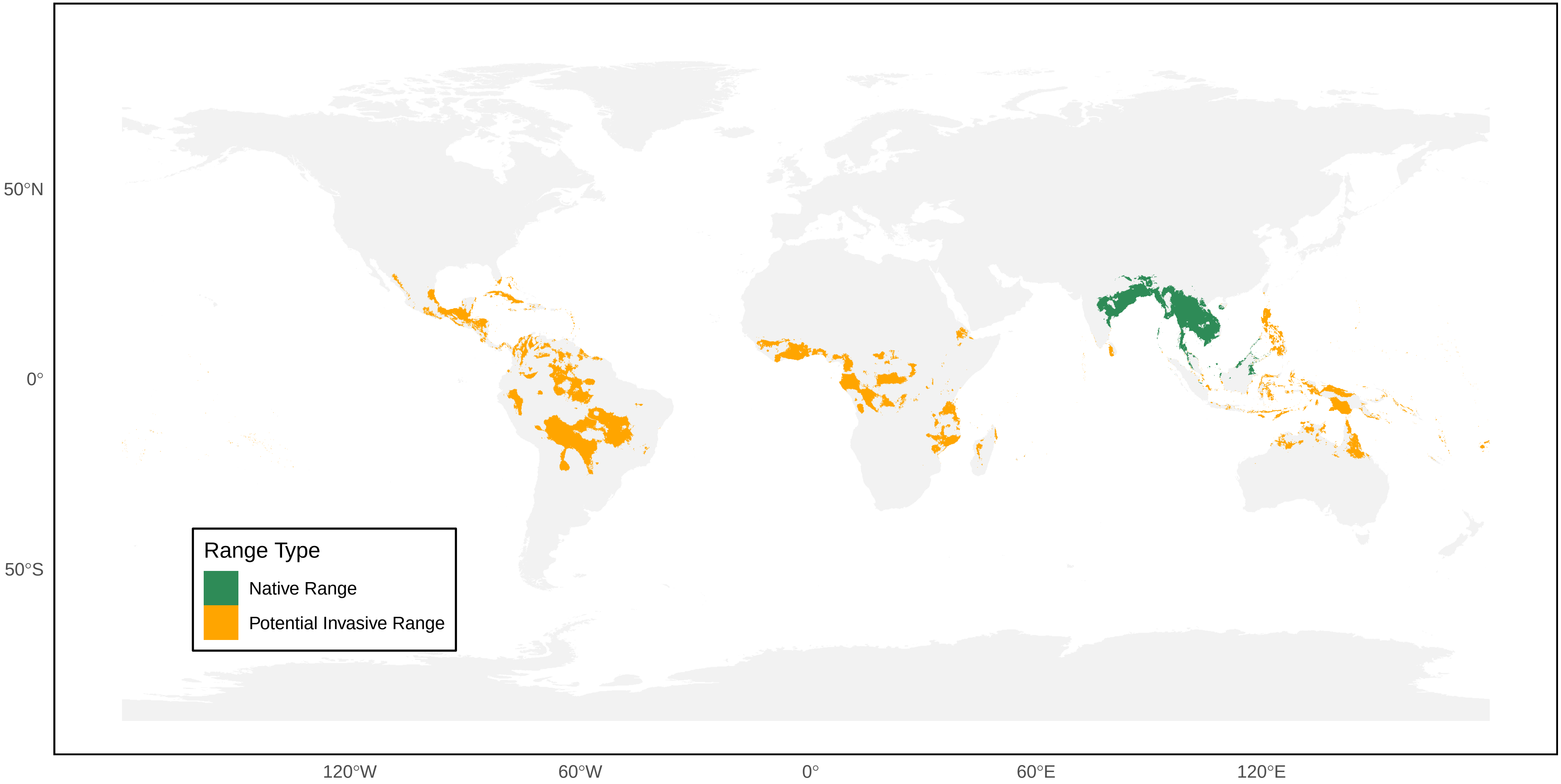

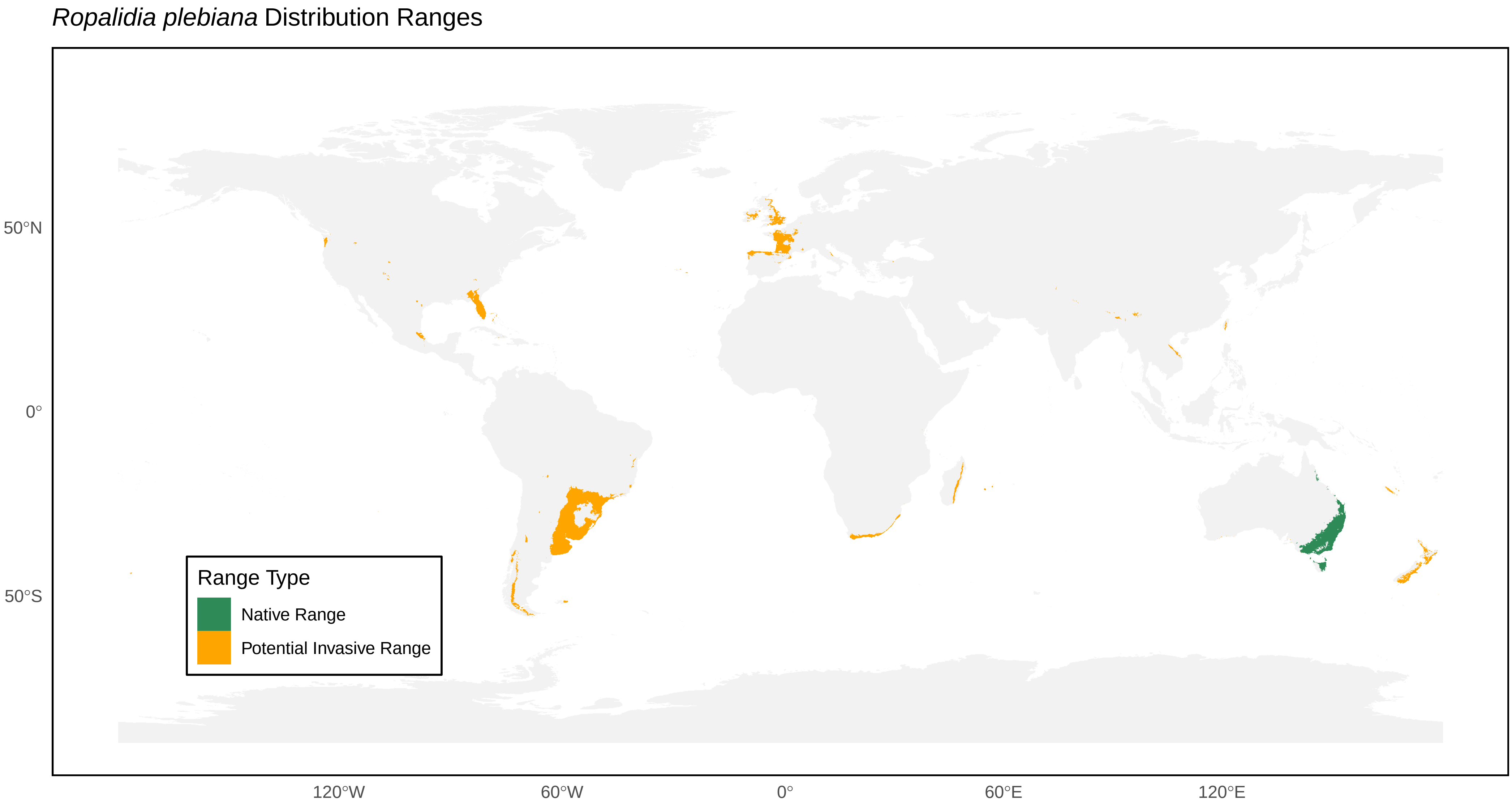

*Ropalidia plebiana* Distribution Ranges
Range Type
Native Range
Potential Invasive Range
50°N
0°
50°S
120°W
60°W
0°
60°E
120°E

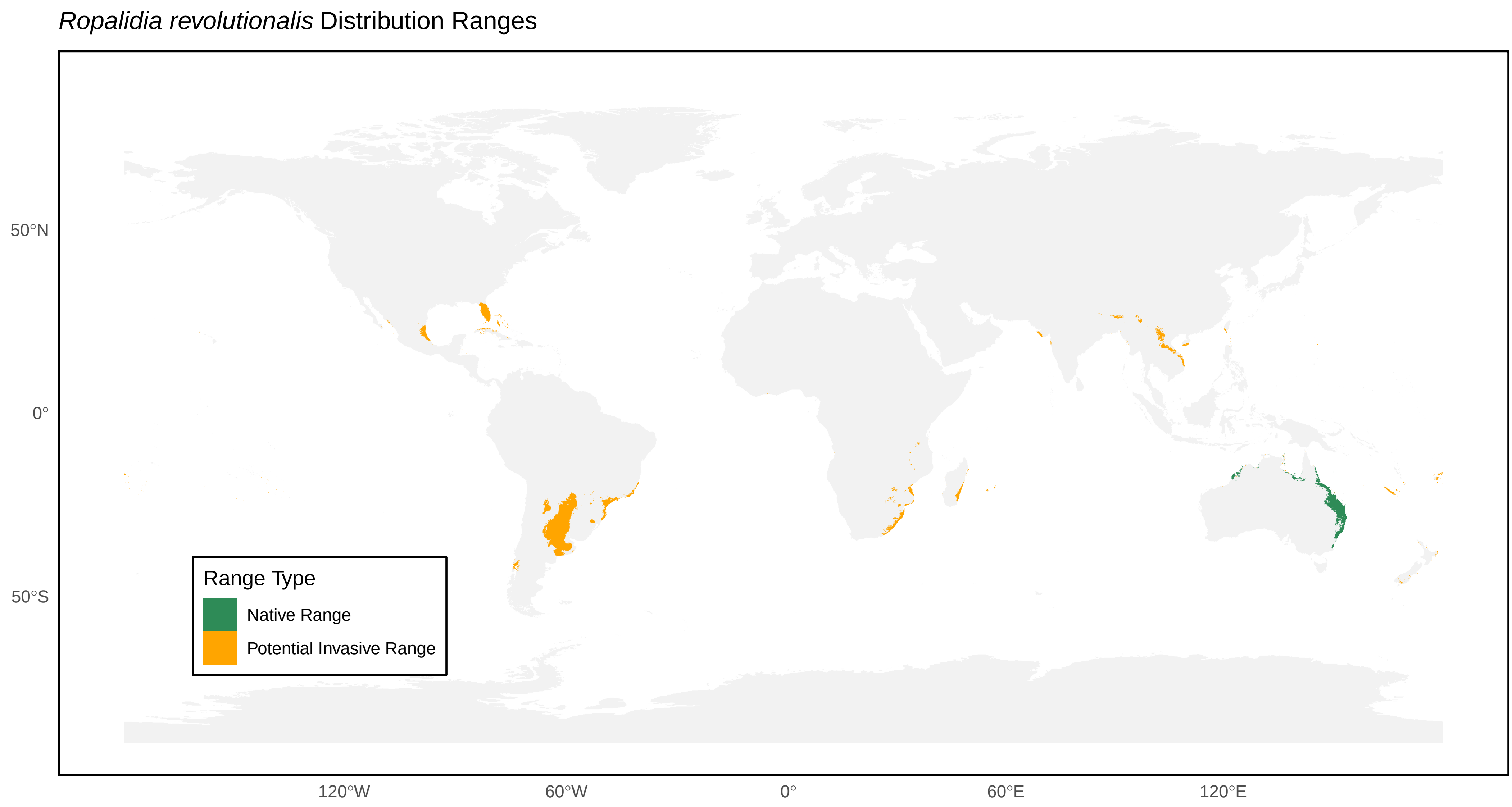

*Ropalidia revolutionalis* Distribution Ranges
50°N
0°
50°S
120°W
60°W
0°
60°E
120°E
Range Type
Native Range
Potential Invasive Range

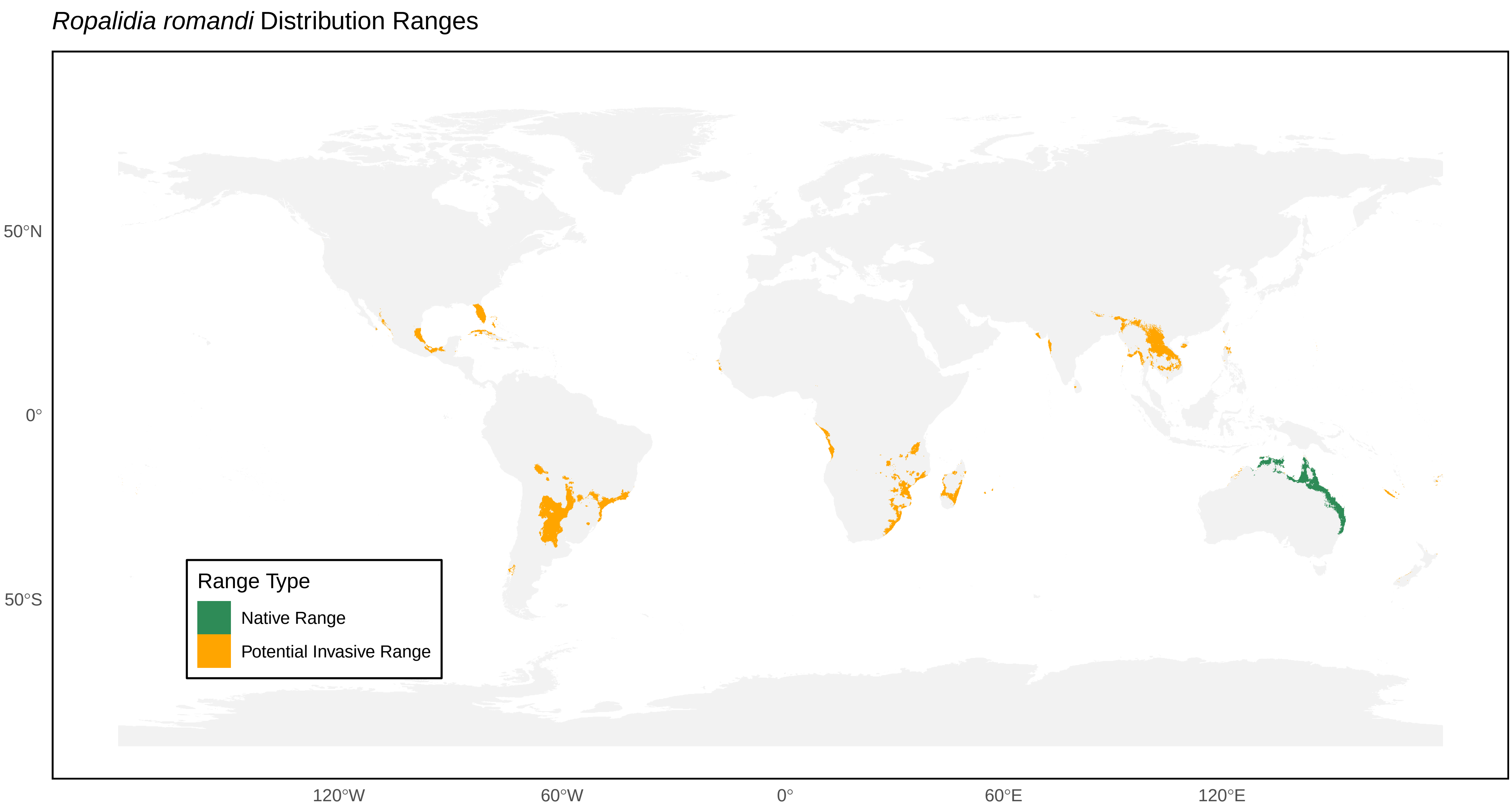

*Ropalidia romandi* Distribution Ranges
50°N
0°
50°S
120°W
60°W
0°
60°E
120°E
Range Type
Native Range
Potential Invasive Range

*Ropalidia rufoplagiata* Distribution Ranges

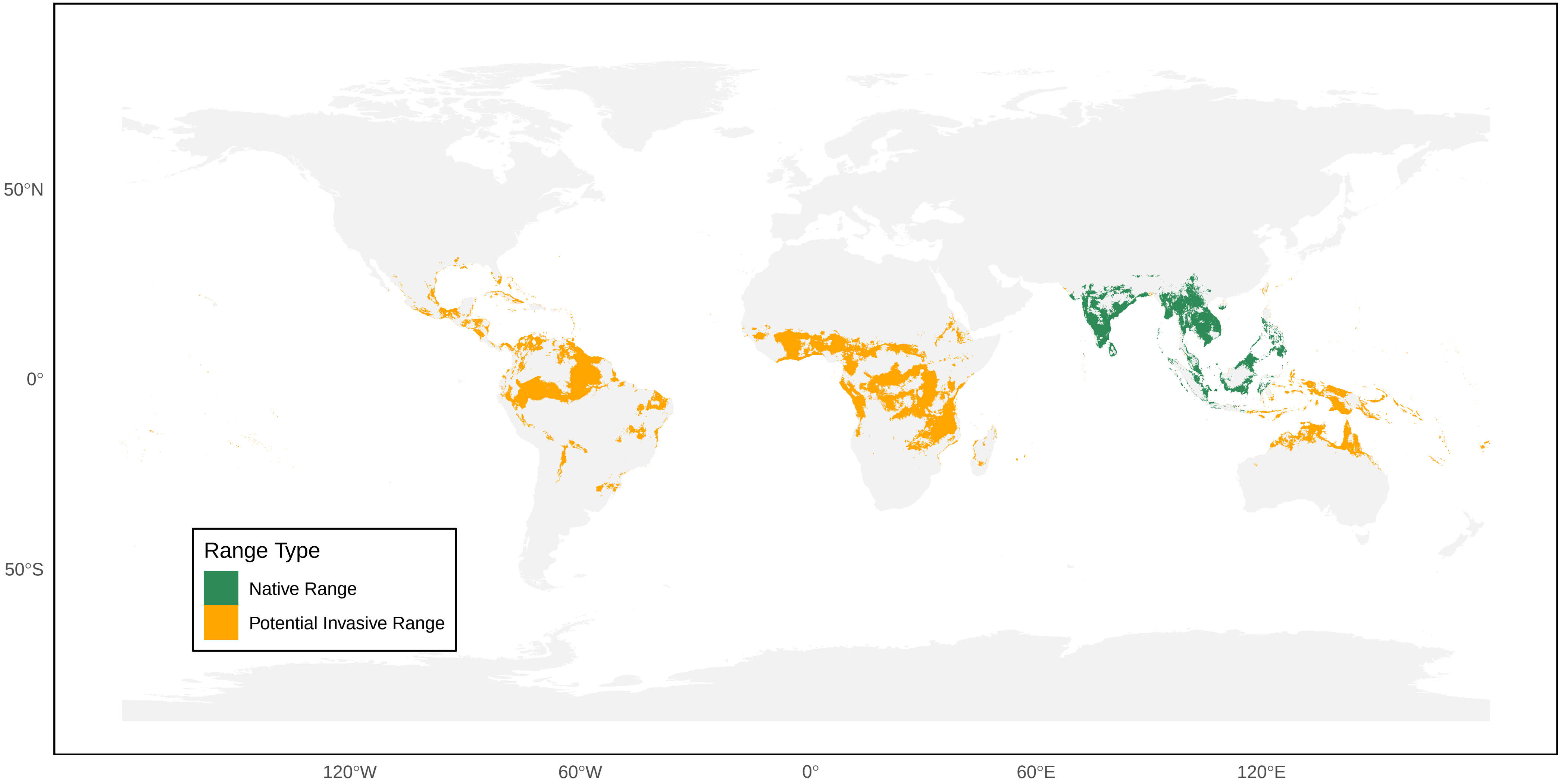

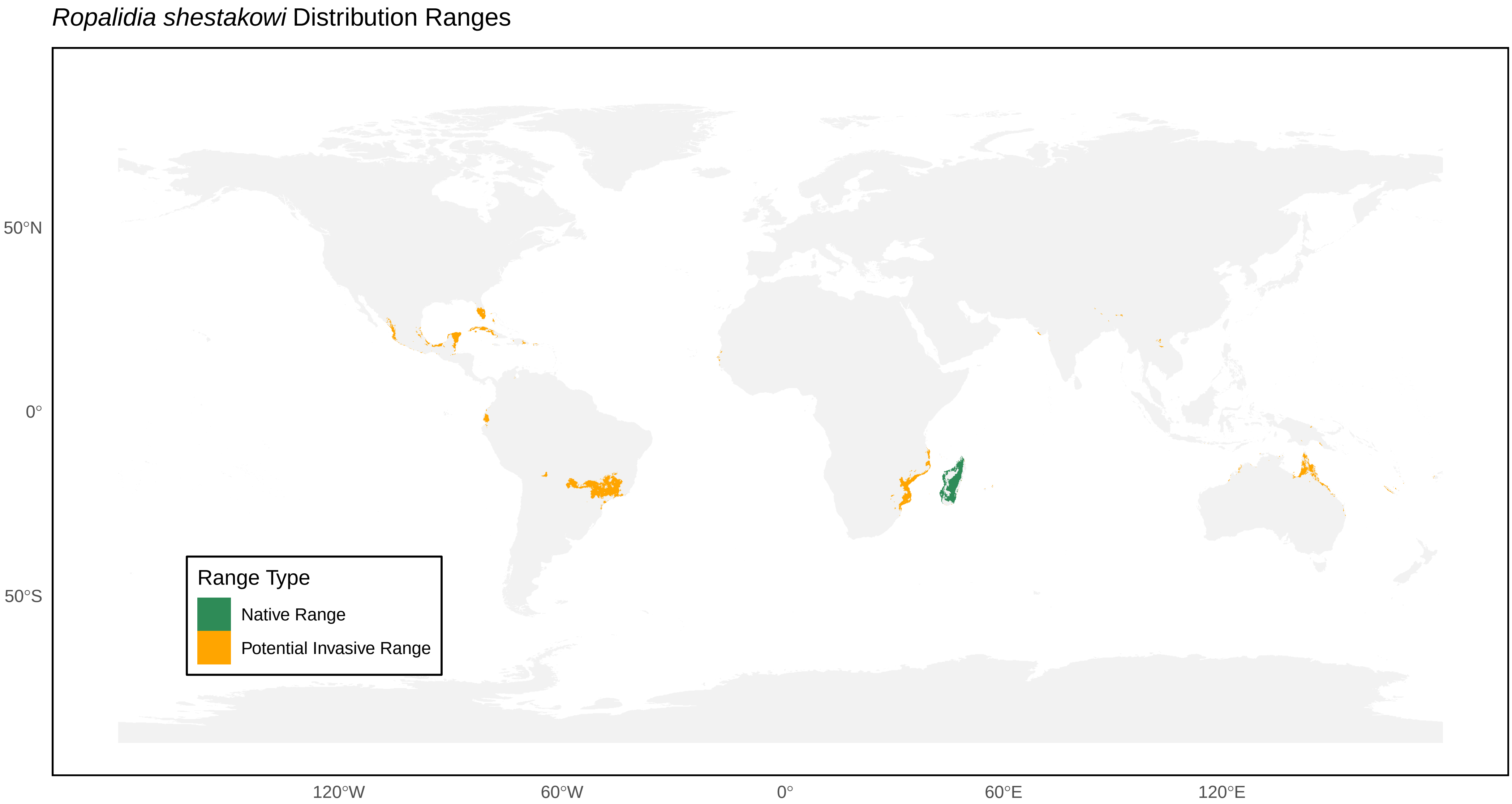
*Ropalidia shestakowi* Distribution Ranges
50°N
0°
50°S
120°W
60°W
0°
60°E
120°E
Range Type
Native Range
Potential Invasive Range

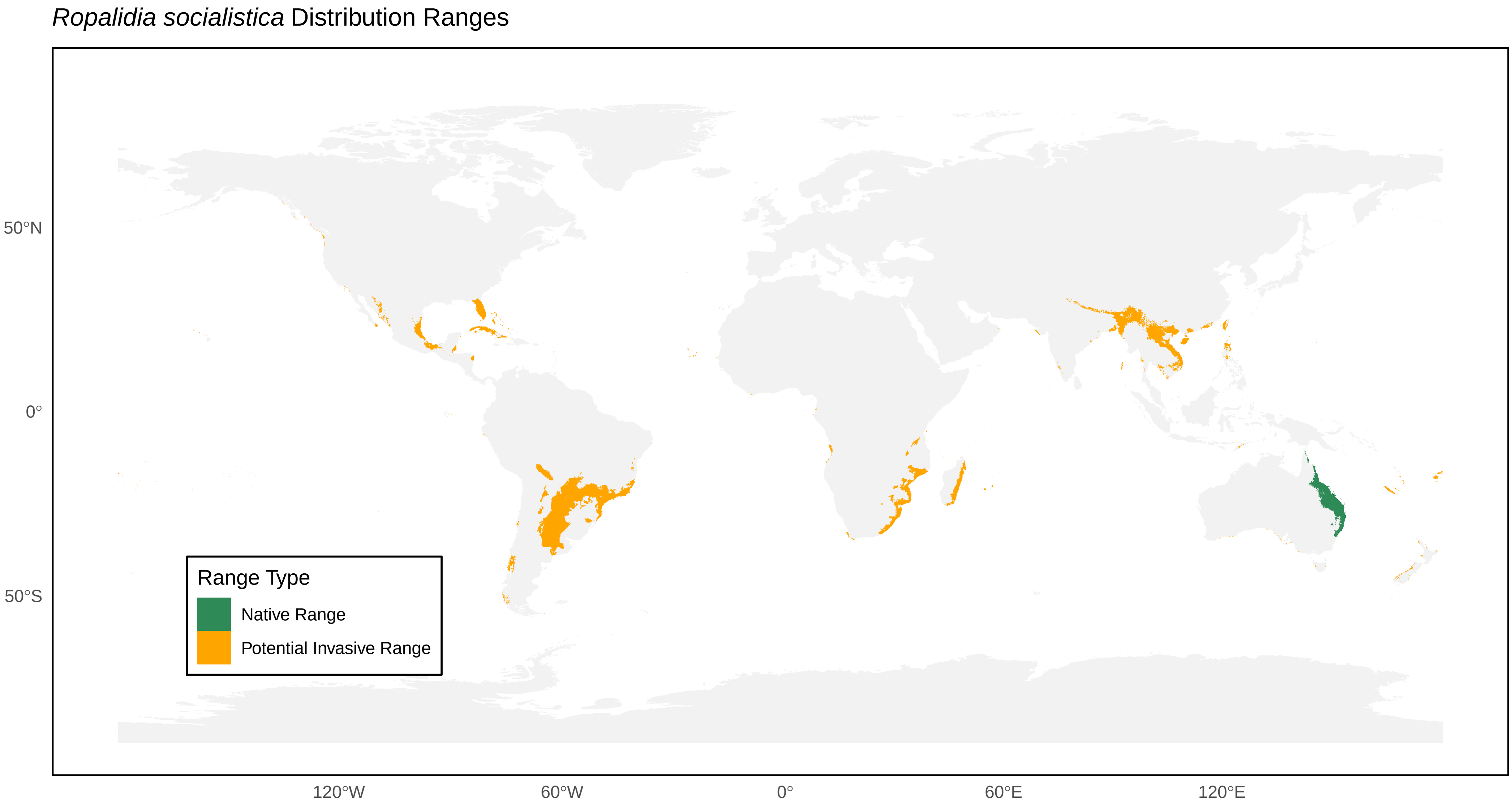

*Ropalidia socialistica* Distribution Ranges
50°N
0°
50°S
120°W
60°W
0°
60°E
120°E
Range Type
Native Range
Potential Invasive Range

*Ropalidia stigma* Distribution Ranges

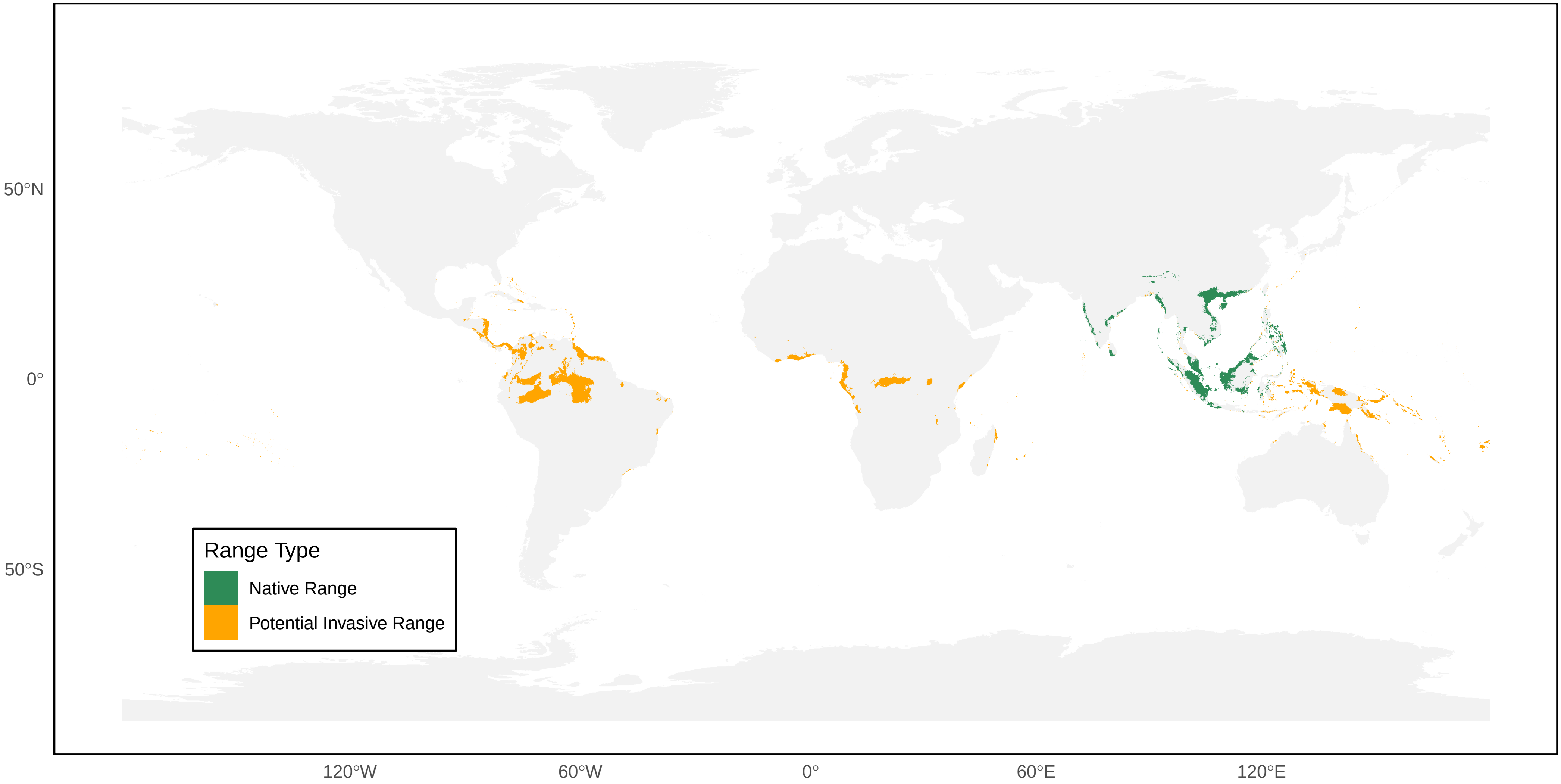

*Ropalidia sumatrae* Distribution Ranges

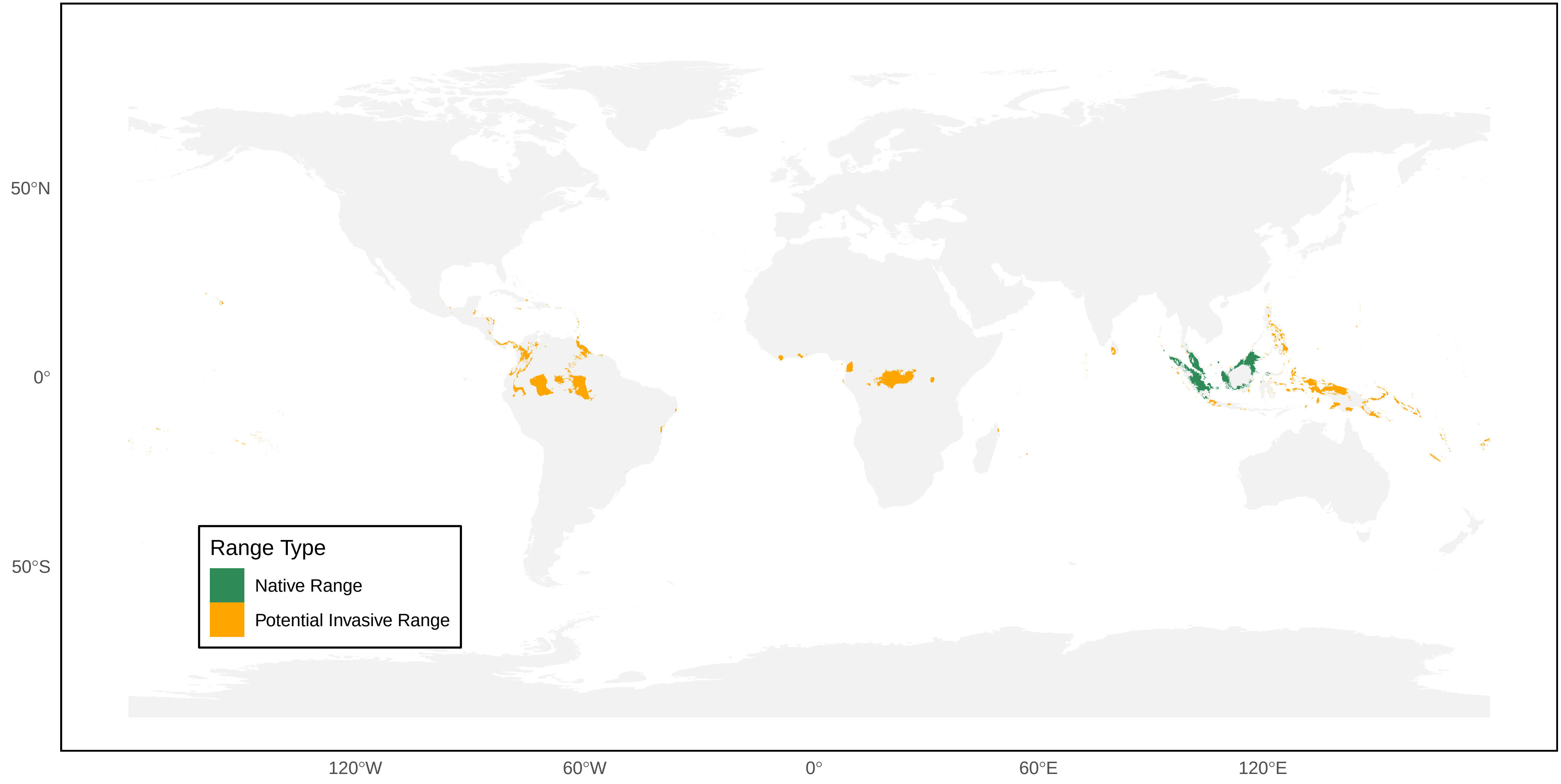

*Ropalidia taiwana* Distribution Ranges

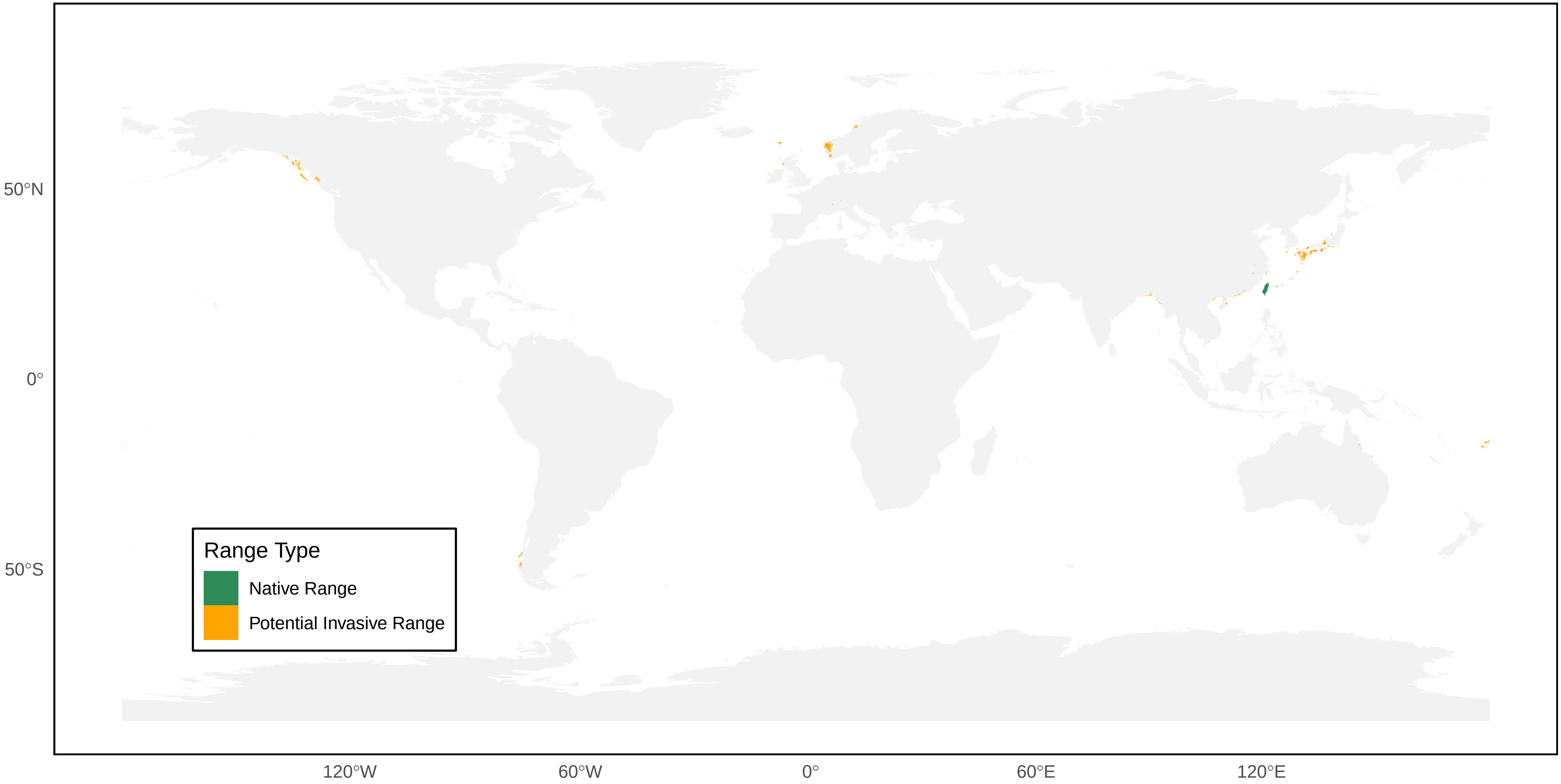

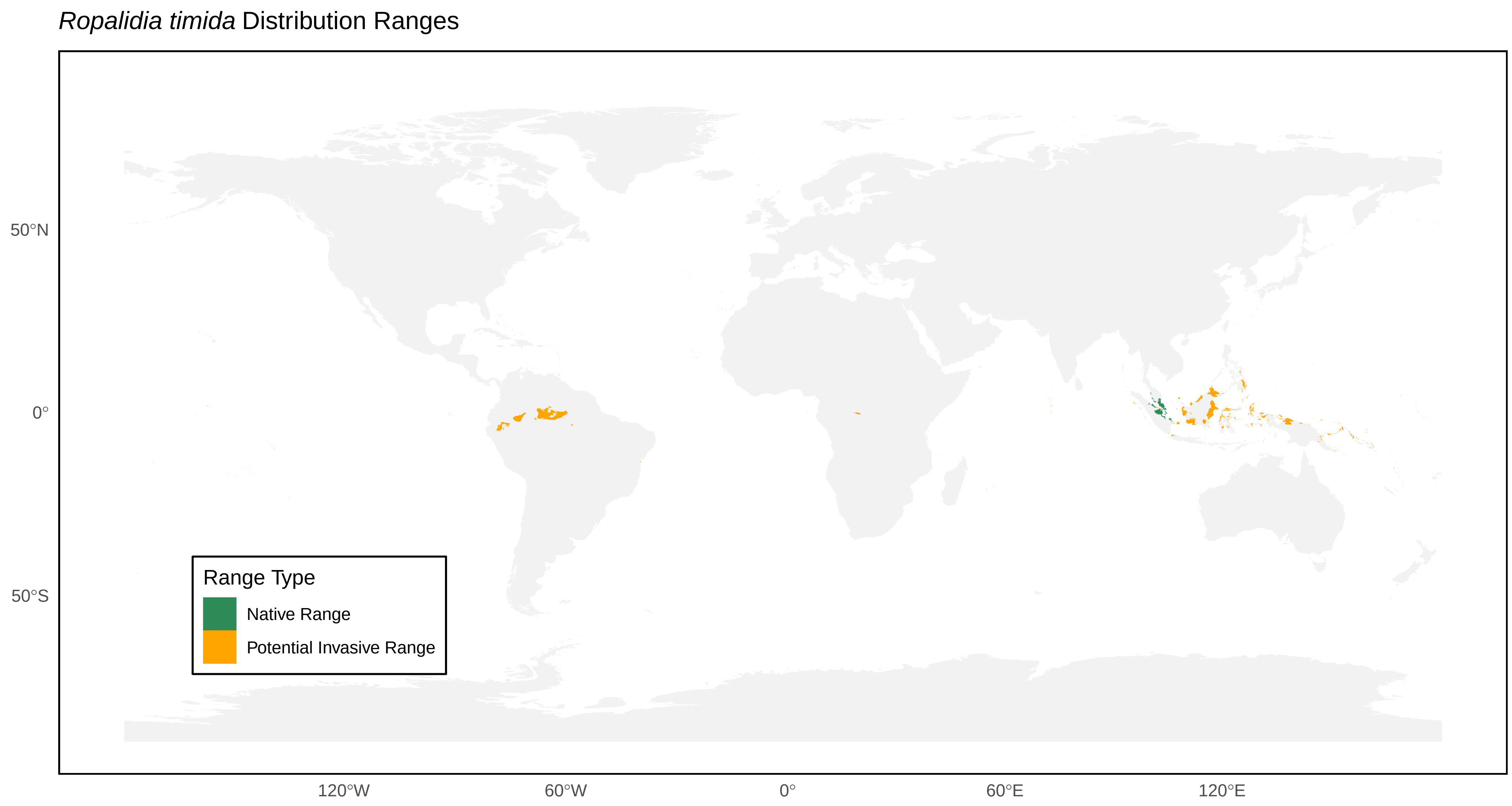
*Ropalidia timida* Distribution Ranges
50°N
0°
50°S
120°W
60°W
0°
60°E
120°E
Range Type
Native Range
Potential Invasive Range

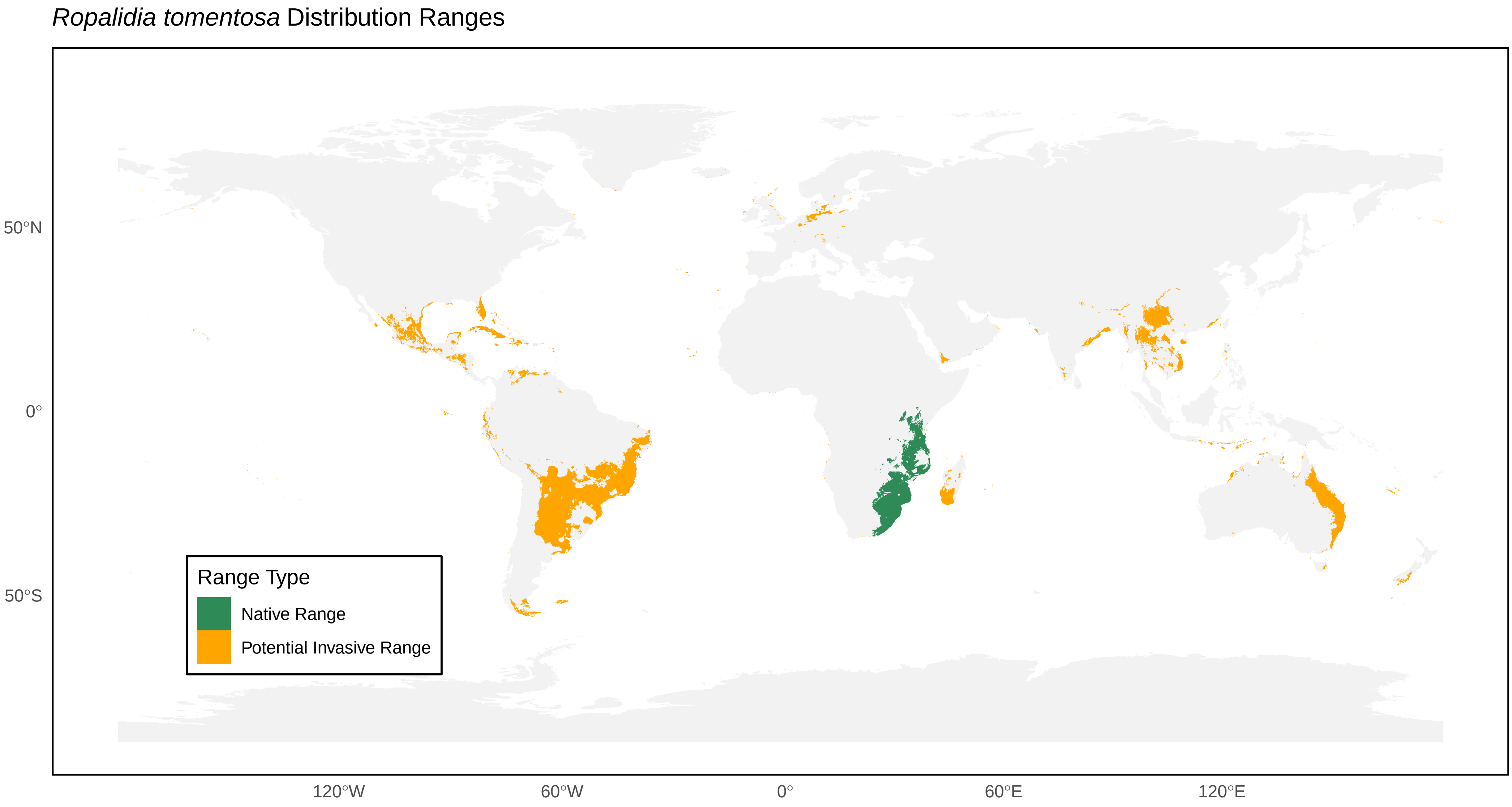

*Ropalidia tomentosa* Distribution Ranges
50°N
0°
50°S
120°W
60°W
0°
60°E
120°E
Range Type
Native Range
Potential Invasive Range

*Ropalidia variegata* Distribution Ranges

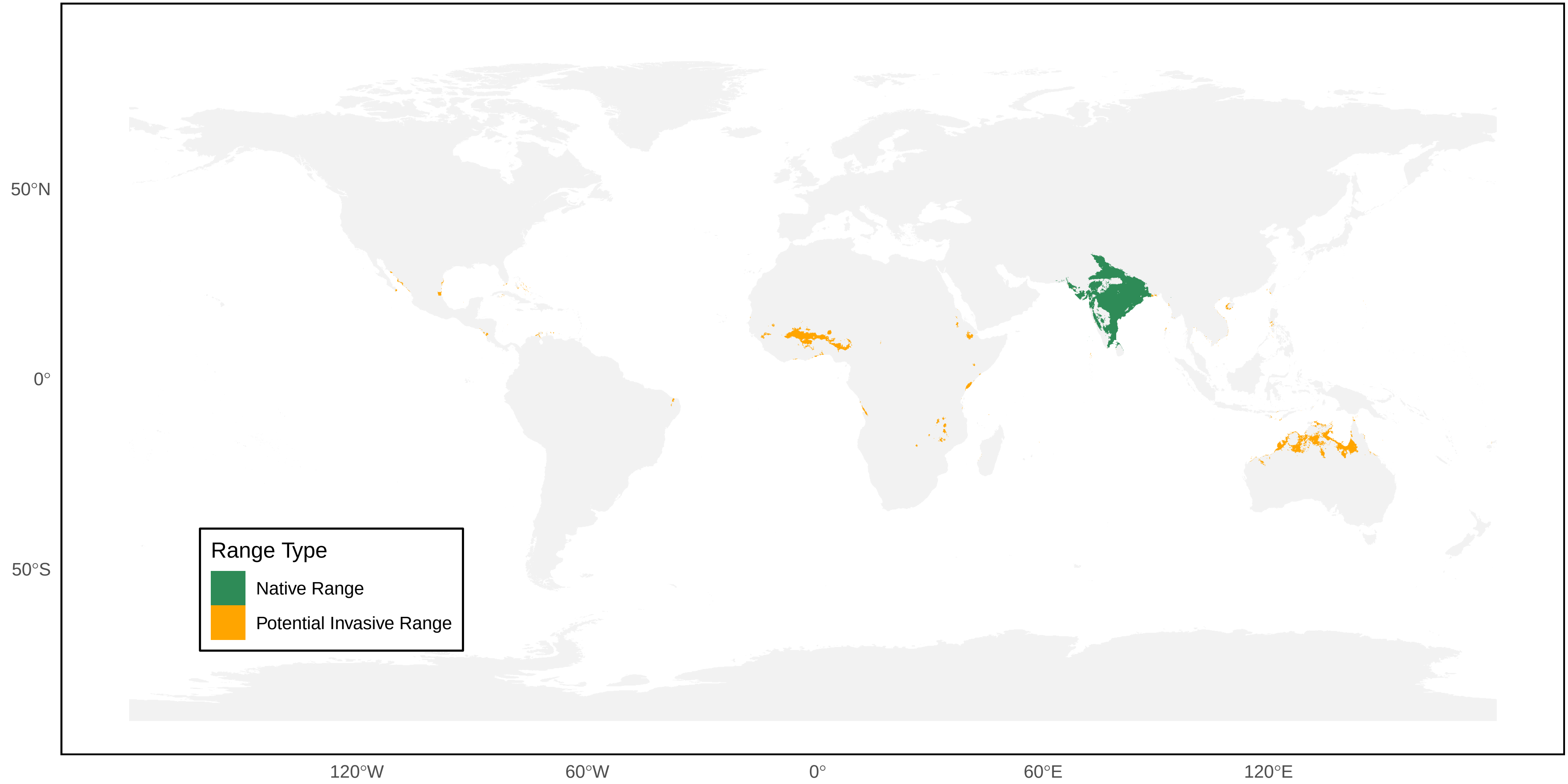

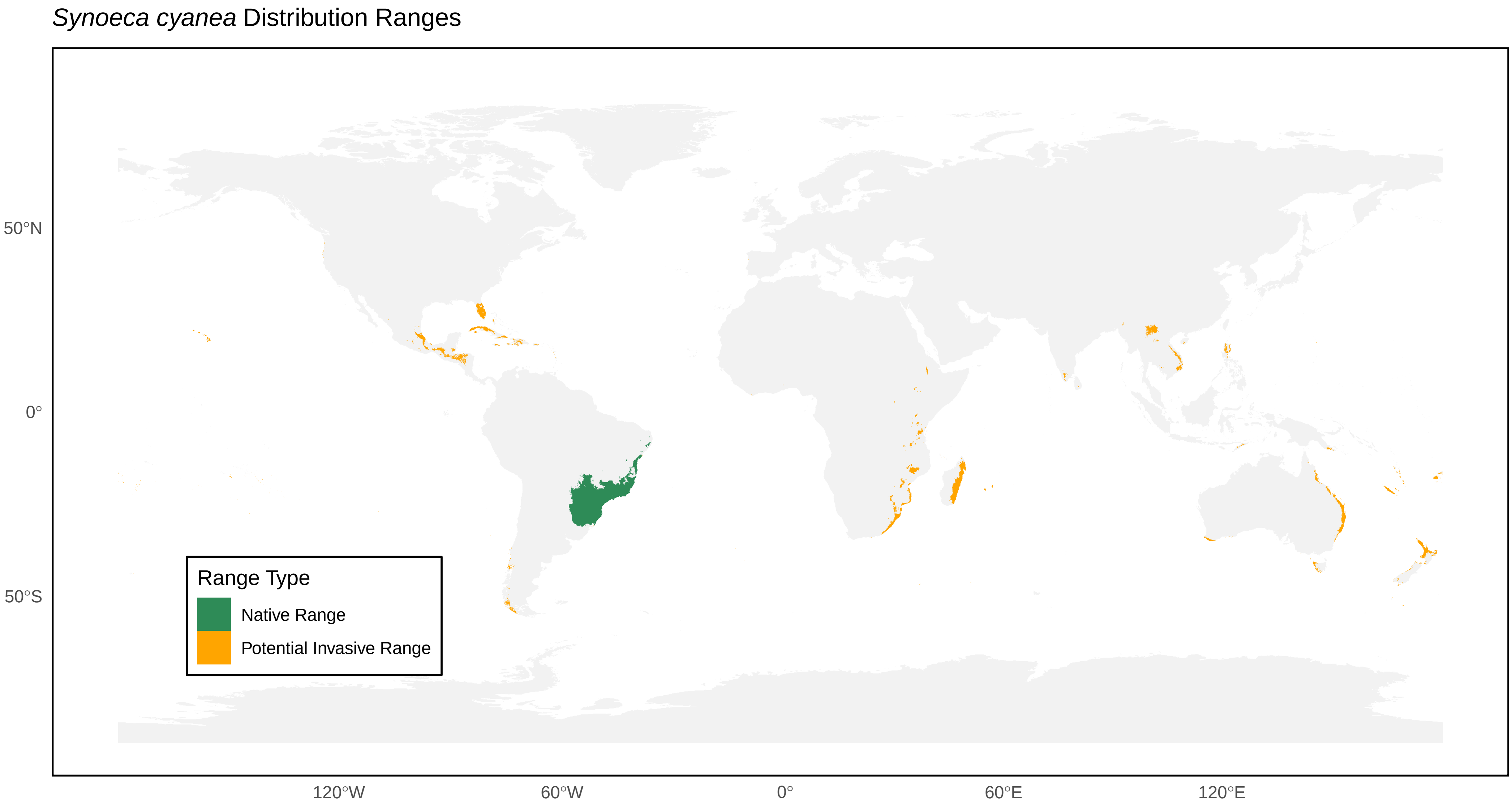

*Synoeca cyanea* Distribution Ranges
50°N
0°
50°S
120°W
60°W
0°
60°E
120°E
Range Type
Native Range
Potential Invasive Range

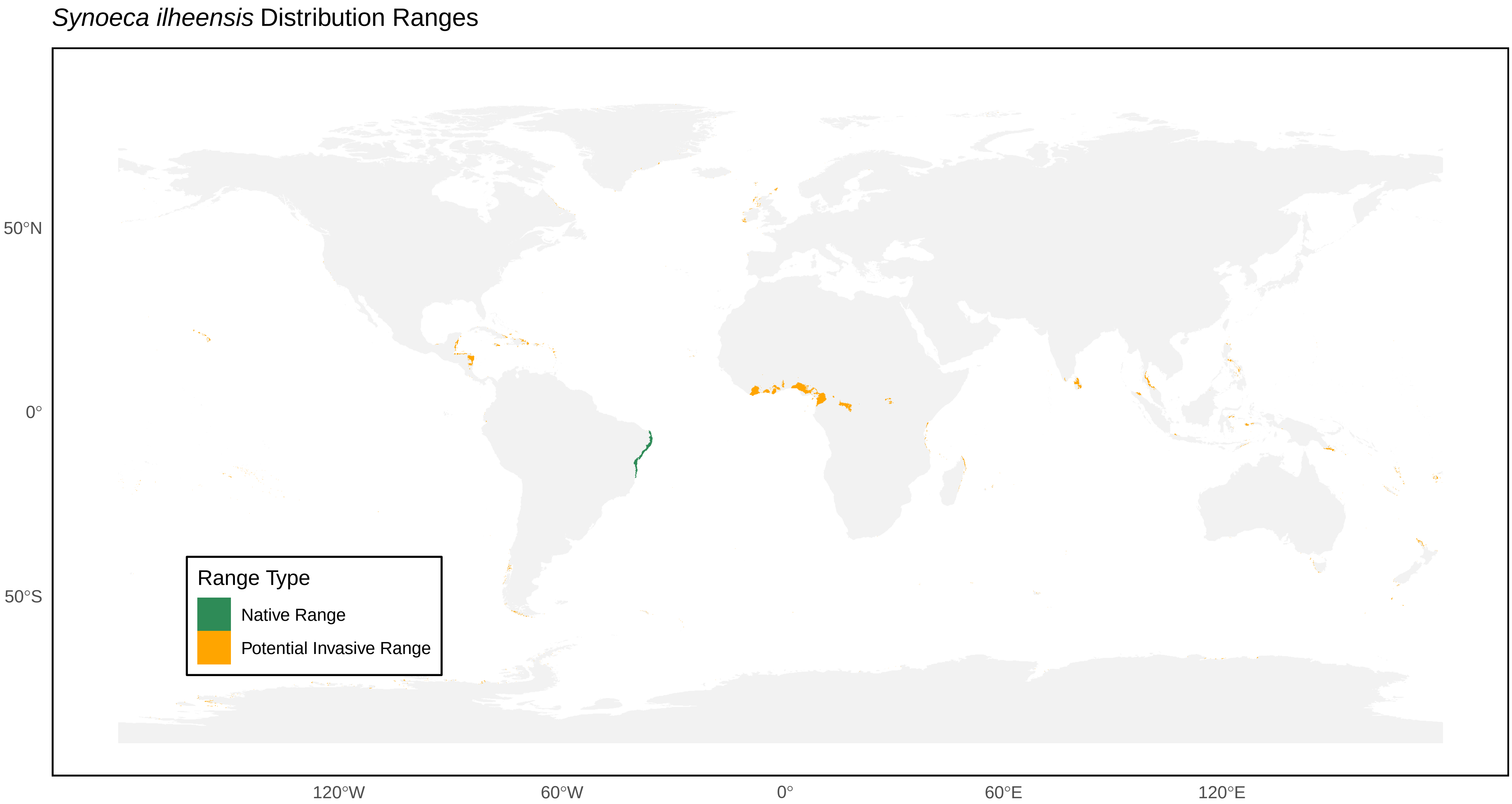

*Synoeca ilheensis* Distribution Ranges
50°N
0°
50°S
120°W
60°W
0°
60°E
120°E
Range Type
Native Range
Potential Invasive Range

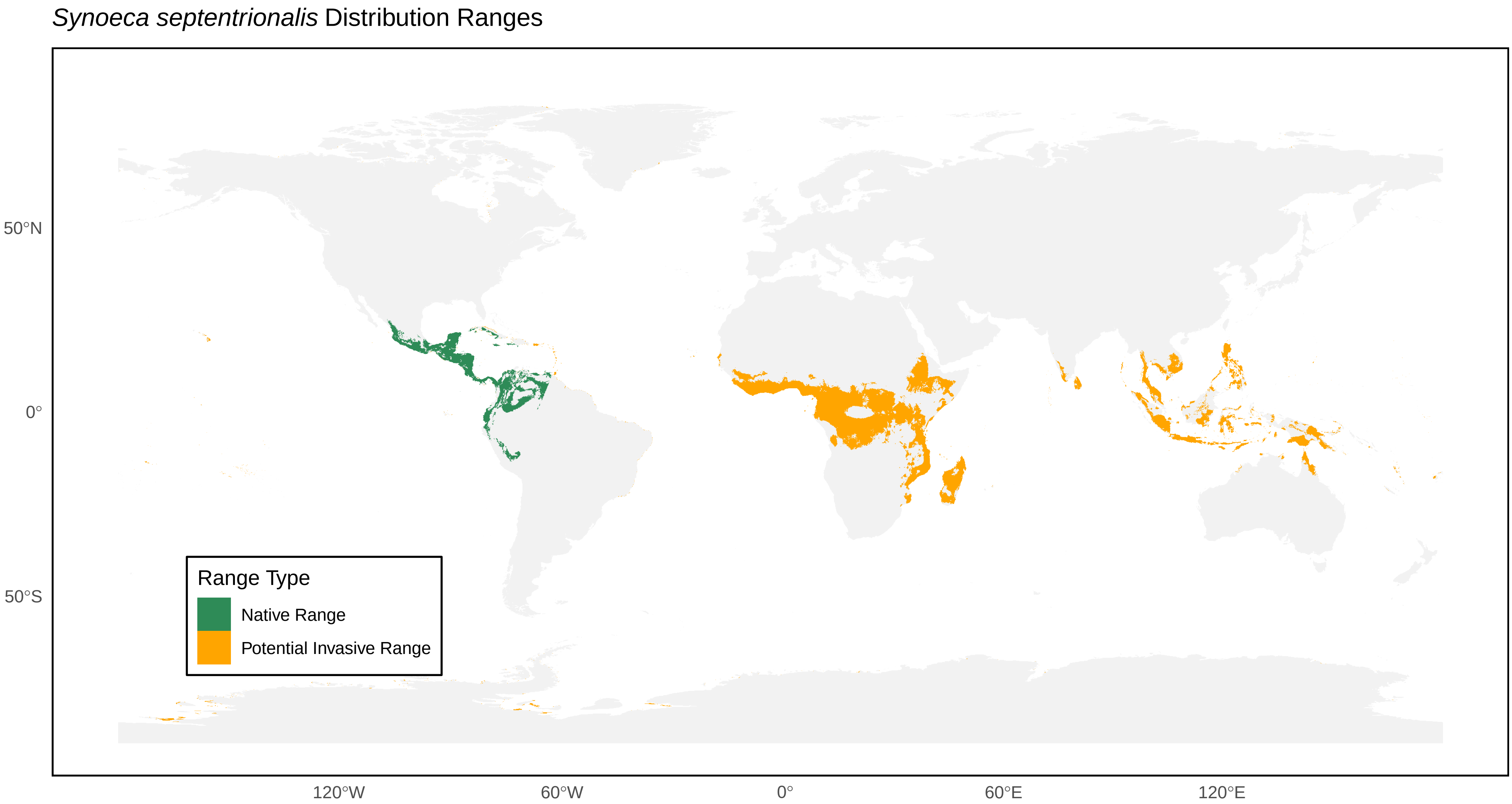
*Synoeca septentrionalis* Distribution Ranges
50°N
0°
50°S
120°W
60°W
0°
60°E
120°E
Range Type
Native Range
Potential Invasive Range

*Synoeca surinama* Distribution Ranges

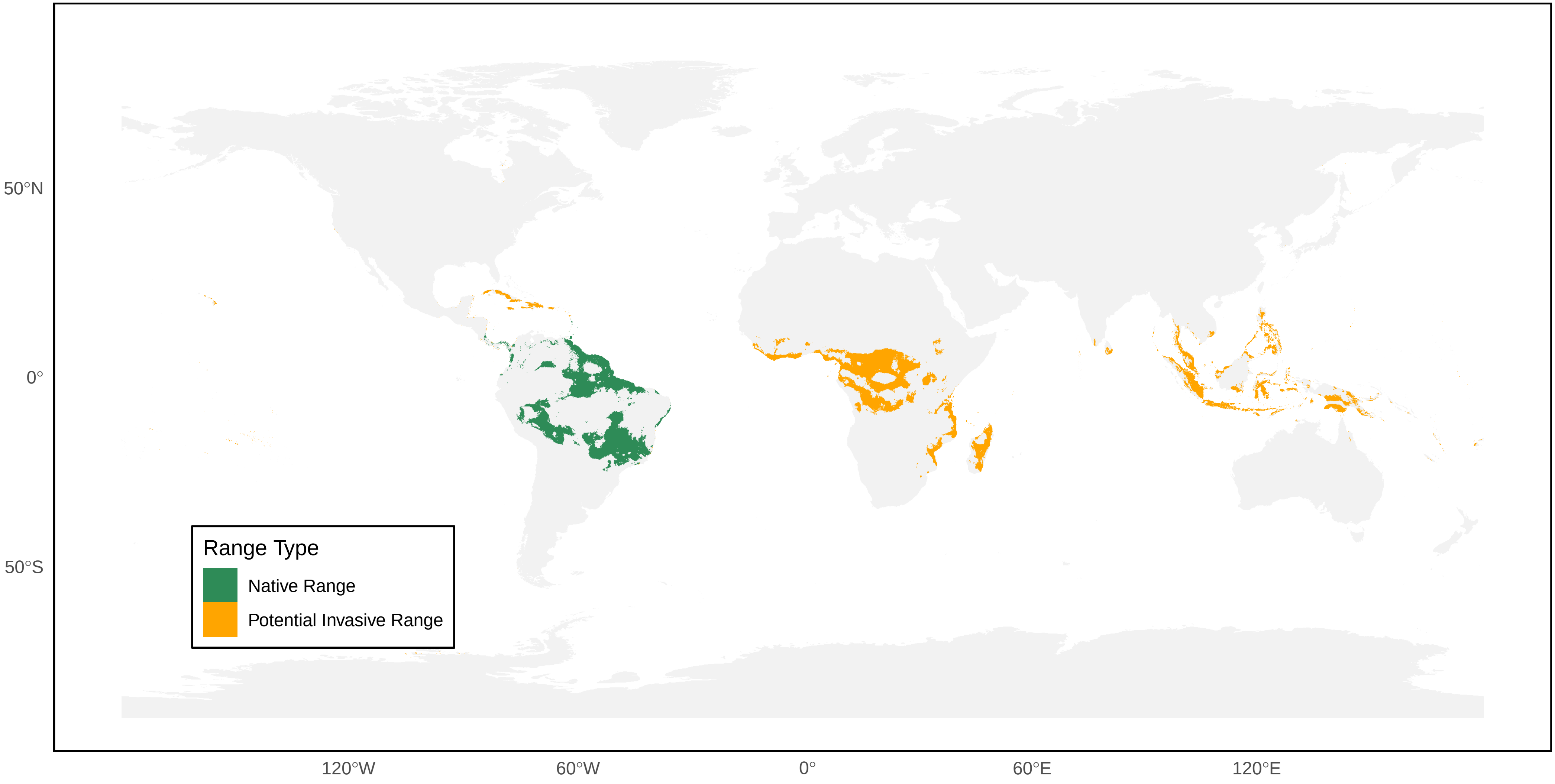

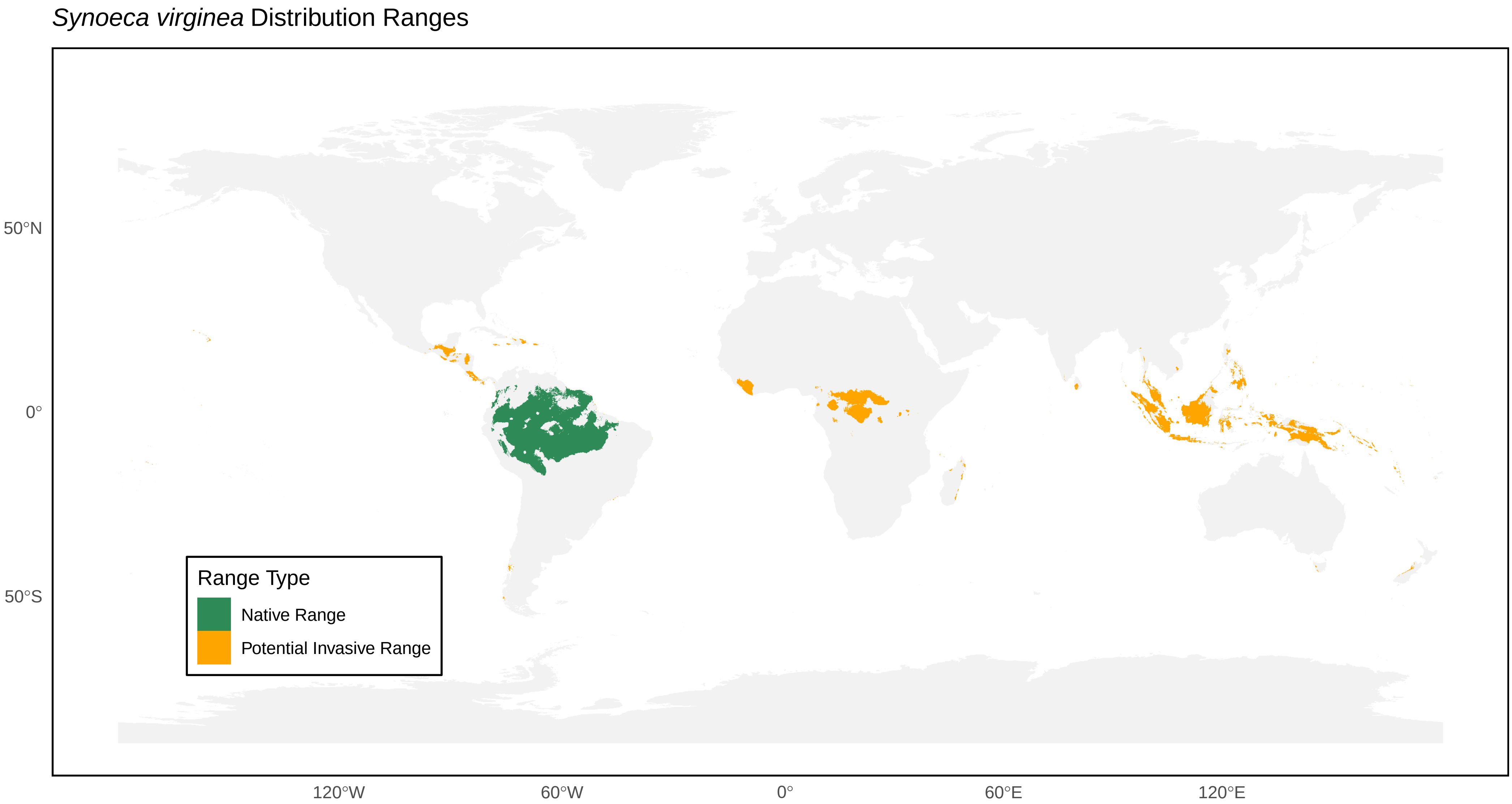

*Synoeca virginea* Distribution Ranges
50°N
0°
50°S
120°W
60°W
0°
60°E
120°E
Range Type
Native Range
Potential Invasive Range

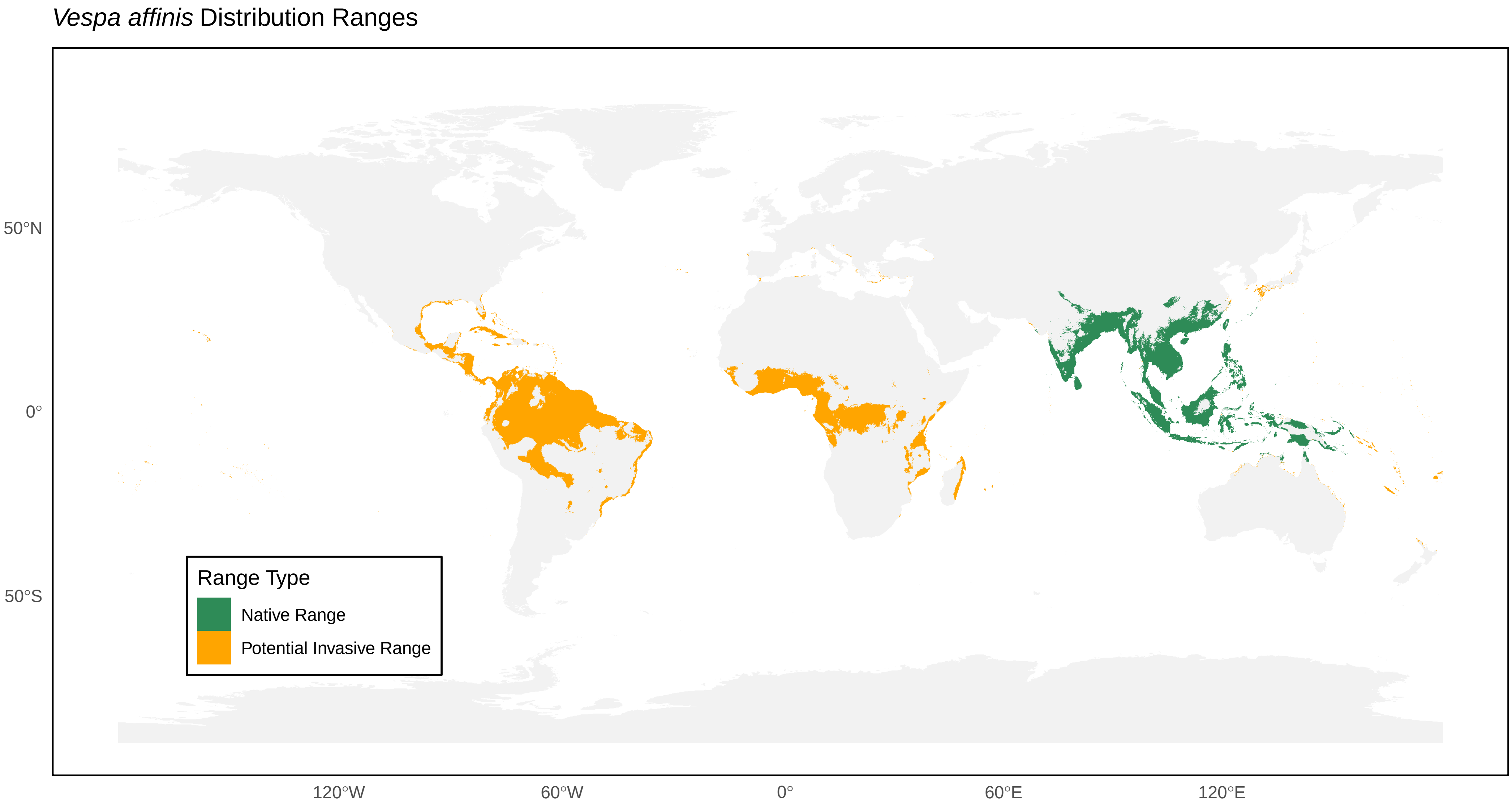

*Vespa affinis* Distribution Ranges
50°N
0°
50°S
120°W
60°W
0°
60°E
120°E
Range Type
Native Range
Potential Invasive Range

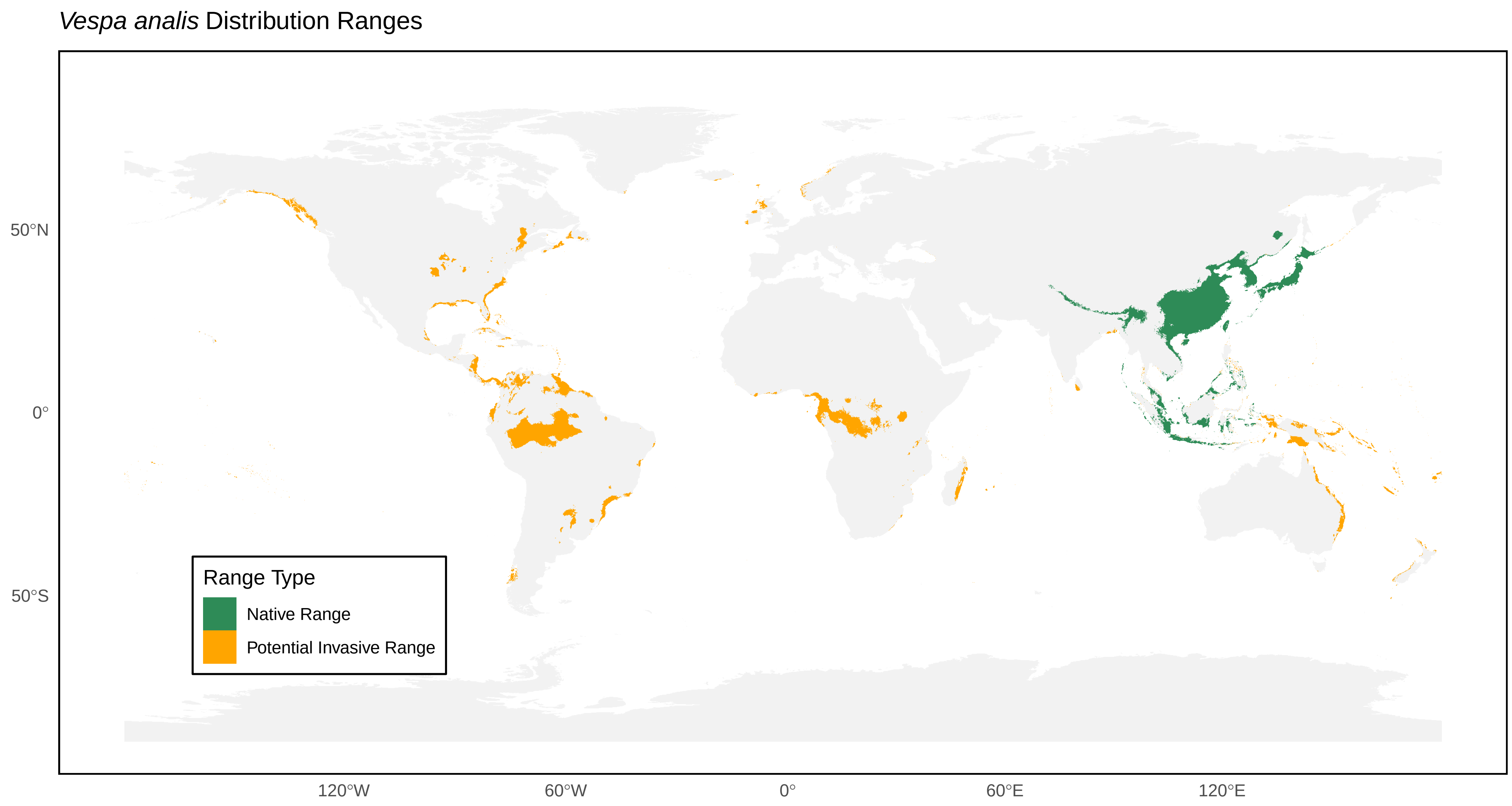

*Vespa analis* Distribution Ranges
50°N
0°
50°S
Range Type
Native Range
Potential Invasive Range
120°W
60°W
0°
60°E
120°E

*Vespa basalis* Distribution Ranges

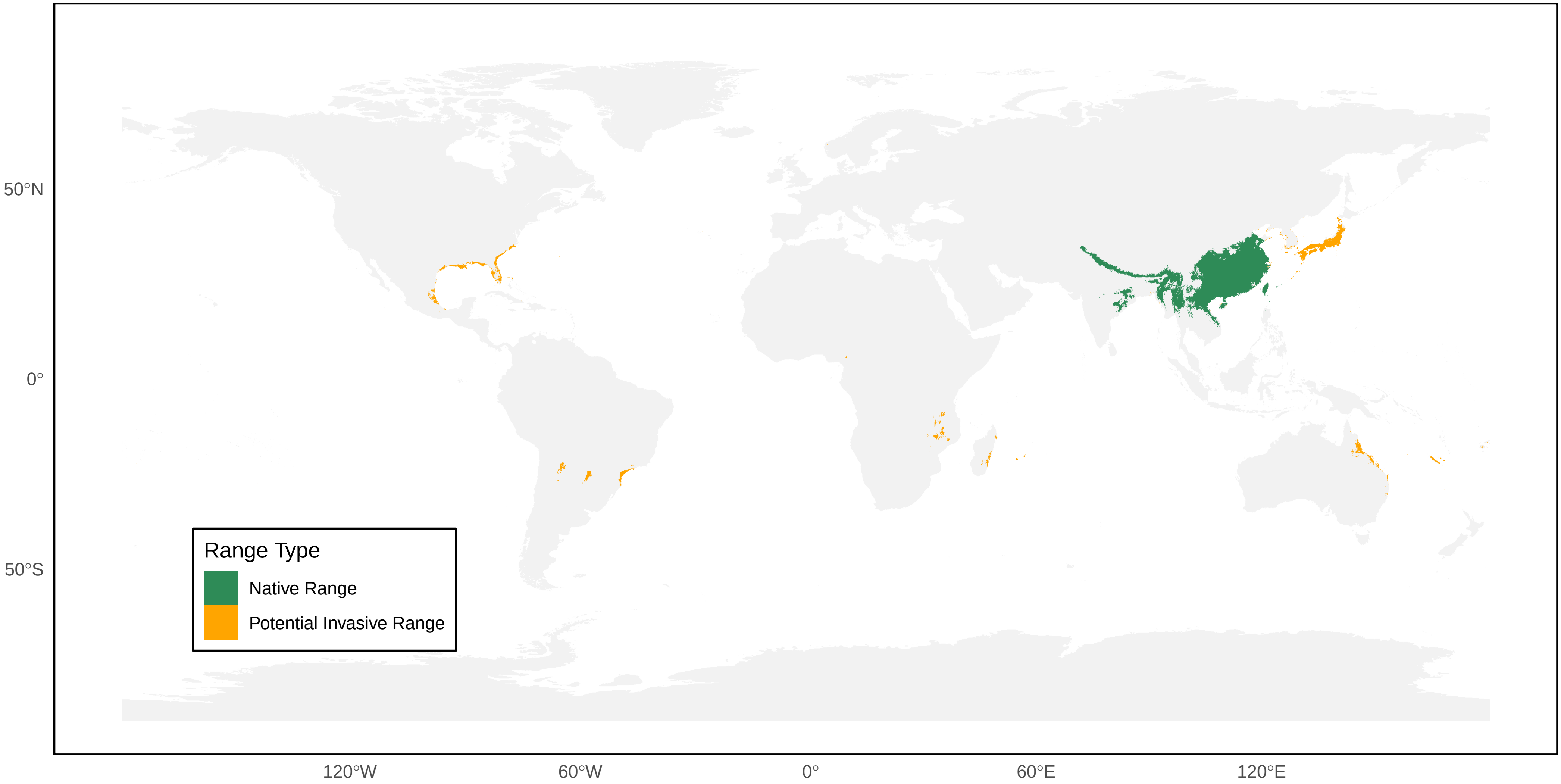

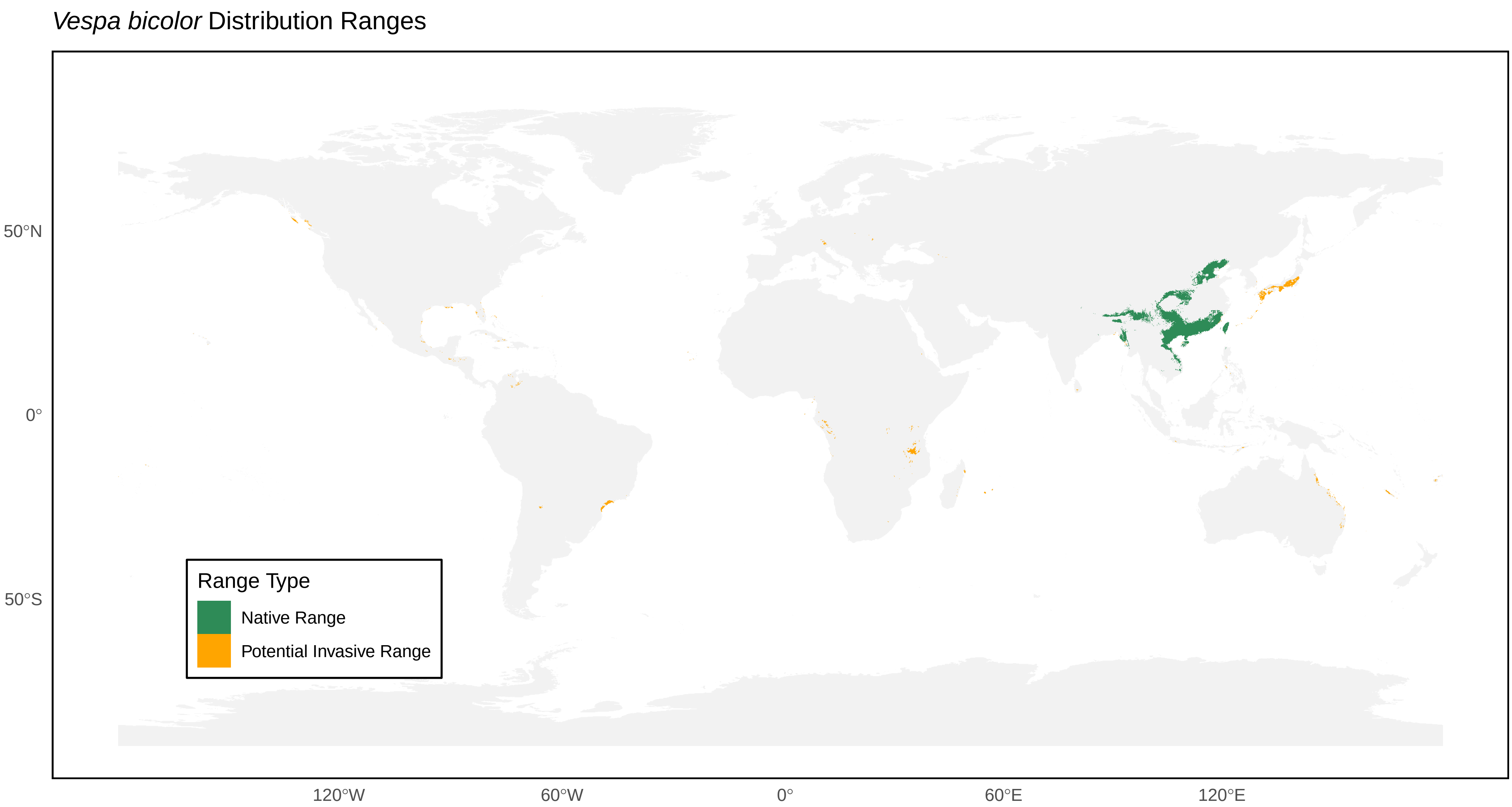
*Vespa bicolor* Distribution Ranges
50°N
0°
50°S
120°W
60°W
0°
60°E
120°E
Range Type
Native Range
Potential Invasive Range

*Vespa binghami* Distribution Ranges

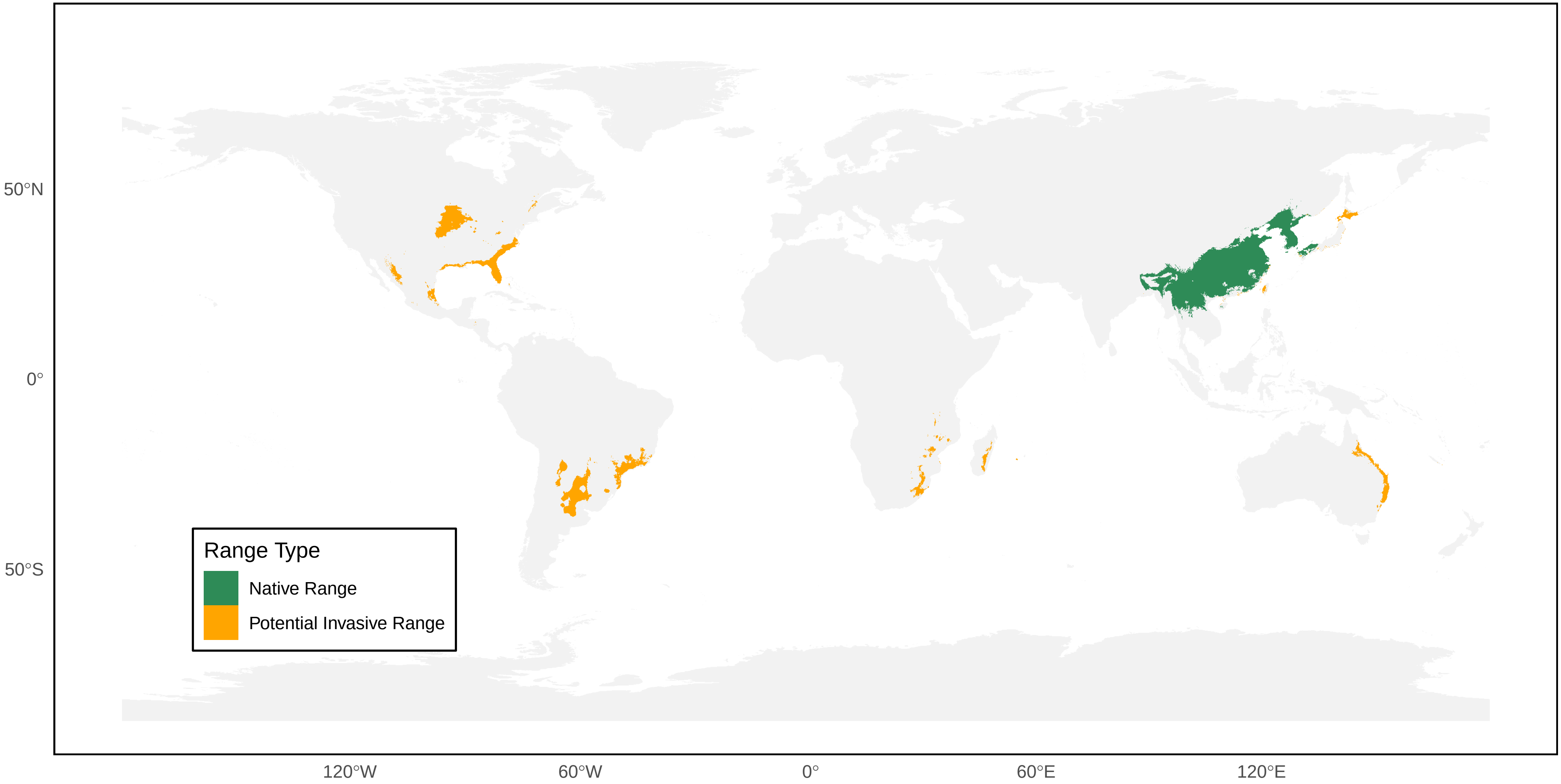

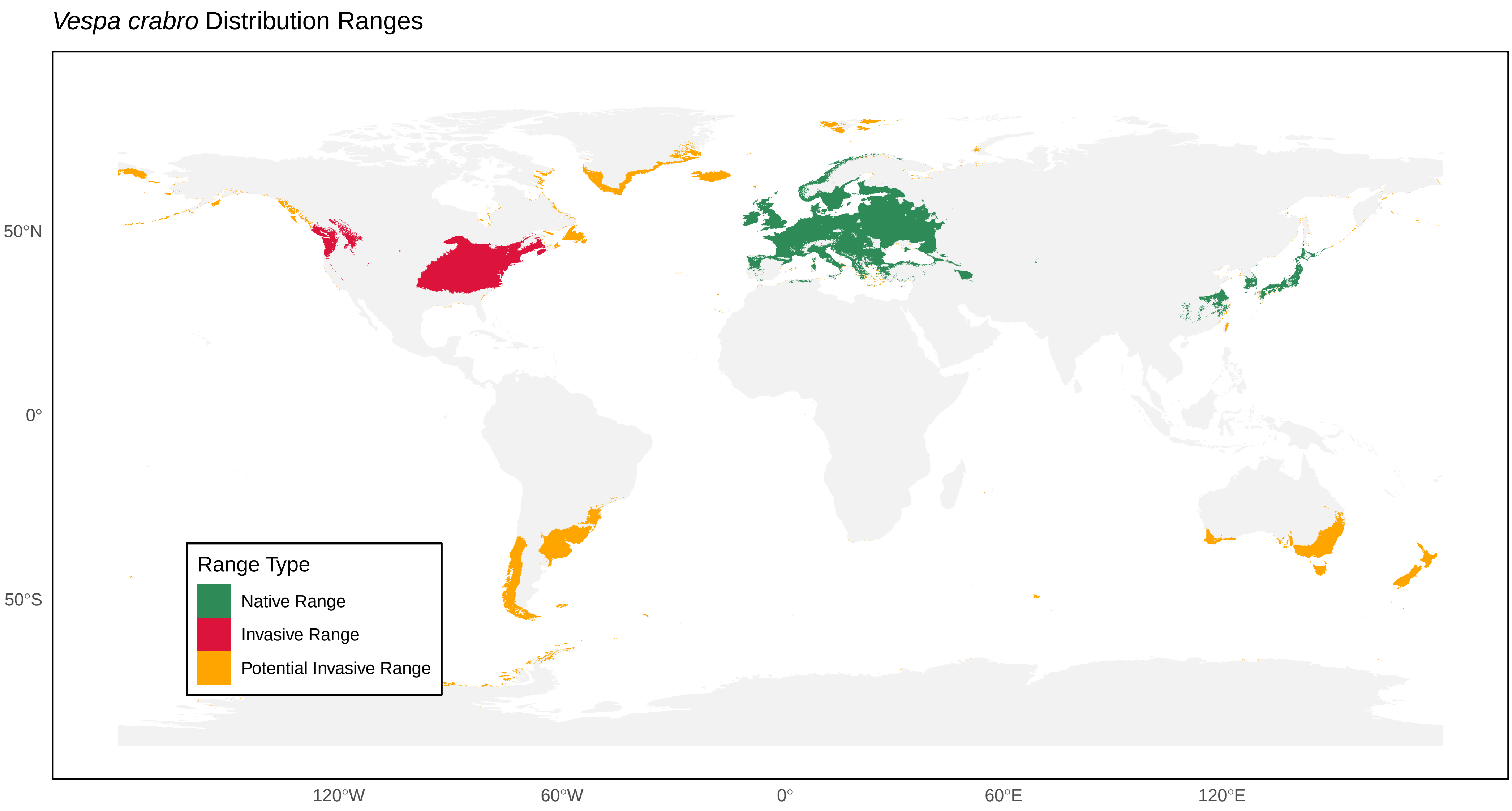
*Vespa crabro* Distribution Ranges
50°N
0°
50°S
120°W
60°W
0°
60°E
120°E
Range Type
Native Range
Invasive Range
Potential Invasive Range

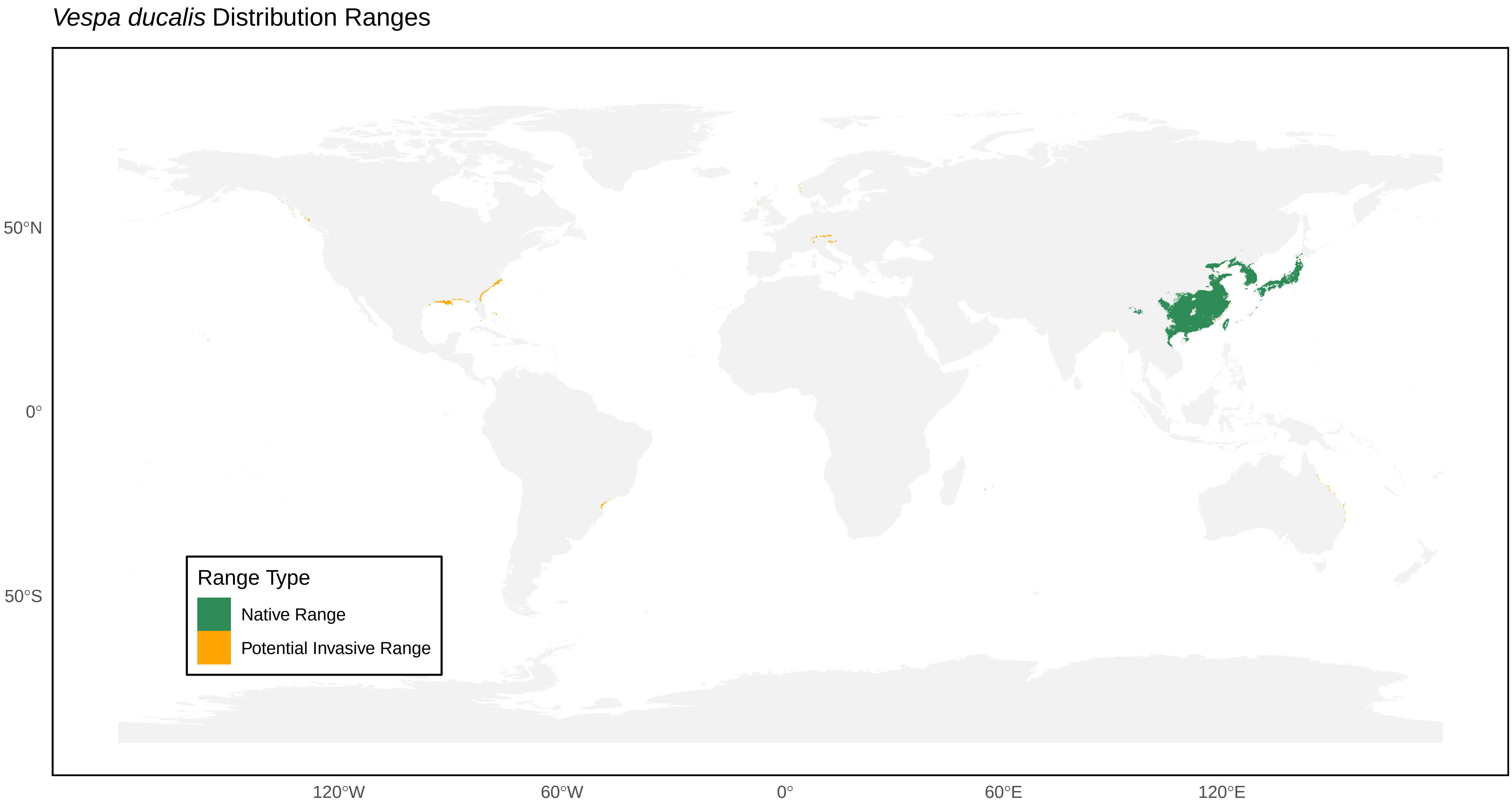
*Vespa ducalis* Distribution Ranges
50°N
0°
50°S
120°W
60°W
0°
60°E
120°E
Range Type
Native Range
Potential Invasive Range

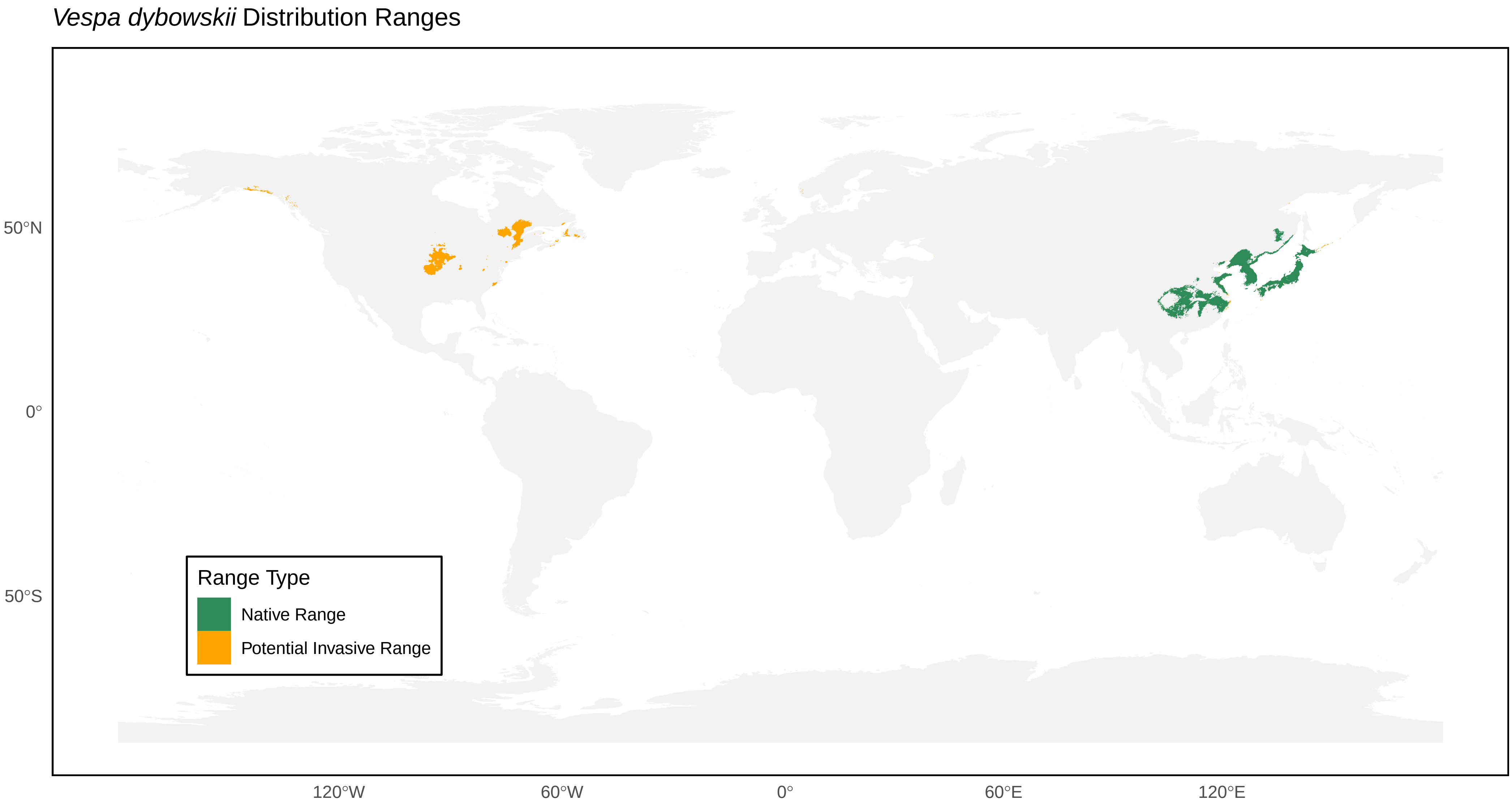

*Vespa dybowskii* Distribution Ranges
50°N
0°
50°S
120°W
60°W
0°
60°E
120°E
Range Type
Native Range
Potential Invasive Range

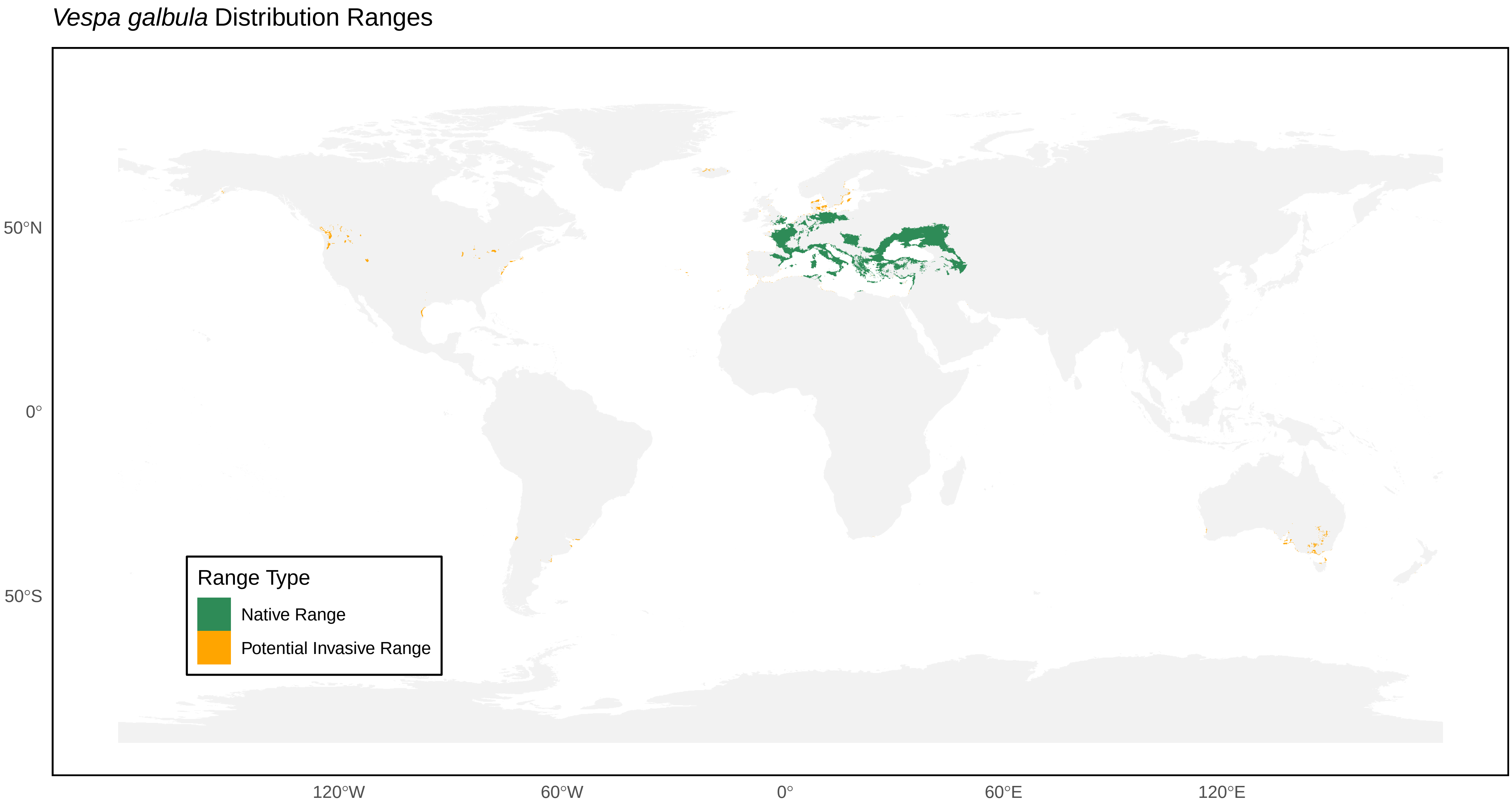

*Vespa galbula* Distribution Ranges
50°N
0°
50°S
120°W
60°W
0°
60°E
120°E
Range Type
Native Range
Potential Invasive Range

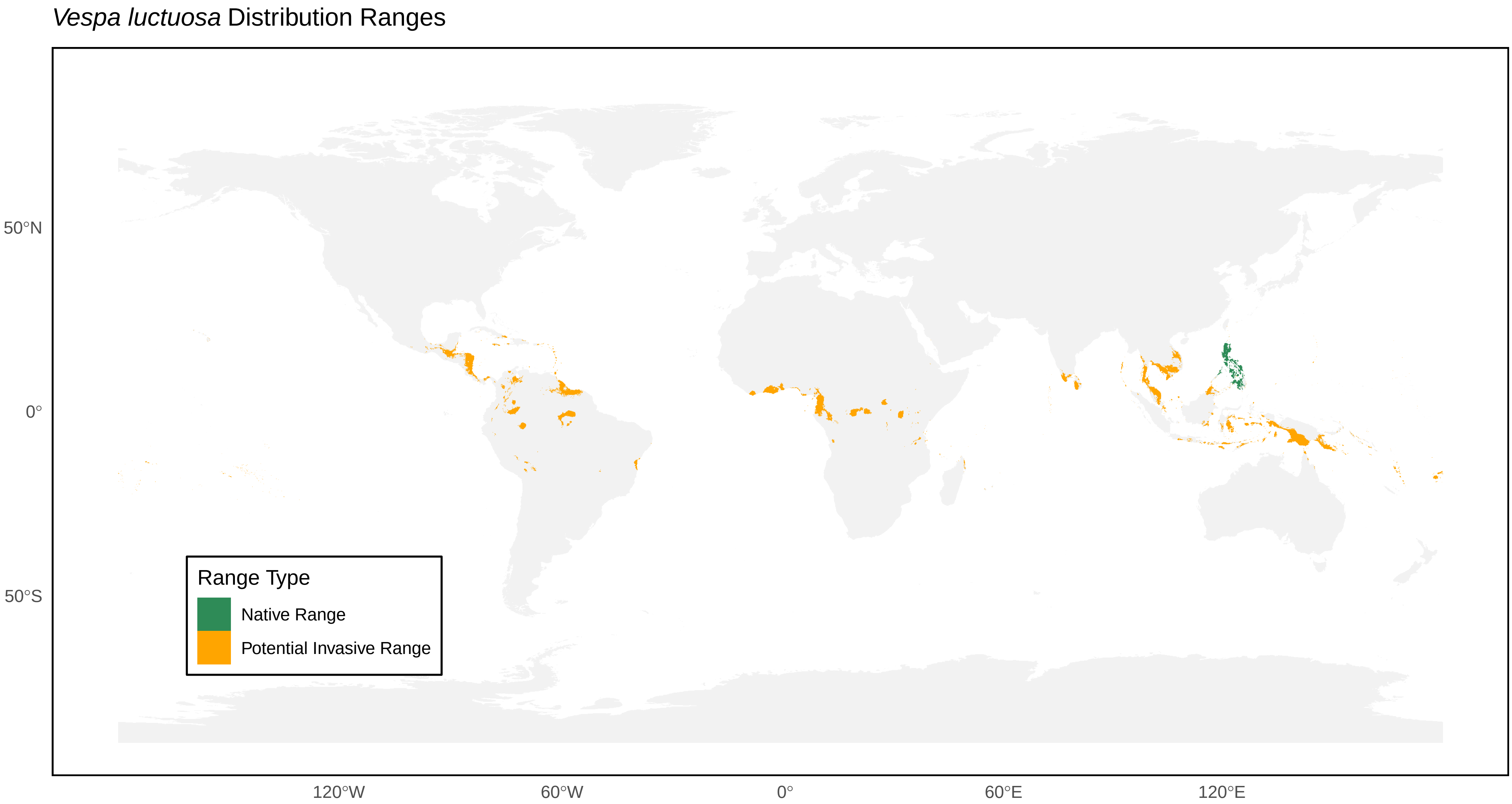

*Vespa luctuosa* Distribution Ranges
50°N
0°
50°S
120°W
60°W
0°
60°E
120°E
Range Type
Native Range
Potential Invasive Range

*Vespa mandarinia* Distribution Ranges

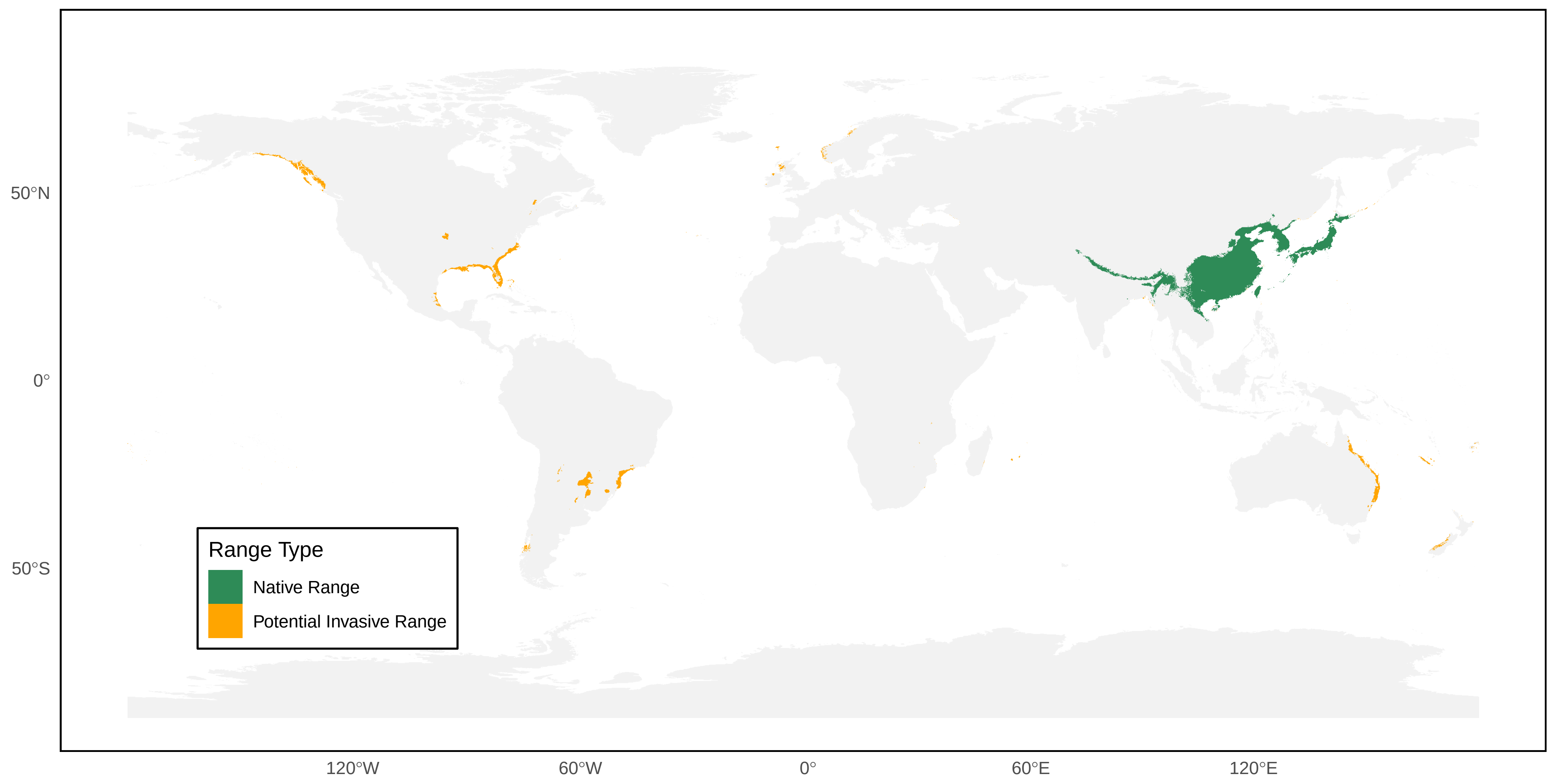

*Vespa mocsaryana* Distribution Ranges

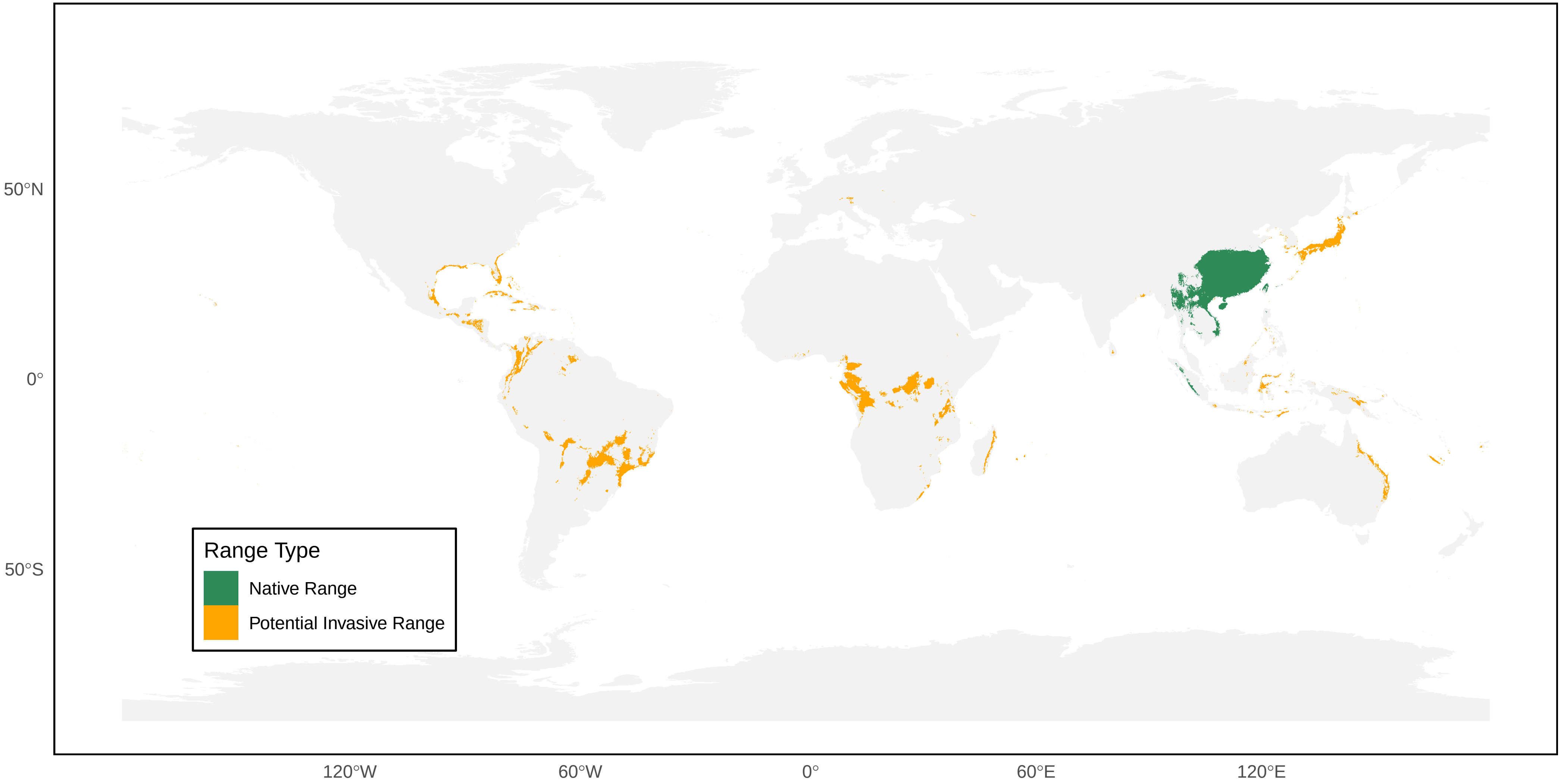

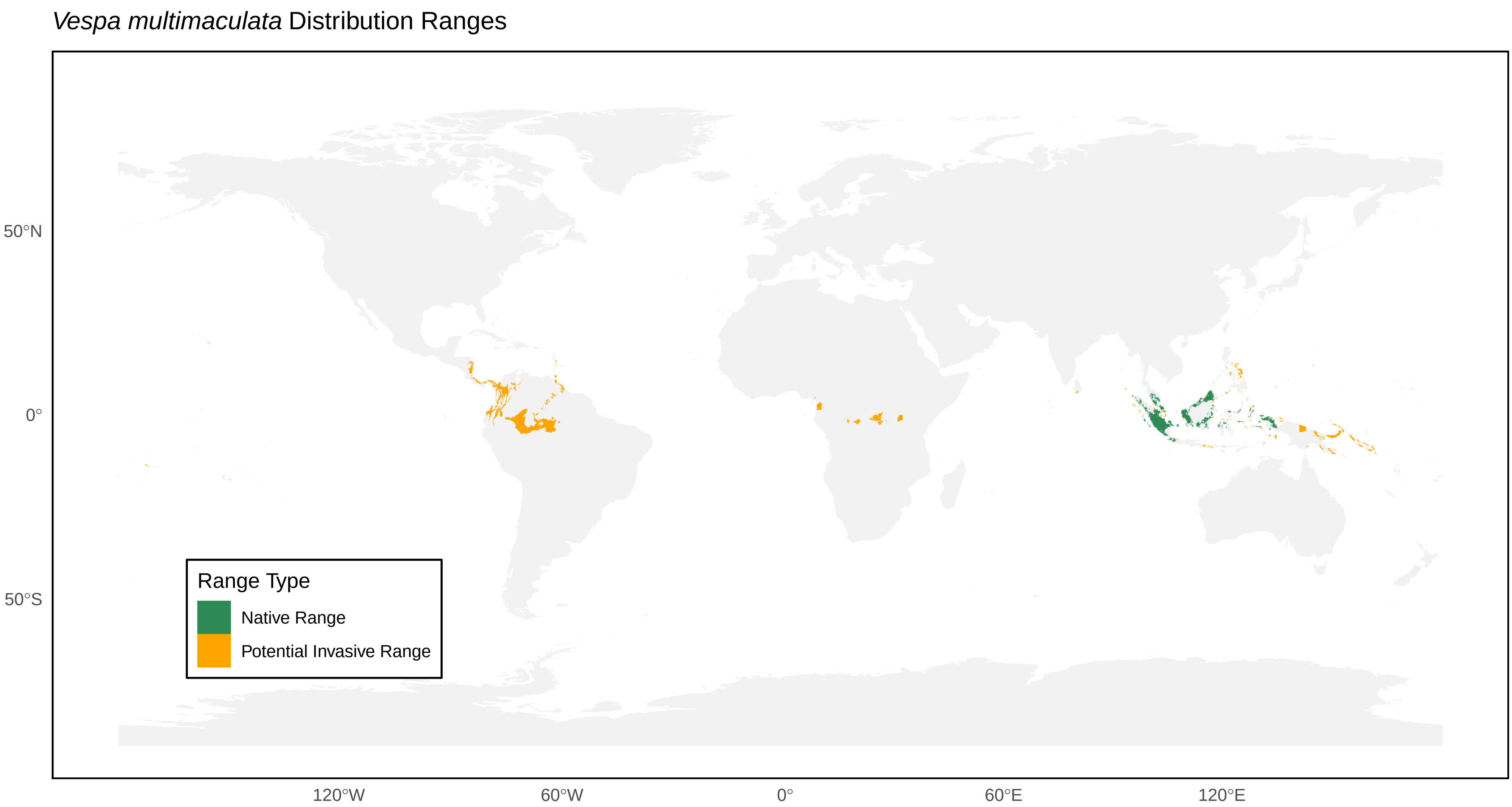
*Vespa multimaculata* Distribution Ranges
50°N
0°
50°S
120°W
60°W
0°
60°E
120°E
Range Type
Native Range
Potential Invasive Range

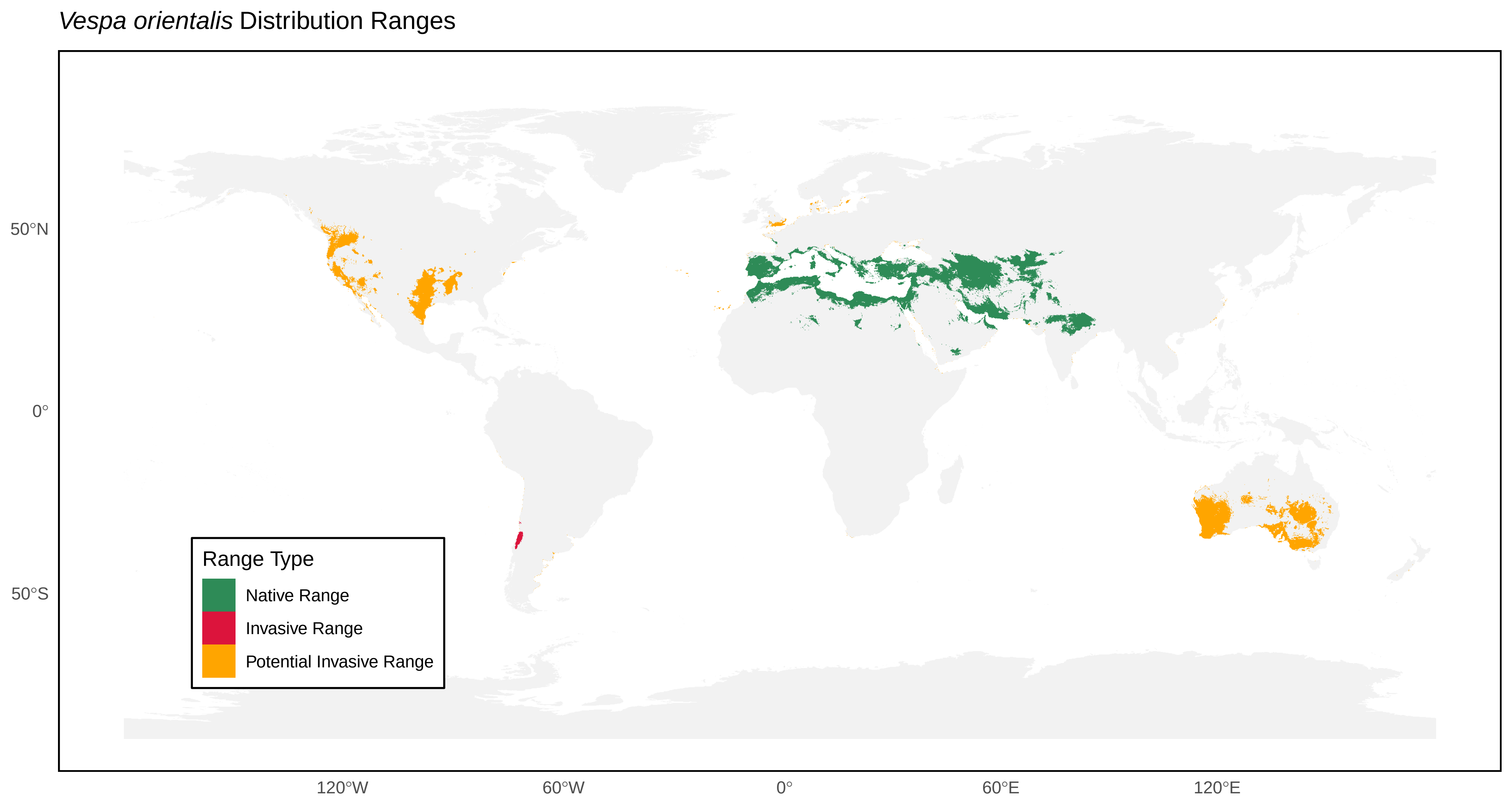

*Vespa orientalis* Distribution Ranges
50°N
0°
50°S
120°W
60°W
0°
60°E
120°E
Range Type
Native Range
Invasive Range
Potential Invasive Range

*Vespa simillima* Distribution Ranges

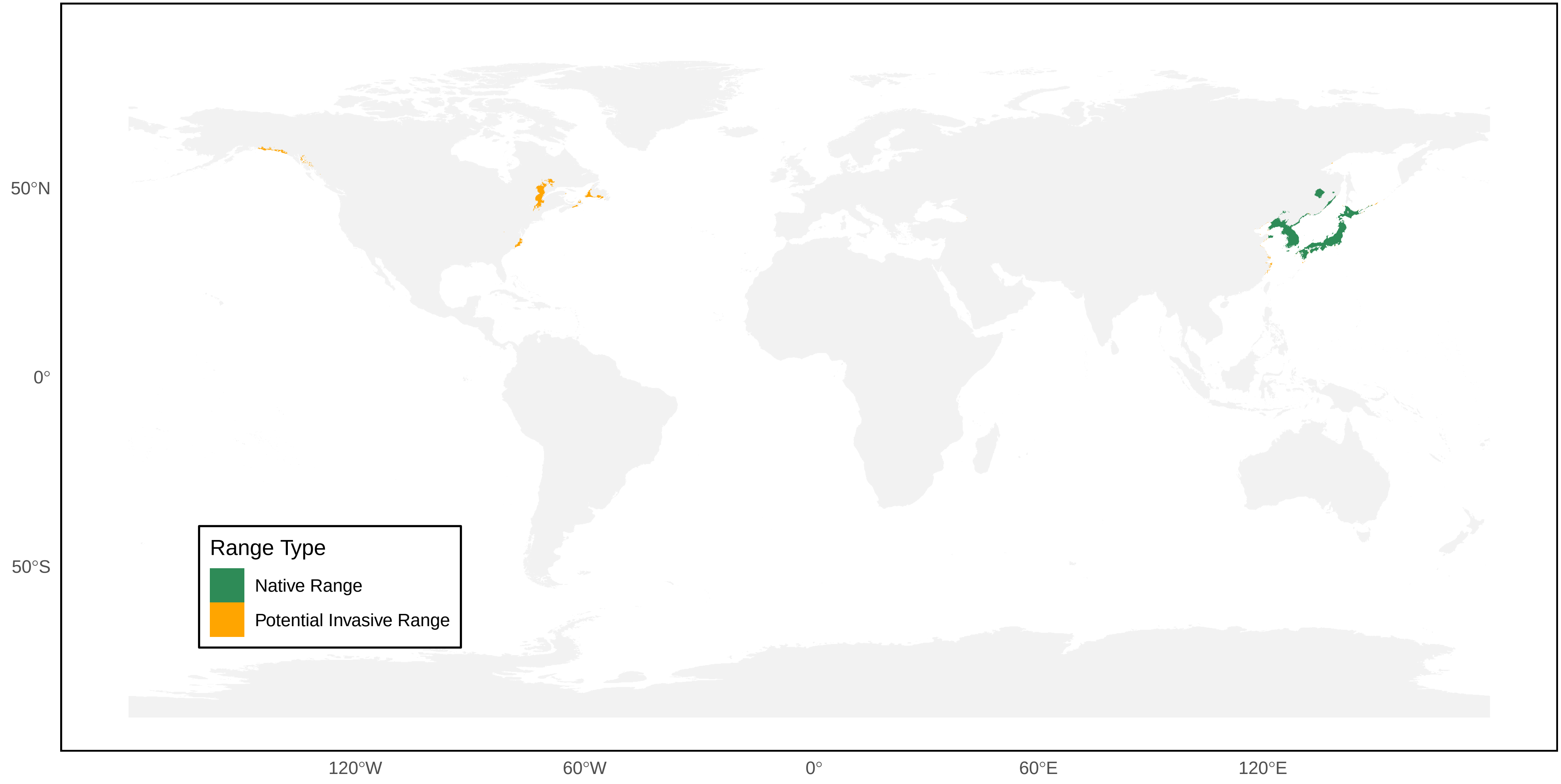

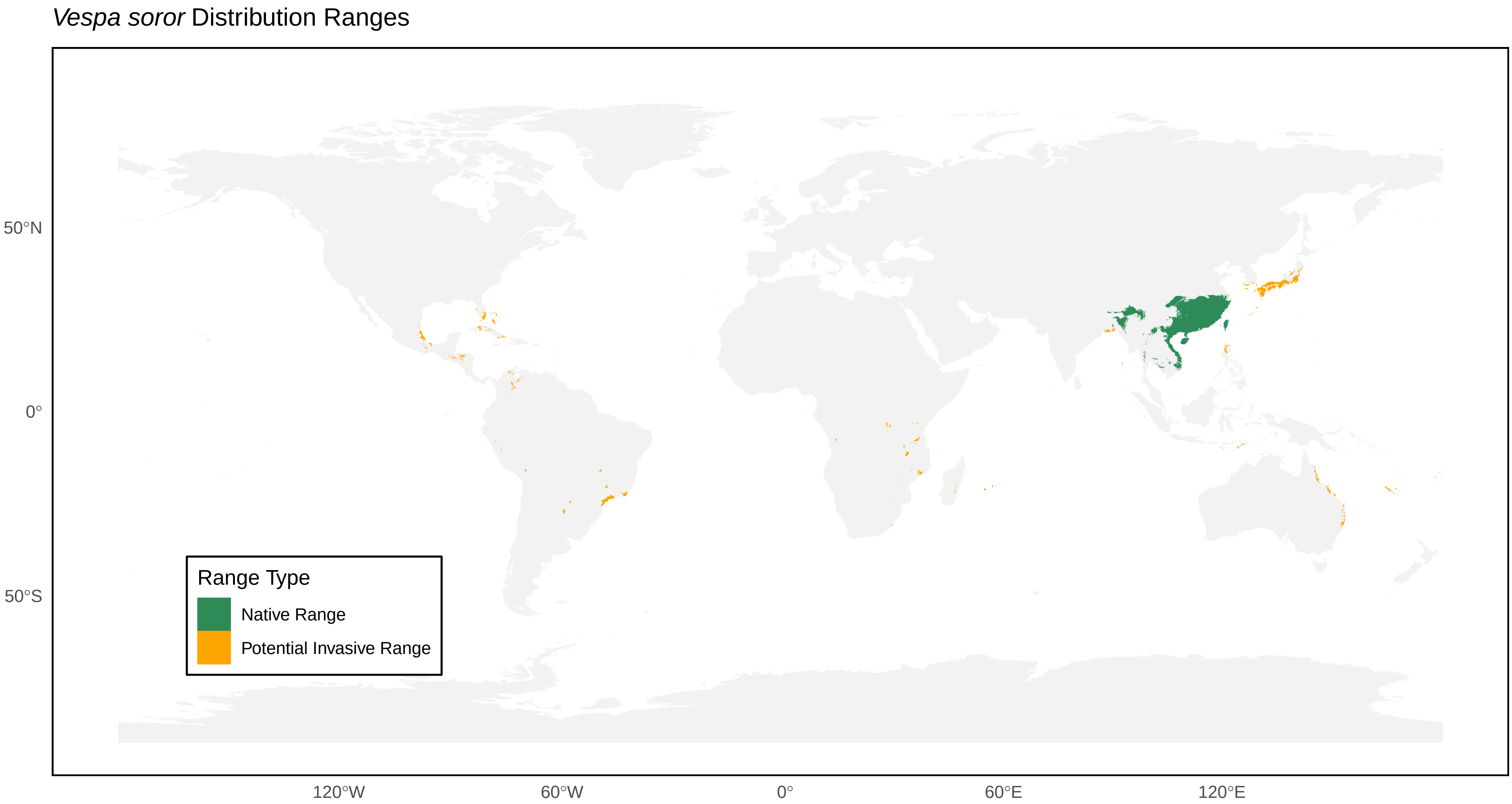

*Vespa soror* Distribution Ranges
50°N
0°
50°S
120°W
60°W
0°
60°E
120°E
Range Type
Native Range
Potential Invasive Range

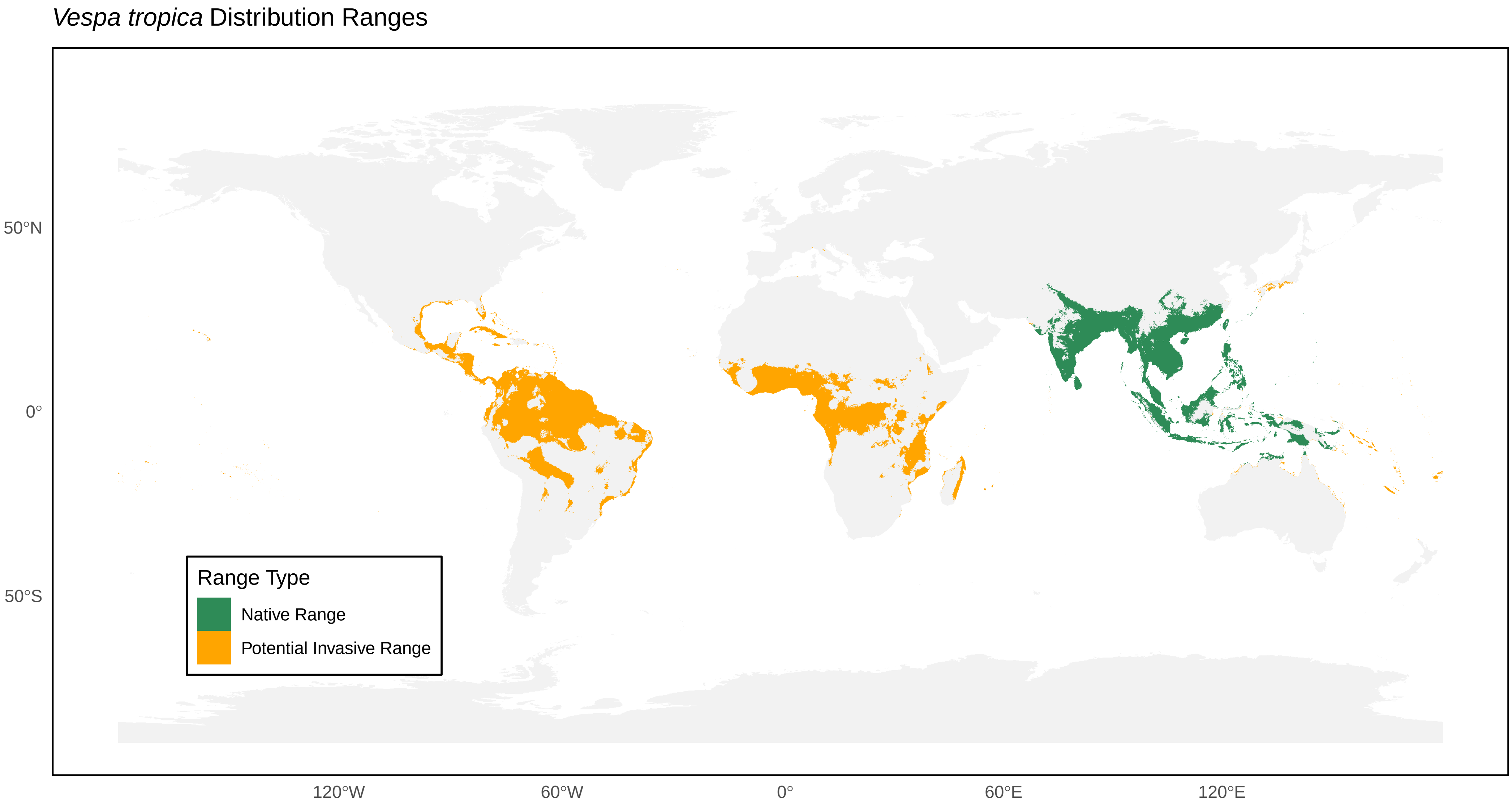

*Vespa tropica* Distribution Ranges
50°N
0°
50°S
120°W
60°W
0°
60°E
120°E
Range Type
Native Range
Potential Invasive Range

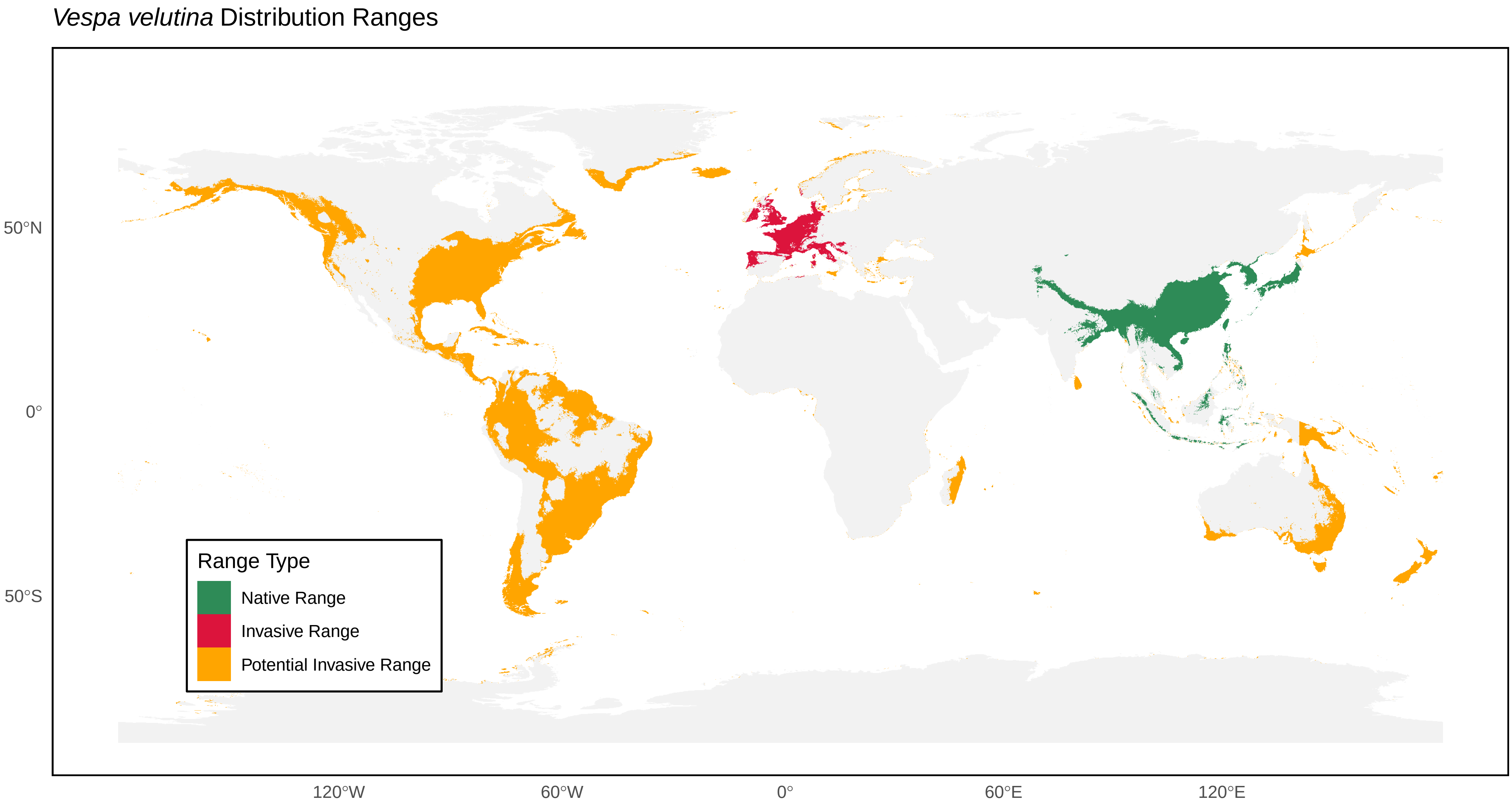
*Vespa velutina* Distribution Ranges
50°N
0°
50°S
120°W
60°W
0°
60°E
120°E
Range Type
Native Range
Invasive Range
Potential Invasive Range

*Vespa vivax* Distribution Ranges

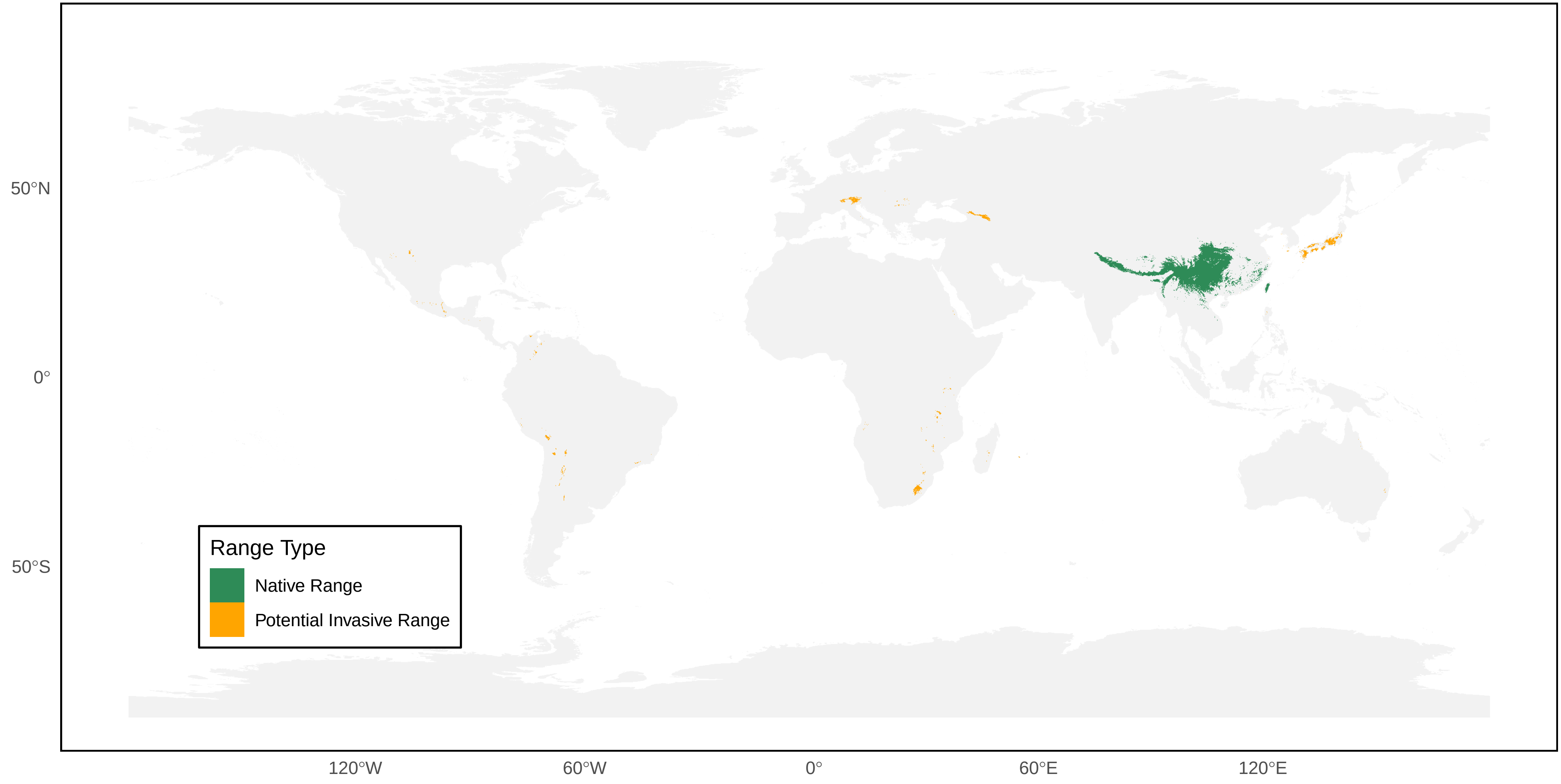

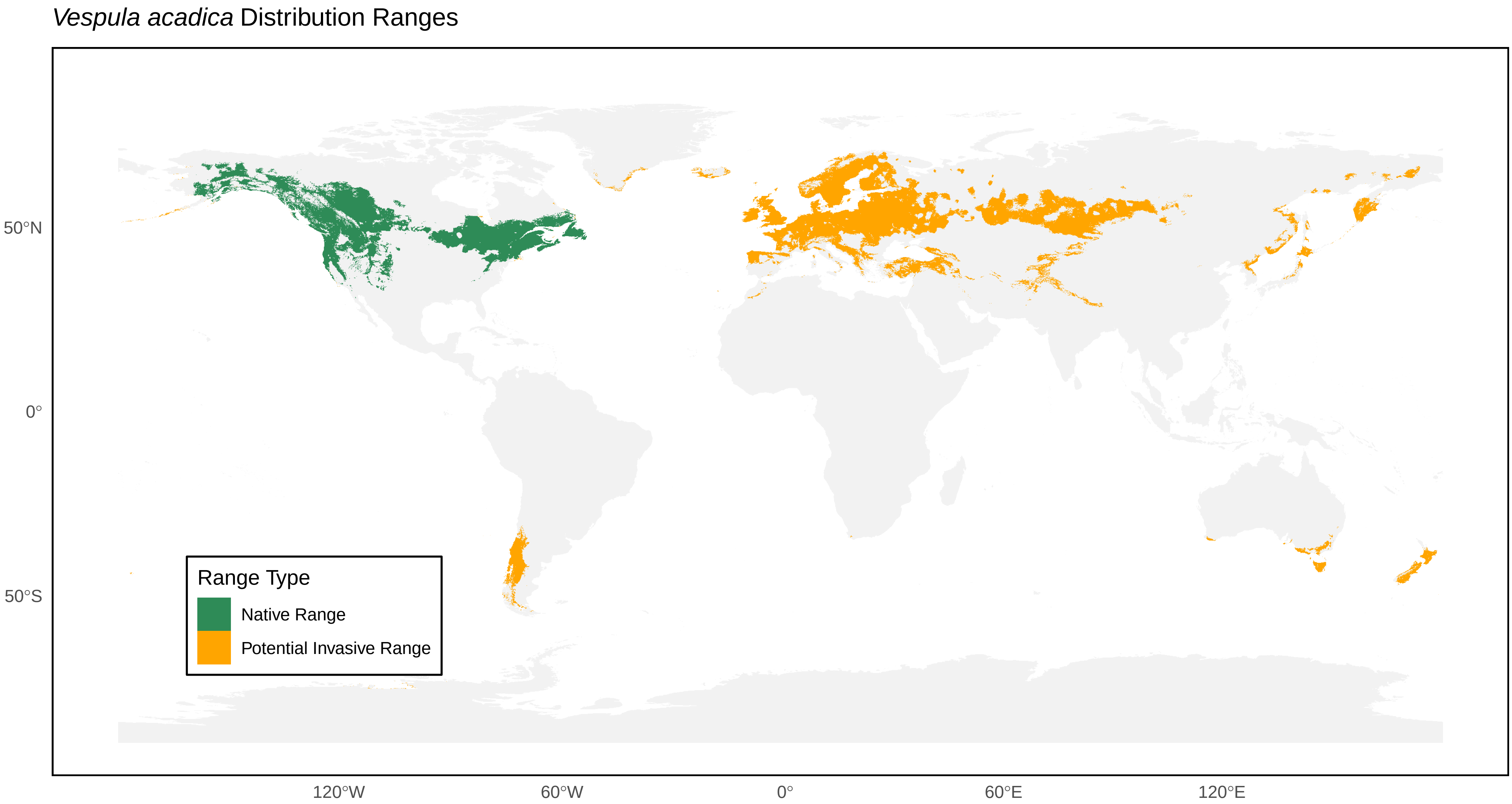
*Vespula acadica* Distribution Ranges
50°N
0°
50°S
120°W
60°W
0°
60°E
120°E
Range Type
Native Range
Potential Invasive Range

*Vespula alascensis* Distribution Ranges

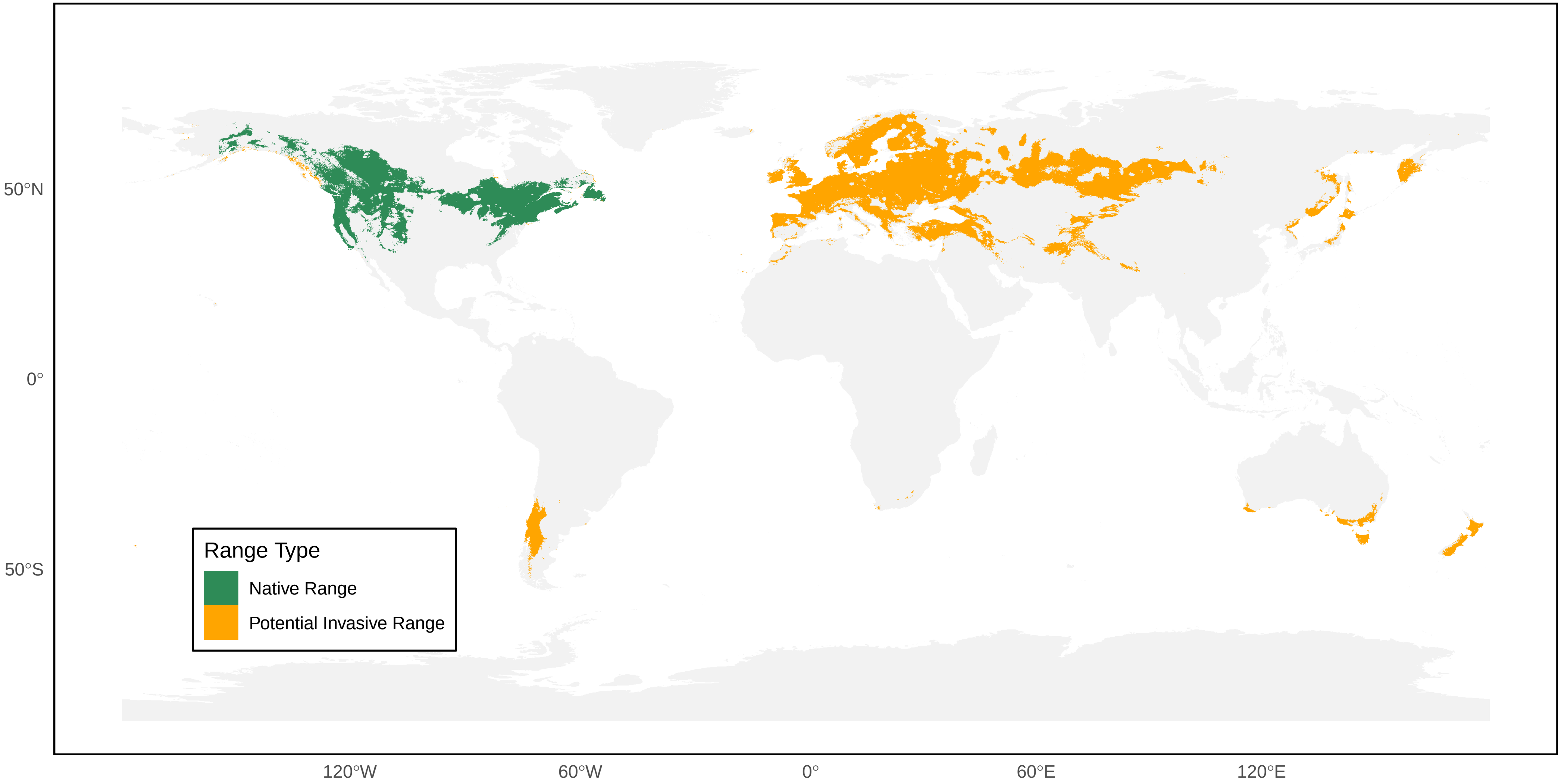

*Vespula arisana* Distribution Ranges

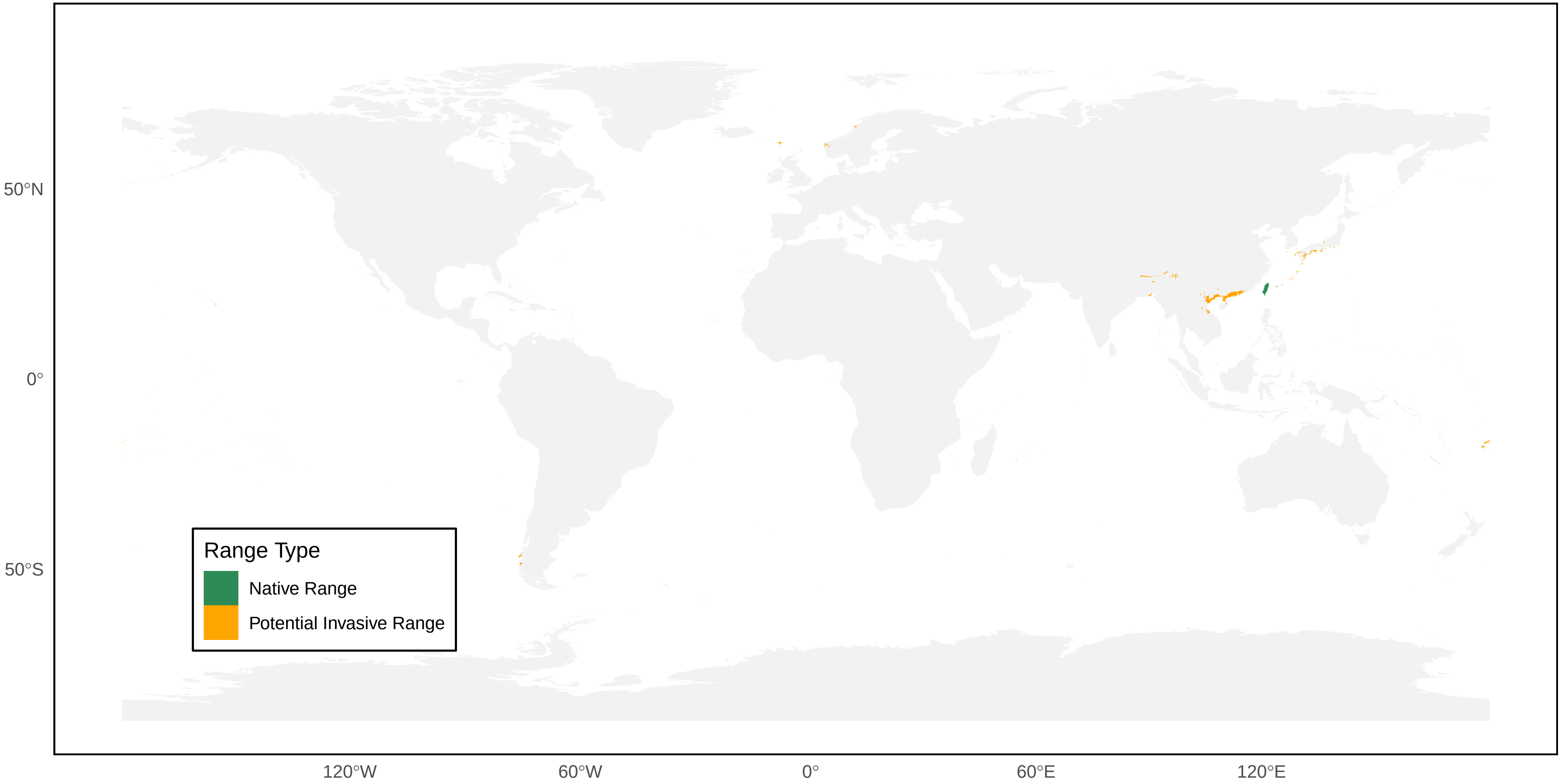

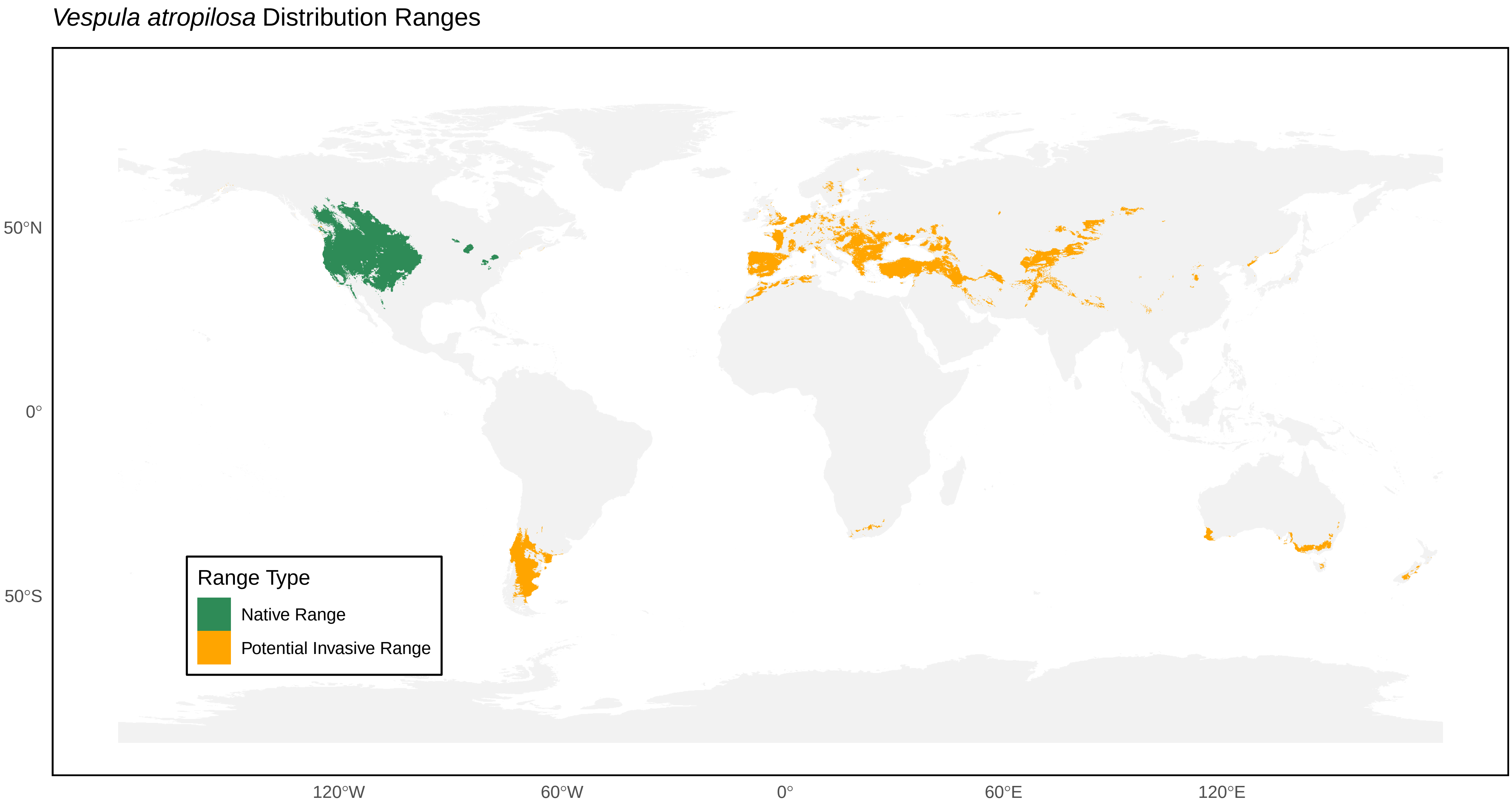
*Vespula atropilosa* Distribution Ranges
50°N
0°
50°S
120°W
60°W
0°
60°E
120°E
Range Type
Native Range
Potential Invasive Range

*Vespula austriaca* Distribution Ranges

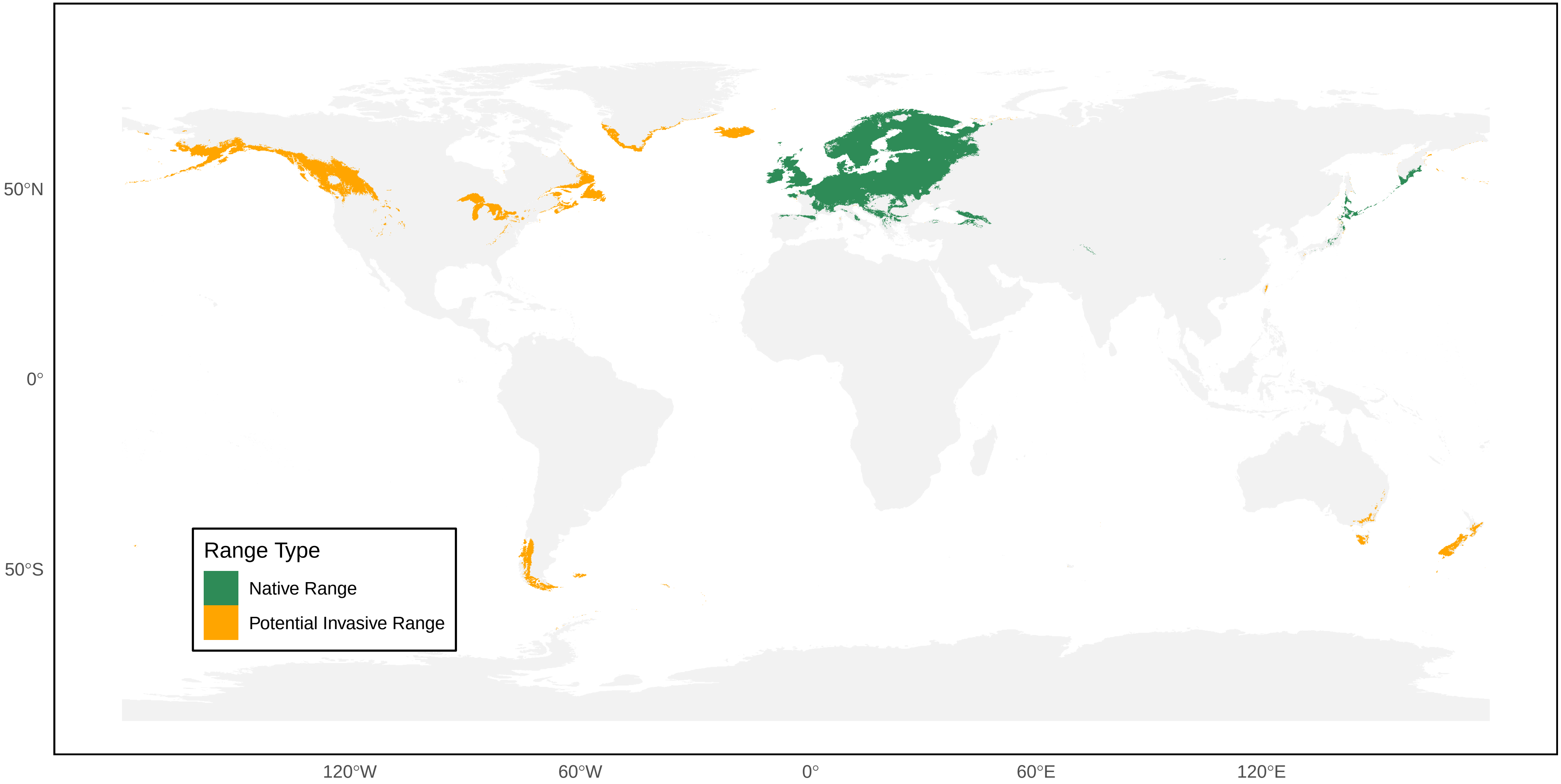

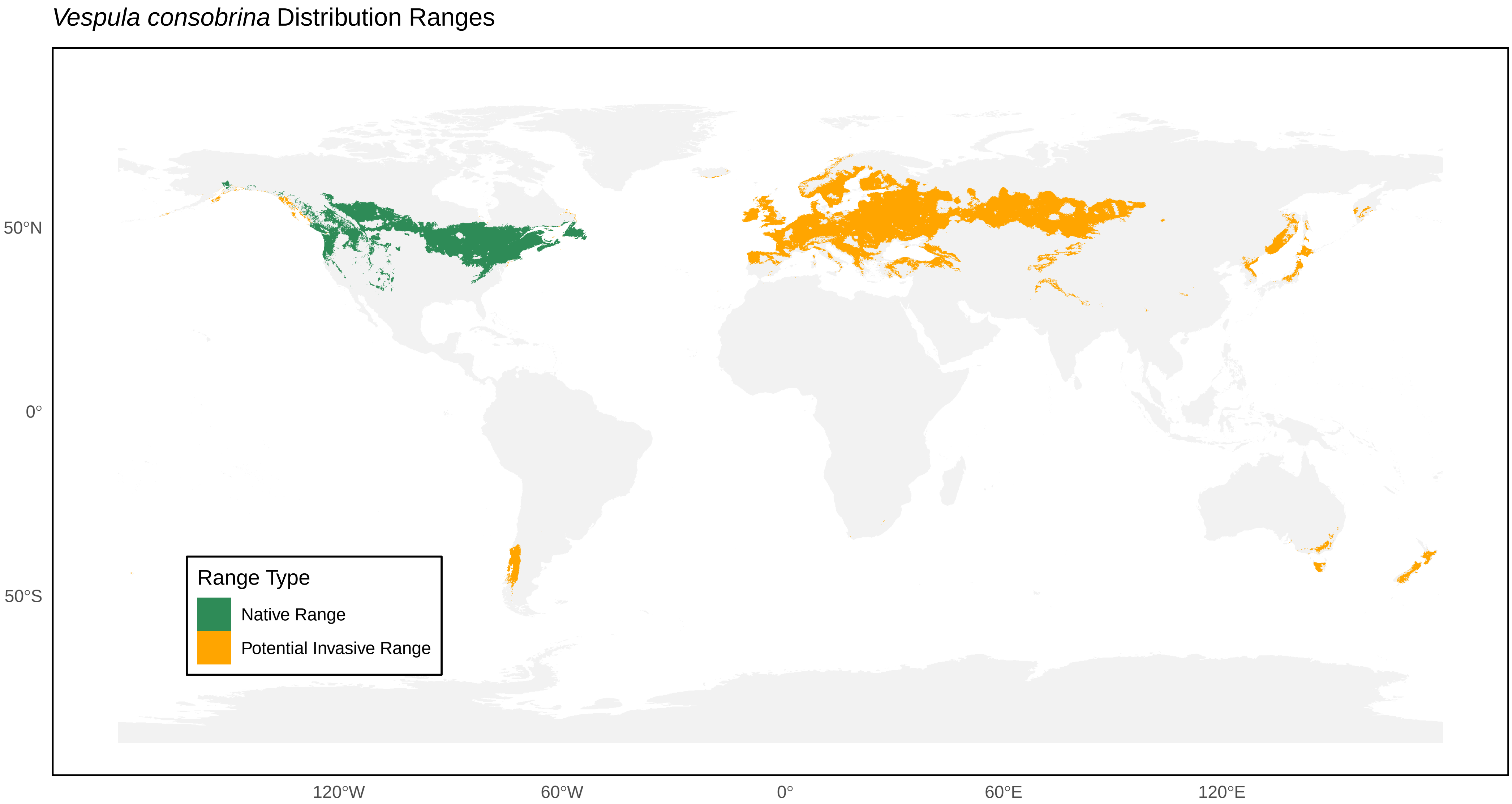
*Vespula consobrina* Distribution Ranges
50°N
0°
50°S
120°W
60°W
0°
60°E
120°E
Range Type
Native Range
Potential Invasive Range

*Vespula flaviceps* Distribution Ranges

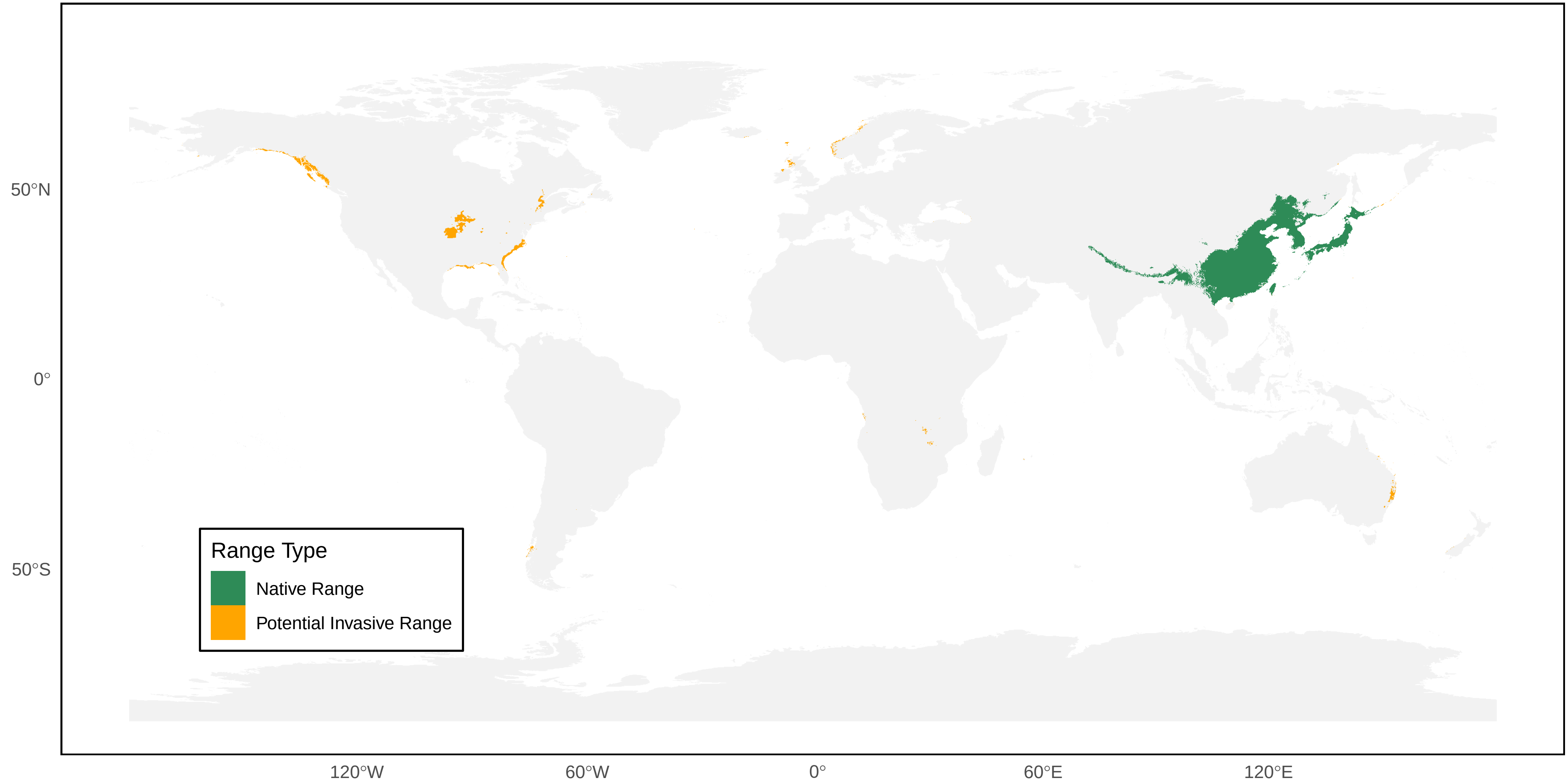

*Vespula flavopilosa* Distribution Ranges

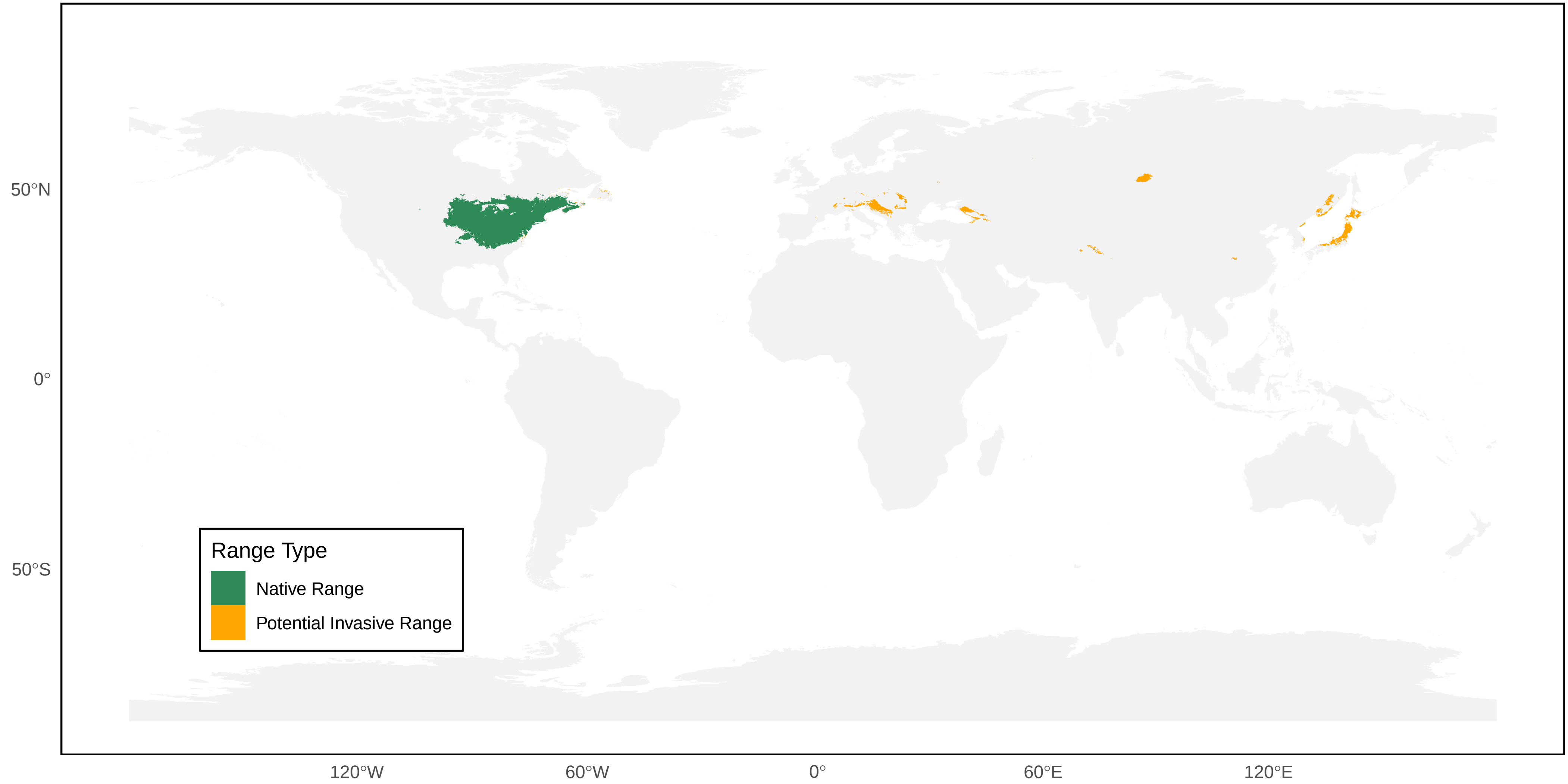

*Vespula germanica* Distribution Ranges

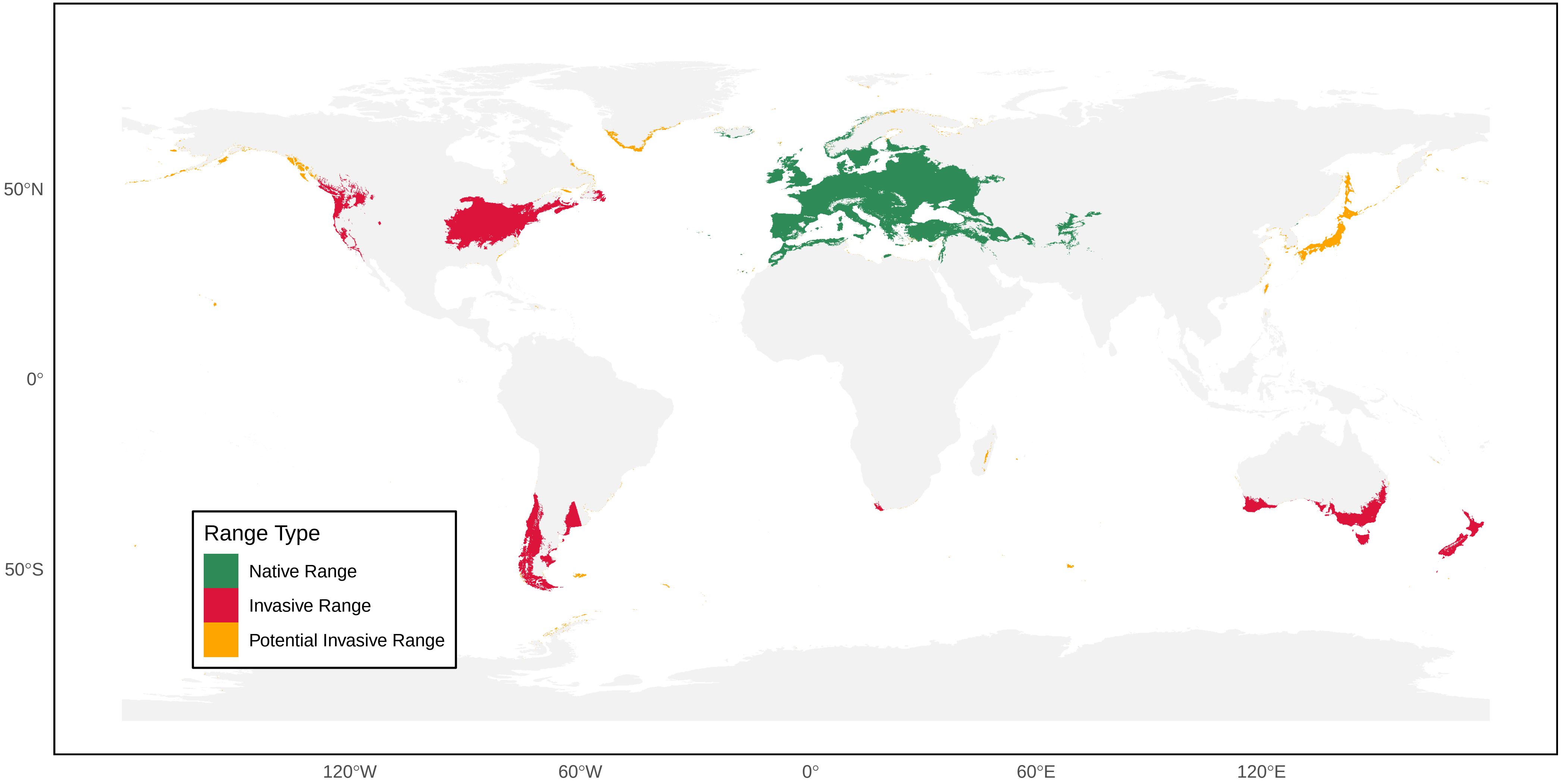

*Vespula infernalis* Distribution Ranges

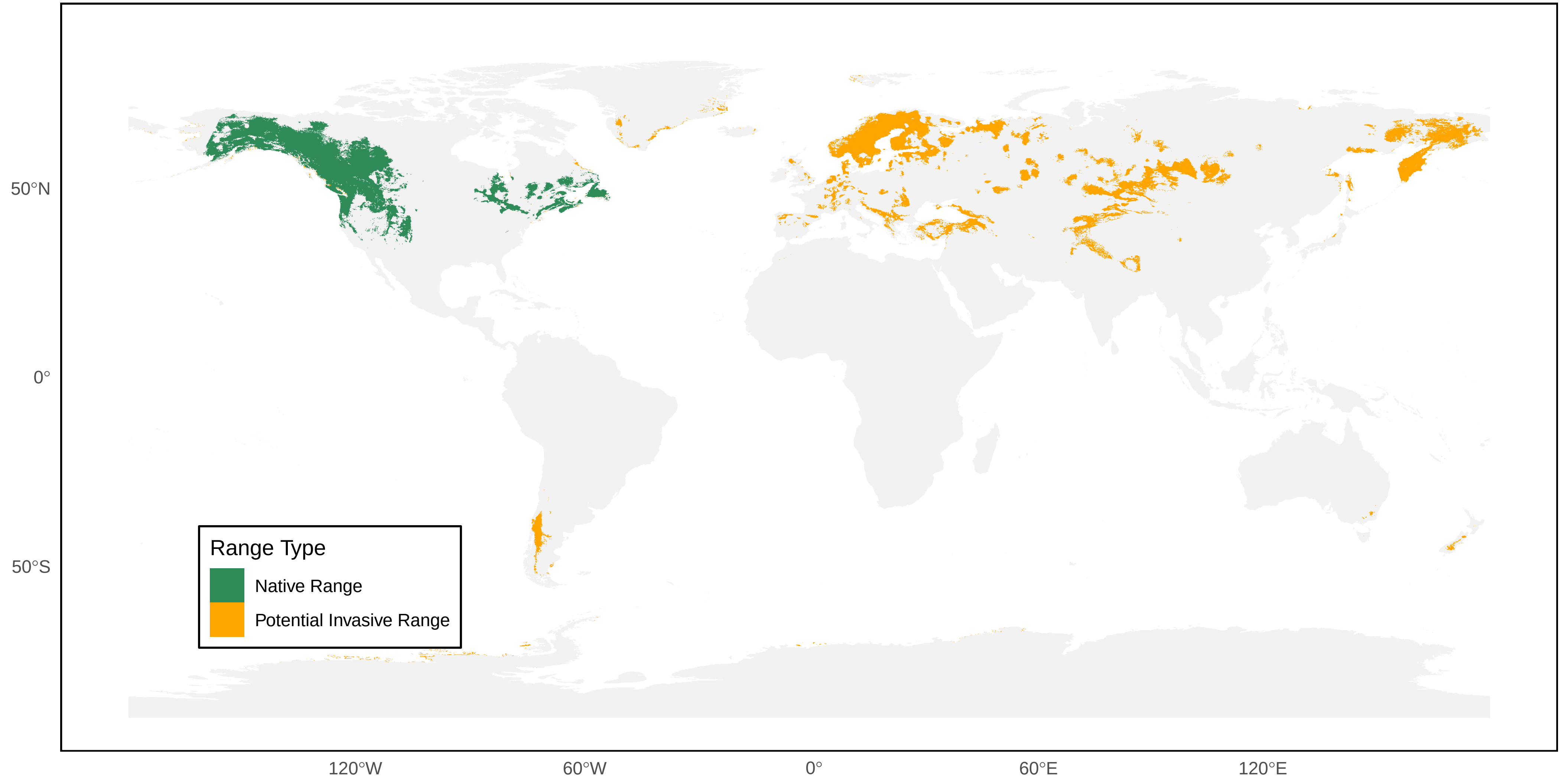

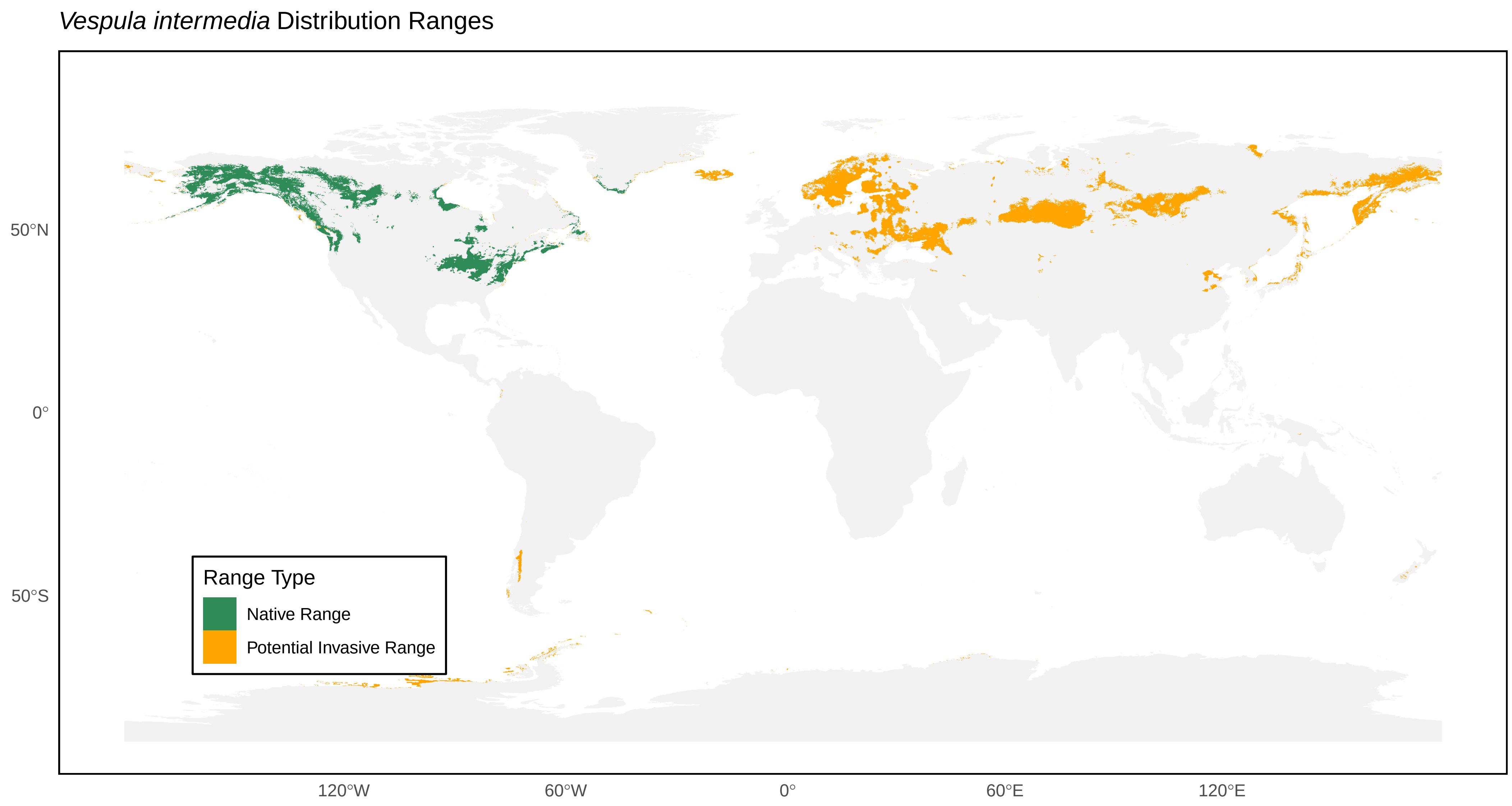

*Vespula intermedia* Distribution Ranges
50°N
0°
50°S
120°W
60°W
0°
60°E
120°E
Range Type
Native Range
Potential Invasive Range

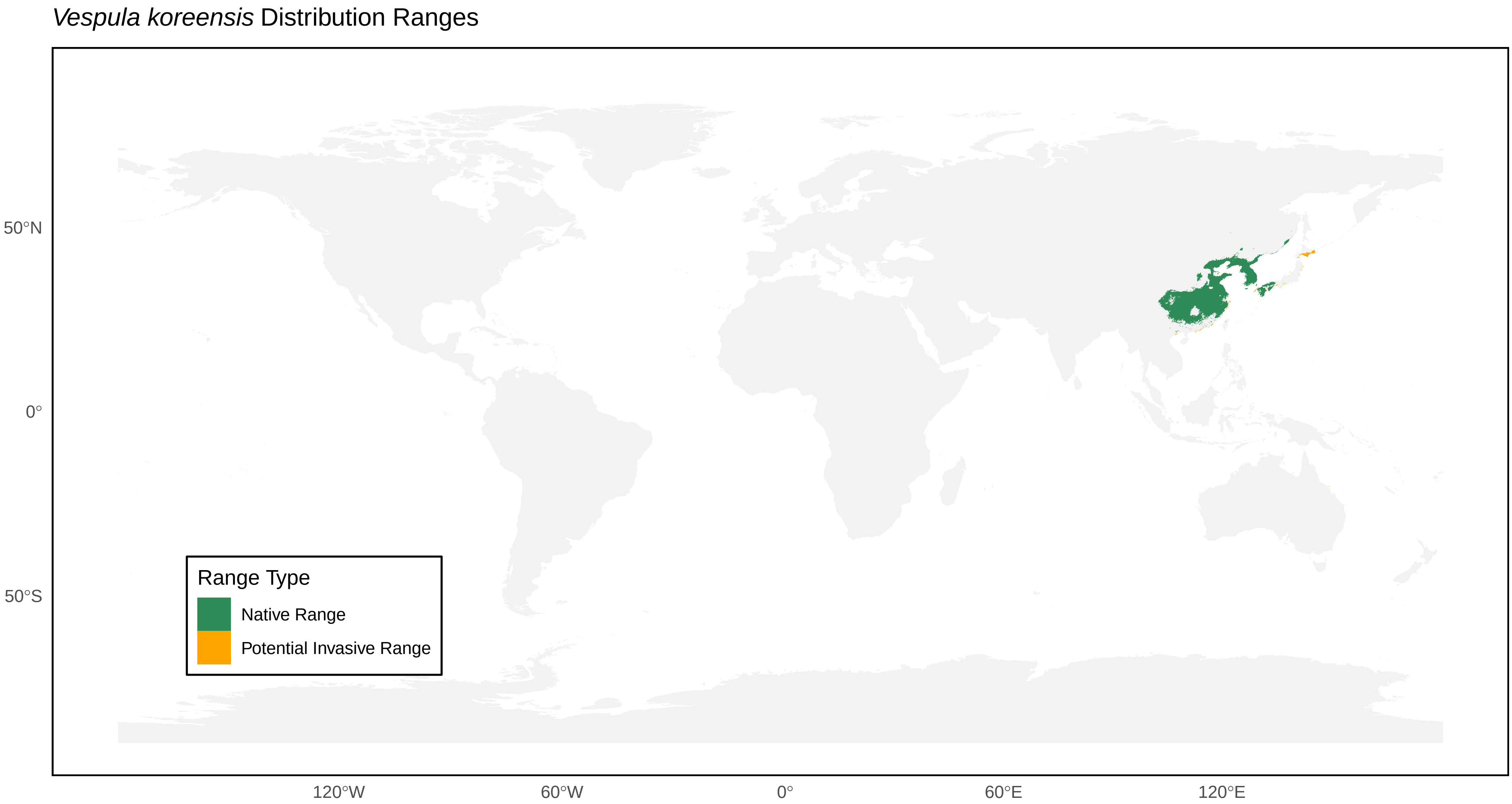
*Vespula koreensis* Distribution Ranges
50°N
0°
50°S
120°W
60°W
0°
60°E
120°E
Range Type
Native Range
Potential Invasive Range

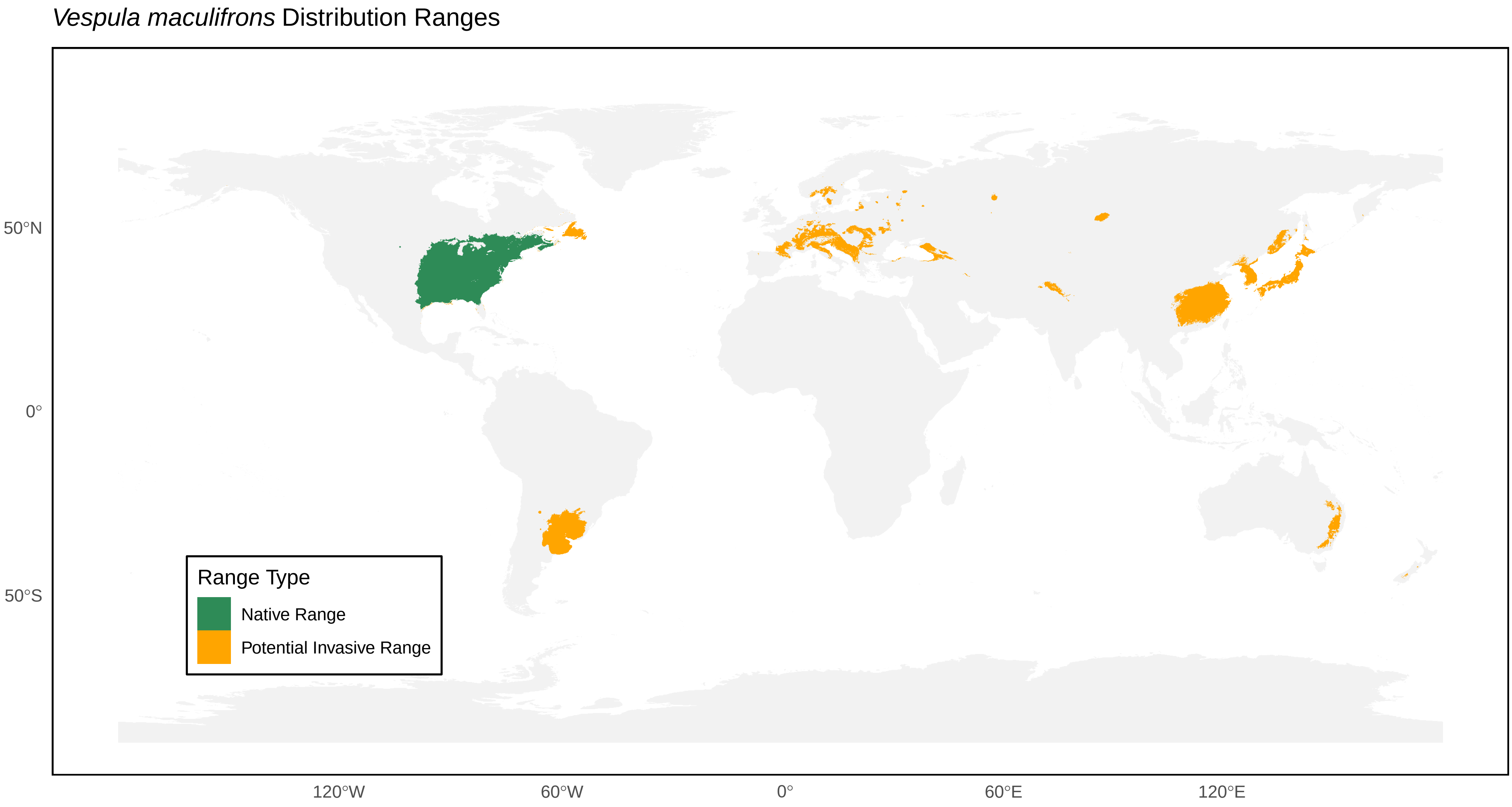
*Vespula maculifrons* Distribution Ranges
50°N
0°
50°S
120°W
60°W
0°
60°E
120°E
Range Type
Native Range
Potential Invasive Range

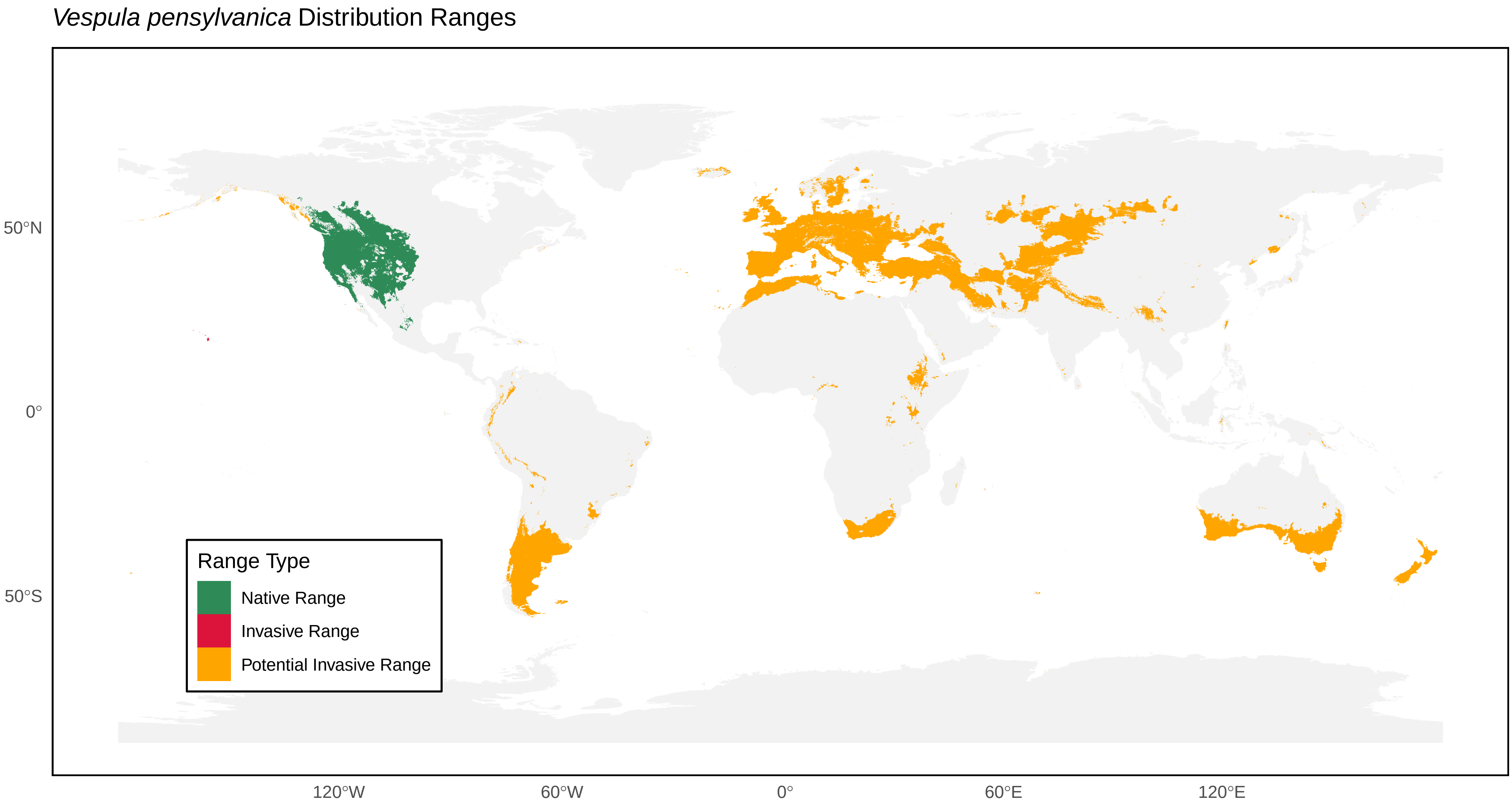

*Vespula pensylvanica* Distribution Ranges
50°N
0°
50°S
120°W
60°W
0°
60°E
120°E
Range Type
Native Range
Invasive Range
Potential Invasive Range

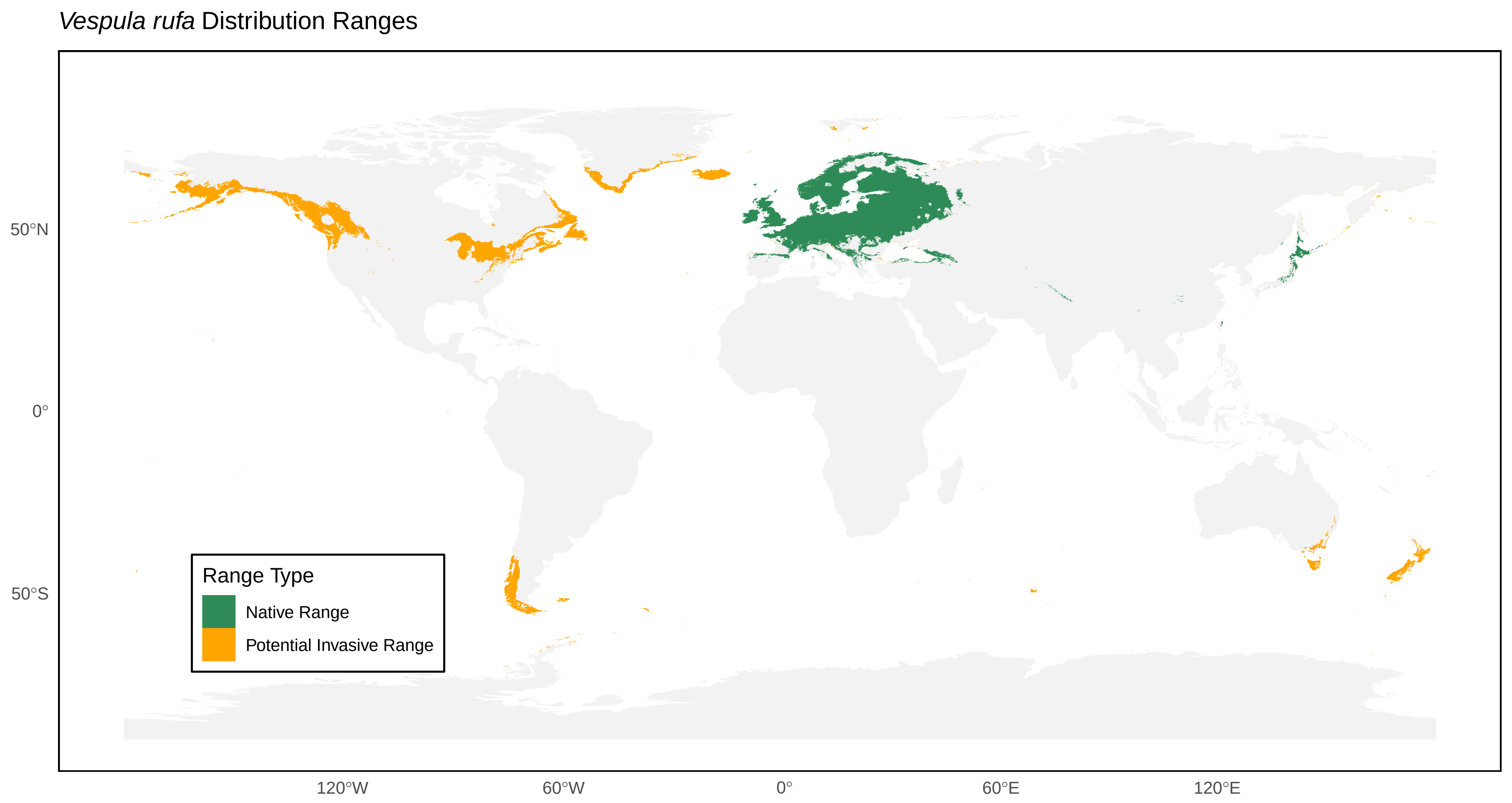
*Vespula rufa* Distribution Ranges
Range Type
Native Range
Potential Invasive Range
50°N
0°
50°S
120°W
60°W
0°
60°E
120°E

*Vespula shidai* Distribution Ranges

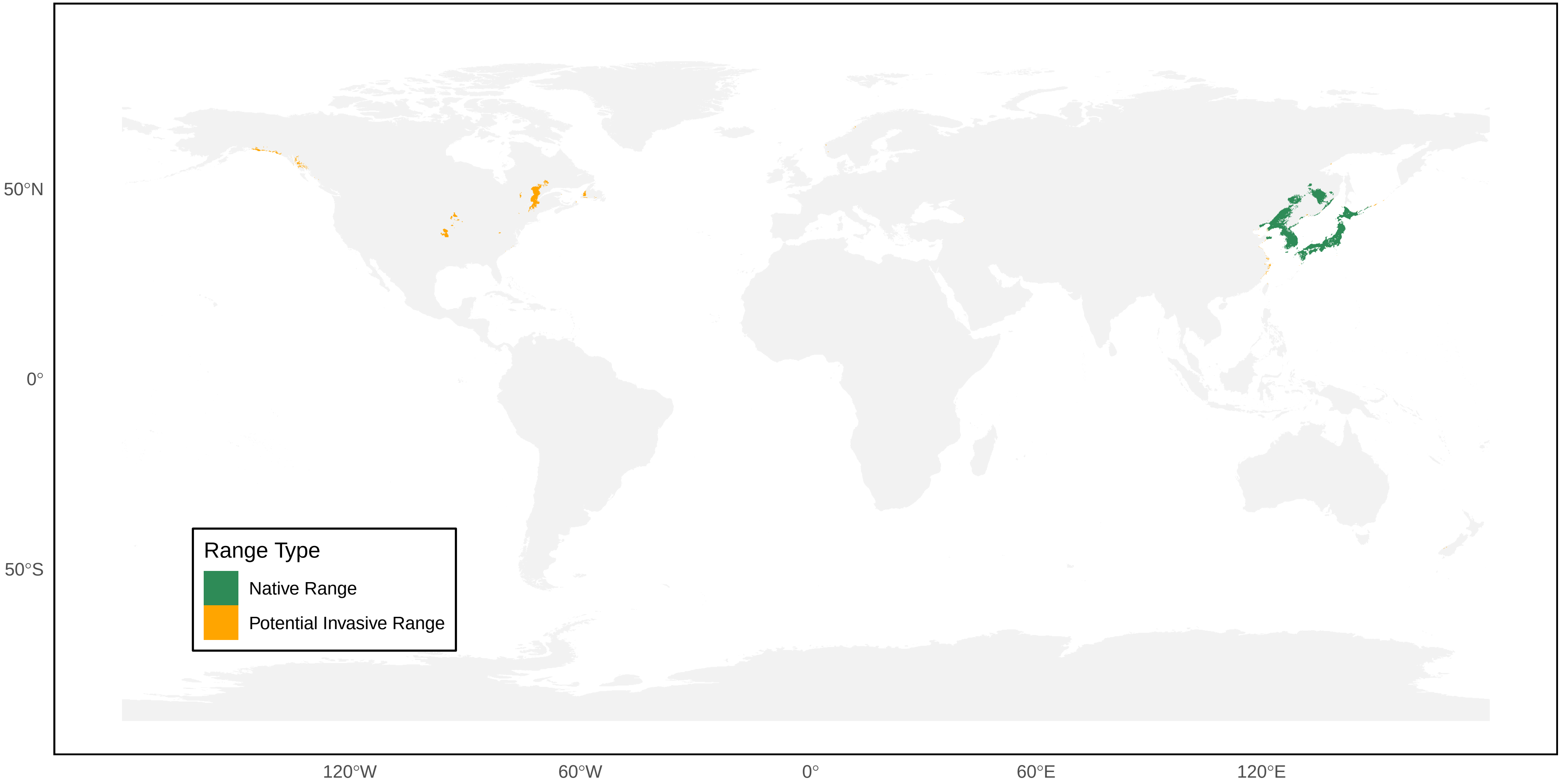

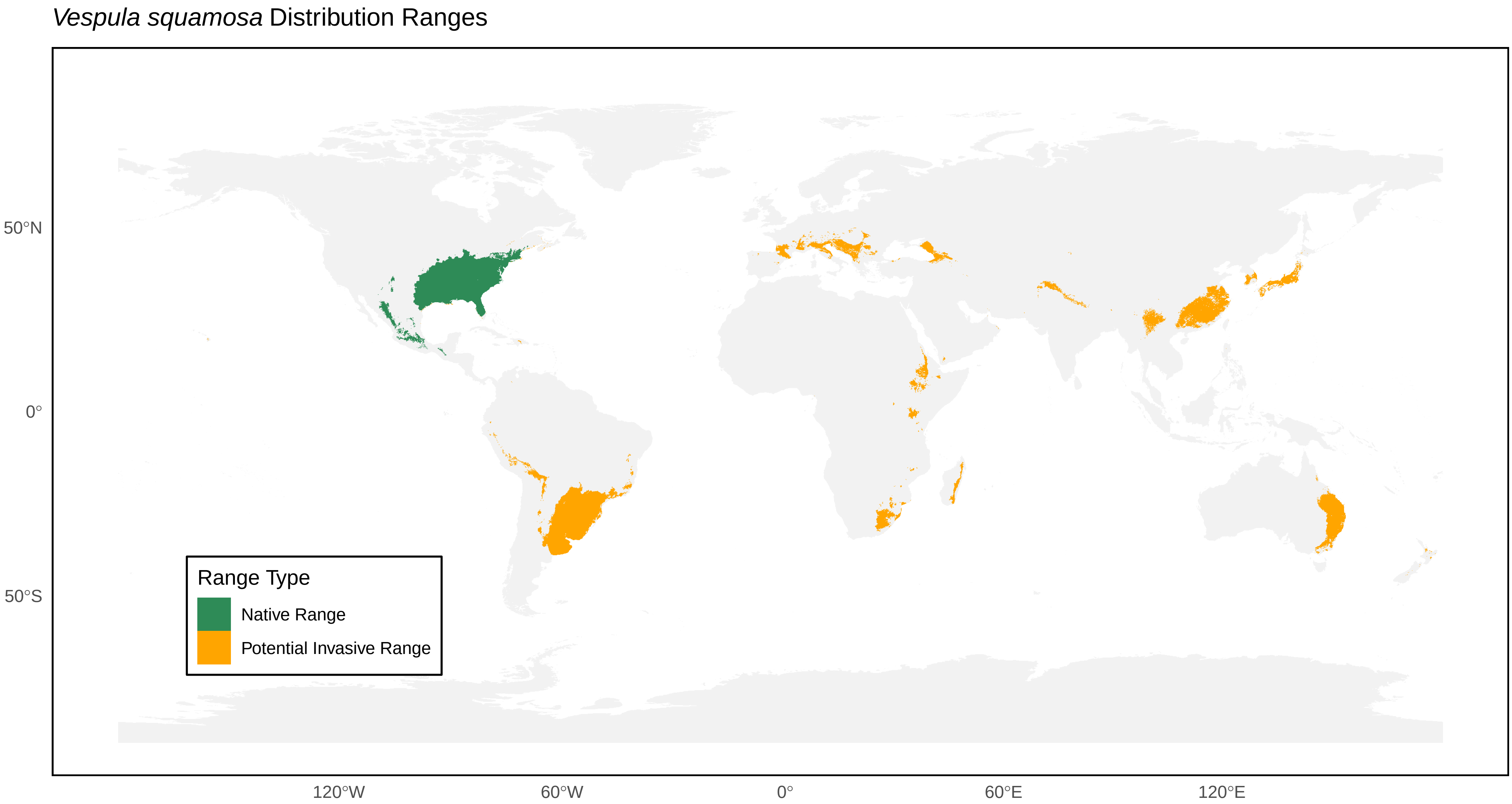
*Vespula squamosa* Distribution Ranges
50°N
0°
50°S
120°W
60°W
0°
60°E
120°E
Range Type
Native Range
Potential Invasive Range

*Vespula sulphurea* Distribution Ranges

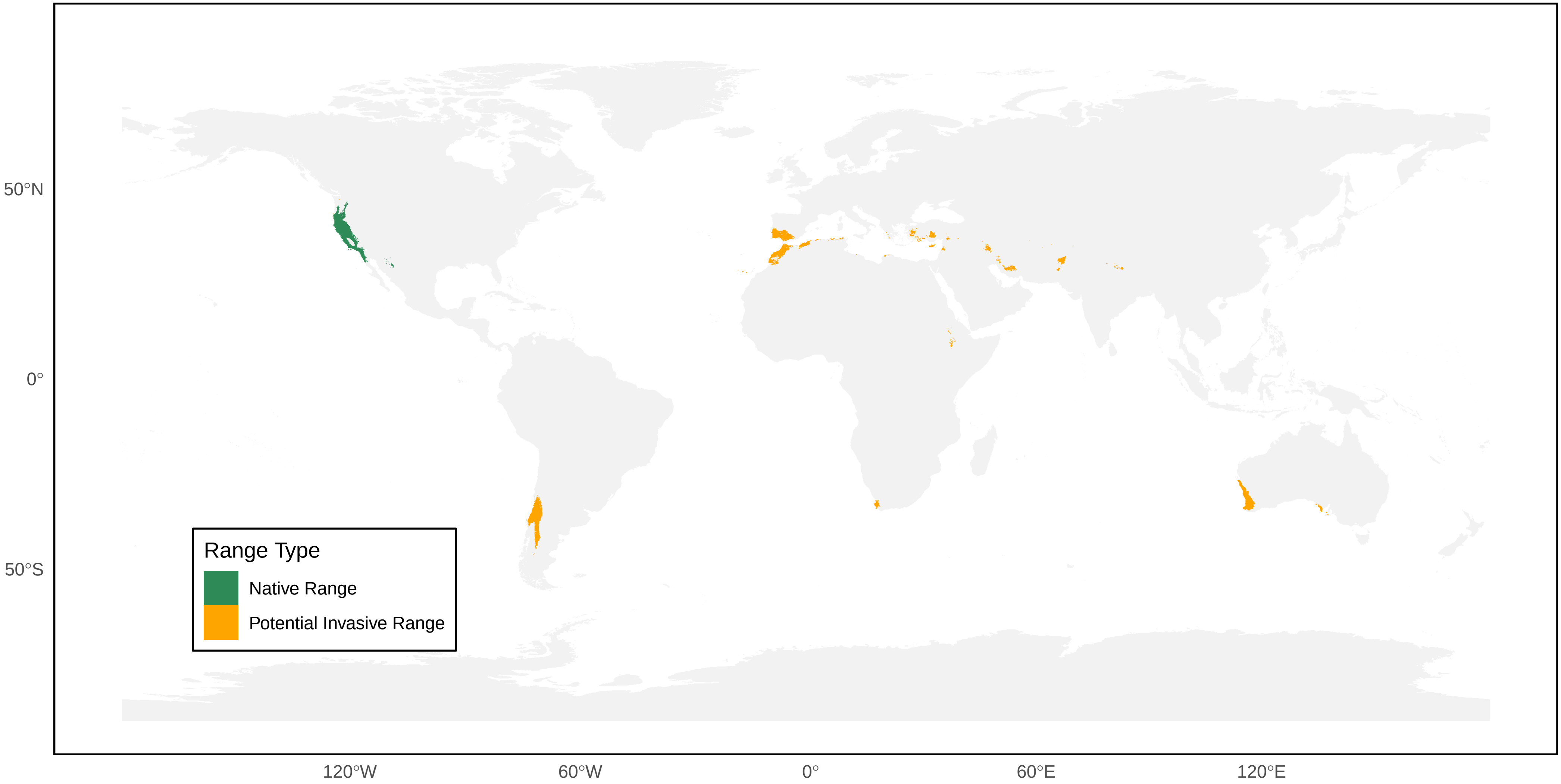

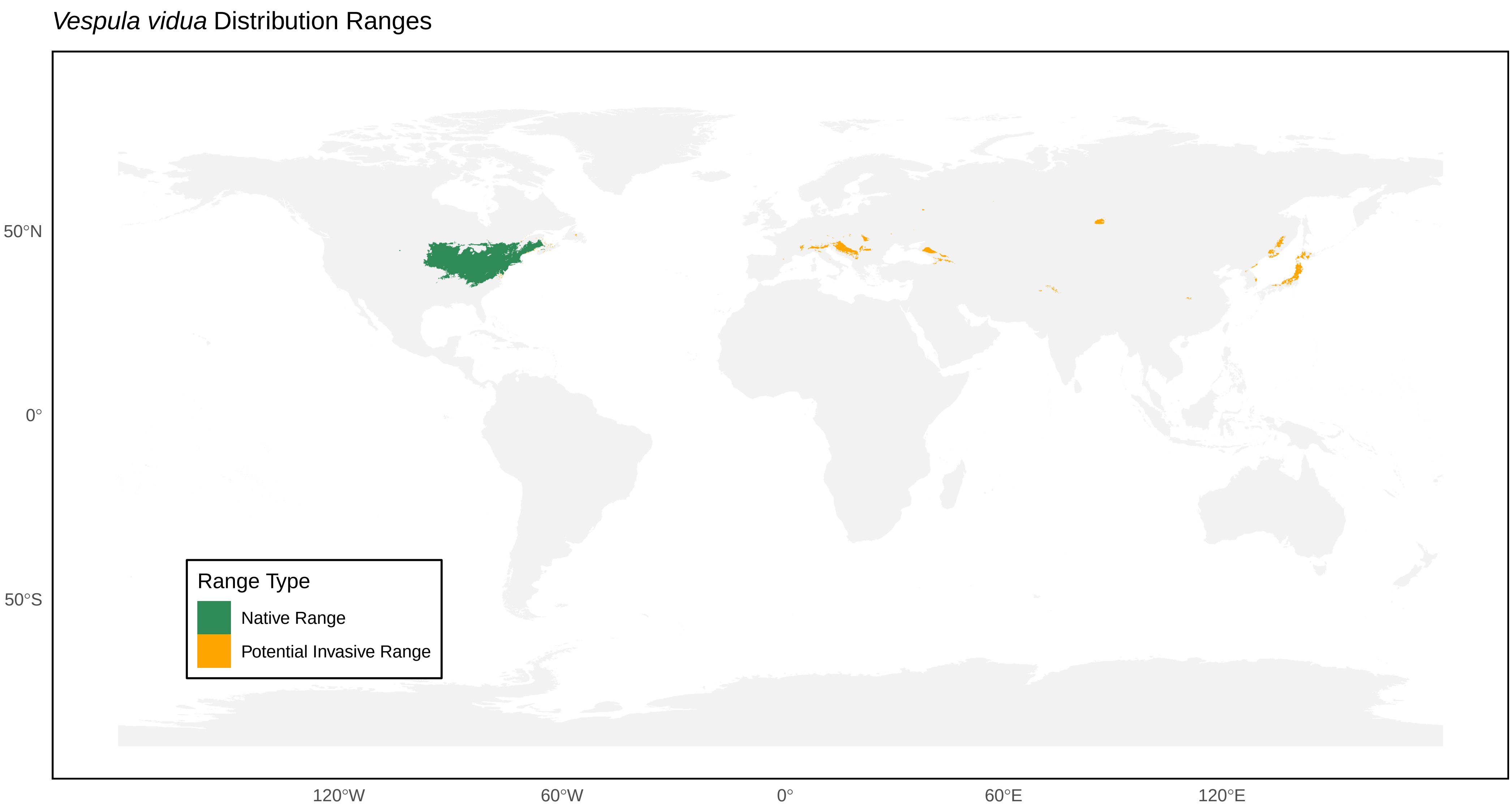
Vespula vidua Distribution Ranges
50°N
0°
50°S
120°W
60°W
0°
60°E
120°E
Range Type
Native Range
Potential Invasive Range

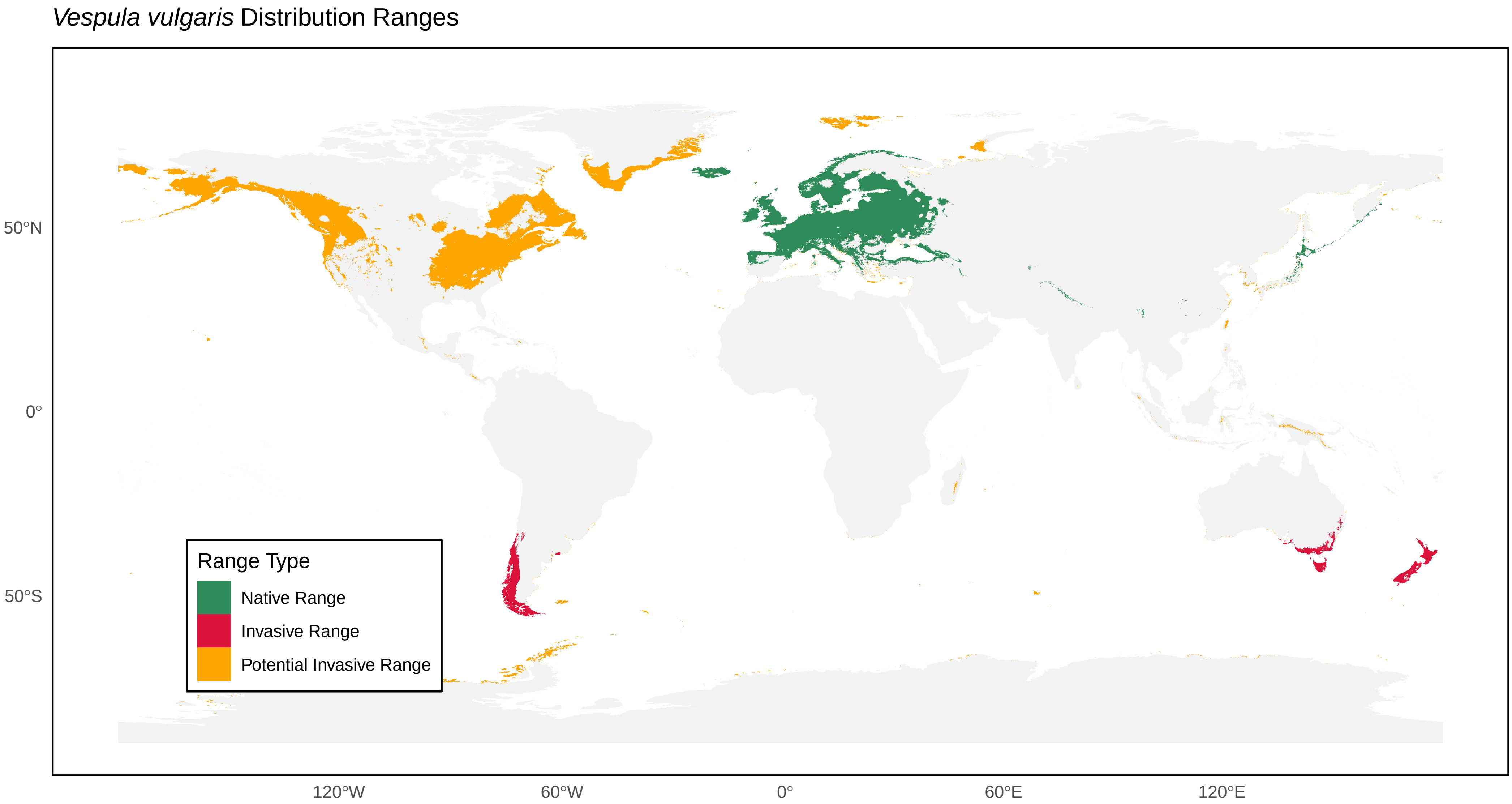
*Vespula vulgaris* Distribution Ranges
Range Type
Native Range
Invasive Range
Potential Invasive Range
50°N
0°
50°S
120°W
60°W
0°
60°E
120°E
